# Postglacial genomes from foragers across Northern Eurasia reveal prehistoric mobility associated with the spread of the Uralic and Yeniseian languages

**DOI:** 10.1101/2023.10.01.560332

**Authors:** Tian Chen Zeng, Leonid A. Vyazov, Alexander Kim, Pavel Flegontov, Kendra Sirak, Robert Maier, Iosif Lazaridis, Ali Akbari, Michael Frachetti, Alexey A. Tishkin, Natalia E. Ryabogina, Sergey A. Agapov, Danila S. Agapov, Anatoliy N. Alekseev, Gennady G. Boeskorov, Anatoly P. Derevianko, Viktor M. Dyakonov, Dmitry N. Enshin, Alexey V. Fribus, Yaroslav V. Frolov, Sergey P. Grushin, Alexander A. Khokhlov, Kirill Yu Kiryushin, Yurii F. Kiryushin, Egor P. Kitov, Pavel Kosintsev, Igor V. Kovtun, Nikolai P. Makarov, Viktor V. Morozov, Egor N. Nikolaev, Marina P. Rykun, Tatyana M. Savenkova, Marina V. Shchelchkova, Vladimir Shirokov, Svetlana N. Skochina, Olga S. Sherstobitova, Sergey M. Slepchenko, Konstantin N. Solodovnikov, Elena N. Solovyova, Aleksandr D. Stepanov, Aleksei A. Timoshchenko, Aleksandr S. Vdovin, Anton V. Vybornov, Elena V. Balanovska, Stanislav Dryomov, Garrett Hellenthal, Kenneth Kidd, Johannes Krause, Elena Starikovskaya, Rem Sukenik, Tatiana Tatarinova, Mark G. Thomas, Maxat Zhabagin, Kim Callan, Olivia Cheronet, Daniel Fernandes, Denise Keating, Candilio Francesca, Lora Iliev, Aisling Kearns, Kadir Toykan Özdoğan, Matthew Mah, Adam Micco, Megan Michel, Iñigo Olalde, Fatma Zalzala, Swapan Mallick, Nadin Rohland, Ron Pinhasi, Vagheesh Narasimhan, David Reich

**Author notes:** Contributed equally (Tian Chen Zeng, Leonid A. Vyazov, Alexander Kim).

## Abstract

The North Eurasian forest and forest-steppe zones have sustained millennia of sociocultural connections among northern peoples. We present genome-wide ancient DNA data for 181 individuals from this region spanning the Mesolithic, Neolithic and Bronze Age. We find that Early to Mid-Holocene hunter-gatherer populations from across the southern forest and forest-steppes of Northern Eurasia can be characterized by a continuous gradient of ancestry that remained stable for millennia, ranging from fully West Eurasian in the Baltic region to fully East Asian in the Transbaikal region. In contrast, cotemporaneous groups in far Northeast Siberia were genetically distinct, retaining high levels of continuity from a population that was the primary source of ancestry for Native Americans. By the mid-Holocene, admixture between this early Northeastern Siberian population and groups from Inland East Asia and the Amur River Basin produced two distinctive populations in eastern Siberia that played an important role in the genetic formation of later people. Ancestry from the first population, Cis-Baikal Late Neolithic–Bronze Age (Cisbaikal_LNBA), is found substantially only among Yeniseian-speaking groups and those known to have admixed with them. Ancestry from the second, Yakutian Late Neolithic–Bronze Age (Yakutia_LNBA), is strongly associated with present-day Uralic speakers. We show how Yakutia_LNBA ancestry spread from an east Siberian origin ∼4.5kya, along with subclades of Y-chromosome haplogroup N occurring at high frequencies among present-day Uralic speakers, into Western and Central Siberia in communities associated with Seima-Turbino metallurgy: a suite of advanced bronze casting techniques that spread explosively across an enormous region of Northern Eurasia ∼4.0kya. However, the ancestry of the 16 Seima-Turbino-period individuals—the first reported from sites with this metallurgy—was otherwise extraordinarily diverse, with partial descent from Indo-Iranian-speaking pastoralists and multiple hunter-gatherer populations from widely separated regions of Eurasia. Our results provide support for theories suggesting that early Uralic speakers at the beginning of their westward dispersal where involved in the expansion of Seima-Turbino metallurgical traditions, and suggests that both cultural transmission and migration were important in the spread of Seima-Turbino material culture.

## Main

Long-distance similarities in language and shared material culture spanning thousands of kilometers across North Eurasia in the Early and Middle Holocene have been suggested to reflect a not only short-range neighbor-neighbor interactions, but also mobility in individuals’ lifetimes^1,2^. These similarities across the forest zone of North Eurasia (also known as the taiga belt) have prompted diverse theories, ranging from extreme diffusionism (such as the “Circumpolar Stone Age” ^3^), to rapid longitudinal transmission of “ideas, materials, and peoples” through the taiga belt, to latitudinal contacts facilitated by major northward-flowing rivers such as the Irtysh, Ob’, Yenisei, and Lena.^a^

Uralic languages—spoken today in Central Europe (Hungarian), around the Baltic Sea (Finnish, Estonian and Saami), in Eastern Europe (Komi, Udmurt, Mari, and the Mordivinic languages Moksha and Erzya), and western, central, and far northern Siberia, including the Taimyr Peninsula (Khanty, Manis, Selkup, Nenets, Enets and Nganasan)—are one such Trans-continental connection. However, intense debate remains about the geographic location of the homeland of the Uralic languages, the time frame of their dispersals, and the extent to which their ancient speech communities are discernible in the archaeological record. Castrén in the 19^th^ century proposed an origin in the Altai-Sayan mountains, but later scholars preferred locations further west: between the Ob’ and the Yenisei in West Siberia, in Eastern Europe near the confluence of the Volga and Kama Rivers, or even near the Baltic Sea ^4^. Genetic analysis has shown that all present-day Uralic-speaking populations (except for Hungarians) differ from their Indo-European speaking neighbors in having substantial Siberian-associated ancestry (ranging from ∼2% in Estonians to almost all the ancestry of Nganasans), mirrored in uniparental markers by a high frequency of Y-chromosome haplogroup N lineages originating in Siberia^5^. Time transects of ancient DNA showed that this ancestry was intrusive in Europe, arriving after ∼3.5kya in Karelia ^6^ and ∼2.6kya in the Baltic region ^7^ in the regions where Uralic languages are now spoken. However, while genome-wide ancestry from Yamnaya steppe pastoralists has been identified as a “tracer-dye” that can highlight population movements associated with the spread of the Indo-European languages, no corresponding ancestry (or ancestries) have been identified in the ancient DNA record that may a similar role to highlight an analogous set of movements for Uralic populations. Efforts to discern these patterns are made difficult by sampling gaps in ancient DNA, combined with the disruptive effects of migrations in the last few thousand years associated with the spread of Indo-European, Turkic and Mongolic languages that have made it difficult to reconstruct reliable population histories based on patterns of variation in present-day people^8,9^.

We screened samples from 209 individuals from across Northern Eurasia from the Mesolithic (∼11kya) to the Bronze Age (∼4.0 kya) using in-solution enrichment for more than 1.2 million single nucleotide polymorphisms (SNPs), and present genome-wide data from 181 of them that passed quality controls (Methods; Extended Data Figure 2). We generated 76 new radiocarbon dates from these samples. However, North Eurasian forager remains are often strongly affected by freshwater reservoir effects that can cause dates to be overestimated by as much as a millennium ^10^; thus, chronological inferences must also be guided by archaeological context (SI Section IX). These new data fill in key gaps in space and time from the Volga-Ural region to the Lena River Valley in Eastern Siberia (Figure 1). We highlight five major findings (Box 1; Extended Data Figure 1), and provide the detailed evidence for each claim in the next sections.

**Figure 1.**
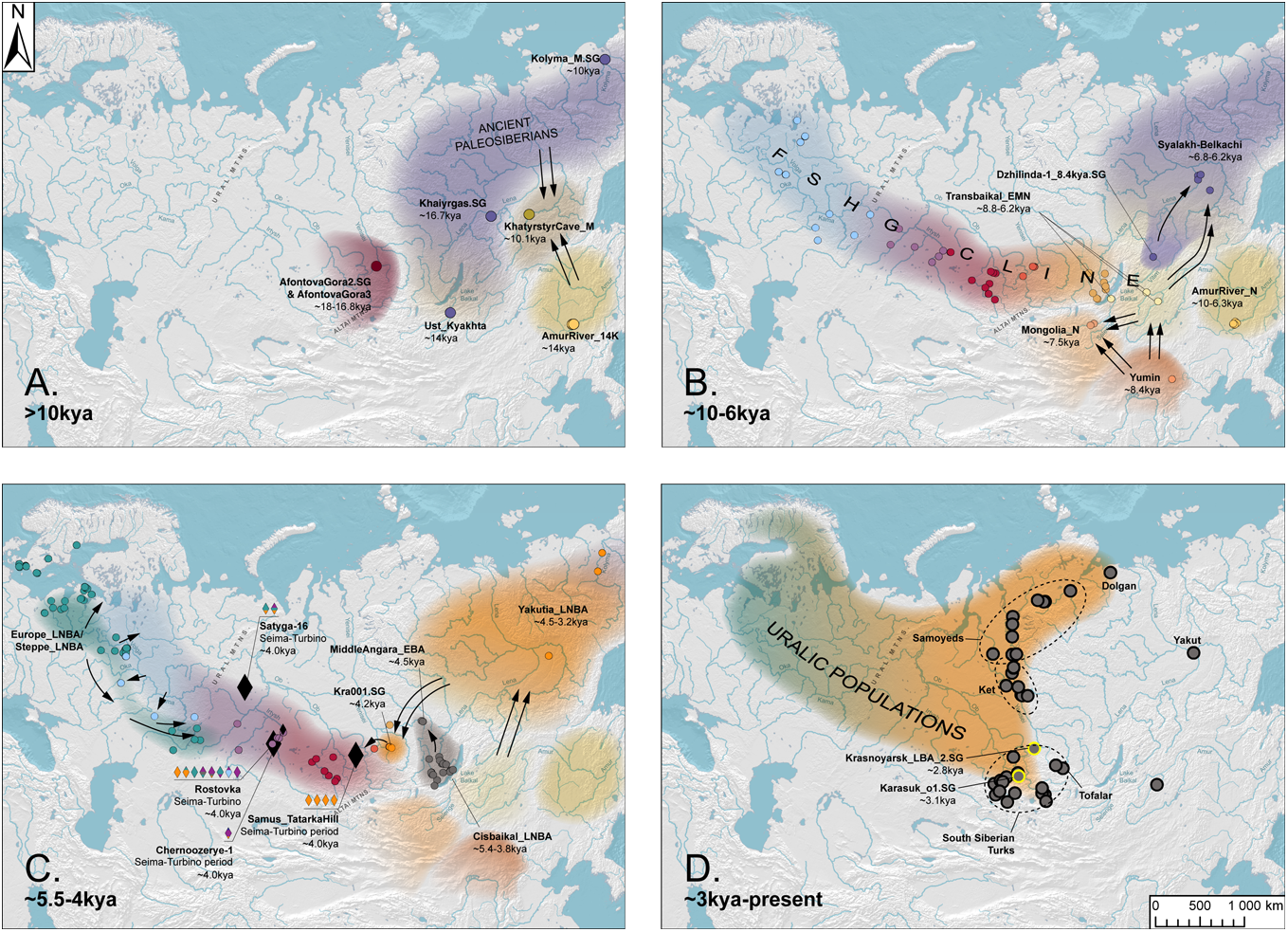
The Forest-Steppe Hunter-Gatherer Cline and its legacy through admixture in ancient northern Eurasia. (Top) Map. Sampling locations of all individuals that fall on the FSHG cline, as well as of selected samples not falling on the cline that are mentioned in the text. (Center) PCA. We project ancient data and present-day shotgun data onto variation from 122 genotyped present-day Eurasian and Native American populations selected to have minimal sub-Saharan African and Oceanian admixture (SI section III). Note the occurrence of a continent-spanning Forest-Steppe Hunter-gatherer cline (FSHG cline), as well as a cline for Uralic populations that stretches from European and Bronze Age Steppe populations to present-day Nganasans, Yakutia_LNBA individuals, and the individuals from the ST-period site of Tatarka. (Bottom) Admixture Proportions. The first row of bar graphs presents qpAdm estimates of ancestry related to four sources (*Russia_AfontovaGora* for ANE ancestry, *China_AmurRiver_LPaleolithic_19K* for East Asian ancestry, *Russia_HG_Elshanks* for EHG, and *Romania_IronGatesMesolithic* for WHG) for all populations on the FSHG cline. 84 out of 93 have passing models (p>0.01); populations that do not have a star above the admixture bar plot. In these cases we show the model with the highest p-value. The second row of graphs display estimated admixture proportions for all 8 sources listed in the legend (i.e., expanding to include *Tarim_EMBA1, Altai_N_old, Iran_GanjDareh_N* and *CHG*); a pink dot above the admixture bar plot indicates that all passing *qpAdm* models have *Tarim_EMBA* included in the sources. A cross above the bar plot indicates that individuals from the population were used as a source (*Altai_N_old; Russia_MiddleVolga_Elshanka_Chekhalino_4_10kya*). For the two populations from the Kuznetsk-Altai Neolithic culture, this produces a 100% A*ltai_N* admixture proportion by default. For the Elshanka individual, we replaced the EHG source with Russia_Veretye_Mesolithic.SG.

### Box 1.

**1)** A Pleistocene population related to Native Americans that we call “Ancient Paleosiberians” (APS), mixed with two East Asian ancestry sources— “Inland Northeast Asian-related” and “Amur Basin-related” —to contribute to later populations throughout Siberia.
**2)** Early pottery users in a latitudinal belt across Northern Eurasia in the early-to-mid Holocene (∼10-5kya) including the forest-steppe and the forest belt immediately adjacent to it, constitute a continent-spanning east-west genetic cline comprising Eastern European-Hunter-Gatherer (EHG), Ancient North Eurasian (ANE), and East Asian ancestries. This *“Forest-Steppe Hunter-Gatherer cline”* (*FSHG* cline), began to dissolve due to population replacements in the Mid-Holocene (∼5kya).
**3)** A genetic turnover ∼5.4kya saw the emergence of a population, *Cisbaikal_LNBA* to the west of Lake Baikal. This ancestry spread from the Cis-Baikal region to the Yenisei region by the end of the Late Bronze Age ∼3.1kya. Today, the presence of this ancestry is strongly associated with Yeniseian-speaking populations and those likely to have mixed with them historically. We suggest that this ancestry was likely dispersed by population movements that spread Yeniseian languages.
**4)** A genetic turnover by ∼4.5kya saw the emergence of a population in Northeast Siberia, *Yakutia_LNBA*. Today, this ancestry tends to be the only East Asian ancestry present among Uralic-speaking populations, a striking feature not shared by any other ethnolinguistic grouping. This ancestry appears in the Krasnoyarsk region along the Upper Yenisei, far to the Southwest of Yakutia, by ∼4.2kya alongside subclades of Y-chromosome haplogroup N found at high frequency among present-day Uralic-speaking males as far as the Baltic Sea. We suggest that this ancestry was likely dispersed by population movements that spread Uralic languages.
**5)** Individuals associated with the *Seima-Turbino* (ST) *phenomenon*—an archaeological term for the sudden appearance of a distinct suite of bronze artifacts across an enormous expanse of Northern

## Origins and Genetic Legacy of Ancient Paleosiberians

To provide an overview of the genetic variation in this region through space and time, we performed unsupervised analyses using Principal Component Analysis (PCA) and ADMIXTURE (Fig. 1; SI Section IV). These reveal that individuals from a broad belt of pottery-using foraging cultures in the forest-steppes and southern forest zone that stretched across Northern Eurasia from the Early-to-Middle Holocene (∼10-5kya) form a vast genetic gradient stretching over 7,000 km that no longer exists today, which we term the Forest-Steppe Hunter-Gatherer (FSHG) cline, linking genetically West Eurasian hunter-gatherers from the Eastern European plain to genetically East Asian hunter-gatherers of the Transbaikal region via a chain of genetically intermediate populations (Fig. 1, Extended Data Fig. 4,5,6; SI Section IV; here, we use the term East Asian to refer to the East Eurasian lineage in East and Southeast Asians but not Australasians or South Asians ^11^). However, many groups deviate from the cline. Populations from deeper into the forest zone of Northeast and Central Siberia (i.e, in a region bounded by the Yenisei River in the west, the Bering Straits in the east, and Lake Baikal in the south) are genetically similar to eastern FSHG populations in being intermediate between ANE (Ancient North Eurasians ^12^) and East Asian populations (in PCA: Extended Data Fig. 4; in qpAdm: SI Data S2 Table 1), but deviate away from the FSHG cline in being shifted towards present-day Native Americans and Arctic populations on both sides of the Bering Straits in PCAs (Fig. 1; Extended Data Figure 6, 7). Individuals from the Cis-baikal region from later than ∼5.4kya (i.e., in the Late Neolithic and Bronze Age) also deviate from the cline in other dimensions (Extended Data Figure 6).

To investigate these differences, we performed statistical analyses of population structure to infer a sequence of ancestry changes in Northeast and Central Siberia and the Baikal region (from ∼17kya to ∼4.0kya) that strongly coincides with changes in material culture. We defined genetic groups across all 100 Holocene individuals (newly reported as well as previously-published) from Northeastern Siberia and adjacent parts of East Asia (Cis-Baikal and Transbaikal regions), by clustering them using f_4_-statistics with a procedure first used in ^14^ (SI Section V; Data S1 Table 4 & 5), and then assigning labels using archaeological and geographic information. The procedure defined seven genetic cluster (five multi-member and two single individuals), which we call, in chronological order: MiddleLena_KhatystyrCave_M_10.2kya (a newly-reported early Holocene individual from Khatystyr Cave along the Middle Lena in Southern Yakutia, of unclear cultural affiliation ^15^; M stands for Mesolithic), MiddleVitim_Dzhilinda1_M_N_8.4kya (at the Mesolithic-Neolithic boundary, from the Dzhilinda-1 site along the Vitim river attributed to the Ust-Yumurchen culture ^14,16,17^; N stands for Neolithic), Transbaikal_EMN (individuals spanning ∼8.8-6.2 kya from the Early and Middle Neolithic Kitoi culture east of the Baikal; EMN stands for Early/Middle Neolithic), Cisbaikal_EN (individuals spanning ∼8.0-6.6kya from the Early Neolithic Kitoi culture west of the Baikal), Syalakh-Belkachi (individuals spanning ∼6.8-6.2 kya from the Early Neolithic Syalakh and Middle Neolithic Belkachi cultures in the Middle Lena Basin), Cisbaikal_LNBA (individuals spanning ∼5.1-3.7kya from the Late Neolithic and Bronze Age Serovo, Isakovo, and Glazkovo cultures west of the Baikal), and Yakutia_LNBA (∼4.5-3.2kya, associated chiefly but not exclusively with the Late Neolithic and Bronze Age Ymyyakhtakh culture). The remaining individuals, which we excluded from the following analyses, were genetically intermediate between these seven population groupings and plausibly represent admixtures between the groups we analyzed. Of these genetically defined groupings, only the Cisbaikal_EN and Transbaikal_EMN populations from the Kitoi culture along the forest-steppe belt lie on the FSHG cline, but the other groups do not (Fig. 1; Extended Data Fig 6, 7). By adding three older individuals from the region--MiddleLena_Khaiyrgas_16.7kya ^14^, Selenge_Ust-Kyakhta_14kya (∼14kya ^18^), and Kolyma_M_10.1kya (∼10.1kya ^19^)—to the seven populations defined by our procedure, we produce a ten-member temporal transect of genetic populations in Northeast Siberia and adjacent parts of East Asia, stretching from ∼17kya to 4kya, described in further in Extended Data Fig 2.

To investigate the deep population history of Eastern Siberia, we used qpAdm to model each target population in the ten-population transect as derived from ones preceding them or contemporary to them with an “outgroup rotation” technique (SI Section VI.A), whereby groups not used as sources for modeling are included as outgroups (i.e., included in the “right” set) to increase statistical leverage to reject models^20^. We found one or a small number of similar passing models for every target population in this ten-population transect (p>0.05). *qpAdm* analyses testing many hypotheses have the potential to produce false positives^21^, so our qpAdm procedure should be treated as a model-rejection and not as a model-selection procedure. Nevertheless, the broad interpretation of the qpAdm results in the following section is supported by patterns in simple f_4_ statistics involving distal populations likely to be relevant to the peopling of Northeast Siberia (Fig. 2A & B), and we emphasize results at this level of precision (SI Section VI.A.iv).

**Figure 2.**
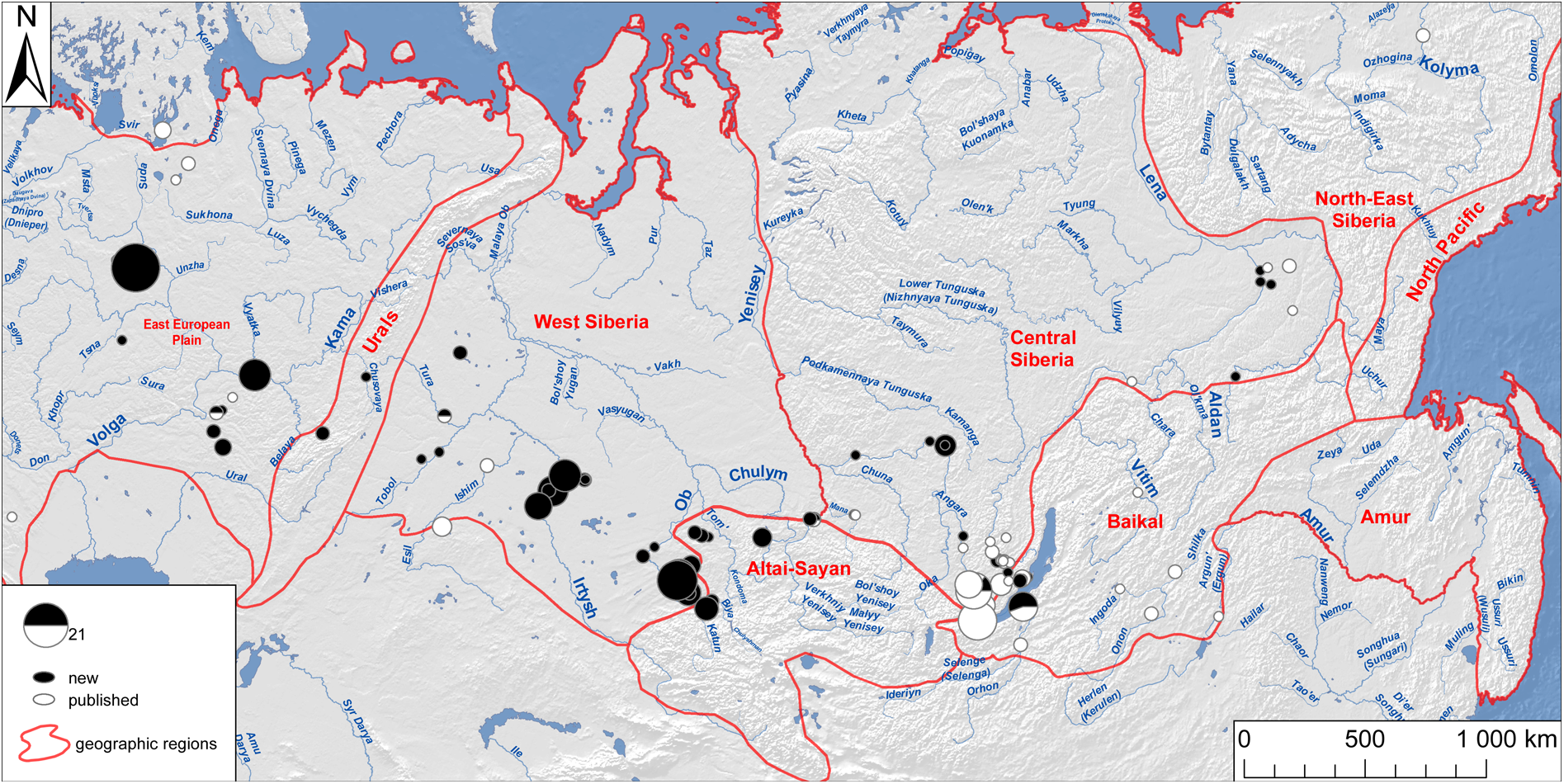
Middle Holocene populations and the admixture events that formed them. (A, Top) Statistics of the form f4(Ethiopia_4500BP, Target, China_Paleolithic, Yana_UP) on the x-axis, plotted against f4(Ethiopia_4500BP, Target, X, Yana_UP) on the y-axis, where X are ancient Native Americans or ancient populations from the Bering Straits. The Target population’s position on the y-axis is proportional to its ratio of ANE and East Asian ancestry. Notice that the following four ancient populations: *Kolyma_M_10.1kya, MiddleVitim_Dzhilinda1_M_N_8.4kya*, *Syalakh-Belkachi,* and *Yakutia_LNBA* are consistently shifted to the left of the curve formed by all other populations, indicating that these four populations share more drift with ancient Bering Straits populations than other populations with similar rations of ANE and East Asian ancestry (be they populations on the *FSHG* cline, Cisbaikal_LNBA, Selenge_Ust-Kyakhta_14kya, or MiddleLena_Khaiyrgas_16.7kya). Results with a more complete set of ancient Native Americans and Beringians are presented in Fig. S7. **(B, Bottom Left)** Statistics of the form F4(Ethiopia_4500BP, X, China_NEastAsia_Inland_EN, China_AmurRiver_Mesolithic 14K) on the x-axis, plotted against F4(Ethiopia_4500BP, X, China_Paleolithic, MA1_HG) (left top) and F4(Ethiopia_4500BP, X, China_Paleolithic, Peru_Laramate_900BP) (left bottom) on the y-axes, where X are ancient populations in Northeast Asia and Siberia. These statistics indicate that substructure exists within the East Asian ancestry of admixed groups in Northeast Asia and Siberia, with differentiation between an Inland East Asian-related source (proxied by the *Yumin* hunter-gatherer under the population label *China_NEastAsia_Inland_EN*), and an Amur-River-related source (represented by the population *China_AmurRiver_Mesolithic_14K*). Populations from the Amur River region always have high affinity with *China_AmurRiver_Mesolithic_14K*, while populations on the Mongolian Plateau and the Baikal area share more affinity with the Yumin hunter-gatherer. The earliest strongly East Asian individual in Northeast Siberia, the Mesolithic *MiddleLena_KhatystyrCave_M_10.2kya*, is extremely Amur-River-related, but other Northeastern Siberian groups rich in APS ancestry, such as *MiddleVitim_Dzhilinda1_M_N_8.4kya, Kolyma_M_10.1kya*, and *Syalakh-Belkachi*, tend to be almost equally related to both Inland and Amur- related sources. The sole exception is *Cisbaikal_LNBA*, which is strongly Inland Northeast Asian-related. Affinity to *China_NEastAsia_Inland_EN* also increases in agriculturalist populations of the Yellow River Valley as one moves westwards, suggesting that Inland-East Asian-related populations were more widely distributed in western regions of continental East Asia. **(C, Bottom Right)** Schematic of population relationships in Northeast Asia and East Siberia. This diagram summarizes the inter-population relationships from ∼17kya to ∼4kya in our ten-member Siberian transect as deduced from qpAdm and presented in SI Section VI.A.iii and summarized in SI Section VI.A.iv. Some major features of this diagram are: 1) That the MiddleLena_Khaiyrgas_16.7kya population is a near-unadmixed representative of an “Ancient Paleosiberian” (APS) population with Native American affinities, 2) that APS ancestry persisted through two routes, 3) that the East Asian ancestry of Siberian populations derives from an Amur Basin-related source and an Inland East Asian-related source.

The term “Ancient Paleosiberians” (APS) was used to designate the ancestry found in the Kolyma_M_10.1kya individual ^19^. Here we broaden this term to designate a pre-Holocene population, admixed between ANE and East Asian ancestries, which gave rise to Native Americans and which also played a key role in the formation of all later groups in Northeast Siberia and along the Bering Straits (including in the ancestry of Kolyma_M_10.1kya itself). We find that that the oldest individual from our transect, the previously-published MiddleLena_Khaiyrgas_16.7kya.SG (from a site along the Middle Lena in Yakutia, attributed to the Dyuktai culture ^14,22^) is a nearly unadmixed representative of APS ancestry. When other pre-Holocene Eurasians are placed as outgroups in qpAdm, MiddleLena_Khaiyrgas_16.7kya and Native Americans (here proxied by high-coverage ancient Peruvians ^23^) are the best representatives of each other’s ancestry, and MiddleLena_Khaiyrgas_16.7kya may be modeled as descending completely from a Native American-related source (SI Section VI.A.ii.a, VI.A.iv.b)—making it the first individual sampled from the ancient DNA record of Eurasia to have this ancestry in near-unadmixed form. Such Native-American-related APS ancestry may have spread with the “Beringian tradition” of lithics rich in microblades and wedge-shaped cores in the Northeast Siberian Upper Paleolithic ^24^. It admixed with East Asian ancestry but still persisted at a high levels in the later two individuals Selenge_Ust_Kyakhta_14kya (from just south of Lake Baikal on the Selenge River, from a site with lithics in this same tradition ^18^) and Kolyma_M_10.1kya (from a site close to the Bering Straits, ^19,25^; SI Sections VI.A.ii.b-c, VI.A.iii.b,VI.A.iv.c). However, these two individuals share even more drift with Native Americans when compared with Khaiyrgas, implying that the APS source in them was more closely related to Native Americans than the APS source in Khaiyrgas. Further west, admixture between APS and an ANE-related source formed the ANE-rich Altai_N population on the FSHG cline (associated with the Neolithic Kuznetsk-Altai culture of the Upper Ob and Altai foothills) by the early Holocene (∼9kya; SI VI.A.ii.d-e, VI.A.iii.c-d)—a population that may have mediated the spread of APS and ANE ancestries to groups further west in the FSHG cline, discussed in the next section.

Prior work has shown that “Neosiberian” ancestry related to East Asians increased at the expense of APS ancestry in Northeast Siberia over the Holocene, from ∼10kya to ∼3kya ^19^. We further infer that the East Asian ancestry in Northeast Siberians can be traced to at least two distinct sources: Inland Northeast Asian-related ancestry which we proxy in our analyses by the Yumin individual (under the population label China_NEastAsia_Inland_EN) from Inner Mongolia (∼8.4kya ^26^), and Amur River-related ancestry, represented by pre-Holocene hunter-gatherers of the Amur Basin (∼14kya, under the population label China_AmurRiver_14K ^11^). The oldest sample in our Siberian transect with high East Asian and low APS ancestry, MiddleLena_KhatystyrCave_M_10.2kya, from a site along the Aldan tributary that empties into the Middle Lena, had extremely strong affinities to Amur River hunter-gatherers (SI VI.A.ii.c, VI.A.iii.b, VI.A.iv.e, Fig. 2B), but subsequent Early Holocene populations from further south (including the Kitoi- associated Transbaikal_EMN and Cisbaikal_EN fat ∼8.8-6kya, and Mongolia_N_North at ∼7.5kya from the Mongolian Plateau) have increasing affinities to the Inland Northeast Asian source (SI VI.A.ii.e, VI.A.iii.d, VI.A.iv.e, Fig 2B). We find that all populations of the FSHG cline East of the Altai, including Cisbaikal_EN, plausibly derive their East Asian ancestry from the Transbaikal_EMN population on which the cline terminates—a source intermediate in affinity between the Inland and Amur-related sources. But non-FSHG populations high in East Asian ancestry (such as Cisbaikal_LNBA or Holocene foragers from the Amur River Basin) deviate from this pattern. Thus, FSHGs and non-FSHGs may be differentiated by their mix of East Asian ancestries (SI VI.A.iv.d).

By the mid-Holocene in the Cis-Baikal region, ancestry from the Cisbaikal_LNBA cluster (∼5.1-3.7kya) replaced that of the Cisbaikal_EN cluster (8-6.6kya), in a population turnover coinciding with the transition from the Early Neolithic Kitoi culture to the Late Neolithic and Bronze Age Serovo, Isakovo and Glazkovo cultures. The incoming Cisbaikal_LNBA population is much higher in APS ancestry than Cisbaikal_EN and is distinctive in deriving its East Asian ancestry from a strongly Inland-related source (Fig. 2B, SI VI.A.ii.f, VI.A.iii.a). While the only fitting qpAdm models have the APS ancestry coming from an Ust-Kyakhta_14kya-related population, we caution against overinterpreting this result due to the time gap separating the two populations. Despite their elevated APS ancestry, Cisbaikal_LNBA does not have increased shared drift with populations in the Americas or the Bering Straits when compared to other groups with similar ratios of ANE to East Asian ancestry (such as Ust-Kyakhta_14kya itself, Khaiyrgas_16.7kya.SG or FSHG populations from the Upper Yenisei region; Fig.1 center, Fig. 2A). Instead, it has high shared drift with populations from Central Siberia and especially the Yenisei River Basin (Extended Data Figure 8; SI Figure 77), and the analyses we present in upcoming sections show that ancestry from Cisbaikal_LNBA may be the first of two conduits by which APS ancestry persisted into present-day populations (i.e. it is a “Route 1” population, Fig. 2C, 3B; SI Section VI.A.iv.d & VI.D).

**Fig. 3.**
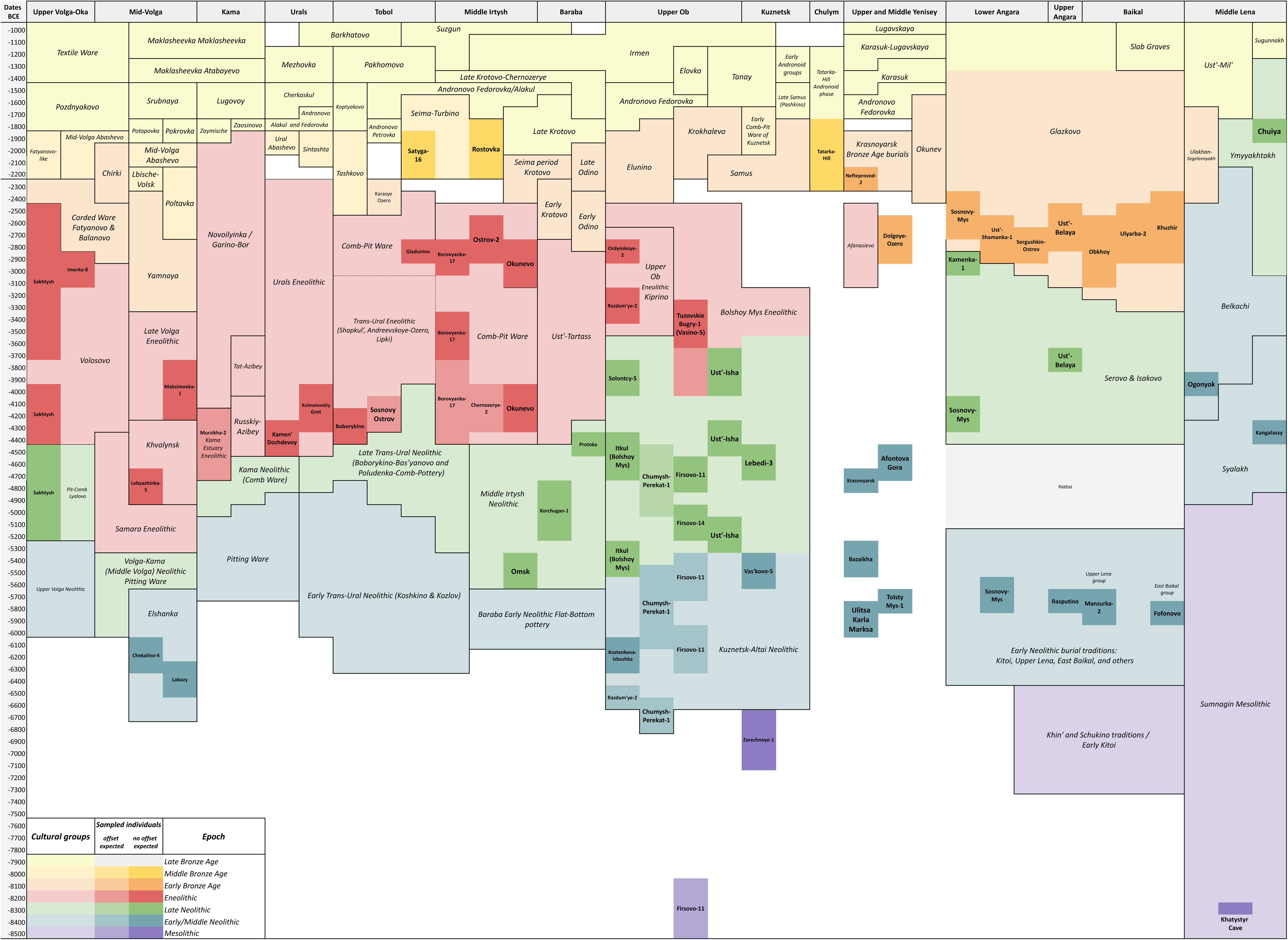
| **The Contribution of Yakutia_LNBA to Modern and Historic Period Admixed Inner Eurasian (AIEA) populations. (A) PCA of f_4_-statistics.** The input into the PCA are f_4_-statistics of the form f4(*Ethiopia_4500BP, AIEA, AG3, EA*) where EA are all East Asian population from the following set: [*China_AmurRiver_N, Mongolia_N_North, Transbaikal_EMN, Cisbaikal_LNBA, Yakutia_LNBA*]. PC1 is strongly correlated with proportions of West Eurasian versus East Asian ancestry, while PC2 highlights genetic similarity to *Yakutia_LNBA* (left). At any proportion of West Eurasian ancestry, ancient and present-day Uralic-speaking populations are especially shifted in PC2 in the direction suggesting disproportionate *Yakutia_LNBA-*relatedness for their levels of East Asian ancestry. In contrast, PC3 highlights genetic similarity to Cisbaikal_LNBA (right), and Yeniseians, South Siberian Turks, Samoyeds, and two Late Bronze Age East Asian outliers along the Upper Yenisei (from ∼3.0-2.9kya, RISE497.SG/Russia_Karasuk_o1.SG, and RISE554.SG/Russia_UpperYenisei_Krasnoyarsk_LBA_o1.SG, potentially from the Lugavskaya culture) are the most shifted in PC3 in the direction suggesting disproportionate shared drift with Cisbaikal_LNBA ancestry, a result corroborated by ADMIXTURE in Fig. 3B (see next). The biplot displaying loadings at left (PC1 vs PC2) has been scaled to 2x its original size, while that at right (PC2 vs PC3) has been scaled to 10x original size. **(B) The distribution of Cisbaikal_LNBA ancestry among present-day populations.** Populations with >4% Cisbaikal_LNBA ancestry are highlighted in large black dots. The positions of the two ancient outliers high in Cisbaikal_LNBA ancestry, potentially from the Lugavskaya culture of the Minusinsk Basin, are highlighted in two white stars. **(C) qpAdm and ADMIXTURE results.** The top row displays qpAdm results for AIEA populations (SI Section VI.C.ii). One orange dot above the bars for an AIEA population indicates that all East Asian ancestry in that population can be modeled as deriving from *Yakutia_LNBA*; two orange dots indicate that—additionally—all passing models include *Yakutia_LNBA* among the sources. We also performed a second set of qpAdm with Cisbaikal_LNBA among the references and sources (SI Section VI.D.ii); a grey dot indicates that all passing models for that population include *Cisbaikal_LNBA* in the sources. The bottom row displays ADMIXTURE results. Notice how almost all Uralic populations have their East Asian ancestry near-exclusively assigned to the Yakutia_LNBA component, and how Yeniseians, South Siberian Turks and Samoyeds are the only populations to have appreciable levels of the Cisbaikal_LNBA-related component. Lastly, the two Late Bronze Age outliers along the Upper Yenisei (from ∼3.0-2.9kya, RISE497.SG/Russia_Karasuk_o1.SG, and RISE554.SG/Russia_UpperYenisei_Krasnoyarsk_LBA_o1.SG) are the only individuals to have the overwhelming majority of their ancestry assigned to the Cisbiakal_LNBA component, in line with f4-statistics (see previous).

North of the Baikal region, in Northeast Siberia and its adjacent regions, the strongly Amur-Basin-related individual MiddleLena_KhatystyrCave_M_10.2kya was followed by the strongly APS-related MiddleVitim_Dzhilinda1_M_N_8.4kya. The increase in APS ancestry possibly results from admixture from a Kolyma_M_10.1kya-related source (SI VI.A.ii.d, VI.A.iii.c). APS ancestry then declines with admixture from East Asian sources over the Early and Middle Holocene, in a set of population turnovers coinciding with transitions between archaeological cultures. The first turnover occurred with the appearance of the Syalakh-Belkachi population (∼6.8-6.2kya, with ∼20% admixture from an East Asian source from the Baikal region admixing into the preceding MiddleVitim_Dzhilinda1_M_N_8.4kya). This was followed by a second turnover with the appearance of the Ymyyakhtakh-associated Yakutia_LNBA population (∼4.5-3.2kya, with ∼50% admixture from Transbaikal_EMN admixing into the preceding Syalakh-Belkachi population). Strikingly, this sequence of four populations from the Lena and Kolyma River Valleys in far Northeast Siberia: Kolyma_M_10.1kya, MiddleVitim_Dzhilinda1_M_N_8.4kya, Syalakh-Belkachi, and Yakutia_LNBA—includes all the ancient Siberian individuals that are shifted towards Native Americans and Bering Straits populations in PCAs. This is corroborated by f_4_-statistics that show they share more drift with ancient and present-day Bering Straits populations than other groups with similar ratios of ANE to East Asian ancestry (e.g. Khaiyrgas_16.7kya, Ust_Kyakhta_14kya, all FSHG populations east of the Altai, or Cisbaikal_LNBA) do (Fig 2A). Using qpAdm, we infer that the third member of this sequence—the Syalakh-Belkachi population—made a major (∼70%) contribution to populations of the Arctic Small Tool Tradition of North America (SI VI.B.i, ^27^, e.g. the Paleo-Eskimo Greenland_Saqqaq.SG, also reported in ^14^); such Paleo-Eskimo-related ancestry (which is by extension Syalakh-Belkachi-related) then persisted into all later ancient and present-day groups such as Eskimo-Aleuts, Chukotko-Kamchatkans and Yukaghirs on both sides of the Bering Straits (SI VI.B.ii). This may account for these Northeast Siberians’ unique trans-Beringian genetic connections and may represent the second major route by which APS ancestry is mediated into present-day populations (i.e. they are “Route 2” populations, Figure 2C, SI Section VI.A.iv.d). We also find that these affinities do not extend to ancient Athabaskans (SI VI.B.iii-iv), an observation that is incompatible with earlier inferences from our group suggesting that Athabaskans and Paleo-Eskimos derive ancestry from the same migration from Eurasia ^28,29^. Instead, this supports suggestions of multiple late Holocene migratory movements into the Americas from Eurasia, ^19,30,31^ further detailed in our Discussion.

## An Early Holocene Forest-Steppe Hunter-Gatherer Cline

From ∼10-4kya, all 150 newly reported and 81 previously published individuals from the North Eurasian forest-steppe and the Southern forest zone adjacent to it are part of the FSHG cline. This cline is visible in ADMIXTURE (Fig. 1 bottom, third row; Extended Data Fig. S4-6), and in PCA (Fig. 1, center) as an arc connecting pottery-using Eastern European foragers to their counterparts in the Transbaikal region through a chain of intermediate populations. The center of this cline lies close to the Ancient North Eurasian (ANE) individual Afontova-Gora 3 (AG3), and also Tarim_EMBA (an early Bronze Age population from the Tarim Basin that may descend from hunter-gatherers of Central Asia, ^32^) (Fig. 1, center; Extended Data Fig. S4-6).

To prepare FSHG individuals for formal genetic analyses, we grouped them, first by site, then (for those with direct radiocarbon dates) by time-period, and finally according to their genetic similarity in unsupervised analyses (ADMIXTURE: SI Section IV; Extended Data Fig. 10; and PCA: Fig 1, Extended Data Fig. S4-6; resulting labels are listed in Fig. 1, bottom & SI Data S1, Table 3). In each site, those individuals for which we have no radiocarbon dates are merged into a generalized grouping that lacks a time-period designation in its group label. The great majority of the resulting FSHG “genetic populations” can be modeled in qpAdm as admixtures of four ancestries (84 out of 93 populations P>0.01): Western Hunter-Gatherer ancestry (WHG, represented by hunter-gatherers from Serbia ∼10kya, ^33,34^), EHG ancestry (represented by a newly-published individual, I6413, from ∼10kya from a site of the Elshanka culture, the oldest pottery-using culture in Eastern Europe), ANE ancestry (represented by AG3 ^12^), and East Asian ancestry (represented by a ∼19 kya individual from the Amur Basin ^11^; Figure 1 bottom, distal qpAdms; SI Section VI.E.i, Data S2 Table 1). Starting from the western end of the FSHG cline, hunter-gatherers from the Baltic to the Urals, attributed to (in the west) the Early Neolithic Elshanka, Late Neolithic Pit-Comb Ware/Lyalovo, and Eneolithic Volosovo cultures, and (in the east) to the Samara Eneolithic, Kama Estuary Eneolithic and Eneolithic Ural cultures, have mostly EHG-related ancestry, with low levels of WHG-related ancestry, in line with previous findings ^35,36^. Eastwards across the Urals, in populations of the Tobol and Middle Irtysh Early Neolithic and of the succeeding circle of Eneolithic West Siberian cultures using Comb-Pit Ware pottery, EHG ancestry admixed with ANE ancestry and low levels of East Asian ancestry. These populations are genetically similar to adjacent Botai-attributed individuals from northern Kazakhstan (∼5.4-5.1 kya) ^8^. Further east, individuals from the Altai foothills and the upper Ob, from the Kuznetsk-Altai culture spanning the Early Neolithic and Eneolithic (from sites such as Firsovo-11, Tuzovskie-Bugry-1 and Ust’-Isha), can be modeled as two-way admixtures of ANE and East Asian ancestry. This continues into individuals from Neolithic sites of the Upper Yenisei and Kan River Basin from sites without clear cultural attribution, where ANE ancestry declines and East Asian ancestry increases. The gradient extends into the Kitoi culture of the Baikal region, through the previously discussed Cisbaikal_EN population, to terminate in the Transbaikal_EMN population that is almost completely East Asian in ancestry.

Because the latest individual in the archaeogenetic record that has near-unadmixed ANE ancestry (AG3 at ∼16kya ^12^) is much older than any individual or population comprising the FSHG cline, we attempted to find proximal sources for the ANE in ANE-rich FSHG populations (i.e., all FSHGs west of the Altai). We find that two potential proximal sources successfully account for all their ANE in qpAdm (even with AG3 in the references; Figure 1 bottom, second row; SI VI.E.ii): first, a source comprised of the oldest individuals (∼9kya) from the Kuznetsk-Altai Neolithic (Altai_N_old); and second, the Tarim_EMBA population (∼4kya)^32^. Tarim_EMBA postdates FSHG populations, but ADMIXTURE and PCA suggest gene flow between a source related to them and FSHGs in West Siberia (Fig. 1, center; Fig. 1, bottom, third row; Extended Data Fig. 10). Outgroup rotation also shows that models without a Tarim_EMBA-related source tend to fail for West Siberian FSHGs when Tarim_EMBA are placed in the references (Fig. 1, bottom, center row; SI Section VI.E.ii). It has been suggested that populations genetically related to Tarim_EMBA lived in Central Asia before the arrival of pastoralism during the Bronze Age ^32^; ancestry from this source may have contributed to FSHG populations in West Siberia, explaining our results, a scenario made even more plaubsible by the recent discovery of an individual with this hypothesized profile from Mesolithic Tajikistan ^37^. Therefore, two sources may have admixed into the ANE-rich populations in the center of the FSHG cline: a Tarim_EMBA-like population from Central Asia, and a population like that of the later Kuznetsk-Altai Neolithic of the Altai region.

The FSHG cline was marked by genetic stability from ∼10kya to ∼5kya, providing evidence for continued genetic exchange between neighboring populations along the cline during this period. However, the archaeogenetic record shows that it fragmented in the Mid-Holocene following migrations from both West and East (Extended Data Fig. 1). From the West, these migrations brought Steppe_EMBA ancestry associated with Yamnaya pastoralists, followed by Europe_LNBA ancestry associated with the expansion of the Fatyanovo culture into the Volga basin, and then the closely-related Steppe_MLBA ancestry in populations of the Sintashta culture and of the Andronovo area ^36,38,39^. From the East, these migrations introduced Cisbaikal_LNBA ancestry at ∼5.4 kya. Subsequently, admixture between Steppe_MLBA and East Asian ancestries gave rise to admixed groups across much of Northern Eurasia and Central Asia, culminating in the multiple genetic clines found among present-day Turkic-, Mongolic-, Tungusic, Uralic- and Yeniseian-speaking populations, who retain little ancestry from the FSHG cline ^8^ (Fig 3C, bottom row; Extended Data Fig 4; Fig S5). To evaluate the contribution that FSHG and East Siberian populations made to the genetic formation of later populations across Northern Eurasia, we genetically analyzed a set of AIEA (Admixed Inner Eurasian) populations—our term for ancient and present-day Uralic, Turkic, Mongolic, Tungusic and Yeniseian-speaking populations plus pastoralists of the Late Bronze Age and Iron Age, including Scythians, Sarmatians, and Xiongnu ^40–43^. We replicate the finding that FSHG populations contribute little to the genetic formation of later AIEA populations. Instead, the two latest populations of our East Siberian transect, Cisbaikal_LNBA and Yakutia_LNBA, played an important role in the genetic formation of Yeniseian- and Uralic-speaking populations respectively.

## Cisbaikal_LNBA ancestry is a tracer-dye for prehistoric mobility associated with the spread of Yeniseian languages

The Cisbaikal_LNBA population (∼5.1-3.6 kya) is very rich in APS ancestry, occupies a distinct position in PCA (Extended Data Fig 7C) and has a uniquely strong affinity to Inland Northeast Asians (Fig. 2B, SI Section VI.A.ii.a, VI.A.iv.a). While other APS-rich groups from Northeast Siberia are more closely related to Arctic populations on both sides of the Bering Straits (i.e., “Route 2” populations), Cisbaikal_LNBA is unique among APS-rich groups in sharing more genetic drift with present-day populations of the Yenisei Basin, suggesting that it may be a conduit by which APS ancestry persisted into present-day populations of that region (“Route 1”, Fig. 2A, Fig. 3B; Extended Data Fig. 8; SI VI.A.iv.a).

In ADMIXTURE, the Cisbaikal_LNBA component appears at significant levels only in the Yeniseian- speaking Kets, in Samoyeds, and in Siberian Turkic-speaking populations all from the Yenisei Basin (Fig 3B & C, bottom row). In a PCA of f_4_-statistics designed to be sensitive to different types of East Asian ancestry, Yeniseian-, Samoyedic- and South Siberian Turkic-speakers are shifted systematically in the direction produced by increased shared drift with Cisbaikal_LNBA over other East Asian ancestries (Fig. 3A; SI Section VI.D.i). In qpAdm, models for all Yeniseian, Samoyedic and Siberian Turkic populations consistently fail when Cisbaikal_LNBA is retained in the references; for them, Cisbaikal_LNBA ancestry is required as a source in all passing models (Fig. 3C, top row; SI Section VI.D.ii, Fig. S12; Extended Data Fig 1D)—a link not found among other ethnolinguistic groupings. This link is also reflected in Y- chromosomal haplogroups: sequences ancestral to the haplogroup Q-YP1691, found at very high frequencies in Kets and at lower frequencies in neighboring Samoyedic and Siberian Turkic populations such as Selkups and Tuvans ^35,44–46^ have been recovered in the archaeogenetic record thus far only from male members of the Glazkovo culture that belong to the Cisbaikal_LNBA population (SI Section VIII).

Ethnolinguistic and historical data indicate that South Siberian Turks have assimilated Yeniseian speakers, beginning with the arrival of the Yenisei Kyrgyz to the southern Yenisei Basin in the 6^th^ century CE and continuing through the Russian colonization ^47–51^. The remaining Siberian Turkic languages—Yakut and Dolgan—are spoken today by populations whose ancestors migrated within the last two millennia from the region where South Siberian Turks live today, providing a plausible explanation for why these groups also carry Cisbaikal_LNBA ancestry ^52^. Further north, ethnographic records indicate that Samoyedic-speaking groups have had cordial relationships with Yeniseian speakers, with much intermarriage ^48^.

In our analysis of AIEA, we unexpectedly found that two previously-published ^40^ East Asian outliers (RISE497.SG or Russia_Karasuk_o1.SG, & RISE554.SG or Russia_UpperYenisei_Krasnoyarsk_LBA_o1.SG) from the Late Bronze Age (∼3.0-2.9kya) in the Minusinsk Basin along the Upper Yenisei, can be modeled in qpAdm with very high Cisbaikal_LNBA ancestry (∼85-95%, Fig. 3B; SI VI.D.iii). f_4_-statistics and ADMIXTURE suggest that they have by far the strongest genetic affinity to Cisbaikal_LNBA among all modern or ancient AIEAs (Fig. 3A, C; Extended Data Fig. 5). These genetic outliers (labeled as being from the Karasuk culture in the original publication, but which our archaeological research indicates may instead be assigned to the Lugavskaya culture; SI Section II.G), show that people with very high levels of Cisbaikal_LNBA ancestry were present along the Upper Yenisei by the Late Bronze Age ∼3.0kya, near the geographic region where Cisbaikal_LNBA is maximized in populations today (Fig. 3B)—a geographic distribution which may be explained by partial genetic persistence of such groups into the Iron Age and Medieval period. Furthermore, the Ket themselves are known to have reached their current position along the Middle Yenisei in a recent Northward expansion, while all six other documented (and now extinct) Yeniseian languages were found in regions further south along the Yenisei River, in regions that are now occupied by South Siberian Turkic populations where Cisbaikal_LNBA ancestry peaks in the present day (Fig. 3B) ^53,54^. These results suggest that, at least within the last three millennia, Cisbaikal_LNBA ancestry has come to be strongly correlated with the movements of Yeniseian speakers and their closest ethnic contacts.

Individuals of the Cisbaikal_LNBA population first appear in the Late Neolithic Serovo and Isakovo cultures in the Cis-Baikal region ∼5.4kya. Virtually all individuals from the succeeding Glazkovo culture as late as ∼3.8kya also fall into this genetic cluster (SI VI.B). This ancestry subsequently appeared among individuals in the forest zone of the Middle Angara (which drains out of Lake Baikal and leads into the Yenisei) in the Early Bronze Age ∼4.8kya (SI VI.D.iii), around when Glazkovo pottery appeared in the region ^55,56^. It has been suggested that the distribution of Yeniseian hydronyms ^57,58^ points to a Yeniseian homeland in the Cis-Baikal region several millennia before present ^54^, an inference compatible with our results. Cisbaikal_LNBA ancestry may thus trace the movements and contacts of Yeniseian speakers even further into prehistory.

## Yakutia_LNBA ancestry is a tracer-dye for prehistoric mobility associated with the spread of Uralic languages

We next characterized a group of individuals we term Yakutia_LNBA, who have a distinct position on PCAs (Fig. 1, Extended Data Fig. 6) and are among the APS-rich populations sharing elevated drift with Arctic populations on either side of the Bering Straits (Figure S7). The oldest Yakutia_LNBA individual dates to ∼4.5kya in the Lena River Valley, and individuals from this group persist to at least ∼3.2kya in that region. Yakutia_LNBA can be modeled as a product of admixture between the preceding local Syalakh-Belkachi population with admixture from populations of the Kitoi culture of the Transbaikal (∼50% Syalakh-Belkachi + ∼50% Transbaikal_EMN; SI VI.A.ii.f); all males of Yakutia_LNBA carry subclades of haplogroup N also found in the Transbaikal_EMN population (SI VIII). Such genetic links are consistent with theories of the origin of the Ymyyakhtakh culture ^59^, to which all individuals in the Yakuta_LNBA cluster are assigned—except a single individual recovered from the Krasnoyarsk-Kansk forest-steppe far to the southwest of the Lena River Valley (Kra001.SG from the Nefteprovod-2 site ∼4.2kya ^14^), in a location otherwise dominated by populations from the FSHG cline. The existence of this individual suggests that populations with Yakutia_LNBA ancestry had dispersed from Northeast Siberia to the forest-steppes North of the Altai-Sayan region shortly before the start of the second millennium BC, paralleled by the appearance of Ymyakhtakh pottery in this region at that time^55^.

We show that Yakutia_LNBA-related ancestry is strongly associated with all ancient and present-day Uralic-speaking populations. In ADMIXTURE at K = 18, a component maximized in Yakutia_LNBA appears that peaks in Nganasans among present-day populations and accounts for almost all the East Asian ancestry in Uralic-speaking populations. Non-Uralic AIEAs have either no Yakutia_LNBA, or other East Asian components in addition to Yakutia_LNBA, such as those maximized in Mongolia_N_North, China_Yellow_River_MN, or Amur Basin foragers (Figure 3C, bottom row; Extended Data Fig. 10).

In a PCA of f_4_-statistics, we find that Uralic speakers are consistently shifted in a direction indicating increased affinity towards Yakutia_LNBA compared to other East Asian ancestries (Fig. 3A; SI Section VI.C.i). Furthermore, we find using a second set of f_4_-statistics that, at any level of East Asian admixture, the AIEA group with the highest affinity to Yakutia_LNBA over other East Asian ancestries is always a Uralic-speaking population (Extended Data Fig. 9; VI.C.i). Lastly, qpAdm analysis shows that models for Uralic speakers—without exception—require Yakutia_LNBA as a source, which can almost always account for all their East Asian ancestry (Fig. 3C, top row; SI Vi.C.ii). This contrasts with other ethnolinguistic groupings among AIEAs, such as Scythians, Sarmatians and their predecessors in the Late Bronze Age (who are often modeled as deriving all their East Asian ancestry from a Mongolia_N_North-related source) or Turkic, Mongolic and Tungusic populations (who have additional ancestry from East Asian agriculturalists or Amur Basin hunter-gatherers).

These observations suggest that the genetic formation of Uralic-speaking populations involved an episode of gene flow from a population with Yakutia_LNBA ancestry. In contrast, the genetic formation of non-Uralic AIEAs either did not involve populations with Yakutia_LNBA ancestry, or involved additional episodes of gene flow from populations of the Eastern Steppes or East Asian agriculturalists that did not affect the ancestors of present-day Uralic-speakers. Additional evidence for a link between Yakutia_LNBA and Uralic populations lies in uniparental markers: Male individuals of the Yakutia_LNBA cluster carry Y-chromosomes under subclades of haplogroup N that are present at high frequency in present-day speakers of Uralic languages ^5^.

## The Seima-Turbino Phenomenon involved the movement of both people and ideas, and was implicated in the initial westward dispersal of Yakutia_LNBA ancestry

Populations from Western Siberia to the Upper Volga as late as the MBA (associated with cultures such as the Fatyanovo, Sintashta and Andronovo) do not show any Yakutia_LNBA ancestry ^5–7^, but present-day Uralic populations from the same regions do, suggesting an east-to-west spread of this ancestry into Western Siberia and Eastern Europe in the MBA at the earliest (∼4kya). This process, which eventually reached the shores of the Baltic Sea, would have involved Yakutia_LNBA ancestry partially replacing Steppe_MLBA ancestry (that was predominant in populations of the Sintashta, Srubnaya, Andronovo, and related cultures of the Late Bronze Age in Eastern Europe and Central Asia), and Europe_LNBA ancestry (that was predominant in populations of the Corded Ware horizon, including the Fatyanovo culture ^36,38,39^). This process was potentially accompanied by the dispersal of Y-haplogroup N, which— despite its very high frequency in Uralic populations in Eastern Europe and West Siberia today (∼20-100%) ^5,60^—is completely absent from the archaeogenetic record of these regions prior to the arrival of Yakutia_LNBA ancestry from the East ^6^. In this section, we present new genomic data supporting the hyothesis that the earliest stages of this westward dispersal of Yakutia_LNBA ancestry, found together with subclades of Y-chromosome haplogroup N, may have taken place within cultural contexts that produced the Seima-Turbino (ST) phenomenon.

The ST phenomenon refers to the sudden appearance of a very similar suite of bronze artifacts manufactured with advanced casting techniques that spread rapidly (in a century or so) into many cultures spanning a vast region of Northern Eurasia, from China to the Baltic Sea. Archaeologists credit this trans-cultural phenomenon ^61–67^ for the introduction of metallurgy into Eastern Eurasia and the dissemination of advanced casting methods for tin bronze into Europe ^65,68^. ST items are noted for their technological sophistication and aesthetic refinement (Fig 4C); most are weapons, but some are striking objects of probably ritual or religious significance. Most items are isolated finds from sites attributed to a wide range of cultures within the distributional area of ST artifacts, but many others also come from standalone necropolises across Western Siberia and Eastern Europe (Figure 4A): large complexes of burials and, more often, cenotaphs (empty ritual graves) with extremely rich collections of ST artifacts and casting molds. The distribution of ST objects, with massive concentrations in exceptional ceremonial sites that punctuate an enormous trans-cultural scatter of metal items, has fueled speculation about the social nature of the ST phenomenon, as well as the identity of their bearers ^61,62,69,63,67^. So far, the only conclusive evidence found for the manufacture of ST bronze artifacts (such as casting molds) are from residential sites of metal-using fisher-foragers in the Ob-Irtysh basin and the region between the Upper Ob and Upper Yenisei—an extraordinary cultural association that has occasioned much comment ^63,70,71^.

**Fig. 4.**
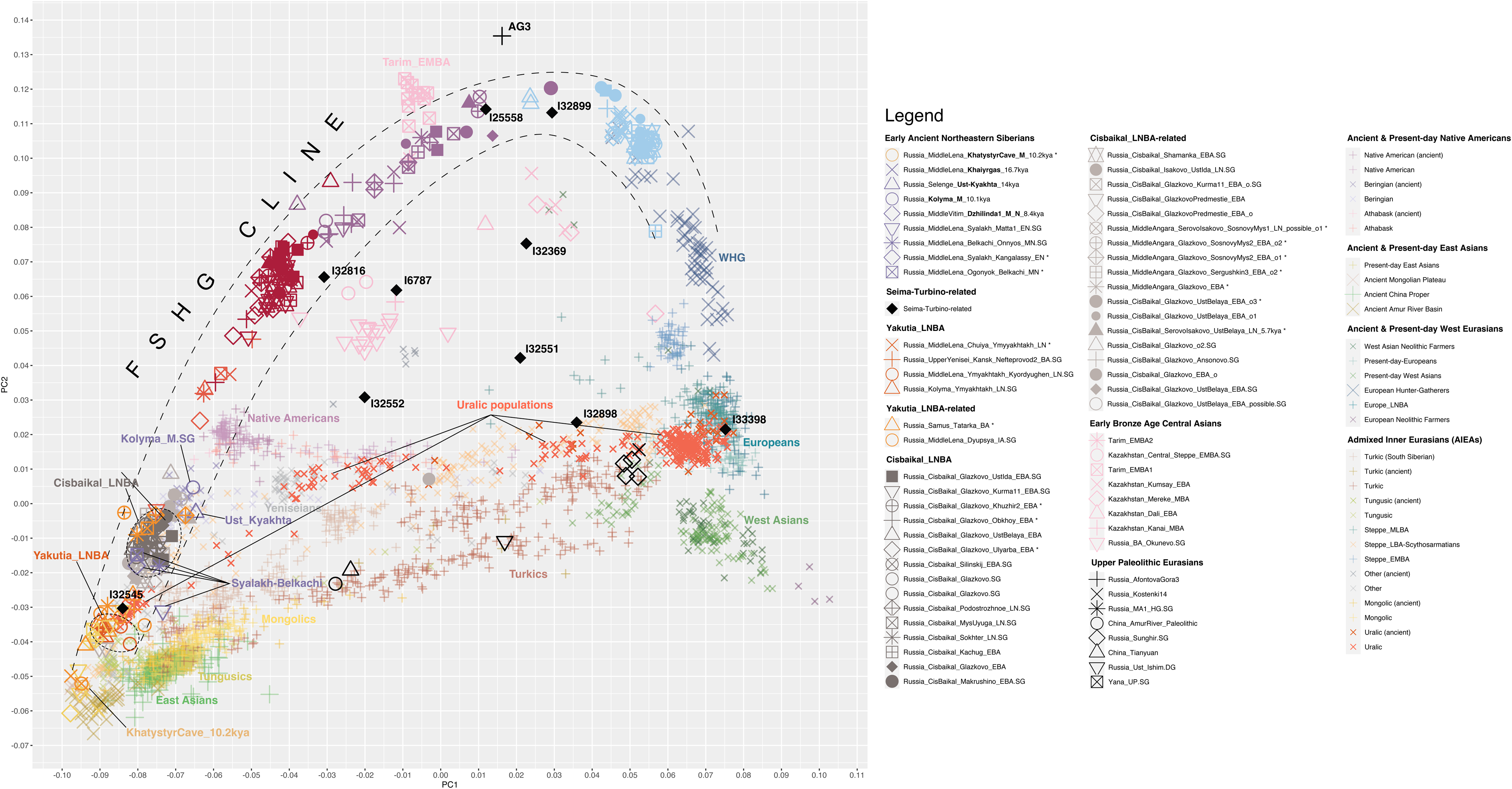
| **The Seima-Turbino Phenomenon, and Genetic Analysis of Seima-Turbino period individuals. (A) A map of Seima-Turbino sites and finds.** The sites of Chernoozerye-1, Rostovka, Satyga-16 and Tatarka, from which archaeogenetic samples were sequenced for this study, are marked in red circles. Important Seima-Turbino sites are numbered: 1, Seima. 2, Reshnoe. 3, Turbino. 4, Kaninskaya Cave. 5, Satyga-16. 6, Rostovka. 7, Samus-4. 8, Shaitanskoe Ozero. 9, Tatarka. **(B) ADMIXTURE and qpAdm proportions for all 11 of our Seima-Turbino-period individuals.** 8 individuals are from Rostovka, 2 from Satyga-16, 1 from Chernoozerye-1 and 4 from Tatarka. The **top row** of bars displays a selection of qpAdm models using a distal set of admixture sources, with the simplest passing (p>0.01) qpAdm model always displayed (if multiple equally simple qpAdm models pass at the p>0.01 level, the one with the highest p-value is displayed). This set of qpAdm are performed with the same set of sources and references as for the AIEA qpAdms in Fig 3B. No model passes for one individual (I32899) at the p>0.01 level, indicated by an asterisk above the bar. The **center row** of bars displays qpAdm results for a more proximal set of sources, with the individuals from Tatarka used as the source for Yakutia_LNBA ancestry and populations from Late Neolithic or Eneolithic Western Siberia (between the Urals and the Altai) as the source for FSHG ancestries; all ST individuals are successfully modeled under this setup. Compared to the setup for the qpAdm in the top row, *Yakutia_LNBA* has been added to the references. Ancestry from Late Neolithic or Eneolithic hunter-gatherers from West Siberia is ubiquitous among ST individuals (SI Table VII-B). Ancestry from the population at Tatarka also suffices to account for all the Yakutia_LNBA ancestry of the ST individuals even when Yakutia_LNBA is left among the references. As in figure 3B, samples that can be modeled with their East Asian ancestry derived completely from a *Yakutia_LNBA*-related source (Yakutia_LNBA in the distal model or *Russia_Samus_Tatarka_MBA* in the proximal model) are highlighted by a single orange dot above their qpAdm bar charts, while those that require such a in every passing model are highlighted by two such orange dots; all individuals from Tatarka, Chernoozerye-1 and Satyga-16 as well as 4/9 individuals from Rostovka require such ancestry. Additionally, one individual (I32816) requires *Cisbaikal_LNBA* ancestry among the sources when *Cisbaikal_LNBA* is added to the references as part of outgroup rotation, as highlighted by the gray dot; the model displayed for this individual is the simplest passing model that contains both *Cisbaikal_LNBA* and *Yakutia_LNBA* ancestry among the sources in a setup with *Cisbaikal_LNBA* added to the outgroup rotation. Lastly, in both sets of qpAdm, the individual (I6787) from Chernoozersky-1 requires contribution from a source from far West of the Urals (*WHG* ancestry) which is present in all passing models (SI Table VII-B). All details for the qpAdm analyses performed on these ST individuals can be found in SI section VII. The **bottom row of bars** displays ADMIXTURE proportions at K=18. These results are strikingly congruent with qpAdm. **(C) Characteristic Seima-Turbino artifacts.** 1. Double-bladed dagger with a ring-shaped pommel, robbery find, unknown provenance (probable Omsk region or Rostovka). 2. Double-bladed dagger with a horse figurine on the pommel, an accidental find near Shemonaikha, East Kazakhstan. 3., 5., 7. Crook-backed knives with figurines on pommels: 3. from Seyma; 5. from Elunino-1, burial 1, 7. from Rostovka, burial 2. 4. Scapula-shaped celt with goat image, Rostovka, cluster of finds near burial 21. 6. double-bladed plate dagger with a double elk-head figurine pommel, an accidental find near Perm’ (probably in association with the Turbino site). 8. Top of staff with a horse figurine, an accidental find near Omsk. 9a. & 9b. Single-ear long spearhead with a relief figurine of a *Felidae* predator (tiger or mountain leopard) on the socket (9a. the spear tip,10b. the detail of the socket), an accidental find near Omsk.

We generated genome-wide data from 16 individuals from four ST-period burial sites, three in the Ob-Irtysh Basin between the Ural and Altai mountains in Western Siberia, and one in the region between the Upper Yenisei and Upper Ob, just north of the Altai-Sayan mountains. Radiocarbon dating and archaeological context indicate they date to a tight interval around 4.0 kya (SI Section IX; Data SI Table 4). From the Ob-Irtysh Basin, we sequenced 9 individuals from the ST necropolis of Rostovka, on the banks of the middle Irtysh, and 2 from the ST necropolis of Satyga-16, east of the Mid-Ural Mountains. We add to this samples from two sites that have less direct evidence of involvement with the ST phenomenon, but are cotemporaneous to it and that our genetic analyses suggest may be connected with it: one individual from the burial site of Chernoozerye-1^b^, located close to Rostovka, and four males from a previously undescribed burial site, Tatarka Hill^c^ on the Chulym tributary, between the Upper Ob and Upper Yenisei ^72^. In ADMIXTURE and PCA, the mostly male individuals from the ST necropolises of Rostovka and Satyga-16, and the individual from the ST-period burial of Chernoozerye-1, are extremely heterogeneous, harboring highly variable proportions of three major and two minor sources of ancestry that we describe below (Figure 4E, bottom row; Fig 1, center; Extended Data Fig. 4, 5). In contrast, the four ST-period individuals from Tatarka Hill are genetically homogeneous, very similar to Yakutia_LNBA, and indeed can be modeled with near-complete descent from Yakutia_LNBA in qpAdm (SI Section VI.C.iii).

Distal qpAdm shows that the three major ancestries at the genetically heterogeneous sites of Rostovka, Satyga-16 and Chernoozerye-1 are 1) ancestry from variable positions in the ANE-rich central section of the FSHG cline, 2) Steppe_MLBA ancestry, and 3) Yakutia_LNBA ancestry. These ancestries occur as unadmixed individual representatives of each ancestry type, or intermingled within admixed individuals (2-way: FSHG ancestry + Steppe_MLBA, or 3-way: FSHG ancestry + Steppe_MLBA + Yakutia_LNBA; Figure 4B, top row). While the sample size is small, we find it striking that both individuals from Satyga-16 further west are highly admixed (carrying all three ancestry types), contrasting with the tendency among people in Rostovka (4/9 single ancestry: 2 FSHG, 1 Yakutia_LNBA and 1 Steppe_MLBA; 1/9 2-way admixed; SI VII).

We next analyzed these same ST individuals with proximal qpAdm. We first find that the Yakutia_LNBA ancestry in Rostovka, Satyga-16 and Chernoozerye-1 can be sourced from a population related to the four unadmixed Yakutia_LNBA males at Tatarka Hill in the region between the Upper Yenisei and Upper Ob, with no evidence of additional mixture from Yakutia_LNBA populations in Northeast Siberia (Fig 4B, middle row). Such a genetic link is reinforced by the presence at Rostovka of the subclade of haplogroup N (N-L1026, in individual I32545, a male with near-unadmixed Yakutia_LNBA ancestry), which is also carried by all four males from Tatarka Hill (SI VIII). The specific subclade of the unadmixed Yakutia_LNBA individual from Rostovka (N-L1026 > Z1936) is widespread among present-day Uralic populations from East of the Urals to the Baltic Sea, but attains maximal frequencies (up to ∼40%) towards the west, in Baltic Finnic populations such as Finns, Veps and Karelians ^60^. Second, we find that the FSHG ancestry of ST individuals can be modeled as coming in large part from preceding, local Late Neolithic and Eneolithic populations of West Siberia (Fig 4E, middle row). This may suggest that individuals from Rostovka with unadmixed FSHG ancestry may derive from the metallurgical foragers of the nearby Odinovo and Krotovo cultures—the cultures best known to have engaged in the systematic casting of ST artifacts ^67^. However, some individuals require additional FSHG ancestry from EHG-related or Altai_N-related sources, suggesting that admixture also occurred from FSHG populations from further afield.

The three major ancestry sources are accompanied by two minor ancestries: non-Yakutia_LNBA East Asian ancestry, and WHG-related ancestry (Figure 4E). We find in both distal and proximal qpAdm that the ST-period individual from Chernoozerye-1 (I6787), located close to Rostovka, consistently requires a large fraction of WHG-related ancestry in fitting models, in addition to Yakutia_LNBA and FSHG-related ancestry (SI VIII)—a result also corroborated by ADMIXTURE (Fig 4E, bottom row). This implies recent ancestry from at least as far west as the Baltic Sea ^73^, in a remarkable case of an individual whose recent ancestry traces to at least three hunter-gatherer source populations from widely-separated regions of Eurasia. Similarly, two individuals from Rostovka, I32816 and I33369, have ancestry from far to the east, in the former case definitively from a Cisbaikal_LNBA-related source, possibly from foragers of the contemporaneous Glazkovo culture.

Such genetic heterogeneity suggests that ST artifacts in at least Western Siberia were manufactured, exchanged, and saw their dispersal within some set of sociocultural institutions or contexts that could integrate people from multiple populations from across a potentially continent-spanning network into social groups, such that they were interred together at single sites. This social or institutional context must have been able to support their admixed descendants, while sustaining—for a considerable time— continued contact and intermarriage with the geographically distant and culturally disparate source populations, to account for the complex multi-way admixtures we find in single ST-period individuals. Our sampling captures several snapshots of this ongoing social process, which created a distinctive pattern of human mobility that might be a genetic correlate to what archaeologists have found for the ST, which is a slow decay in artifact similarity across geographic distances unusual among cultural groups of the period^61^.

## Discussion

Linguistic transmission, especially in large-scale societies, need not involve the movement of people or genetic admixture. An observation that populations speak related languages need not imply that they will genetically resemble each other more than populations that do not ^74^. However, linguistic transmission across community boundaries in smaller-scale prehistoric societies is likely to have required at least some degree of human mobility that might be visible as genetic admixture, because language dispersal and shift in such contexts is most plausibly accompanied by substantial movements of people ^75^. Such patterns suggest that Yakutia_LNBA ancestry may have spread in episodes of human mobility that were associated with the prehistoric dispersal of Uralic languages, in the same way that the appearance of Yamnaya/Steppe_EMBA ancestry may be correlated with migrations responsible for the expansion of the Indo-European languages (a “tracer-dye” ^38^). Likewise, Cisbaikal_LNBA ancestry may be connected to the spread of Yeniseian languages.

Yeniseian languages are related to the Na-Dene languages in North America under the Dene-Yeniseian hypothesis ^76^. We investigated this connection by using qpAdm to distinguish sources of APS ancestry in ancient (<4kya) Siberian and American Arctic groups that have been connected to present-day Yukaghirs, Chukotko-Kamchatkans, Eskimo-Aleuts, and Athabaskans (SI Section VI.B). We find strong evidence that all such ancient groups show at least partial descent from Paleo-Eskimo-related populations (represented by Greenland_Saqqaq.SG), and by extension Syalakh-Belkachi and other “Route 2” populations, *except* Athabaskans from ∼1.1kya (SI VI.B.iiii-iv), consistent with some previous inferences ^19,30,31^, and contradicting other work, including from our own team ^77^. We instead find weak evidence that ancient Athabaskans may require a small quantity of ancestry from a population related to Cisbaikal_LNBA—genetic evidence for the linguistic hypothesis of a distinctive link between Yeniseian and Na-Dene languages. This suggestive result awaits corroboration with further sampling and more sensitive analytic methods.

There is little agreement among archaeologists about the social processes that drove the sudden spread of ST artifacts across such a wide range of cultures ^65,63,66,67,78,64^. The extreme genetic diversity and heterogeneity of ST necropolises in our sample is at the very least inconsistent with a concept of a single, homogeneous ST “people” (as proposed in e.g. ^69^, also see references in ^63^). Instead, it suggests that active participation of multiple social groups that were genetically and culturally distinct was an essential part of the ST phenomenon itself—a conclusion consistent with the rest of the inventory found in ST necropolises, such as pottery (which displays similarities to that produced by West Siberian foragers ^61,62,69,78^ and that of the forest-steppes around Tatarka Hill ^56^), artifacts of flint, bone, or jade (which displays similarities to those produced by cultures of far Northeast Siberia and Lake Baikal), and metal items from non-ST traditions (which may have been produced in the Sintashta and especially the closely-related Abashevo cultures) ^61,62,69,78^. These three sources of material culture closely parallel the three major genetic ancestries in our sample.

Also remarkable is the prevalence at ST sites of ancestry from multiple foraging populations across Northern Eurasia, from the Cis-Baikal to as far West as the Baltic Sea. This highlights the involvement of populations with a foraging mode of subsistence in this network, in line with recent work highlighting the wide and culturally transformative reach of metal exchange networks in the Bronze Age ^79–81^, as well as the oft-neglected sociopolitical dynamism and economic complexity found across hunter-gatherer populations ^82^.

Extreme diversity should not obscure the fact that the ST phenomenon was the context within which Yakutia_LNBA ancestry first dispersed westwards—almost to the Urals—for the first time, as suggested by the presence of such ancestry at all four ST sites we sampled (Tatarka, Rostovka, Satyga-16, and Chernoozerye-1). The presence of unadmixed Yakutia_LNBA ancestry at Tatarka Hill, to the west of the Kra001 individual from two centuries earlier, shows that the Yakutia_LNBA ancestry that penetrated onto the forest-steppe of this region, far Southwest of the Northeast Siberian forest zone, may have persisted to later contribute to the Yakutia_LNBA ancestry in ST necropolises. Intriguingly, Kra001 is just one out of a group of burials at Nefteprovod-2 dated from ∼4.2 to ∼3.9kya that show strikingly similar burial rites as the individuals at Tatarka Hill (SI Section II.F.iii.b-c). This confluence of cultural and genetic similarities suggests that a coherent and culturally distinctive population mediated the intrusion of Yakutia_LNBA ancestry westwards into the Krasnoyarsk-Kansk forest-steppes before 4.2kya, which then persisted in the region and later genetically impacted ST necropolises. Additionally, a material counterpart to the genetic link between some individuals from ST necropolises and populations of Northeast Siberia can be found in suits of armor made of bone plates, which have been found from the Glazkovo and especially the Ymyyakhtakh cultures. One set was buried with a male of the Yakutia_LNBA population mentioned in this study: N4a1.SG from the Kyordyughen site ^83^; others hail from the Upper Yenisei region where Kra001.SG and the Tatarka individuals were situated ^83^, and three were buried in Rostovka, one with an admixed male reported in our study (I32816 from Grave 33 ^61^) that bore both Yakutia_LNBA and Cisbaikal_LNBA ancestries.

Linguists have shown that Uralic languages have on the order of hundreds of Indo-Iranian loanwords directly inherited from the Proto-Uralic speech community or from the speech communities right after its breakup ^84,85^. The Indo-Iranian expansion has been linked to the spread of Steppe_MLBA ancestry from the Sintashta population in the Trans-Ural region into other parts of Central and West Asia (where it persisted into later populations on the steppes, the Iranian plateau and Afghanistan that are historically attested as being Iranic-speaking ^41–43,86–88^), and also further into South Asia, where it persists into present-day Indo-Aryan groups ^39^. Our findings from Rostovka and Satyga-16, showing contact and admixture between a Steppe_MLBA population (which, from archaeological considerations, is plausibly that of the Abashevo culture ^61^) and Yakutia_LNBA populations, provides an attractive hypothesis regarding the context in which this linguistic exchange could have first begun.

Despite the fact that Uralic languages, distributed from Western Siberia to Central Europe, are geographically separated from languages of the Eastern Steppes and far Northeast Siberia, linguists have discovered traces of ancient contact with Yukagiric and Eskimo-Aleut languages on the one hand ^89–92^, and members of the “Altaic” language area (Mongolic, Tungusic, and especially Turkic) on the other. In the latter case, the high level of typological similarity with Uralic languages has been repeatedly emphasized ^93,4,94,92,95^. As a resolution to these conundrums, linguists have suggested a recent eastern origin of the population associated with the later expansions of Uralic speakers (e.g., A “pre-proto-Uralic spoken further east… probably somewhere… near both Mongolia and the watershed area between the Yenisei and the Lena, possibly as recently as 3000BC” ^89^)—a scenario very compatible with our results.

A large team of Uralicists recently proposed that early Uralic populations were involved in the ST phenomenon, which catalyzed a rapid westward expansion ∼4 kya along river networks ^85^. While our results are consistent with this scenario, they cannot more precisely inform the question of the location of the Uralic homeland, as they are compatible with multiple hypotheses. A number of linguistic considerations (e.g., that proto-Uralic lacks a developed metallurgical vocabulary ^85,92^) may indicate that the proto-Uralic speech community slightly predates the ST. In any case, our analyses suggest that, if not proto-Uralic, at least early Uralic-speaking communities (as well as Indo-Iranian-speaking communities) were among the many groups that interacted to support the long-distance emigration and sociocultural exchanges that created the intriguing and widely-dispersed sets of material remains that form the ST phenomenon. We summarize our arguments linking our genetic results to the prehistoric dispersal of the Uralic languages in Box 2.

### Box 2.

Samples from the Late Bronze Age, Iron Age, and Medieval period of the Yenisei River Basin, the Altai region, Western Siberia, and Eastern Europe would be critical in connecting more ancient populations with historically attested Uralic-speaking groups. Such sampling would paint a clearer picture of the dispersals of later, admixed populations carrying Yakutia_LNBA ancestry, and might provide an opportunity to trace such movements to their origin. This would allow for a more precise determination of the archaeological identity of the proto-Uralic speaking community, and illuminate the relationship between it and the wider social world of the West Siberian Bronze Age within which it was possibly embedded.

## Methods

### Sampling of ancient individuals

Descriptions of the archaeological and cultural contexts for all ancient samples analyzed in this study, including their grave position within archaeological sites, their grave numbers and burial inventory, as well as archaeological publications describing the sites themselves (where available), are provided in Supplementary information section I. The skeletal samples are under the stewardship of the co-authors or museum collections that are listed in SI Data Table 2, column G. Samples may be accessed by their skeletal code listed in Supplementary Data S1 Table 2, along with the contact details of the persons stewarding the samples.

### Sampling of present-day individuals

We newly genotyped 229 present-day individuals from 10 ethnolinguistic groups using the Affymetrix Human Origins SNP array. All DNA samples were collected with informed consent for broad studies of population history and full public release of de-identified genetic data. All newly reported data are represented by co-authors of this study who were involved in sample collection.

### Ancient DNA data generation, bioinformatic processing, and quality control

Approximately 40mg of bone powder was collected in a clean room from skeletal remains, after which DNA was extracted using a protocol that retains short and damaged DNA fragments ^96,97^. The bone powder was collected from petrous bones, long bones, teeth, and ossicles. Individually-barcoded double ^98,99^ and single-stranded libraries ^100^ were built after incubation with uracil DNA glycosylase (UDG treatment, to reduce errors characteristic of ancient DNA damage,). We performed in-solution enrichment for ∼1.24 million SNPs (“1240k enrichment”, ^101^) and also enriched for the mitochondrial genome ^102^. Two rounds of enrichment were performed, after which sequencing was performed on the Illumina NextSeq 500 or HiSeq X 10 instruments.

The resulting read pairs were separated using library-specific barcode pairs or index pairs (for double-stranded and single-stranded libraries respectively) and merged prior to alignment. Read pairs were merged if 1) 15 or more base pairs (bp) overlap, 2) at most one mismatch occurred and base quality was at least 20, 3) at most three mismatches occurred and base quality was lower than 20. The resulting sequences were aligned to the human genome reference sequence (hg19) ^103^ and the mitochondrial RSRS genome using samse from bwa-v.0.6.1 ^104,105^. Duplicated reads were removed if they shared start and stop positions, orientation, and (for double-stranded libraries) barcode pairs. Analysis was performed on sequences at least 30 bp in length. We trimmed 2 bp from the ends of each read to reduce deamination errors. For each sample, we merged the sequences from all libraries. Most of the sequences used for population genetic analysis were constructed by randomly sampling at each SNP on chromosomes 1-22 and X, with a mapping threshold of 10 and base quality 20.

We flagged as “questionable” libraries that had evidence of contamination based on the upper bound of the match rate to the mitochondrial consensus sequence (assessed using contamMix v1.0-10, ^106^) being less than 95%; we also flagged as “critical” libraries if this value was less than 90% (Supplementary Data S1 Table 2). We also flagged as “questionable” males with evidence of high polymorphism on the X chromosome (lower bound of the 95% confidence interval for mismatch rate >1%), or as “critical” (if >5%), estimated using ANGSD v0.923 ^107^. For high-coverage contaminated individuals, we generated alternative sequences restricting to molecules showing signs of characteristic ancient DNA damage (designated by a suffix “_d” in the Genetic ID of the sample in Supplementary Data S1 Table 2).

For a subset of 15 individuals with high percentages of human DNA, we generated shotgun sequences (designated by the suffix “.SG” in Supplementary Data S1 Table 2) using the pre-enrichment libraries. We carried our sequencing on an Illumina HiSeq X Ten instrument. These shotgun sequences were used for analysis only in PCAs (Supplementary Materials Section III).

### Uniparental analysis

Mitochondrial haplogroups were determined with Haplogrep v2.1.1 ^108^. Y-chromosome haplogroups were evaluated using the methodology described in ^109^, section S5, using both targeted and off-target SNPs. Allelic status was determined by majority rule.

### ADMIXTURE and PCA

All relatives and shotgun sequences were excluded from ADMIXTURE analysis. For relative pairs or groups, the lower-coverage individual was excluded.

We used ADMIXTURE ^110^ after pruning SNPs with high missingness in plink (using option --geno 0.5 ^111^), after which 597,573 autosomal SNPs were retained. We used K=18 as the first K value where Yakutia_LNBA and Cisbaikal_LNBA were clearly separated from East Asian components characteristic of FSHG populations (e.g. the components maximized in Mongolia_N_North and AmurRiver_14K). Further details of our application of ADMIXTURE can be found in Supplementary Materials Section IV.

We pruned individuals from PCA analysis if they were found to be a first degree relative of another individual in the dataset with high coverage. PCA was performed using smartPCA in the EIGENSOFT package ^112^, using numoutlier: 0 and lsqproject: YES for three out of four PCAs. Further details on our PCAs can be found in Supplementary Materials Section III.

### qpAdms and F4 statistics

All F4-statistics were calculated using the *qpDstat* package of *ADMIXTOOLS* ^13^ with the f4mode: YES parameter. Further details of each set of F4 statistic calculations can be found where they are presented, in Supplementary Materials sections V.B, V.C, VI.A.iv, VI.C.i, and VI.D.i.

All qpAdm analyses were run using the R package ADMIXTOOLS2 ^113^. Precalculated *f_2_*-statistics, used to speed up the process of *f4*-ratio estimation central to *qpAdm*, were performed allowing for maximal missingness = .99 over multiple datasets. Further details for each set of qpAdm can be found in Supplementary Materials sections VI.A.i, VI.B, VI.C.ii, VI.D.ii, and VI.E.i.

### Relatedness and Runs-of-Homozygosity

We looked for kinship relationships between the individuals included in our study. We computed pairwise allelic mismatch rates in the autosomes by randomly sampling one DNA sequence at each ‘1240k’ polymorphic position, following the same strategy as in ^114, 115^ and ^116^, which is similar to that in ^117^. We then estimated relatedness coefficients *r* for each pair as in ^114^:

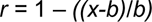

with *x* being the mismatch rate of the pair under analysis and *b* the base mismatch rate expected for two genetically identical individuals from the population under analysis, which we estimated by computing intra-individual mismatch rates. We also computed 95% confidence intervals using block jackknife standard errors over 5 Megabase (Mb) blocks.

## Data Availability

Bam files of aligned reads for the 181 newly published ancient individuals and 15 newly reported whole genome sequences can be obtained from the European Nucleotide Archive (accession no. to be made available upon publication). SNP array genotype data for 181 newly reported modern individuals can be obtained from a permanent link at the Dataverse repository (accession no. to be made available upon publication).

## Acknowledgments

We dedicate this paper to Oleg Balanovsky, who had a leading role in the collection of present-day samples newly reported in this study, and who would have been an author had he not died tragically in 2021. We thank Nicole Adamski, Rebecca Bernardos, Neil Bradman, Andrey A. Chizhevsky, Matthew Ferry, Eldar Idrisov Judith Kidd, Sergei V. Kuz’minykh, Kirsten Mandl, Pagbajabyn Nymadawa, Olga Poshekhonova, Harald Ringbauer, Ludmila Saroyants, Kristin Stewardson, Svetlana Tur, and Yuldash Yusupov, and Zhao Zhang, for wet laboratory or bioinformatic support, for sharing samples we analyzed, or for critical comments. A.A.T. acknowledges support from the Russian Science Foundation (project #22-18-00470). M.Z. acknowledges support from the Collaborative Research Grants Program #091019CRP2119 to Nazarbayev University. The research of G. Boeskorov was conducted within the framework of the scientific research program of the Diamond and Precious Metals Geology Institute, Siberian Branch of the Russian Academy of Science. Research by Alexander Stepanov, Elena Solovyeva and Viktor Dyakonov was carried out within the research program of the Institute of Archeology and Ethnography of the Siberian Branch of the Russian Academy of Sciences “North Asia in the Stone Age: cultural dynamics and ecological context” (FWZG-2022-0003). We are grateful to the Museum of the Institute of Plant and Animal Ecology UB RAS for sharing samples. P.F. was supported by Czech Science Foundation (project no. 21-27624S), and by the Czech Ministry of Education, Youth and Sports (program ERC CZ, project no. LL2103). P.V., R.P., and D.R. were supported by John Templeton Foundation grant 61220. P.V. and D.R. were supported by gifts from J.-F. Clin. D.R. was supported by National Institutes of Health grant HG012287; by the Allen Discovery Center program, a Paul G. Allen Frontiers Group advised program of the Paul G. Allen Family Foundation; and is an Investigator of the Howard Hughes Medical Institute. This article is subject to HHMI’s Open Access to Publications policy. HHMI lab heads have previously granted a nonexclusive CC BY 4.0 license to the public and a sublicensable license to HHMI in their research articles. Pursuant to those licenses, the author-accepted manuscript of this article can be made freely available under a CC BY 4.0 license immediately upon publication.

## Extended Data Figure Legends

**Extended Data Figure 1.**
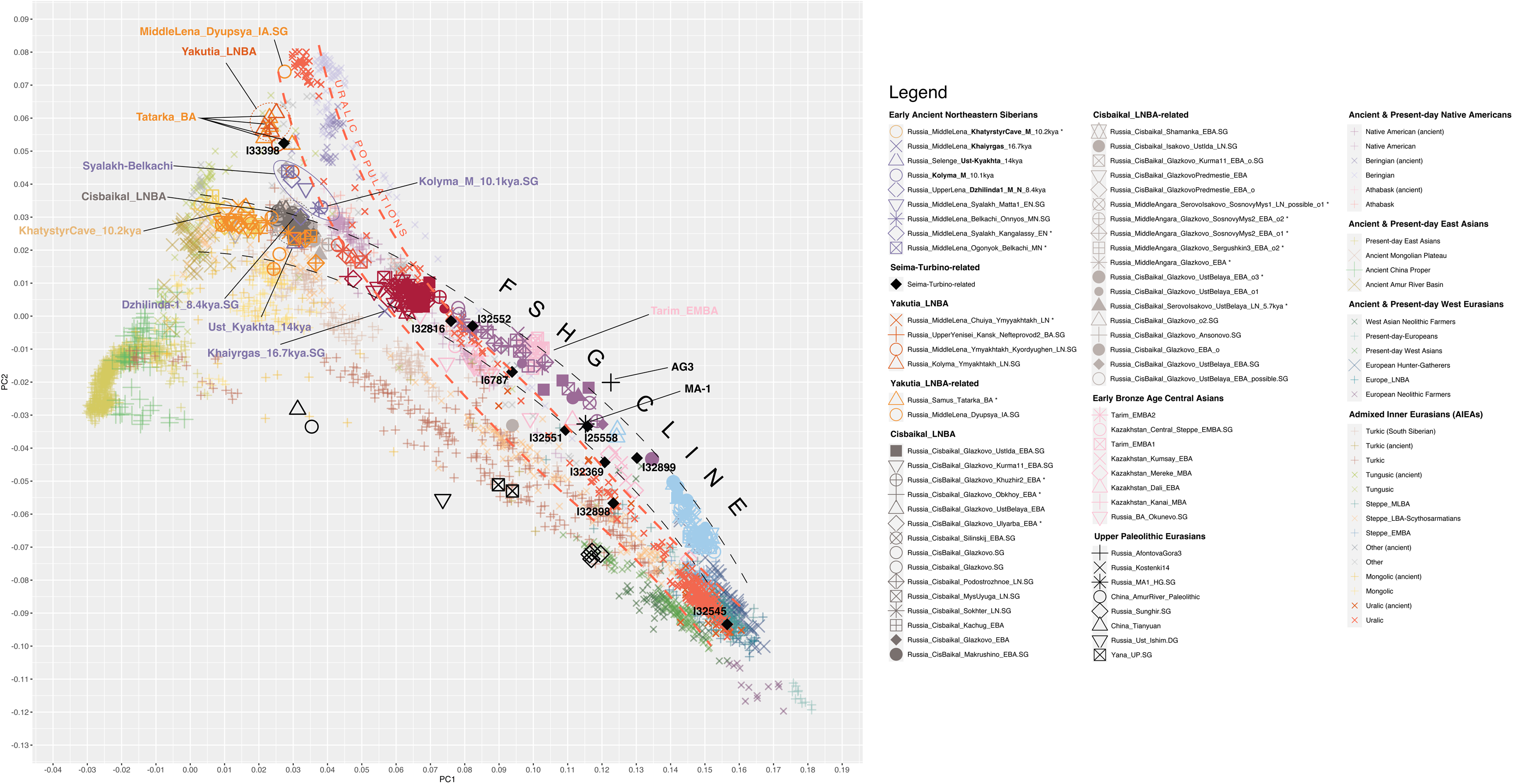
Summary of Genetic Changes Taking Place in Northern Eurasia. Panel A shows the widespread distribution of individuals with Ancient Paleosiberian (APS) ancestry in Siberia before the Holocene, >10kya. Panel B shows the formation of the FSHG cline by ∼10kya, and the formation of the population on its eastern terminus (Transbaikal_EMN) through admixture between Amur River and Inland East Asian ancestries. Panel C shows the emergence of CIsbaikal_LNBA and Yakutia_LNBA in genetic turnovers in the Cis-baikal and Northeastern Siberian regions in the Mid-Holocene, and the genetic diversity of Seima-Turbino period individuals ∼4.0kya. Panel D shows the genetic gradient between West Eurasian ancestry and Yakutia_LNBA formed by present-day Uralic populations, along with all locations from which present-day populations with Cisbaikal_LNBA ancestry have been sampled (grey dots ringed with black), alongside the geographic locations of two late Bronze Age/early Iron Age individuals (grey dots ringed with yellow) with >90% Cisbaikal_LNBA ancestry.

**Extended Data Figure 2.**
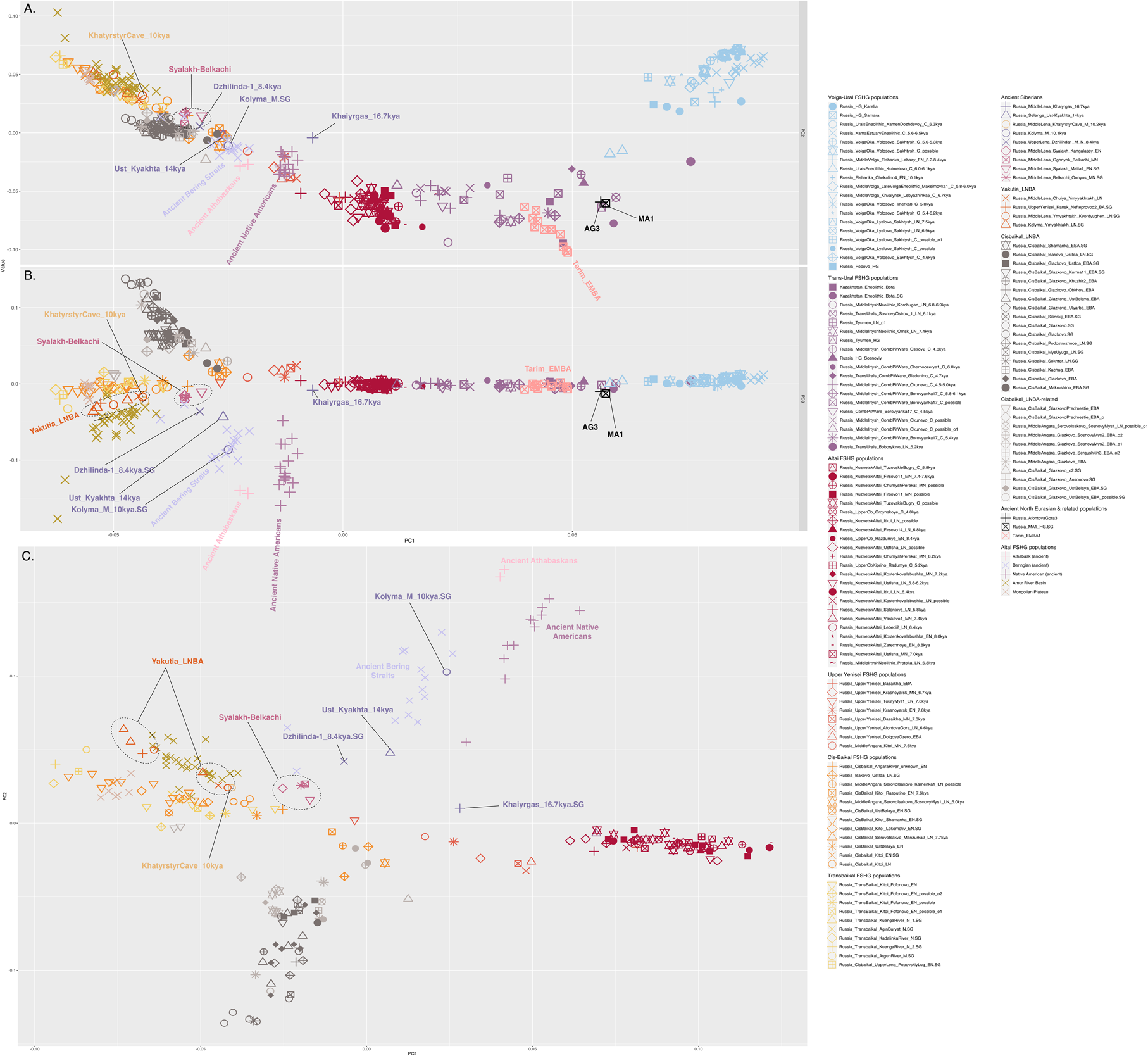
Sites with newly-reported samples. This map displays all sites from which samples that fall in all major populations that are the subject of focus in in this paper have come. These include all sites 1) whose samples fall on the FSHG cline, 2) whose samples fall in the Cisbaikal_LNBA cluster or are admixed with it, 3) whose samples fall in the Yakutia_LNBA cluster or are admixed with it, 4) who are a part of the ten-population East Siberian transect described in our qpAdm modelling, and 5) who are Seima-Turbino period individuals. Each site is represented by a pie chart, whose size is proportional to the number of individuals from that site; the white fraction represented previously-published samples, and the black fraction represents newly-published samples. Our sampling helps to fill important geographic and temporal lacunae.

**Extended Data Figure 3.**
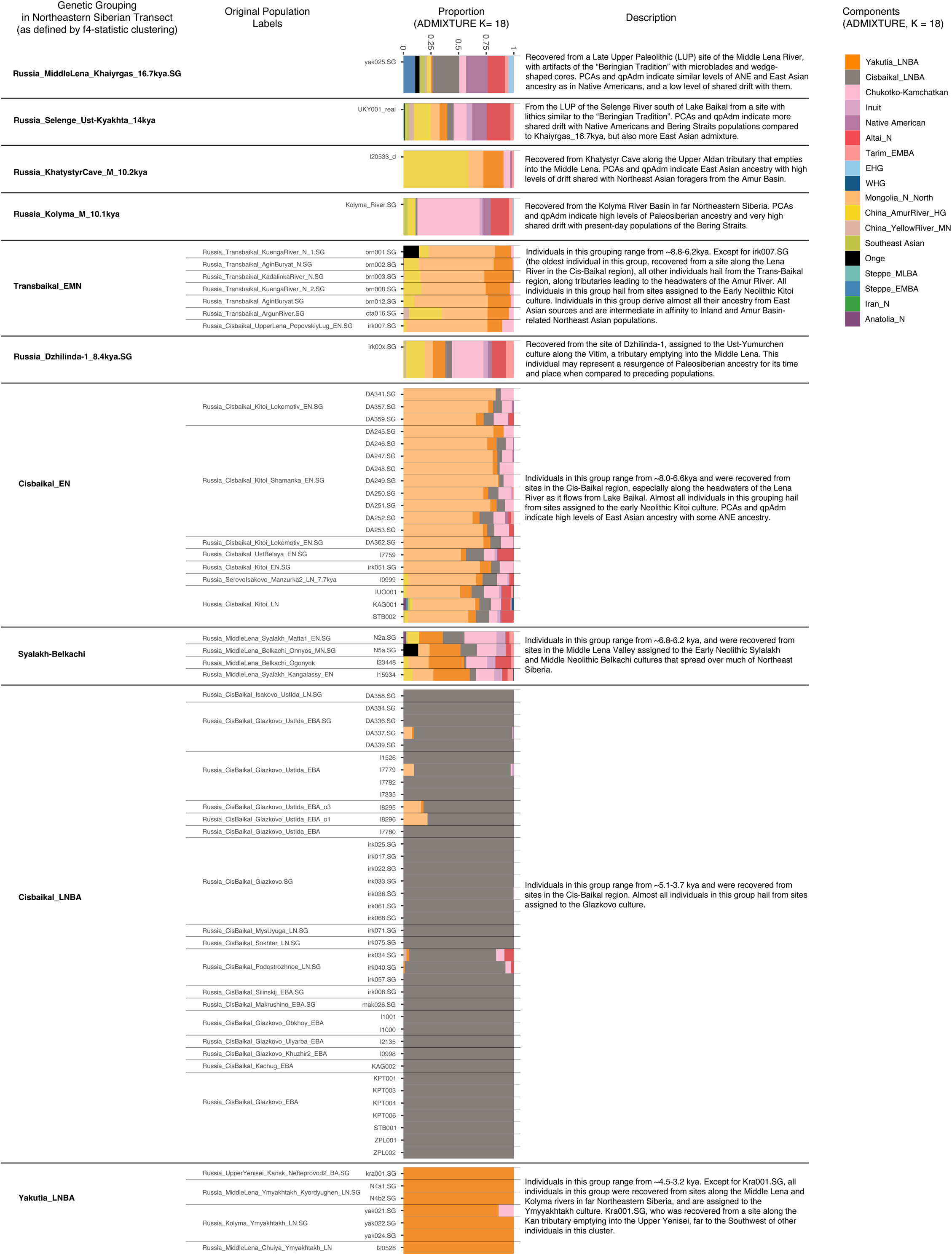
Chronology of sites and cultures in each geographic region. This table summarizes the temporal and geographic disposition of cultures in from the Mesolithic to the Late Bronze and Iron Ages across Northern Eurasia. Sites whose samples are analyzed in our paper are highlighted in darker boxes, within the containing boxes that indicate archaeological cultures. Sites whose colors are darker are those that we believe, based on radiocarbon, isotopic, and archaeological evidence, can be more securely dated.

**Extended Data Figure 4.**
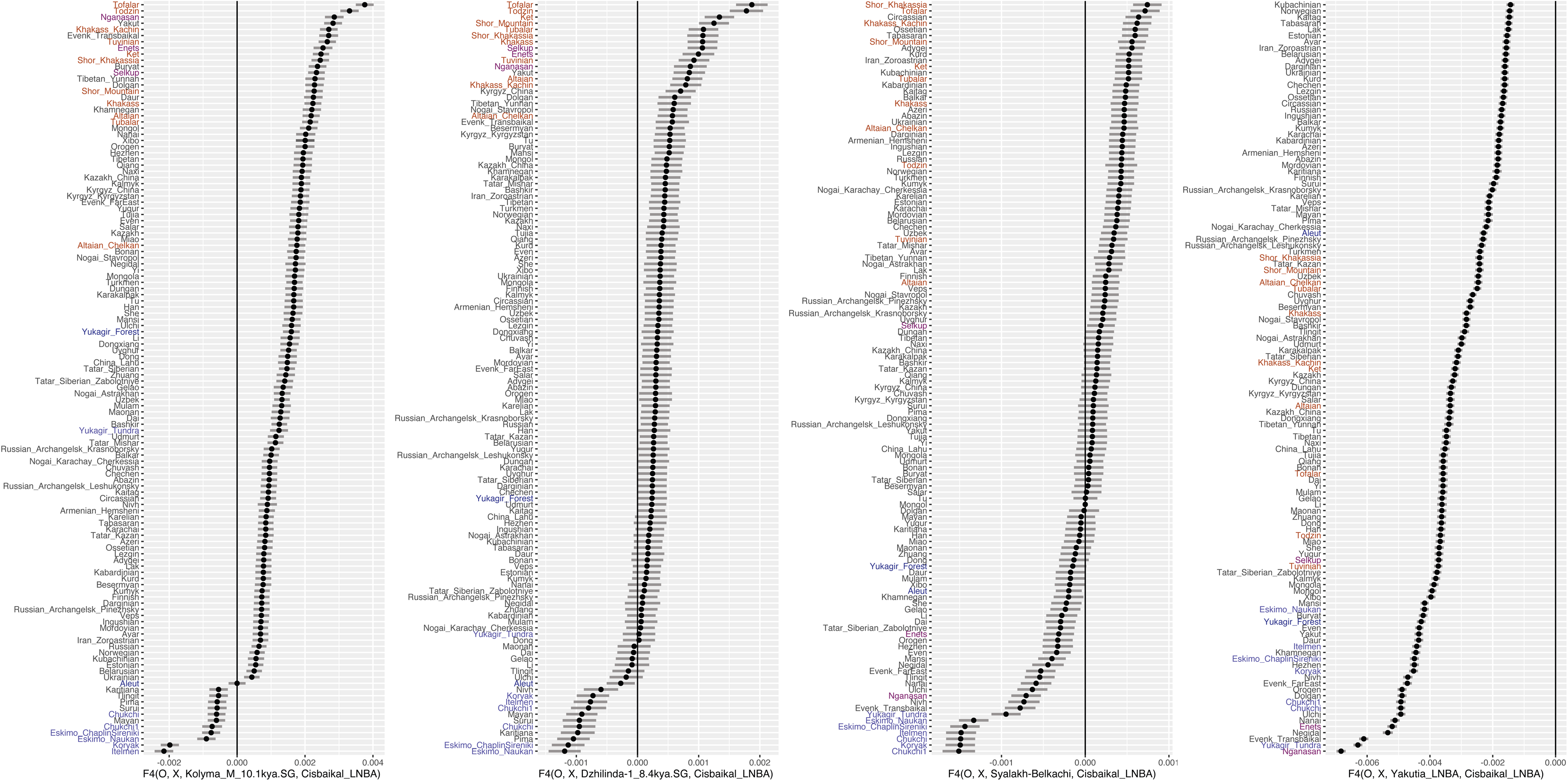
PCA with target populations projected onto ancient populations with an especially high fraction rich in ANE ancestry. To further illuminate the role that levels of *ANE* ancestry plays in generating variation among the populations we analyze, we use as a basis for another projection 71 shotgun-sequenced ancient individuals from across Eurasia, of which a large proportion are enriched in *ANE* ancestry and fall outside the range of present-day variation (e.g. individuals from populations like *Tyumen_HG.SG* or *Kazakhstan_Botai.SG*; for a full list, see SI Section III). The forest-steppe-hunter-gatherer cline forms a curved arc stretching from *EHG* populations to present-day East Asians; the center of the arc dominated by populations rich in *ANE* ancestry is moved toward the positive direction in PC2. The individual furthest along the positive direction in PC2 is AG3. Clines formed by later Inner Asian populations, such as present-day Uralic, Turkic, and Mongolic speakers, as well as Late Bronze Age and Iron Age steppe populations such as Scythians and Sarmatians, are distinguished from the *FSHG* cline by their much lower values along PC2, suggesting a much lower level of ANE ancestry. This PCA shows that populations along the FSHG cline, remaining stable for many millennia, were substantially outside the range of present-day genetic variation in Northern Eurasia.

**Extended Data Figure 5.**
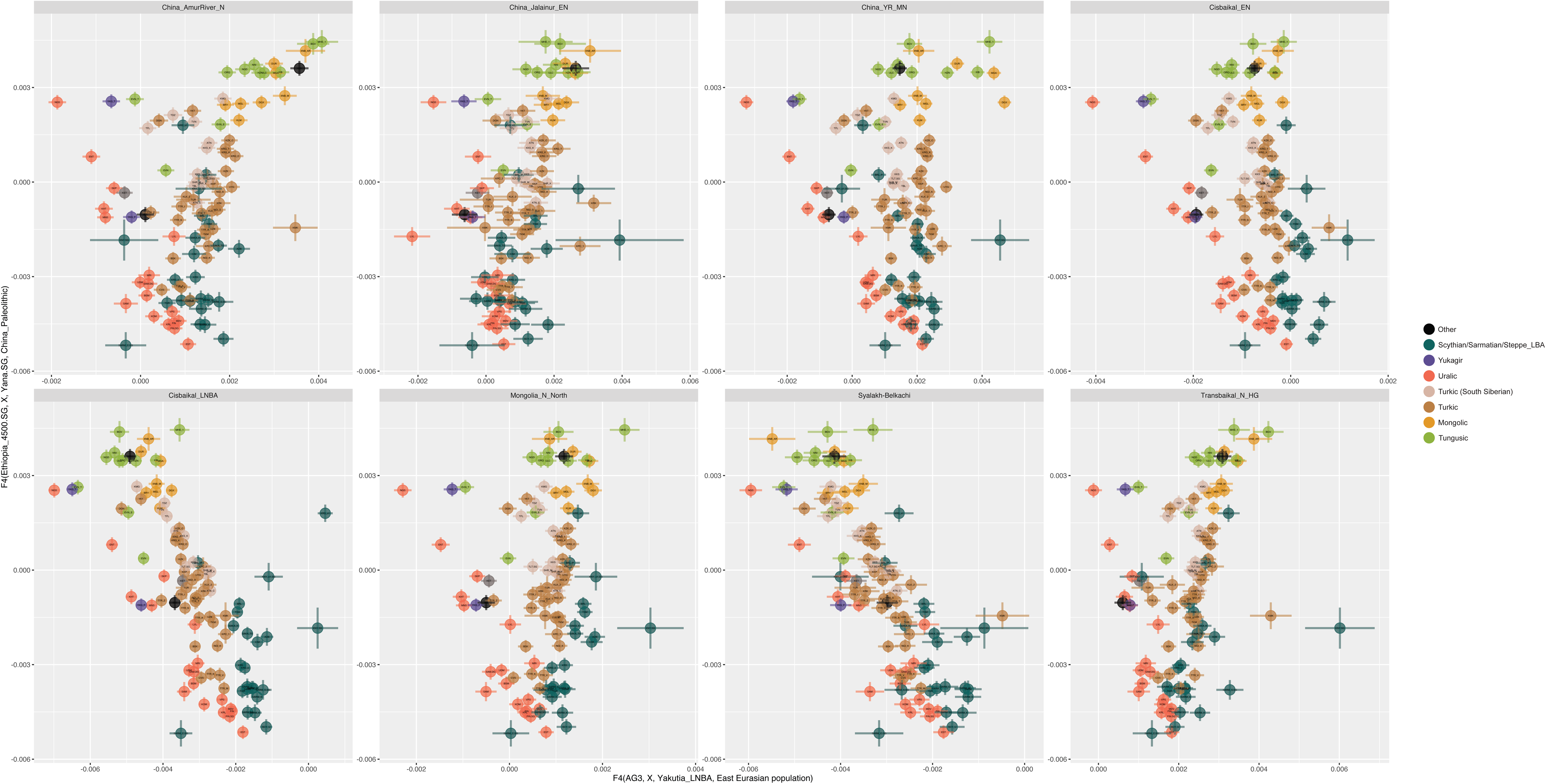
PCA focusing on East Eurasian populations. To further uncover possible structure among the East Asian ancestries within the populations that we analyze, we constructed a third PCA, using as a basis 37 East Asian present-day populations that have minimal West Eurasian admixture, and a single West Eurasian population (Norwegian), all genotyped on the Affymetrix Human Origins array. For a full list of the populations used, refer to SI Section III. We projected all other shotgun-sequenced and hybridization-captured ancient and present-day individuals onto this basis. Once again, the forest-steppe-hunter-gatherer cline forms a curved arc stretching from West Eurasian populations to present-day East Asians, with the center of the arc deflected toward the AG3 individual. East Asian populations are now differentiated along PC2, with Southeast Asians and East Asian agriculturalists taking on especially negative values along that dimension, populations from the Amur River Basin taking on intermediate values, followed by populations on the Mongolian Plateau and surrounding areas. A large gap then separates these populations from *Yakutia_LNBA* and Russia_Tatarka_BA, which take on very positive values along PC2, close to present-day Nganasans and a genetically very similar Iron-Age individual from Yakutia who clusters with Nganasans in the previous two PCAs (Yakutia_IA.SG; also see Extended Data Fig. 10). As one moves East along the *FSHG* cline, their positions along PC2 tend to converge to the values found among populations of the Mongolian Plateau. In contrast, the Dzhilinda1_M_N_8.4kya and Kolyma_M_10.1kya individuals, and the Syalakh_Belkachi, *Yakutia_LNBA* and Russia_Tatarka_BA populations do not fall on the *FSHG* cline and are shifted in the positive direction on PC2, toward the positions occupied by Nganasans, Beringian populations, and Native Americans. Lastly, Uralic populations possess the most positive values among PC2 when compared to Turkic, Mongolic and Tungusic populations.

**Extended Data Figure 6.**
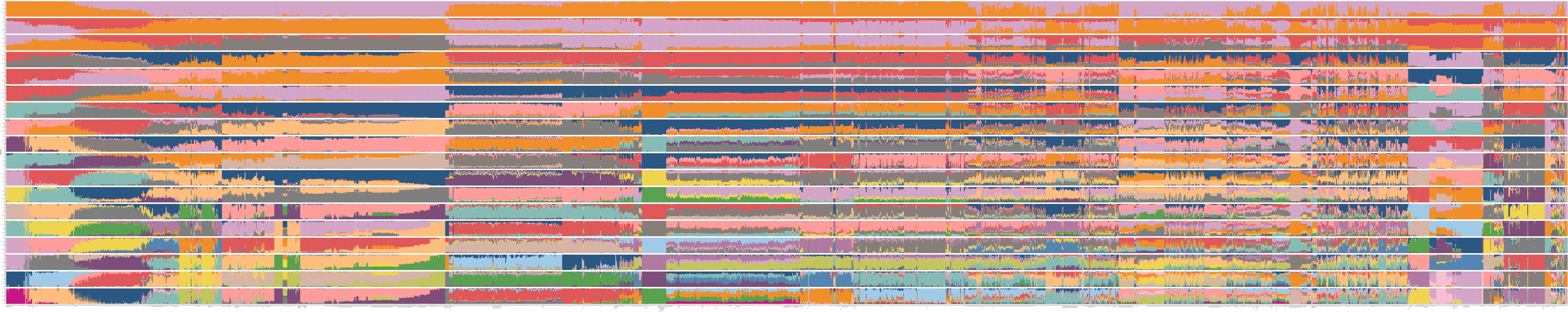
PCA focusing on ancient individuals from Northern Eurasia and the Americas. To understand structure among *FSHG* populations and non-*FSHG* Siberians, we constructed two PCAs with ancient individuals including all individuals from the *FSHG* cline, ancient non-*FSHG* Siberians, and a selection of ancient Beringians and Native Americans. Notably, all these populations possess combinations of only *WHG*, *EHG*, *ANE* and East Asian ancestries. No individuals were projected in these PCAs. The first PCA (Extended Data Fig. 6A) includes all individuals in the set, and the second (Extended Data Fig. 6B) includes only individuals East of the Altai mountains. We notice in the first PCA that 1) the forest-steppe-hunter-gatherer cline again forms a curved arc stretching from West Eurasian populations to East Asian populations along PC1 and PC2. In PC3, populations along the *FSHG* cline also form a straight line. However, populations rich in East Asian ancestry are differentiated along PC3, with individuals and populations within or closely related to the *Cisbaikal_LNBA* cluster having the most positive values, followed by those in the *Transbaikal_EMN* cluster and populations of the Mongolian Plateau, followed by individuals and populations in the *Yakutia_LNBA* cluster, followed by those from the Amur River Basin, followed by populations from the Bering Straits and the Americas. Notably, all individuals along the *FSHG* cline, including individuals rich in East Asian ancestry (e.g. *Cisbaikal_EN, Transbaikal_EMN*, and all *FSHG* individuals from the Krasnoyarsk region) form a straight line in PC3, suggesting a constant source of East Asian ancestry at the East Asian terminus of the *FSHG* cline. *2) Khairygas_16.7kya* occupies a central position among the other groups rich in East Asian ancestry in East Siberia, Beringia and the Americas, suggesting a lack of shared drift with later populations of the Bering region or the Americas. The situation is different for later populations: *Kolyma_M_10.1kya* falls among ancient Beringian populations, while the more East Asian-admixed *Ust_Kyakhta_14kya* and *Dzhilinda1_M_N_8.4kya* occupy a position in between *Syalakh-Belkachi* and ancient Bering Straits populations, with the even more East Asian-admixed *Syalakh-Belkachi* population showing even less of this displacement towards ancient Bering Straits populations. We find an extremely similar pattern in the second PCA (Extended Data Fig. 6B), except with an opposite ordering of the clusters along PC3. Our results suggest that the distinctions we discover between groupings produced by the clustering analyses in SI Section V can be recovered in PCA analyses aimed at recovering fine-scale structure, despite underlying similarities in deep ancestry in populations in East Siberia, Beringia, and the Americas—all the products of admixture between *ANE* and East Asian-associated ancestry.

**Extended Data Figure 7.**
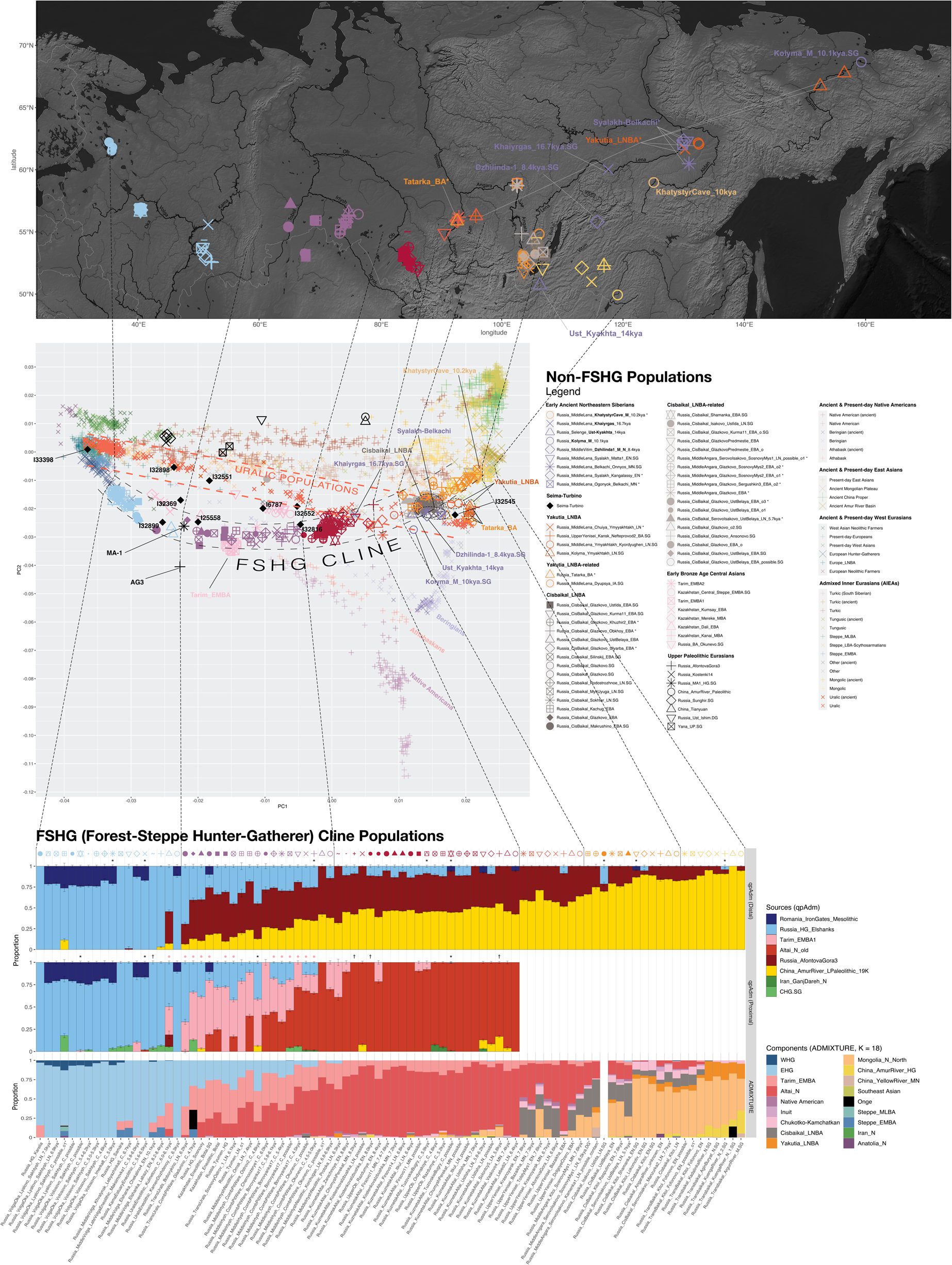
Populations created by genetic grouping procedure applied over Northeast Siberians. Details of populations created by the grouping procedure applied to individuals in Northeastern Siberia.

**Extended Data Figure 8.**
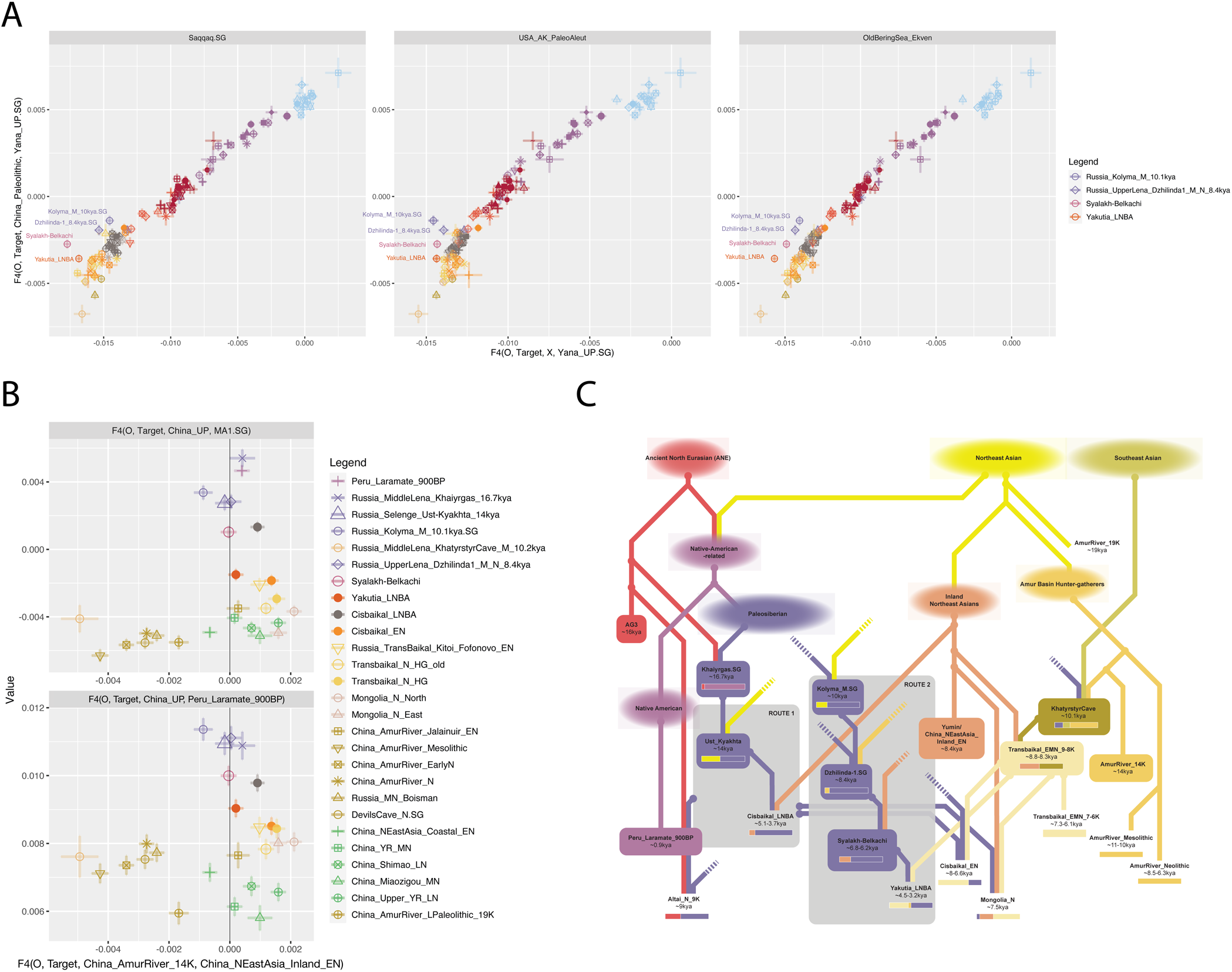
Statistics of the form f4(Ethiopia_4500BP.SG, Target, “Route 2” population, Cisbaikal_LNBA). Central Siberian populations from the Yenisei Basin (including Kets and South Siberian Turks) are highlighted in brown, while Arctic North American and Asian populations on either side of the Bering Straits populations are highlighted in blue. Bering Straits populations that are heavily European-admixed (Aleut and Yukagir_forest) are colored dark blue, while Samoyedic populations (Enets, Selkup, and Nganasan) are colored violet. Despite the similarity of the APS-rich populations in this comparison (all being admixtures between APS ancestry and East Asian ancestry), present-day groups of the Bering Straits are always closer to groups with “Route 2” APS ancestry (i.e., Kolyma_M_10.1kya → Dzhilinda1_8.4kya → Syalakh-Belkachi → Yakutia_LNBA), while Central Siberian populations of the Yenisei Basin are always closer to Cisbaikal_LNBA. For the version including a comparison with Ust_Kyakhta, refer to SI Section VI.A.iv.d.

**Extended Data Figure 9.**
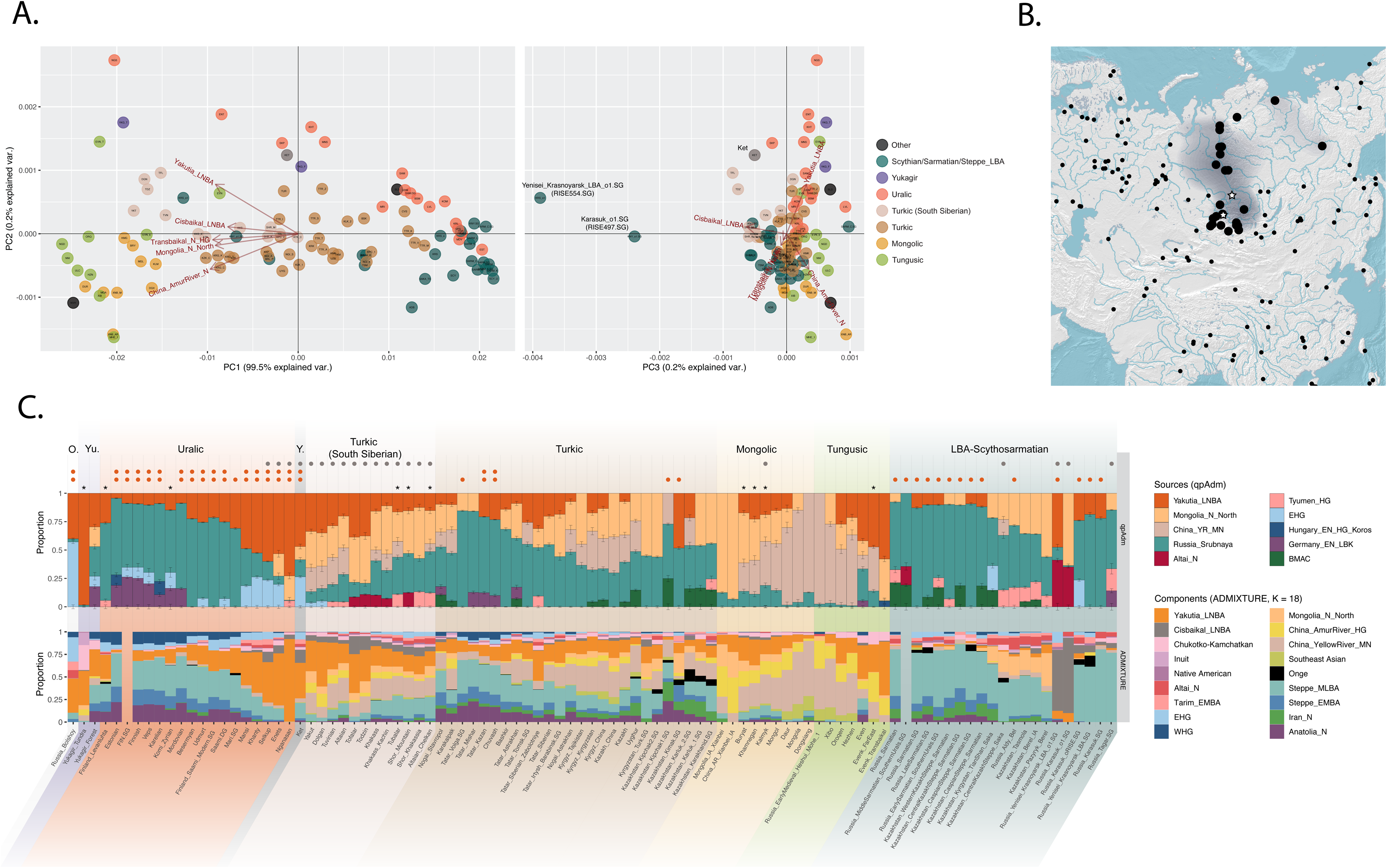
| F4 statistics of the form f4(*Ethiopia_4500BP.SG, X, Yana.SG, China_Paleolithic*) plotted against f4(*AG3, X, Yakutia_LNBA, East Eurasian Population*). *China_Paleolithic* includes the Tianyuan and Armur_River_33K genomes, East Eurasian Population is some population grouping in Siberia or Northeast Asia other than *Yakutia_LNBA*, and X are Admixed Inner Eurasian populations (AIEA populations) including ancient Central Asian nomads from the Late Bronze to Iron Age down to the Scytho-Sarmatian period, as well as modern or ancient populations that speak languages from the Yukaghiric, Yeniseian (Kets), Uralic, Turkic, Mongolic, Tungusic, and Nivkh language families. Modern Uralic-speaking populations, and ancient putatively Uralic-speaking populations uniformly prefer *Yakutia_LNBA* to other East Asian ancestries no matter the other population used in the comparison. Furthermore, at any level of admixture between East and West Eurasian ancestries, the population with the greatest affinity to *Yakutia_LNBA* is always a Uralic-speaking population. F4-statistics therefore highlight the connection between Uralic populations and Yakutia_LNBA ancestry over other sources of East Asian ancestry.

**Extended Data Figure 10.**
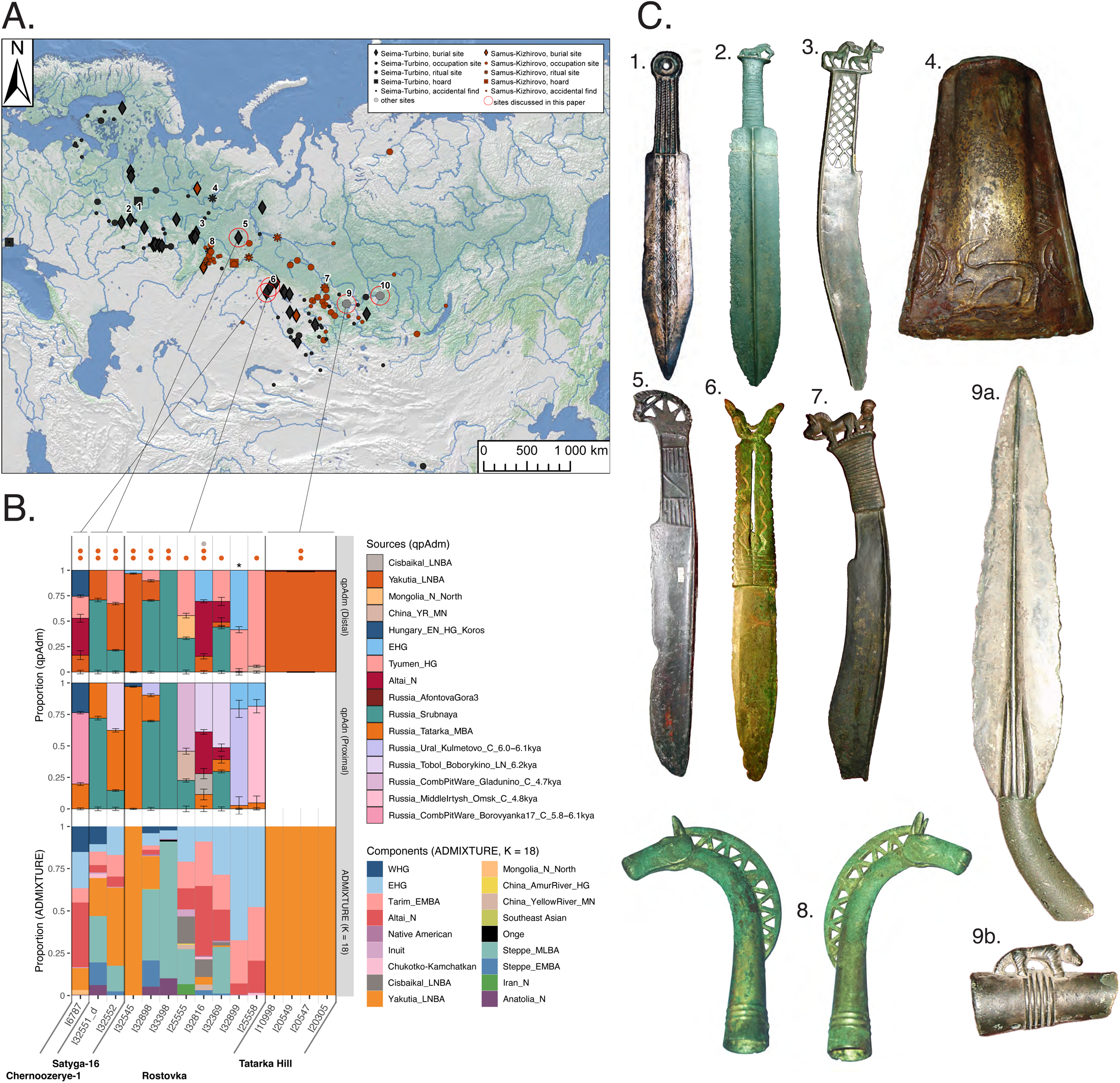
ADMIXTURE results. For details, refer to SI Section IV.

## Supplementary Files

Supplementary File 1

Supplementary text sections I-XI, figures S1-S12, and Supplementary Tables VI-A-1 to VI-D-1. Includes discussion of archaeological context, details of sample preparation, details of population genetic analysis using PCA, *ADMIXTURE*, *qpAdm* and other formal methods such as *f_4_*-statistics, relatedness analysis, uniparental markers, and also linguistic discussion and archeological interpretation.

Supplementary Data S1

Excel spreadsheet with details of all published and newly reported individuals used in this analysis; each table is on a different sheet. Table 1 lists all present-day individuals used in any analysis. Table 2 lists all ancient individuals used in analysis, as well as six additional newly reported individuals that were detected as first-degree relatives of other individuals with higher coverage (I11747, I13100, I11744,). Due to the large number of analyses and the different roles that any given individual sample may play in each analysis, Table 3 presents individual-level information about the clustering results, group labels (which determine the individual’s role in populations in, e.g., *qpAdm* models) and groupings (e.g., for color and point-shape assignment in PCAs) for each ancient and present-day individual across any analysis conducted in this this paper. Table 4 presents C14 and isotopic data for newly-published samples. Table 5 presents a synchronization table for all newly-published samples and their cotemporaneous archaeological entities.

Supplementary Data S2-S7

Excel spreadsheets with details about *qpAdm* and *f_4_* analyses.

**Supplementary Data S2** presents results (all models, whether passing or failing, of the combinatorial rotating outgroup *qpAdm* protocol) for all *qpAdm* sets for *FSHG* populations and other ancient Siberians that are combinations of *EHG*, *ANE* and East Asian ancestry. There are two sets of results in two tables (separate sheets). Details on the procedures used to produce the results can be found in SI Section VI.E.

**Supplementary Data S3** presents results (all models, whether passing or failing, of the combinatorial rotating outgroup *qpAdm* protocol) for all *qpAdm* sets for East Siberian populations, aimed at investigating their mutual relationships, and of which the results are summarized in Fig. 2C. There are 16 sets of results, on 16 sheets labelled “Table 2A” to “Table 9”. In addition, Table 1 presents the individuals assigned to each population used across all *qpAdm* models, and Table 10 presents a corresponding set of *f_4_*-statistics. Details on the procedures used to produce the *qpAdm* results can be found in SI Section VI.A, and details on the *f_4_*-statistics can be found in SI Section VI.A.iv, under the subheading “Ancient Paleosiberian ancestry persists in Selenge_Ust-Kyakhta_14kya and Kolyma_M_10.1kya, but with East Asian admixture and possible Native American backflow”.

**Supplementary Data S4** presents results of the clustering analysis in SI Section V. Table 1 presents cluster memberships, Table 2 and 3 present the results of the clustering procedure detailed in SI Section V for the individuals listed in VI-B and VI-C, respectively, and Table 4 presents the results of *qpAdm* analyses testing group homogeneity described in VI-B and VI-C.

**Supplementary Data S5** presents results of the *qpAdm* analyses (all models, whether passing or failing, of the combinatorial rotating outgroup *qpAdm* protocol) described in SI Section VI.B, where ancient populations from either side of the Bering Strait are modeled as descending from the previously modeled populations in Eastern Siberia. Table 1 presents the individuals assigned to each population used across all qpAdm models. There are three sets of results shown in Tables 2, 3A, and 3B.

**Supplementary Data S6** presents results of the *qpAdm* analyses (all models, whether passing or failing, of the combinatorial rotating outgroup *qpAdm* protocol) described in SI Section VI.C and VI.D, where the *Yakutia_LNBA* and *Cisbaikal_LNBA* admixture proportions among AIEA populations are estimated using *qpAdm*. There are five sets of results shown in Tables 1A (basic *qpAdm* models described in Section VI.C.i), 1B (without shotgun-sequenced individuals in references or sources), 2 (with an Amur-River-related population added to the sources and references), and 4 (with *Cisbaikal_LNBA* added to the references and sources). In addition, it contains a table with *qpAdm* results for the *Russia_Samus_BA_Tatarka* population (Table 3) using a setup described in SI Section VI.C.iii.

**Supplementary Data S7** presents the results of the qpAdm analyses (all models, whether passing or failing, of the combinatorial rotating outgroup *qpAdm* protocol) described in SI Section VII, investigating the ancestry of ST individuals. Table 1 lists the individuals used in group labels and in the calculation of f2 statistics. Table 2 lists the results of the distal qpAdms, and Table 3 the proximal qpAdms. Table 4 lists the results for qpAdms for two individuals (I25555 and I32816) with *Cisbaikal_LNBA* added to the sources and the references as part of outgroup rotation.

## Footnotes

a Refer to SI Section I.A. for a guide to the geography and geographic terms used in this paper.

b This individual, judging from isotopic values indicative of a resevoir effect (SI Data 1 Table 2/4; SI Section IX), and the fact that ST materials are in probable association (SI Section II.F.iii.a)—is a possible ST-period burial from Chernoozerye-1, a cemetery of the Late Krotovo culture. To see the complete discussion of why we think this individual is ST, refer to SI section II.F.iii.a.

c Our samples can be securely dated to ∼4.0kya, cotemporaneous with the Seima-Turbino phenomenon, because of their C14 dates and accompanying isotopic data indicating a low likelihood of reservoir effects (SI Data 1 Table 4; SI Section IX).

