## Supplementary Information for "Postglacial genomes from foragers across Northern Eurasia reveal prehistoric mobility associated with the spread of the Uralic and Yeniseian languages"

II.D.i.b) Trans-Ural Eneolithic (Russia_CombPitWare_Eneolithic) ... 97
*II.D.ii. Ural Mountains and Cis-Urals ... 98*
II.D.ii.a) Ural Early Eneolithic Burials (Russia_UralsEneolithic) ... 98
II.E. TEMPERATE ZONE OF EASTERN EUROPE ... 101
*II.E.i. Lower Kama and Middle Volga Region ... 101*
II.E.i.a) Elshanka Early Neolithic culture (Russia_Elshanka_EN) ... 101
II.E.i.b) Khvalynsk Eneolithic culture (Russia_Khvalynsk_Eneolithic) ... 104
II.E.i.c) Eneolithic at the Kama Estuary (Russia_KamaEstuaryEneolithic) ... 106
II.E.i.d) Volga Late Eneolithic burials (Russia_LateVolgaEneolithic) ... 110
*II.E.ii. Volga-Oka Region ... 113*
II.E.ii.a) Lyalovo Pit-Comb Ware Late Neolithic culture (Russia_UpperVolga_Lyalovo_LN) ... 113
II.E.ii.b) Volosovo Eneolithic culture (Russia_Volosovo_Eneolithic) ... 120
II.F. SEIMA-TURBINO-PERIOD INDIVIDUALS ... 128
*II.F.i. Overview of the Seima-Turbino phenomenon ... 128*
*II.F.ii. Seima-Turbino necropolises (burial sites) ... 129*
II.F.ii.a) Rostovka burial site ... 129
(1) Burials at the Rostovka Seima-Turbino necropolis ... 133
II.F.ii.b) Satyga-16 burial ground ... 135
(1) Burials at the Satyga-16 Seima-Turbino necropolis ... 136
*II.F.iii. Peripheral Area of the Seima-Turbino Phenomenon ... 138*
II.F.iii.a) Chernoozerye-1 burial site ... 138
II.F.iii.b) Tatarka Hill site ... 139
(1) Burials at the Tatarka Hill Site ... 140
II.F.iii.c) The Nefteprovod-1 and Nefteprovod-2 sites (Anzhevskiy complex), archaeologically related to the site of Tatarka Hill ... 141
II.G. KRASNOYARSK CISBAIKAL_LNBA-RICH OUTLIERS ... 143
II.H. REFERENCES (FOR SECTION II) ... 145

**III. POPULATION GENETIC ANALYSIS: PCA ... 158**

**IV. POPULATION GENETIC ANALYSIS: ADMIXTURE ... 160**

**V. POPULATION GENETIC ANALYSIS: FORMAL GROUPING OF INDIVIDUALS IN NORTHEASTERN SIBERIA INTO POPULATIONS ... 160**

V.A. UNGROUPED PRE-HOLOCENE INDIVIDUALS FROM SIBERIA ... 161
V.B. GROUPING INDIVIDUALS FROM THE TRANSBAIKAL REGION AND EASTERN SIBERIA ... 162
V.C. GROUPING INDIVIDUALS FROM THE CISBAIKAL REGION AND WESTERN SIBERIA ... 165

**VI. POPULATION GENETIC ANALYSIS: QPADM & FORMAL STATISTICS ... 169**

VI.A. THE DEEP POPULATION HISTORY OF EASTERN SIBERIA ... 169
*VI.A.i. Modelling Approach and Parameters ... 169*
*VI.A.ii. Naïve Models ... 172*
VI.A.ii.a) Tranche 1: Khaiyrgas_16.7kya ... 172
VI.A.ii.b) Tranche 2: Ust_Kyakhta_14kya ... 173
VI.A.ii.c) Tranche 3: KhatystyrCave_M_10.2kya, Kolyma_M_10.1kya ... 173
VI.A.ii.d) Tranche 4: Dzhilinda1_M_N_8.4kya, Transbaikal_EMN_old, Altai_N_old ... 176
VI.A.ii.e) Tranche 5: Altai_N, Transbaikal_EMN, China_AmurRiver_N, Cisbaikal_EN, Mongolia_N_North, Syalakh-Belkachi ... 178
VI.A.ii.f) Tranche 6: Yakutia_LNBA, Cisbaikal_LNBA ... 182
*VI.A.iii. Models with Cisbaikal_LNBA in Tranche 3 ... 183*
VI.A.iii.a) Finding a Tranche for Cisbaikal_LNBA ... 183
VI.A.iii.b) Tranche 3: KhatystyrCave_M_10.2kya & Kolyma_M_10.1kya ... 185
VI.A.iii.c) Tranche 4: Dzhilinda1_M_N_8.4kya, Altai_N_old & Transbaikal_EMN_old ... 186
VI.A.iii.d) Tranche 5: Altai_N, Transbaikal_EMN, China_AmurRiver_N, Cisbaikal_EN, Mongolia_N_North & Syalakh-Belkachi ... 190
VI.A.iii.e) Tranche 6: Yakutia_LNBA ... 194

*VI.A.iv. Summary of Population Relationships in Siberia* ... *194*
VI.A.iv.a) Investigating the genetic peculiarities of Cisbaikal_LNBA ... 194
VI.A.iv.b) The Khaiyrgas_16.7kya is a near-unadmixed representative of an "Ancient Paleosiberian" (APS) lineage that retains broad affinities to Native Americans ... 195
VI.A.iv.c) Ancient Paleosiberian (APS) ancestry persists in Ust_Kyakhta_14kya and Kolyma_M_10.1kya, but with East Asian admixture and possible Native American backflow ... 196
VI.A.iv.d) Ancient Paleosiberian (APS) ancestry persisted into late Holocene populations of Northeastern Eurasia and the Americas along two routes. ... 196
VI.A.iv.e) East-Asian ancestry Admixed into Siberian populations during the Holocene along two routes 200
VI.B. PERSISTENCE OF ANCIENT PALEOSIBERIAN ANCESTRY INTO LATER POPULATIONS ALONG THE BERING STRAITS 202
*VI.B.i. Connection between the Syalakh-Belkachi Population and the Saqqaq Paleo-Eskimo* ... *204*
*VI.B.ii. qpAdm models indicate that Ancient Paleosiberian ancestry persisted on both sides of the Bering Strait, especially through Saqqaq.SG* ... *205*
*VI.B.iii. qpAdm models provide evidence for a distinct stream of Ancient Paleosiberian ancestry among ancient Athabaskans* ... *209*
*VI.B.iv. Summary of the role of Ancient Paleosiberian ancestry in population relationships on either side of the Bering Straits* ... *212*
VI.C. AFFINITIES OF YAKUTIA_LNBA ... 214
*VI.C.i. f4-statistics demonstrate that Yakutia_LNBA is especially related to Uralic-speaking populations* ... *214*
*VI.C.ii. qpAdm demonstrates that Uralic-speaking populations uniquely among Inner Eurasian populations draw their East Asian ancestry almost exclusively from Yakutia_LNBA* ... *215*
*VI.C.iii. Modeling the spread of Yakutia_LNBA ancestry to the Volga-Ural region* ... *217*
VI.D. AFFINITIES OF CISBAIKAL_LNBA ... 218
*VI.D.i. $f_4$-statistics demonstrate that Cisbaikal_LNBA ancestry plays an important role among Yeniseian, South Siberian Turkic, and Samoyedic populations* ... *219*
*VI.D.ii. qpAdm models demonstrate that Cisbaikal_LNBA ancestry plays an important role among Yeniseian, South Siberian Turkic, and Samoyedic populations* ... *220*
*VI.D.iii. Cisbaikal_LNBA ancestry expands into the Middle Angara and Krasnoyarsk regions in the Bronze Age 221*
VI.E. MODELING THE FSHG CLINE ... 223
*VI.E.i. Simple qpAdm models demonstrate that FSHG populations can be modeled as a cline of admixed populations grading between East Asian ancestry, ANE, EHG and WHG, with post-LGM ancestry sources* ... *223*
**VII. POPULATION GENETIC ANALYSIS: INDIVIDUALS FROM SEIMA-TURBINO-PERIOD SITES ... 225**
**VIII. Y-CHROMOSOMES HAPLOGROUPS ... 233**
**IX. RADIOCARBON AND STABLE ISOTOPE ANALYSIS ... 233**
**X. REFERENCES FOR SECTIONS III-IX. ... 236**

### I. Geographic Terms and Climatic Context

#### *I.A. Geographic Terms*

The study area covers a vast territory of Northern Eurasia from the Far East to the Baltics, encompassing a diverse range of orographic conditions, climate, and resources. The area can be broadly divided into several sub-areas: Northeastern, Central, Western, and Southern Siberia, and the

East European Plain (Fig. 1).

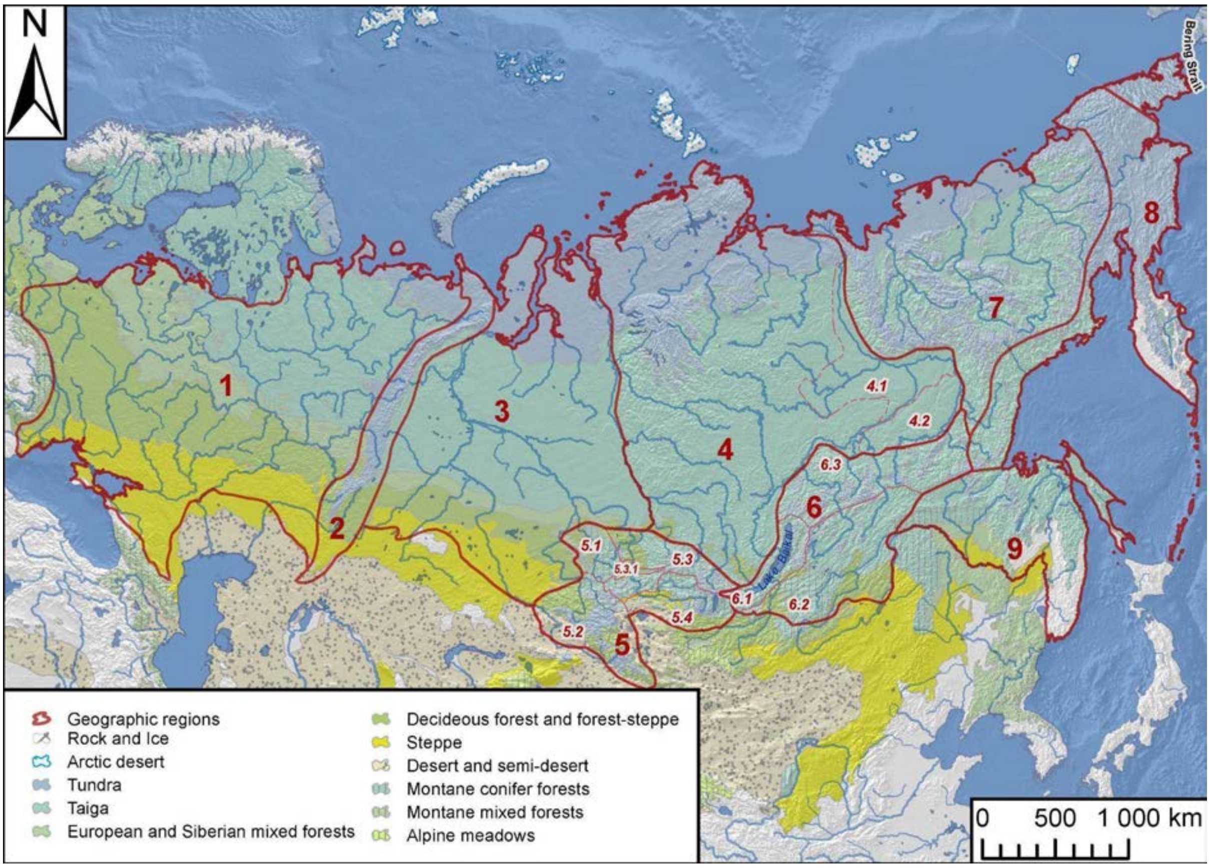

***Fig. S1.*** *Geographic regions of Northern Eurasia. 1 - East European Plain, 2 - Urals, 3 - West Siberia, 4 - Central Siberia (4.1 - Central Yakutia, 4.2 - Aldan), 5 - Altai-Sayan (5.1 - Kuznetsk-Salair, 5.2 - Altai, 5.3 - Sayan with 5.3.1 - Minusinsk Basin, 5.4 - Tyva), 6 - Baikal (6.1 - Baikal, 6.2 - Transbaikalia, 6.3 - Baikal-Stanovoy), 7 - North-East Siberia, 8 - North Pacific, 9 - Amur-Sakhalin (8 and 9 taken together are called Russian Far East) (boundaries of regions and provinces based on Map of the physical-geographical zoning of the USSR 1983, Gvozdevskiy, N. A., & Mikhailov, N. I. (1978); biogeographic regions Olson et al. 2001, generalized).*

**Table S1 | Most common geographic terms in main text**

| Term | Regions (numbered in map) |
|---|---|
| Forest belt | Regions colored as “Taiga” and adjoining areas colored as “European and Siberian mixed forests” in Fig. S1 |
| Forest-steppe belt | Regions colored as “Decideous forest and forest-steppe” in Fig. S1 |
| Northeast Siberia | Region numbered 7 in Fig. S1 |
| Central Siberia | Region numbered 4 in Fig. S1 |
| Altai-Sayan | Region numbered 5 in Fig. S1 |
| Western Siberia | Region numbered 3 in Fig. S1. Contains the Ob and Irtysh Basins (Fig. S2). |
| Cis-baikal | Southernmost part of Central Siberia (region 4), just to the West of Lake Baikal, in Fig. 1, including the headwaters of the Angara and Lena as they leave Lake Baikal (Fig. S3). |
| Trans-baikal | Region numbered 6.2, including the region just below the number 6 in Fig. S1 |

| | |
|---|---|
| Yakutia | Regions numbered 4.1 and 4.2, eastern part of region numbered 4 more generally, plus region numbered 7, in Fig. S1 |

**Northeastern Siberia** (Fig. 1, see 7) encompasses the eastern regions of Siberia up to the Lena River, as well as the river basins that flow into the Laptev Sea, East Siberian Sea, and Okhotsk Sea. The area is characterized by a complex relief, mainly composed of ridges and plateaus, which descend like an amphitheater to the north, giving way to the Yana-Indigirka and Kolyma lowlands. The climate changes from arctic to subarctic, always continental, with very cold winters and cool summers. This region is one of the "poles of cold" of the Northern Hemisphere, with winter temperatures around -50°C (the official minimum in Oymyakon is -67.8°C). Permafrost is widespread everywhere across the region. The arctic and shrubby tundra in the north of Northeastern Siberia is replaced by taiga in the south, dominated by larch forests that can tolerate the harsh conditions of the area. In the southern part of Northeastern Siberia, the larch taiga is replaced by mixed forests of larch and other coniferous trees such as spruce and pine.

**Central Siberia** (also called Middle Siberia, Fig. 1, see 4), is located in the center of North Asia between the major Siberian rivers Lena and Yenisei. The Central Siberian Plateau is the main relief structure of the area, which is bordered in the south by the Eastern Sayan Mountains and the Baikal ranges. The area includes also the North Siberian Lowland and Taimyr (with the Byrranga Mountains) in the north. ***Yakutia*** (Fig. 1, see 4.1) as a vast territory within Central Siberia is often considered within the administrative framework, but in our study it is basically understood as a flat area in the middle and lower reaches of the Lena River, the lower reaches of the Aldan River and the Vilyui Basin. One of the region's features is the Siberian Traps - a huge lava field that erupted millions of years ago, cut through by narrow canyon-like river valleys (the Putorana Plateau). The climate is sharply continental, with a large gradient of winter and summer temperatures, and frosty and dry winters, especially severe in the center of the area in Yakutia; permafrost is widespread almost everywhere. Central Siberia is isolated from the inflow of air masses from the Pacific and Atlantic Oceans, so precipitation comes only from the northwest and its amount is reduced in the easterly direction. The Taimyr Peninsula and the North Siberian Lowland are occupied by arctic tundra and shrubby tundra. Most of the region is dominated by taiga, mainly larch forests, with an admixture of pine and dark coniferous. The most southern part of Central Siberia and the upper reaches of the Angara, flowing from Baikal, is usually called *Cis-Baikalia,* is occupied by southern taiga landscapes with humid forests of Siberian cedar and spruce.

**Western Siberia** (Fig. 1, see 3) is located in the western part of North Asia and covers a vast territory, from the Ural Mountains to the Yenisei River basin. The region is characterized by a slightly undulating plain (the West Siberian Plain), interspersed with low hills and ridges. In this flat landscape, only the Siberian Uvaly band is most noticeable - these are low moraine hills formed after one of the Quaternary glaciations and stretching in a sub-latitudinal direction throughout Western Siberia. The climate of Western Siberia is sharply continental, with large seasonal and diurnal temperature fluctuations. Winters are long, severe, and dry, with temperatures frequently dropping below -30°C. Summers are relatively short, warm, and humid. The region receives low annual precipitation, mainly during the summer months, but it does not have time to evaporate or enter the rivers. Therefore, this region contains the largest wetland areas in the world. Western Siberia is home to a diverse range of natural zones, from tundra and forest-tundra in the north to steppe in the south. The vegetation of Western Siberia is dominated by taiga forests, primarily consisting of coniferous trees such as spruce, fir, pine, and larch. The forested area in the center of the West Siberian Plain is interspersed with huge bogs and other wetlands. In the south, the forest gives way to the forest-steppe zone, characterized by a mixture of birch-pine forest and meadows. The steppe region of Western Siberia is primarily comprised of grasslands, which gradually become drier to the south.

Western, Central, and Northeastern Siberia are bordered to the south by **Southern Siberia Mountains** rugged terrain, which comprises two big regions **Altai-Sayan** (Fig. 1, see 5) and **Baikal** (Fig. 1, see 6) with several specific provinces features. Altai-Sayan region includes *Kuznetsk-Salair* mountain-hollow area (5.1); the *Altai* ridges (5.2); the Western and Eastern *Sayan* ridges (5.3) with the *Minusinsk Basin* (5.3.1.); and arid *Tyva* basin surrounded by ridges (5.4). Some mountain ranges, such as the Mongolian Altai and the Eastern Sayan, extend further to the south and are gradually replaced by the Mongolian Plateau, which descends to the plains to the east. In a broad sense big Baikal region, directly includes the *Baikal* province (6.1) with mountain ranges and basins directly bordering Lake Baikal from the northwest and southeast; *Trans-Baikalia* (6.2), which comprises the territory east of Lake Baikal and includes the Vitim Plateau, the mountain ranges in the Selenga River basin, and further southeast the Yablonevy, Daursky, Chersky, and Nerchinsky ranges; and *Baikal-Stanovoy* also called North Trans-Baikalia (6.3) further extends on the area of Stanovoy ranges and Aldan highlands. In the south, Trans-Baikalia passes into the Mongolian steppes, and in the north-east Baikal-Stanovoy borders the Amur region and the Far East. The mountains of Southern Siberia serve as a natural barrier between the westerlies zone (North Eurasia) and the monsoon zone (South and Southeast Asia). The climate in Southern Siberia Mountains is continental and can be quite harsh, but it is also diverse depending on the relief. Due to the great distance from the ocean, winds from the Atlantic arrive having already lost a significant amount of moisture. Nevertheless, the western and northern slopes of the mountains in Southern Siberia receive a significant amount of precipitation, which is why large Siberian rivers such as the Ob, Yenisei, and Amur originate in this region. The slopes of the mountains are covered with dense dark coniferous and deciduous-coniferous forests, which provide a variety of habitats. In contrast, the southern ranges and intermountain basins receive less moisture and are occupied by dry steppes and even semi-deserts. Additionally, the intermountain basins (like Minusinsk Basin) experience specific winter temperature regimes, as cold air flows down the slopes of the mountains and stagnates there, resulting in very severe frosts and calm weather. Long, frosty winters contribute to the formation of permafrost in this region. Summer in Southern Siberia is short, but still relatively warm.

The **Ural Mountains** (Fig. 1, see 2) are a natural boundary between Europe and Asia, separating the Siberian territories from the **East European Plain** (Fig. 1, see 1). The extensive East European (Russian) Plain is a vast lowland covering a significant part of Eastern Europe. It extends from north to south from the Barents and White Seas to the Black and Caspian Seas, and from the coast of the Baltic Sea to the Ural Mountains. In the northwest, it is bounded by the Scandinavian mountains, in the southwest by the Sudeten Mountains, in the southeast by the Caucasus, and in the west, the Vistula River serves as a conventional boundary of the plain. Its relief is characterized by a gently rolling terrain with a slight slope towards the north and northwest. The plain has a flat or slightly convex surface, with occasional low hills and ridges. The East European Plain is cut by a network of rivers, including the Volga, the Dnieper, and the Don, which form extensive river valleys and lowlands. The relief of the East European Plain is largely a result of glacial and alluvial processes that took place during the Pleistocene. The plain is divided into several sub-regions with unique topography. The northern part of the plain is dominated by vast wetlands and swamps. The central part is characterized by rolling hills and valleys, while the southern part is a vast flat plain with occasional low ridges. The vegetation zones of the East European Plain are expressed especially distinctly and sub-latitudinally located, and they vary from north to south, reflecting changes in temperature, precipitation, and soil. The northern part of the plain is dominated by tundra and taiga, while the central part is covered by cool mixed and then deciduous forests. The southern part of the plain lies in the forest-steppe zone, where grassy steppe gives way to forests of birch, oak, and pine. The forest-steppe zone is particularly diverse, with rich flora and fauna. The steppe belt is characterized by low landscape, interrupted only by occasional hills and river valleys. In the extreme south, in the Caspian lowland, the vegetation has a semi-desert appearance.

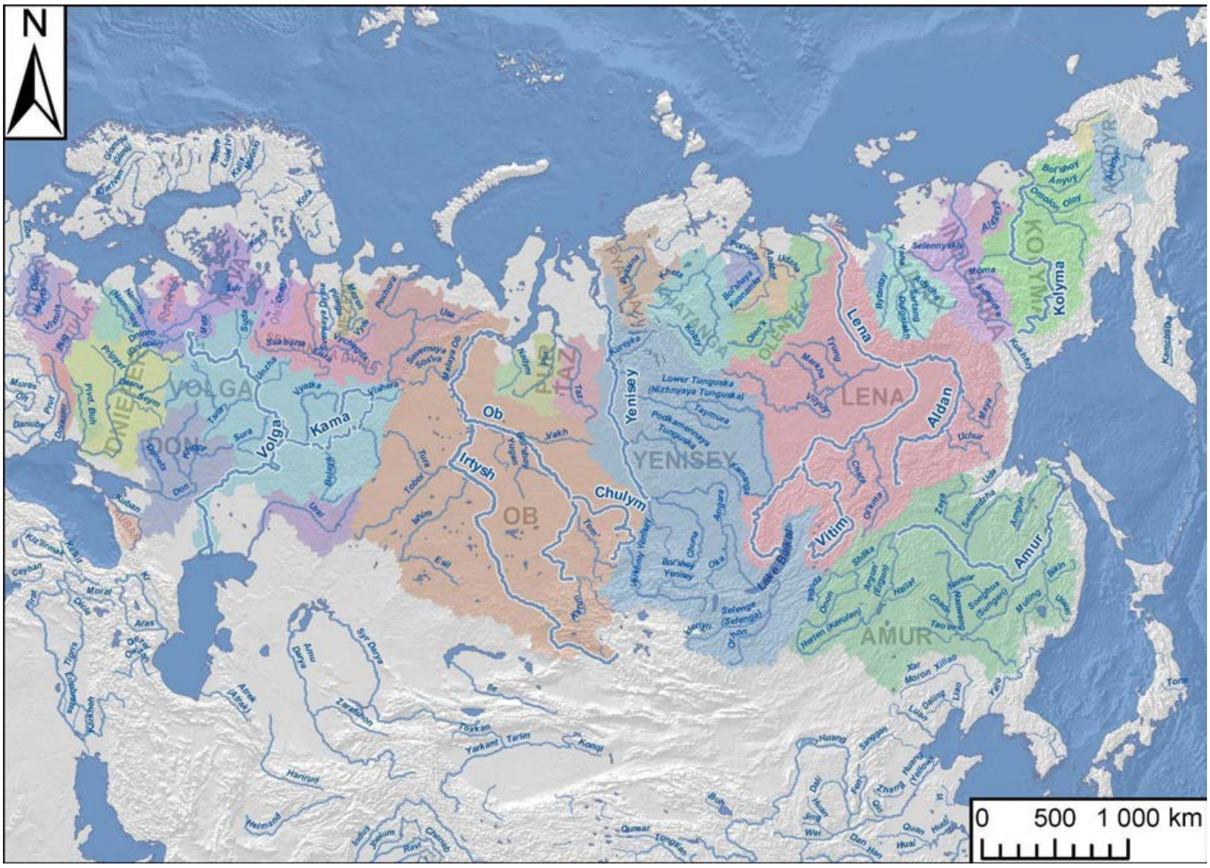

***Fig. S2**. Rivers and river basins of Northern Eurasia. The rivers discussed in the article are highlighted.*

#### *I.B. Climatic and Biogeographic Zones*

Main climatic and biogeographic zones of vast territories of Eastern Europe and Siberia are:

**The tundra zone** covers the northernmost part of Eastern Europe and Siberia, stretching from the Arctic Circle to the northernmost coastlines. Tundra is a distinctive habitat characterized by its frigid climate, low biodiversity, short growing season, and poor soil drainage. Permafrost is everywhere except the extreme west of the zone. Many small thermokarst lakes and vast wetlands are impassable in summer. The tundra biome experiences extremely low temperatures and strong winds, creating harsh conditions that limit growth of most plants and animals. The climate is too harsh for trees to grow, and thus tundra biome is characterized by low-growing plants such as mosses, lichens, and dwarf shrubs that are well adapted to the nutrient-poor soils. Insects, rodents, and migratory birds are common in the tundra during the short summer months when there is a burst of vegetation growth, but many animals migrate to warmer regions during the long winter season.

**The boreal zone** covers the northern part of Eastern Europe and Siberia, stretching approximately from the Arctic Circle to the southern boundary of the taiga. The climate is harsh and cold, with long, severe winters and short, cool summers. The boreal zone is characterized by its extremely cold temperatures, with record lows reaching -54°C in some areas of northeastern Russia. The main factor in extremely cold winters in the boreal zone is the Siberian High - a large high-pressure system that forms over Siberia during the winter months, typically from October to April. The Siberian High has a significant effect on the climate, as it brings cold, dry air from the Arctic region and spreads it across much of Siberia and East Asia, and even often determines the weather in Eastern Europe. In the summer months, the Siberian High weakens and moves northward, allowing warmer, moister air to move into the region. Precipitation along the boreal zone is regulated by westerlies and therefore decreases from west to east. The dominant biome in this zone is the taiga, cold coniferous or cool mixed forests. The soil under forests is often acidic, soddy-podzolic, or podzolic. The taiga is dominated by closed, dense forests in the warmer southern regions of the biome. In the colder, northern regions, where the short growing season and low temperatures limit species diversity and tree density, sparse northern taiga forests predominate. Undergrowth is generally absent or weakly developed due to the low levels of light that penetrate the forest canopy. The ground is often covered with a monotony of grass-shrub layers and green mosses. Shrub species such as juniper, honeysuckle, and black currant, as well as dwarf shrubs like blueberries and lingonberries, and herbs such as oxalis and wintergreen, are not abundant.

In the north of Europe, spruce forests are prevalent in the taiga, and towards the south, the forest becomes mixed with deciduous species. In the taiga of the Urals and Western Siberia, light coniferous forests of Scots pine with an admixture of spruce, fir, and Siberian pine dominate. Central and Northeastern Siberia are home to a sparse larch taiga with an undergrowth of elfin Siberian pine and Daurian rhododendron.

**The temperate zone** covers the central part of Eastern Europe and south Urals. Further to the east this zone wedges out; it does not exist in Siberia and reappears only in the southern part of the Far East (in the Amur and Primorye regions). The temperate zone of Eastern Europe is characterized by mild to cool summers and cold winters, with a relatively narrow range of temperatures throughout the year. This region is influenced by both Atlantic and continental air masses, leading to variable weather patterns. Precipitation is moderate and distributed evenly throughout the year, with higher amounts in the western regions. The temperate zone in the Far East has a temperate monsoon climate, with very humid summers. The dominant biome in this zone is the temperate deciduous forest and cool mixed forest, consisting mainly of deciduous broadleaf trees such as oak and maple, as well as mixed forests of conifers and broadleaves. The deciduous forests of Eastern Europe are known for their high biodiversity. The vegetation cover is typically characterized by a multi-layered structure, with an upper layer of tall trees, a middle layer of shrubs, and an understory of herbaceous plants. The forest floor is usually covered with a thick layer of leaf litter and humus, which helps to retain moisture and nutrients. In some areas, the vegetation may also include patches of meadows or wetlands. The soils are typically nutrient-rich and have a high capacity for retaining water, which contributes to the growth of diverse plant communities. The dominant soil types in these forests are generally well-drained and moderately to highly fertile, such as luvisols and cambisols.

To the south, as the climate becomes more continental, forest zones gradually turn into **forest-steppes**. Here, areas of deciduous forests in Eastern Europe or aspen-birch forests in Western Siberia alternate with rich forb-grass meadows on chernozems. The forest-steppe zone has a favorable climate with long, cold winters, but comparatively long and warm summers and a good balance between precipitation and evaporation. Forest-steppe is an important and unique ecosystem, providing various niches and habitats for both humans and wildlife alike.

**The steppe zone** covers the southern part of Eastern Europe and Siberia, characterized by a continental climate with hot summers and cold winters. The steppe belt in the East European Plain stretches from the Carpathians to the southern Urals and the Caspian lowland and further east through Kazakhstan to the Altai mountains, and the fragmented steppes areas continues in the mountain basins of Southern Siberia. Its southern boundary is marked by the foothills of the Caucasus Mountains, while in the north it gradually gives way to the forest-steppe zone. The climate in this zone is continental with hot and dry summers and cold winters. The natural vegetation is dominated by grasses and herbs adapted to arid conditions, with a few sparse shrubs and trees found in river valleys and sheltered areas. Dry grass steppes (feather grass, fescue, couch grass) predominate, and in more humid areas - forb-cereal steppes. As a result of the decay of the rich grass cover in the steppes, chestnut and the most fertile chernozem soils were formed. In the past, the steppe belt was an important area for nomadic pastoralism, and it remains a region important for farming today. The climate along the steppe zone of Eurasia changes from west to east as the region transitions from the Atlantic influence to a more continental regime and falls under the influence of the Siberian High. In the western part of the steppe zone, the climate is characterized by mild, humid winters and warm summers, with a more even distribution of precipitation throughout the year. Moving eastward, the climate becomes increasingly continental, with colder winters and hotter summers, and a more pronounced seasonal distribution of precipitation, with more rainfall in the summer months.

#### *I.C. Climate change during the Late Pleistocene and the Early Holocene*

There is relatively poor information about how the climate and vegetation of the vast region of Siberia have changed over time, and thus the dynamics of paleoecological changes and their chronology can only be given in broad strokes.

##### I.C.i. Late Pleistocene Northern Asia

Based on data compiled from North Eurasia, Tarasov et al. (2000) concluded that the distribution of biomes during the Late Glacial Maximum (18 thousand calibrated years before present, abbreviated as ka BP) differed significantly from that of modern biomes. The taiga belt was greatly reduced and fragmented, and cool mixed and temperate deciduous forests were absent in most areas where they now exist. The tundra belt expanded, and the steppe was the dominant vegetation type across northern Eurasia south of approximately 57°N, directly adjacent to the tundra to the north.

During the Late Pleistocene (21-14ka BP), steppe was widespread in central Yakutia and northeastern Siberia (Cao et al., 2019), reflecting the expansion of tundra-steppe as indicated by the abundance of forb species revealed by ancient sediment DNA between 46 and 12.5 ka on the Taymyr Peninsula (Jørgensen et al., 2012). Relatively closed landscapes with mixed coniferous and deciduous broad-leaved forests were present only in southeastern Siberia, northeastern China, and the Baikal region (Cao et al., 2019). The melting of permafrost caused elevated lake levels, which have been widely documented in Mongolia between 17 and 10 ka BP. Sandy dune formation occurred in western Mongolia during this time, along with development of steppe vegetation. Similar processes likely occurred in the Steppe Altai and south of Western Siberia (Rudaya et al., 2020).

##### I.C.ii. Early Holocene in Northern Asia

At the last deglaciation and the beginning of the Holocene, the tundra-steppe was replaced by light taiga in southern Siberia and by tundra in northern Siberia. In addition to the general increase in temperatures at the onset of the Holocene, the summer monsoon-associated maximum of precipitation around 10.8–10.5 ka BP was pronounced in Southern Siberia, northward and eastern of the Steppe Altai, and north-west China (Rudaya et al., 2020). A change in primary vegetation in Northeastern China occurred in the early Holocene (11.5 and 10.5 ka), caused by rapid increase in abundance of temperate deciduous trees, which may reflect the warmer climate and enhanced summer monsoon (Hong et al., 2009; Liu et al., 2014).

Although the extent of non-tree vegetation decreased in Siberia during the early Holocene warming (12–7 ka BP), the boreal vegetation did not show a corresponding increasing trend. The extensive and deep permafrost in northern Asia provoked a lag in vegetation response to climate change of several millennia. During the early Holocene, only sparse larch forests could survive on these shallow active-layer permafrost regions. It was not until 8 ka BP that evergreen conifer species, dominated by pine and spruce, grew in abundance, which is consistent with the increase in permafrost-free conditions (Li et al., 2022). During the early Holocene (12-8 ka BP), a sharp increase in Northern Hemisphere temperature caused permafrost to thaw rapidly, resulting in the formation of thermokarst lakes transforming landscapes in the vast expanses of Siberia. In the south of Western Siberia, a sharp reorganization of landscape appearance occurred about 11.2-10.1 ka BP when the proportion of birch forests gradually increased although steppes still prevailed in the landscape. Pine forests began to play an important role in vegetation at about 9.6 ka cal BP, apparently spreading from the Urals refugium (Ryabogina et. al., 2020).

In contrast to Central and Northeastern Siberia, the western transfer of heat and precipitation from the Atlantic began to have a more significant impact on the climate of the East European Plain from the very beginning of the Holocene. The shift from tundra and steppe vegetation to a more forested landscape is manifested during the early Holocene as a rapid increase in the abundance of trees and shrubs, including birch, pine, and oak. From 11 to 9 ka BP taiga in Europe retreated northwards; this is reflected in the appearance of deciduous taxa at higher elevations and latitudes (Binney 2017).

The global climatic event at 8.2 ka BP is relatively poorly documented in Siberia due to the rather low temporal resolution of study sites, so its impact on habitat conditions in all regions of Siberia is difficult to trace.

#### I.C.iii. Middle Holocene in Northern Eurasia

By the middle of the Holocene, an atmospheric circulation similar to the modern one had been established. The long-term general warming (Atlantic optimum) in the middle of the Holocene led to intensification of westerlies' impact on most of Europe and Siberia, against a backdrop of weakening of the Siberian High. The arrangement of biome borders across North Eurasia closely resembled the modern pattern by 8 ka BP (Binney et al., 2017). By this time, the mosaic landscapes of the early Holocene were finally structured, and a clear boundary emerged between the steppe and forest belts of western and central Asia.

During the Holocene optimum (7.5-4 ka BP), the southern boundary of surficial permafrost receded northward to near the Arctic Circle. After a rapid thawing period, there was a relatively slow and decreasing rate of thaw in the mid-late Holocene (7-2 ka BP) (Li et al., 2022). The presence of boreal evergreen conifers in the area that is now dominated by larch forests indicates that mid-Holocene winters were warmer than present in central and southern Yakutia (Tarasov et al., 1998). Around 6 ka BP, the northern forest limit in Siberia had extended poleward by several hundred kilometers north of the modern forest limit, specifically in the Yamal and Taymyr peninsulas. The mid-Holocene warming was most pronounced in Eastern Europe, where tundra was probably absent. Taiga and cool conifer forests occurred further north than the modern range of European Russia, while cold mixed forests were present in the extreme northwest of the region (northern Karelia and Kola). However, taiga had reached its current range in the continental interior (Central and Southern Siberia and northern Mongolia) (Tarasov et al., 1998).

In the southern regions, humidification in the Mid Holocene was not elevated everywhere. Winter precipitation in Central Asia remained relatively low until approximately 8 ka BP before increasing at an almost linear rate to the present level (Liu et al., 2014). Optimal conditions for vegetation and plant diversity in the Steppe Altai and Southern Siberia were recorded between 7.5–7 ka BP and ended 3.6–2.7 ka BP. A warm and arid interval was also recorded in the pollen record of the Baraba forest-steppe (Western Siberia) 7–5 ka BP. The maximum extent of mixed coniferous-deciduous forest in the south and southwest of Southern Siberia has been recorded between 7.2 and 2.7 ka BP (Rudaya et al., 2020).

In the Western Siberia forest-steppe, the driest phase of the Holocene was recorded between 7.1 and 5.5 ka BP. It was associated with an expansion of pine forests and adverse conditions for leaved trees but without signs of an increase in steppe plots in landscapes (Ryabogina et. al., 2020).

In the Southern Ural, a climatic optimum as a warm but also rather humid time was recorded at 7.4–6.3 ka BP. Detailed reconstructions of vegetation and climate dynamics in the forest-steppe interface zone of the Eastern European Plain demonstrated trajectories to temperatures warmer than the present level (by 1–1.4 C) at 7–4.8 ka BP. However, the forest-steppe boundary remained near its present position during this period because annual precipitation was similar to the current values, about 500 mm per year (Novenko et al., 2016). On the other hand, there is evidence of drier conditions that resulted in the formation of mid-Holocene buried humus soils in the forest belt (Aleksandrovskiy and Chichagova 1998). Paleohydrological studies in the central part of European Russia also showed that runoff in the Don and Dnieper basins was 40% less than modern values (Sidorchuk et al. 2012).

As insolation decreased and the impact of westerlies weakened, various regions experienced a shift towards cooler climates, dating back to about 5 ka BP. However, moisture dynamics during this period have been less clear (Wanner et al., 2008).

The treeline in Northeastern and Central Siberia retreated to the south between 5 and 3 ka cal yr BP (Binney 2017), while the cooling after 4.3 ka BP stimulated the freezing of wet mires in the middle taiga zone and formation of palsa bogs in the northern taiga and forest-tundra in Siberia. Against this background, the role of the light larch taiga increased, while the conditions for the dark coniferous taiga worsened.

After boreal woodland expansion the onset of the Middle Holocene in northern Mongolia, at the end of the middle Holocene (4.5 ka BP), there was a decrease in precipitation and changes in vegetation towards the steppe. Humidity increased again only during the late Holocene in concert with dropping temperatures (Hurka et al., 2019).

The Steppe biome in Southern Siberia and Steppe Altai also expanded as aridification began around 4.8 ka BP. Despite the gradual decrease of precipitation and the intensification of aridity, the interval of 4.5–2.5 ka BP cannot be described as xerothermal (Rudaya et al., 2020).

Climate deterioration in the south of the Ural Mountains started earlier, around 6.3 ka BP. But in the Western Siberia forest-steppe, a reduction in arid conditions started only from 5.5 to 4.9 ka BP, likely due to decreased evaporability from cooling, but there is little evidence of a significant increase in precipitation. A gradual rise in forest-steppe lake levels started only around 4.9 thousand years ago, likely due to increasing rainfall, and the proportion of birch in the forests increased while the proportion of pine decreased (Ryabogina et. al., 2019; 2020). The ratio of pine to birch in Siberian forest-steppe typically reflects changes in humidity levels, with an increase in the proportion of birch forests during more humid periods with elevated water tables, and a displacement of birch by pine during dry periods. Humid conditions were not constant, as there was also an impulse towards dryness at about 3.5–3.3 ka BP (Ryabogina et. al., 2019; 2020).

The expansion of pine in the steppe belt of Western Siberia and Kazakhstan occurred from the southern Ural and West Siberia to northern and central Kazakhstan after 5.5 ka BP. The vegetation cover approached its present-day appearance as pine spread throughout the steppe zone. During a short aridization period ca. 3.5 ka BP, forest-steppe and northern steppe biomes moved beyond their present-day borders to the north. Later, the late Holocene was characterized by the dominance of steppe with birch and pine forests along rivers (Hurka et al., 2019).

In the Eastern European Plane, cooling with increasing precipitation from 4.8–2.5 ka BP were partially responsible for changes in vegetation within the forest-steppe boundary. Woodland cover expanded, reaching 40% of the area at about 3.7 ka BP years ago. The proportion of broadleaf trees increased, spruce gradually penetrated the forests, and alder became abundant. A widespread shift of forest into forest-steppe and steppe areas has been traced by several pieces of evidence (Novenko et al., 2016). Climate reconstructions show that the subsequent phase (3.7-1.7 ka BP years ago) was marked by climate cooling and an increase in precipitation, consistent with other evidence for climate cooling in Northern and Central Europe around this time (Wanner et al., 2008).

#### I.D. References (for section I)

Li, W., Tian, F., Rudaya, N., Herzschuh, U., and Cao, X. (2022). Pollen-Based Holocene Thawing-History of Permafrost in Northern Asia and Its Potential Impacts on Climate Change. Frontiers in Ecology and Evolution, 10. https://doi.org/10.3389/fevo.2022.894471

Tarasov, P. E., Volkova, V. S., Webb, T., Guiot, J., Andreev, A. A., Bezusko, L. G., Bezusko, T. V., Bykova, G. V., Dorofeyuk, N. I., Kvavadze, E. V., Osipova, I. M., Panova, N. K., and Sevastyanov, D. V. (2000). Last glacial maximum biomes reconstructed from pollen and plant macrofossil data from northern Eurasia. Journal of Biogeography, 27(3), 609-620. https://doi.org/10.1046/j.1365-2699.2000.00429.x

Tarasov, P. E., Webb, T. III, Andreev, A. A., Berezina, N. A., Bezusko, L. G., Blyakharchuk, T. A., Bolikhovskaya, N. S., Cheddadi, R., Chernavskaya, M. M., Chernova, G. M., Dorofeyuk, N. I., Dirksen, V. G., Elina, G. A., Filimonova, L. V., Glebov, F. Z., Guiot, J., Gunova, V. S., Harrison, S. P., Jolly, D., . . . Zernitskaya, V. P. (1998). Present-day and mid-Holocene biomes reconstructed from pollen and plant macrofossil data from the former Soviet Union and Mongolia. Journal of Biogeography, 25(6), 1029-1053. https://doi.org/10.1046/j.1365-2699.1998.00236.x

Rudaya, N ., Krivonogov, S., Słowiński, M., Cao, X., and Zhilich, S. (2020). Postglacial history of the Steppe Altai: Climate, fire and plant diversity. Quaternary Science Reviews, 249, 106616. https://doi.org/10.1016/j.quascirev.2020.106616

Cao, X., Tian, F., Li, F., Gaillard, M.-J., Rudaya, N., Xu, Q., and Herzschuh, U. (2019) Pollen-based quantitative land-cover reconstruction for northern Asia covering the last 40 ka cal BP, Clim. Past, 15, 1503–1536, https://doi.org/10.5194/cp-15-1503-2019

Jørgensen, T., Haile, J., Möller, P., Andreev, A., Boessenkool, S., Rasmussen, M., Kienast, F., Coissac, E., Taberlet, P., Brochmann, C., Bigelow, N.H., Andersen, K., Orlando, L., Gilbert, M. T. P., and Willerslev, E.: A comparative study of ancient sedimentary DNA, pollen and macrofossils from permafrost sediments of northern Siberia reveals long-term vegetational stability, Mol. Ecol., 21, 1989–2003, 2012.

Hong, B., Liu, C., Lin, Q., Yasuyuki, S., Leng, X., Wang, Y., Zhu, Y., and Hong, Y.: Temperature evolution from the δ18O record of Hani peat, Northeast China, in the last 14000 years, Sci. China Ser. D, 52, 952–964, 2009.

Liu, Z., Wen, X., Brady, E. C., Otto-Bliesner, B., Yu, G., Lu, H., Cheng, H., Wang, Y., Zheng, W., Ding, Y., Edwards, R. L., Cheng, J., Liu, W., and Yang, H.: Chinese cave records and the East Asia Summer Monsoon, Quaternary Sci. Rev., 83, 115–128, 2014.

Ryabogina N.E., Afonin A.S. and Ivanov S.N. Late Glacial and Holocene in the south of Western Siberia: geochemical indices and pollen data in Kyrtyma Lake sediments. IOP Conf. Ser.: Earth Environ. Sci. 438 012023. https://doi.org/10.1088/1755-1315/438/1/012023

Binney, H., Edwards, M., Macias-Fauria, M., Lozhkin, A., Anderson, P., Kaplan, J. O., Andreev, A., Bezrukova, E., Blyakharchuk, T., Jankovska, V., Khazina, I., Krivonogov, S., Kremenetski, K., Nield, J., Novenko, E., Ryabogina, N., Solovieva, N., Willis, K., and Zernitskaya, V. (2017). Vegetation of Eurasia from the last glacial maximum to present: Key biogeographic patterns. Quaternary Science Reviews, 157, 80-97. https://doi.org/10.1016/j.quascirev.2016.11.022

Liu Z. Y., Wen X. Y., Brady E. C. et al. Chinese cave records and the east Asia summer monsoon. Quaternary Science Reviews. 2014. Vol. 83. P. 115–128.

Novenko, E.Y., Tsyganov, A.N., Rudenko, O.V. et al. Mid- and late-Holocene vegetation history, climate and human impact in the forest-steppe ecotone of European Russia: new data and a regional synthesis. Biodivers Conserv 25, 2453–2472 (2016). https://doi.org/10.1007/s10531-016-1051-8

Aleksandrovskiy AL, Chichagova OA (1998) Radiocarbon age of Holocene paleosols of the East European forest–steppe zone. Catena 34:197–207

Sidorchuk A, Panin A, Borisova O (2012) River runoff decrease in North-Eurasian plains during the Holocene optimum. Water Resour 39:69–81

Wanner, H., Beer, J., Bütikofer, J., Crowley, T. J., Cubasch, U., Flückiger, J., Goosse, H., Grosjean, M., Joos, F., Kaplan, J. O., Küttel, M., Müller, S. A., Prentice, I. C., Solomina, O., Stocker, T. F., Tarasov, P., Wagner, M., and Widmann, M. (2008). Mid- to Late Holocene climate change: An overview. Quaternary Science Reviews, 27(19-20), 1791-1828. https://doi.org/10.1016/j.quascirev.2008.06.013

Hurka H., Friesen N., Bernhardt K.-G., Neuffer B., Smirnov S., Shmakov A., Blattner F. The Eurasian steppe belt: Status quo, origin and evolutionary history. Turczaninowia, 2019. T. 22, № 3. C. 5-71 https://doi.org/10.14258/turczaninowia.22.3.1

Ryabogina, N. E., Afonin, A. S., Ivanov, S. N., Li, H., Kalinin, P. A., Udaltsov, S. N., and Nikolaenko, S. A. (2019). Holocene paleoenvironmental changes reflected in peat and lake sediment records of Western Siberia: Geochemical and plant macrofossil proxies. Quaternary International, 528, 73-87. https://doi.org/10.1016/j.quaint.2019.04.006

Olson, D. M., Dinerstein, E., Wikramanayake, E. D., Burgess, N. D., Powell, G. V. N., Underwood, E. C., D'Amico, J. A., Itoua, I., Strand, H. E., Morrison, J. C., Loucks, C. J., Allnutt, T. F., Ricketts, T. H., Kura, Y., Lamoreux, J. F., Wettengel, W. W., Hedao, P., Kassem, K. R. 2001. Terrestrial ecoregions of the world: a new map of life on Earth. Bioscience 51(11):933-938.

Map of the physical-geographical zoning of the USSR. Scale: 1:24000000. 1983. Atlas of the USSR. Maps of Nature. Physical-geographical zoning / Main Directorate of Geodesy and Cartography under the Council of Ministers of the USSR. Moscow. 1983.

Gvozdevskiy, N. A., & Mikhailov, N. I. (1978). Physical Geography of the USSR. Asian Part. 3rd ed., revised and enlarged. Textbook for students of geographical faculties of universities. Moscow: "Mysl".

### II. Archaeological context

#### II.A. East part of Central Siberia (Lena River Basin)

##### II.A.i. Middle Lena

###### II.A.i.a) Sumnagin Mesolithic culture in the Middle Lena Region (Russia_MiddleLena_M)

In 1964-1965, Yu.A. Mochanov discovered the first sites of the Sumnagin culture on the Aldan River and identified them as a specific Paleolithic culture of the Late Holocene in 1966. Later, this culture was defined as the Latest Paleolithic bu Yu. Mochanov. The multiphase sites Ust-Timpton-1 (horizons 4A to 4G), Belkachi-1 (layers 8 to 20), Sumnagin I (layers 17 to 28) are described as the main sites attributed to the Sumnagin culture. The area of the Sumnagin culture covers a vast territory in Northeast Asia from Taimyr (Tagenar-6) and the upper reaches of the Vilyui (Ust-Chirkuo) in the west to Chukotka in the east (Panteleikha-3), from the coast of the Arctic Ocean in the north (Siktyakh), including adjacent islands (Zhokhovo) and including basins of the Aldan, Vitim, and Olekma Rivers in the south and possibly even further to the south. According to numerous radiocarbon dates, the Sumnagin culture existed during the period from 10/9.5 to 6.5 kya (Mochanov, Fedoseeva, 1976). According to the recently updated chronological data, the age of the Sumnagin culture is determined as 8700–4800 calBCE (History of Yakutia, 2020).

The economy of the Sumnagin-attributed population was based on elk and deer hunting. About 90% of flint tools are made on blades extracted from prismatic cores. The blade tools include inserts, retouched knives, scrapers, angles and side burins, borers, etc. More than half of blade tools are made on microliths with a width of less than 0.4 mm. There are bone tools for inserting flint blades. Of the large tools, diabase axes-head (including ones with "ears") and adzes are typical (Mochanov, 1969; 1977; Mochanov, Fedoseeva, 1976; 2013). At the Zhokhovo site, evidence of sled dogs was obtained (Pitulko, Kasparov, 2017). Meanwhile, until now, no burials of the Sumnagin period were discovered. In this regard, a Khatystyr man for the first time can give an impression of the origins of the Sumnagin culture from the region to the south of Yakutia. For this period, in the north, from the location of Duvanny Yar in the lower reaches of the Kolyma, an isolated find of a fragment of a female skull and a series of remains at the Zhokhovo site are known (Питулько и др., 2015; Питулько, Павлова, 2015; Sikora et al., 2019).

###### (1) Khatystyr-Cave site

The Khatystyr Cave is situated on the right bank of the Aldan River, approximately 2 km from the village of Khatystyr. It is a limestone cavern characterized by a shallow entrance in the form of a hole, which developed at the location of a deep fissure.

*(a) single burial (individual ID I20533)*

In 1962, human bones were accidentally discovered by A. Ivanov, a foreman of the Aldan timber industry enterprise, in a cave near the village of Khatystyr in Yakutia's Aldan region. The skeleton, found lying on its back not far from the entrance and adjacent to the wall, was accompanied by evidence of a fire. Further exploration of the cave revealed a collection of bear, wolf, and fox bones within its depths. Initially, the bones were presumed to be recent, prompting examination by criminal investigators and, at their request, by geologist B.S. Rusanov. Subsequently, archaeologists Yu.A. Mochanov and S.A. Fedoseev also examined the cave and the skeleton. Based on their preliminary findings, the presence of modern objects (such as a metal button or buckle) near the human remains led them to conclude that the bones held no archaeological significance. Consequently, the study of the Khatystyr individual was discontinued, and the skeleton was transferred to the Geological Museum of the Yakut branch of the USSR Academy of Sciences (now the Geological Museum of the Institute of Geology of Diamond and Precious Metals of the Siberian Branch of the Russian Academy of Sciences). Nonetheless, some of the bones were retained by B.S. Rusanov for radiocarbon dating conducted in the 1970s. The results revealed remarkable antiquity of the skeleton, dating back 9800 years and making it, at the time, the oldest discovery not only in Yakutia but also in Eastern Siberia as a whole. Regrettably, these findings were not published in scientific journals and thus failed to attract the attention of researchers.

In 2017, V.M. Dyakonov reevaluated the bone remains, obtaining an AMS date of 9100 yBP. Dates were also obtained for the bones of a bear and a dog, which were slightly more recent. The scull from the burial is dated to 8285-8211 calBCE (a combination of two dates from human bones, 8287–8012 calBCE (8985±30 BP, PSUAMS-8447), and 8291-8022 calBCE (9010±30, IAAA-170069), the date of human bones is confirmed by dated dog scull from the same layer in the cave, 8284-7975 calBCE (8980±30, IAAA-183039). The dates correspond with the early period of the Sumnagin Mesolithic culture, dated to 10700-6800 calBP (8700–4800 calBCE) (Moiseyev et al. 2023).

The find of a dog scull in the bottom layer of the Khatystyr Cave (Moiseyev et al. 2023) relates the Khatystyr burial with the Early Neolithic burials with Canidae recorded in the Baikal area. At least three of them are to be listed: the Olkhon Island of the Baikal Lake, a collective burial 26 of the Shamanka-2 site, and a Late Mesolithic burial Lokomotiv-R8 that contained human bones and bones of a wolf, dated to 6374-6031 calBCE (TO-11558) (Bazaliyskiy 2012).

##### II.A.i.b) Syalakh Neolithic culture (Russia_MiddleLena_SyalakhBelkachi_EN)

As a specific culture of the Early Neolithic, the Syalakh was first identified by Yu.A. Mochanov in 1964–1966 at the Aldan multiphase sites (Mochanov, 1966; 1969). One of the first-discovered sites attributed to the Syalakh culture, the site Syalakh, which gave its name to the Early Neolithic culture of Yakttia, was discovered as early as in 1943 by A.P. Okladnikov 90 km south of Zhigansk. The main sites of the Syalakh culture are the multi-layered sites of Sumnagin-1 (layers 11 to 16), Belkachi I (layers 6-7), Ugino-1 (layer 5), Ust-Mil-1 (layer 5) on the Aldan River, Kurung-2 (layer 4), Bolshaya Kyuske (layer 6), Tensik-2 (layer 3) on the Olekma River, Khatyngnakh-2 (layer 4, 5), Suldyukar (layer 3), Talanda-2 (layer 7) on the Vilyui River, Siktyakh-1 (layer 7) on the Lena River. The area of the Syalakh culture covers a vast territory in Northeast Asia from Taimyr to Chukotka. It was formed as a result of the migration of tribes from the Cisbaikal region. Numerous radiocarbon dates obtained date the Syalakh culture in the range from 4900 to 3500 calBCE (Mochanov, Fedoseeva, 1976; 2013; Alekseev, 1987; 1996).

Together with the Syalakh culture, pottery with imprints of braided net, fully polished tools, axes and stepped adzes, and knives, were introduced to Yakutia for the first time. Among other toos, double-sided retouched arrowheads and spears, knives, adzes with core-shaped handles spread. Harpoons stand out from bone tools. Hunting for elk and deer among the Syalakh tribes was supplemented by lake and river fishing.

Numerous pieces of rock art of the Early Neolithic with images of elk and deer were discovered. The dating of these rock arts to the Early Neolithic is confirmed by sacrificial offerings in the form of flint arrowheads, scrapers, and other items that are typologically interpreted as Syalakh (Okladnikov, Zaporozhskaya, 1972; Kochmar, 1994).

Burials in the Syalakh culture were not known until the Matta burial was discovered in 1996 (Pestereva, Stepanov, Dyakonov, 2016; Zubova, Batanina, Panov, Stepanov, Kishkurno, 2017). The last two out of three radiocarbon dates showed age of 5475-5071 cal BCE (6328±81 BP,NSKA-1663) and 4442-4257 calBCE (5940±30 BP,Beta-422229) (Kılınç et al., 2021), which generally refers the Matta burial to the turn of the Early Neolithic and Mesolithic in Yakutia.Kangalassy site

The burial was accidentally discovered by workers while digging a utility pit in the center of the village of Kangalassy. Kangalassy is situated at the foot of a hill, directly beneath the high left bank of the Lena River. The area is a small hill near the mouth of the Zolotinka River.

*(a) single burial (individual ID I15934)*

During digging of a cesspit in 1958, workers came across human bones and discovered a skull, a bone knife with stone blades, four bone arrowheads, a bone awl, and several bone fragments, presumably from needles. Archaeologists were called to the site, and I.D. Novgorodov conducted excavations of the preserved part of the burial. The following information was obtained.

The burial was made at a depth of 1.2 meters from the modern surface. The bones were covered with an organic layer, possibly birch bark. The burial was oriented with the head to the east. On the femur and tibia bones, "in neat rows along the skeleton," small beads made of mollusk shells were found, marking the sewn clothing. According to the workers, similar beads were also found on the ribs.

Next to the tibia bones, on the left side of them, an irregularly shaped circle made of thick shell with two holes in the center was discovered. Three jade tools, ten arrowheads, a small flat stone fragment with a blade on one side (a cutting tool?) made of dark material, a whetstone broken into five pieces, two bone awls, and a bone knife smaller than the first one were also found in the same area. The finds were sent to a museum and initially attributed to the Bronze Age. However, after reevaluation and obtaining radiocarbon dates, researchers attributed the Kangalassy burial to the Belkachi culture. The latest obtained dates in combination (4354-4260 calBCE (2468±20 BP) [R_Combine: (5515±25 BP, PSUAMS-9131), (5400±30 BP, IAAA–170064)] suggest an earlier age and cultural affiliation with the Syalakh culture for the burial. Taking into account its relatively early chronological attribution, we refer to the Kangalassy burial as to Syalakh-Belkachi complex.

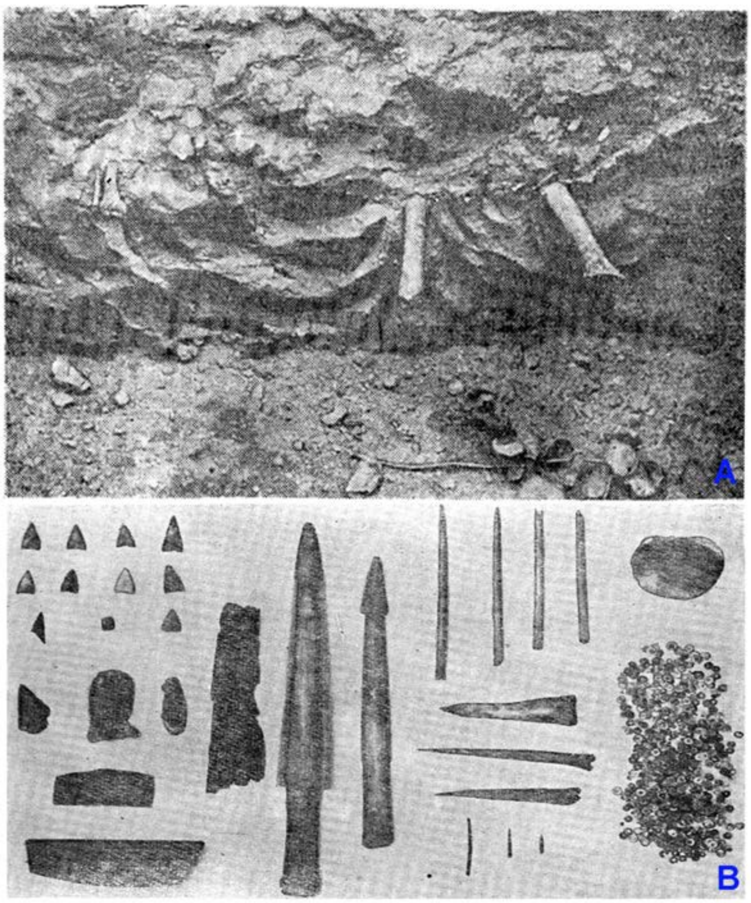

***Fig. S3.*** *The Kangalassy burial.* ***A*** *- burial at the moment of discovery;* ***B*** *- grave goods (Novgorodov, 1960).*

##### II.A.i.c) Belkachi Neolithic culture (Russia_MiddleLena_Belkachi_N)

The possibility of accurate identification of the Belkachi culture appeared only in 1964-1966 with the discovery of multilphase sites on the Aldan River, primarily the sites of Belkachi-1 and Sumnagin-1 (layers 9 to 10). At the Belkachi-1 site, under clear stratigraphic conditions, the remains of the Belkachi culture occurred in layers 4 and 5, overlaid by the Ymyyakhtak assemblage (layer 3) and underlaid by layers of the Syalakh culture (layers 6 to 7) (Mochanov, 1969; Mochanov, Fedoseeva, 1976; 2013).

The same stratigraphic situation was observed at the northernmost multiphase site in Yakutia, Siktyakh-1 on the Lower Lena, where the Belkachi materials in the 6th layer, between the Syalakh and Ymyyakhtakh assemblages occurred. The other sites of the Belkachi culture, in addition to Belkachi-1 and Sumnagin-1, on the Aldan River, are Ust-Timpton 1 (layer 3), Bilir (layer 3), Ust-Mil-1 (layer 4), Ugino-1 (layer 4), on the Vilyui River, Ust-Chirkuo (layer 2), Talanda-2 (layers 4 to 6), Khatyngnaakh (layer 2), Suldyukar (layer 2). The obtained radiocarbon dates date the Belachi culture to the period from 5200±100 to 4100±100 BP (Mochanov, Fedoseeva, 1976; 2013). Recently updated dates show the chronological interval of 4200-2100 calBCE (History of Yakutia, 2020).The area of the Belkachi culture also covered the extensive territory of Northeast Asia – from the tributaries of the Yenisei River and the extreme limits of Chukotka to the Amur in the south.

Distinctive features of material culture are pottery with impressions of a cord, a pointed or rounded bottom, with a rim ornamented with smooth or comb-shaped depressed belts. Some vessels have a molded rim. Stone tools are characterized by polished stepped adzes, chipped and partially polished axes, adzes with ledges-ears, arrowheads with asymmetric stingers or stalked arrowheads on plates, multifaceted cutters-drills, beak-shaped combined tools, polished knives, bilaterally retouched spearheads and darts, bone liner tools (вкладышевые орудия) ets.

Monuments of rock art are more numerous in comparison with the Early Neolithic. The rock art is still characterized by images of animals - elks, deer, slightly different in details and styles from the drawings of the Early Neolithic. Apparently, the first anthropomorphic images appeared during this period. Under the petroglyphs there are sacrificial offerings made of stone and bone tools, characteristic of the Belkachi culture (Okladnikov, Zaporozhskaya, 1972; Kochmar, 1994).

In the Belkachi culture one Tuoy-Khaya cemetry on the Vilyui River is known (Fedoseeva, 1968; Mochanov, Fedoseeva, 2013). On the Lena River the burials of the Kangalassy, Uolba, Ogonyok are known, on the Amga R. – Khayirgas and Onnyos, Dzhikimda burial – on the Olyokma R., Rodinka burial – Kolyma R. A distinctive and most important feature of the Belkachi burials is the filling with ocher. The grave inventory contains bone and stone inventory, among which are Belkachin flint arrowheads, bone composite arrowheads, polished adzes, bone composite tools, daggers, piercings, awls, bone jewelry, rich decorations for clothes made of shells, small plastic figurines (small sculptures), various animals, including a dog, as in the Onnyos burial.

One of the main problems of the Belkachi culture is their ancestral genetic connection with the modern population of the north of Eastern Siberia and Alaska. Yu.A. Mochanov assumed a connection between the Belkachi population and the Na-Dene Indians, and at the same time, assimilation processes with the Ymyyakhtakh people were assumed (Mochanov, 1969; Mochanov, Fedoseeva, 2013).

The Belkachi culture (4200–2100 calBCE) is among the largest in Northeast Asia, with its technocomplex being present not only in Yakutia but also in other regions such as Lower Amur, Northern Sakhalin, Taimyr, and Chukotka. Some opinions suggest that cultural influences of the Belkachi culture of taiga hunters and fishermen extended to America, playing a role in the development of the Arctic small tool (also known as Paleo-Eskimo or Paleo-Inuit, Flegontov et al. 2019) tradition in Alaska, Canada, and Greenland. Belkachi burials have been discovered along the Lena, Vilyui, Olekma, Amga, and Kolyma rivers. These burials share several common characteristics, including soil grave pits without masonry, backfilling with ocher, extended supine position of the deceased, and the presence of abundant accompanying grave goods, often including bones of animals and birds (Alekseev et al., 2022).

##### (1) Ogonyok site

The burial known as Ogonyok was discovered in October 2016 within the premises of a summer cottage in Yakutsk, located 8.7 km south-southwest of the city center, during the excavation of a utility pit using an excavator. The burial is situated on the southeastern slope of the second floodplain terrace of the Lena River. In close proximity to the burial site lies the Late Neolithic site Ogonyok, discovered in 2001, which is attributed to the Ymyyakhtakh culture. The burial itself is situated in the northern part of the terrace, alongside the habitation area. The territory of the monument, once covered by a pine forest, is now largely occupied by suburban structures (Alekseev et al., 2022).

###### *(a) single burial (individual ID I23448)*

Upon initial examination, it was observed that the burial had been severely damaged by the power shovel. The skull and artifacts were recovered from the pit's bottom and spoil. The skull, in a deteriorated state, retained the parietal and frontal bones, while the facial and occipital parts were missing. The interior surface of the vault exhibited dense red ochre pigmentation. Additionally, a human tibia with traces of ochre and a fragment of a collarbone were found at the pit's bottom.

Excavations focused on the surviving portion of the grave. This allowed to conclude tentatively that the grave pit had an oval contour. An ochre layer was identified within the preserved portion of the grave's fill and on its base. Based on the impression of the skull on the pit wall, it was deduced that the individual was interred on their back, with the head oriented towards the southeast, facing the nearby oxbow lake upstream of the Lena River. The grave goods discovered *in situ* included an ochre-dyed bone needle located to the right of the individual's head. Within the fill of the grave pit, two fragments of small animal bones, likely remnants from a funerary feast, were uncovered. Detailed sieving of the spoil from modern construction yielded accompanying materials from the Ogonyok burial. These included a core, three flint knife-like plates, an adze, a fragment of a polishing stone, a fragment of a composite arrowhead with a flint insert, and an anthropomorphic figurine carved from mammoth tusk.

Four radiocarbon dates were obtained from the Ogonyok burial through AMS at the Institute of Accelerator Analysis, Japan. The human skull bone was dated to 4241-4001 calBCE (IAAA-170070) and the tibia show the date  4050–3967 calBCE (5210±20 BP, IAAA-170063). Additionally, two AMS 14C dates were obtained from animal bones found in the accompanying inventory of the burial. One of them, derived from a lynx canine, yielded a date of 3970–3790 calBCE (5090±30 BP, IAAA-162746). Similarly, the other date obtained from a reindeer bone was 3960–3790 calBCE (5070±30 BP, IAAA-162747). It is worth noting that the dates obtained from the animal bones are essentially identical (Alekseev et al., 2022). The intervals of dates obtained from human and animal bones do not overlap and we suggest better using the combination of only animal-derived dates to prevent possible freshwater reservoir offset of C14 dates, which is often recorded in the Lena region. Thus, the burial is dated to 3957-3799 calBCE (5080±22 BP) [R_Combine: (5090±30 BP, IAAA-162746), (5070±30 BP, IAAA-162747)].

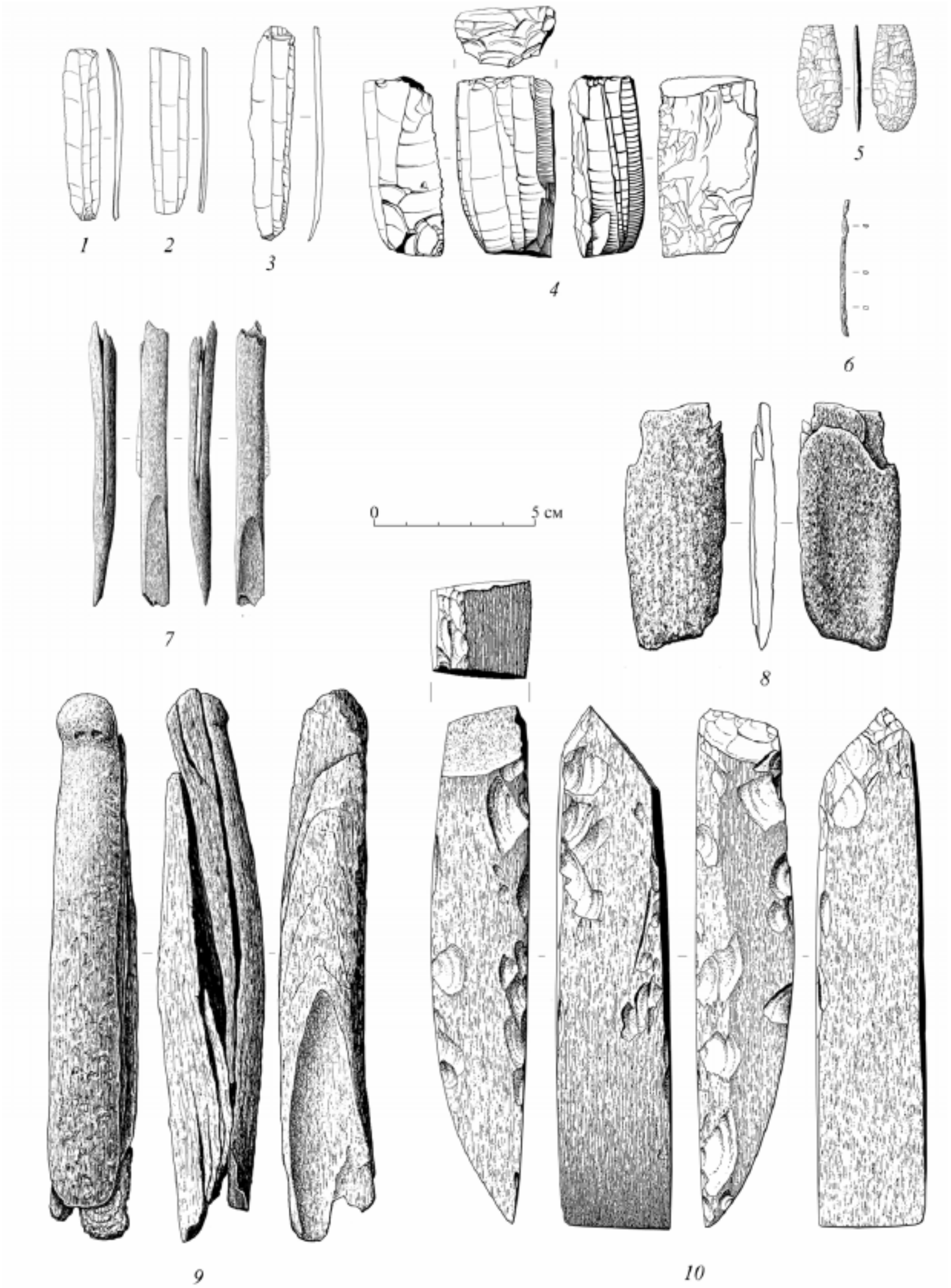

***Fig. S4.*** *Grave goods from the Ogonyok burial.* ***1-3*** *- blades;* ***4*** *- prismatic core;* ***5*** *- flint arrowhead;* ***6*** *- bone needle;* ***7*** *- bone composite arrowhead with a blade in the groove;* ***8*** *- bone polisher;* ***9*** *- anthropomorphic figurine made from a mammoth tusk;* ***10*** *- polished chisel (Alekseev et al., 2022).*

##### II.A.i.d) Ymyyakhtakh Neolithic culture (Russia_MiddleLena_Ymyyakhtakh_LN)

One of the first Late Neolithic sites was discovered by A.P. Okladnikov in 1942 on the Lena River, at the Ymyyakhtakh site, which later became the eponym of a special culture. Based on the materials of this site, A.P. Okladnikov singled out a special stage of the Late Neolithic of the Middle Lena cultural region [Okladnikov, 1950; 1955]. For the first time, a pure complex of the Ymyyakhtakh culture was identified by Yu.A. Mochanov in 1964-1965 at the multilayer sites of Belkachi I and Sumnagin I on the Aldan River [Mochanov, 1969]. The age of the Ymyyakhtakh culture is determined by radiocarbon dates 4100±100 – 3300±100 BP [Mochanov, Fedoseeva, 1976]. According to the latest updated data, the age of the Ymyyakhtakh culture is determined as 2950-1150 calBCE [History of Yakutia, 2020, p. 478].

This culture occupied a vast area. According to the researcher of Chukotka M.A. Kiryak: "The expansion of the Ymyyakhtakhs was global" [Kiryak, 1993]. In the west, the Ymyyakhtakh culture almost reached the Yenisei River, in the north it occupied Taimyr, in the east it extended to the Northern Okhotsk and Eastern Chukotka regions, in the south – to the Baikal and Transbaikalia.

At the Ymyyakhtakh sites, characteristic multilayer waffle, ribbed, and smooth-walled ceramics with an admixture of organic matter (animal wool, grass, and needles) were found in clay dough [Mochanov, 1969; Fedoseeva, 1980]. In addition to clay and organic matter, the dough contains sand, gruss, fireclay, small pebbles and crushed quartzite. An obligatory element of the ornament of this pottery is a belt of through rounded holes. The artistic ornament on the vessels also includes incised geometric patterns in the form of intersecting lines, zigzags and triangles.

A characteristic feature of the stone inventory of the Ymyyakhtakh people was the thoroughness and filigree processing of tools, and the clear consistency of their forms. From various types of stone, they made all kinds of products – arrowheads and spears, scrapers, cutters, inserts, knives, abrasives, axes, adzes, etc. Ymyyakhtakh people were familiar with all the basic stone processing techniques mastered in previous eras – retouching, polishing, drilling, sawing, separation of plates from cores, etc. In general, stone processing reached its peak during this period. A significant place in the inventory of the Ymyyakhtakh people was occupied by tools made of bone and horn, from which hunting and fishing tools, weapons, protective weapons, jewelry, and household items were made [Fedoseeva, 1980].

In the rock art of the Late Neolithic, images of a person acquired paramount importance, animalistic drawings receded into the background. Anthropomorphic figures are shown standing, dancing, in motion, in action, with raised arms, half-sitting. All of them are stylized to some extent. The drawings were made in red ocher of various shades [Okladnikov, Zaporozhskaya, 1972; Kochmar, 1994].

Two large Ymyyakhtakh cemetery are known – Chochur-Muran and Deering-Yuryakh on the Middle Lena [Fedoseeva, 1980; Fedoseeva, 1988; 1992; Mochanov, Fedoseeva, 2013], as well as several single and double burials – Ichchilyakh on the Lower Lena [Okladnikov, 1946], Vilyui Shosse in Yakutsk [Dyakonov et al., 2003; Dyakonov, Kuzmin, Hodgins, 2012]), burials of Pomazkino, Kamenka 2 on the Kolyma River [Kashin, Kalinina, 1997; Kashin, 2013; Kilinc et al., 2021]. Burial of Ergetsy is known in Vilyui River [Arkhipov, unpublished]. In Central Yakutia the Kyordugen burials are known, in which a warrior with a shield made of bone plates and armor made of horn plates was buried, provided vivid material that allows us to reconstruct the details of the social structure and military affairs of the ancient population of Yakutia at the end of the Stone Age [Alekseev, Zhirkov, Stepanov, Sharaborin and Alekseeva, 2006; Fyodorova, 2008; Stepanov, Kuzmin, Hodgins and Jall, 2012; Kilinc et al., 2018; 2021].

It is believed that the Ymyyakhtakh people took a direct part in the formation of the Yukagir and Nganasan ethnic groups and the Yukagir culture (Kiryak, 1993; Khlobystin, 1998; Everstov, 1999; 2014).

##### (1) Chuiya site

The Chuiya site is situated on a cape-like ledge of a 6-meter terrace along the lakeshore, which is intersected by a modern highway. This cape extends between two expansive basins. Lake Chuiya occupies the southern basin, while the village of Chuyayu is situated in the northern basin. Located 300 meters west of this cape, the Maya River emerges from Lake Chuiya, flowing westward, bypassing the village of Maya, and eventually merging into the larger Lake Maya. The Maya-Chuyuyinskaya lake system forms part of the Lena-Amga interfluve, characterized by typical landscapes of alas-lakes and erosion-accumulative landforms. In 1992, residents of the village of Chuiya discovered human bones in an erosive thermokarst ravine, formed as a result of road construction in the 1970s (Stepanov et al., 2021).

*(a) destroyed burial (individual ID I20528)*

The burial site was investigated by ethno-archaeologists N.P. Prokopiev and E.K. Zhirkov. Unfortunately, the skull and part of the upper shoulder level were lost, likely due to thermal erosion of the terrain following bulldozing during road construction. Part of the skeleton was exposed on the eroded surface of the ravine slope, and the femur was discovered by local children. Ten meters west of the burial, at a depth of 27 cm, in a small rounded pit, a human mandibula was found, accompanied by an Ymyyakhtakh-type needle case made of thin-walled hollow bone with decorative horizontal carved lines" (Prokopiev, 1994).

Initially, in 1992, the investigators assumed that the mandibula and the skeleton belonged to the same burial, and the burial was tentatively assigned to the Neolithic period based on the presence of the needle case. However, after excavations conducted in 2010, and considering analogies in the position of the side-oriented skeleton, it was proposed that the burial might be attributed to the Early Iron Age. This hypothesis was subsequently supported by radiocarbon dating, which dated the postcranial skeleton of the buried individual to the Early Iron Age, specifically between 753 and 409 calBCE. The mandibula underwent two separate dating processes, both indicating a Neolithic age: 1957–1750 calBCE (3540±30 BP, Beta-432254) (Stepanov et al., 2021) and 1882–1699 calBCE (3475±25 BP, PSUAMS-9018). These two age determinations give in combination the date 1888-1750 calBCE.Therefore, two burials were discovered in the erosional thermokarst ravine: one from the early Iron Age and the other (the mandibula and the needle case) and from the Late Neolithic Ymyyakhtakh culture. The second burial underwent archaeogenetic analysis (individual ID I20528).

#### *II.B. South part of Central Siberia and the Baikal Region*

##### II.B.i. Transbaikalia, Cisbaikalia, and Upper Angara

###### II.B.i.a) Early Neolithic in the Transbaikalia (Russia_Fofonovo_Kitoi_EN)

###### (1) Fofonovo burial site

The Fofanovo burial ground is a significant and ancient burial complex in the Baikal region, consisting of three distinct groups of burials from different periods, namely the Early Neolithic and the Bronze Age. The site was initially explored in 1926 by A.P. Okladnikov, followed by excavations conducted by M.M. Gerasimov in 1936 and 1959, A.P. Okladnikov in 1948 and 1950, V.P. Konev from 1987 to 1991 and 1996, and since 2007 by E. Zhambaltarova. The cemetery is situated on the right bank of the Selenga River, approximately 0.5 km upstream from the village of Fofanovo, in the Kabansky district of the Republic of Buryatia. The burials are found on the southwestern, southeastern, and eastern slopes of Fofanovskaya mountain.

M.M. Gerasimov and E.N. Chernykh, based on the stratigraphy, planigraphy of the burial ground, and the characteristics of the burial rites, identified three distinct groups of burials.

The first group comprises 28 Early Neolithic (Kitoi) burials: 1-7 (1959), 1-17 (1987-1989 and 1996), 1, 3-5 (2007, 2008, 2013), and a ritual object with a bear skull (2008). This group is characterized by the absence of tomb structures, shallow burial depth (ranging from 0.5 to 1.4 m from the contemporary surface), oval and approximately rectangular pit shapes, deceased individuals placed on their backs or sides with bent legs, southeast orientation of the bones, and the presence of ocher in the backfill. Radiocarbon dates based on bone samples from these burials fall within the range of 7036-6073 calBCE (7610±210, GIN-4477) to 5471-5215 calBCE (6350±50 yBP, GIN-4128) (Mamonova, Sulerzhitsky, 1989).

The second group of the Fofanovo burial ground consists of 29 Glazkovo burials: burials 8-36 (1959) and 6 (2013). Many of these burials feature stone embellishments above the pits. The deceased individuals are positioned on their backs in an extended or bent leg position. The burial pits are typically long (up to 2.8 m) and narrow (0.5-0.8 m) in shape. The grave goods include arrowheads, scrapers, knives, horn and bone artifacts, metal tools, ceramic fragments, and more. Radiocarbon dating of bone samples from this group of burials yielded dates ranging from 2196-1937 calBCE (3670±40 BP, GIN-4473) to 2908-2352 calBCE (4100±100 BP,GIN-4803) (Mamonova, Sulerzhitsky, 1989, pp. 23-24).

The cultural and chronological attribution of the third group of burials, consisting of burials 37-41 (1959), remains undetermined. These burials lack tomb structures, and the pit contours were not clearly delineated. The skeletons were found at a depth of 0.45-1 m from the modern soil level. The deceased individuals were laid on their backs with bent knees, and the orientation varies from northeast to southeast. Only a needle case and fragments of a bone-based dagger were discovered in the graves of this group. According to M.M. Gerasimov and E.N. Chernykh, these burials were made at a later time than the Glazkovo burials (Zhambartalova and Volkov, 2016; Lbova et al., 2008; Gerasimov and Chernykh, 1975).

*(a) imprecisely known burials (individual IDs I12488, I8501, I12961, I8535)*

We conducted sequencing on four individuals originating from the Fofonovo burial site, specifically recorded as coming from **excavation 4, burial 1** (individual ID I12488); **excavation 4, burial 7** (I8501)**; excavation 4, burial 10** (I12961)**; and excavation 7** (I8535). Regrettably, these burials cannot be precisely matched with existing published sources, precluding a comprehensive assessment of their archaeological context. The individual ID I12961 underwent radiocarbon dating, yielding an estimated age range of 5883-5720 cal BCE (6905±35 BP, PSUAMS-13169), which aligns with the established chronological framework of the site and suggests Early Neolithic (Kitoi) attribution of the studied burials.

#### II.B.i.b) Kitoi burial tradition and other Early Neolithic Burial Traditions (Russia_Cisbaikal_Kitoi_MN)

##### (1) Rasputino site

The village of Rasputino is situated on the left bank of the Angara River, 450 km downstream from the city of Irkutsk, on a 15-meter terrace that extends from the mouth of the Igirma River and extends far beyond the village. Upstream from the village, the terrace is intersected by numerous waterholes and rises steeply above the floodplain. It gradually merges with the upland of the bedrock coast to the southeast. Over the years, extensive plowing has caused significant damage to the terrace slopes and resulted in the destruction of ancient burials. However, two burials were accidentally discovered by local residents on the sides of waterholes.

*(a) burial 2(?) (individual ID I13679)*

The skeletal code of the individual ID I13679 in the Tomsk Anthropological Collection refers to **mound 13, burial 2**, excavated by G.P. Sosnovskiy in 1921. However, G.P. Sosnovskiy never excavated any mounds and even burials in Rasputino; he visited the village and collected information about two burials found by E.G. Rasputin, a resident of the Rasputino Village, in 1903 (Okladnikov 1976). We suppose mislabeling happened and treat individual I13679 as a skeleton from burial 2 at the Rasputino site.

According to G.P. Sosnovsky's account, which was collected from E.G. Rasputin in 1921, the burial was initially found in 1903, located one verst (~1.1 km) from the village upstream along the Angara River, in a hollow on the cultivated land of E.G. Rasputin, approximately 1/2 arshin (~35 cm) from the first burial.

The bones of the skeleton from this burial were found to be stained red with bloodstone. The burial orientation was similar to the first one, with the skull facing northwest and the body in a supine position. According to G.P. Sosnovsky, the following items were discovered in this burial:

1. A bone needle case made from a tubular bird bone, containing two bone needles decorated with a longitudinal ornament of zigzag rows of dashes.
2. Fragments of a tetrahedral bone stick with 57 parallel notches on one of the narrow faces (possibly a counting stick or "tag"). One end of the stick is tapered.
3. Bone awls (4 pieces).
4. A curved dagger made of horn, serving as a piercing tool.
5. Two small wedge-shaped jade axes.
6. Two oblong jade knives with pointed edges.
7. A jade chisel.
8. Two sandstone whetstones.
9. Two oblong pebbles used as pestles for grinding pigments.
10. A chisel-shaped bone tool.
11. Pieces of red ocher.
12. Several fragments of bone tools and unfinished bone artifacts.
13. Additionally, two wild boar tusks were found resting on the neck of the skeleton. The artifacts were positioned along the sides of the skeleton (Okladnikov 1976).

Unfortunately, the collection is lost. The funeral rites and the typology of the finds support the Kitoi attribution of the burial (Georgievskaya 1989). The burial is dated to 5731–5635 calBCE (6800B±30 BP, PSUAMS-9118).

##### (2) Manzurka-2 site

The burial ground was on the Manzurka River, on the second terrace above the floodplain. The elevation of the terrace ranged from 1 to 10 m above the water's edge. In 1974, A.P. Okladnikov conducted excavations, uncovering 5 burials in total (Aseev, 2009).

###### *(a) burial 2 (individual ID I0999)*

**Burial 2 (**Individual ID I0999) Burial 2 (Individual ID I0999) was positioned on the slope of the terrace, approximately 100 m south of burial 1. Three small stones protruding from the turf marked its presence on the surface. Upon removing the sod, the complete outline of the mound became visible. The mound exhibited a semi-oval shape and was constructed using small, slightly rounded pieces of limestone compactly laid out in a single layer. The dimensions of the stone mound measured 90 cm from north to south and 50 cm from west to east. Subsequent excavation beneath the mound's base, at a depth of 10 to 12 cm, revealed the femurs and leg bones, indicating that the buried individual was laid on their back and oriented towards the north. The upper half of the skeleton was absent, suggesting evidence of looting. However, within the pelvic region on the right side, a miniature arrowhead with a tooth-like decoration and deep recesses just above the straight attachment on the tip was discovered. It is presumed that this tooth-like feature served as a means to secure the tip to the shaft, which was split from above. Furthermore, during the excavation beneath the skeleton, a small plate knife made of green

Baikal jade was found. One half of the oval blade exhibited delicate retouching on both sides (Aseev, 2009). The burial is radiocarbonally dated to 5831–5673 calBCE (6870B±25 BP, PSUAMS-4215).

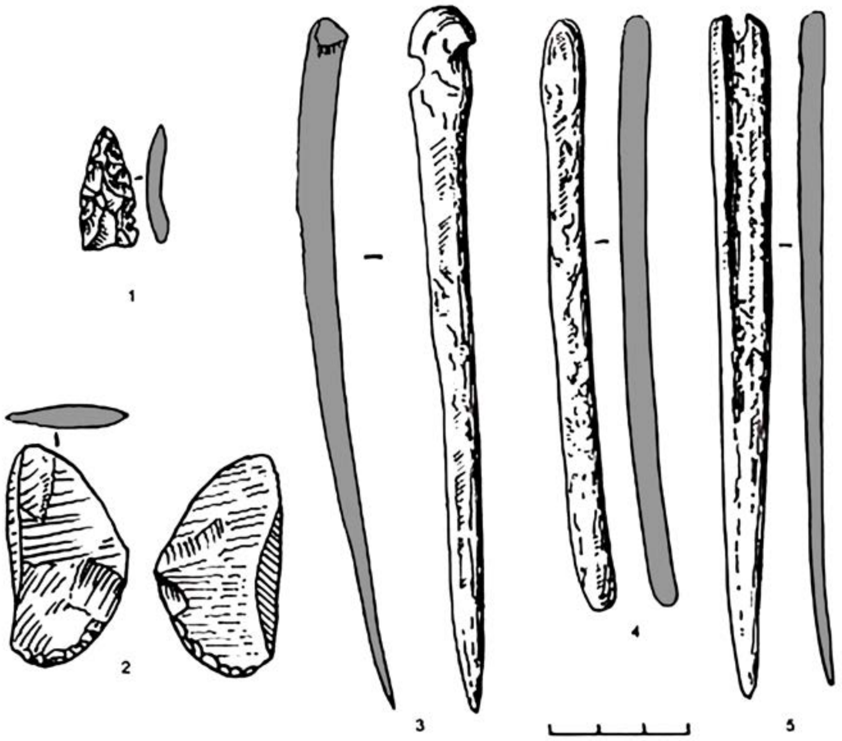

***Fig. S5.*** *Burial 2 at the Manzurka-2 site. Grave goods from the burial (Aseev, 2009).*

#### II.B.i.c) Serovo and Isakovo Neolithic burial traditions (Russia_Cisbaikal_SerovoIsakvo_N and Russia_AngaraRiver_SerovoIsakvo_N)

##### (1) Ust'-Belaya site complex

Situated at the confluence of the Belaya River into the Angara River, approximately 110 km northwest of Irkutsk, the Ust-Belaya site represents a multi-phase archaeological complex. Notably, it was subjected to investigations by various scholars at different times. N.N. Gurina and L.Ya. Krizhevskaya conducted initial investigations in 1957, followed by G.I. Medvedev and G.M. Georgievskaya in 1962, and subsequently by G.M. Georgievskaya from the late 1960s to the 1990s (Georgievskaya 1989).

The site has yielded a diverse assemblage of over 20 cultural ceramic complexes, primarily associated with the Neolithic epoch, alongside artifacts from the Bronze and Iron Ages. Notably, burials have been identified in both the northern and southern sectors of the site. The northern area revealed four investigated burials, while the southern sector unveiled and excavated 17 burials.

These burials encompass distinct groups that are attributed to the Kitoi, Serovo-Isakovo, and Glazkovo complexes. The Kitoi burials were published by Georgievskaya in 1989.

###### *(a) Unknown/unidentified burials (Individual ID I8296 and I8298)*

We publish genetic results from two burials from the Ust'-Belaya site, individual IDs I8296 and I8298. Unfortunately given the state of the records, it is not possible to attribute them to specific burials from this site. Radiocarbon analysis of individual ID I8296 yielded an estimated age range of 2860-2570 calBCE (4105±20 BP, PSUAMS-5480), while individual ID I8298's dating indicated an age range of 3790-3690 calBCE (4970±20 BP, PSUAMS-5481). The date of ID I8298 aligns within the chronological span of the Late Neolithic burial practice of the Serovo and Isakovo traditions, while the date of individual ID I8296 suggests the Glazkovo cultural attribution.

#### II.B.i.d) Glazkovo Early Bronze Age culture (Russia_Cisbaikal_Glazkovo_EBA and Russia_AngaraRiver_Glazkovo_EBA)

##### (1) Obkhoy burial ground

The Obkhoy burial ground, spanning the Early Neolithic to Bronze Age, was brought to light during archaeological investigations along the Upper Lena River by A.P. Okladnikov in 1971. It occupies the southwestern portion of the right estuarine region of the valley, with an elevation of approximately 15 meters above the water level of the Kulenga. Upon initial examination, the site had already experienced partial degradation due to human activity.

Based on distinct burial customs and accompanying artifacts, Okladnikov attributed this site to the Glazkovo culture, although no publications were made on the findings. Resuming research efforts in 2017-2018 under the guidance of V.I. Bazalinsky, further exploration was conducted. During the 2018 phase of the project, two burials, presumably from the Early Neolithic period, were identified along the borders of the previous excavations.

###### *(a) Burial 7 (Individual ID I1000) and burial 13 (individual ID I1001)*

We publish genetic results for two burials from the Obkhoy burial ground, burial 7 (individual ID I1000) and burial 13 (individual ID I1001). The skeleton from burial 7 is dated to 2872-2581 calBCE (4132±32 BP) [R_combine: (4100±40 BP, Poz-83436); (4180±50 BP, GIN-4121)](Mamonova and Sulerzhitskiy, 1989). Human bones from burial 13 are dated to 3087-2895 calBCE (4350±36 BP) [R_combine: (4340±50 BP, Poz-83437); (4360±50 BP, GIN-4122)] (Mamonova and Sulerzhitskiy, 1989).

##### (2) Ulyarba site

The Ulyarba burial ground is situated in the Olkhonsky district of the Irkutsk region, 188 km northeast of the city of Irkutsk and 4 km southwest of the village of Sarma. The ancient burials were positioned on the slope of the mountain, at the base of Cape Antukhay on the northwest coast of Mukhor Bay in the Small Sea of Lake Baikal. L.P. Zyablin discovered and partially investigated the burials in 1959, with a total of 28 burials excavated. Based on the burial locations and certain ritualistic characteristics, L.P. Zyablin identified four burial groups as Ulyarba 1-4 cemeteries. Within the Ulyarba-1 burial ground, the discoverer attributed 5 burials (1-3, 16, 19), four of which were aligned in a row along the slope at its base. The primary concentration of burials, situated 50 m up the slope from the Ulyarba-1 burials on the next terrace-like ledge, was designated as Ulyarba-2. L.P. Zyablin identified Ulyarba-3 (burials 25-28) 15 m to the north of the latter, and Ulyarba-4 (burials 21-22) 60 m to the west. The Ulyarba 1-2 cemeteries were assigned to the Bronze Age, Ulyarba-3 to the Serovo culture of the Neolithic period in the Baikal region, while the dating of the Ulyarba-4 site was not determined. In subsequent years, the Ulyarba-4 burials were attributed to the Slab Graves of the Early Iron Age. In 1976, O.I. Goryunova resumed excavations of the Ulyarba burial ground, unearthing 6 burials, and the site was later studied by V.I. Smotrov in 1983 (6 additional burials were unearthed). Based on their locations, the excavated burials from 1976 and 1983 were assigned to the Ulyarba-2 burial ground, although some of them, due to the characteristics of their burial complexes, can be traced back to the Neolithic period (Serovo culture). Since the Neolithic and Bronze Age burials were situated on the same territory, O.I. Goryunova chose not to separate them into distinct burial grounds, but considered them as a single chronologically diverse necropolis. In total, 40 burials from the Neolithic to Paleometallic periods were unearthed at the burial ground throughout the years of investigation (Goryunova 2002).

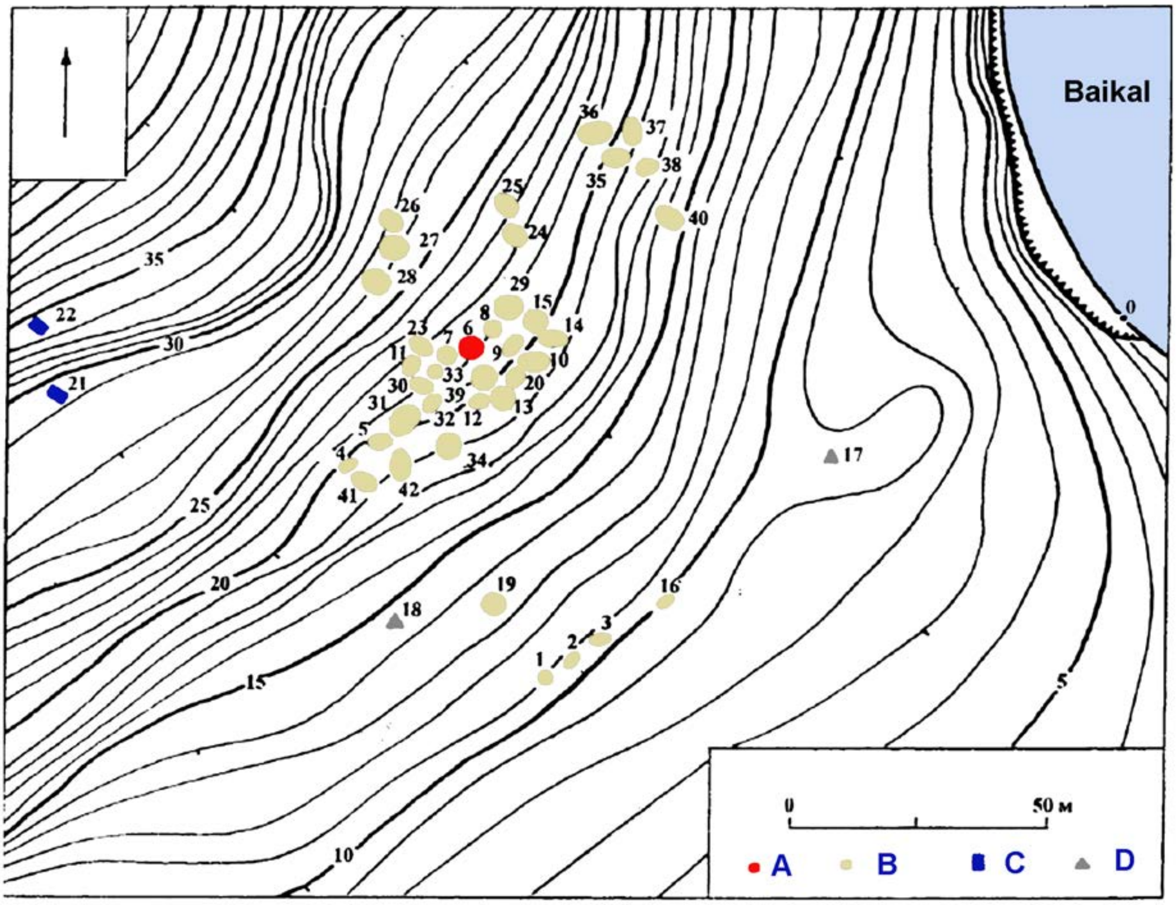

***Fig. S6.*** *Ulyarba site.* ***A*** *- burial 6;* ***B*** *- other Neolithic and Bronze Age burials; C - Early Iron Age burials; D - rocks (Goryunova 2002, modified).*

*(a) Burial 5 (Individual ID I2133)*

Burial 5 (Individual ID I2133) enclosed with a rubblework structure aligned in the southwest-northeast direction, constructed by arranging multiple layers of rock slabs. The burial was also covered by horizontally laid stone slabs inside the grave pit. The grave pit itself follows the west-southwest-east-northeast orientation. The accompanying funerary assemblage is not specified (Goryunova 2002). The burial is dated to 2623–2458 calBCE (4000±35 BP, Poz-82201).

*(b) burial 6 (individual ID I2135)*

**Burial 6** (Individual ID I2135) enclosed with a rubblework structure aligned in the southwest-northeast direction, constructed by arranging multiple layers of rock slabs. The grave pit itself follows the west-southwest-east-northeast orientation. Within the grave, the remains of an adult individual were found, positioned supine with flexed knees, and adorned with a significant application of ocher pigment. Although the accompanying funerary assemblage was present, its composition is not specified (Goryunova 2002). The burial is dated to 2864–2496 calBCE (4092±32 yBP, OxA-33162).

**(3) Khuzhir site**

*(a) burial 2 (individual ID I0998)*

From the Khuzhir burial site, we publish genetic data on the individual ID I0998 from **Burial 2** (Individual ID I0998), dated to 2836–2468 calBCE (4040B±35 yBP, Poz-83426). A combination with a previously published date from the same burial (2871–2610 BCE (4140±50 yBP, GIN-4098) (Mamonova and

Sulerzhitskiy, 1989) demostrates chronological interval 2851-2491 calBCE (4073±29 BP) [R_Combine: (4040±35 BP, Poz-83426), (4140±50 BP, GIN-4098)].

#### II.B.ii. Lower and Middle Angara

##### II.B.ii.a) Early and Middle Neolithic sites (Russia_MiddleAngara_Kitoi_MN)

###### (1) Sosnovy Mys site complex

The Sosnovy Mys site (or archaeological site complex) was first discovered and explored by N.I. Drozdov in 1974-1975. The term refers to a multiphase occupation site and two burial grounds, Sosnovy Mys 1 and 2. The site complex is located on a 6-8 meter alluvial terrace on the left bank of the lower part of Sosnovy Island, stretching along the course of the Angara River, opposite the mouth of its right tributary, the Kata River. The island's surface is densely covered with pine forest, moss, and grassy litter.

***The Sosnovy Mys-1 burial ground*** was explored and excavated by N.I. Drozdov in 1974-1975, and the excavations revealed 8 burials. One more burial was excavated in 1987 by V. I. Privalikhin. Archaeological excavations resumed in 2011-2012 as part of preparations for the flooding of the Boguchanskaya hydroelectric power station, uncovering over 8,000 m2 of the site. The investigations revealed a complex multi-layered habitation area and a burial ground spanning different periods, totaling 22 burials. Currently, there are 31 excavated burials at the site, with at least 16 of them belonging to the Neolithic period (Savelyev et al., 2020; Petrova et al., 1976; Privalikhin et al., 2013).

*The* ***Sosnovy Mys-2 burial ground.*** In 1982, V.I. Privalikhin conducted reconnaissance investigations in the lower section of the island, leading to the identification of a new archaeological site named Sosnovy Mys-2. These investigations yielded the discovery of three burials, with one attributed to the Bronze Age, another to the Early Iron Age, and a third burial, potentially dating back to the Neolithic period, which unfortunately had been destroyed. The Sosnovy-Mys-2 burial ground is situated on a 10-12 meter terrace on the left bank of the island, 2 km above its lower end, where Sosnovy Mys-1 is located (Savelyev et al., 2020; Petrova et al., 1976; Privalikhin, 1998; Privalikhin et al., 2013).

###### *(a) Sosnovy Mys-1, burial 5 (individual ID I2134)*

**Burial 5** (individual ID I1234) was discovered in 1974 at a depth of 1.19 meters from the surface. The skeletal remains were found in a supine position with the body fully extended. The upper portion of the skeleton was partially disturbed by the grave pit of burial 4, while the lower part was covered with a layer of ocher. The funerary assemblage associated with the burial includes a flint knife, an unfinished arrowhead made of chalcedony, an ornament crafted from a musk deer tusk, a fragment of a bone projectile point, and a pendant made from a deer tusk (Petrova et al., 1976). There were no traces of rubblework above the grave, and it was overlaid by another burial, where the rubblework was recorded. This fact supports the attribution of burial 5 at Sosnovy Mys-1 to the earliest phase of the site, probably related to the Kitoi period. The burial is dated to 5718–5556 calBCE (6710B±40 BP, OxA-33163).

##### II.B.ii.b) Serovo and Isakovo Neolithic burial tradition (Russia_MiddleAngara_SerovoIsakovo_LN)

###### (1) Sosnovy Mys site complex

*(a) Sosnovy Mys-1, burial 9 (individual ID I1961)*

**Burial 9** (Individual ID I1961) began at a depth of 25 centimeters from the soil surface, a compact arrangement of stones in the shape of an elongated oval was discovered, stretching from the southeast to the northwest and encircled by a ring of large stones and slabs. Digging further, at a depth of 70 centimeters from the contemporary soil surface, the skeletal remains of a buried man were uncovered. The preserved dimensions of the grave pit measure 133x64x42 centimeters. The skeletal position of the man indicates that he was laid on his back, with arms extended parallel to the body, in an extended posture. The burial orientation was observed with the head facing southeast and the legs pointing northwest, corresponding to the downstream direction.

Accompanying the male skeleton were 16 grave goods, comprising various items and tools crafted from stone, horn, bone, and ceramics:

1. A chisel made of silicified shale, measuring 16.7 x 4.7 x 3.2 centimeters, was found on the bones of the chest and the thoracic spine and was turned with the dorsal side up.
2. Three sherds of a single pottery vessel adorned with dotted-comb ornamentation and pits.
3. A perforator fashioned from the slate bone of an adult elk.
4. Two antler rods of rectangular cross-section, with unknown purpose.
5. Nine flint arrowheads.
6. A beaver chisel, presumed to be an ornament or an amulet.

The presence of a rubblework, the specific skeletal position of the man on his back with extended arms, and the orientation with the head to the southeast, together with the assemblage of accompanying artifacts, lead us to attribute this burial to the spectrum of sites associated with the Serovo Neolithic culture of the Baikal region (Privalikhin et al., 2013). The date of the burial falls between 4239–4002 calBCE (5297B±25 BP) [R_combine: (5320B±40 BP, Poz-82202); (5283B±32 BP, OxA-33488)].

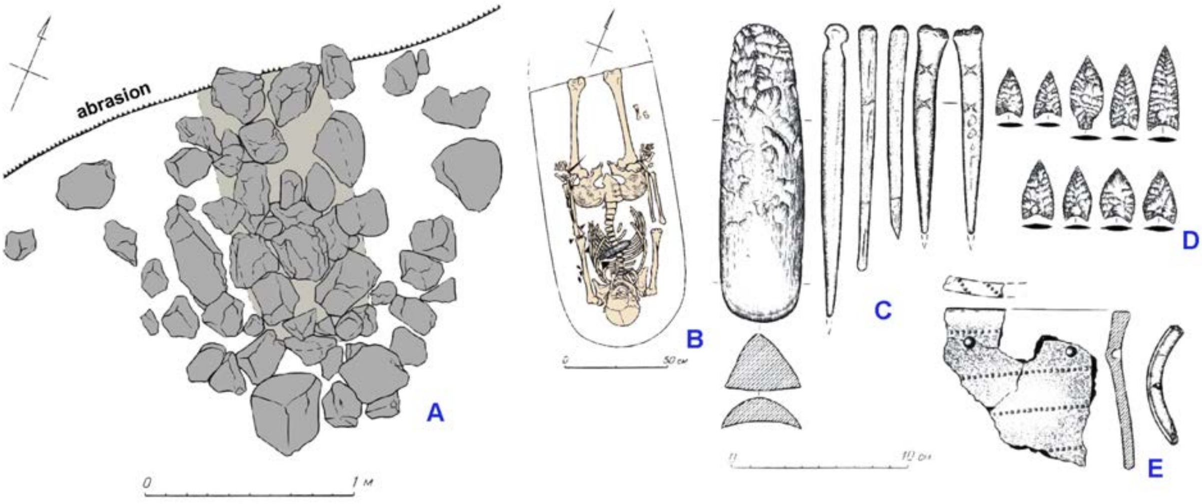

***Fig. S7.*** *Burial 9 at Sosnovy Mys.* ***A*** *- stone mound above the grave pit;* ***B*** *- burial 9 after excavation;* ***C*** *- chisel and perforators;* ***D*** *- arrowheads;* ***E*** *- pottery (Privalikhin et al., 2013, modified).*

*(b) unidentified burial (individual ID I11107)*

Another sampled individual from Sosnovy Mys-1 has individual ID I11107, but the burial number is unidentifiable. It was excavated in 1974 by N.I. Drozdov, and may come from **burials numbered from 1 to 4, or 6**.

There is no description available for burial 1 (Petrova et al., 1976).

Burial 2 consisted of a random heap of bones from a single individual, suggesting dismemberment or re-burial as part of the burial rite. Additionally, a vertebral column of another individual was discovered in the northeast corner of the grave pit. The grave goods associated with this burial included 17 pendants made from tubular bone and two points made from slate bone of a deer.

Burial 3 was unfortunately destroyed due to slope erosion, leaving no further information available.

Burial 4 was positioned within the grave pit of burial 5. The buried individual was found lying on their back with bent limbs, within a grave pit that measured 0.67 meters deep. During the clearing process, a bone needle case was uncovered beneath the pelvic bone.

Burial 6 was located directly under the rubblework boulders, without any evident grave pit. It consisted of the skeleton of one individual and the skull of another. The skeleton was positioned on its right side, with the arms bent at the elbows and lying parallel to the right side of the body, and the legs bent at the knee and hip joints. The tibia of the right leg had been cut open. The burial was double, as indicated by the presence of skull 2 next to skull 1. The buried individuals were positioned on their backs with their legs bent at the knees. Five flint arrowheads were found between the bones, and a fragment of a large flint spearhead was discovered near one of the skeletons.

Given that all the described burials exhibit similar burial rites, resembling burial 9, it is probable that individual ID I11107 can be associated with the Serovo culture of the Baikal region.

**(2) Kamenka-1 site**

The archaeological site of Kamenka-1 is situated on a limestone cape positioned between the channels of the Angara and Kamenka rivers. The cliff rises approximately 21 meters above the water level and extends for over 150 meters in a west-to-east direction, displaying an approximatelytriangular shape on the ground. The upper platform of the cape is relatively flat, covered with turf, and slopes downward towards the west, widening to about 100 meters. In 1997, A.L. Zaika uncovered a burial on the eastern edge of the cape.

*(a) single burial (individual ID I10900)*

At a depth of 25 centimeters, stonework consisting of medium-sized rock fragments was discovered. The rubblework exhibited an elongated shape, with its long axis aligned along a northeast-southwest line. Below the second layer, a well-preserved complete skeleton of an adult male, estimated to be 25-30 years old and 166 centimeters tall, was revealed.

Reconstruction of the funeral rite suggests that following the burial activities and placement of grave goods, the body of the deceased was initially positioned along the sides of the grave and subsequently placed on top, covered by large, flattened rock slabs. The stones were predominantly arranged in a vertical manner. The material used was likely comprised of large rock fragments that were exposed and uncovered during excavation. After the placement of the larger stones, smaller boulders were used to line the grave. Soil was carefully selected and poured over the masonry, forming a tomb mound characterized by dark-gray humus sandy loam. At a later stage, a funeral fire, possibly a ceremonial component, was laid in the northern part of the grave mound, specifically at the center of the mound's edge.

The skeleton was found in a fully anatomical order, lying in an extended position on its back. The orientation of the burial placed the individual's head towards the northeast (downstream the Kamenka River) and the feet towards the southwest, downstream he river bend. The burial location is in proximity to hangars located in this section of the channel. The head of the buried person rested on a stone "cushion" formed by an inclined sandstone slab of non-local origin. Due to subsidence (vertical downward displacement of stones) of the masonry, the skull had turned to the left. Similarly, the middle part of the spinal column exhibited slight curvature and shifted towards the left side. The right arm extended alongside the body, with the humerus positioned within a recess formed by the rocks that delineated the grave pit. The phalanges of the hand were scattered near the right tibia. The left arm of the buried person was slightly bent at the elbow and placed on the belt. The bones of the hand and the phalanges of the fingers were randomly arranged in the pelvic region. The legs were parallel, extended, and brought together at the level of the feet. The bones of the feet were in a mixed state and were situated within a recess formed by the rocks.

Various grave goods were discovered in association with the burial. These include a polished knife with a concave blade (item 1), 32 fragments of horn used for bow lining found along the left side of the body, three stone arrowheads to the left of the right ulna (item 4), a retouched stone knife above the right pelvic bone and a polished adze below the left pelvic bone (items 2 and 3), a massive bone arrowhead near the northern wall of the grave pit at the belt level (item 5), two fragments of a bone puncture among the phalanges of the left hand in the pelvic area, three additional bone punctures located between the femurs (items 8-10), a bone harpoon positioned with its point downward near the buried individual's left thigh, and two overlaid bone tool blanks (items 11 and 12). Fragments of a broken earthen vessel were found near the northern wall of the grave at the knee level (item 13), and an accumulation of flakes and chips (40 pieces) along with bone objects including two awls, two rod blanks, and a prismatic core were discovered in the lower region of the right leg (item 14). Lastly, a horn imitation of a bear's claw or a boar's tusk was found next to the lower part of the right leg (item 15).

The burial is attributed to the Serovo culture and dated to the period earlier than 2750±120 calBCE, based on a date from charcoal from a campfire which overlaid the burial (4700±120 BP, COAH-3780) (Zaika, 2009).

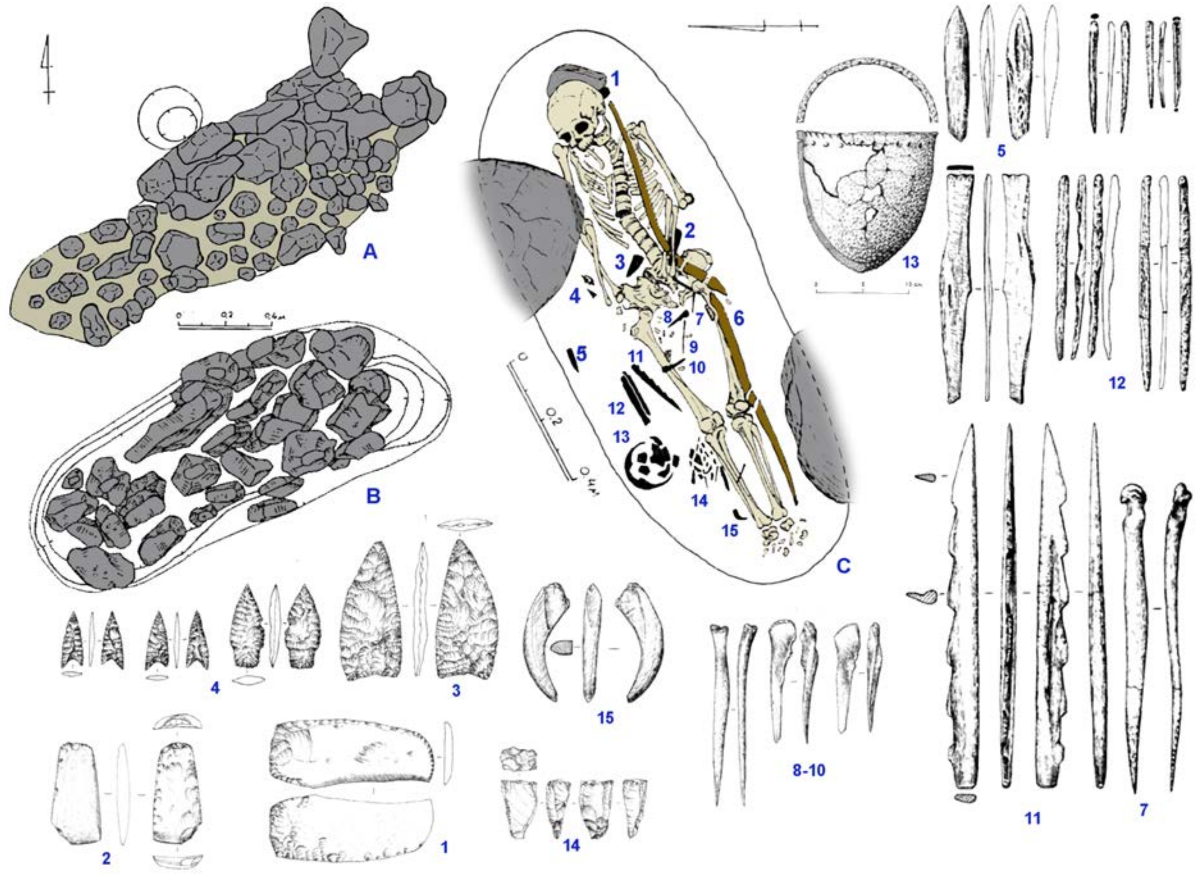

***Fig. S8.*** *Burial at the Kamenka-1 site.* ***A*** *- rubblework, upper level;* ***B*** *- rubblework, lower level;* ***C*** *- burial:* ***1*** *- polished knife;* ***2*** *- chisel;* ***3*** *- retouched bifacial knife;* ***4*** *- stone arrowheads;* ***5*** *- bone arrowhead;* ***6*** *- antler bow plates;* ***7-10*** *- bone perforators;* ***11*** *- harpoon;* ***12*** *- bone tool blank;* ***13*** *- pottery vessel;* ***14*** *- accumulation of flakes, chips, cores, and bone awls;* ***15*** *- antler pendant (Zaika, 2009, modified).*

#### II.B.ii.c) Glazkovo Early Bronze Age culture (Russia_MiddleAngara_Glazkovo_EBA)

##### (1) Sergushkin Ostrov site complex

Sergushkin Island (Sergushkin Ostrov), situated along the Angara River, spans 12 km and is densely covered with forest. The burial ground is located within a multi-layered site, discovered by V.I. Privalikhin in 1974, in the central part of the island on its southwestern coast, approximately 2.5 km above the lower tip of the island. A total of seven burials were unearthed and examined within the area. Three burials (1, 4, 7) can be dated back to the early Iron Age, corresponding to the existence of the Tsepan archaeological culture. Three burials (2, 3, 5) are attributed to the Bronze Age, while one burial (6), characterized by a cremation rite, dates back to the 13th century CE (Privalikhin, 2009).

###### *(a) Sergushkin-3, burial 2 (individual ID I5280)*

**Burial 2** (Individual ID I5280) was identified by the presence of grave masonry stones protruding from beneath the turf on the gentle slope of the terrace, which gradually descends towards the river on the northwestern side of the island. A cluster of 20 masonry stones formed a small irregular oval shape measuring 114x93 cm. At a depth of 25 cm below the surface level, beneath a large masonry stone situated on the western side, the crushed skull of a male individual, aged 30-35 years, was discovered. The burial position was on the left side, with the right temporal side of the skull exhibiting fractures caused by a blow from above with a stone. The lower portion of the stone corresponded precisely in shape and size to the dent on the skull. The skeleton of the individual was found in a crouched position on the left side, with the knees bent and the arms pressed against the chest. The hands were positioned near the eye sockets and lower jaw. The general orientation of the skeleton was with the back facing the river and the head pointing downstream. The femurs, tibias, humerus, and radius bones were fractured. Additionally, a bone needle with a deliberately broken eye and sting, measuring 6.9x0.2 cm and up to 1.5 cm across, was found at the knee joint of the right leg. This burial is attributed to the Glazkovo culture of the early Bronze Age (Privalikhin, 2009).

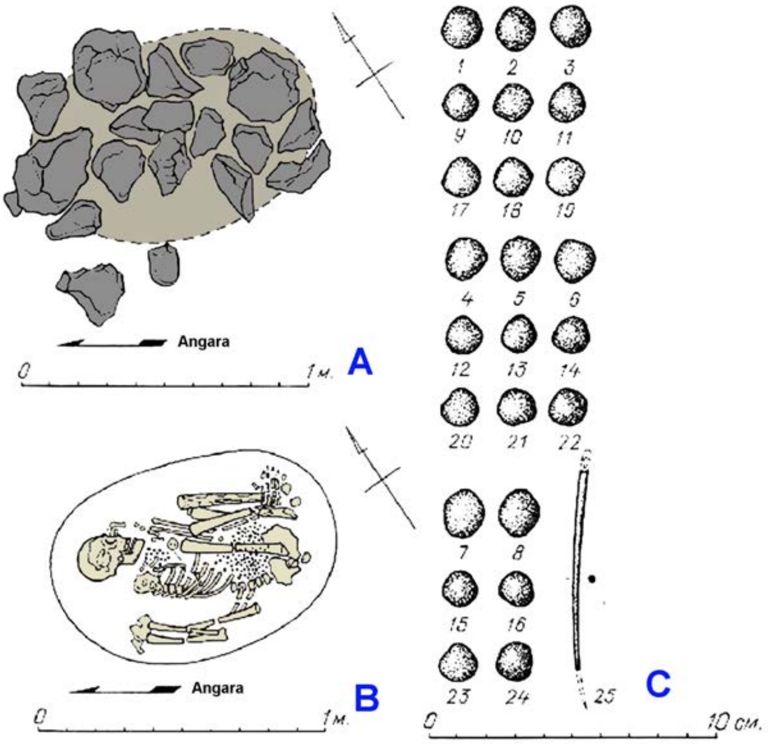

***Fig. S9.*** *Burial 2 at the Sergushkin-3 site.* ***A*** *- rubblework;* ***B*** *- burial;* ***C*** *- grave goods (Privalikhin, 2009, modifued).*

*(b) Sergushkin-3, burial 5 (individual ID I20310)*

**Burial 5** (Individual ID I20310) was uncovered at a depth of 45 cm below the modern surface level, following the removal of the Early Iron Age layer. The burial was covered by 54 stones and slabs forming grave masonry. It had an elongated oval shape, oriented along a southeast-northwest line, measuring 200 cm in length and 155 cm in width. At a depth of 110 cm from the contemporary soil surface, within an oval grave pit, the skeleton of a female individual aged 55-60 years was discovered. The skeleton was positioned on the left side, in a crouched posture, with the arms and legs tightly bent at the elbow and knee joints, pressed against the chest. The orientation of the skeleton was with the back facing the river and the head aligned with its course, i.e., to the northwest. A rich array of accompanying items was found within the grave, including an earthenware vessel with handles, a needle case made of bird bone, a bone awl-piercer, two bone needles, three discs of calcite resembling marble, a ring made of white jade, thirteen large beads of calcite resembling marble, a roe deer bone, two flakes of siliceous rock, and fish vertebrae. This burial is attributed to the Glazkovo culture of the early Bronze Age (Privalikhin, 2009).

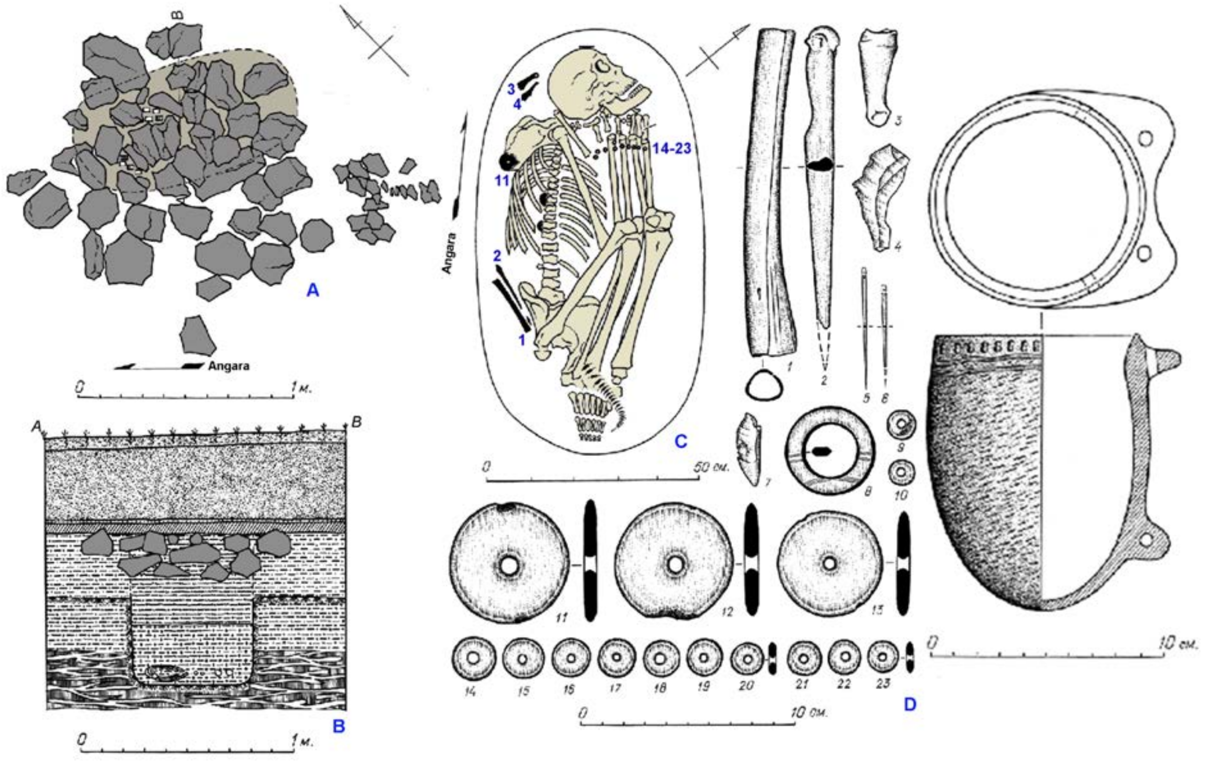

**Fig. .** Burial 5 at the Sergushkin-3 site. ***A*** *- rubblework;* ***B*** *- stratigraphic profile of the burial;* ***C*** *- plan of the burial;* ***D*** *- grave goods (Privalikhin, 2009, modified).*

##### (2) Ust'-Shamanka-1 site

*(a) single burial (individual ID I2143)*

The site was excavated in 1978 by V.N. Sokolov, who unearthed six burials, compactly placed in rows. Archaeological artifacts found in graves demonstrate similarity to the Early Bronze Age sites of the Angara region, such as Sosnovy-Mys and Sergushkin-3 (Dudarek, Lokhov 2014). Only one individual **(individual ID I2143)** was sequenced. Unfortunately, the excavators did not label the anthropological material from the site and we failed to identify the burial number to provide detailed observation of the archaeological context. However, the combination of two radiocarbon dates show chronological range of 2841-2494 calBCE (4071±21 BP) [R_Combine: (4040±25 BP, PSUAMS-9085); (4130±35 BP, Poz-82203)]. The chronological attribution of the burial as well as presence of the Glazkovo-attributed graves at the site makes the Glazkovo attribution of the studied material the best guess.

##### (3) Sosnovy-Mys site complex

See above.

*(a) Sosnovy Mys-2, burial 1 (individual IDs I20308 and I20309)*

**Burial 1:** The construction of the burial consisted of an oval elongated rubblework oriented along the east-west axis. The outer edges of the construction comprised the largest and heaviest boulders and slabs, while the central part was filled with smaller stones.

At a depth of 65 cm from the soil surface and 26 cm from the lower level of the construction, human bones (individual ID I20309) were discovered in a rounded grave pit measuring 88 x 80 cm and located 60 cm deep. The skull, along with the bones of the arms, legs, pelvis, vertebrae, and ribs, were neatly arranged at the bottom of the grave pit, without exhibiting signs of proper anatomical articulation. It is presumed that initially, the corpse of a 20-25-year-old individual remained exposed to the open air for a certain period. After the decay of muscle ligaments and tissues, the bones, together with accompanying grave goods, were collected and buried in the ground.

Burial 1, with its grave pit, cut through and disturbed another earlier burial, likely dating back to the Late Neolithic period. Within the bottom of the disturbed Neolithic burial's grave pit, located on the southeast side of burial 1, fragmented bones of a lower leg (individual ID I20308) and calcaneal bone were found.

Within the fill of burial 1's grave pit, which contained small charcoal fragments, various grave goods were discovered near the skull, between the skeleton's bones, and underneath them. These grave goods included ceramics, stone and bone tools, as well as stone ornaments and animal tooth aplliques (Privalikhin, 1998).

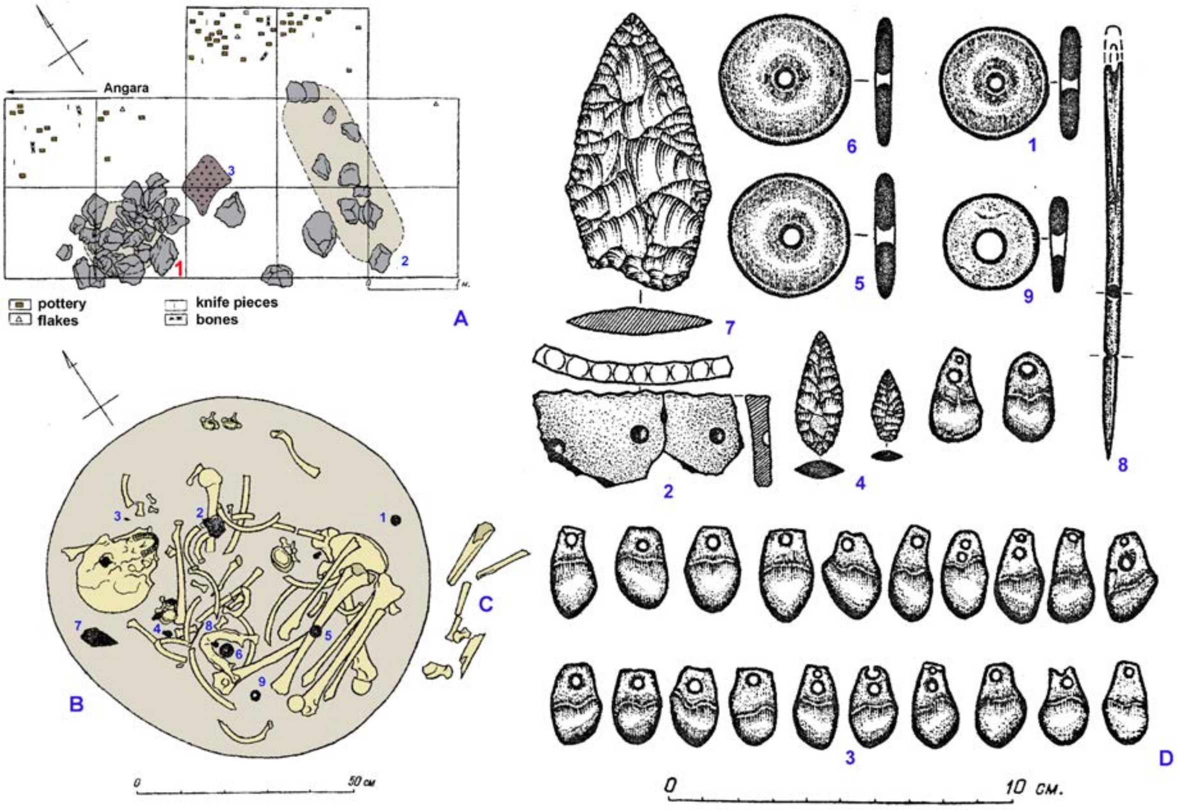

***Fig. S10.*** Burial 1 at the Sosnovy-Mys-2 site. ***A*** *- excavation plan,* ***1*** *- rubblework over burial 1,* ***2*** *- burial 2,* ***3*** *- fireplace;* ***B*** *- burial* ***1****, skeleton attributed to Glazkovo culture (I20309);* ***C*** *- human bones (I20308) from the destroyed burial (possibly Late Neolithic?) to the southeast of the Glazkovo grave pit;* ***D*** *- grave goods:* ***1, 5, 6, 9*** *- perforated calcite rings,* ***2*** *- pottery,* ***3*** *- deer teeth appliques,* ***4*** *- arrowheads,* ***7*** *- flint spearhead,* ***8*** *- bone needle (Privalikhin, 1998, modified).*

### II.C. South part of West Siberia Altai-Sayan

#### II.C.i. Yenisei River Basin

##### *II.C.i.a)* Yenisei Forest-Steppe Neolithic (Russia_Krasnoyarsk_N and Russia_UpperYenisei_N)

###### (1) Afontova Gora Neolithic burials

The Neolithic burial ground is situated in the vicinity of a Paleolithic site Afontova Gora-2.

The multilayer Paleolithic site was initially discovered by I.T. Savenkov in 1884. In 1924, N.K. Auerbach, G.P. Sosnovsky, and V.I. Gromov conducted excavations and investigated the first burial, which was possibly attributed to the Neolithic. Another nearby burial was unearthed in 1932 by A.F. Katkov and V.G. Kartsov from the Krasnoyarsk Museum. In 1937, A.P. Okladnikov and a museum employee V.I. Neshumaev excavated another burial located 3 meters away. Additionally, in 1977, schoolchildren transferred artifacts from a destroyed burial at Afontova Gora to the museum.

All the burials, comprising a minimum of 5 individuals, are positioned on the edge of the Mount Afontova slope, approximately 20 meters from the Yudin museum-estate. The estimated area of the Neolithic burial ground on Mount Afontova is between 200 and 300 square meters. The burials themselves are compactly located within a few meters' distance from each other. The skeletons are oriented perpendicular to the Yenisei River but parallel to the ancient terrace of the Kacha River.

Among the significant grave goods found at the site are bone pendants depicting ducks, needles and needle cases, awls adorned with anthropomorphic images, harpoons, mother-of-pearl disc-shaped beads, stone leaf-shaped arrowheads, and clay round-bottomed vessels with surface ornamentation.

###### *(a) Neolithic burial C (individual ID I11102)*

In 1932, A.F. Katkov excavated a **Neolithic burial at Afontova Gora** (Individual ID I11102). The grave was situated on a small hill that terminates in cliffs on both the southwestern and southern sides. No discernible traces of the grave pit or burial structures were found. The burial was located at a depth of 40-85 cm, likely within a hole oriented from south to north, inferred from the alignment of the bone remains (skull, pelvis, thigh). The skeleton exhibited a slight reddish discoloration, possibly due to the presence of ocher.

The grave goods recovered from the burial include pear-shaped pendant decorations, animal teeth fashioned in the form of ducks, and flat iridescent disc-shaped beads.

From the 1932 excavation conducted by A.F. Katkov on Mount Afontova, a braincase belonging to an individual aged 30-35 has been preserved. The craniological features of the braincase suggest a combination of Caucasoid and Mongoloid traits. According to A.F. Katkov, the burial can be dated to the early stage of the Paleometallic Age and is contemporaneous with the burial at Bazaikha, as revealed by excavations led by S.M. Sergeev (Okladnikov, 1949, 1957; Vdovin et al., 2016, Savenkova and Makarov, 2018; Berdnikov et al., 2022).

Radiocarbon dating conducted on the human skeleton recovered during A.F. Katkov's 1932 excavation yielded a calibrated date between 4673–4461 calBCE (5720±25 BP, PSUAMS-7666).

##### (2) Tolsty Mys-1

The Tolsty Mys-1 site is situated on the first floodplain terrace, approximately 8 meters above the right bank of the river. This location comprises Neolithic cultural layers. In 2010, excavations were conducted in a 300-square-meter area within the settlement.

###### *(a) single burial (individual ID I11102)*

During the excavation of the Neolithic horizon, **a single burial** (individual ID I11102) was documented. Within the base of the grave pit, the skeletal remains of a young man aged approximately 20-25 years (age estimation by Tatyana Savenkova) were found. The burial was performed in an extended supine position, with the head oriented in the downstream direction. The preservation of the bones is poor, and the temporal region of the skull exhibits damage. A patch of pink ocher was discovered in the vicinity of the skull, suggesting its ceremonial use.

In the occipital section of the burial, the fragmented remains of a ceramic vessel displaying imprints of a "braided net" design on its exterior surface were found. Some fragments from this vessel were recovered from the fill material within the grave pit. It remains unclear whether the vessel is directly associated with the burial or relates to a separate settlement layer. (Grevtsov et al., 2010; Grevtsov, 2010). The burial is AMS-dated to 5757–5636 calBCE (6825±30 BP, PSUAMS-7730).

##### (3) Krasnoyarsk finds

###### *(a) Krasnoyarsk find (individual ID I11101)*

The burial was initially discovered in June 1955 by a geologist, A.P. Balykov, on the left bank of the Yenisei River, near the summer dachas of the employees of the City Department of Public Education (GORONO) of Krasnoyarsk. It was situated on the first floodplain terrace, approximately 8 meters above the river's edge, directly opposite Sosnovy Island. In the same year, the excavation was conducted by Z.K. Glusskaya. The museum collection was analyzed by an anthropologist V.P. Alekseev in 1960. A reconstruction of the skull found within the burial was undertaken by M.M. Gerasimov and was subsequently reexamined by V.A. Dremov and T.M. Savenkova. The archaeological artifacts were studied by N.P. Makarov, I.M. Berdnikov, and A.S. Vdovin.

The **single burial** (individual ID I11101) contained the remains of a woman estimated to be 30-35 years old. The body was interred in a supine position, with the orientation across the river, the head facing north, and the feet pointing south. Grave goods recovered from the burial included ornaments comprising flat, mother-of-pearl rounded beads, pendants crafted from animal teeth, plates made of boar tusks, and a needle case with threaded ends (Glusskaya, 1963; Vdovin and Makarov, 2016; Savenkova and Makarov, 2018; Alekseev, 1960; Solodovnikov et al., 2020; Dryomov, 1988; Kozintsev, 1974). The burial is dated to the period between 4793–4612 calBCE (5845±30 BP, PSUAMS-7549).

###### *(b) Ulitsa Karla Marksa (Karl Marx Street, Krasnoyarsk) (individual ID I11103)*

During the archaeological monitoring of earthworks for the reconstruction of the heat collector in Karl Marx Street in Krasnoyarsk in July 2000, several significant archaeological complexes were discovered by A.Yu. Tarasov.

One of these discoveries occurred on the southern side of the collector trench, approximately 21 meters north of the northern facade of a residential building at Karl Marx Street 21. At a depth of 1.64 meters from the present-day surface, an area stained with ocher was identified. Further excavation revealed that the postcranial human skeleton was situated in soil saturated with ocher. Unfortunately, it was not feasible to fully expose the trench side due to a trading pavilion located above the burial. As a result, anthropological materials were extracted by carefully excavating the trench wall.

The excavated cranium originates from a **single burial** (individual ID I11103), devoid of any accompanying grave goods. Radiocarbon dating places this burial within the 95% confidence interval of 5962–5735 calBCE (6945±30 BP, PSUAMS-7550).

*(c) Bazaikha burial ground, unnumbered burial (individual ID I11104)*

The Bazaikha burial ground, situated in the "Bor" area on the right bank of the Yenisei River near the mouth of the Bazaikha River, is currently located within the city limits of Krasnoyarsk. It was initially discovered by I.T. Savenkov in 1883. During the excavations carried out from 1883 to 1885, Savenkov revealed several burials, tentatively attributed to the Neolithic period, with one burial containing a celt axe, suggesting a Late Bronze Age date. In 1920, S.M. Sergeev excavated another burial, which is currently under study by T.M. Savenkova, N.P. Makarov, and A.S. Vdovin using materials from the museum collection.

Based on the available data, the Bazaikha burial ground encompassed at least seven burials, with six excavated by I.T. Savenkov and one by S.M. Sergeev.

One of the **burials in Bazaikha** contained the skeletal remains of a woman (individual ID I11104) in a supine position with her head oriented to the west. Grave goods discovered in the burial included flat, rounded mother-of-pearl beads, pendants crafted from animal teeth, and boar tusk plates (Vdovin and Makarov, 2016; Savenkova and Makarov, 2018; Alekseev, 1961; Debets, 1948; Dus, 1923). Radiocarbon dating places the burial within the 95% confidence interval of 5471–5231 calBCE (6385 ± 25 BP, PSUAMS-7674).

#### II.C.i.b) Yenisei Forest-Steppe Eneolithic/EBA (Russia_UpperYenisei_EBA)

##### (1) Dolgoye Ozero burial ground

The Dolgoe Ozero (Dolgoe Lake) burial ground is situated in the vicinity of Kansk, near a dune in the northwestern section of the quarry adjacent to the Dolgoe Lake. It was initially discovered by G.A. Maksimenkov in 1958 during a reconnaissance survey.

At a depth of 70 cm below the soil surface, a human humerus protruded from the ground on the cliff face. Below it, within the screened area, a skull was found, accompanied by a puncture mark from an elk's slate bone. Located 50 cm to the west were two lower jaws, upper limb bones, and fragments of skulls. Further clearing of the quarry cliff at a depth of 55-60 cm revealed three damaged grave pits, consisting of two single burials and one double burial.

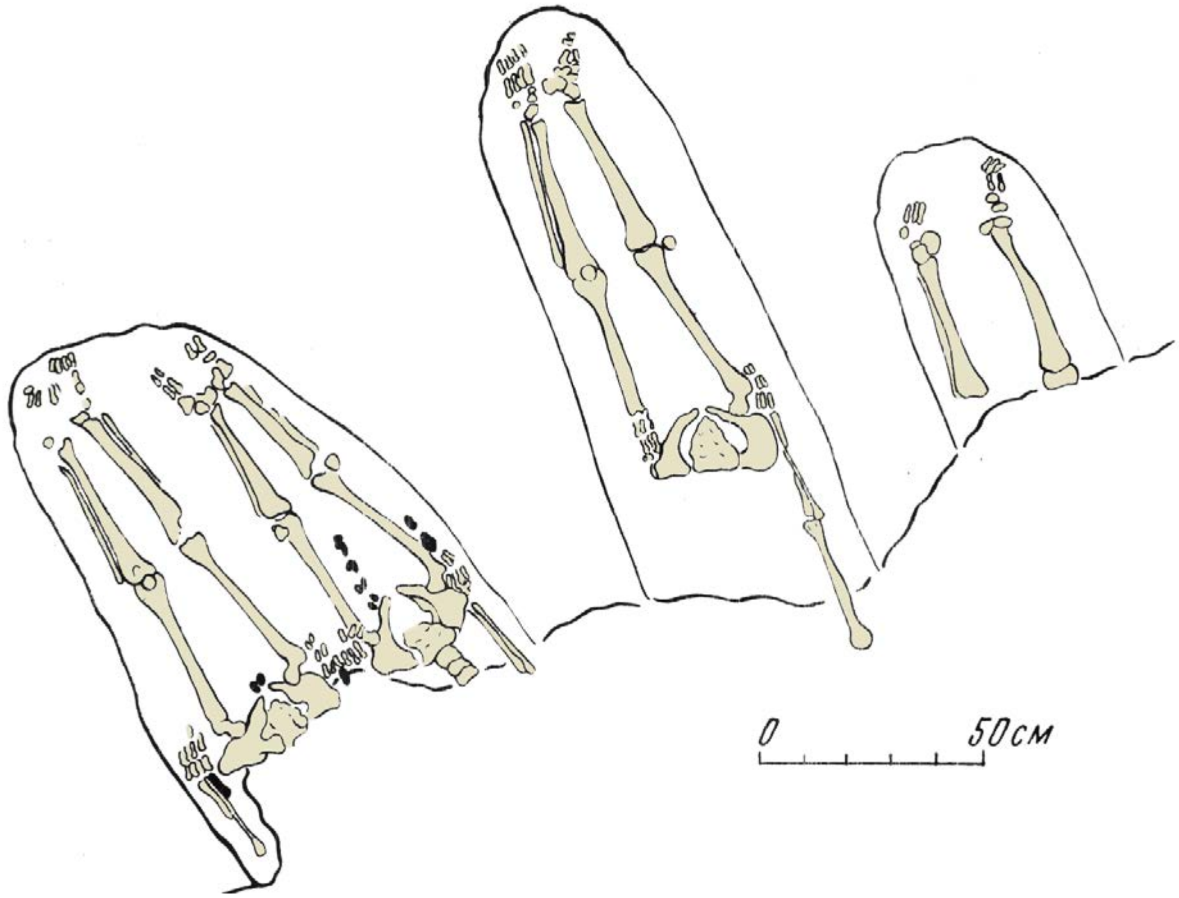

***Fig. S11.*** *Partially preserved burials at the Dolgoye Ozero site (Maksimenkov, 1964, modified).*

*(a) unidentified burial (individual ID I1522)*

The preserved parts of the grave pits measured up to 140x50 cm. Within them, four disarticulated skeletons were discovered, positioned supine with their heads oriented towards the southeast and their feet perpendicular to the oxbow lake channel, with their heads pointing northward. Grave goods recovered from these burials included three perforators crafted from elk's slate bone, pendants made from deer teeth, flint flakes, and a fragment of ocher. The skulls were found outside burials. M.M. Gerasimova conducted an examination of the skulls, revealing the following results: skull 1 belonged to a woman aged 25-35 years; skull 2 belonged to a male aged 20-25 years; skull 3 belonged to a female aged 30-35 years; and skull 4 belonged to a male aged 40-45 years (Maksimenkov, 1964; Gerasimova, 1964; Timoschenko, 2013).

The scarcity of the records makes it infeasible to determine which of the skulls and burials the individual ID I1522 pertains to. The burial is radiocarbon-dated to 2879–2586 calBCE (4150±35 BP, Poz-82193).

#### II.C.ii. The Upper Ob and Kuznetsk Depression

In 2019, a project for the preparation and publication of the "History of Altai" was implemented, in which one of the volumes reflected the results of the study of the prehistoric epoch and presented materials from archaeological sites attributed to the Neolithic period within four stages of its development (early, developed, late, and final). However, no emphasis was placed on their correlation with previously identified cultures (History of Altai, 2019, pp. 52–95). This circumstance is related to the fact that none of the previously presented concepts can be accepted as a comprehensive model for summarizing the accumulated information. There are more than enough reasons for such a situation. For example, all culture names are given based on regional designations, which do not correspond to the archaeological principles of studying sources and only demonstrate a geographical factor, hindering a genuine understanding of the specific characteristics of the accumulated materials (as a result, there are currently two chronologically distinct Upper Ob cultures in West Siberian archaeology). However, the main problem lies in the limited extent of interdisciplinary research on the still relatively small array of available data, which were obtained from dispersedly identified sites of different categories (burial grounds, settlements) and chance finds. For instance, only in the past decade have AMS dating methods begun to accumulate, allowing for a more precise determination of the chronological range of specific archaeological objects. Other important findings, obtained through the use of high-tech equipment, including paleogenetic analyses, which are only partially reflected in this article, will become significant as well. The correlation between the materials from burial grounds and settlement complexes will be of particular importance. Given the briefly mentioned information, substantial prospects for further scientific research work are evident.

An overview of the process of studying Neolithic sites and the identification of cultures in the territory of the southern part of West Siberia and Altai is reflected in the recently published volume "History of Siberia," which examines two regions: Upper Ob and Altai-Sayan Plateau (History of Siberia, 2022, pp. 216–218). The beginning of a systematic understanding can be attributed to M.N. Komarova, who in 1956 identified the Kiprino and Irba stages in the development of the Neolithic in the forest-steppe region of Upper Ob. The first generalization was made by V.I. Matyushenko, who in 1969 distinguished and characterized the Upper Ob Neolithic culture. The position of M.A. Anikovich is noted, who in the same year and on the same territory "…defined a group of sites associated with the Baikal Neolithic" (History of Siberia, 2022, p. 217). In the 1970s, V.I. Molodin refined the existing periodization, identifying the Zavyalovo and Kiprino stages in the development of the Upper Ob culture and pointing out an intercultural group of sites. In 1978, A.P. Okladnikov and V.I. Molodin identified the Kuznetsk-Altai culture of the Neolithic period based on materials from individual burial grounds and the study of inscriptions. This perspective was critiqued in the second half of the 1980s by Yu.F. Kiryushin, who attributed some sites to his previously identified Bolshoy Mys culture of the Eneolithic period, primarily based on the study of the ceramic complex.

By the end of the 20th century, the number of archaeological materials attributed to the Neolithic period significantly increased. This was particularly true for the steppe zone of the left bank of Upper Ob, where more than 150 settlement sites with characteristic finds and individual burials are known. This allowed A.L. Kungurov, A.V. Onnikov, and A.A. Tishkin to identify the Rubtsovsk Neolithic culture based on the eponymous site, which yielded a significant volume of artifacts, the study of which continues to the present. The rationale for this culture was reflected in A.V. Schmidt's dissertation. The Neolithic sites of the forest-steppe zone on the right bank of Upper Ob demonstrate their own specific features, and the main burial grounds (Firsovo-11 and 14, Tuzovskie Bugry, Chumysh-Perekat, etc.) are considered in this article. Importantly, they show similarities with the left bank sites. This circumstance allows us to consider them all as part of a single cultural and historical entity, within which individual cultures with their distinctive stages should be considered.

Planned and multi-year excavations in the territory of the Mountainous Altai in its northern part have allowed for the study of well-stratified settlement complexes and the identification of the early, developed, late, and final stages of the Neolithic period. Within the framework of the late stage, Yu.F. Kiryushin and K.Yu. Kiryushin proposed the differentiation of the Middle Katun culture, although the researched settlements and sites are actually located in the lower reaches of the Katun River.

Another historiographical aspect highlighted is the Ural-West Siberian Neolithic community (History of Siberia, 2022, p. 218), which has not seen further development.

In the mentioned volume "History of Siberia," characteristics of the Upper Ob Neolithic culture, Kuznetsk-Altai culture, and Middle Katun culture are presented (History of Siberia, 2022, pp. 260–268, figures 155–159). These cultures possess their own local peculiarities, but they require more detailed examination at a contemporary level, as previous hypotheses have already run their course. One example of this is the rejection of dating the petroglyphs to the Neolithic period, attributed to the Kuznetsk-Altai culture. It is necessary to abandon all previously proposed cultural designations that do not adhere to the methodological principles of archaeological research and have exhausted their potential. This is evidenced, in particular, by new insights into ethnocultural processes in the Baraba and Middle Irtysh regions, where several archaeological cultures of the Neolithic period have been identified (History of Siberia, 2022, pp. 248–259).

This review and the new scientific information, including paleogenetic data, indicate the need for the study of Neolithic materials in the south of West Siberia and Altai at a contemporary research level, with a purposeful search for other archaeological sites and excavations of foundational monuments.

#### II.C.ii.a) Kuznetsk-Altai Neolithic / Upper Ob Neolithic / Bolshoy Mys Eneolithic sites (Russia_KuznetskAltai)

##### (1) Razdum'ye-1 site

Razdum'ye-1 is a Medieval hillfort located on an elevated promontory on the right bank of the Ob River. The site was extensively investigated and excavated by Alexei Umansky between 1960 and 1966. The archaeological exploration revealed a complex and multi-phased assemblage spanning from the Neolithic to the late Middle Ages. In 1965, two Neolithic/Eneolithic burials were excavated at the site (Umansky, 1987).

###### *(a) Early Neolithic burial 1(6) (individual ID I12695)*

**Burial 6** (individual ID I12695), designated firstly as burial 1, and this code is still in use in the Tomsk anthropological collection, but the burial is recorded as grave 6 in a publication by Umansky in 1987) was excavated in 1965. The grave pit had an oval shape with dimensions of approximately 1.2x0.75 meters. Its depth reached 1.3 meters from the present-day surface. The southwestern part of the grave had collapsed into the river. The deceased individual, identified as a male aged 50-60 years, was found in a supine position with the head oriented to the north. The arms were extended, while the leg bones were missing. The bones exhibited slight traces of ochre staining. Small fragments of charcoal and sheep's astragalus were discovered in the grave fill. Traces of wood ashes indicated the presence of grave siding and a vertical post positioned behind the skull of the deceased. No grave goods were present in the burial (Umanskiy 1987). It is dated to 6571–6424 calBCE (7635B±35 BP, PSUAMS-9117).

##### (2) Zarechnoye-1 site

Zarechnoye-1 consists of a group of burial mounds excavated by Viktor Zakh in 1980 and 1983. The site is located on a small promontory on the high left terrace of the Inya River. The burial ground contains approximately 80 mounds, grouped into two clusters. Excavations covered an area of approximately 900 square meters.

The excavations at Zarechnoye-1 revealed burials belonging to the Irmen culture beneath the mounds. Additionally, an Andronovo flat burial ground, a Samus settlement complex, and five burials dating back to the Neolithic and Early Bronze Ages were discovered. Two Neolithic burials were found beneath mound 2, which was constructed during the Irmen period (Zakh, 1985).

*(a)	mound 2, Early Neolithic burial 3 (individual ID I2138)*

**Burial 3 of mound 2** (individual ID I2138) The burial was partially overlapped by a later grave attributed to the Irmen culture. The boundaries of the grave pit were not traced. The skeleton was placed in a vertical, standing position, fronting east. Its upper part was damaged by the later grave. The skull was located 0.4 m to the southeast from the remaining bones of the skeleton and was sprinkled with ocher. The buried person was a woman about 25 years old. The right side of the chest was abundantly covered with ocher. During excavation, in the infill of the grave on the left side of the skeleton, two microblades and a stone axe made from a slate flake were found. Near the right pelvic bone, a fragment of a bird bone polished on both sides and a bone tool made from split bone were recorded (Zakh, 1985). The burial has been dated to 7035-6651 calBCE (7920±50 BP, Poz-82211).

*(b)	mound 2, Early Neolithic burial 4 (individual ID I2139)*

**Burial 4 of mound 2** (individual ID I2139) was located at a distance of 1 to 3 meters from burial 3. The grave pit was not clearly identified, as it was partially covered by an Irmen culture burial. The buried individual, a male estimated to be 25-30 years old, was found in a supine position with the head oriented to the north, parallel to the edge of the terrace. A piece of animal bone was discovered on the right side of the chest (Zakh, 1985). The burial has been dated to 7035–6651 calBCE (7920±45 BP, OxA-33165).

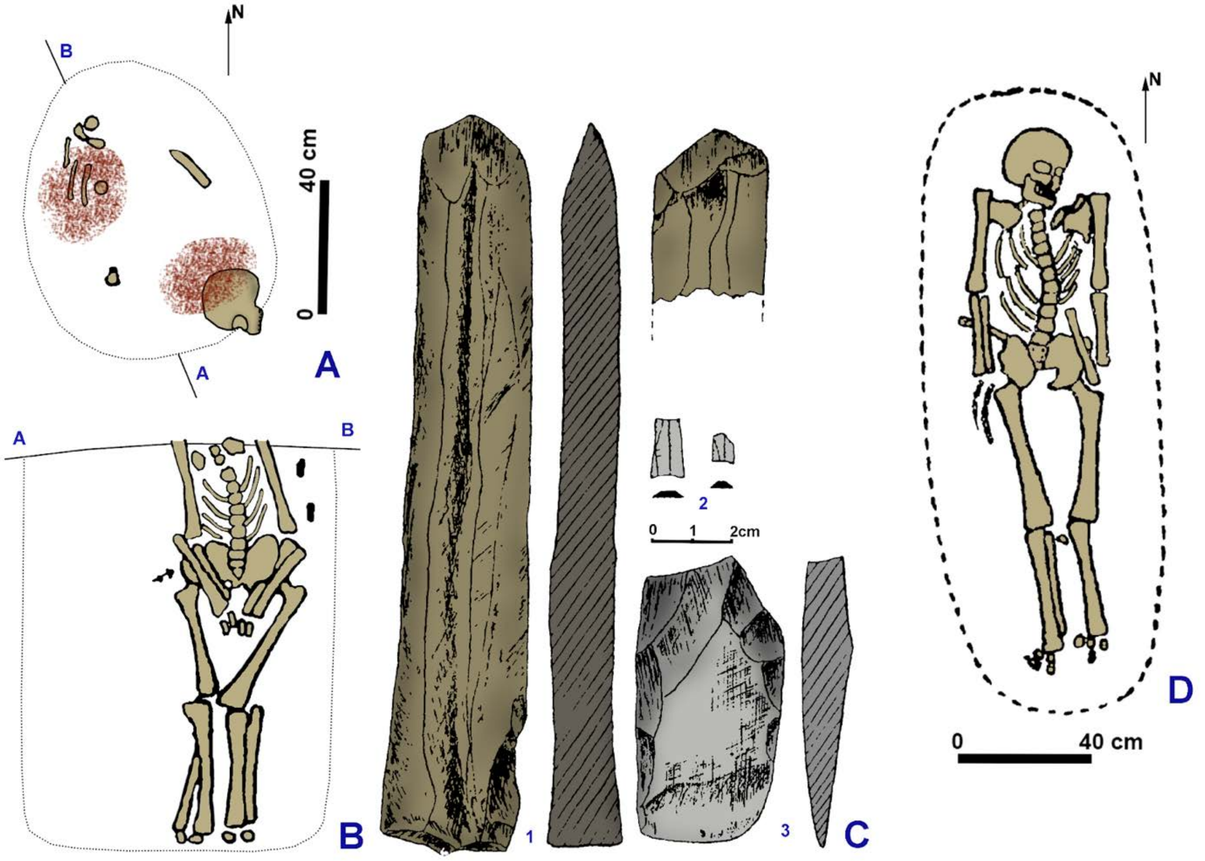

***Fig. S12.** Early Neolithic burials at the Zarechnoye-1 site (Zakh, 1985, modified). **A**: mound 2, burial 3, plan; **B**: mound 2, burial 4, section; **C**: artifacts from burial 3, **1** - bone tool, **2** - blades, **3** - stone axe; **D**: mound 2, burial 4, plan.*

##### (3) Firsovo-11 site

The Firsovo-11 cemetery is situated on the right bank of the River Ob, opposite the city of Barnaul. AT this location, the right bank of the Ob features a wide, swampy floodplain measuring up to 7 km, adorned with numerous oxbow lakes and remnants of the original coastline in the form of crests and mounds. The surrounding vegetation consists of meadows with thickets of shrubby willows, aspens, and birches. During periods of flooding, the water level in the Ob rises by 1.5–3 m, submerging the floodplain almost entirely and approaching the bedrock bank upon which the monument is situated.

The cemetery was initially investigated in 1977 by A.L. Kungurov and V.B. Borodaev, and later excavated by Ya.V. Frolov in 1993–1994. The monument spans an area of 0.3 hectares. Eight burials were discovered at the site, including five individual graves, two paired graves, and one collective burial. The burials are arranged in two rows, oriented along a southeast-northwest axis. The deceased individuals within the graves were found to be oriented with their heads pointing northward or northeastward. Grave depths varied from 0.4 to 1.7 m.

Thirteen AMS dates were obtained, ten for human bones and teeth, and three for animal remains. The results indicate the presence of a reservoir effect, with an average offset of approximately 700 years. Some of the dates have been published (Kiryushin et al. 2021a, Kiryushin et al. 2021b). The age of the Neolithic burials corresponds to the middle of the 5th millennium BC. The "Neolithic core" of the site comprises burials 9, 14, 15, 16, 17, 41, and 42, with burial 18 belonging to an earlier period.

Noteworthy artifacts found at the site include ornaments crafted from animal teeth (bear, wolf, horse, etc.). These tooth ornaments serve as chronological indicators, offering the potential for specialized dating methods to investigate the reservoir effect further (Kiryushin et al., 2021a, 2021b).

The Neolithic burials at the Firsovo-11 site form a cohesive cultural and chronological group together with graves 1 and 13 of the Itkul (Bolshoi Mys) cemetery. With time, this group may expand to include burials 2, 4, 6, 10, and 14 of the Itkul (Bolshoi Mys) cemetery.

The designation of the Neolithic burials at the Firsovo-11 site as belonging to the Bolshoi Mys culture of the Eneolithic period (Kiryushin, 2002) requires thorough reassessment. The other names used to label the Neolithic cultures in the region include Upper Ob and Kuznetsk-Altai Neolithic cultures.

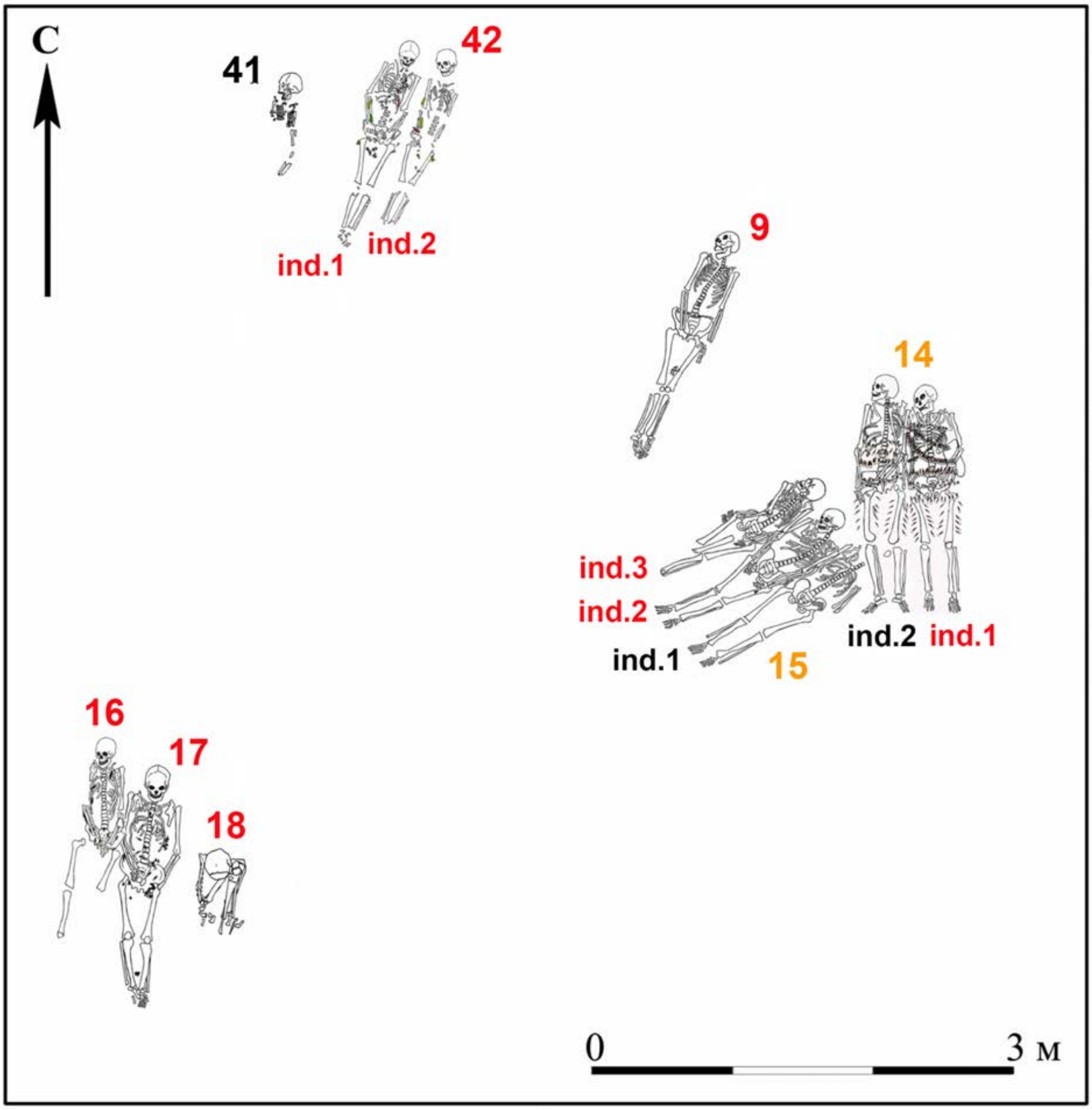

***Fig. S13.*** *Neolithic burials at Firsovo-11. Involved individuals are labeled in red.*

*(a) burial 9 (individual ID I0996)*

**Burial 9** (individual ID I0996) was unearthed in 1993 during an excavation of the lower section of an Early Iron Age burial mound. It was located 2 meters southwest of burial 42 and 1-1.5 meters northwest of burials 14 and 15, situated in the central part of the "northeastern row." The condition of the grave stain was indistinct and difficult to decipher. The grave had a depth of 1.7 meters from the contemporary ground surface and exhibited an elongated shape, aligned along the south-southwest-north-northeast axis. The deceased individual, identified as a male aged 40-50 years, was found in a supine position with the head oriented towards the north-northeast.

The burial was accompanied by grave goods, which included nine triangular stone arrowheads with either a straight or notched base. Additionally, there were 22 bone patches discovered: 21 of them exhibited elongated outlines with subrectangular or oval shapes and a hole in the upper part, while one patch had a round shape with a hole in the center. It should be noted that the skull showed signs of deformation due to the pressure exerted by the overlying soil (Kiryushin et al., 2021a, 2021b).

A combination of two radiocarbon dates from the burial shows interval 5479-5377 calBCE (6469±18 BP) [R_Combine: (6500±24 BP, MAMS-48706); (6435±25 (PSUAMS-4214)].

*(b) burial 14 individuals 1 (ID I13096) and 2 (I0997)*

**Burial 14** was uncovered in 1993, positioned 1.5 meters southeast of burial 9 and partially intersecting with burial 15 in the eastern section of the "northeastern" row. The state of the grave stain was indecipherable and offered no readable information. Within the burial, located at a depth of 0.8 meters from the current ground surface, the remains of two individuals were discovered, interred "shoulder to shoulder."

***Individual 1*** (individual ID I13096) (a male, aged 20-30 years) was found in a supine position with the head oriented towards the north. The skull exhibited deformation resulting from the pressure exerted by the soil. An analysis of a human ulna from skeleton 1 provided a date of 6330-5913 calBCE (7222±82 BP, GV-02887).

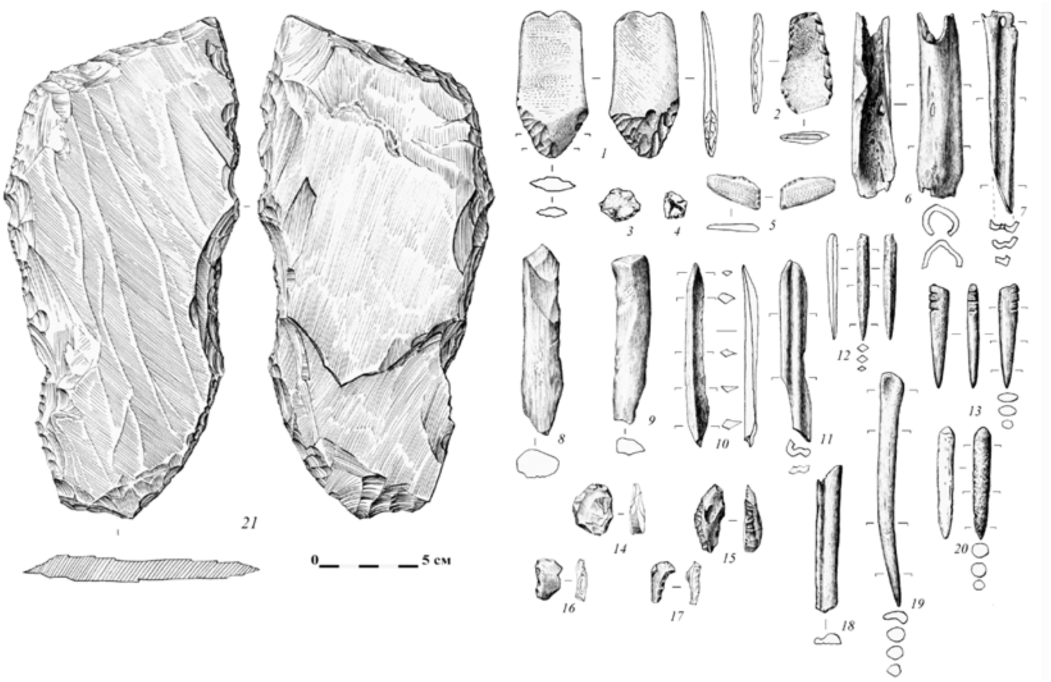

***Fig. S14.*** *Grave goods from burial 14 at Firsovo-11.* ***1-13*** *- skeleton 1; 1****4-20*** *- skeleton 2;* ***21*** - burial *14.* ***1*** *- polished knife;* ***2*** *- polished axe;* ***3, 4*** *- retouched flakes;* ***5*** *- fragment of a polished knife;* ***6*** *- fragment of a tubular bone;* ***7-9, 11*** *- tool blanks;* ***10, 12, 13, 19, 20*** *- blades;* ***14, 15*** *- scrapers;* ***16*** *- retouched flake;* ***17*** *- serrated and notched tool;* ***18*** *- fragment of a bone tool.*

***Individual 2*** (individual ID I0997) (presumably a young woman, aged 18-20 years) was also lying extended on her back with the head facing north. Similar to the previous individual, the skull displayed deformation due to soil pressure.

Burial 14 represents the most affluent interment within this necropolis. The accompanying funerary inventory comprises a multitude of bone and stone artifacts. Notably, the skeletons were adorned with necklaces composed of animal teeth that had been drilled at the roots and organized in clusters (Kiryushin et al., 2021a; 2021b).

Radiocarbon dating was conducted on a fragment of an elk tooth from skeleton 1's necklace, resulting in a date of 4708-4547calBCE (5778±28 BP, GV-03572). A date from human bones shows much earlier chronology, 5617-5479 calBCE (6595±30 BP, PSUAMS-4119), most likely due to freshwater reservoir effect. Due to observed δ13C/δ13N ration, we expect freshwater reservoir offset in the date obtained from human bones and prefer to use the date from the elk tooth.

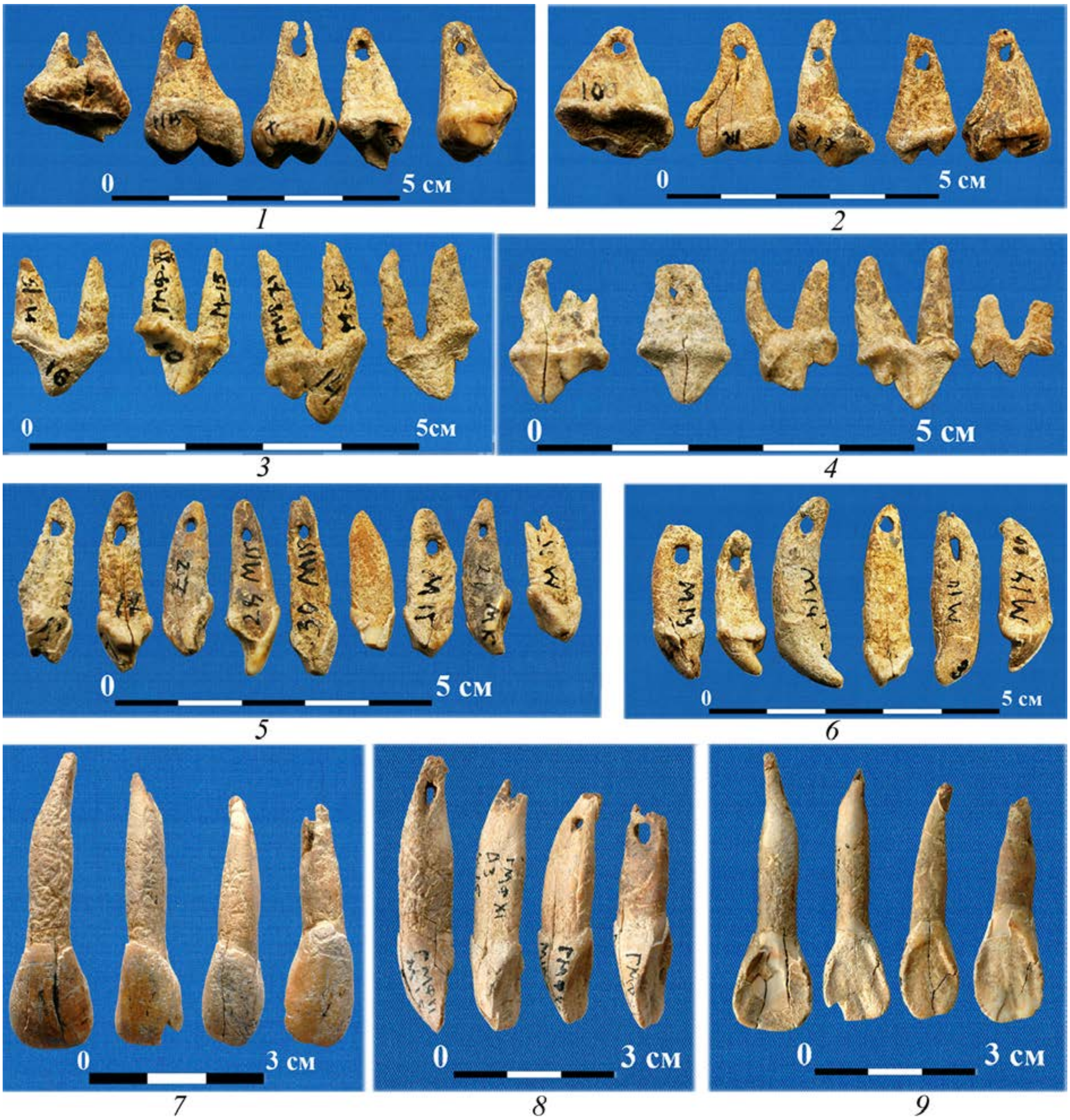

*Fig. S15. Ornaments and costume details from burial 14 at Firsovo-11.*

*(c) Burial 15, individuals 2 (ID I11744) and 3 (ID I10963)*

**Burial 15,** situated at the southeastern end of the "northeastern" row, has experienced partial disturbance due to burial 14. The grave pit was not recorded. This burial represents a collective interment of three elderly males, positioned in an extended supine position, placed adjacent to each other ("shoulder to shoulder"), with their heads oriented toward the east-northeast. The bottom of the grave was located at a depth of 0.87 meters from the contemporary surface.

***Individual 2*** <u>(individual ID I11744)</u> (a male, aged 45-55 years) was laid out fully extended on his back, with his head facing the east-northeast. The skull has suffered deformation due to the weight of the soil.

***Individual 3*** (individual ID I10963) (an elderly male, aged 55-65 years) was also laid out fully extended on his back, with his head facing the east-northeast. The skull shows deformation caused by the pressure of the surrounding earth. The bone condition is deteriorated.

A triangular stone arrowhead with a notched base was discovered on the left femur of skeleton 2 (Kiryushin et al., 2021a; 2021b) (Kiryushin et al., 2021a, 2021b).

Individual 2 has been dated to 5711-5558 calBCE (6705±30 BP, PSUAMS-8933), freshwater reservoir effect is expected. In another paper, skeleton 1 was dated to 5702-5485 calBCE (6684±39 BP, UBA-22954) and skeleton 3 to 5734-5521 calBCE (6723±68 BP, GV-02888) (Kiryushin et al., 2021c). The combination of dates shows interval 5665-5561 calBCE (6700±23 BP) [R_Combine: ind. 1 (6684±39 BP, UBA-22954); ind. 2 (6705±30 BP, PSUAMS-8933); ind. 3 (6723±68 BP, GV-02888).

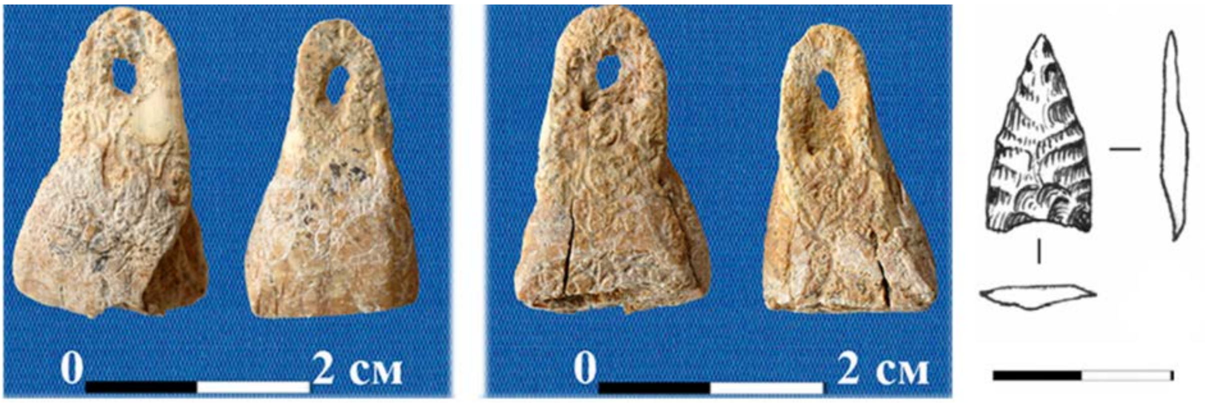

***Fig. S16.*** *Ornaments from a horse chisel burial 17 and an arrowhead from burial 15 at Firsovo-11.*

*(d) burial 16 (individual ID I10962)*

**Burial 16** (individual ID I20962) is situated at the far northwestern end of the second row. The actual grave pit was not identified or delineated. During the excavation of the area, the bones were discovered at a depth of 0.4 meters from the contemporary surface. The deceased individual was interred at the boundary between the humus layer and the underlying subsoil sand.

The deceased, a male estimated to be 30-40 years old, was laid out in a fully extended supine position, with his head facing northward, slightly inclined towards the east. The skull exhibits deformation due to significant deterioration and preservation issues. The bone condition is also poor.

Accompanying the burial are several grave goods: three arrowheads with leaf-like shapes (resembling double-edged points), a bone point, a stone knife resting on a silicified slate slab, and eight pendant teeth.

Based on an analysis of a fragment of human bone, a date was determined: 5478-5372 calBCE (6459±28 BP, GV-03573).

*(e) burial 17 (individual ID I11741)*

**Burial 17** (individual ID I11741) was positioned within the second row, situated between burials 16 and 18. Burial 16 partially ovarlaid burial 17. The precise outline of the grave pit has not been determined. The excavation reached the bottom of the burial at a depth of 75 centimeters from the contemporary surface. The skeletal remains of a woman, estimated to be between 45 and 55 years old, were discovered within the burial. The deceased was laid out in a fully extended supine position, with her head facing northward, slightly deviating towards the east. The skull exhibits deformation due to poor preservation, and the bone material is in a deteriorated condition.

Accompanying grave goods include 13 stone arrowheads with either straight or notched bases, as well as 34 pendants made from animal teeth, each featuring a drilled hole in the root.

A date of 5478-7361 calBCE (6448±28 BP, GV-03575) was obtained through analysis of a fragment of human bone. Additionally, a date of 4782-4553 calBCE (5818±28 BP, GV-03574) was obtained from a fragment of a bear's tooth.

*(f) burial 18 (individual ID I11742)*

**Burial 18** (individual ID I11742) occupies the extreme southeast position within the second row, although the precise outline of the grave pit has not been identified. The interred male individual, estimated to be between 35 and 45 years old, was placed in a seated position within the grave. The skull was discovered at a depth of 0.5 meters from the modern surface and exhibits deformation resulting from the pressure of the surrounding soil. The excavation reached a depth of 1 meter from the modern surface, revealing a burial richly adorned with ocher. The sole funerary item present in the burial is a knife-shaped piece.

The cranial dimensions of the interred individual, differing from those of others buried at the site, further underscore the chronological and ritual distinctiveness of this burial.

Based on fragments of the radius bone, a date of 8561-8016 calBCE (9106±80 BP, GV-02889) was obtained.

As a working hypothesis, it is proposed that the very old date for burial 18 at Firsovo-11 is not an artefact, suggesting that this burial indeed belongs to the final Mesolithic or early Neolithic period. Burial 18 at Firsovo-11 represents one of the few early Holocene burials where the deceased was interred in a seated position (Kiryushin et al., 2021a, 2021b).

*(g) burial 42, individuals 1 (ID I12225) and 2 (ID I12224)*

**Burial 42** was discovered in the northern part of the first row, located 2 meters northwest of burial 9 and 0.5 meters east of burial 41. The exact boundaries of the grave pit have not been determined. The depth of the grave is 0.7 meters from the modern surface. Within the burial, the remains of two individuals were found.

***Individual 1*** (Individual ID I12225), identified as a man between 45 and 55 years old, was laid out in a supine position with his head oriented to the north-northeast. The skull exhibits deformation caused by natural processes and the pressure exerted by the surrounding soil. The bone preservation is in poor condition. The burial assemblage includes stone and bone artifacts, as well as clothing adornments made from animal teeth that were drilled into the root (Kiryushin et al., 2021a, 2021b). A date of 5623-5363 calBCE (6534±72 BP, GV-02890) was obtained from a fragment of human bone.

***Individual 2*** (Individual ID I12224), a woman aged 40 to 50 years, was also laid out in a supine position with her head oriented to the north-northeast. Similar to the previous individual, the skull shows signs of deformation resulting from natural processes and soil pressure. The bone preservation is also in poor condition. The accompanying grave goods consist of eleven stone arrowheads with a notched base, two small polished stone axes, a split beaver chisel, and a fragment of a bone artifact (Kiryushin et al., 2021a, 2021b). A date of 5609-2390 calBCE (6404±28 BP, GV-03576) was obtained from a fragment of human bone. Additionally, a date of 4685-4501 calBCE (5745±28 BP (GV-03577) was obtained from a fragment of a bone artifact.

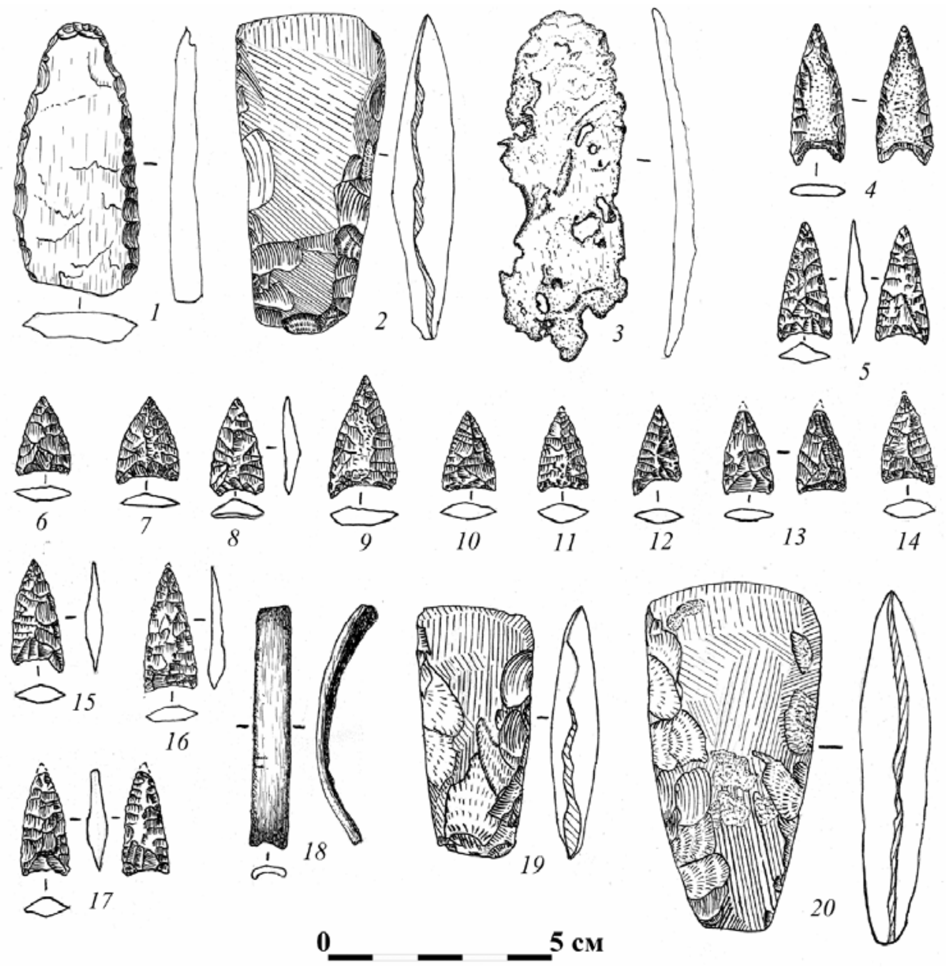

***Fig. S17.*** *Grave goods from burial 42 at Firsovo-11.* ***1-9*** *- skeleton 1;* ***10-20*** *- skeleton 2.* ***1*** *- knife on a silicified shale;* ***2, 19, 20*** *- hatchets;* ***3*** *- bone slab;* ***4-17*** *- an arrowhead;* ***18*** *- a bracelet made of beaver incisors.*

##### (4) Chumysh-Perekat-1 burial site

The Chumysh-Perekat burial ground, discovered in 2012 by S.P. Grushin, was extensively studied between 2014 and 2019 by a joint expedition from Kemerovo State University and Altai State University, led by A.V. Fribus and S.P. Grushin.

Chumysh-Perekat is situated on a prominent cape along the right bank of the Chumysh River, a right tributary of the Ob River. This area in the Western Salair region marks the boundary between the forest-steppe and mountain taiga landscape zones. An area of 1,718 square meters was investigated within the monument, revealing a total of 36 burials, of which 12 belong to the Neolithic period.

The Neolithic burials form distinct rows extending southeast from the western edge of the cape into the terrace. The Neolithic graves were characterized by inhumation. The grave pits are shallow, and their size and shape are not clearly discernible. With the exception of one grave, all burials are individual in nature. Burial 8 represents a double burial. The interred individuals were laid out in a supine position with their knees pressed against each other, and their heads oriented towards the northeast. The grave goods associated with these burials are not numerous and consist of stone and bone tools, ornamental items crafted from animal teeth, and a bone wand bearing an image of an elk's head. Notably, the artifact assemblage includes stone and bone implements, jewelry, and artistic objects, such as a fragment of a stone figurine depicting a fish of the Baikal type.

Four radiocarbon dates were obtained from human bones at the site, at laboratories in St. Petersburg and Tomsk.

Based on the available radiocarbon dates, the Chumysh-Perekat monument can be attributed to the 6th – the first half of the 5th millennium BC. The burial ground exhibits significant similarities to the Firsovo-11 site in terms of burial practices and the complex of clothing items. It also shares similarities with other sites such as Solontsy-5, Ust-Isha, and Big Cape, as indicated by the funeral rite and the assortment of cultural artifacts (Kungurova, 2000; 2005). Currently, it is not possible to discern distinct stages in the functioning of the cemetery (Fribius and Grushin. 2017a, 2017 b, 2020).

*(a) Burial 6 (individual ID I11745)*

**Burial 6** (individual ID I11745): The exact shape of the grave pit, as in other cases, could not be determined. The dimensions are estimated as follows: length 196 cm, width 78 cm, and depth approximately 35 cm from the modern surface. The bottom of the pit exhibited a slight depression from head to feet and had a southwest-northeast orientation. The burial pertains to an adult individual, whose skeleton was laid out elongated on the back, with arms alongside the body and legs slightly bent at the knees, and the head facing northeast. Burial 6 was disturbed during the construction of burial 7. It is an individual burial. The only grave item found was a bone arrow or dart tip, hexagonal in shape and measuring 10.8 cm in length. Radiocarbon dating yielded an age of 5970-5414 calBCE (6739±130 BP, IMCES-14C1005). (Fribus, Grushin, 2020).

*(b) Burial 7 (I13097)*

**Burial 7** (individual ID I13097)**:** Based on the position of the preserved bones, it appears that burial 7 partially intersected burial 6, resulting in damage to the skeleton of grave 6. The dimensions of the grave pit could only be approximated: length 103 cm, width 55 cm, and depth from the modern surface ranging from 45 to 55 cm. The orientation of the grave was northeast-southwest, and similar to burial 6, it exhibited a depression from head to feet. The bottom of burial 7 was situated 20 cm below the level of the bottom of burial 6. This grave represents an individual burial of a child. Analysis of the *in situ* bones, including the skull, some thoracic bones, and leg bones, suggests that the interred individual was laid out elongated on their back, possibly with a slight deviation to the right side, and with the head pointing northeast. No grave goods were present. The determined dated of the human bones from the burial is 6742-6509 calBCE (7810±35 BP, PSUAMS-13976).

*(c) Burial 9 (I12626)*

**Burial 9** (individual ID I12626)**:** The shape of the grave pit could not be determined. The orientation was southwest-northeast, and the depth from the subsoil level was 22 cm. It was an individual burial of a man aged 45–55 years, who was laid out stretched on his back, with the head pointing northeast. The skull exhibited a slight turn to the right. The legs were brought together at the knees, and the arms were positioned along the body. Many phalanges of the fingers and bones of the feet were missing.

*(d) Burial 12 (I13098)*

**Burial 12** (individual ID I13098)**:** The shape of the grave pit could not be determined. The orientation was southwest-northeast, and the depth from the subsoil level was 10 cm. The burial was individual and partially disturbed. The bones preserved *in situ* indicate that the buried man, approximately 20–25 years old, was laid out stretched on his back, with the head pointing northeast. The skull and lower jaw were displaced towards the legs, and the right arm was slightly bent at the elbow, possibly with the hand placed on the pelvis. The bones of the left hand, pelvic and femur bones, and foot bones were missing. The bones on the left side of the thoracic region were displaced. Radiocarbon dating of human bones provided an age of 6058-5663 calBCE (6974±110 BP, IMCES-14C1003) (Fribus, Grushin, 2020).

Alternatively, the human bones from the grave were dated to 6392-6095 calBCE (7400±35 BP, PSUAMS-9042). The combination of these two dates provides a date of 6059-5663 calBCE (6974±111 BP) [R_Combine: (6974±110 yBP, IMCES-14C1003); (7400±35 BP, PSUAMS-9042)].

*(e) Burial 13 (I10961)*

**Burial 13** (individual ID I10961): The exact shape of the grave pit could not be determined. It had a southwest-northeast orientation and a depth of 20 cm from the subsoil level. This grave represents an individual burial, where the deceased was laid out stretched on their back, with the head pointing northeast. The skull exhibited a slight turn to the right, and the bones of the legs were tightly pressed together. The arms were positioned alongside the body. Most of the bones from the feet and hands were missing. Adjacent to the skeleton, at a distance of 20 cm on both the right and left sides, three vertically positioned wood pieces were discovered, possibly indicating elements of a tomb construction. The fragments measured 1-2 cm in diameter, with surviving lengths of approximately 10-12 cm. Grave goods accompanying the burial included a stone lanceolate biface knife, a flake, a bone dagger tip, a stone pendant resembling an elk's tooth with decorative patterns, a fish-shaped pendant made of mother-of-pearl, a beaver chisel, 33 pendants made from animal teeth (possibly dog or wolf) with holes in the root sections, and a bird's metacarpal bone serving as a buckle. Based on the burial materials, a reconstruction of the costume elements was undertaken (Fribus, Grushin, 2017).

The burial was radiocarbon-dated to 4991-4342 calBCE (5770±150 BP, Le-11465).

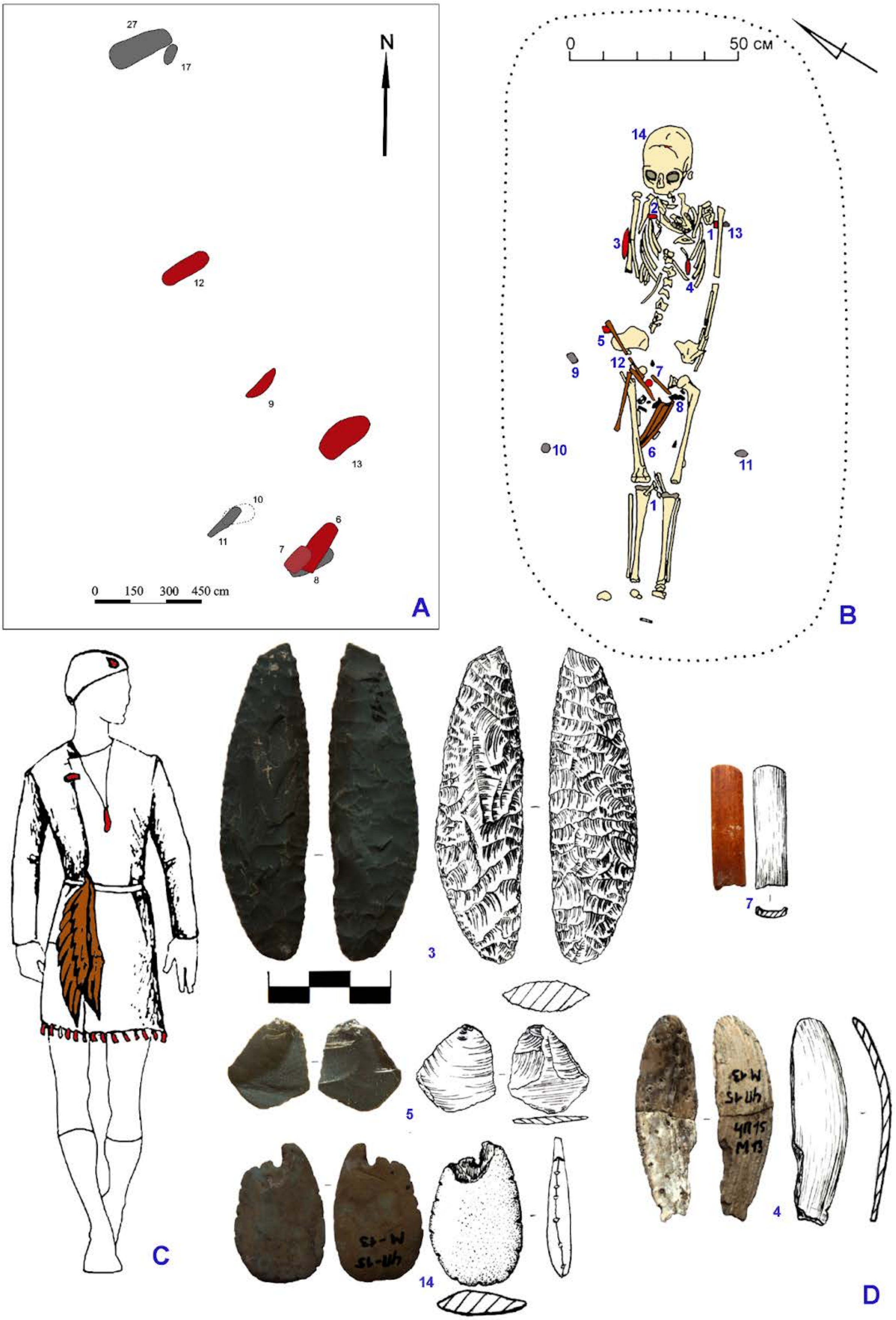

27
17
N
12
9
13
10
11
6
7
8
0 150 300 450 cm
A
0
50 см
14
2
1
13
3
4
5
9
12
7
8
10
6
11
1
B
C
3
7
5
4
14
D

*Fig.* ***S18****. Chumysh-Perekat burial site. A: plan of cemetery, sequenced burials are marked in red; B: burial 13 at Chumysh-Perekat; C: reconstruction of the costume; D: artifacts from the grave. C, D: 1 - animal tooth, 2 - beaver incisor, 3 - knife, 4 - fish-shaped pendant, 5 - flake, 6 - bone dagger tip, 12 - metacarpal bones of a bird; 7 - perforated bone, 8 - animal teeth, 9-11 - wood pieces, 13 - charcoal, 14 - stone pendant.*

##### (5) Vas'kovo-4 site

The Vaskovo-4 burial site is situated on an elevated cape-like area near the village of Vas'kovo in the Promyshlennovsky district of the Kemerovo region. It is located at the confluence of the Inya River and its small right tributary, the Tykhta River. The arrangement of the burials within the site remains unpublished. The site was accidentally discovered in 1963 due to river bank erosion and subsequently investigated in 1964 by Yu.M. Borodkin. Unfortunately, the field documentation and a significant portion of the artifact collection from the site have been lost. Partial publications of the materials exist (Borodkin 1967, 1972, 1976), which are widely known and referenced in various comprehensive studies (Molodin 1977, 1992; Anikovich 1969; Bobrov 1988, etc.). The cultural attribution of the site remains controversial (Zah 2003).

Initially, the burial ground consisted of at least three burials, one of which was found in a deteriorated state. Gender and age characteristics of the individuals are not specified. It is likely that all burials were of adult individuals. The positioning of the corpses within the graves is not detailed in the publications. However, in all burials, the individuals were oriented with their heads to the east, perpendicular to the headwaters of the small tributary, and with their feet downstream in relation to the main body of water, the Inya River. Ocher was found covering the human remains in all burials (Marochkin 2014).

###### *(a) Neolithic burial 1 (individual ID I2074) and 3 (I2075)*

According to the available publication (Borodkin 1972), no burials lacked grave goods. However, due to the fragmented nature of the available information, it is not possible to differentiate the burials based on the extent of their accompanying inventories. From the detailed description of **burial 1** (individual ID I2074), it can be inferred that the nature and quantity of grave goods varied significantly between burials.

The spatial arrangement of the grave goods within the burial can only be determined with certainty in the case of burial 1. In most instances, the objects were positioned in the pelvic area (5 objects), followed by the chest area (2). There was one recorded case of objects placed near the head and hips (Marochkin 2014).

**Burial 1** (individual ID I2074) is dated to 5610–5376 calBCE (6520B±40 BP, Poz-83514), and **burial 3** (individual ID I2075) is dated to 5551–5369 calBCE (6500±40 BP, Poz-83403).

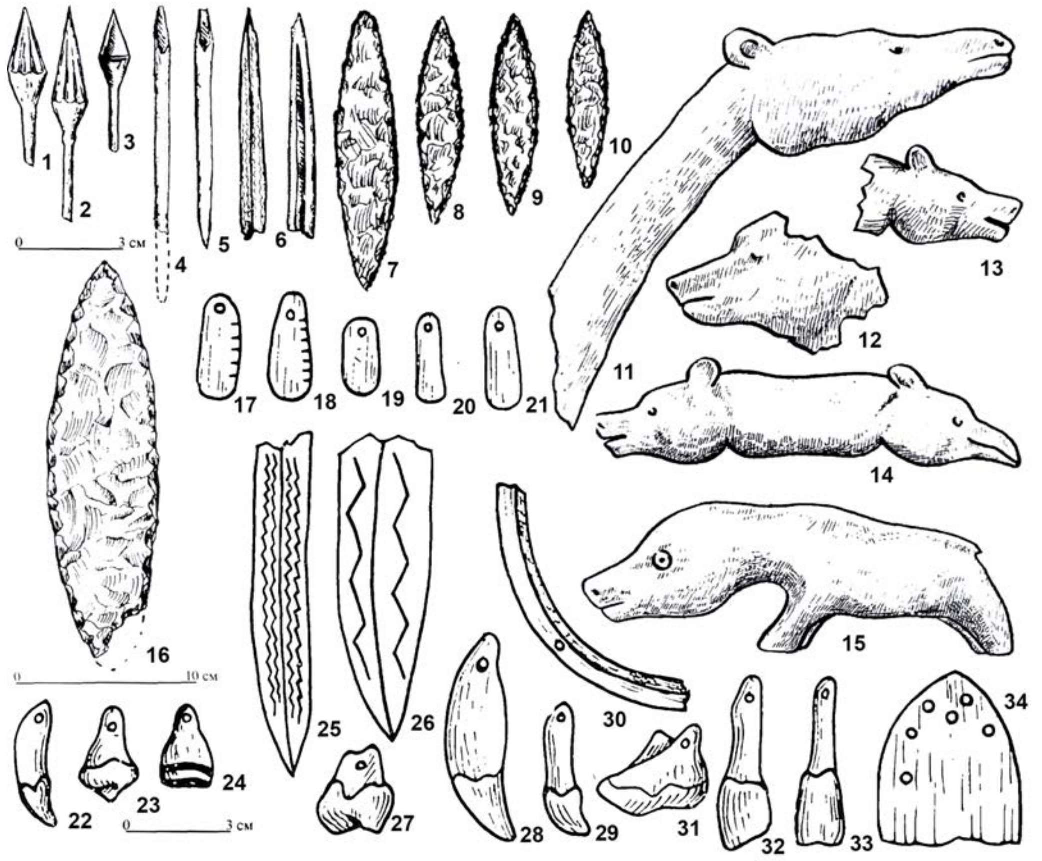

***Fig. S19.*** *Grave goods from the burials at Vas'kovo-4. Attribution of the goods to specific burials is unknown.* ***1-3, 7-10*** *- arrowheads;* ***4-5*** *- pointed rods;* ***6*** *- knife base;* ***11-15*** *- zoomorphic figurines (no scale);* ***16*** *- bifacial knife;* ***17-21*** *- plate pendants;* ***22-24, 27-29, 31-33*** *- pendants made of animal teeth;* ***25-26, 30, 34*** *- other items made of bone and antler (Marochkin 2014, Borodkin 1976).*

##### (6) Archaeological site complex at the Itkul' Lake

The complex of Neolithic archaeological sites at the Itkul' Lake includes two burial grounds, ***Bolshoy Mys*** and ***Kostenkova Izbushka***.

The ***Bolshoy Mys burial site*** is situated across from the village of Vershinino, approximately 4 km from the village of Pleshkovo. It was initially discovered and partially damaged during economic activities in 1954 by A.D. Sambursky. Subsequent investigations were carried out in 1962 by B.Kh. Kadikov and V.I. Molodin. The findings from the excavations have been fully published (Dremov, 1997; Kiryushin et al. 2000; Molodin, 1999).

The original number of burials at the site was at least 18, including two burials that were destroyed by A.D. Sambursky. The burial ground is located at the end of a promontory that extends from the northeastern shore of Lake Itkul towards its center. According to Yuri Kiryushin, during the Neolithic-Eneolithic period, the area was an island (Kiryushin, Maloletko, 1986).

The burials are arranged in three rows across the cape, following a northwest to southeast line. The northeastern row consists of burials 14-16. Burials 1-8 and 10-13 are grouped in the central row. The southwestern row includes the destroyed burials discovered earlier, as well as burials 9 and 17. The excavations have investigated 16 burials, and the preservation of the remains in all of them is poor. Burial 17 was partially disturbed by a badger's burrow. Men of various ages, ranging from 30 to 70 years old, were buried in single graves 1-3, 9, 11-13, and 15. Individual burials of women (aged 18-55) include graves 4, 14, and 16. Female burials 5-6 and 7-8 appear to be paired burials (Kiryushin, Kungurova, Kadikov, 2000). Burial 17 is also a pair, but there is no consensus regarding the age and sex of the individuals buried within it. According to V.I. Molodin, it consists of a man (about 18 years old) and a woman (about 25 years old). However, V.A. Dremov believes that both buried individuals are women, aged 40 and 25, respectively. Individual burial 10 contained the remains of a child aged 1.5-2 years old.

The characteristics of the grave pits are not described in the publications, with only mentions of visual observations regarding filling spots (Kiryushin, Kungurova, Kadikov, 2000). Based on photographic illustrations of the excavation process, it can be inferred that the human remains were placed in shallow, approximately rectangular soil pits that corresponded in size and proportions to the size of the individuals being buried.

In all cases, the skeletons were positioned on their backs, in an extended posture. Burials 1-4 and 11-13 exhibit a distinctive characteristic - the hands are placed on the pelvis, and the shins are flexed. This positioning is not consistently observed in burials 8, 10, 14, and 17. The state of preservation in other burials does not permit a definitive determination of the presence or absence of this characteristic.

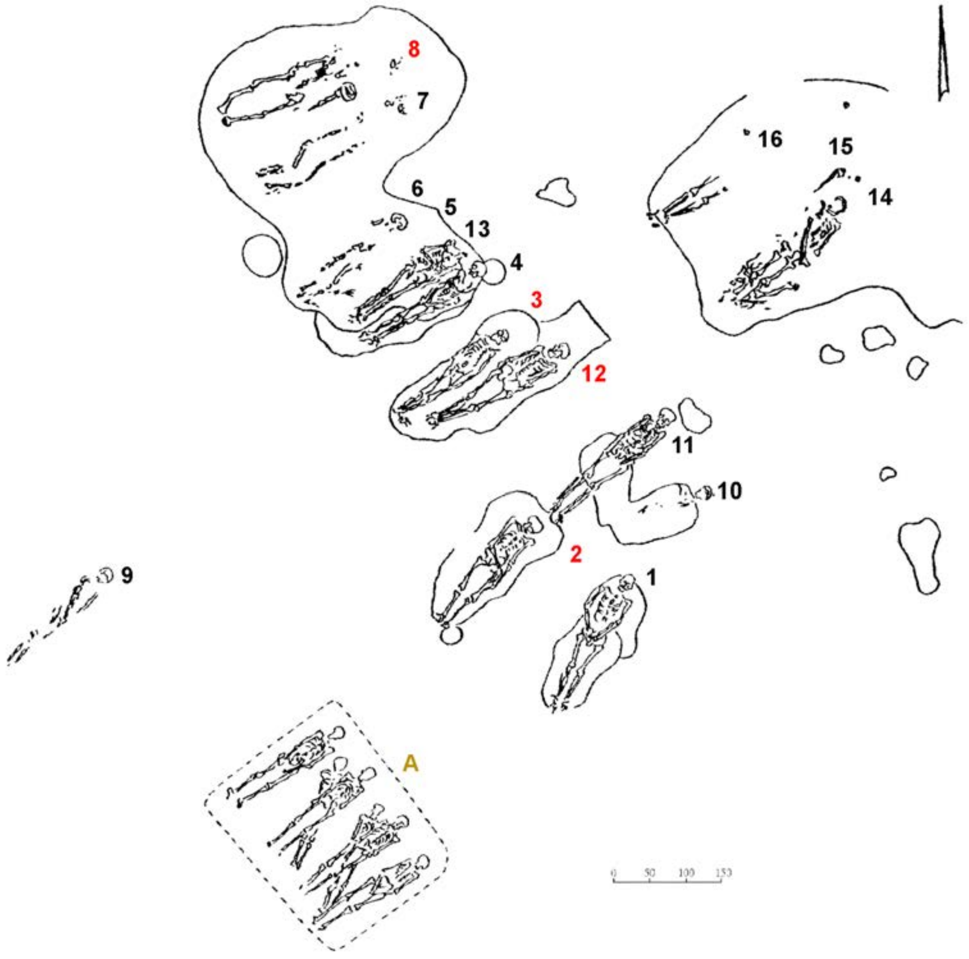

***Fig. S20.*** *Bolshoy Mys burial ground. The burials containing sequenced individuals are labeled in red.* ***A*** *- approximate position of the burial destroyed in 1954 during the construction of a dugout building (an individual from this burial was sequenced) (Kiryushin 2002).*

The ***Kostenkova Izbushka burial ground*** is located in close proximity to the village of Pleshkovo and is within direct line of sight from the Bolshoy Mys burial ground. It was discovered and studied in 1981 by Yuri Kiryushin during excavations of the Bronze Age settlement bearing the same name. The materials from the early complex have been extensively published (Kiryushin, 1983; Kiryushin, Kungurova, Kadikov, 2000). The early burials at the site are associated with the Bolshoy Mys Eneolithic culture (Kiryushin, 1986; Kiryushin, Kungurova, Kadikov, 2000). Given the current water level in Lake Itkul, the burial ground is confined to a cape-shaped ledge along the shore of Dergach Bay. Unfortunately, there is no available data regarding the planigraphy of the monument. The burial pits have not been explicitly recorded. The individuals were interred in a supine position. Burial 4 is notable for the partial disturbance of anatomical integrity, with only the long bones of the skeleton preserved. A distinctive feature of the remains from burial 5 is the positioning of the left hand on the pelvis (as the bones of the right hand have not survived), and a slightly flexed position of the legs relative to the upper part of the skeleton (Marochkin 2014).

*(a) Itkul' (Bolshoy Mys), burial 2 (individual ID I11003)*

**Burial 2** <u>(individual ID I11003)</u>: The grave had irregular oval outlines. Within it lay the skeleton of a man aged 30-35, positioned supine with his head facing northeast. His left arm extended alongside the torso, while the right arm was bent at the elbow, with the ulna bones resting on the pelvis and the hands clasped together. The skull exhibited a tilt to the left. Notably, the bones of the feet were absent. A red-brown schist flake was discovered approximately 5-6 cm deeper under the lower jaw of the buried individual. Adjacent to the upper part of the right humerus was a stone hatchet. A grinding rectangular block was found near the left fold on the interior. The majority of the jewelry was concentrated around the neck area and the region of the right pelvic bone. Hanging pendants made of large deer teeth were positioned along the shoulder line, beneath the lower jaw. On the left side of the chest, two large teeth from a bear and a fox were placed. Pendants crafted from small teeth of mustelids and artiodactyls (roe deer) were situated near the head of the right femur (3 pieces), beneath the right wing of the pelvis, on the ulna bones of the right hand, and under the sacrum. The height of the buried individual was measured at 1.65 m (Kiryushin et al. 2000).

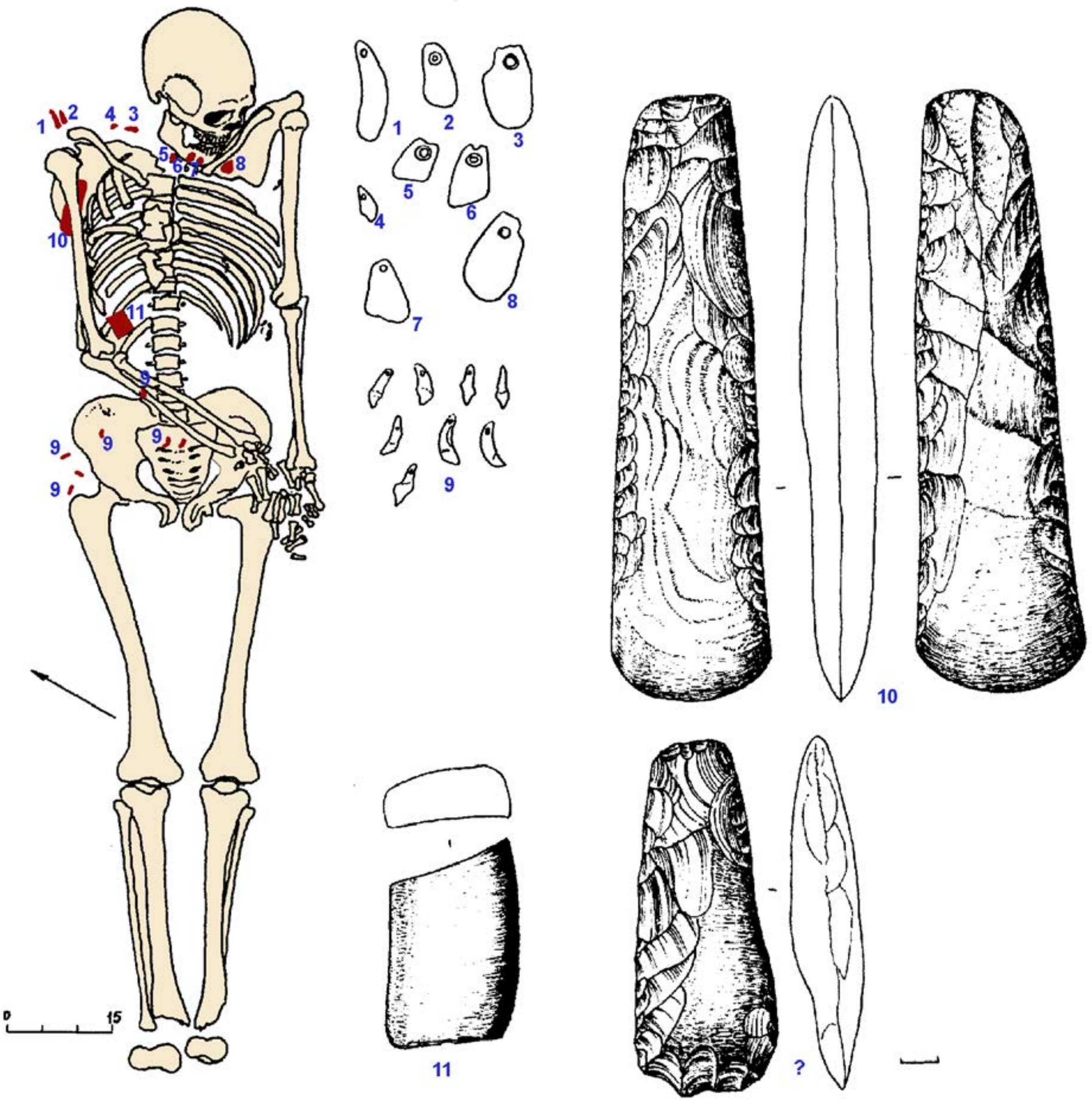

***Fig. S21.*** *Burial 2 at Itkul' (Bolshoy Mys).* ***1-11*** *- grave goods:* ***1-9*** *- pendants;* ***10*** *- stone hatchet;* ***11*** *- grinding block* (Kiryushin et al. 2000).

*(b) Itkul' (Bolshoy Mys), burial 3 (I13676)*

**Burial 3** (individual ID I13676): Located in the central row of the burial ground, this burial is paired with burial 12. The skeleton of a man aged 25-30 was found in an extended supine position with the head oriented towards the northeast. His arms were extended alongside the body, slightly bent at the elbows, with the wrists resting on the pelvic bones. Only a few phalanges of the hands and feet were preserved, and they were found in a disordered state. The right ulna bone was flexed. The clavicles, ribs, and sternum exhibited displacement, and the lower jaw was wide open. Within the burial, a small quartzite flake was discovered between the humerus and the scapula. In the area of the sternum, incisors of a rodent (beaver) were found, including a whole tooth and a fragment, as well as a badger's tooth. The total length of the skeleton was measured at 1.65 m (Kiryushin et al. 2000).

*(c)    Itkul' (Bolshoy Mys), burial 8 (I1964)*

**Burial 12** (individual ID I12874): Positioned in the central row and forming a pair with burial 3, this burial had a rectangular shape in the northeastern portion of the grave pit. Based on the fillings of the graves, it is evident that burial 3 was made subsequent to burial 12. The skeleton within burial 12 was found in anatomical order and belonged to a man aged 60-70. He was buried in an extended supine position with the head facing northeast. The ribs on the left side displayed slight displacement, and the phalanges of the arms and legs were in a disordered state. The hands were slightly bent at the elbows, directed with the wrists towards the hip and pelvic joints. Pieces of red stone were found on the outer side of the right pelvic bone (Kiryushin et al. 2000).

*(d)    Itkul' (Bolshoy Mys), burial 12 (I12874)*

**Burial 8** (individual ID I1964): This burial is paired with burial 7, as the bones of the individuals in both burials were located at a distance of 0.25 m from each other and at a depth of 0.85 m. The skeleton of a woman aged 25-35 was found in anatomical order, lying stretched out on her back with her head facing east, and her arms placed alongside her body. The right arm was slightly bent at the elbow. The skull was crushed and displaced towards the left shoulder, while the lower jaw remained in an anatomical position. The skeleton exhibited the absence of many small and thin bones, such as ribs, joints, and phalanges. In proximity to the right knee, a stone rod of a fishing hook was discovered at a distance of 0.50 m. Jewelry items including incisors of a beaver or marmot (13 pieces), a deer tooth pendant, and a roe deer tooth were found in the neck region and to the left of the vertebra on the chest (Kiryushin et al. 2000).

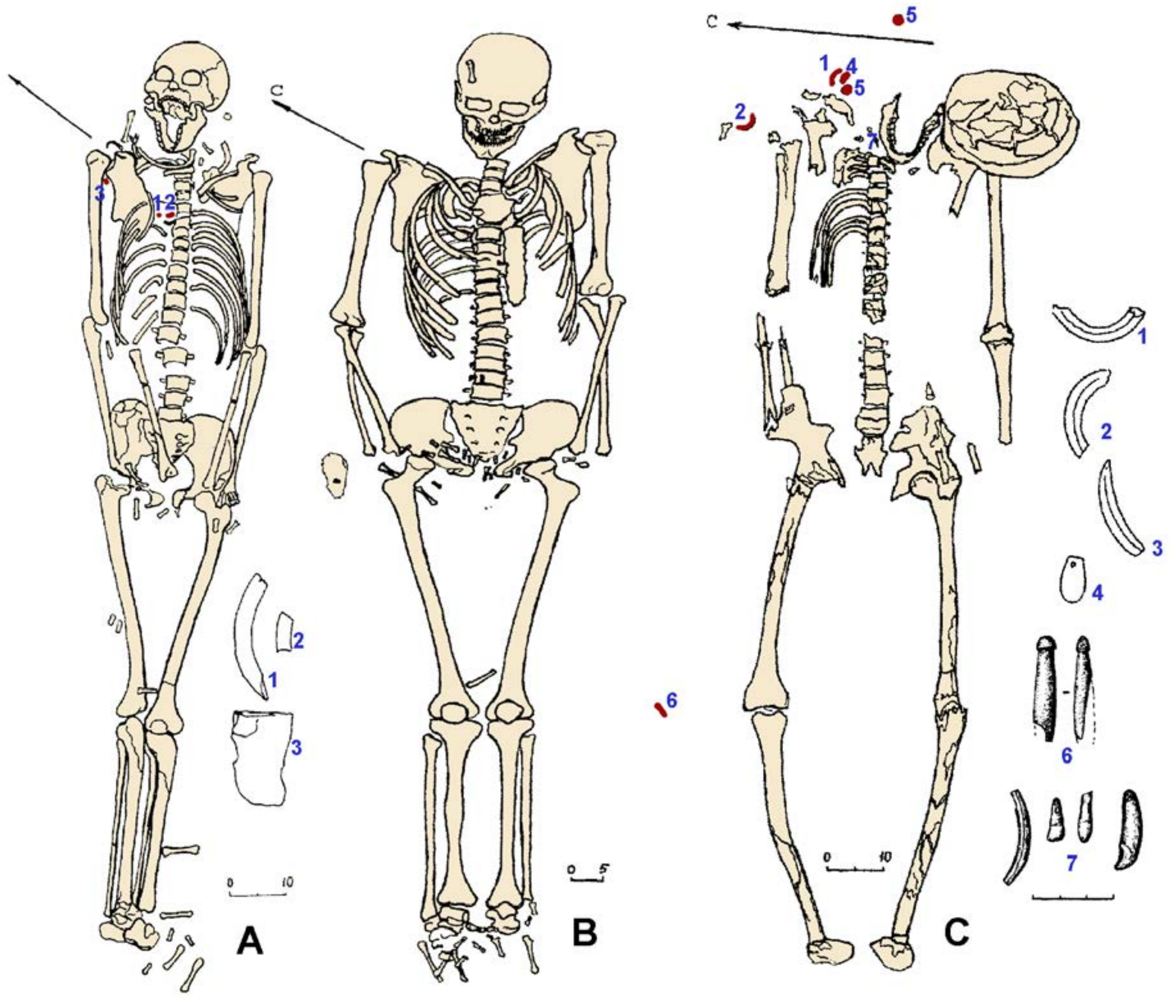

***Fig. S22.*** *Burials 3, 12, and 8 at Itkul' (Bolshoy Mys). **A** - burial 3: **1-2** - teeth of beaver and badger; **3** - quartzite flake; **B** - burial 12; **C** - burial 8: **1-3** - beaver's and badger's incisors; **4** - pendant; **5** - gypsum desert roses; **6** - fragment of a fishing hook; **7** - perforated animal teeth.*

*(e) Itkul' (Bolshoy Mys), Burial 17 individual 1 (I12873)*

**Burial 17**: The burial was excavated by V.I. Molodin in 1976. Two individuals were buried at a depth of 1 m from the present surface. The skeletons were positioned supine with their heads facing northeast, and their arms extended alongside their bodies. The upper part of both skeletons had been disturbed by a badger hole, with the skulls of both individuals found in different places within the hole.

***Skeleton 1*** (referred to as "A", individual ID I12873), according to the assessment of anthropologists V.P. Alekseeva and N.N. Mamonova, belonged to an 18-year-old young man, while according to V.A. Dremova, it belonged to a mature woman. The genetically determined sex of this individual is male.

*Skeleton 2* (referred to as "B") belonged to a 25-year-old woman. Next to the right hand of the deceased, a large bone needle case made from the tubular bone of an animal was found.

On Skeleton 1, a significant number of decorations were discovered in the area of the chest and neck, as well as on both skeletons in the waist area, below the pelvic bones and hands, and around the wrists. These decorations, which were likely sewn onto their clothing, included bone pendants with hanging holes, pendants in the form of stylized fish, drop-shaped ("petal") pendants, marmot incisors, threads made from tubular bones, nacre clothing appliques made from shells. V.I. Molodin, the researcher of the burial, noted the similarity of the costumes worn by both deceased individuals (Kiryushin et al. 2000).

*(f) Itkul' (Bolshoy Mys) burial ground, waste of dugout (burial A) (I1962)*

One more sample from Itkul' (Bolshoy Mys) comes from the **waste of a dugout** (individual ID I1962) that destroyed a Neolithic burial in 1954. According to the description made by the developer, he unearthed two or four skeletons oriented to the northeast. This burial is labeled as A. No specific details of the funeral rite were recorded, and human remains were collected in the course of the exploration of the dugout waste by archaeologists. The individual is dated to 6226–6031 calBCE (7260B±40 BP, Poz-82197).

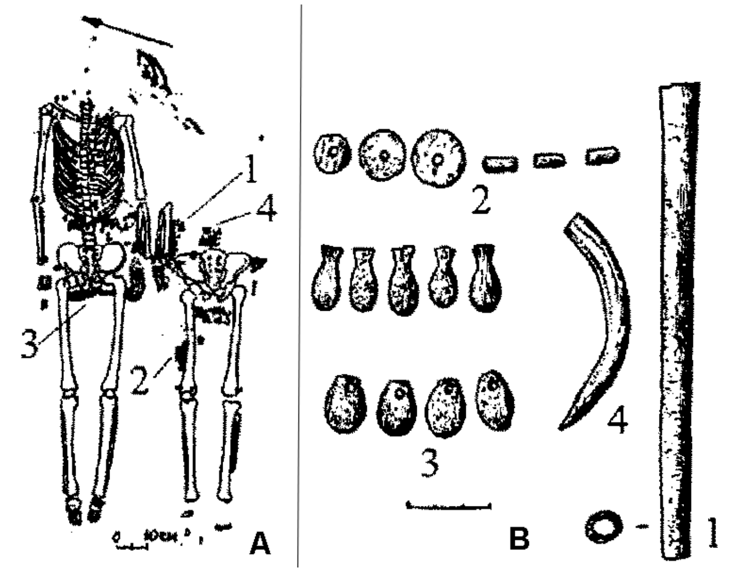

**Fig. *S23.*** *Burial 17 at Itkul' (Bolshoy Mys).* ***A*** *- burial 17;* ***B*** *- grave goods:* ***1*** *- needle case;* ***2*** *- nacre clothing appliques;* ***3-4*** *- pendants.*

*(g) Kostenkova-Izbushka burial ground, burial 3 (I2137)*

**Burial 3** (individual ID I2137) was attributed to the Late Bronze Age, and the information about it has not been published in any papers describing the Neolithic burial site at Kostenkova Izbushka. However, the radiocarbon date of the human bone sample revealed it is Neolithic, 6226–6031 calBCE (7260±40 BP, Poz-82197), and the burial must be treated together with burials 4 and 5 from the same site.

*(h) Kostenkova-Izbushka burial ground, burial 3 (I2137) and 5 (I20312)*

**Burial 5** (individual ID I20312) was situated within the cultural layer of the settlement at a depth of 0.5 m, resulting in indistinct outlines of the grave spot. The grave contained an incomplete skeleton of an approximately 40-year-old man, interred on his back with his head oriented towards the southwest. The skeletal positioning appeared somewhat unnatural. The left leg was extended in an east-west direction, while the right leg was bent at the knee. The southwest direction was determined based on the orientation of the pelvis. The right arm and the bones of the right half of the torso were missing. The left arm was bent at the elbow, with the wrist resting on the pelvic bones. In the vicinity of the base of the skull and the cranial-maxillary cavity, a biconical bone arrowhead was discovered. Fragments of vessel walls were also found within the grave. Adjacent to the right knee, an upper jaw of a beaver was present, while the second upper jaw of a beaver was located to the left of the skull in the chin area.

The deceased individual had been struck by an arrow with a bone tip, which was buried alongside him. The shot was fired at close range from above, descending downwards. The arrow penetrated through the mouth, resulting in the breakage of the top two right incisors and the dislodgement of the bottom three incisors. The impact was so forceful that it caused the jaw, where the tooth sockets were situated, to collapse from the inside. It appears that the arrow pierced the larynx, grazed the cervical vertebrae, and continued towards the left. There are visible marks of the impact on the cervical vertebrae. In this case, distinguishing between posthumous ritual and the act of killing, along with its potential nuances, is quite challenging. The arrowhead type is similar to the needle-shaped tips found at the Shigir culture sites. It exhibits a round cross-section, wider in the region of the tip. Comparable examples include an arrowhead from burial 6 of the Protoka burial ground (Polosmak, Chikisheva, Balueva, 1989).

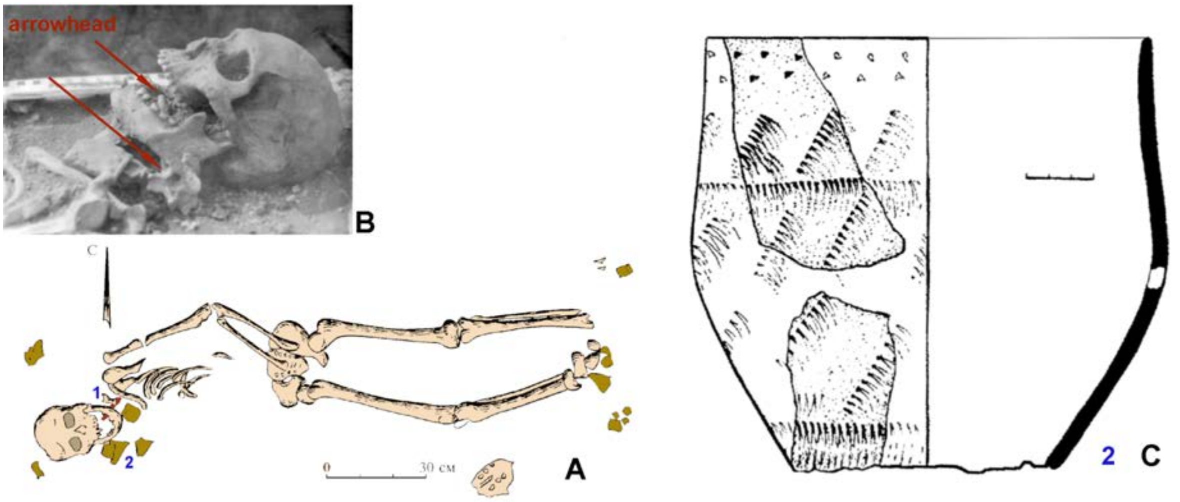

***Fig. S24.*** *Burial 5 at Kostenkova Izbushka.* ***A*** *- burial 5 (**1** - arrowhead; **2** - pottery vessel);* ***B*** *- position of the arrowhead in the burial;* ***C*** *- pottery vessel.*

##### (7) Ust'-Isha site

The Ust-Isha burial ground is situated near the village of Most-Isha, on a cape-shaped section of the elevated riverbank of the Ishi River, where a small stream flows into it. The site was initially discovered and investigated by B.Kh. Kadikov in 1961, with the majority of the findings already published.

The cemetery comprises a total of 14 burials, of which 11 are securely attributed to the Neolithic period. Two additional burials, with the skeletons positioned in a crouched posture on the left side (with no detailed description available), are presumed to also belong to the Neolithic period (Marochkin, 2014). Lastly, one female burial dates back to the early Iron Age. The burials are arranged in a single row, oriented along a northwest-southeast line, perpendicular to the cape's direction (southwest-northeast).

Several burials (1, 2, 3, 12) have suffered partial destruction due to activities related to the early Iron Age settlement and burial ground. Burials 1, 4, 8, and 9 contain the remains of males aged 30-60 years old. Females (about 30 years old) are interred in burials 2, 6, and 7. Burials 3, 5, and 12 contain the remains of children and adolescents, with their sex undetermined. Burials 2, 3, and 12 are double burials, while the remaining burials are single. The primary burial structure consists of soil pits, whose dimensions and proportions are determined by the size of the interred remains. In some cases, shale deposits were observed at the filling level along the pits' perimeters. The most common position for the corpses is extended on their backs. Burials 7 and 8 exhibit a distinctive position, with the hands placed on the pelvic bones and the shins tightly drawn towards the body. Two burials were made in a crouched position. Evidence of multi-stage burials (lack of joints and small skeletal elements) can be observed in burials 12 (a, b) and 7 (adolescent burials). Additionally, the teenage burials at Ust-Isha show indications of partial cremation.

All burials are oriented with the feet pointing downstream of the main reservoir, along the cape's alignment. Regarding cardinal directions, there is minimal variability: all male burials and one child's burial exhibit a northeast-facing orientation, while female remains are positioned with their heads to the east (two burials) and east-southeast (one burial) (Kiryushin, Kungurova, Kadikov, 2000; Marochkin, 2014).

*(a)    burial 4 (individual ID I20314).*

**Burial 4** (individual ID I20314)**:** The skeleton of a 30-35-year-old man was found at a depth of 0.45 m from the surface, lying on his back with his head to the northeast. The back of his head was elevated, and his arms were positioned alongside his body. Adjacent to the waist, a large stone dagger and a bone slotted dagger were discovered. Beneath the left leg, a flake and a bone awl were found. Numerous bone clothing appliques and pendants were present in the skull and torso region. The skull displayed 8-shaped appliques. Fish-shaped ornaments adorned the left side of the body, forming chains and rows. These decorations encircled the left arm and were most densely concentrated in the upper part of the sacrum, demarcating the upper third of the pelvis. Clothing appliques made from small incisors of a rodent (possibly marmot or beaver) extended from the left elbow to the middle of the humerus. Similar decorations were also observed on the middle section of the right humerus. Large animal teeth were arranged in a chain-like pattern, extending from the back side, spanning from the right shoulder to the waist, and forming a diagonal row between the human hips. Several specimens were noted beneath the pelvic bones. The height of the buried individual was estimated at 1.60 m (Kiryushin, Kungurova, Kadikov, 2000). The burial is dated to 4442–4266 calBCE (5500B±25 BP, PSUAMS-9052).

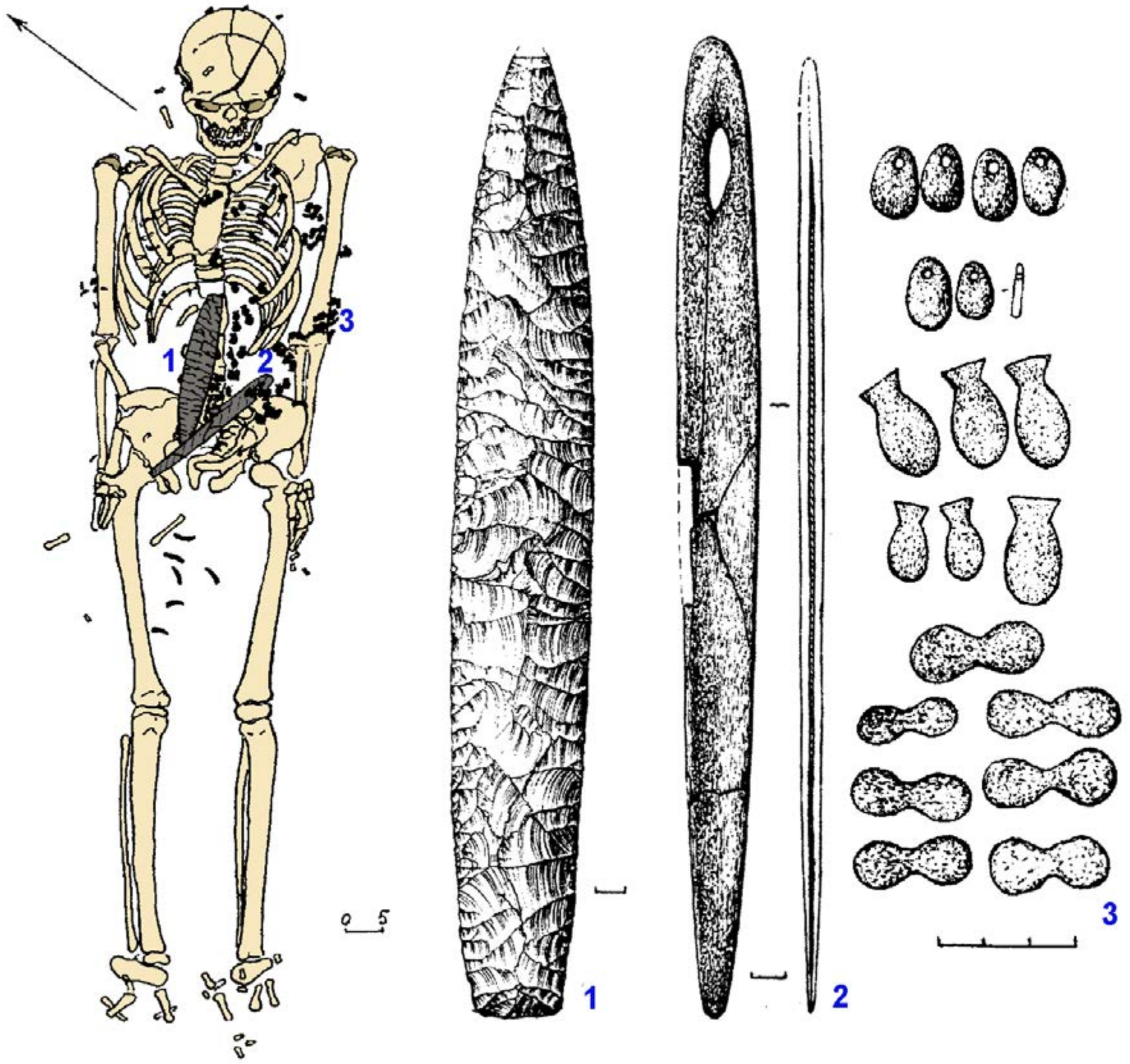

***Fig. S25.*** *Burial 4 at Ust'-Isha.* ***1*** *- stone dagger;* ***2*** *- bone dagger;* ***3*** *- pendants.*

*(b) burial 6 (Individual ID I1963).*

**Burial 6** (individual ID I1963): This burial was found at a depth of 0.30 m from the modern surface. The skeletal remains were highly dispersed and incomplete within the grave. However, the arrangement of the arm, sacrum, pelvis, and skull suggests an extended supine position for the interred individual. The skull, which was fragmented, lay on the right side. Presumably, the deceased was placed with their head pointing northeast. The burial lacked any ornaments or objects, with the exception of a dart tip located 25 cm to the left of the burial. The total height of the buried individual was 1.52 m. The gender was indeterminate, while the estimated age ranged from 40 to 50 years (Kiryushin, Kungurova, Kadikov, 2000). The burial is dated to 3957–3796 calBCE (5076B±29 BP) [R_combine: (5490B±40 BP, Poz-82209); (4585B±40 BP, OxA-33424)].

*(c) burial 7 (Individual ID I2142).*

**Burial 7** (individual ID I2142): The grave stain was traced from a depth of 0.28 m from the modern surface. The skeleton of a 20-30-years-old woman was discovered at a depth of 0.70 m from the surface, in a supine position with her head to the southeast. The left arm was extended alongside the body, while the right arm was slightly bent at the elbow and placed on the pelvis. The woman's skull was crushed and turned towards the left side. The costal bones were poorly preserved, the pelvic bones were crushed, and the spinal bones were missing. In the neck area, several pendants made of deer and elk teeth were found. Two pendants discovered in an empty lower chest appeared to be part of a previous cluster. The length of the skeleton was 1.62 m. The skulls of burials 6 and 7 were located approximately 0.40 m apart from each other (Kiryushin, Kungurova, Kadikov, 2000). The burial is dated to 5300–5032 calBCE (6200B±40 BP, Poz-82207).

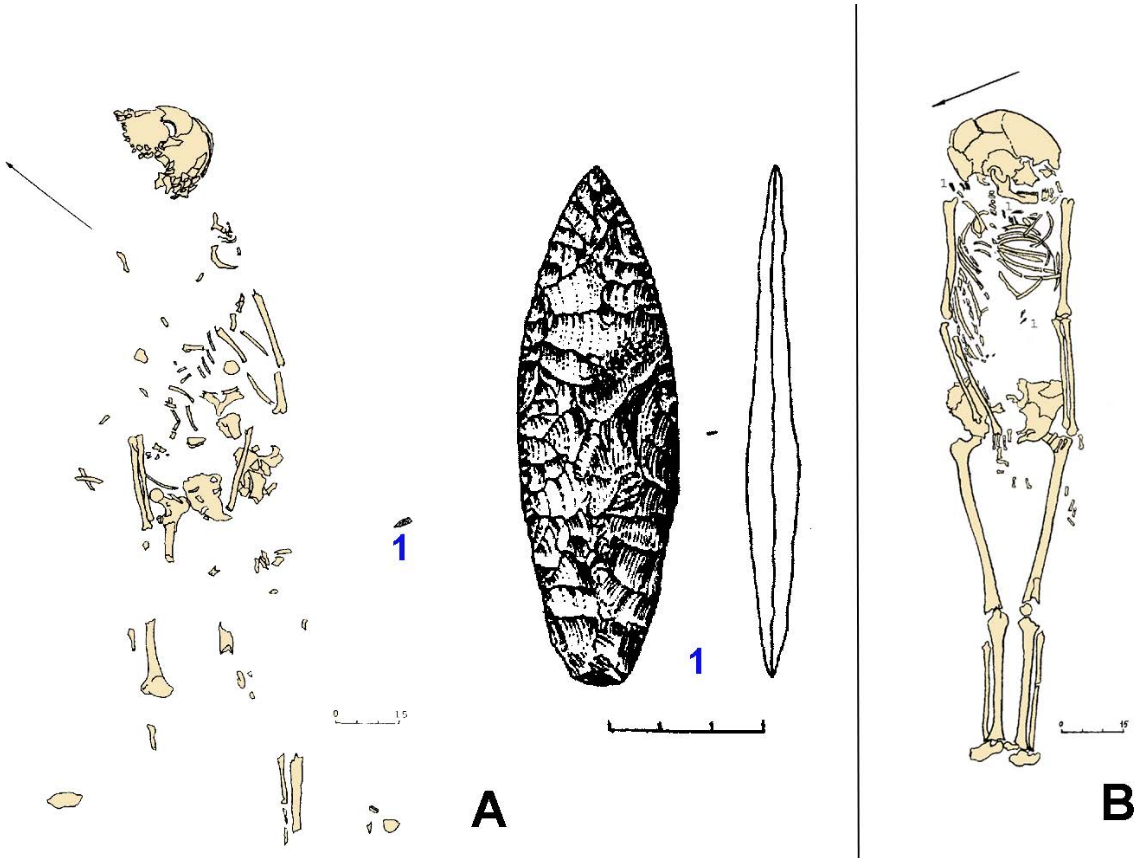

***Fig. S26.*** *Burials 6 and 7 at Ust'-Isha.* ***A*** *- burial 6 (**1** - point of a javelin);* ***B*** *- burial 7.*

*(d) burial 8 (Individual ID I12876)*

**Burial 8** (individual ID I12876): A 25-30-years-old man was interred in a supine position with his head facing northeast and his arms extended alongside his body. The right shoulder was inwardly rotated, while the head was slightly bowed and turned to the left side. Within the waist region of the skeleton, four daggers were present, arranged in pairs. In the center was a shortened stone dagger and a large slotted bone dagger, while on the right side, two identical slotted daggers were positioned. A compact cluster of 16 arrowheads was situated 30 cm from the left femur. The burial exhibited an abundance of decorative elements such as beads, bone patches, and pendants. A scattering of beads was found opposite the left humerus, within a 15 x 30 cm section. Along the arm, clusters of pendants made from large incisors of maral, elk, beaver, and two hollow bones of a bird were observed. A total of 3-4 clusters, resembling "ligaments" of large incisors, were noted. Beads were also present in the area spanning from the waist to the middle of the thigh bones. Positioned between the femurs in the upper part, two flaps of denticulate shells were found. On the right side, descending in a line from the belt area to the upper third of the pelvis, eight-shaped bone clothing appliques were present. The left half of the skeleton was adorned with "petal" pendants made of deer teeth, descending along the left arm, curving around it, and tightly clustered in the elbow area. Double pendants crafted from elk teeth were positioned beneath the skeleton along the spine. Concentrated on the skull, specifically along the back of the head, forehead, and crown, were clothing appliques formed from the teeth of a small artiodactyl animal (roe deer). Within the cluster on the frontal part, 15 such clothing appliques were observed. The height of the buried individual was estimated at 1.70 m (Kiryushin, Kungurova, Kadikov, 2000).

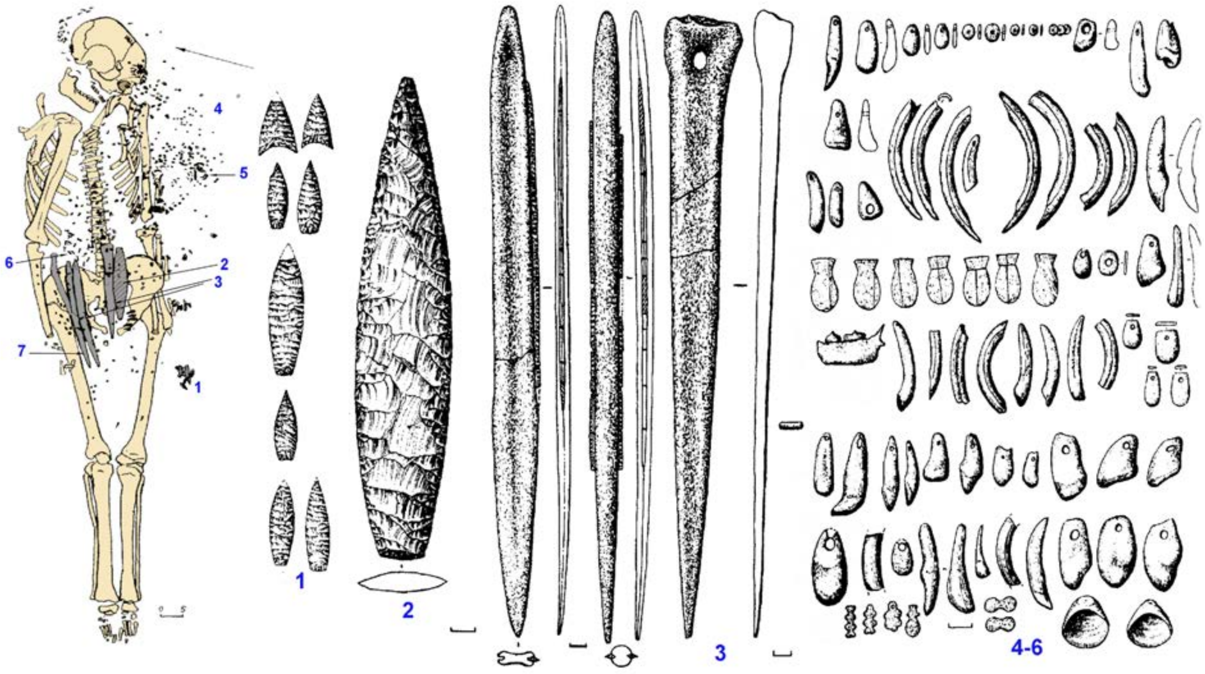

***Fig. S27.*** *Burial 8 at Ust'-Isha.* ***1*** *- arrowheads;* ***2*** *- stone dagger;* ***3*** *- bone daggers;* ***4*** *- nacre beads and pendants made of stone and bone;* ***5*** *- beaver's incisors;* ***6*** *- eight-shaped pendants;* ***7*** *- shell.*

*(e) Burials 10 (Individual ID I1966) and 11 (Individual ID I1965).*

**Burials 10** (individual ID I1966) **and 11** (individual ID I1965) have not been described properly in the publications. In one of them, a skeleton was placed in a crouched position on its left side with its head to the north-northeast. Another one contained a skeleton in a supine position with his legs bent and put on their right side, oriented to the west-northwest (Kiryushin, Kungurova, Kadikov, 2000). Burial 10 is dated to 4361–4174 calBCE (5450B±40 BP, Poz-82208), and burial 11 – to 4037–3797 calBCE (5120B±40 BP, Poz-82206).

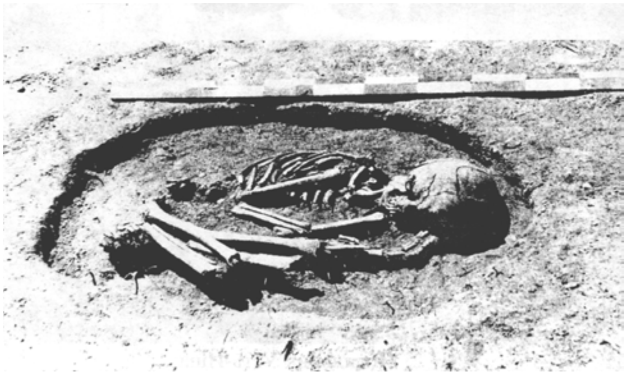

***Fig. S28.*** *Burial 10 or 11 at Ust'-Isha.*

#### (8) Firsovo-14 burial site

The Firsovo-14 burial site is situated on the right bank of the Ob River, opposite Barnaul, 1 kilometer west of the village bearing the same name. It is located in the forest-steppe zone of Altai, on a promontory that rises four meters above the floodplain of the Ob. The site is positioned above the oxbow lake of the Losikha River, which is a right tributary of the Ob.

Since 1987, a total area of over 5,000 square meters has been excavated at the site, and approximately 400 burials have been investigated. Among these burials, around 200 belong to the Andronovo culture, about 20 are attributed to the Irmen culture, and over 100 are associated with the Staroaleyka culture. Additionally, single graves representing other periods and burial traditions have been discovered at the site. Over the years, one Korchazhkino burial and two burials displaying the Sargary appearance from the Late Bronze Age were excavated here (Kiryushin et al. 2013).

##### *(a) Neolithic burial 267 (individual ID I11754)*

**Burial 267** (Individual ID I11754) was excavated in 1996 at the end of the promontory. The exact outline of the grave pit could not be determined. The skeletal remains were found at a depth of 0.4 meters from the modern surface, within a layer of yellow sandy loam. The deceased was positioned on their right side, with the head facing east. The legs were flexed at the knees and pressed against the body, causing the feet to be near the pelvic bones. The arms were bent at the elbow joints and pressed against the chest. The right hand was positioned near the face, while the left hand was beneath the skull. The overall posture of the skeleton can be described as resembling the "fetal position”. The positioning of the legs and back clearly indicates the limited space within which the body was placed. The placement of the head appears to be unnatural. It is likely that these factors can be attributed to the size of the grave pit, against which the deceased's knees and back were pressed. One should not exclude the possibility that prior to burial, the individual was tightly wrapped or sewn into a shroud. The buried individual was laid with their feet perpendicular to the flow of the Ob River. In relation to the nearby Losikha oxbow lake, the deceased was positioned parallel to the channel, with their back facing the water, head elevated, and feet pointing downstream.

Artifacts made of bone, antler, and stone were discovered within the grave alongside the deceased. All of the findings were grouped together in one location, near the chest and between the arms. The accompanying funerary inventory consists of the following items:

1. a 16.5 cm long polished bone "point";
2. a 12.1 cm long polished antler spatula;
3. a fragment of a polished antler rod, measuring 5.1 cm in length with a cross-section of 0.9x0.6 cm;
4. a distal fragment of a harpoon featuring teeth from an animal's rib on one side;
5. the proximal part of a flake with finely retouched lateral edges;
6. an abrasive tool measuring 7.1 cm in length, with a side face width of 1.8–2.2 cm and a working surface width of 1.3 cm. It is made of fine-grained sandstone;
7. a polished knife measuring 7.9 cm in length, 1.4–2.2 cm in width.

The compact arrangement of the accompanying inventory suggests that all of these items could have been contained within some type of handbag, which was placed in the hands of the deceased (Kiryushin et al. 2013). The burial is dated to 4994–4797 calBCE (6005±35 BP, PSUAMS-8988). Alternatively, two other dates for the human bones were obtained in another paper, 5318-4846 calBCE (6166±96 BP, NSKA-01942), and 5206-4935 BP (6100±25 BP, IGAN-5831) (Kiryushin et al. 2021). The combination of three dates provides an age of 5194-4905 calBCE (6073±20) [R_Combine: (6005B±35 BP, PSUAMS-8988); (6166±96 BP, NSKA-01942); (6100±25 BP, IGAN-5831)].

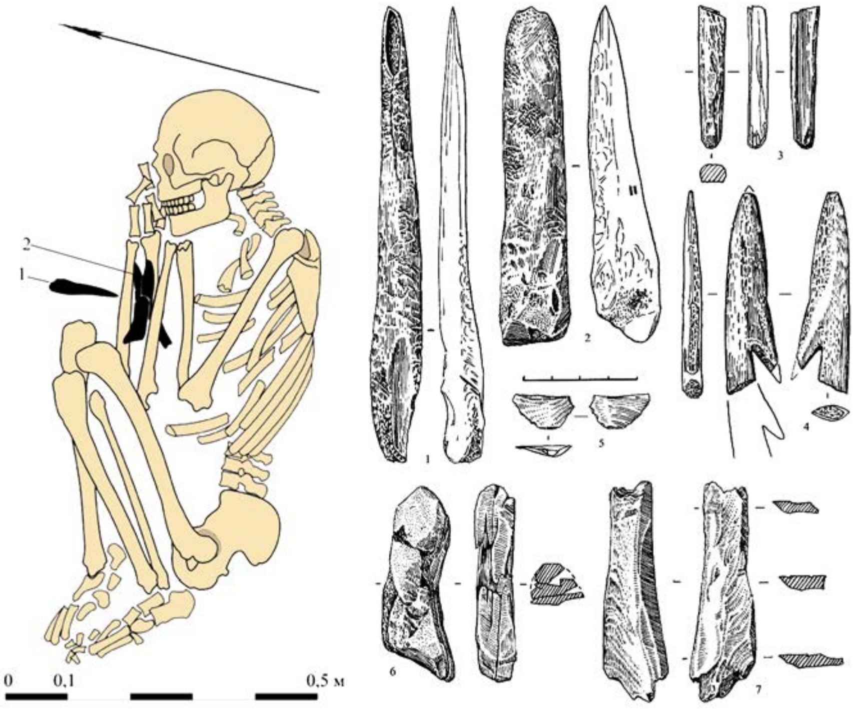

***Fig. S30.*** *Burial 267 at Firsovo-14.* ***1*** *- antler spatula;* ***2*** *- grave goods.*

##### (9) Lebedi-2 site

The Lebedi-2 cemetery is situated on a narrow promontory of the left bank of the middle Inya River. This promontory comprises the second terrace above floodplain. It is formed by loamy, sandy, and sand deposits (Bobrov 1988).

The cemetery consists of six burials and one pit containing crushed animal bones and artifacts (referred to as Pit 5). Burials 1 and 2 were partially damaged due to construction activities. Burials 1, 2, and 7 contain the remains of men over 60 years of age. Burial 3 belongs to a teenager aged 12-13, and burial 4 belongs to a child of approximately 8 years old (in both cases, the sex is undetermined). Burial 6 contains the remains of a woman over 60 years old. Burial 2 is a double burial, while the rest are single burials. The nature of Pit 5 is unclear and its purpose is uncertain (possibly a cenotaph?).

The spatial distribution of the burials follows an axis subordinated to the direction of the cape. The burials are arranged in a row, with two distinct groups within the row: the western group (burials 6 and 7) and the eastern group (burials 1-4). Pit 5 is adjacent to the eastern group.

The burial structures consist of elongated, subrectangular, or oval earth pits. There is no discernible pattern in their form variation. The size and proportions of the pits correspond to the size of the individuals interred within them, with the pits for adolescent burials being shorter. Pit 5 shares similar proportions and shape with the burial structures, although this cannot be confirmed for burial 2 due to its destruction. All other burials are characterized by the placement of the remains in an extended position on their backs. Burials 3 (children), 6 (adult female), and 7 (adult male) exhibit a specific detail where the hands are positioned on the pelvis, and the legs are tightly pressed against the shins.

The remains of the adults (burials 1, 2, 6, and 7) are oriented with their heads towards the northeastern sector in relation to the cardinal points, and with their feet downstream relative to the river. The remains of the children (burials 3 and 4) are oriented with their heads towards the northwestern sector, perpendicular to the river flow and perpendicular to the burials of adults. Pit 5 is aligned along the southwest-northeast axis with its longer side, resembling the orientation of the adult burials (Marochkin 2014).

*(a) burial 7 (individual ID I2076)*

**Burial 7** (individual ID I2076) is a narrow, almost rectangular pit; its dimensions are 2.1 x 0.57 m. The bones belonged to a man of about 60 years. He was interred extended and supine, with the head to the northeast. The position of the arms should be noted. They are extended along the body but the hands are turned toward each other and lie on the pelvis. The accompanying inventory has retained its initial position. The headdress decoration consisted of five beaver incisors (they lay on the forehead), while on each side were sewn six bear teeth. A row of animal teeth (bear, moose, and wolf) was placed along the skeleton, beginning with the neck vertebrae. At the pelvis, this row went to the right toward the radius, then turned to the center of the body, going to the left knee. Such a position conforms with clothing opened below the belt and wrapping to the right. The accompanying inventory was found on the pelvic bones (i.e., it was placed on the belt of the interred). On the left, with point up, lay a wide, leaf-shaped flint dagger (labelled "3" in Fig. ). The placement of the fingers on the left hand does not permit a conclusion that the handle was grasped. Between the hands was placed, bottom up, a round-bottomed vessel of small dimensions. It was decorated with one vertical zigzag line executed by strokes of a narrow, sharpened stick, and another line that was incised (labelled "3" in Fig. ). By the vessel and partly on the base of the dagger was a polished knife with a concave blade edge. It was of green serpentine (labelled "4" in Fig. ). Under it lay a small bone spoon or spatula with a poorly preserved straight base. To the right of the objects described was placed a small, polished axe (labelled "1" in Fig. ). In addition, two beaver incisors were found on the belt (Bobrov 1988, Marochkin 2014). Burial 7 is dated to 4689–4403 calBCE (5700B±50 BP, Poz-83432).

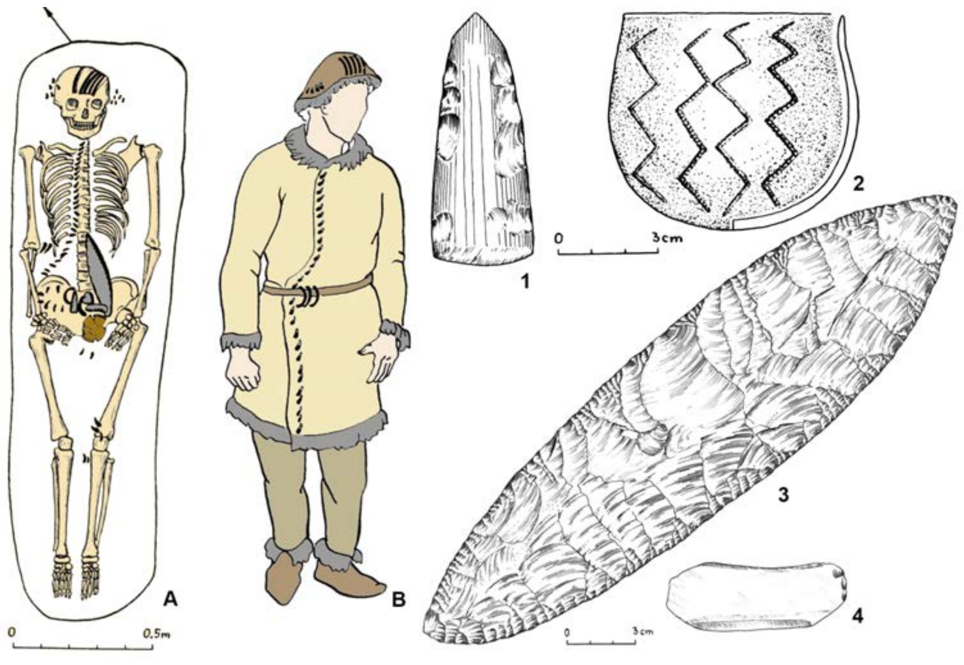

***Fig. S31.*** *Burial 7 at Lebedi-2.* ***A*** *- burial 7;* ***B*** *- reconstruction of clothing;* ***1*** *- polished axe;* ***2*** *- pottery vessel;* ***3*** *- leaf-shaped flint dagger;* ***4*** *- polished knife (Bobrov 1988).*

##### (10) Solontsy-5 site

The Solontsy-5 burial ground is a component of the Solontsy archaeological site complex, situated near the village of Pilno on a narrow cape-like area where the Biya River and its tributary, the Chapshushka River, converge. The complex of sites has been known since the 1920s, with the initial excavation of two Neolithic burials conducted in the 1970s by V.D. Lunev (Kungurova 2005). The primary burial complex was unearthed in the early 2000s during N.Yu. Kungurova's investigation of the Neolithic settlement at Solontsy-2. A comprehensive publication of the materials has been accomplished (Kungurova, Chikisheva 2002; Kungurova 2003, 2005).

The burial ground consists of nine burials, one of which (burial 9) suffered partial disturbance due to medieval settlement activities. Burials 1, 3, 4, 5, and 7 contain the remains of men ranging in age from 30-40 (burials 1 and 4) to 60 years (burials 3 and 7). In one instance, the burial of a teenager was classified as male (burial 5). The remains from burials 2 (about 60 years old), 6 (indeterminate), 9 (about 50 years old), and 8 (about 12 years old) were identified as female.

All burials are individual, except for paired burials 1 (a man and a child) and 8 (two teenage girls). The burials are arranged in two rows aligned along the northwest-southeast axis (burials 1, 3, 5, 7, 4 and 6, 9, 8, 2). Unfortunately, there is no available data regarding the nature of the burial structures as the excavation did not permit the identification of burial pits.

Two distinct positions of the deceased are evident. The first position is extended on the back (burials 1, 4-9). This variant may also encompass the paired burial 8, where the remains of two individuals are superimposed. Burials 1, 4, 8, and 9 exhibit a particular feature with the hands placed on the pelvis and the legs brought together at the shins. The second position is a flexed posture on the left side (burials 2 and 3).

Among the nine burials, six demonstrate anatomical disruption and the loss of small bones, indicating potential multi-stage ("secondary") burials. All burials display a consistent orientation with the heads facing the east-northeast sector and the feet positioned downstream the primary water reservoir, the Biya River. There are variations in orientation between men and women, with men predominantly oriented towards the east-northeast and women towards the east (Marochkin 2014, Kungurova 2003).

###### *(a) burials 1 (individual ID I2072) and 7 (I2073)*

**Burial 1:** The burial site contained the remains of an adult male, estimated to be 35-40 years old, buried in an extended supine position with his head facing northeast. The burial was located at a depth of 0.75 meters from the soil surface. Adjacent to the bones of the right hand was the skeleton of a 1.5-year-old child.

***Skeleton 1*** (Individual ID I2072): The man's skull had two stone slabs closely positioned against its occipital surface. Along the edge of one of the slabs, a large chopping tool called a biface was placed. A musk deer tusk with notches along its concave edge was placed on this slab. On the right side of the skull, there was a dense accumulation of 73 flakes. A polished axe was found beneath the lower jaw, resting on the upper ribs. Under the left shoulder bone, a dense accumulation of items was discovered, including a pestle with an expanded base, a long stone bar, five bone harpoon tips, one bone end plate for a bow, five stone arrowheads and their fragments, a bone point, and a stone rod for a hook. Underneath these items and the humerus bone, there was a dense accumulation of 34 flint flakes, including a scraper and four retouched flakes. Closer to the stone slab, two stone fishing rods were located. Near the ulna bone of the left hand, two harpoon tips were superimposed, along with a horn chipper adjacent to them, and below that, a narrow polished adze-shaped tool. Two bone rods with hook points inserted into them were found on the phalanges of the left hand. On the right side, from the lower ribs to the pubic bones, two large stone biface blades were positioned: a long dagger and a wide asymmetrical knife. A scraper made of sandstone tiles lay beneath the center of the spinal column. At the distal end of the left tibia, a piece of stone was present. Near the skull, shoulder, and hip joints, musk deer fang points were discovered. Decorative items made of large deer teeth and petal-shaped bone stripes adorned the neck and chest. Additional bone stripes from the teeth of large ungulates were located in the pelvic region and between the femurs.

***Skeleton 2***: The child's skeleton, located near the male, was slightly turned to the left, with the lower part facing the man's forearm bones. The child was buried in an extended supine position. Between the two skeletons, a bone dagger with stone inserts was placed. The bones of the child's left hand were positioned beneath it, indicating that the right hand may have been resting on top (only the ulna bones without phalanges have been preserved). The tibia bones were also found beneath the dagger and partially on it. Under the dagger, near the right knee, there was a cluster of 11 stone arrowheads and their fragments. A valve of a large shell was discovered at the articulation of the right thigh and pelvis, with two more valves of the same type located in the center of the lower part of the chest and near the left shoulder joint (likely originally placed on the chest). Four stone flakes were situated under the mandibula. A piece of hematite (red mineral paint) was found beneath the spine in the waist area. The burial also contained split teeth of a large ungulate, positioned under the dagger, on the bones of the left hand, on the ribs of the left side of the chest, and between them.

The items discovered in this burial comprise a diverse and rich assortment. Comparative morphological analysis of the items from burial 1 indicates their similarity and possible cultural and chronological affinity with artifacts from the Neolithic Burial at Ust-Aleika-5 (Barnaul Ob) (Borodaev et all, 2022). Burial 1 is dated to 3958–3713 calBCE (5050B±40 BP, Poz-83497).

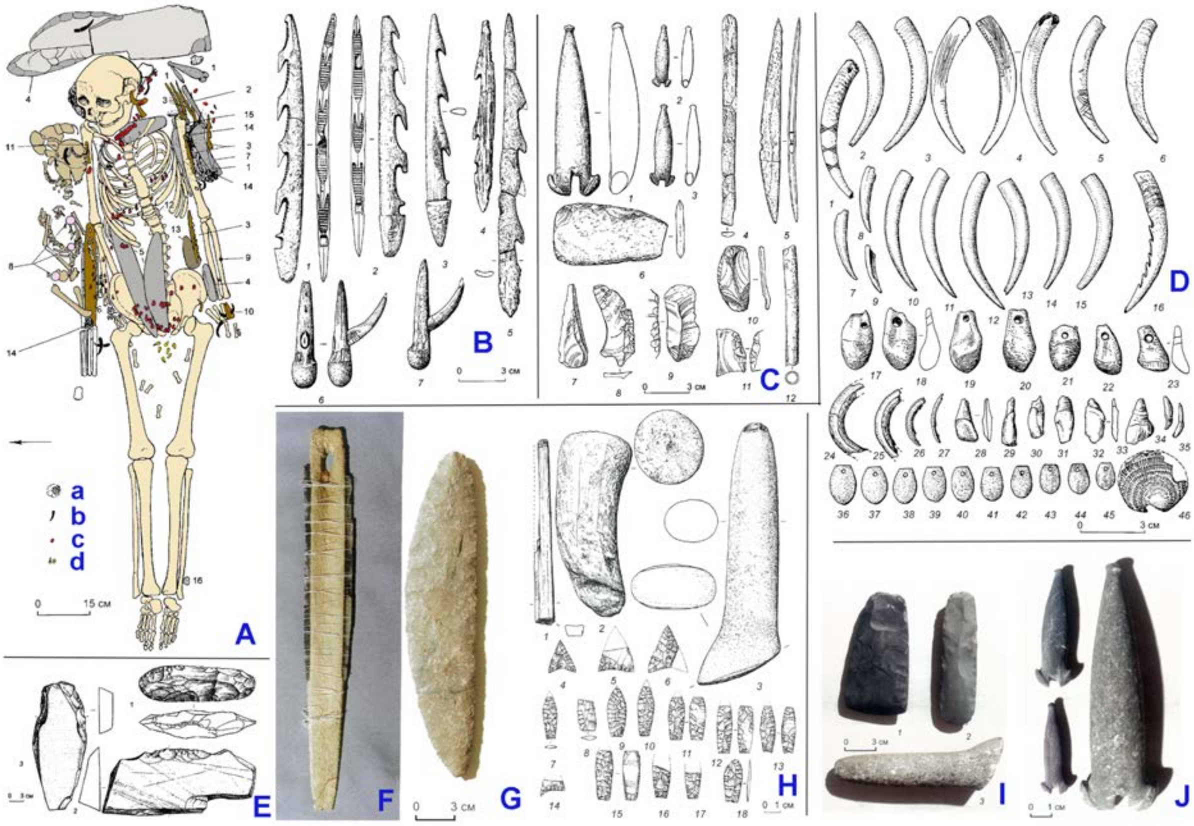

***Fig. S32.*** *Burial 1 at Solontsy-5.* ***A*** *- burial 1 (dark color - child bones, light color - adult bones):* ***a*** *- flakes;* ***b*** *- musk deer tusks;* ***c*** *- bone appliques;* ***d*** *- teeth appliques;* ***1*** *- fishing rods;* ***2*** *- grinding stone;* ***3*** *- harpoon tips;* ***4*** *- chisels;* ***5*** *- knives;* ***6*** *- slotted dagger;* ***7*** *- "small flat iron" with a handle;* ***8-9*** *- shells;* ***10*** *- bone hooked rods;* ***11*** *- bones;* ***12*** *- scraper;* ***13*** *- fabricator;* ***14*** *- arrowheads;* ***15*** *- scraper made from a flake;* ***16*** *- stone;* ***B*** *- bone tools;* ***C*** *- various grave goods;* ***D*** *- ornaments, pendants, and*

*appliques; **E** - bifacial axe and stone plates; **F** - slotted bone dagger; **G** - stone dagger; **H** - stone implements and arrowheads; **I** - photo of stone tools; **J** - photo of stone fishing rods (Kungurova 2003, modified).*

*(b) burial 7 (individual ID I2073)*

**Burial 7** (individual ID I2073): The burial consisted of the remains of a 55-60-year-old man buried at a depth of 0.70 meters from soil surface. The position of the body was supine, with the head oriented towards the east and the arms extended along the sides. The facial bones were compressed within the skull, and the lower jaw exhibited downward and leftward displacement. A circular scarlet pigment spot was observed in the vicinity of the skull and to its left side. Positioned on the left side of the burial was a large stone biface dagger. Additionally, a stone arrowhead was discovered in the lower lumbar region of the spine, with its point directed towards the vertebra. The burial exhibited an abundance of decorative items made from animal teeth and bone petal patches, which were arranged in ribbons. A small flake and a fragment of ferruginous rock used as a pigment were also recovered from the burial. The estimated length of the skeleton ranged from 175 to 180 cm (Kungurova 2005). Burial 7 is dated to 3971–3802 calBCE (5105B±25 BP, PSUAMS-4120).

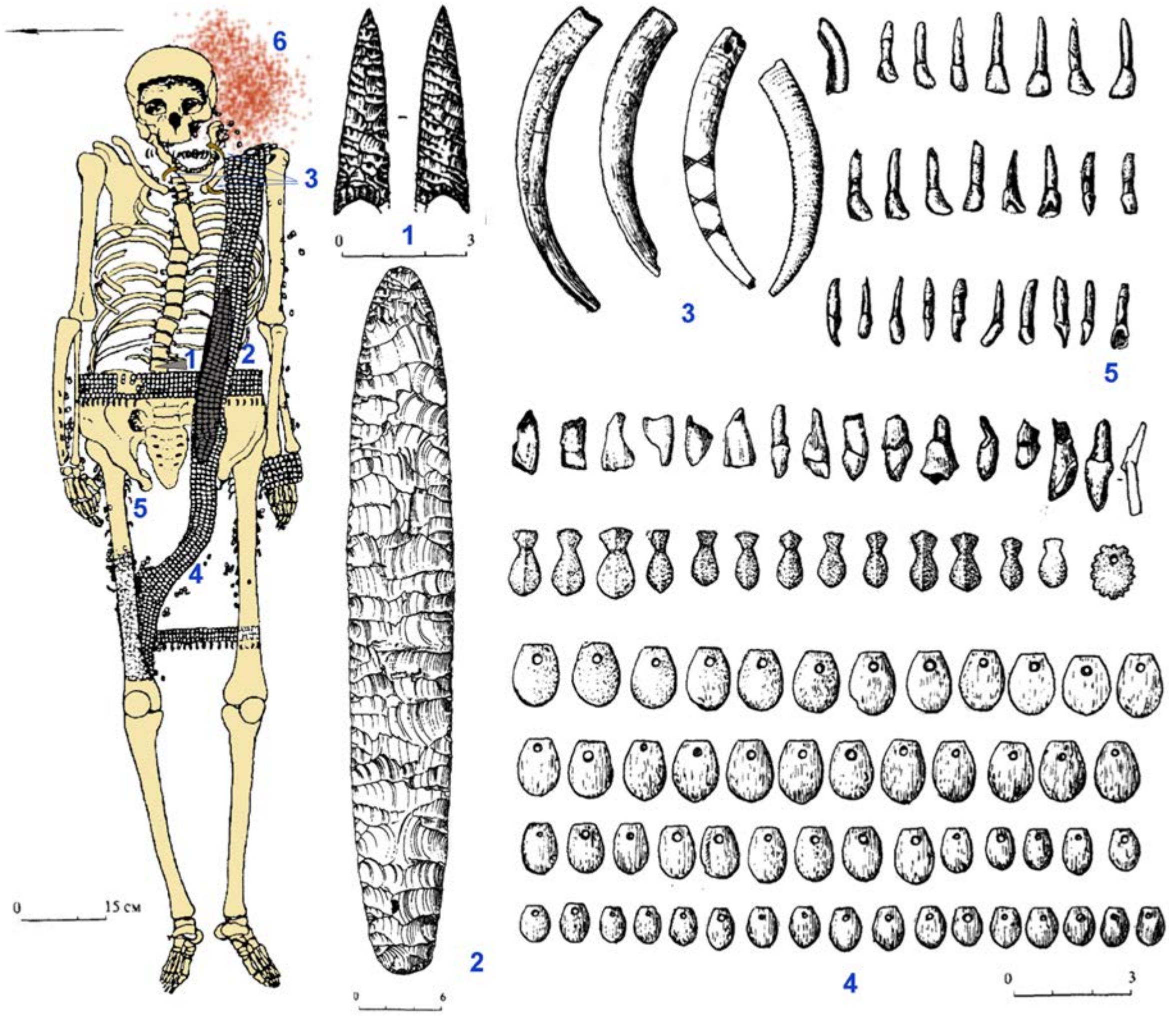

***Fig. S33.** Burial 7 at Solontsy-5. **1** - arrowhead; **2** - bifacial dagger; **3** - musk deer tusks; **4** - appliques; **5** - animal teeth; **6** - ocher spot (Kungurova 2005, modified).*

##### (11) Tuzovskiye Bugry-1 Late Neolithic-Eneolithic site

The Tuzovskie Bugry-1 burial ground, also known as Vasino-5, was discovered in 1999 by D.A. Pugachev and subsequently investigated in 2000 by M.T. Abdulganeev.

The site is situated on the right bank of the Ob River, characterized by a wide floodplain (up to 12 km) with swampy areas and numerous oxbow lakes. These features are interspersed with crests and hillocks that emerge from the bedrock bank. The surrounding vegetation consists of meadows with dense thickets of shrubby willows, aspens, and birches.

Excavations at the site covered an area of 174 square meters, revealing a total of 37 burials. Among these, 19 can be attributed to the Eneolithic and Early Bronze Age periods. Unfortunately, some of these graves were destroyed due to the construction of an early Iron Age necropolis. The Eneolithic burials were arranged in four rows oriented along a southeast-northwest axis, each row containing 5-6 burials. Most of the deceased individuals were found in an extended supine position, with their arms lying alongside their bodies and their heads oriented towards the northeast. The majority of the burials were single, while three were paired and two were collective, containing three and five individuals, respectively.

Thus far, 12 radiocarbon AMS dates have been obtained, with 11 derived from human bones and teeth, and one from a mountain goat tooth. These results have raised the issue of a reservoir effect, which needs further investigation. Noteworthy chronological indicators in the archaeological context include decorations crafted from mollusk shells (Dentalium and Corbicula) as well as animal teeth (such as musk deer, mountain goat, or ram). Based on the available evidence, the Eneolithic burials can be dated to 3600-3200 calBCE.

One specific burial, grave 34, can be attributed to the Early Bronze Age. Burials 4 and 18 lacked any accompanying inventory, likely due to disturbance during the construction of early Iron Age graves. Notable artifacts found in these contexts include decorations made from mollusk shells (Dentalium and Corbicula), which were imported (Kiryushin et all, 2011), and animal teeth (musk deer, mountain goat, or ram) (Kiryushin et all, 2012). Burial 7 yielded pottery of Bolshoy Mys and Comb-Pit pottery types within its fill (Kiryushin and Kiryushin, 2015).

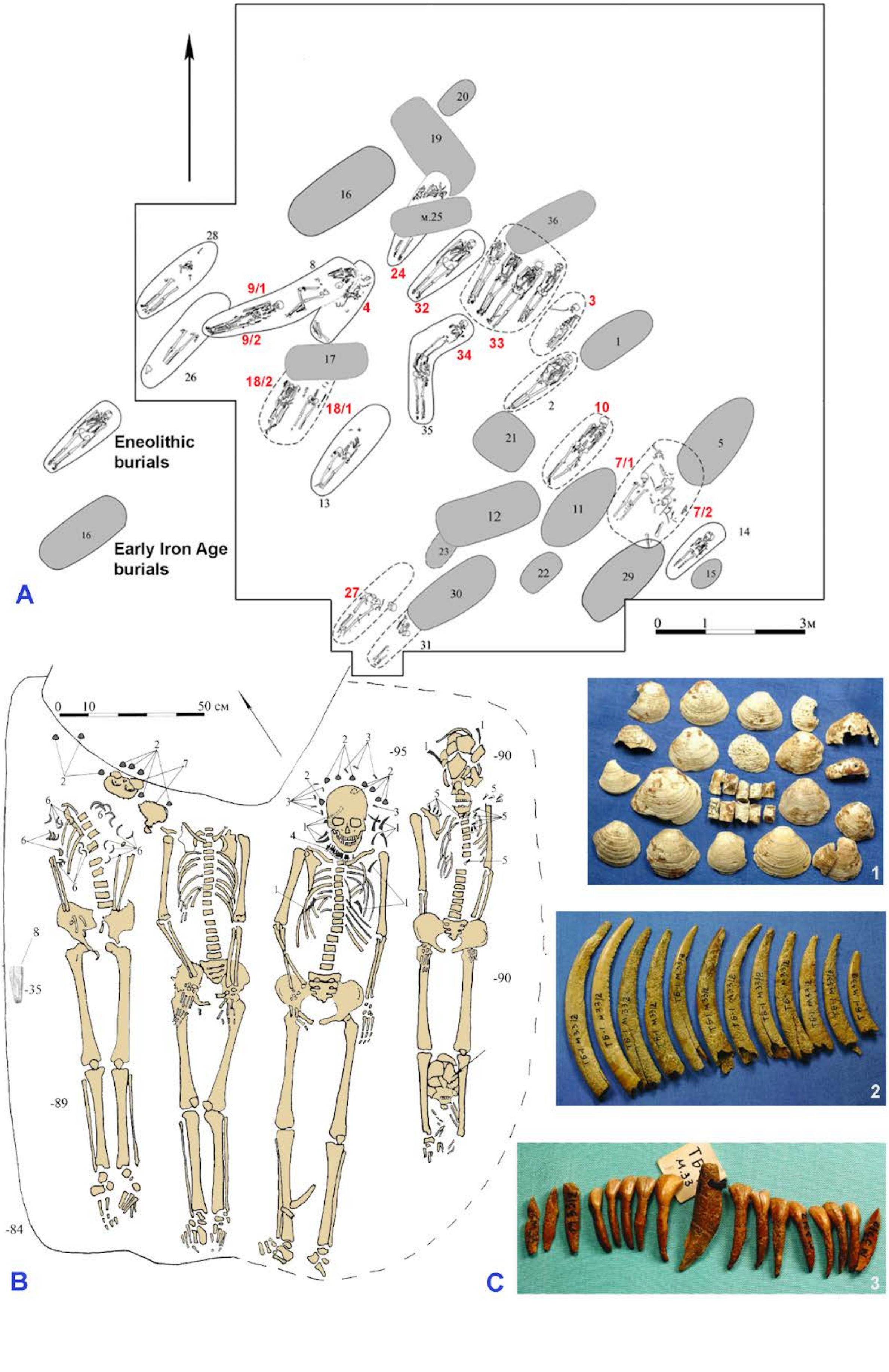

Eneolithic burials
Early Iron Age burials
A
B
C
0 1 3м
0 10 50 см
9/1
9/2
4
24
32
33
3
34
18/2
18/1
10
7/1
7/2
27
м.25
-95
-90
-89
-84
-35

***Fig. S34.*** *Tuzovskiye Bugry-1 burial ground. A: a plan of the cemetery, sequenced samples are labeled in red; B: burial 33: 1 - tusks of musk deer, 2 - Corbicula shells, 3 - Dentalium shells, 4 - necklace with 18 small animal incisors, 5 - necklace with 6 small animal incisors, 6 - necklace with 25 marmot incisors, 7 - skull fragments; C: grave goods from burial 33: 1 - mollusk shells, 2 - tusks of musk deer, 3 - pendant from the upper fang of a badger and small incisors of a musk deer (Kiryushin et al., 2011, 2012, 2015, modified).*

*(a) burial 3 (individual ID I11747)*

**Burial 3** (individual ID I11747) was situated adjacent to burial 33 and marked the southeastern end of the "first" row. It was found at a depth of 0.6 meters from the current surface, and no distinct grave stain was visible. The buried individual, a male aged 35-45 years, was interred in a supine position with the head oriented to the northeast. Some bones were missing, and the remaining ones were disordered. The burial took place long after the individual's death. The grave goods included two *Corbicula* shells, a horn point, and a pendant made from a deer tooth with a perforation.

*(b) burial 4 (I12323)*

**Burial 4** (individual ID I12323) was located in the northwestern part of the second "central" row and was partially intersected by burial 8. It had a southwest to northeast orientation along its long axis, with a depth ranging from 0.45 to 0.6 meters. The precise shape of the grave pit could not be determined. The buried individual was a man aged 40-55 years. Similar to other burials, this one also took place long after death and lacked any grave goods.

(c) *burial* 7 Individuals 1 (*ID I11750) and 2 (ID I11748*)

**Burial 7** was situated in the southeastern part of the second "central" row, between burials 10 and 14. Its approximate dimensions were 2.15 x 1.4 meters, with the long axis half aligned with a southwest to northeast line. Burial 7 had been disturbed by burials 5 (east corner), 29 (south corner), and possibly burial 11 (west corner). The burial occurred long after the individuals' deaths. The grave pit contained disarticulated bones of a man aged 30-40 years (individual 1, individual ID I11750) and a woman aged 40-55 years (individual 2, individual ID I11748), with the anatomical order disrupted. Grave goods included a bone harpoon and fragments of Colletopterum sp. Additionally, two AMS dates were obtained based on a fragment of a human radius bone: 3810–3700 calBCE (5005±25 BP,IGAN-5832) and 4450–4030 BCE (5409±93 BP, NSKA-01943). A combination of these two dates (5409±93 BP, (NSKA-01943) yield an age of 3949-3714 calBCE (5034±25 BP) [R_Combine: (5005±25 BP, IGAN-5832); (5409±93 BP, NSKA-01943)].

*(d) burial 9, Individuals 1 (ID I12628) and 2 (ID I10965)*

**Burial 9**, along with burial 8, was positioned between the second and third rows, cutting through parts of burials 4 and 26. It had a west-to-east orientation with a slight deviation to the north. This was a paired burial, with the skeletons of an adult male aged 40-55 years (individual ID I12628) and a child aged 4-6 years (individual ID I10965) located at the bottom of the grave. Both individuals were found in a supine position, with their heads pointing northeast. The adult was missing the left humerus and some ribs, which had fallen to the left due to the construction of the neighboring burial 8. The child's body was positioned atop the adult, with the skull resting on the adult's abdomen and the leg bones positioned between the adult's legs. The grave goods consisted of 15 ornaments made from Dentalium shells. Child skeleton from the grave was dated to 4037-3807 calBCE (5135±25 BP, PSUAMS-8879), and the adult skeleton to 3958-3798 calBCE (5080±25 BP, PSUAMS-9038). A combination of probability densities yields a date of 3970-3804 calBCE (5108±18 BP) [R_Combine: (5135±25 BP, PSUAMS-8879); (5080±25 BP, PSUAMS-9038)].

*(e) Burial 10 (I12627)*

**Burial 10** (individual ID I12627) was situated in the second "central" row, between burials 2 and 7. The skeleton of an adult woman aged 35-45 years was found at a depth of 0.6 meters from modern surface. The burial was in an extended supine position, with the head oriented to the northeast. The skull was turned to the left, and the hands were positioned under the pelvis. Grave goods included 105 ornaments made from incisors of a mountain goat or ram. Two AMS dates were obtained: one based on human bones, yielding a result of 4045-3804 calBCE (5146±37 BP, GV-03580), and the other based on a tooth of a mountain goat or ram, yielding a result of 3517-3358 calBCE (4637±37 BP, GV-03579). We expect freshwater reservoir offset in the date obtained from human bones and attribute the burial to the chronological period determined by a date from the animal tooth.

***(f)** burial 18, mandible 1 (individual ID I17582) and mandible 2 (I13260)*

**Burial 18** (**mandible 1:** individual ID I17582; **mandible 2**: individual ID I13260) was located in the "third" row between burials 26 and 13, and it had been partially disturbed by burial 17 from the Early Iron Age. The excavated area measured approximately 1.25 x 1 meters, with its long axis aligned with a southwest to northeast line. At the bottom of the grave, at a depth of 0.81 meters, were the incomplete skeletons of three individuals: two adults and one child. One of the buried individuals was determined to be a male aged 25-35 years, while the age of the child was estimated to be 8-12 years. It was challenging to ascertain the sex and age of the third skeleton, but it is likely that it belonged to a woman.

*(g) burial 24 (I17682)*

**Burial 24** (individual ID I17682) was situated adjacent to burial 32, with the northeastern part of burial 24 disturbed by burial 19 and the central part destroyed by burial 25. The exact dimensions of burial 24 were challenging to determine. It had an elongated oval shape, with approximate dimensions of 2.05 x 0.6 meters along its long axis aligned with the southwest to northeast line. The depth from the modern surface was 1 m, while it was 0.25 m below the mainland. Two burials were recorded: an adult woman aged 35-45 years (along the southeastern wall) and an infant (to the left of the adult near the northwestern wall). The adult's remains were missing the skull, left arm, radius bones, right hand, lower part of the chest, pelvic bones, both femurs, and upper part of the right tibia, which had been severed. The adult was placed in the grave in a supine position with the head oriented to the northeast. The child's body was positioned with the head pointing northeast, likely lying on the left side. Only a crushed skull and part of the chest were preserved from the child's remains. Burial goods included a fragment of a stone tool and 17 shell ornaments (round beads with a central hole) made from *Colletopterum* sp. shells. An AMS date of 3951-3656 (5016±54 BP, GV-03581) was obtained from human bones.

*(h) burial 27 (I11751)*

**Burial 27** (individual ID I11751) was located in the "third" row between burials 13 and 31. The northeastern part of the grave had been completely destroyed, and its outline was untraceable. The filling of the grave was only observed at the mainland level from a depth of 0.97 meters at the bottom. The fixed part of the grave had dimensions of 1.4 x 0.55 meters along its long axis aligned with the southwest to northeast line. At the bottom, the lower part of an adult woman's skeleton (aged 30-40 years) was found, including the legs, pelvic bones, hands, and a broken radius of the left hand. Based on the positioning of the remains, the individual was laid in a supine position with the head oriented to the northeast. Burial goods consisted of fragments of a bone item (possibly bow plates), pebbles with fine scratches, a retouched flake, 11 shell fragments of *Colletopterum* sp., and two *Corbicula* shells. Two AMS dates were obtained from human bones: 5055+57 yBP (GV-03582) and 5004+36 yBP (UBA-22957).

*(i)* *burial 32 (I0994)*

**Burial 32** (individual ID I0994) was situated in the northwestern part of the "first" row, between graves 24 and 33. The filling of burial 32 consisted of light-gray sandy loam, and its depth from the modern surface was 0.7 meters, while it was 0.1 meters below the mainland. At the bottom, the skeleton of an adult individual was found in a supine position with the head oriented to the northeast. Grave goods included 15 *Corbicula* shells, with fragments of two additional shells discovered in the grave fill. Additionally, a chisel made from the tusk of a mountain goat or ram and bones from an animal no smaller than a ram were present. An AMS date of 3932-3535 calBCE (4914±56 BP, GV-03583) was obtained from human bones. Another radiocarbon determination dates the burial to 3986-3715 calBCE (5090±50 BP, Poz-83434). A combination of probability densities yields a date of 3946-3659 calBCE (5013±38 BP) [R_Combine: (4914±56 BP, GV-03583); (5090±50 BP, Poz-83434)].

*(j)* *burial 32* (Individual IDs I0995, I13100, I13811))

**Burial 33** contains the remains of four adults and one child, all buried in an extended supine position with their heads facing northeast. Three of them were sequenced; unfortunately, we could not find data which makes possible the exact identification of the skeletons within grave 33 corresponding to our genetic sequence.

Skeleton 1 was positioned along the southeastern wall. The lower part of the ulna and radius bones were covered by the pelvic bones, while the hand bones were placed on the femurs. The skull is fragmented. Between the tibia bones, the skeleton of a baby was found. Adjacent to the adult's skull, two large curved fangs from an animal were discovered. Additionally, a necklace made of 16 small incisors of the same animal was found around the shoulder and neck area, as well as on the upper ribs. Skeleton was dated to 3626-3198 calBCE (4649±54 BP, GV-03584).

Skeleton 2 (located near the northwest wall) was positioned close to the center of the grave. The bones of the left hand were extended along the skeleton, while the bones of the right hand were placed on the pelvis. Ten piercings of spadefoot mollusk shells were found around and beneath the skull, along with three fragments and nine intact shells of bivalve mollusks. Furthermore, ten large curved fangs from an animal were present, along with two additional fangs located on both sides of the spine, near the lower ribs. Eighteen small incisors of the same animal were found predominantly near the cervical vertebrae, with some on the right side. Skeleton was dated to 3939-3634 calBCE (4937±56 BP, GV-03585).

Skeletons 3 and 4, situated near the northwest wall of the grave, were damaged during the construction of burial 36. In the case of skeleton 3, the bones of the hands were found on the pelvis, while the skull was missing.

Skeleton 4 was positioned along the northwestern wall, with the skull and humerus absent. The upper part of the chest and spine had shifted toward the northwest. Similar to skeleton 3, the bones of the hands were on the pelvis. Among the ribs, 25 small incisors from the same animal were discovered.

Above the bottom of the grave (at a depth of 0.5 m from the soil surface), fragments of a skull were found, accompanied by seven shells. Additional skull fragments were uncovered at the southwestern wall of the grave, at a depth of 0.25 m, and a broken abrasive tool was found at a depth of 0.35 m.

The sequenced individual ID I0995 was dated to 4220-3801 calBCE (5170±50 BP, Poz-83435), the individual ID I13100 was dated to 3947-3713 calBCE (5030±25 BP, PSUAMS-9043).

For general dating of burial 33, we prefer to use the date from human bones of Skeleton 1 (3626-3198 calBCE (4649±54 BP, GV-03584)) as the only one corresponding with the dates from animal bones found in grave 10 (3517-3358 calBCE (4637±37 BP, GV-03579)). Other radiocarbon dates were pprobably affected by the freshwater reservoir offset. The high level of fish consumption in the Tuzovskiye Bugry Eneolithic population group is supported by stable isotopic values.

*(k) burial 34 (I13099)*

**Burial 34** (individual ID I13099) was situated in the central part of the second "central" row, between burials 4 and 2. The grave pit had an approximately rectangular shape, with dimensions of 1.3 x 0.6 meters along its long axis aligned with the southwest to northeast line. The southwest edge of burial 34 intersected with the northern edge of burial 35. The filling of burial 34 consisted of light-gray sandy loam, and its depth from the modern surface was 1.09 meters, while it was 14 centimeters below the mainland. At the bottom of the grave, the skeleton of an 8-10 year old child was found in a supine position with the head oriented to the northeast. The skull was slightly turned to the left, and the left arm extended along the body, while the right arm was bent at the elbow, indicating that the missing hand should have been positioned on the shoulder. The feet were placed where the skull of the individual buried in burial 35 would have been. Grave goods included lead earrings and two flat-bottomed vessels.

*(l) Unidentified burial (I11405)*

The list of sequenced individuals from Tuzovskiye Bugry-1 includes individual ID I11405, which is associated with **burial 5** based on its skeletal code. However, archaeological attribution places burial 5 in the Early Iron Age, while individual I11405 shows close familial connections to Eneolithic burials at Tuzovskiye Bugry-1. In light of this discrepancy, we propose that a mislabeling may have occurred, and we consider individual I11405 as originating from an Eneolithic-period burial in a grave with an unidentified number.

#### II.C.ii.b) Upper Ob Eneolithic sites (Russia_UpperObKiprino_Eneolithic)

##### (1) Razdum'ye-1

The Razdum'ye-1 site was described earlier (SI Section II.D.ii.a) 1)).

*(a) burial 7 (individual ID I13678)*

**Burial 7** (individual ID I13678) represents an irregular oval pit with an extension in the southwestern half. Its dimensions are approximately 1.55x0.75-1.20 m, and it has a depth of 1.35 m from the level of the modern surface. The grave is oriented along a north-northeast to south-southwest line and likely experienced damage in ancient times.

Near the center of the grave, at the bottom, a skull was discovered, while a tooth belonging to an individual aged 40-45 years old was found near the western wall. In the northeastern part of the grave, close to the wall, the remains of a pillar have been preserved. The cranium exhibits a pale pink coloration.

Behind the cranium, in a pile, various artifacts were found, including a bone harpoon, two bone retouchers, a grinding plate, fragments of a river mollusk shell, bone plates, small animal bones, and stones. Among the stones, a stone sculpture was discovered. Furthermore, a rough arrowhead made of quartzite (?) with a triangular-shaped tang outline was found. Towards the southwestern end of the grave, two grinding plates, three stones, three fragments of a bone plate, a roe deer horn, and a shard of a thick-walled vessel with rough craftsmanship were uncovered. The vessel shard showed traces of surface smoothing using a notched spatula or a bundle of dry grass (Umanskiy 1987). The burial is dated to 3361–3102 calBCE (4525B±20 BP, PSUAMS-9045).

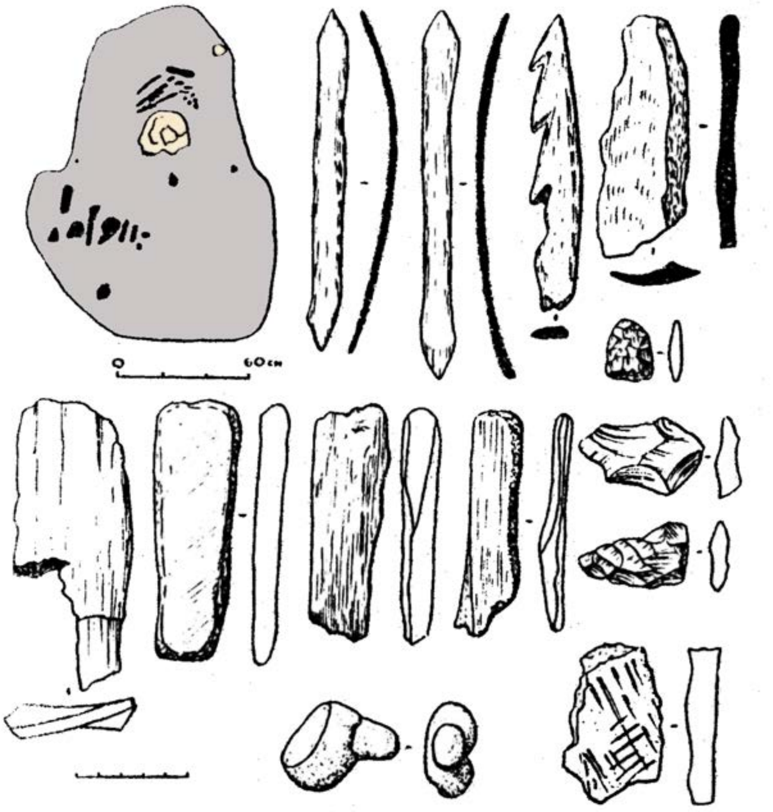

***Fig. 35.*** *Burial 7 at the Razdumye-1 site (Umanskiy 1987, modified).*

##### (2) Ordynskoye-1 burial site

The Ordynskoye-1 burial ground is located on the left bank of the Ob River valley, now the Ob reservoir, on a cape-shaped ledge of the terrace, at the entrance to the Ordynsky Bay. Western part of the site was excavated by M.P. Gryaznov in 1953-1954, and the eastern part was excavated by T.N. Troitskaya in 1973-1977.

The main part of the burials of the burial ground belongs to the Late Krotovo and Irmen. At least three burials were defined by T.N. Troitskaya and V.I. Molodin as an earlier phase of the burial ground (Troitskaya 1973) and classified as the Kiprino stage of the Upper Ob culture (Molodin 1977), attributed by modern researchers to the Eneolithic (Kiryushin 2008).

###### *(a) mound 11, burial 4 (individual ID I11002)*

Burial 4 under mound 11 was partially destroyed as a result of coastal erosion. The depth of the grave was 25 cm. The skeleton lay on its left side, with its head to the northeast. Burial goods are represented by a polished stone plate. The burial was attributed to Kiprino on the basis of the find of similar plates in other early burials on the site, which also contained Kiprino pottery (Troitskaya 1973). The date of the burial is 2907-2704 calBCE (4235±25 BP, PSUAMS-9034).

#### II.C.iii. Middle Irtysh and Baraba forest-steppe

##### II.C.iii.a) Middle Irtysh Neolithic culture (Russia_MiddleIrtyshNeolithic_LN)

###### (1) Omsk Neolithic occupation site

The archaeological complex known as the "Omsk Site" is situated in the former floodplain of the left bank of the Irtysh River within the city of Omsk. The estimated area of this complex spans at least 12 hectares. Excavations conducted at the site have revealed layers dating from the Paleolithic to the Middle Ages, representing different phases of occupation. Archaeological investigations have been ongoing at this site for over a century.

During rescue excavations in 1988-89, Boris Konikov discovered a Neolithic burial ground within the Omsk site. Five burials were unearthed at depths exceeding one meter from the contemporary surface. Three of these burials had experienced significant disturbance, while the other two remained undisturbed. The skeletons were positioned in a supine orientation with their heads facing north-northeast. The discovered artifacts exhibited close parallels with Neolithic sites in the Omsk Irtysh region. Within the cultural layer of the Neolithic settlement at the Omsk Site, a well-preserved dwelling pit was identified. Additionally, a substantial collection of stone tools, including cores, blades, and scrapers, was attributed to the Neolithic period (Metel, 2016; Konikov, 2016).

*(a) The Omsk site, burial 2 (individual ID I1959)*

The female **burial 2** <u>(individual ID I1959)</u> was made in a supine position. Grave goods are represented by two drilled wolverine fangs, two retouched plates, scraper on a flake (Konikov, 2016). The burial is dated to 5612–5381 calBCE (6530B±40 BP, Poz-82200).

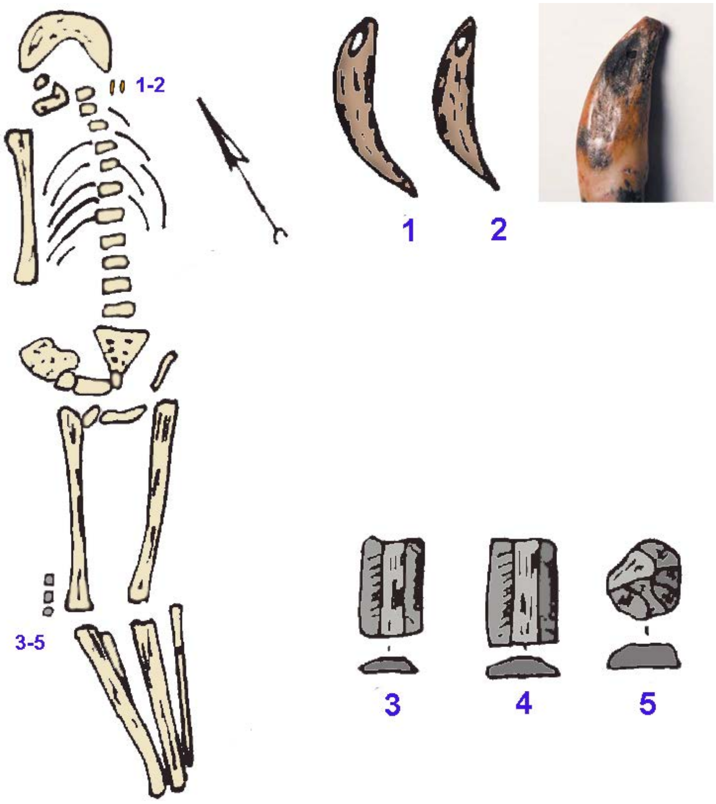

***Fig. S36.*** *Burial 2 at the Omsk site.* ***1, 2*** *- wolverine fangs,* ***3, 4*** *- blades,* ***5*** *- scraper (Konikov, 2016, modified)*

##### (2) Korchugan-1 site

The Korchugan-1 archaeological complex comprises a series of multi-temporal settlement and burial sites, encompassing five distinct phases. These include two burial grounds, namely Korchugan-1a and Korchugan-1b, as well as three settlement complexes known as Korchugan-1c, Korchugan-1d, and Korchugan-1e. Among these components, the Korchugan-1 burial ground is situated along the shores of the oxbow of the Kryuchnoye lake, on the elevated Korchugan terrace remnant. This remnant is characterized by steep banks, rising 2 to 5 meters above the floodplain.

Specifically, Korchugan-1a is a Neolithic soil burial ground, located approximately 8.5 meters from the northern edge of the terraces and 3.5 meters from the western edge. The necropolis consists of three burials, designated as burials 2, 3, and 7, which are arranged in a linear fashion along the NW-SE line. In terms of its spatial arrangement, the Korchugan-1a burial ground holds a central position within the broader complex of the site (Marchenko, 2009).

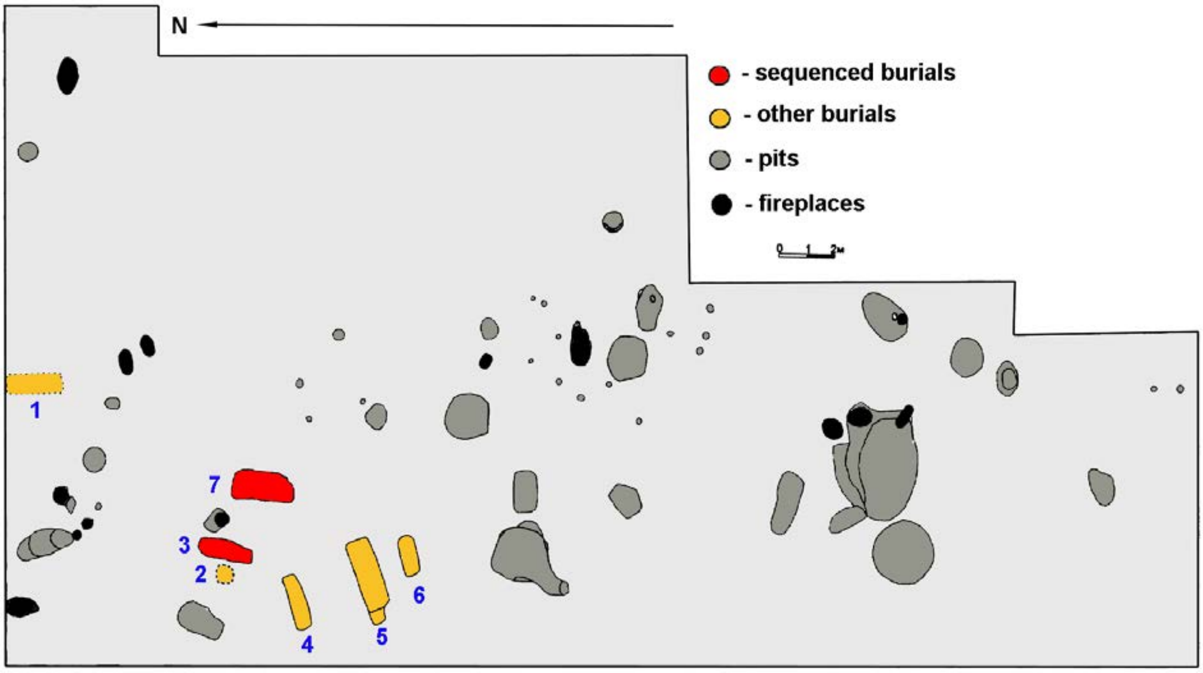

***Fig. S37.*** *The Korchugan cemetery (Marchenko, 2009, modified).*

*(a) burial 3 (individual ID I0991)*

**Burial 3** <u>(individual ID I0991)</u>: The grave pit of Burial 3 has an oval shape, measuring 2.2 x 0.75 x 0.5-0.17 meters, with its longitudinal axis aligned along the NNE-SW direction. The walls of the pit are vertical, and the bottom has a complex shape. Consequently, the deceased individual was positioned in a grave where the upper body was raised. The burial was conducted following the burial rite, with the body extended on the back and oriented towards the NNE. The right arm of the deceased was slightly bent at the elbow, with the hand placed in the pelvic area, while the left arm was positioned under the pelvic bones. A stone axe was placed beneath the skull, and a small stone flake was found near the right hip joint (Marchenko, 2009). The burial is dated to 5206-4799 calBCE (6060±50 BP, Poz-83427).

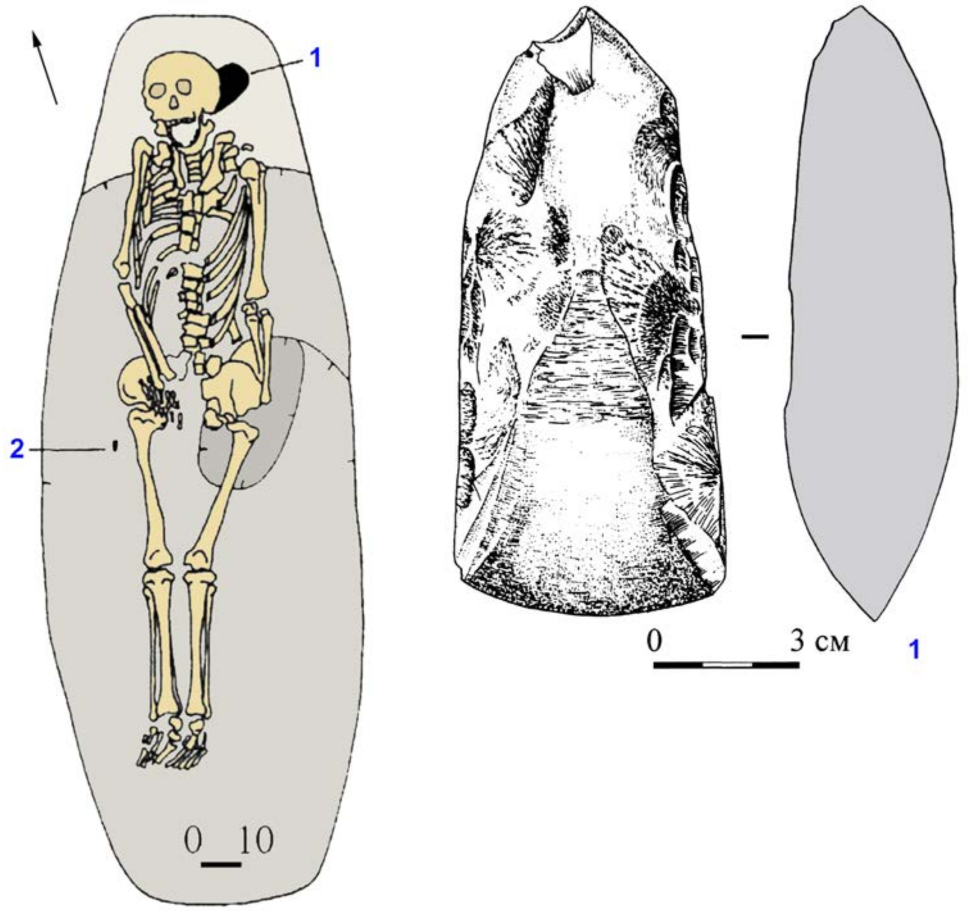

***Fig. S38.*** *Burial 3 at the Korchugan-1 site.* ***1*** *- stone axe,* ***2*** *- flake (Marchenko, 2009, modified).*

*(b) burial 7 (I0992)*

**Burial 7** (individual ID I0992): The grave pit of Burial 7 has an irregular shape, measuring 2.5-1.75 x 0.52 x 0.5-0.4 meters, and its longitudinal axis is aligned in proximity to the north-south direction. The burial was conducted according to the burial rite, with the body extended and the head positioned towards the north. Due to the uneven bottom of the pit, the upper part of the body and the head were elevated compared to the legs. The bones of the hands were connected to the pelvic bones. Notably, pendants made of rodent incisors, possibly from squirrels, were discovered near the right knee. In addition, a necklace consisting of nine miniature bone figurines of birds, crafted in a flat style, was found in the chest area. One of the pendants in the necklace had a drop-shaped bone pendant with a hanging hole, making it the tenth component of the necklace (Marchenko, 2009). The burial is dated to 5002-4727 calBCE (5990±50 BP, Poz-83428).

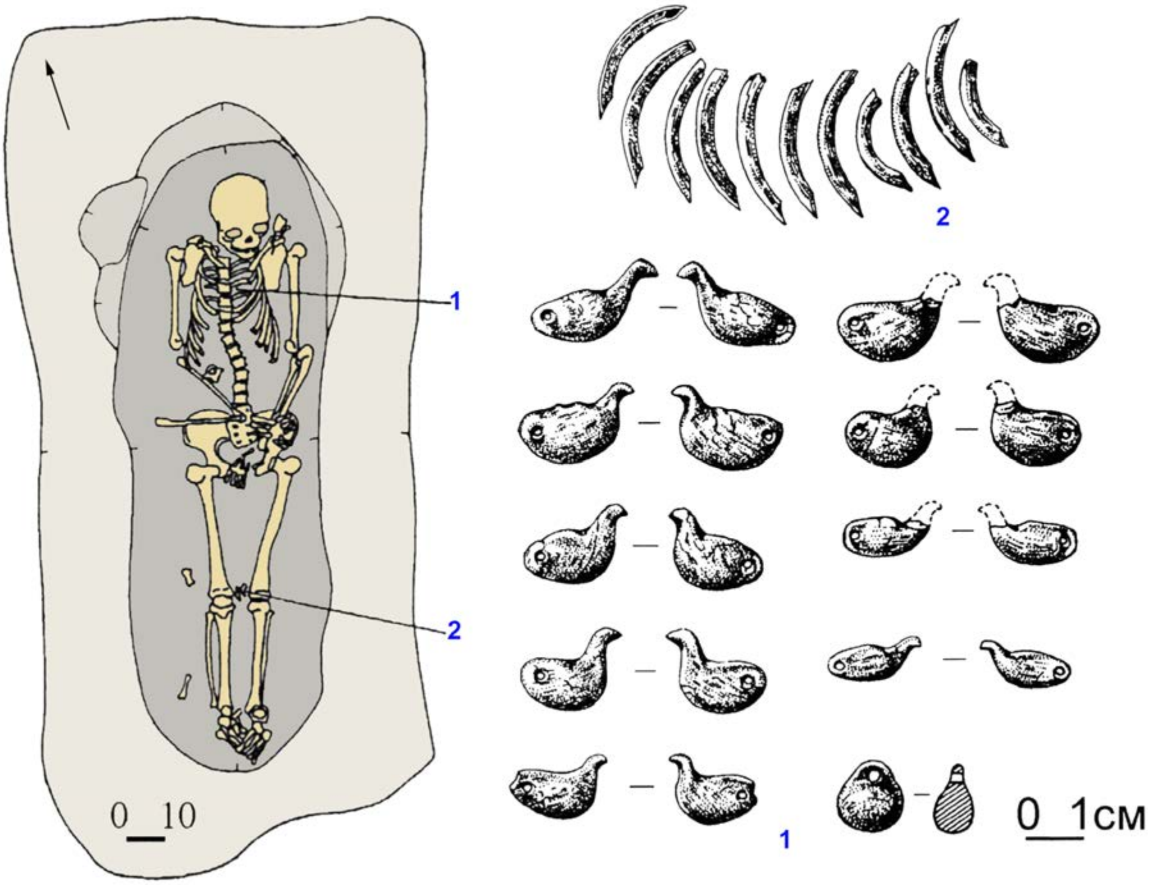

***Fig. S39.*** *Burial 7 at the Korchugan-1 site.* ***1*** *- necklace,* ***2*** *- rodent incisors (Marchenko, 2009, modified).*

##### (3) Protoka site

The Protoka burial ground was initially discovered by Viktor Zakh in 1980. Situated on the northwestern periphery of the Baraba forest-steppe, it encompasses nine burial mounds positioned along the periphery of the Kryuchnoe lake terrace, an oxbow lake formed by the Tara River. In 1986, Natalya Polosmak conducted excavations at the site, focusing on five burial mounds originating from the Bronze and Early Iron Ages. It was during the excavation of mound 5 that Neolithic burials were unexpectedly unearthed.

The burials at Protoka were classified as secondary, as the bones were largely found in disarray, although some elements retained their anatomical position. This arrangement suggests that the composition and distribution of the bone accumulations might be connected to the level of skeletal preservation at the time of interment. The spatial arrangement and characteristics of the bone accumulations indicate that the deceased bodies were likely buried after a significant period of decomposition. In total, 14 burials were identified, containing the remains of 26 individuals, encompassing men, women, and children (Polos'mak et al., 1989).

###### *(a) mound 5, burial 3, individual B (individual ID I7616)*

**Burial 3** (individual ID I7616). The burial comprises the remains of three individuals arranged in a side-by-side position, consisting of two adults and a child aged approximately 7.5-8 years old.

The upper portion of *skeleton B* was preserved in anatomical order down to the pelvic bones. The interred individual, identified as a male between 40-45 years old, was positioned supine with an extended body posture, and the head oriented towards the north-northwest. Adjacent to the right hand of skeleton B, the skeletal remains of a child (skeleton C) were discovered. Notably, fragments of the child's skull (C) were found resting on the left clavicle of skeleton B. Additionally, a polished stone chisel was discovered in close proximity to these remains. Roughly 20 cm away from skull B, a damaged skull (skeleton A) belonging to a man aged 22-26 years was found.

Within the burial context, four pendants crafted from stone and bone were unearthed. These pendants were originally sewn onto clothing and were discovered in the vicinity of the chest area of skeleton B. A whetstone, a fragment of a vessel, and two bone arrowheads were positioned near the waist level of the same individual's spine. Furthermore, a fragment of pottery was found in close proximity to skull A (Polos’mak et al., 1989).

The burial has been dated to 4446–4349 calBCE (5555±20 BP, PSUAMS-4856), providing a chronological context for the interment.

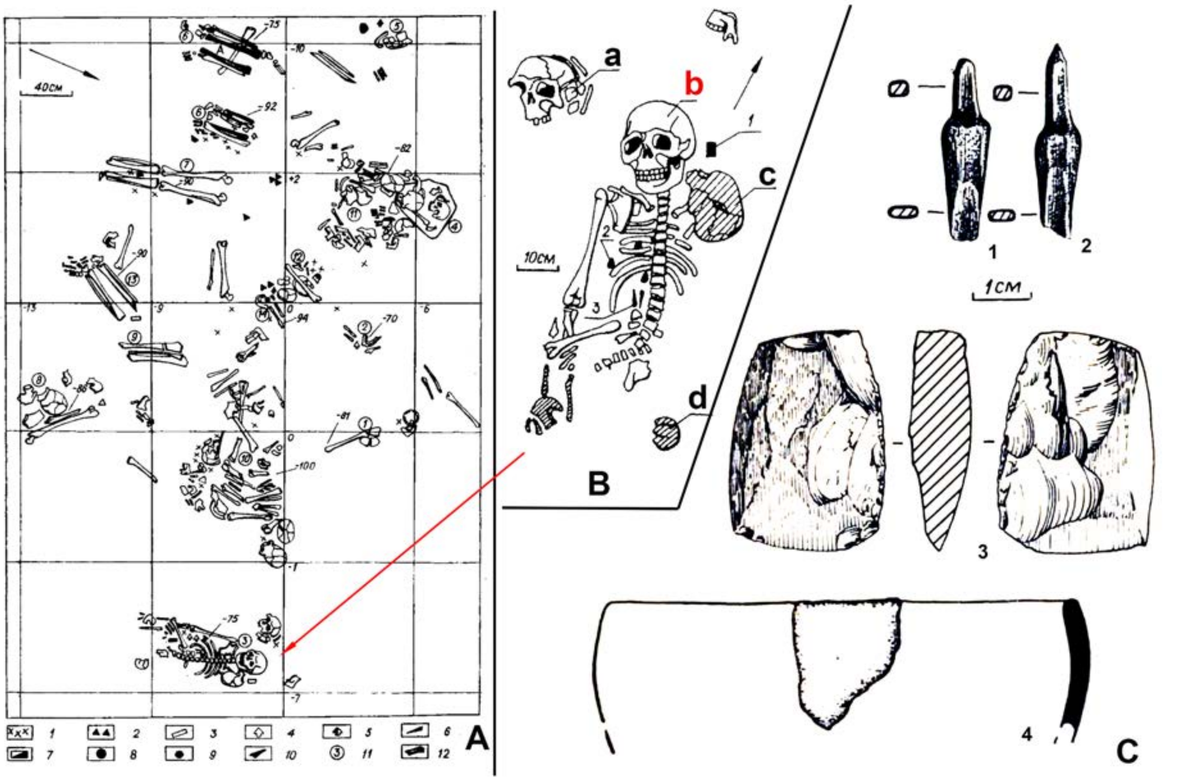

***Fig. S40.*** *The Protoka cemetery.* ***A*** *- Neolithic burials covered by mound 5 (**1** - ceramics; **2** - pendants; **3** - adze; **4** - arrowhead (bone); **5** - arrowhead (stone); **7** - abrasive; **8** - core, **9** - vessel, **10** - a jaw of a dog, **11** - burial number; **12** - bones of the lower horizon; **a** - a vessel; **b** - knife-like plate; **h** - core);* ***B*** *- burial 3 (**1 -** chisel; **2** - pendants; **3** - arrowheads (bone); cross-hatched bones of a child's skeleton);* ***C*** *- grave goods from burial 3 (**1,2** - arrowheads; **3** - chisel; **4** - pottery sherd) (Polos’mak et al., 1989, modified).*

##### II.C.iii.b) Eneolithic sites with Comb-Pit Ware (Russia_CombPitWare_Eneolithic)

###### (1) Eneolithic phase of the Borovyanka-17 site

The Borovyanka-17 burial ground was discovered in 1999, revealing burials spanning various archaeological epochs from the Eneolithic to the Middle Ages. Some of the materials from early burials have been previously published (Khvostov, 2000; 2001). Through archaeological investigations, over twenty burials and pits dating back to the Eneolithic period have been unearthed at the burial ground. The remarkably well-preserved state of the skeletons in most burials allows for a clear identification of the primary burial practices of the Eneolithic population in the forest-steppe zone of the Middle Irtysh region.

Single burials are the most common, while paired and collective burials occur less frequently. The number of individuals interred in collective burials ranges from three to eight. The deceased were typically laid on their backs, often with bent legs, and their heads were oriented towards the northeast, occasionally facing north (burials 81, 82). A notable characteristic of the funeral rite is the use of ocher to fill the burial or surrounding the deceased. In some instances, sizable pieces of ocher were placed near the head of the buried individual (burial 83). The majority of the deceased were buried through inhumation, while evidence of fire in funeral rites is less common, such as partial burning of the skeleton (burial 36), charred planks over the burial entrance pit (burial 83), or embers in the grave filling. The presence of light ash bedding in several burials (burials 12, 19, 36, 46) likely indicates its role in the fire cult. Cremation was observed only in one case (burial 84).

Another distinctive feature of the burial rite is the intentional disturbance of anatomical integrity. For example, in burial 64, containing a double burial, the skulls of the buried children were separated. In tiered burial 36, the tibia bones of the lower skeleton were cut off along with the foot bones and placed on top of the femurs, while in burial 46, the skull was missing. Burial 4 likely contained a special hole for burying the skull adjacent to the skeleton. Additionally, several instances of secondary burials were recorded, where the deceased were dismembered and placed within some form of container before interment. There are also burials of skulls, sometimes arranged in groups (burial 83).

Several burials (burials 42, 45, 67, 72) contained pits filled with fish scales, bones, and gills. The association of these pits with funeral rites is indisputable: in collective burial 72, such a hole was located between the heads of the buried individuals, while in burial 42, it was found near the feet. In burial 67, scattered human bones were discovered alongside intact fish skeletons.

The burial goods found within the burials vary in their abundance, ranging from richly furnished burials (burials 14, 36) to interments with no burial goods. Deliberate destruction of items is a characteristic practice. Miniature ceramic vessels were recovered from two burials (burials 12, 66).

For the dating of the Eneolithic phase of the Borovyanka-17 cemetery, we use seven radiocarbon dates, obtained from human bones. Based on them and taking into account the δ15C/δ15N ratios observed in the samples from the site, we suggest two sub-phases in the Borovyanka-17 Eneolithic cemetery, the earlier one, dated to 3700-3300 calBCE, and the later one, which is attributed to 2900-2800 calBCE.

*(a) burial 4 (individual ID I6962)*

**Burial 4** (individual ID I6962) was identified as an oval spot measuring 220x90 cm, characterized by gray-brown sandy soil and elongated in the southwest-northeast direction. Another dark gray sandy loam area was observed on the south side (burial 15). Excavation revealed a burial pit with elongated proportions and rounded corners, measuring approximately 230x95-100 cm, oriented in the north-northeast-south-southwest direction. The northeastern part of the burial had been disturbed by a badger's hole, resulting in the destruction of the upper portion of the skeleton. The southwestern and central parts of the burial had a flat bottom and sheer walls, with a depth of 20-22 cm from the subsoil level (43-45 cm from the present surface). The northeastern part of the burial was deeper, with a hole depth reaching 40-44 cm from the subsoil level (60-65 cm from the present surface). The lower part of the skeleton remained undisturbed, and the bones were well-preserved. The deceased individual was buried in a supine position with bent legs, and the head was oriented towards the northeast. The leg bones were shifted to the left of the spinal line. The left arm extended alongside the body, while the right arm was slightly bent at the elbow, with the hands positioned at the pelvis. The leg bones exhibited slight staining with ocher, and ocher also covered the bottom of the grave pit. The skull had been displaced and was found within the fill of the badger hole, approximately 10-15 cm north of the grave pit boundary.

The grave goods discovered in this burial consisted of 56 well-preserved bone drop-shaped pendants. All the pendants were concentrated in the groin area of the deceased, with the highest concentration observed between the femurs and within the pelvic cavity.

The burial is dated to 4045–3819 calBCE (5160±25 BP, PSUAMS-4841).

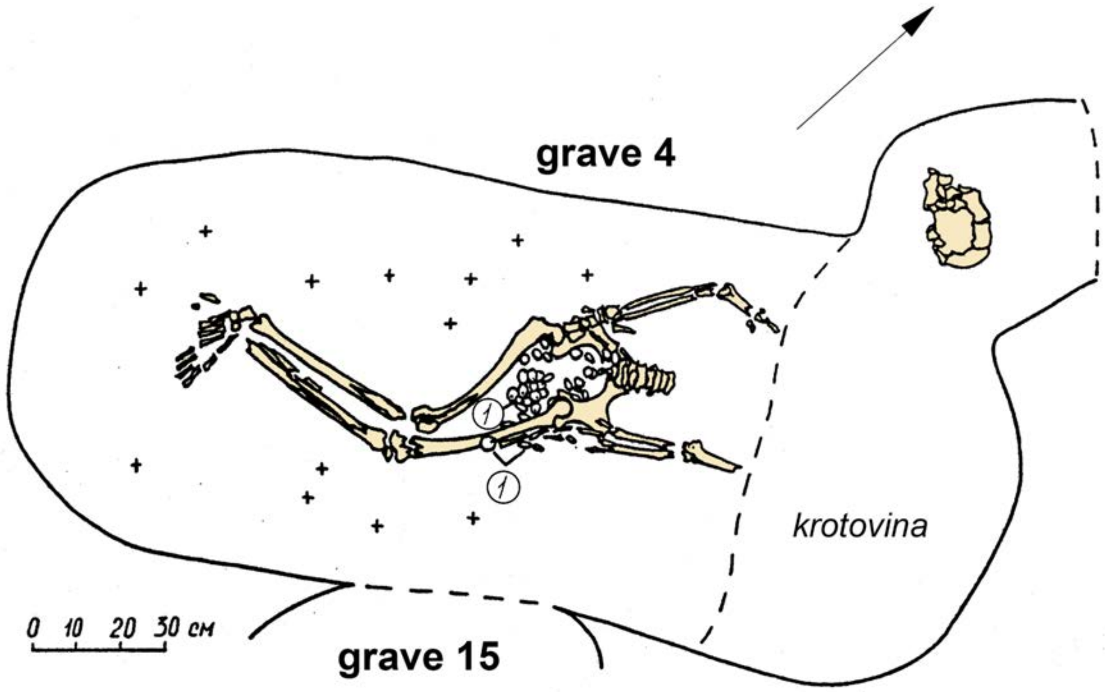

***Fig. S41.*** *Burial 4 at Borovyanka-17.*

*(b) burials 36C (Individual ID I6963) and 42 (Individual ID I6831 & I6999)*

Burials 42 (individual ID I6831 & individual ID I6999) and 36C (individual ID I6963) were was were initially identified as a dark gray amorphous sandy loam area at the subsoil level. Within the infill, two rims of the Comb-Pit pottery, three flakes, and a stone scraper were discovered. The large grave pit, measuring up to 465x185-190 cm, was divided into two parts: the northeastern section (burial 36) and the southwestern section (burial 42).

Burial 36C (individual ID I6963). Upon the complete excavation of burial 36, it became evident that traces of light-gray ash sandy loam were present at the bottom of the grave. Further excavation of this layer revealed the presence of another skeleton (burial 36C). As a result, burial 36 was identified as a tiered burial.

The skeleton in burial 36C was situated in a pit measuring 225x73-78 cm, aligned along the NNE-SSW axis, and covered by a thick layer of light ash sandy loam. The skeleton (individual ID I6963) was well-preserved and positioned on its back, with the head facing north-northeast. The tibias were intentionally separated and placed alongside the femur. Based on the vertical positioning of the shoulder blades, it appears that the deceased was tightly wrapped or bound. The leg bones, arm bones, and skull showed signs of burning. A significant number of bone beads (several hundred pieces) were discovered in the chest and belt areas. The largest accumulation of beads was found in the chest region. Additionally, petal-shaped bone pendants (six specimens) were located to the right of the pelvis (near the right femur). Two bone beads were found between the femurs and under the pelvis. Pendants made of hare incisors were also present.

After thorough cleaning of the tiered burial, the depth of the grave pit in the subsoil was determined to be 84-88 cm (116-120 cm from the surface).

The burial is dated to 4320-4052 calBCE (5340±30 BP, PSUAMS-3929).

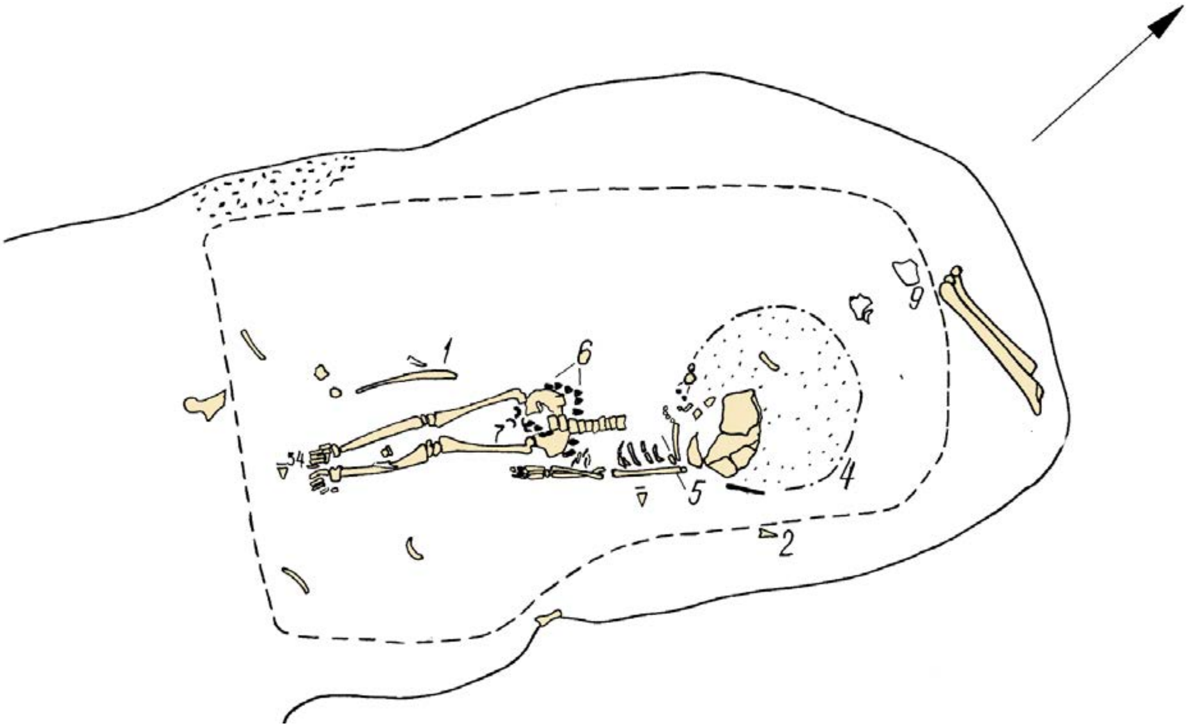

*Fig. S42.* *Burial 36C at Borovyanka-17.*

**Burial 42** (individual IDs I6831 & individual ID I6999). The amorphous grave pit 42 extended along the NE-SW axis, measuring 270x160 (190) cm. Its walls exhibited sheer profiles, and the bottom gently sloped lower towards the NE. The pit's depth from the surface ranged from 47 to 53 cm. Unfortunately, the skeletal remains of the two interred individuals (A and B) suffered severe damage due to badger burrowing and plowing. The south-western part of the grave was destroyed by burial 62, which is attributed to the Late Krotovo culture based on details of burial rites and presence of bronze artifacts.

***Skeleton A*** from grave 42 was situated in the northwestern area of the pit. The preserved elements included the skull and upper chest components (collarbone, portions of ribs, and humerus). Based on the extant skeletal fragments, the deceased was interred supine, with the head oriented to the NE. Notably, a slate arrowhead with a notch (1) was uncovered 20 cm southeast from the buried individual's skull. An animal bone was also discovered 10 cm northwest from the right humerus (3).

Adjacent to the southeastern wall of the grave, the remains of ***skeleton B*** were encountered, albeit highly deteriorated. Solely the pelvic bones, fragments of tubular bones, and small, unidentifiable bones were preserved. Some of these bones lay on the surface or slightly above its level. The positioning of the pelvic bones suggests that the deceased was laid to rest supine, with the head directed to the northeast. No accompanying artifacts were found in association with this individual. Notably, a hole measuring 30x15 cm was observed 20 cm northwest of the pelvis, containing fish scales within a layer approximately 10-15 cm thick.

One of the two skeletons from grave 42 was sequenced (individual ID I6831) and dated to 3985-3804 calBCE (5125B±25 BP, PSUAMS-4829).

Another sequenced individual (individual ID I6999), according to its label, comes from burial 62, which cuts burial 42 and contains Late Krotovo artifacts. However, the radiocarbon date of individual ID I6999, 3620-3375 calBCE (4710B±20 BP, PSUAMS-4340), makes impossible its Late Krotovo attribution. We assume that the dated and sequenced individual labeled as burial 62 originally came from the Eneolithic burial 42 and was displaced to a later context by pedoturbation processes. Noticeably, this individual demonstrates the lowest value of δ15N and the highest value of δ15C, which makes the radiocarbon dates obtained from the human bones of this individual the most reliable.

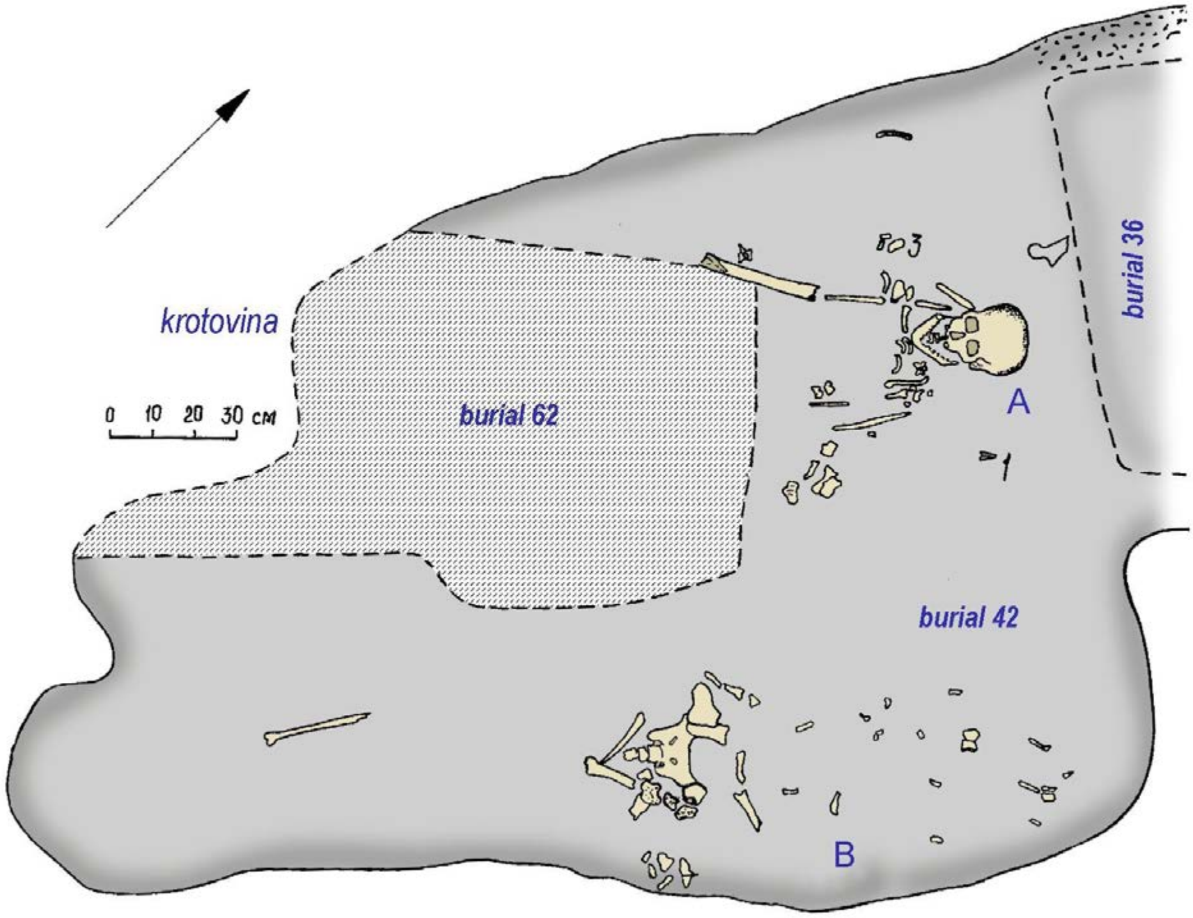

*Fig. **S43**. Burial 42 at Borovyanka-17.*

*(c) burials 66C (Individual ID I6951)*

**Burial 66** (individual ID I6951) was identified as an oval-shaped spot of dark-gray sandy loam, slightly narrowed in the northeastern part, measuring 220x120-140 cm. It was oriented along the northeast-southwest line. Within the spot's infill, two fragments of Comb-Pit pottery, a sherd of a Chernoozerye-type vessel, and a substantial amount of charcoal were discovered. The grave pit itself had an oval shape, measuring 220x110-135 cm and was oriented along the northeastNE-southwest line. The depth of the grave pit in the subsoil was 33-37 cm, equivalent to 55-60 cm from the modern surface. The walls were inclined, and the bottom had a slight drop towards the northeast, with a difference of approximately 5-8 cm.

In the western corner of the grave, a dense cluster of human bones was found. Upon further excavation, it was revealed to contain the bones of 7-8 individuals showing signs of deliberate dismemberment, including partially arranged spines and evidence of bone damage. Near the cluster, slightly to the northeast, the remains of two poorly preserved human skulls and a femur were discovered. The sequenced individual ID I6951 comes from skull labelled “C” (Cyrillic “B”). Additionally, poorly preserved fragments of a human skull and a tubular bone (possibly a femur) were found in the northeastern part of the grave, just above the surface level. Fragments of burnt wood and an accumulation of small charcoals were recorded in the southern corner of the grave.

Based on the condition of the remains, it appears that the deceased were first dismembered and then, after some time, buried in some form of container, possibly a sack. The skulls were placed near the bones. In the western corner of the grave, adjacent to a cluster of bones, a stone flake was found. Beneath the bones, a small pottery vessel with a sharp bottom and with jar-shaped form was discovered. Fragments of Comb-Pit pottery were found in the southern and northern corners of the grave, while a scraper was located at the bottom of the grave near the skull in the northeastern part.

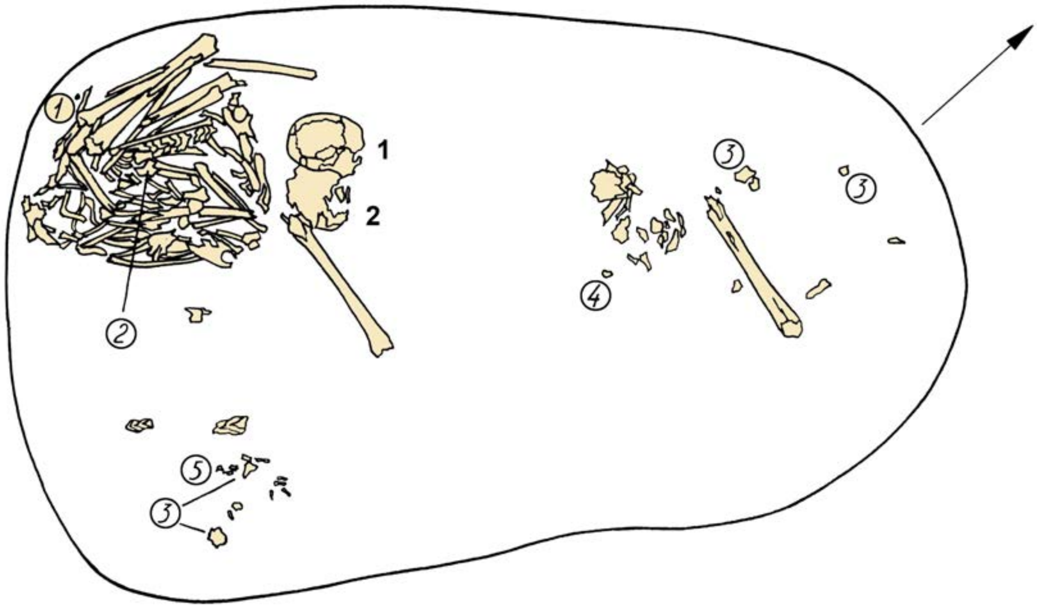

***Fig. S44.*** *Burial 66 at Borovyanka-17.*

*(d) burial 83 (skull F – Individual ID I6829, skull D - Individual ID I6828)*

**Burial 83** (individual ID I6828, individual ID I6829) was identified as an elongated spot of black humus sandy loam, elongated along the northeast-southwest line, expanding towards the southwestern part, with dimensions of 355-365x140-220 cm. An accumulation of fish scales, measuring 40x25 cm and 3-5 cm thick, was found at the southeastern side of the wall. The filling of the grave contained a fragment of Comb-Pit pottery, a knife-shaped piece, and numerous small charcoal fragments. The bottom of the grave was flat, with a slight rise towards the southwestern part, while the walls were slightly inclined and nearly sheer. A ledge was observed in the western corner, and remains of a burnt board, measuring 95 cm in length and 3-10 cm in width, were discovered near the southeastern side.

A total of eight individuals were found in the burial, including three postcranial skeletons (1-3) and five skulls (A-H). Skull F belonged to sleleton 2, and skull G belonged to skeleton 1. Skeletons 1 and 2 were laid in supine positions, while skeleton 3 consisted of a cluster of human bones intentionally broken and chopped.

*Skeleton 2*, found in the northeastern part of the grave, was in a supine position with the head facing northeast. The skull (*scull F*, “E” in Cyrillic) (individual ID I6829), left femur, tibia, and partial foot bones were well-preserved. Traces of ocher, up to 3 cm thick, were observed around the skull, pelvis, and ankle bones.

The *skulls D* (individual ID I6828) and E (“Г” and “Д” in Cyrillic) were excavated at the northwestern wall of the grave. They were positioned on their bases.

The burial is dated to 4214-3958 calBCE (5210В±30 BP, PSUAMS-3920), based on radiocarbon date of skull F.

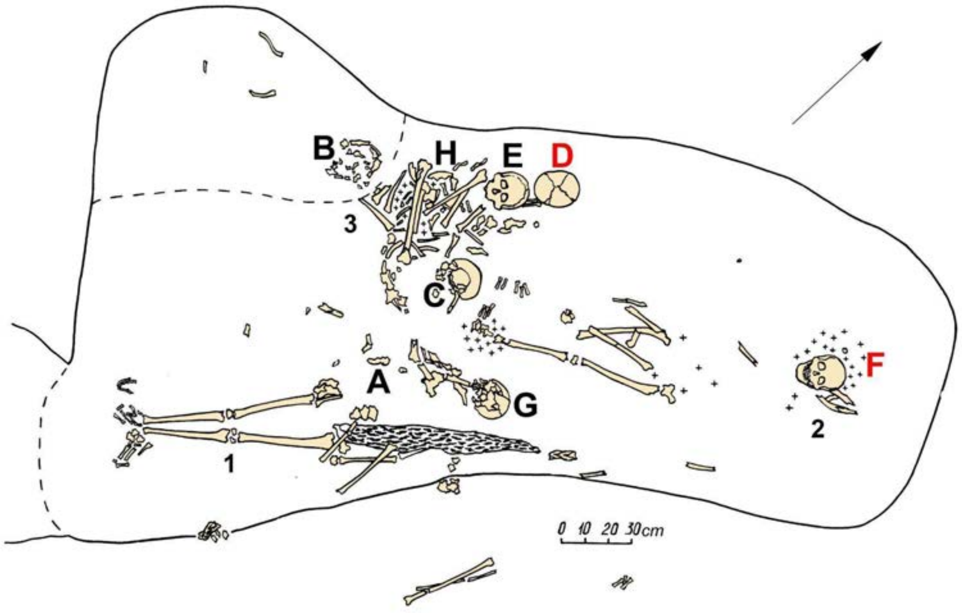

***Fig*. *S45***. *Burial 83 at Borovyanka-17. Sequenced skulls D and F are labeled in red.*

*(e) unidentified burials (I6827, I6830, I6959)*

Three skeletons from Borovyanka-17 lack burial numbers. Individual ID I6830 is dated to 2911-2881 calBCE (4270±20 BP, PSUAMS-4828), ID I6827 - to 3937-3654 calBCE (4990±30 BP, PSUAMS-3919).

The individual ID I6830 is dated to a later period than all other skeletons from Borovyanka-17. Noticeably, it has a substantially different ratio of δ15N/δ15C from other burials on the site, which may reflect differences in consumption and economy.

**(2) Eneolithic phase of the Okunevo site complex**

The archaeological site complex located in the Tatarsky Uval (ridge) area, collectively known as "Okunevo," was investigated by V.A. Mogilnikov, B.A. Konikov, A.I. Petrov, and V.I. Matyushchenko. Excavations were conducted in various areas, identifying separate sites within the complex: the Okunevo-3, 3a, 4 cemeteries, the fortified settlement 5, and the cemeteries 7, 7a.

**Okunevo-3**, situated in the southern part of the Tatar ridge on a small promontory with a height of 10 meters, was studied by V.A. Mogilnikov (1972) and B.A. Konikov (1976-1977). The graves of the Neolithic, Early Bronze Age, Early Iron Age, and Middle Ages have been investigated at this site.

**Okunevo-3a** is a group of three burial mounds located between the Okunevo-3 burial ground (to the south) and the Sargat culture settlement Okunevo-9b, which dates back to the 1st millennium BCE. Excavations were conducted by V.A. Mogilnikov (1972) and V.I. Matyushchenko (1997).

**Okunevo-4** is a kurgan cemetery consisting of numerous small mounds, adjacent to the southern side of the Okunevo-3. Burial mounds investigated at this site yielded materials from the 16th-17th centuries, associated with the southern Khanty population.

Occupying a promontory 10 meters high above the floodplain at the northern end of the Tatar ridge, **Okunevo-5** was studied by B.A. Konikov. The site revealed materials from the Stone Age to the Middle Ages, including a Bronze Age burial ground and a transitional sacrificial site from the Bronze to Iron Ages (Early Iron Age). It encompasses a settlement, a potential Early Bronze Age settlement, and a potential cult site from the Early Iron Age.

**Okunevo-7** (including its western edge, Okunevo-7a) is situated in the northern and central parts of the Tatarsky Uval, south of Okunevo-5. It was excavated by A.I. Petrov (1980), V.A. Mogilnikov (1981), and V.I. Matyushchenko (1985, 1987-1991). Described as an flat burial site, it contains burials spanning the late Neolithic to the late Middle Ages. The estimated area of the necropolis is 1000 square meters, with approximately one-third explored.

The Neolithic to Early Bronze Age burials are dispersed among the Okunevo-3, 5, and 7 sites, which can be considered part of a single necropolis (Topleko, 2015; Bolshanik et al., 2001).

We sequenced 11 individuals from the Neolithic-Early Bronze Age burials at Okunevo. The chronological attribution of the site is based on 6 newly obtained radiocarbon dates, which fall into two periods. The earliest date was obtained from burial 189, attributing the initial subphase of the cemetery to 4228-3979 calBCE. Other burials belong to later subphase, which spans from 3089 to 2784 calBCE. The chronological differences are reflected in burial rites and artifact assemblage of the studied burials. The earliest grave 189 contained a secondary burial with traces of postmortem decomposition of the skeletons, while the later graves yielded bronze artifacts and pottery.

*(a) burial 47B (individual ID I6846)*

Grave 47, oriented in a north-south direction, was identified on the ground as a subrectangular feature measuring 295 x 100 cm, with a depression of 10-15 cm observed on the western side. The feature was filled with dark, highly humus-rich sand. Within the southwestern corner of the burial pit, two distinct shelves were identified at depths of 35 cm and 50 cm below the surface, respectively. Along the southern wall, another 10 cm wide shelf was observed, situated 30 cm below the ground level. This burial feature yielded the discovery of two interments.

Burial B, also oriented along the north-south axis, exhibited an elongated subrectangular shape and was situated at a depth of 60 cm. Its dimensions measured 295 x 100 cm. At the southern extremity of the grave, a conspicuous ocher deposit, measuring 15 x 20 cm with a maximum thickness of 5-7 cm, was prominently evident below the ceiling. A similar ocher deposit, measuring 25 x 15 cm, was also identified at the northern end of the grave, positioned to the left of the cranial remains. The ceiling structure was preserved in the southern half of the grave, occupying an area measuring 170 x 100 cm. The ceiling was constructed from longitudinally arranged blocks, measuring 15-20 cm in width and up to 10 cm in thickness.

Upon removal of the ceiling, the skeletal remains of the primary individual were uncovered, positioned supine and fully extended, with the head oriented to the north. Unfortunately, the skeletal preservation was notably poor, with the skull fragmented into numerous pieces, and the bones of the thoracic and lumbar regions of the skeleton in a deteriorated state. In the southern portion of the grave, beneath the ceiling, poorly preserved fragments of a child's cranial remains were also encountered. Additionally, fragments of a bone ring and an asymmetric stone knife were recovered from this layer within the grave. The grave fill yielded 25 fragments from three to five vessels adorned with Comb-Pit and Krotovo culture motifs (Matyuschenko, Polevodov 1994).

*(b) burial 59 (individual ID I6834)*

Burial 59 was situated within the upper subsoil layer at a depth of 40 cm from the surface, with approximate dimensions of 180 x 50 cm. It was oriented along a north-south axis, with a slight deviation towards the northeast and southwest. Within this grave, an articulated skeleton was unearthed, positioned in an extended supine orientation with the head directed to the north. The preservation of the remains was limited to the following elements: the skull (in a state of poor condition), several bones of the upper thoracic region, the left humerus, both ulnas, the lower portion of the vertebral column, the pelvis, and the larger bones of the lower extremities.

Adjacent to the head, positioned on the right side, was the poorly preserved lower fragment of a flat-bottomed vessel adorned with vertical zones featuring comb stamp decorations. Additionally, a copper plate torque was noted to be placed around the neck of the deceased individual. Fragments of ceramic artifacts were recovered from the fill material within the grave (Matyuschenko, Polevodov 1994).

Burial 59 is dated to 2909-2784 calBCE (4250±20 BP, PSUAMS-4831).

*(c) burial 62 (individual ID I6847)*

Burial 62, with approximate dimensions of 170 x 75 cm, was oriented along a north-south axis. Within the northern half of the grave pit, the remains of a skeleton were preserved, represented primarily by the skull, facing southward. To the north of the skull, the remains of tubular bones were discovered, albeit in a poorly preserved state. Surrounding the skull and bones, a discernible darker humus stain was observed.

Located to the south of the skull, a notable cluster consisting of 40 ring-shaped horn beads and a spectacle-shaped bronze pendant was unearthed. At the southern terminus of the grave, a clay tetrahedral flat-bottomed vessel was uncovered, accompanied by a bronze tetrahedral awl. The vessel in question exhibits characteristics of a squat closed jar, characterized by a swollen body. It features vertical lines of stick impressions that divide the middle portion of the body, delineated by double horizontal lines of oblique short impressions. The bottom of the vessel is adorned with radially arranged imprints (Matyuschenko, Polevodov 1994).

Burial 62 is dated to 3082-2910 calBCE (4370±25 BP, PSUAMS-3923).

*(d) burial 65 (individual ID I6950)*

Burial 65 was identified as an elongated oval area with light humified soil containing numerous ocher inclusions. The dimensions of the grave pit measure 265 x 70 cm, with a depth of 90 cm, and it is aligned along a north-south axis.

Within the grave, a skeleton was discovered, positioned in an extended supine posture with the head oriented toward the north. The left hand of the individual rested upon the pubis, while the right hand was placed adjacent to the right thigh. Notably, the feet were absent, and the condition of the skull was severely compromised, having been crushed. The lower jaw was located to the left of the skull.

The entire expanse of the grave exhibited a layer of ocher, with a more concentrated presence observed around the areas corresponding to the legs and head. Adjacent to the right hand of the deceased, a stone knife was recovered (Matyuschenko, Polevodov 1994).

*(e) burial 69 (individual ID I6835)*

Burial 69 was uncovered at a considerable depth of 150 cm beneath the surface. The grave pit exhibited distinct characteristics, manifesting as a well-defined rectangular shape measuring 220 x 90 cm, with a depth of 80 cm. Notably, a significant portion of the skeleton was concealed beneath a substantial layer of large stone blocks. The stone slabs employed in the grave's construction measured approximately 25 cm in thickness, though some regions exhibited thinner dimensions, ranging between 10-15 cm. Their width varied between 10-20 cm.

Within the confines of the grave, a relatively well-preserved skeleton was situated in a supine position with its head oriented toward the north. To enhance the burial, a fine layer of ocher had been meticulously distributed across the entirety of the bone assemblage. In proximity to the tibia bones, the remains of an infant were discovered, though they were severely deteriorated. Alongside this infant burial, a large quartzite side-scraper, exhibiting signs of double-sided craftsmanship, was situated by the right hand. Positioned beside the left hand, another side-scraper crafted from the same quartzite material was found. Additionally, the grave yielded a clay sphere and two fragments of early ceramics (Matyuschenko, Polevodov 1994).

*(f) burial 79 (individual ID I6837)*

Burial 79 exhibited a pronounced elongated shape, extending from the south to the north, with rounded corners. Its dimensions measured 385 x 95 cm, and the depth reached 35 cm. The primary component, situated to the north, encompassed a standard cavity measuring 290 cm in length. Adjacent to this segment, on the southern side, lay a relatively smaller pit, measuring 100 cm in length and 25 cm in depth.

The principal grave contained the mortal remains of an individual who had been interred in a supine position with the head oriented to the north. Remaining from the skeletal structure were remnants of a fragmented skull, segments of both larger ulnas, minor fragments of the pelvis, femurs, and a solitary left fibula. Traces of ocher were detected in the vicinity of the skull and around the abdominal area.

The deceased had been furnished with a suite of accompanying artifacts. To the northwest of the skull, a stone flake and a fragment of a stone axe were situated. Positioned on the eastern side of the skull lay a stone whetstone, two pieces of quartzite, placed one atop the other, bearing a striking resemblance to rock crystal fragments, along with a diminutive pebble. Adjacent to the right temple of the deceased, a second comparable pebble was discovered, alongside an artifact crafted from a branch of a petrified tree infused with iron oxide. The latter artifact took on the form of an elongated, mildly curved rod, with one side potentially displaying deliberate modeling. Its appearance bore resemblance to an animal's visage: it featured a protrusion extending from the forehead to a muzzle-like structure, with a slightly projecting lower jaw. On the left side, there existed a longitudinal groove demarcating the mouth. The remaining portion of the artifact exhibited significant deformation, likely attributable to natural fracturing. The rod measured 14.5 cm in length and 1.8 cm in width. Interposed between these items were a scraper, a chert fragment, and assorted flakes. Notably, in the southern segment of the grave, closer to its northern extremity, a cluster of human teeth was observed within a 30 x 20 cm ocher deposit (Matyuschenko, Polevodov 1994).

*(g) burial 168 (individual ID I6833)*

Burial 168, aligned along a north-south axis, possessed dimensions measuring 230 x 85 cm and reached a depth of 10 cm. This burial contained a fragmented and poorly preserved skull, alongside the remains of the left humerus, both femurs, and the left tibia. The deceased was interred in a supine position, fully extended, with the head oriented towards the north.

Within the confines of the grave, several artifacts were uncovered, including a stone semi-disk situated in the pelvic region and a bronze ring positioned near the right ulna. The semi-disk, crafted from limestone slabs and displaying significant damage along its entire periphery, exhibited a semi-lunar configuration, with one side appearing smooth and the other rounded. Notably, a central aperture was present within the arc, likely intended for suspension or attachment purposes (Matyuschenko, Polevodov 1994).

The burial is dated to 3089-2917 calBCE (4385±20 BP, PSUAMS-4830).

*(h) burial 172 (individual ID I6964)*

The precise delineations of this grave could not be ascertained. The mortal remains of the individual were discovered approximately 15 cm above the subsoil level, positioned in a supine orientation with the head oriented toward the north. Some fragments of the skull, along with a portion of the right humerus and damaged femurs, have survived. The right femur, a fragment of a tibia, and assorted bone fragments were recovered from the western portion of the grave. Notably, a fragment of a bone ring was located amid the leg bones.

In proximity to the burial, an pottery vessel was unearthed. This vessel, characterized as a flat-bottomed, partially enclosed jar with a slightly everted rim featuring a protrusion along the edge. The ornamentation comprised two horizontal bands embellished with broad-toothed comb impressions, interspersed with vertical lines created by comb impressions. Between these pairs of vertical lines, oblique comb hatching was applied. Additionally, a horizontal herringbone pattern, produced using the same comb impressions, adorned the central region of the vessel's body. The vessel's base exhibited a random arrangement of comb impressions and perforations (Matyuschenko, Polevodov 1994).

Burial 172 is dated to 3013-2902 calBCE (4340±20 BP, PSUAMS-4850)

*(i) burial 180 (individual ID I6836)*

The precise boundaries of the grave enclosure eluded identification; therefore, its estimated dimensions were roughly 110 x 55 cm, with a depth of approximately 10 cm. Intriguingly, it was revealed that this particular grave served as an overlay for another grave, designated as Grave 181.

Within this grave, a juvenile interment was discovered, arranged in a lateral recumbent position with the body fully extended and oriented in a northerly direction. Notably, the preservation of the remains was limited to the left side of the damaged cranium, fragments of the vertebral column, and portions of the femur.

Remarkably, the grave assemblage encompassed vestiges of bronze artifacts, including a distinctive bronze hryvnia fashioned in the form of a curved plate. This hryvnia was recovered in close proximity to the cranium, positioned around the neck of the interred individual (Matyuschenko, Polevodov 1994).

*(j) burial 189 (individual ID I6848)*

Burial 189, oriented along the NE-SW axis, exhibited an elongated configuration, extending beyond the confines of the excavation squares. In the southern portion, the spot's width measured between 150-170 cm, while in the northern half, it ranged from 110-120 cm. Excavation efforts unveiled that this location had once accommodated a trench roughly 110 cm wide and 75 cm deep below the subsoil level. Adjacent to the ditch's western margin, on the southern side, small depressions in the subsoil, each measuring up to 35 cm in diameter, were observed.

The contents of the ditch consisted of two distinct horizons. The uppermost layer, approximately 60 cm thick, comprised humified soil devoid of any accompanying objects. Below this, a lower layer of dark humus-rich soil was encountered, with a minimal thickness that transitioned into humified sandy loam. The ditch's base remained uniformly level.

Notably, at the northern extremity of the ditch, situated 120 cm from the excavation's northern wall, a sudden drop in the floor's elevation, by 25 cm, formed a pit measuring 110 x 120 cm in size. This depression reached a depth of 65 cm below the ditch floor and 120 cm relative to the upper horizon of the mainland. Intriguingly, this pit was replete with compact humus-laden sandy loam, yet it yielded no associated objects of archaeological significance.

Within the confines of the ditch, two interments were discerned, albeit slightly displaced to the southeast. These interments are referred to as the "southwestern" (1) and "northeastern" (2) individuals.

Individual 1 (ID I6848), positioned in the southwestern portion, appeared to have been interred in a supine posture. The cranium, severely crushed and flattened, was oriented towards the right. Notably, the right shoulder and forearm bones were conspicuously absent. The right thigh exhibited a slight displacement to the right, with the bones of the right shin situated beneath the pelvis and adjacent to the right thigh's head. Remarkably, the bones of the right foot were positioned near the thoracic ribs, suggesting that the right leg may have been flexed at the knee, placing the foot in the vicinity of the buttocks. In contrast, the left thigh displayed a displacement to the left, with its head positioned beneath the pelvis. The left tibia overlapped the lower extremity of the left thigh. Notably, the left leg may have initially been inverted, and the phalanges of the left foot and other bones were located slightly to the left. The arrangement hints at the possibility that the corpse may have been bound before interment. Adjacent to the skull, a petal-shaped pendant was discovered.

Individual 2, presumably interred prior to the first, exhibited an unconventional resting position. The cranium was found in an upward-facing orientation. To the south of the skull, the bones of the shoulder and forearm were situated, while to the west, the femurs, lacking their heads, were discovered, with the joints oriented towards the skull. Additionally, the right thigh was superimposed over the left, and below and adjacent to the hip bones, the pelvic bones were found in a disarrayed state, with the right pelvic bone occupying the position of the left, beneath the thigh bones, and vice versa. Phalanges from this interment extended beneath the skull of the first skeleton.

The configuration of the second skeleton may be indicative of a secondary burial event, occurring subsequent to the disruption of ligaments, resulting in the bones being arranged without adherence to a specific order (Matyuschenko, Polevodov 1994).

Burial 189, individal 1 has been dated to 4228-3979 calBCE (5250±30 BP, PSUAMS-3924), demonstrating the earliest date from all radiocarbonally dated skeletons at the cemetery.

*(k) burial 222 (individual ID I6839)*

Burial 222, oriented along the NE-SW axis, exhibited dimensions of 85 x 240 with a depth of 40 cm, presenting a subrectangular outline. It's important to note that the north-eastern wall of the grave has sustained damage and is denoted in the record as a conventional marking. Approximately 30 cm above the ground surface, a layer of calcined soil, measuring 20 cm in thickness, was uncovered in close proximity to the grave.

Within the confines of the grave, poorly preserved remains were discerned, comprising a skull, leg bones, and several vertebrae. Notably, the skull was situated approximately 30 cm above the bottom of the grave, whereas the remaining skeletal elements were positioned at the base. Intriguingly, the skull had become traversed by tree roots, potentially suggesting that these roots elevated the skull to its higher position within the grave. No funerary objects or artifacts were recovered from this burial pit.

Burial 222 hs been dated to 3011-2890 calBCE (4320±25 BP, PSUAMS-3921).

**(3) Ostrov-2 site**

*(a) burial Ostrov-2/1 (I2144)*

From the Ostrov-2 site, we sequenced one individual, from burial 1 (I2144). It is dated to 2911-2677 calBCE (4230±35 BP, OxA-33167).

**(4) Chernoozerye-1 burial site**

The Chernoozerye-1 burial ground was excavated by Vladimir Gening between 1967 and 1969. The site is situated on the periphery of the floodplain terrace of the Irtysh River. This particular area of the terrace has been repeatedly utilized by the ancient population for habitation purposes. Alongside the burials found at the Chernoozerye-1 burial ground, other archaeological sites from the Paleolithic, Neolithic, and pre-Andronovo Bronze Ages have been discovered here. Additionally, a burial ground associated with the Andronoid culture, five small burial mounds, and one large burial mound from the Early Iron Age were found. Furthermore, a burial from the Iron Age was also uncovered at the site. The excavated area of the burial ground covers approximately 3,500 square meters in total (Gening and Stefanova, 1994).

*(a) burial 57 (individual ID I6050)*

**Burial 57** (individual ID I6050). The grave is oval-shaped and oriented in the north-northeast-south-southwest direction, with dimensions of approximately 150x80x10 cm. Based on the well-preserved bones, the deceased individual is identified as a male, estimated to be around 50 years old. The burial position is crouched, with the individual lying on his right side, and the head placed towards the north-northeast. The femurs are positioned at a right angle to the spine, while the lower leg bones are nearly perpendicular to the femurs. The left hand is bent with the palm resting on the stomach, while the right hand is bent with the palm towards the face. No grave goods were present in this burial (Gening and Stefanova, 1994). The burial is dated to 4216-3968 calBCE (5225B±25 BP, PSUAMS-2944); freshwater effect is possible.

#### II.D. Urals

##### II.D.i. Trans-Urals (Tobol River Basin in Western Siberia)

According to current archaeological understanding, the Early Neolithic period in the Trans-Urals (6000–4700 BC) is characterized by the simultaneous existence of two significant cultural traditions, namely Koshkino and Kozlov-Mys (Shorin and Shorin 2021, 2022; Vybornov et al., 2014).

The Late Neolithic period (5000–3950 BC) is represented by two distinct cultural traditions: Boborykino-Bas'yanovo, and Poludenka-Comb-Pottery (Shorin and Shorin 2021, 2022; Shorin et al., 2015; Vybornov et al., 2014). Some researchers also distinguish the specific Sosnovy Ostrov culture (Usacheva 2016), but other treat it as a part of Poludenka-Comb-Pottery tradition.

The Eneolithic period in the Trans-Urals is dated to 4300-3000 BC and is characterized by the coexistence of several cultures identified through the ornamental features of their pottery. These cultures, namely, Shapkul', Andreevskaya (Andreevskoye-Ozero), Lipchinskaya (Lipki), and others, are collectively referred to as the Trans-Ural Eneolithic, which shares material culture characteristics with the Comb-Pit Pottery of southern western Siberia (Mosin, 2008; Epimakhov and Mosin, 2015). Clear boundaries between the Neolithic as well as Eneolithic cultural groups in the Trans-Urals were absent, with numerous transitional types present. The populations of the Trans-Urals during those periods engaged in mobile practices aligned with the annual economic cycle. This involved movement across foothill, forest, and forest-steppe territories, with each community having a network of long-term (winter) and short-term (summer) camps and settlements.

The primary subsistence strategies revolved around hunting and fishing. Hunting targeted animals such as horses, elks, bears, red deer, roe deer, badgers, martens, otters, as well as waterfowl. Fishing activities are inferred from the discovery of variously shaped sinkers among the artifacts, as well as the presence of fish bones and scales.

###### II.D.i.a) Trans-Ural Late Neolithic-Eneolithic (Russia_CombPitWare_Eneolithic)

###### (1) Sosnovy Ostrov occupation site

The site of Sosnovy Ostrov is situated on a sand dune that rises approximately 2 meters above the swampy region of the lake known as Maly Tarman. This lake is part of the Tarman Lakes, which are remnants of a paleochannel of the Iska River, a left tributary of the Tobol River in the Middle Urals.

During the excavations of the site, several cultural assemblages were discovered. Assemblage 1 comprises ceramics and housing structures attributed to the Kozlov culture, which represents the early Neolithic period. Assemblage 2 consists of ceramics and housing structures associated with the Sosnovy Ostrov culture. Within Assemblage 2, a distinct subset labeled as Assemblage 2a was identified, consisting of pottery and Burial 2, both associated with the Sosnovy Ostrov culture of the Late Neolithic. Finally, Assemblage 3 is interpreted as belonging to the Eneolithic period.

*(a) Sosnovy Ostrov, burial 1 (individual ID I12696)*

Three burials were made within the central section of the western wall of a dwelling pit measuring 4 meters in length and 1.3 meters in width, attributed to Assemblage 2, corresponding to the Sosnovy Ostrov culture, shortly after its collapse.

**Burial 1** (individual ID I12696) was discovered in the central area of the dwellingpit, at a depth of 170 cm. Within the burial, the remains of a human skull and small calcified bones, potentially belonging to a complete skeleton, were excavated. The orientation of the pit follows a northeast-southwest line. The grave goods associated with this burial include a polished "hammer" crafted from green stone. The burial has been dated to the period of 4245–4050 calBCE (5320±25 BP, PSUAMS-9040). Cultural attribution of the burial, however, is problematic. It was constructed after the Sosnovy Ostrov house was destroyed but before the site was occupied by the population with Comb-Pit pottery.

Additional burials within the site were unearthed at deeper layers within the dwelling pit (the following two burials were not sequenced).

Burial 2 is located at a depth of 190 cm. At this depth the pit diminishes in size, measuring 1.4 x 1.75 meters. At this level, a human jawbone was discovered. The pit is oriented along the northeast-southwest line. Grave goods include a miniature vessel adorned with a comb stamp, which corresponds to Assemblage 2a of the Sosnovy Ostrov culture.

Burial 3 was excavated at a depth of 220 cm, this grave pit measures 210 cm in length and 80 cm in width. On the southern side, the pit is bounded by a 1-meter-long charcoal deposit from Assemblage 1. Within this section of the grave, the remains of a third burial were found, including a skull, hand bones positioned across the grave pit, and traces of rib fragments. The positioning of the skull and ribs indicates that the head of the interred individual was oriented towards the east-northeast. No grave goods were discovered in association with this burial.

**(2) Boborykino-2 occupation site**

A burial located at the Boborykino Dune was discovered approximately 0.1 km southwest of the former village of Boborykino, situated on a cliff along the left bank of the Iset River in the forest-steppe region of the Tobol River Basin.

Konstantin Salnikov and Otto Bader associated the burial with the Boborykino-2 occupation site due to its spatial proximity, being only 0.2 km away, as well as due to similarities observed in the shapes of pendants and pointed axes found in the occupation layer and as part of the grave goods (Salnikov 1962, Bader 1970). Although the cultural attribution was not explicitly specified, it was mentioned that the burial was of Neolithic origin.

At the Boborykino-2 site, pottery classified as Poludenka-type was attributed to the Neolithic period, while pottery decorated with comb stamps was assigned to the Eneolithic period. Additionally, pottery of the Boborykino type was associated with the Early Bronze Age (Salnikov 1961). Konstantin Salnikov further delineated a chronological sequence of cultural groups recorded at the site, which included the Neolithic, Late Neolithic, Boborykino, Andronovo, and Kamenogorsk cultures, Sarmatian period, and Bakal culture (Salnikov 1961).

*(a) damaged burial at the Boborykino-2 site (individual ID I8280)*

The individual interred in the **damaged burial** (individual ID I8280), estimated to be around 60 years old, was in a supine position, and the bones were painted by ocher. Adjacent to the right side of the skull, the remains of another skull and a jaw belonging to a child aged 10-12 years were discovered.

The bones of a young adult were also found in exposed layers at the river bank, raising the total number of buried individuals nearby the Boborykino-2 site to three.

Among the grave goods, a total of 26 polished bone drop-shaped pendants were unearthed in the vicinity of the neck of the first interred individual. Additionally, 20 flint knife-shaped pieces, measuring up to 20 cm in length, were found, along with two stone axes, three bone beads, and several end scrapers positioned to the right of the skull. Traces of red dye were observed on the tools (Salnikov 1962; Bader 1970; Vokhmentsev 2000). The burial is dated to 4338–4176 calBCE (5410±25 BP, PSUAMS-4278). Based on δ15N/δ15C ratio, we do not expect the freshwater reservoir offset in this date.

Mikhail Vokhmentsev attributed the Boborykino burial to the Eneolithic period based on the presence of drop-shaped pendants, as similar decorations were predominantly found in Eneolithic burials and sites (Vokhmentsev 2000). However, the chronological position of the burial makes the Late Neolithic attribution more plausible.

#### II.D.i.b) Trans-Ural Eneolithic (Russia_CombPitWare_Eneolithic)

##### (1) Gladunino 3 occupation site

The Gladunino-3 site is located on the eastern shore of Lake Voronye, at a height of 2.5 – 3 m from the water surface level. The settlement supposedly occupied an area of 2100 square meters. The peripheral part of the site was excavated, where 5 storage pits, traces of post holes, and a single burial were recorded. Two phases have been identified on the site: 1 - Early Neolithic (the Koshkino and the Kozlovskaya (Kozlov-Mys) cultures), 2 - Eneolithic (the Sosnovy-Ostrov, Shapkul', and Andreevskaya (Andreevskoye-Ozero) cultures) (Shilov and Maslyuzhenko 2002). According to recent studies, the Sosnovy-Ostrov culture is attributed to the Late Neolithic period.

###### *(a) burial at the Gladunino-3 settlement (Iindividual ID 6305)*

The **inhumation burial** (individual ID 6305) discovered at the Gladunino-3 site was made in a rectangular grave pit measuring 219x61x20 cm and aligned along the southwest-northeast orientation. The interred individual was positioned on their back, in a supine position, with the head facing eastward. The anatomical order of the skeleton was maintained, except for the deliberate dissection of the left pelvic bone, which was neatly placed between the ribs and the right half of the skeleton. The fingers of the right hand were flexed, while the left hand was laid in a resting position on the pelvis. Additionally, the left femur exhibited slight rotation.

The grave goods accompanying the burial include:

1. A fragment of a spherical stone mace, crafted from brown iron ore and featuring a perforation.
2. Fragments of pottery decorated by comb stamp patterns and pit impressions, which the excavators attributed to the Sosnovy Ostrov culture.
3. An end scraper made of gray-green jasper.
4. Two blades and two flakes made of grayish flint and quartz (Shilov and Maslyuzhenko 2002; Khohlov and Nechvaloda, 2002).

The burial is dated to 2894–2701 calBCE (4210B±20 BP, PSUAMS-2910). Cultural attribution of the burial is problematic, but its chronological position suggests the Late Eneolithic period when the region was mostly populated by groups with Comb-Pit pottery or closely related to them.

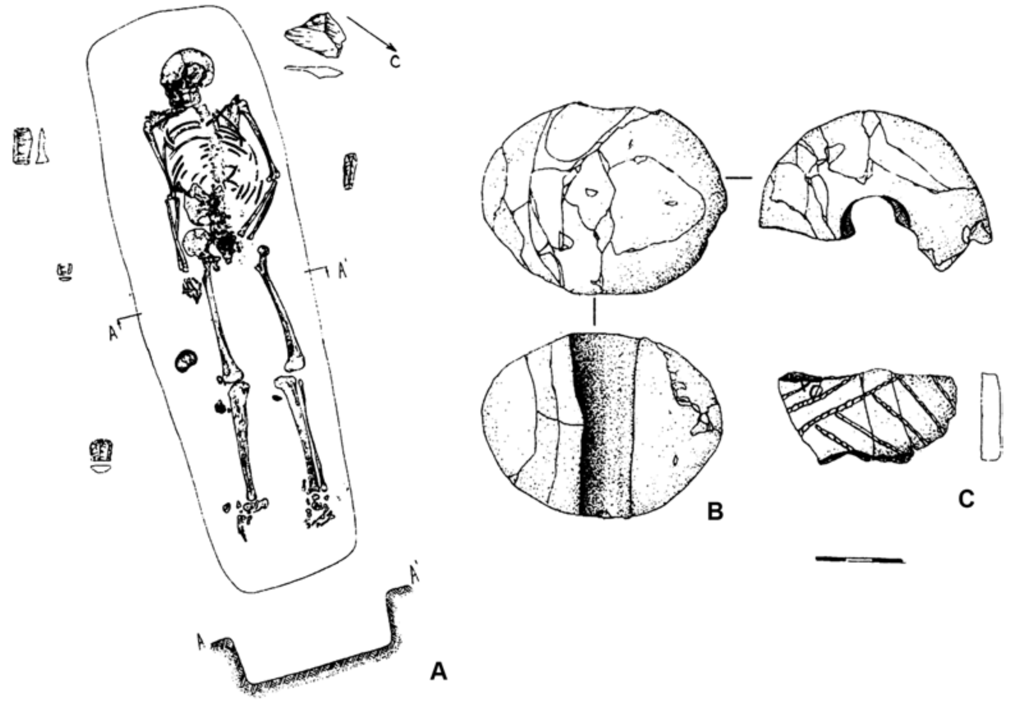

***Fig. S46.*** *The Eneolithic burial at the Gladunino-3 site.* ***A*** *- burial;* ***B*** *- stone mace;* ***C*** *- pottery sherd (Shilov and Maslyuzhenko 2002, modified).*

##### II.D.ii. Ural Mountains and Cis-Urals

In the Urals region, a wide range of archaeological cultures and cultural types from the Neolithic to the Eneolithic period have been identified. These cultures often coexisted contemporaneously, and their main distinguishing feature was the ornamentation found on everyday pottery vessels, such as comb, comb geometric, pricked, incised, false cord, and others.

The burials from the Urals studied in this paper, were typically situated in remarkable locations such as caves, grottoes, and beneath rocky overhangs. These burial sites lacked cultural markers, specifically ceramic vessels, which makes it impossible to attribute them to a specific Ural Neolithic or Eneolithic culture. Therefore, the most appropriate term to describe these burials accurately would be "Ural Early Eneolithic Burials."

###### II.D.ii.a) Ural Early Eneolithic Burials (Russia_UralsEneolithic)

Systematic exploration of grottoes and caves in the Urals began in the 1930s under the direction of Sergei Bibikov. In 1938, a burial was discovered in the Buranovskaya cave, which appeared to be that of a male individual. Accompanying the burial were 35 drop-shaped pendants made of ophite (serpentine). The following year, a female burial was excavated in the rocky canopy of Starichny Greben, which contained 15 pendants made of ophite, similar to those found in the Buranovskaya cave. In the same year, the remains of two individuals were found in the Ust-Kataevskaya-2 cave. One of them, presumably a child aged 5-7 years, was remarkably well-preserved. The child's bones were placed in a recess in an incorrect anatomical order and covered with stones. Among the grave goods in this burial were 20 incisors of a bobak marmot without holes, intensely colored with ocher, as well as 53 conical beads made of *Dentalium* mollusk shells, 36 flat and rounded shell beads, and 32 teardrop-shaped pendants made of gray slate stone (Bibikov 1950).

The next phase of research took place in the 1980s when Yuri Serikov discovered and excavated burials at Kamrn’ Dozhdevoy.

In more recent years, cultural remains have been investigated in the Burmantovsky grotto and Sokolinaya Gora caves, revealing a complex assemblage of artifacts, likely originating from destroyed ground burials. In the Burmantovsky grotto, apart from a human tooth, 69 various pendants made of bone and shell, and 37 arrowheads made of flint and green tuff-porphyrite were found. A similar assortment of artifacts was uncovered in the Zhilische Sokola cave.

No established tradition of funeral rites has been identified; however, several features have been recorded:

1. Inhumation in shallow pits.
2. Burials covered with earth, stones, or interred in grottoes and caves.
3. Evidence of a fire cult associated with the burials.
4. Traces of ocher found in shallow pits (8-30 cm deep), on the bones, and on grave goods in some burials.
5. Skeletons, when laid in the correct anatomical order, displayed a northeast or southwest orientation.

Typical is also the presence of drop-shaped pendants made of stone or bone, as well as jewelry crafted from teeth of various animals, among the grave goods. The general chronological framework for the Ural Early Eneolithic burials is 4500–3900 BCE (Serikov, 1998; Nokhrina and Shorin, 2000; Epimakhov and Mosin, 2015; Shilov and Maslyuzhenko, 2002).

##### (1) Kulmetovskiy-Grot cave

Kulmetovskiy Grot is situated on the right bank of the Ai River, a left tributary of the Ufa River, which in turn is a right tributary of the Belaya River, a left tributary of the Kama River. The cave is located approximately 3-4 kilometers upstream from the village of Kulmetovo. At the entrance of the cave, there is a collection of rock paintings.

###### *(a) burials 1 (individual ID I20893) and 2 (I20948)*

During archaeological excavations of cultural layers dating to the Eneolithic period, fragmented remains of two human skeletons were discovered, documented as **Burial 1** (individual ID I20893) and **Burial 2** (individual ID I20948). Unfortunately, the burials were not excavated carefully, and as a result, the burial rites associated with these individuals remain unknown (Shirokov, 2009). Burial 1 is dated to 4047-3961 calBCE (5195B±25 BP, PSUAMS-9029), and burial 2 to 4315-4052 calBCE (5335B±25 BP, PSUAMS-9031).

##### (2) Kamen’-Dozhdevoy cave

Kamen' Dozhdevoy is a prominent rocky cliff reaching heights of up to 35 meters, situated on the right bank of the Chusovaya River. Within this cliff, one of the notable rifts is located beneath the Kamen’ Dolzhdevoy. The grotto, characterized by its karstic origins, features an entrance situated 14 meters above the water level. In 1988, Yuri Serikov discovered burials within this grotto.

These burials consist of inhumations placed within shallow pits and adorned with a sprinkling of ocher. The first burial pertains to a female individual in a supine position, while the second burial corresponds to a male individual in a crouched position. Notably, both skeletons exhibit an orientation towards the northeast.

Within the male burial, an assortment of artifacts was recovered, including 2 biconical bone arrowheads, 4 stone arrowheads, a stone knife showing signs of grinding, a bone harpoon, a scraper crafted from a chalcedony flake, 10 beaver incisors, one fox tusk, 8 bone ornithomorphic figurine-pendants, and 6 drop-shaped bone pendants made from beaver bones. Additionally, 14 flake-blanks, presumably stored within a bag, were also discovered at the burial site (Serikov, 1993).

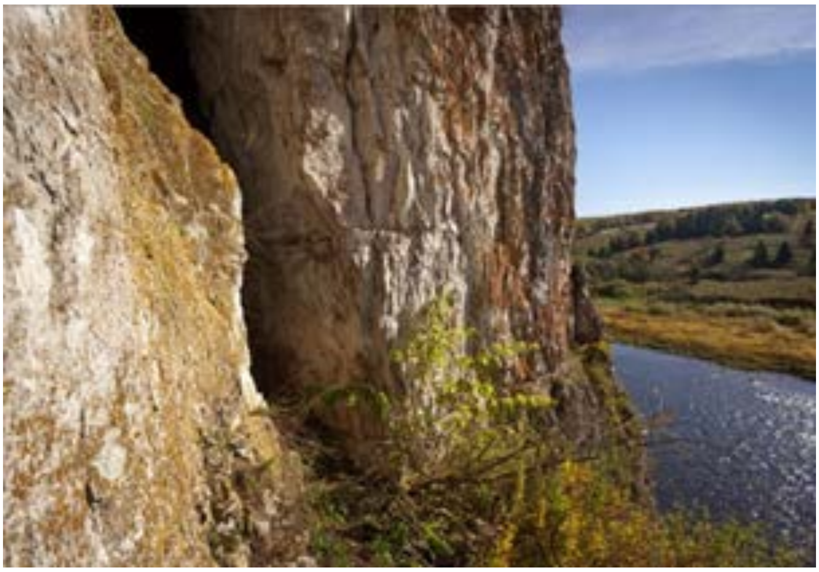

***Fig. S47.*** *The Kamen'-Dozhdevoy cave.*

*(a) burial 1 (individual ID I11637)*

The female burial, designated as **Burial 1** (individual ID I11637), was found in a supine position with an orientation towards the northeast. Among the grave goods accompanying the burial, a substantial assemblage of 76 pendants made of animal teeth was identified. Specifically, these pendants consisted of 73 lower fangs from otters, 2 moose incisors, and a bear tooth. Additionally, the burial yielded undrilled incisors from an elk and a lower canine of a fox (Serikov, 1993). The burial is dated to 4496–4253 calBCE (5530B±60 BP, COAH-5154).

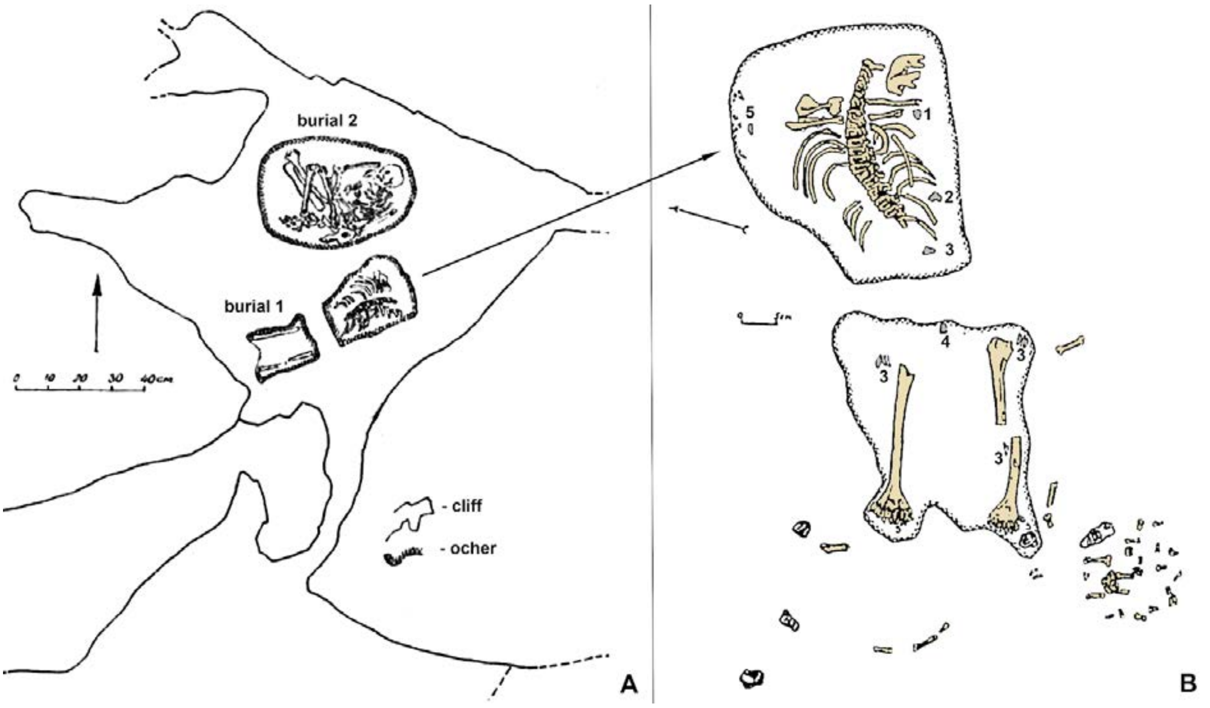

***Fig. S48.*** *Burials 1 and 2 in the Kamen'-Dozhdevoy cave.* ***A*** *- excavation plan;* ***B*** *- burial 1;* ***1, 2*** *- flakes;* ***3*** *- pottery sherds;* ***4*** *- scrappers;* ***5*** *- perforated animal teeth (Serikov, 1993, modified).*

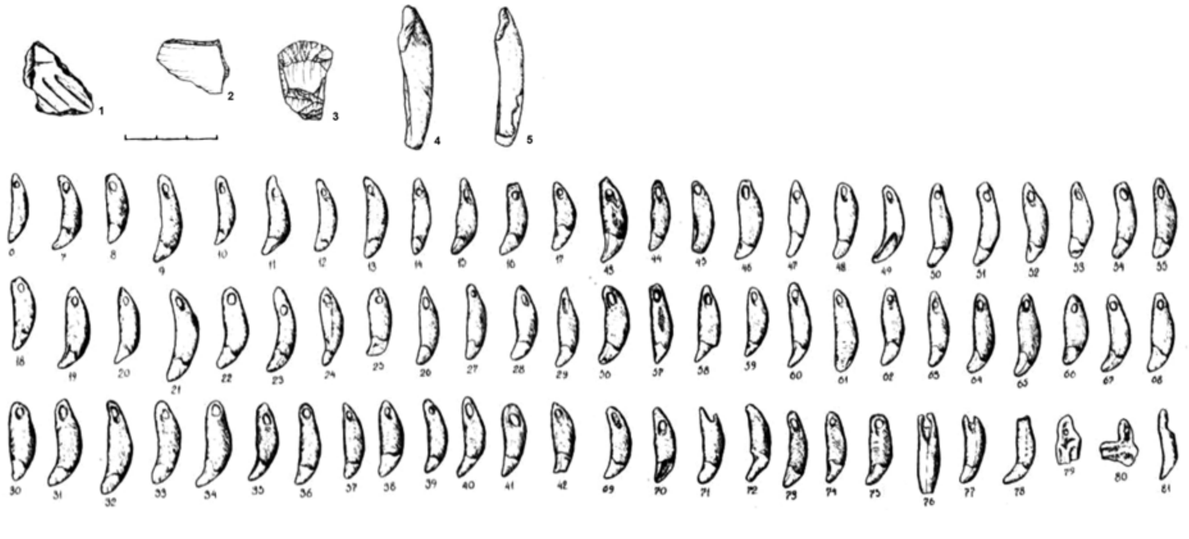

***Fig. S49.*** *Grave goods from burial 1 in the Kamen'-Dozhdevoy cave.* ***1*** *- pottery shard;* ***2*** *- flake;* ***3*** *- scrapper;* ***4, 5*** *- moose incisors (Serikov, 1993).*

### II.E. Temperate zone of Eastern Europe

#### II.E.i. Lower Kama and Middle Volga Region

##### II.E.i.a) Elshanka Early Neolithic culture (Russia_Elshanka_EN)

Elshanka-type sites were initially excavated during the mid-1970s in the forest-steppe and steppe regions of the Urals. The culture derived its name from the eponymous site, Staraya Elshanka-2. Presently, approximately 25 sites in the forest-steppe Volga region have been investigated, all of which contain artifacts associated with the Elshanka culture.

The pottery of the Elshanka culture exhibits characteristic features, including relatively thin walls with a smoothed or burnished outer surface, as well as a smoothed inner surface. Vessels with vertical walls are prevalent, while pointed and flat bottoms are found in equal numbers, with rounded bottoms occurring less frequently. Ornamentation is typically confined to the upper third of the vessel, primarily on the neck and occasionally at the rim's top. The typology of the flint tools remains a topic of debate.

One of the most widely accepted hypotheses proposes that the Elshanka culture emerged in the first half of the 7th millennium BCE. It persisted in the forest-steppe Volga region at least until the middle, and possibly even the third quarter, of the 6th millennium BCE.

The issue of the origins of Neolithization (i.e., the appearance of pottery) in the forest-steppe Volga region remains highly contentious. However, there is a notable resemblance between the ceramics of Central Asia and the pottery of the Elshanka culture.

During its later development stage, the Elshanka population interacted with communities practicing the inscribed pottery tradition in the Lower Volga region. The initial advancement of the Lower Volga population appears to be associated with increased aridization, particularly during the latter half of the 7th millennium BCE.

At the peak of aridization (end of the 7th millennium BCE), the Lower Volga population expanded into the forest-steppe region, leading to the formation of the Middle and Late Neolithic Volga-Kama culture. The first half of the 6th millennium BCE witnessed the coexistence of late Elshanka complexes alongside early Volga-Kama culture complexes, ultimately resulting in the gradual decline of the Early Neolithic ceramic tradition (Andreev et al., 2021).

##### (1) Chekalino-4 occupation site

The Chekalino-4 site is situated on the Sok River, specifically at the northern extremity of the first terrace above the floodplain. It is positioned between an ancient dry channel, which has left behind two remaining lakes, and a flooded floodplain depression that directly adjoins the modern river channel. Excavations have revealed that the site existed in close proximity to water during ancient times. Artifacts from the Neolithic, Eneolithic, and Bronze Ages have been discovered at this location.

Regarding the Neolithic ceramics found at the site, they can be categorized into two distinct cultural and chronological complexes: Elshanka and Volga-Kama. The Elshanka findings are covered by soil that developed on alluvial deposits and are located at the level of the buried soil surface. Radiocarbon dating conducted on pottery sherds has provided a broad chronological range spanning from 8450 BCE to 6010 BCE (Andreev et al., 2018).

###### *(a) The Chekalino-4 settlement, Neolithic burial B (individual ID I6413)*

Approximately 4 meters southeast from the channel bank, an excavated burial, commonly referred to as "**burial B**" (individual ID I6413) in recent publications, was discovered by Andrey Mamonov and Aleksandr Khoklov. The exact outlines of the grave pit were not clearly defined. The skeleton was found just below the upper level of the subsoil. The interred individual was positioned in a crouched posture, with the arms placed beneath the head and the legs bent at the knees. The orientation of the skeleton is to the northwest. The grave goods accompanying the burial consist of an unretouched knife-shaped artifact, which was located at the base of the skull. The absence of soil redeposition indicators in the layers above the burial allows us to attribute it to the Elshanka period (Mamonov, 1995; Vybornov et al., 2023). However, the radiocarbon dating of the burial to 8281–7971 calBCE (8970±30 BP, PSUAMS-4211) appears highly unlikely, as it contradicts at least some dates obtained from mollusk shells found at the same site (Le-4781, 8990±100; Le-4782, 8000±120; Gin-7086, 7950±130). We anticipate the presence of a freshwater reservoir effect in these results and instead rely on the date obtained from charred plant remains in pottery clay from the same site (Poz–42051, 7250±60 BP), which corresponds to the period of 6230–6010 calBCE (Andreev et al., 2021).

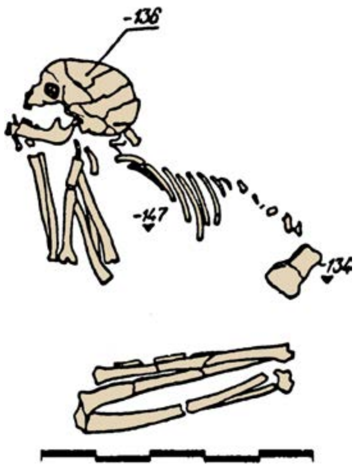

*Fig.* ***S50***. *Burial B at the Chekalino-4 site. 1 - knife-shaped piece.*

##### (2) Labazy burial mounds

The series of burial mounds near the village of Labazy underwent excavation under the supervision of Nina Morgunova in 2007-2008. These Labazy kurgans are situated in the steppe region of the Urals, within the elevated floodplain of the Buzuluk River. The river itself flows approximately 1 km east of the kurgans, characterized by a deeply incised channel and steep, precipitous banks. The surrounding area features a wide, treeless valley adorned with meadow vegetation and sparse shrubs. The necropolis was positioned on the left, higher section of the floodplain. A total of 8 mounds were excavated at the site, with the majority of the burials ascribed to the Late Bronze Age Srubnaya culture. Within mound 4, earlier-age burials were discovered, including one from the early phase of the Srubnaya culture. It appears that the mound was constructed during the Late Bronze Age, potentially covering a cemetery from the Neolithic period (Kuptsova et al., 2019).

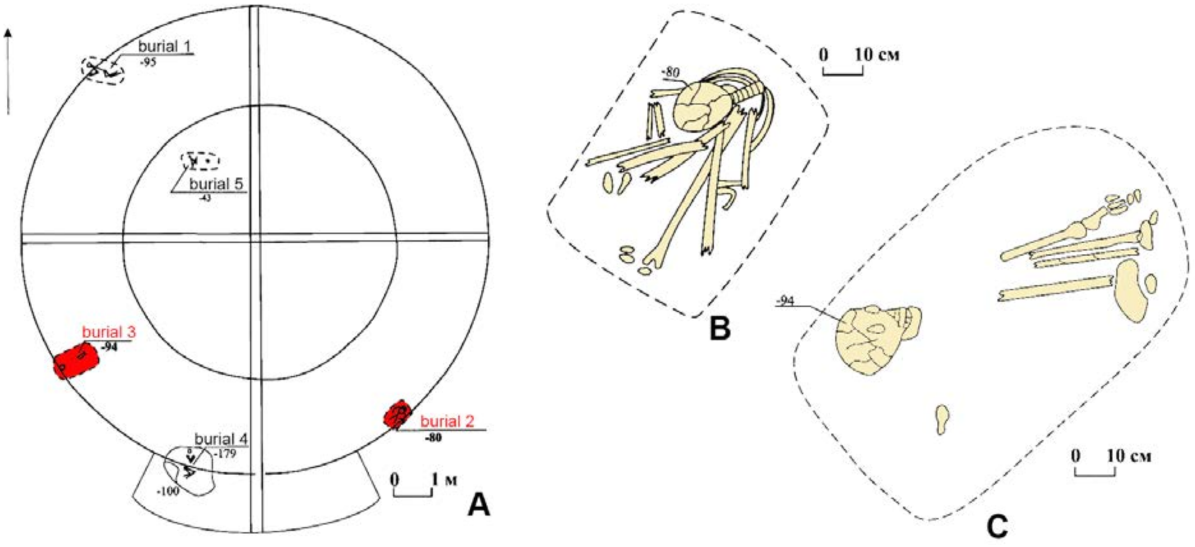

***Fig. S50.*** *Labazy burial mounds.* ***A*** *- plan of mound 4 (the burials that contained sequenced individuals are labeled in red),* ***B*** *- burial 2,* ***C*** *- burial 3 (Kuptsova et al., 2019, modified).*

*(a) Mound 4, burial 2 (individual ID I6916)*

**Mound 4, Burial 2** (individual ID I6916) corresponds to an elderly woman aged over 45. The skull was discovered 5.7 meters southeast of the central point of the mound. The skeleton was positioned in a seated posture, with the legs bent at the knees and the arms bent at the elbows, pulled up towards the knees. The spine exhibits a curved shape, causing the body to tilt forward, with the head resting in the hands. The back of the head is oriented towards the northeast, while the front part faces the southwest. It is possible that the woman was bound during the burial ceremony. No grave goods were uncovered in association with this burial. The radiocarbon dating for this burial indicates a period between 6387–6234 calBCE (7430±30 BP, PSUAMS-4261).

*(b) Mound 4, burial 3 (individual ID I12838)*

**Mound 4, Burial 3** (individual ID I12838) pertains to a male individual aged between 45 and 55 years old. This burial was located in the southwestern sector of the mound, approximately 5.6 meters southwest from the center. The approximate dimensions of the burial pit were 106x63 cm. The skeleton was incompletely preserved, with the skull, leg bones, pelvis, and collarbone being recovered from the grave. The orientation of the skeleton places the head towards the southwest. No grave goods were found in association with this burial. The radiocarbon dating for this burial indicates a period between 6449–6264 calBCE (7525±26 BP) [Radiocarbon dates combined: 7565±50 BP, PSUAMS-7795 7510±30 BP, PSUAMS-4260].

Burials 2 and 3, located in mound 4 at Labazy, are among the oldest burials within the steppe region of the Volga-Ural area. They have been dated to the 7th - beginning of the 6th millennium BCE. This timeframe aligns with the presence of the Elshanka culture in the Volga forest-steppe region. Notably, the Staraya Elshanka-2 site, where Elshanka pottery was first described, is situated only 30 km north of Labazy. This correlation suggests that the individuals interred at Labazy may have belonged to the same population that introduced the earliest Neolithic pottery in the East European Plain (Kuptsova et al., 2019).

##### II.E.i.b) Khvalynsk Eneolithic culture (Russia_Khvalynsk_Eneolithic)

The Khvalynsk Eneolithic culture was first identified by Igor Vasiliev in the late 1970s and early 1980s, based on findings from the Khvalynsk-1 burial site, which was excavated by Igor Vasiliev, Sergey Agapov, and Valentina Pestrikova between 1977 and 1979.

Khvalynsk culture sites have been identified across a vast territory, ranging from the Northern Caspian Sea in the south to the Middle Sura in the north, and from the Moksha River in the west to the upper reaches of the Samara River in the east.

Pottery serves as the most common and recognizable characteristic of the Khvalynsk culture. Various types of vessels are distinguished, including bell-shaped vessels with two variants - straight-walled and with a rounded body and a contracted neck (sac-shaped), spherical vessels, and bowls. A notable feature of Khvalynsk pottery is the thickened collar rim. The majority of vessels are ornamented, with the ornamentation primarily located in the upper portion, extending over the top and the inner surface of the rim, and occasionally covering the entire outer side, including the bottom. Pricking, stamping, and drawing techniques were used for ornamentation.

Khvalynsk sites mainly consist of burial grounds, although some settlements with long-term dwellings are known in the northern part of the region. Burial grounds are generally flat, and burial pits are typically indistinct. The predominant burial position is a crouched position on the back, with the legs bent at the knees, arms bent at the elbows, and hands on the pelvic bones. There are variations in the arm position, with arms either extended along the body, bent at the elbows with hands on the pelvic bones, or bent with hands on the stomach or chest. Skeletons are also found in positions on the left or right side, as well as in a sitting position. Secondary burials, characterized by bones and skulls stacked in a heap, as well as burials imitating the anatomically correct positioning of skeletal bones, have been identified. The orientation of the burials primarily aligns with the northeast, northwest, and north directions, although other orientations are also observed. Ocher use varies from minimal to abundant dressing. Altars containing bones of large and small cattle, horses, and fragments of artifacts have been discovered within the burial grounds.

The chronological timeframe of the Khvalynsk culture is defined as 5000–4500/4300 BCE. The origins of the Khvalynsk culture are subject to debate, with most researchers suggesting connections to the Mariupol group of cultures, particularly the Samara culture in the eastern part of the region. The development of the Khvalynsk tradition is thought to continue within the context of the Yamnaya culture (Korolev and Stavitskiy, 2021b; Kuznetsov, 2020; Morgunova, 2020; Agapov, 2010; Agapov et al.,1990).

##### (1) Lebyazhinka-5 occupation site

The Lebyazhinka-5 settlement is situated on the periphery of the Lebyazhinka farmstead, occupying the first floodplain terrace on the left bank of the Sok River, a tributary of the Volga River. The excavations conducted at the site by Mikhail Turetsky, Nadezhda Ovchinnikova, Aleksandr Khokhlov and others in 1997 have revealed a wide range of materials spanning from the Neolithic period to the late Middle Ages. Among the significant findings, several Eneolithic burials, labeled as Burials 8, 9, and 12, were uncovered (Vybornov et al., 2000, 2023; Shishlina et al., 2017).

###### *(a) Eneolithic burial 12 (individual ID I6908)*

**Burial 12** (individual ID I6908) at the Lebyazhinka-5 site contains five skeletons, with the exact contours of the grave pit remaining elusive. The approximate boundaries of the burial were established based on the arrangement of the skeletons, showcasing variations in their positions. Unfortunately, the paucity of the records means that we cannot confirm which of the skeletons in this grave was sequenced for this study.

*Skeleton 1* was positioned in the western part of the burial as a bundled arrangement, where bones without a skull were concentrated within a small area. This skeleton likely belonged to a mature woman.

*Skeleton 2* was located east of skeleton 1, in a crouched position on its back, with the head oriented towards the north. The skull of this skeleton is preserved, and it belonged to a woman aged 30-40.

*Skeleton 3* was situated east of skeleton 2, lying supine and facing north, although the skull was absent. This skeleton belonged to a man aged 35-40.

*Skeleton 4* was found beneath skeletons 2 and 3. The position of the bones suggests that the individual was laid prone, with only the spine, pelvic bones, and a few ribs remaining. The skull, along with other bones, was missing. The orientation of the buried person has not been determined, and this skeleton belonged to a young man aged 15-17.

*Skeleton 5* partially underlay skeleton 3. It preserved several vertebrae, ribs, clavicle, sternum manubrium, fragments of the left shoulder blade, and pelvis. Among the long bones, the left humerus and the lower half of the right humerus, along with bones from the right forearm, were present. The skull, however, was missing. Presumably, this skeleton was positioned in an extended position on its back, with a probable northward orientation. It belonged to a man aged 20-30.

Burial 12 also contained various artifacts, including 250 marmot incisors, 100 carp pharyngeal teeth, one tubular piercing made of bone or shell, and two flint plates. Ocher was employed in the burial ceremony. Skeletons 4 and 5 possessed the largest number of artifacts, with skeleton 4 exhibiting a greater quantity of marmot incisors and skeleton 5 displaying more fish teeth. These items were likely sewn onto clothing, possibly as adornments on belts or in the region of the spine (Shishlina et al., 2017).

Initially attributed to the Samara culture, the burial was later reevaluated as belonging to the early Khvalynsk culture. It is dated to 4838–4612 calBCE (5865±40 BP, GrA-64051), with a marmot tooth from the grave providing the basis for this dating.

The individual ID I6908 is labeled as one of the skeletons from burial 12, without specification.

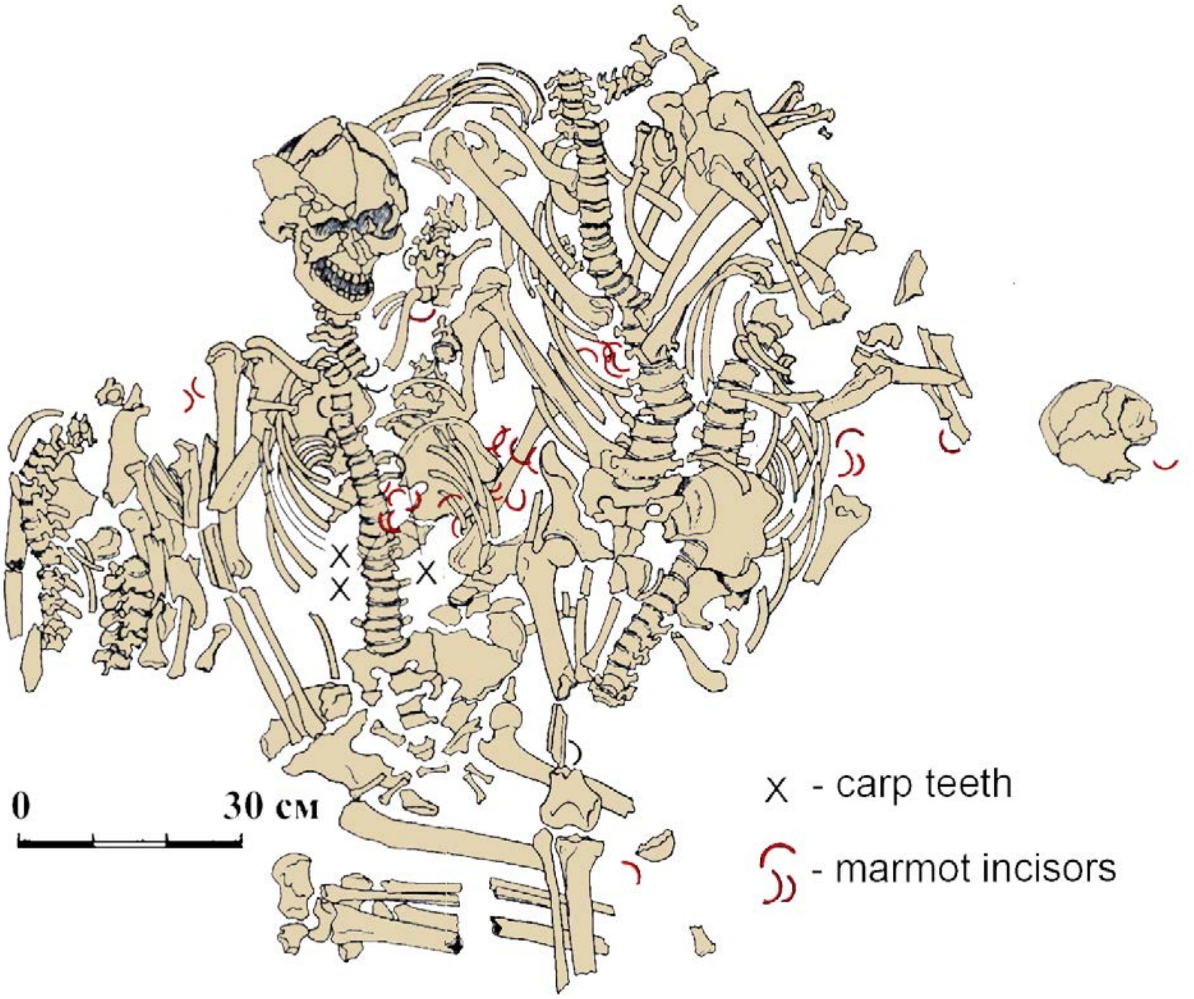

***Fig. S51.*** *Burial 12 at the Lebyazhinka-5 site (Shishlina et al., 2017, modified).*

##### II.E.i.c) Eneolithic at the Kama Estuary (Russia_KamaEstuaryEneolithic)

In the Kama Estuary region, the early phase of the Eneolithic period is attested by four sites belonging to a distinct archaeological group known as the Kama Estuary Eneolithic. This group comprises three burial grounds and one occupation site.

The burial grounds were situated on flat terraces, occasionally with slight natural elevations. They are characterized by rows of graves, with the deceased interred in either a sitting or supine position. Supine burials are typically individual, while sitting burials are collective in nature.

The grave goods found in the Kama cemeteries consist of objects made of stone, bone, and copper. Woodworking and cutting tools, such as trapezoidal adzes and chisels with grooves and distinct handles, are among the most numerous. Numerous cutting tools such as blades, scrapers, incisors, knives, and points have been discovered. Military equipment includes willow-shaped arrowheads and javelin tips, as well as axes and daggers with or without a "button" feature. Hoe-shaped tools likely pertain to agricultural activities. Additionally, a unique stone artifact with a sculptural representation of a phallus was found. Many of the stone tools found in these cemeteries share similarities with those from the Neolithic Kama culture, indicating the continuity of stone tool-making traditions from the local Neolithic period. The presence of new stone-working technologies, such as the use of quartzite and the production of large plates for tool manufacture, along with the discovery of copper artifacts and evidence of emerging craft-based economy, suggests the introduction of innovative ideas associated with the influx of population from the south.

The distinctive pottery of the Kama Eneolithic is characterized by cup-shaped open forms with round bottoms. The clay paste contains organic impurities, and the rims of most vessels are either outwardly flared or inwardly beveled. The primary decorative elements consist of comb stamps and oval depressions, with the most typical ornamental compositions including horizontal zigzag friezes and diagonal impressions of comb stamps, as well as belts of oval pits. These compositions are sparsely placed on the vessel surface. The ceramics of the Kama Estuary Eneolithic display similarities to the pottery traditions of the Bor culture.

Recent studies have dated the Kama Estuary Eneolithic to 4600–4000 BCE. The formation of this cultural group is attributed to the influence of Khvalynsk traditions in the forest zone. It is regarded either as a peripheral northern component of the Khvalynsk culture or as a peripheral southern extension of the Bor culture. In either case, the material culture of the Kama Estuary Eneolithic demonstrates a mixed nature, combining traits from both the Khvalynsk and Bor cultures (Chizhevskiy and Shipilov, 2021; 2018).

###### (1) Murzikha-2 burial site

The Murzikha-2 cemetery was situated on two promontories of the second terrace on the left bank of the Arkharovka River, a left tributary of the Kama River. Currently, the site is submerged under the reservoir formed by the Kama River bridge dam. The majority of the burials at Murzikha-2 date to the Late Bronze Age and Early Iron Age. Eneolithic burials were discovered by Andrei Chizhevsky in 1995, located approximately 35-40 meters from the modern shoreline in the central part of the necropolis.

During the excavation, a total of 18 burials were uncovered, arranged in three rows with 3-6 burials each, along with some individual burials. The rows extend in the northeast-to-southwest direction.

Out of the 18 burials, 12 exhibited identifiable grave pits, which were predominantly round or quadrangular in shape. The depth of the burials was generally shallow, ranging from 4-30 cm, with an average depth of 11 cm.

Approximately two-thirds of the burials were collective, containing two to four individuals. All burials were inhumations, although some individuals were found with evidence of prior cremation within the graves. Ocher was observed in many of the grave pits. The skeletons were positioned in either a sitting or crouched position, with bent legs. Based on the best-preserved skeletons, it can be inferred that the deceased were placed in a sitting position during burial, with their legs half-bent, hands resting on the knees with forearms, and their heads bowed onto their chests.

The orientation of the buried individuals varied, but an eastern direction predominated. In collective burials, two different arrangements were observed: 1) in burials with two or four individuals, the skeletons were placed in a single row, shoulder to shoulder, all facing the same direction; 2) in burials with three individuals, the skeletons were oriented with their heads facing different directions, while their feet pointed towards the center of the grave pit.

The burial inventory of Murzikha-2 was abundant and diverse, comprising items crafted from stone, bone, horn, and copper. Stone artifacts included woodworking and cutting tools, tools for leather processing, weapons and hunting implements, as well as tools associated with land cultivation. Woodworking and cutting tools were the most prevalent category. Military equipment was also abundant, with flint arrowheads discovered in seven burials, along with axes and daggers. The bone inventory was similarly diverse, encompassing weaponry, hunting tools, tools for skin processing, and fishing implements.

Vessels recovered from the burials exhibited comb ornamentation, often arranged in belts of zigzag and oblique lines.

Ornaments were found in 13 burials. The majority of these ornaments (12 burials) consisted of oval, round, and sub-quadrangular pendants made from green phyllite, greenish and gray-black slate, featuring drawings such as parallel lines and horizontal zigzags. Bone and amber pendants were also present. Notably, two graves within the cemetery yielded copper ornaments (Chizhevskiy and Shipilov, 2021; 2018; Chizhevskiy, 2008).

The chronological attribution of the site is based on 8 new radiocarbon dates and more than 10 previously published determinations (Chizhevskiy and Shipilov, 2021), according to them the burial ground is dated to the period from 4700 to 4200 calBCE. However, the composition of δ15N and δ13C reveals a fish-based diet of the population, and the probability intervals of the dates obtained from the same grave do not overlap. These observations may reflect a freshwater reservoir effect-based offset of the age determinations, made on human bones from Murzikha-2.

*(a) Burial 102/ind.1A (Individual ID I8448), ind.1G (Individual ID I8449),*

**Burial 102** (individual ID 102/ind.1A (I8448), 102/ind.1G (I8449)) yielded a variety of pendants crafted from greenish, gray-black, and white serpentine, ranging in size from large (4.5 cm) to small (1.5 cm). Furthermore, within the same grave, a sculptural depiction of a bird's head, possibly that of a goose or swan, was unearthed (Chizhevskiy, 2008). The individual 1A from burial 102 is dated to 4358-4256 calBCE (5470±25 BP, PSUAMS-8281), and skull G from the same burial is dated to 4547-4373 calBCE (5655±25 BP, PSUAMS-8338).

*(b) Burial 104 (I8450)*

Within **burial 104** (individual ID I8450) three harpoons were found, along with another figurine in the shape of a bird's head (Chizhevskiy, 2008). The burial is dated to 4681-4463 calBCE (5730B±30 BP, PSUAMS-8282).

*(c) Burial 118/ind.3 (I8452)*

From **burial 118** (individual ID I8452), an auroch's (*Bos primigenius*) tooth was recovered, along with a sculptural representation of a phallus. The phallus, made of silicified limestone, measured 26.7 cm in length. Similar models made of wood, antler, and bone are characteristic of the Volosovo culture (Chizhevskiy, 2008). The burial is dated to 4532-4358 calBCE (5615B±30 BP, PSUAMS-8283).

*(d) Burial 128/skull 1(?) (I8456), 128/ind.2 (I8455)*

**Burial 128** (individual ID128/skull 1(?) (I8456), 128/ind.2 (I8455)) contained two axes made of soft limestone, both featuring a perforation. One of the axes had a boat-shaped form with an asymmetrical eyelet, while the other was rhomboid-shaped and crafted from a thin plate. Additionally, a dagger made of slab flint was present in the burial. Of particular significance is the discovery of a fragment of a cattle shoulder blade, representing the first reliable evidence of domesticated animals in the Volga-Kama region. This finding provides evidence of the nascent development of a productive economy in the area (Chizhevskiy, 2008). The burial is dated to 4446-4274 calBCE (5515B±30 BP, PSUAMS-8293).

*(e) Burials burials 94/ind.2 (individual ID I11842), 119 (I8453), 123/ind.1 (I8454), 130/ind.1 (I8457), 131 (I11841)*

For **burials 94/ind.2** (individual ID I11842), **119** (I8453), **123/ind.1** (I8454), **130/ind.1** (I8457), **131** (I11841) we do not have detailed archaeological descriptions. According to available data, the burial rites and grave goods from these burials share the same traits as from the described above. Burial 119 is dated to 4653-4452 calBCE (5700±30 BP, PSUAMS-8335), burial 123 - to 4446-4346 calBCE (5550±25 BP, PSUAMS-8292). A significant freshwater reservoir offset is expected in all dates from Murzikha-2 site.

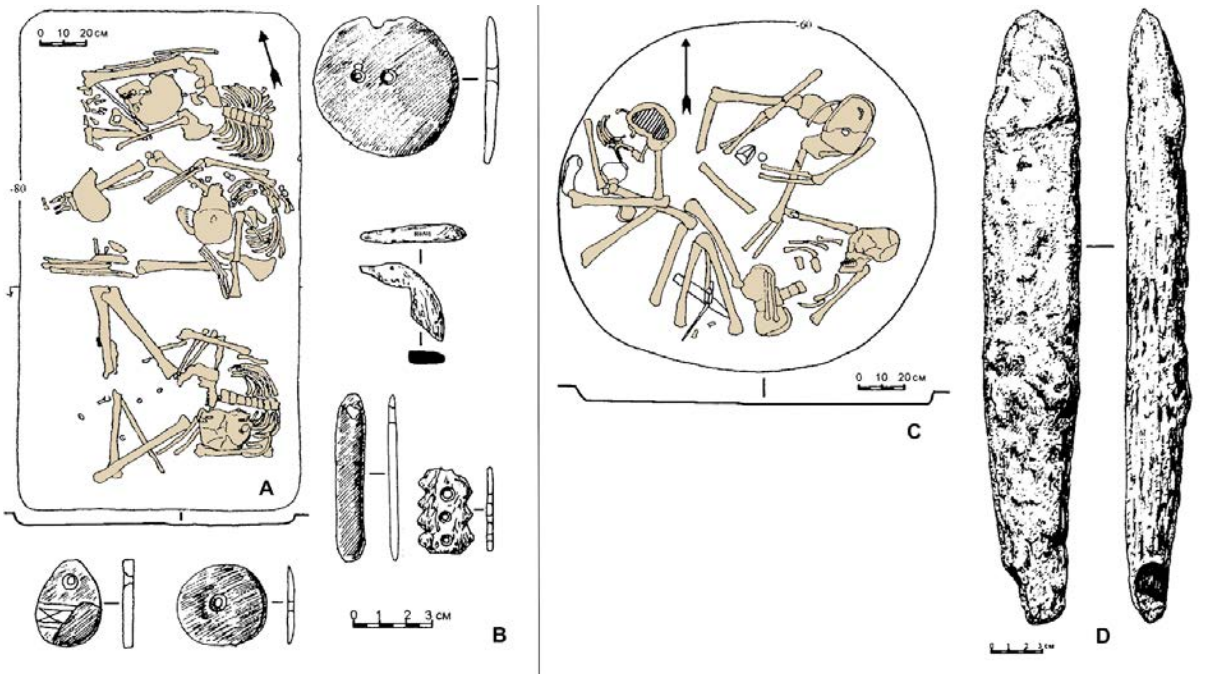

***Fig. S52.*** *Eneolithic burials 102 and 118 at the Murzikha-2 site.* ***A*** *- burial 102;* ***B*** *- grave goods from burial 102;* ***C*** *- burial 118;* ***D*** *- limestone phallus sculpture from burial 118 (Chizhevskiy, 2008, modified).*

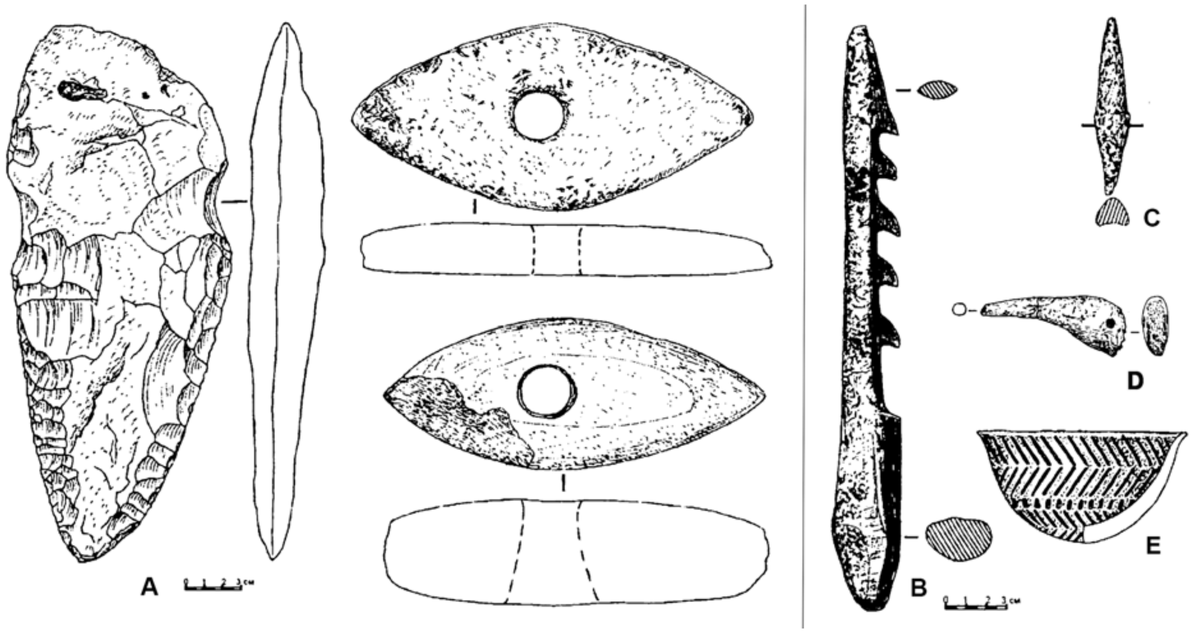

***Fig. S53.*** *Grave goods from Eneolithic burials at the Murzikha-2 site.* ***A*** *- a dagger and stone axes from burial 128;* ***B*** *- harpoon from burial 104;* ***C*** *- antler arrowhead from burial 131;* ***D*** *- antler model of bird's head from burial 104;* ***E*** *- pottery vessel from grave 124 (Chizhevskiy, 2008, modified).*

##### II.E.i.d) Volga Late Eneolithic burials (Russia_LateVolgaEneolithic)

Cultural evolution during the Late Eneolithic in the Middle Volga region was shaped by interaction between the population of steppe origin, characterized by collar ceramics, and descendants of the local Neolithic population. Three distinct cultural groups can be identified, distinguished by their unique ceramic characteristics and origins.

Group 1 comprises sites of the Tok and Chekalino-4 types, where pottery vessels display a gently curved profile, are tempered with crushed shells, and adorned with short to moderately long comb imprints and pits. The cultural traditions of this group likely share a common Volga-Kama Neolithic background, with its development associated with the influence of steppe populations who introduced collared pottery to the region.

Group 2 is represented by sites of the Lebyazhinka type, which later evolved into the "Volosovo-like" tradition observed in Gundorovka. The pottery in this group is tempered with feathers and decorated with moderately long comb imprints and cord impressions. These vessels exhibit similarities to the collared ceramics of the forest-steppe Eneolithic, while also demonstrating typical characteristics of the Volosovo Middle Volga forest zone. The origins of this group can also be traced back to the Volga-Kama culture, but it shows a stronger influence from the Samara culture compared to Group 1.

Group 3 includes sites in the Samara River Valley attributed to the Turganik type and sites in the Sok River Valley associated with the Gundorovka type, also known as the "inner rib pottery" type. Pottery vessels within this tradition feature inner ribs and are tempered with crushed shells and feathers. They are decorated with imprints from fine-toothed combs or plain stamps, as well as hatching and pits. The pottery tradition of this group exhibits similarities to the Sredny Stog, Khvalynsk, and Ivanovka phase of Samara cultures. Vessels with inner ribs are probably derived from those found in the forest-steppe Middle Eneolithic.

Radiocarbon dating places the Late Chalcolithic Mid-Volga pottery within the time frame of 4250–3500 BC, as determined by a series of radiocarbon dates (Korolev, 2021; Shalapinin, 2018).

#### (1) Maksimovka-1 burial ground

The flat burial ground is located on the northwestern outskirts of the village Maksimovka, 480 m east-northeast of the confluence of the Syezhaya and Samara Rivers, a left tributary of the Volga River. The site was discovered in 2017 during salvage excavations, by Viktor Tsibin (Tsibin and Shalapinin, 2018; Korolev and Shalapinin, 2020; Korolev. Et. al., 2021).

##### *(a) Burial 1 (individual ID I8446)*

In **burial 1** (individual ID I8446), 3 skeletons were arranged in a linear formation, extending along the northwest-southeast axis, with numbering from east to west. Unfortunately, because of the paucity of records, we were unable to determine which of the skeletons in grave 1 was sequenced for this study.

*Skeleton 1* was positioned in the southeastern part of the burial. The deceased was laid supine, with the upper body oriented towards the northeast and the legs bent at the knees, pointing westward. The skull fragments were severely damaged and scattered around the chest and pelvis regions. The bones of the chest, arms, and pelvis were also heavily fragmented. The relatively well-preserved right humerus was aligned along the east-northeast to west-southwest axis. A flint scraper was discovered near the left knee of this skeleton (Figure S54., object 8). South of the femur, a stone wringer was found (Figure S54, object 12), and two points were located south of the pelvis (Figure S54, objects 5 and 7).

*Skeleton 2* occupied the central position in the burial and was oriented towards the northeast. The skeleton was supine, with the legs bent at the knees and collapsed towards the west. The skull was situated between the pelvic and chest bones. The shoulders were relatively well-preserved and maintained their anatomical order. The ulna and radius bones, extending along the body, remained intact from the arms. A sandstone pendant was found within the bones of the humerus (Figure S54, 13). A fragment of a point was present in the area of the left shoulder (Figure S54, 6). A flint hole was discovered to the south (Figure S54, 11). A bone tool was located in the vicinity of the pelvis and knees (Figure S54, 10).

*Skeleton 3* was situated in the northwestern part of the burial, with the head oriented towards the northeast. The individual was buried in a supine position, with the legs bent at the knees and falling to the east. The skull was positioned near the chest region, with the crown facing northeast. The shoulder girdle, thorax, and pelvis were in a deteriorated state. The arm bones were extended along the body. Between skeleton 3 and skeleton 2, a fragment of an unadorned vessel wall was discovered.

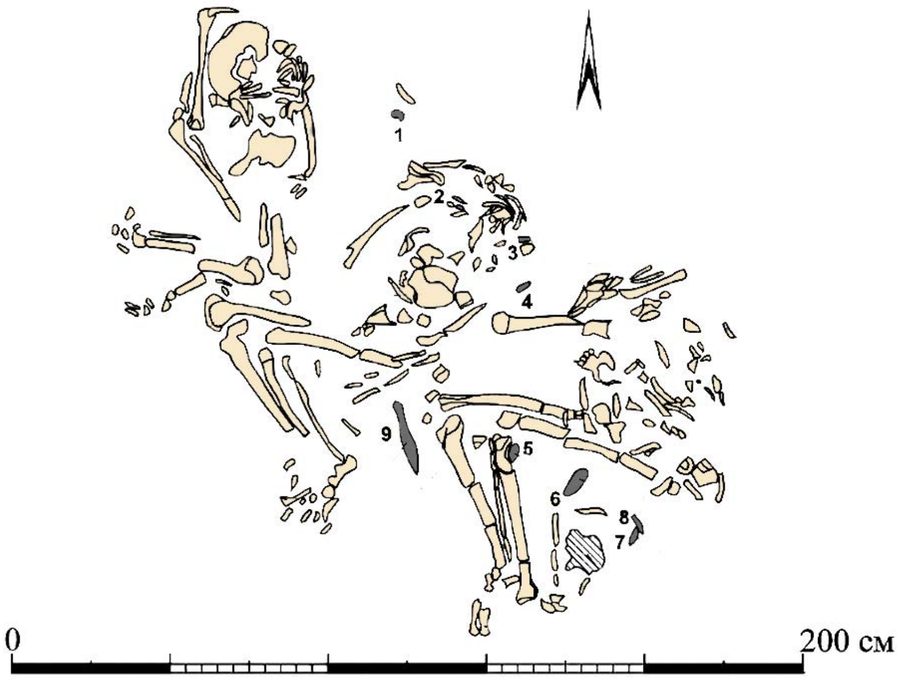

***Fig. S54.*** *Burial 1 at the Maximovka-1 site (Tsibin and Shalapinin, 2018, modified).*

The grave goods found in the burial include a bone tool, a wringer, a sandstone pendant, flint scrapers, piercers, and arrowheads with straight or notched attachments. This burial assemblage shares similarities with the funerary practices observed in the Khvalynsk Eneolithic culture. The presence of arrowheads is a characteristic feature of the Caspian and Altata assemblages of the Volga steppe (Tsibin and Shalapinin, 2018).

The Late Eneolithic burial 1 has been dated to 3946–3711 calBCE (5025±25 BP, PSUAMS-5923).

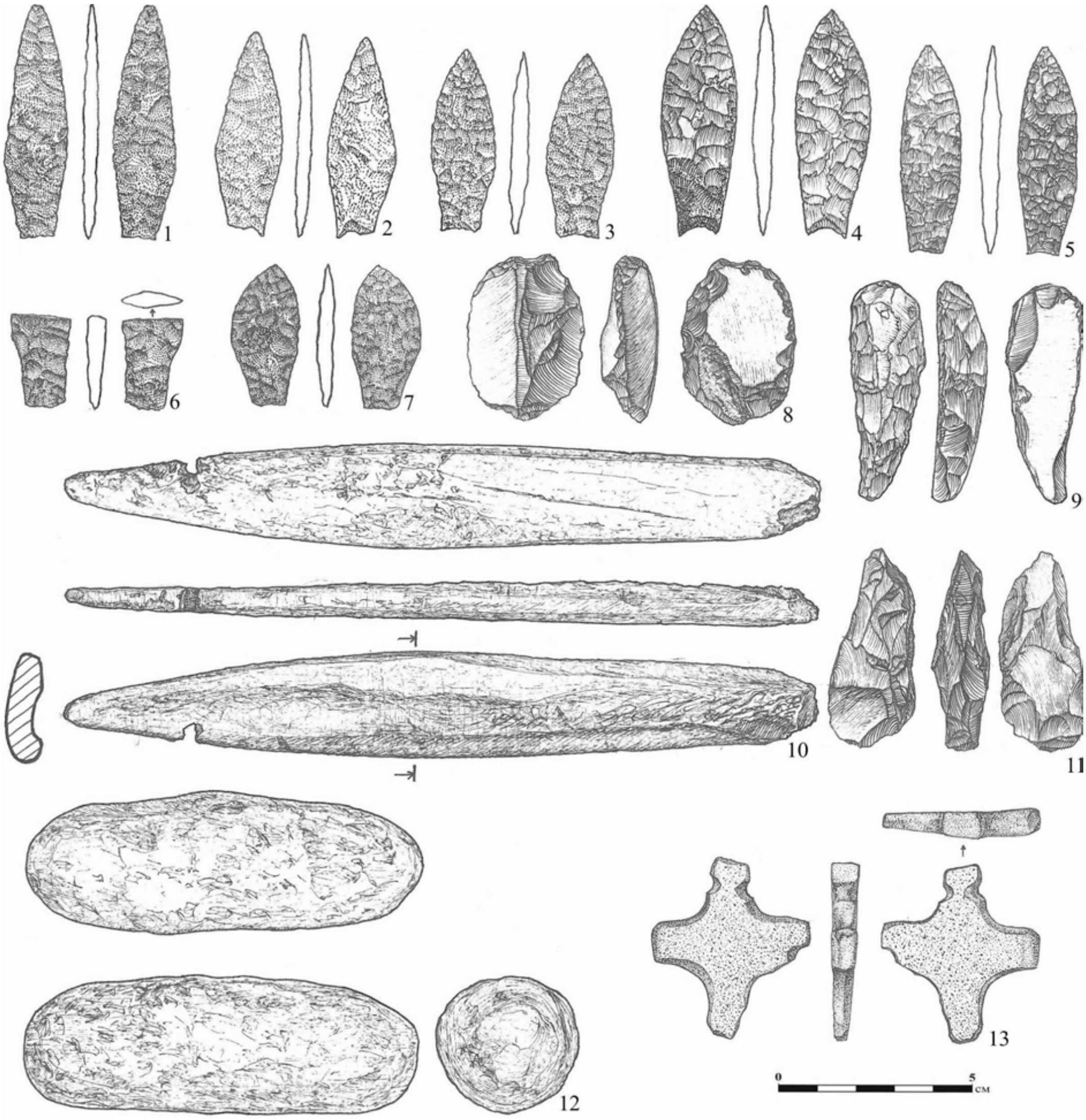

***Fig. S55.*** *Burial 1 at the Maximovka-1 site. Grave goods (Tsibin and Shalapinin, 2018).*

*(b) Unidentifiable burial (I6904)*

**Another burial from Maksimovka-1** (individual ID I6904) lacks detailed archaeological description. It is dated to 4153-3962 calBCE (5205±25 BP, PSUAMS-4255).

#### II.E.ii. Volga-Oka Region

##### II.E.ii.a) Lyalovo Pit-Comb Ware Late Neolithic culture (Russia_UpperVolga_Lyalovo_LN)

The Lyalovo culture was initially defined by Boris Zhukov in 1925 following excavations conducted at the Lyalovo site near Zelenograd in the Moscow region. According to contemporary understanding, the term encompasses Late Neolithic sites within the Volga-Oka interfluve. The Lyalovo tradition is one of several that contribute to the Pit-Comb pottery area, which extended across the forested zone from the

Baltic to the Mid-Volga. Broadly, Lyalovo sites are estimated to date back to 7000–6000 years before present (yBP).

The categorization of the Lyalovo culture primarily relies on distinct characteristics of its pottery tradition. Handcrafted vessels feature rounded (occasionally slightly pointed) bottoms, incorporating crushed granite or sand into clay. The outer surface of the pottery exhibits a continuous zonal ornamentation comprising pits created using a belemnite, various impressions from a comb stamp, and "lunar" impressions. Tool assemblages consist of flint scrapers, knives, and hunting equipment such as arrowheads and spearheads, among others. These tools were produced from flakes, employing double-sided retouching techniques. While bone tools have not been well preserved, some arrowheads and perforators have been recovered. The economy of the Lyalovo culture revolved around hunting (elk, beaver, deer, wild boar, upland and waterfowl species, etc.), fishing, and gathering. Domesticated dogs were also present within the population.

Lyalovo habitation sites are typically located along riverbanks and lakeshores. Extensive elongated houses, both above-ground structures, and semi-dugouts with gable roofs, have been documented. In the vicinity of settlements, ground burials have been discovered, where bodies were interred in a supine position. Grave goods are relatively scarce, primarily comprising tools and ornaments. Four distinct periods have been identified in the development of the Lyalovo tradition, differentiated by specific details of vessel ornamentation: the archaic, early, developed, and late periods.

The Lyalovo culture succeeded the Upper Volga Early Neolithic culture in the Volga-Oka region around 7200–7000 yBP and gradually gave way to the Volosovo Late Neolithic-Eneolithic culture approximately 4100–4000 yBP. Some researchers propose a period of coexistence between the Lyalovo and Volosovo traditions, lasting for several hundred years, during which both populations occasionally occupied the same sites (Gurina and Kraynov, 1996; Engovatova, 2011, 2017; Berezina et al., 2021).

##### (1) Lyalovo phase of the Saktysh archaeological site complex

The Sakhtysh peat bog is situated in the Ivanovo region. It gives rise to the Koika River, which is a left tributary of the Klyazma (Nerl) River. The swamp covers an approximate area of 50 square kilometers and formed in the location of a shallow post-glacial paleolake. A comprehensive scientific investigation of the Sakhtysh archaeological complex commenced in 1962 under the auspices of the Upper Volga expedition, affiliated with the Institute of Archaeology of the USSR Academy of Sciences (now Russian Academy of Sciences, RAS). The project was led by Dmitry Krainov (until 1993) and subsequently by M.G. Zhilin. This endeavor persisted nearly uninterrupted for over forty field seasons, concluding in 2006. During this period, fifteen ancient human habitation sites were identified.

Among the identified sites, five are long-term settlements, namely Sakhtysh-1, 2, 2a, 7, and 8. Additionally, six seasonal sites, including Sakhtysh-3, 4, 9-11, and 14, were discovered, along with four locations yielding individual artifacts, namely Sakhtysh-5, 6, 12, and 13. All of these sites are situated on low, indistinct banks of former river channels and paleolakes, encompassing a segment measuring three and a half kilometers from its source. Throughout the investigations, a total of 149 burials from various epochs and stages of cultural development were unearthed. Chronologically, these burials can be assigned to the Neolithic (20) and Eneolithic periods (128), and the Late Bronze Age (1) (Kostyleva and Utkin, 2010).

###### *(a) Burial 19 (individual ID I8413)*

**Burial 19** (individual ID I8413) was excavated in 1984. The border line of the grave pit was documented at a depth of 55–60 cm. The pit displayed an approximately rectangular shape with rounded corners and a trough-shaped bottom. Its measurements were 2.05 × 0.65 m, with a depth of approximately 20 cm. The well-preserved skeletal remains of a male individual were found lying in a supine position, with the head oriented towards the southwest. The arms were flexed at the elbows, and the hands were positioned atop each other in the waist region. The bones of the right foot were absent, but one of its phalanges was discovered near the right knee. No accompanying grave goods were detected in association with the deceased (Kostyleva and Utkin, 2010). There is no radiocarbon date for this burial yet.

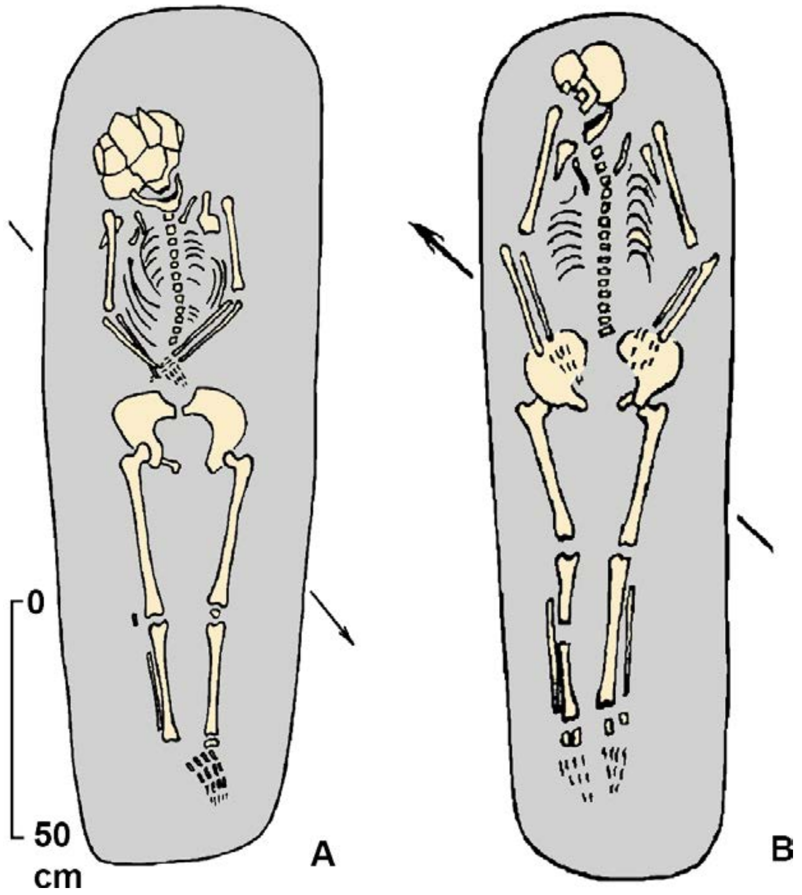

***Fig. S56.*** *Pit-Comb Lyalovo burials at the Sakhtysh-2 site.* ***A*** *- burial 19;* ***B*** *- burial 20 (Kostyleva and Utkin 2010, modified).*

*(b) Burial 20 (I8412)*

**Burial 20** (individual ID I8412) was excavated in 1985. This burial pertained to a young woman and was situated within a an approximately rectangular grave with a trough-shaped bottom, recorded at a depth of 60 cm. The dimensions of the pit measured 1.8 × 0.8 m, with a depth of roughly 20 cm. The infill material consisted of gray loam devoid of any significant findings. The skeleton was well-preserved and positioned in a supine position, facing towards the northeast. The arms were flexed at the elbows, and the hands were clasped on the abdomen. The skull and leg bones exhibited light traces of ocher. No equipment or adornments were present within the burial (Kostyleva and Utkin, 2010). There is no radiocarbon date for this burial yet.

The Lyalovo burial ground at the Sakhtysh-2a site functioned at the initial stage of the development of the Lyalovo culture.

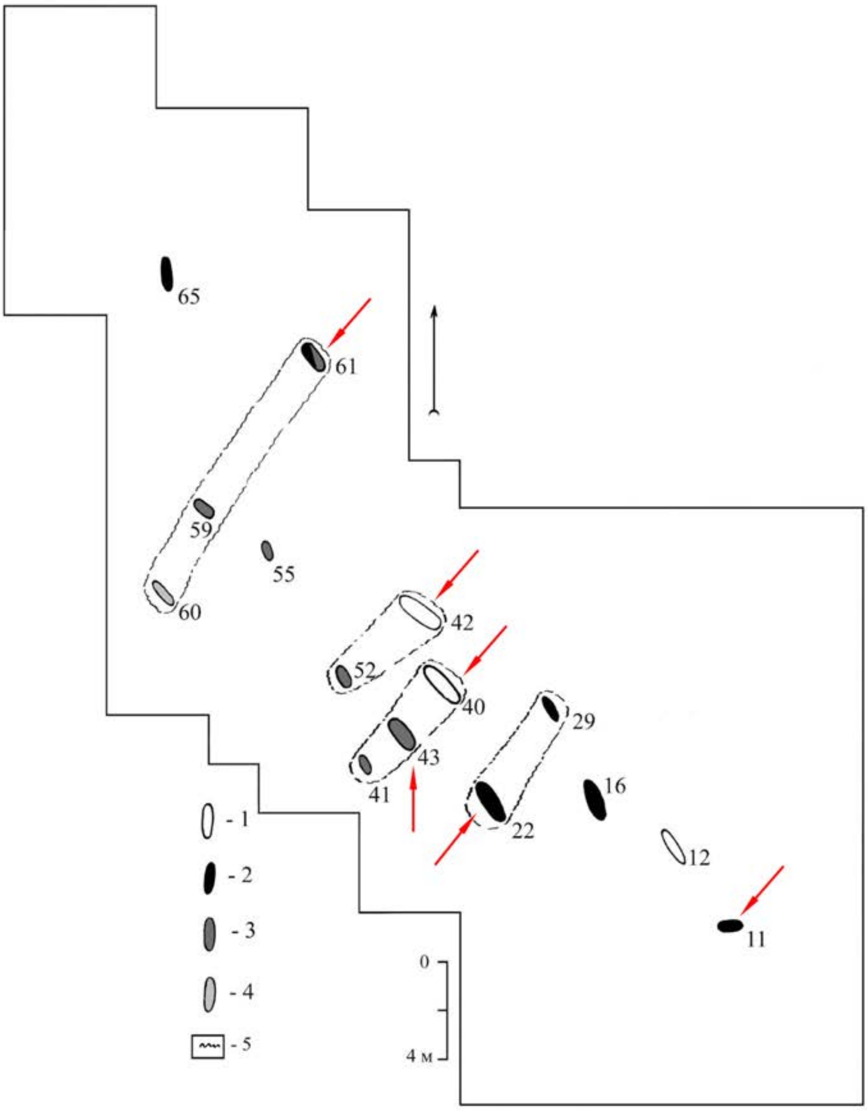

***Fig. S57.*** *Pit-Comb Lyalovo burial site at Sakhtysh-2a.* ***1*** *- male burials;* ***2*** *- female burials;* ***3*** *- child's burial;* ***4*** *- unidentified burial;* ***5*** *- grave rows. Sequenced individuals are marked by red arrows. Burial 11 was attributed to the Lyalovo culture by Kostyleva and Utkin (2010), but we attribute it to the Volosovo culture, according to the crouched position of the body, its west-east orientation, and the genetic profile of the buried individual I8419 (Kostyleva and Utkin 2010, modified).*

The graves are of a rectangular shape with rounded corners or elongated ovals when viewed from above. In cross-sectional and longitudinal views, they appeared as trough-shaped recesses, often displaying an uneven bottom. The dimensions of the graves were determined based on the height of the deceased individuals. No evidence of internal grave structures was discovered within the burials.

The skeletal remains of the interred individuals were remarkably preserved, retaining their anatomical arrangement. However, in some cases, small bones were absent or displaced due to rodent activity. The positioning of the bones suggests that certain corpses were tightly swaddled during burial. In adult individuals, the swaddling primarily encompassed the torso, while the legs remained unbound. This is evident from the unnatural narrowness of the upper portion of the postcranial skeleton, the pronounced compression of the chest, the close proximity of the hand bones to the chest, or their displacement onto the chest area. In burial 43, the child was completely swaddled from head to toe, possibly using animal skins. The sole instance of ocher usage in the funerary practice is discernible in burial 61, where a small area of dark pink paint was observed beneath the female skull.

In single graves, the deceased were laid out in a supine position with their limbs extended. The arm bones were either positioned parallel to the body or slightly flexed at the elbows, while the hands were placed on the abdomen. In burial 42, the left arm was extended, with the hand resting on the pelvic bones, while the right arm was bent at nearly a right angle and positioned on the belt area (Kostyleva and Utkin, 2010).

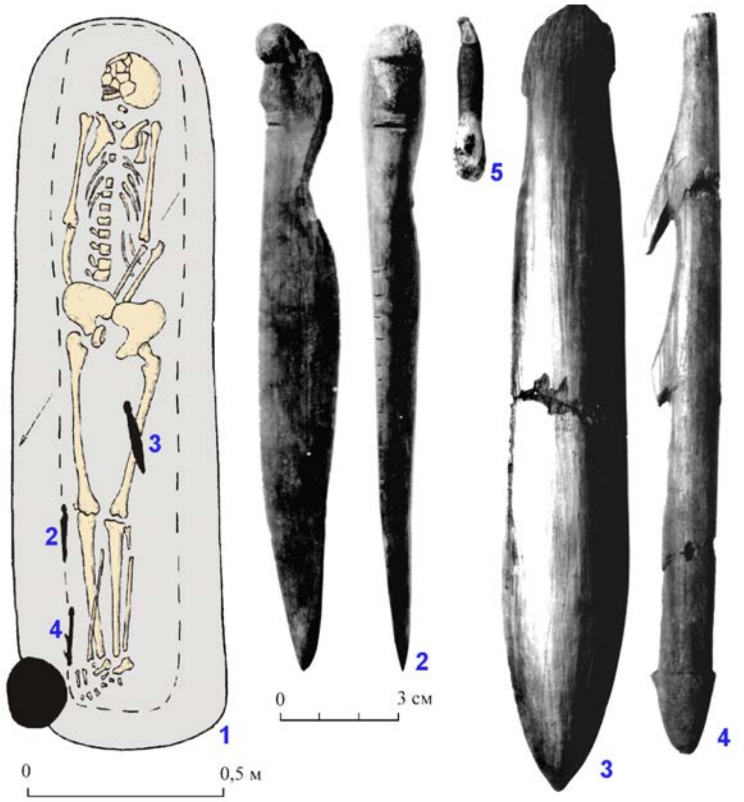

***Fig. S58.*** *Burial 22 at Sakhtysh-2a.* ***1, 6*** *- skeletons;* ***2, 3*** *- daggers;* ***4*** *- harpoon;* ***5*** *- pendant (2–5 - tools made of bone) (Kostyleva and Utkin 2010, modified).*

*(c) Burial 22 (individual ID I8439)*

**Burial 22** (individual ID I8439), consisted of a deceased individual lying in a prone position with an extended body. The arm bones were positioned alongside the body, and the hands were placed under the pelvis. The grave contained several items, including:

1. A wide double-bladed dagger with a lenticular cross-section.
2. A narrow dagger with a hilt adorned with a sculptural depiction of a human face.
3. A harpoon characterized by a short triangular socket, two large beak-shaped teeth, and a point that has been lost over time (Kostyleva and Utkin, 2010). There is no radiocarbon date for this burial yet.

*(d) Burial 40 (I8410)*

In male **burial 40** (individual ID I8410) a short segment from a metapodium of an elk was found parallel to the left humerus, at a certain distance from it (Kostyleva and Utkin, 2010). The burial is dated to

5617-5477 calBCE (6585B±35 BP, PSUAMS-9405).

*(e) Burial 42 (I8411)*

**Burial 42** (individual ID I8411) is dated to 5323-4610 calBCE (6060B±150 BP, GIN-6586).

*(f) Burial 61 (I8416)*

**Burial 61** (individual ID I8416) contained the remains of three individuals. The upper portion of the female skeleton was found in an extended supine position, with the arms positioned alongside the torso and the hands resting on the pelvic bones. Atop the pelvis, the skull of an infant, approximately two years of age, was placed in an elevated position. The remains of the second adult individual, presumably lying prone, were lost between the woman's thighs.

Within burial 61, three bone tools were also unearthed:

1. A piercing tool crafted from a fragment of a narrow animal rib.
2. A knife designed for cutting fish.
3. A long rod-shaped object measuring 19 cm in length and featuring a diameter ranging from 0.8 to 1.4 cm.

Animal tooth appliques, including those from an elk and a bear, were discovered in three burials: 22 (1 ornament), 40 (1 ornament), and 43 (2 ornaments). The placement of these ornaments in relation to the skeletons did not follow a specific pattern (Kostyleva and Utkin, 2010).

The burial is dated to 5633-5483 calBCE (6650B±35 BP, PSUAMS-9406).

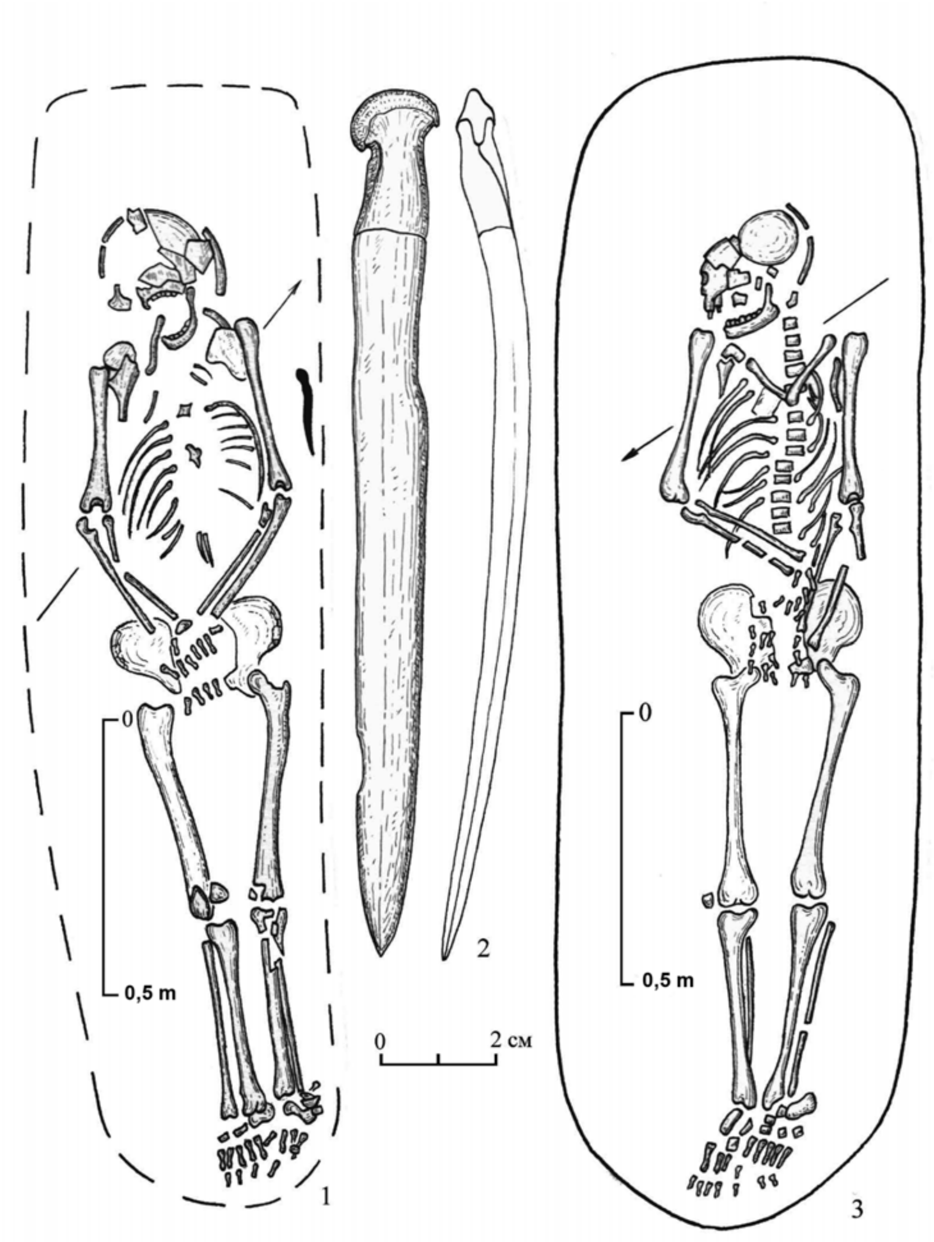

***Fig. S59.*** *Pit-Comb Lyalovo burials at the Sakhtysh-2a site.* ***1*** *- burial 40;* ***2*** *- bone tool (kochedyk) from burial 40;* ***3*** *- burial 42 (Kostyleva and Utkin 2010, modified).*

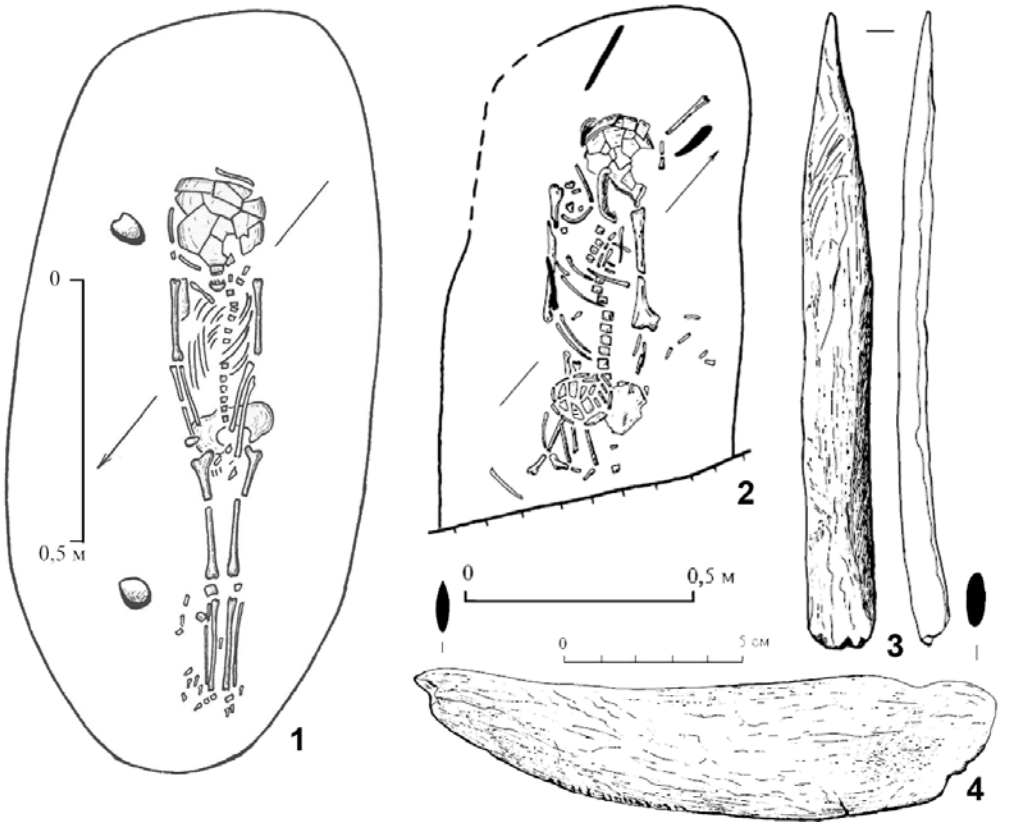

***Fig S60.*** *Pit-Comb Lyalovo burials at the Sakhtysh-2a site.* ***1*** *- burial 43;* ***2*** *- burial 61;* ***3, 4*** *- bone tools from burial 61 (Kostyleva and Utkin 2010, modified).*

##### II.E.ii.b) Volosovo Eneolithic culture (Russia_Volosovo_Eneolithic)

The Volosovo archaeological culture represents a Late Neolithic-Eneolithic tradition found in the forest and forest-steppe regions of the central Russian Plain and the Middle Volga region. It was initially described by Vasiliy Gorodtsov in the early 20th century based on excavations conducted at the Volosovo and Panfilovo occupation sites on the Oka River. These sites revealed similar cultural assemblages of the Neolithic and Eneolithic periods. From the 1920s to the 1950s, there was a debate regarding the chronological position of the Volosovo culture and whether it should be classified as Neolithic or Eneolithic. According to recent understanding, the early Volosovo sites are attributed to the end of the Neolithic period, while the later sites are considered Eneolithic.

The occupation sites of the Volosovo culture include both permanent and seasonal settlements. The typical dwellings were sunken-floor buildings of approximately square shapes, interconnected by passages, and constructed with a frame-pillar structure. Inside the dwellings, a series of open hearths, sometimes up to 11 in number, were located in the center and along the walls. Burials, both individual and collective (involving up to 16 individuals), were found within or near the occupation areas. The bodies were laid in supine, prone, or crouched positions and were often adorned with ocher. Bonfires and bear skulls were also observed between the burials. Grave goods consisted of ornaments made of bone, stone, amber, and occasionally of tools. Pottery vessels had rounded or flat bottoms and were made with the inclusion of shells (in the central Russian Plain) or plant materials (in the Middle Volga region) in the clay. The pottery was decorated on the entire surface or occasionally on the rim. Stone tools were made from flakes or polished stone, while bone tools included harpoons, fishing hooks, arrowheads, and pottery stamps. Ornaments included figurines depicting humans, animals, and birds, crafted from flint, bone, and antler. Amber pendants were widespread even in the eastern regions of the culture.

The economy of the Volosovo culture was based on fishing and hunting. In the later stages, evidence of copper items and indications of copper production was recorded. The Volosovo culture did not remain static throughout its existence. In the 1970s, Alexei Nikitin proposed distinguishing two cultures within the Volosovo tradition: "Volosovo-1" and "Volosovo-2." He associated the first with the Neolithic period, suggesting its origins in northwestern sources from the Baltic and upper Dnieper regions, while he connected the second with the Middle Volga region and attributed it to the Bronze Age. Although this claim was initially criticized, modern researchers tend to support Nikitin's hypothesis. Elena Kostyleva, for example, suggests that the early development of the Volosovo culture can be traced back to the Volga-Oka interfluve and East Baltic Neolithic and to the arrival of population groups from the Eastern Baltic who left burial sites containing amber (Kostyleva and Utkin, 2014). In the late and final stages of the Volosovo culture, noticeable influences from the eastern parts of the East European plain are evident in ceramics and jewelry.

The population of the Volosovo culture is believed by archaeologists to have descended from the bearers of the Pit-Comb Lyalovo and Valdai cultural traditions in the central part of the region, as well as from the Volga-Kama culture population in the Mid-Volga region. During its later stages, the Volosovo was impacted by the Fatyanovo and Balanovo Corded Ware cultures, as well as the Abashevo culture in certain regions. The Volosovo substrate is also suggested to have influenced cultures such as the Chirki, Pozdnyakovo, Fatyanoid, and other cultures of the Central part of the East European plain and the Volga region (Kuz'minykh, 2006; Nikitin, 2021).

##### (1) The Volosovo phase of the Saktysh archaeological site complex

###### *(a) The Sakhtysh-1 settlement, burial 8 (individual ID I8407)*

A group of 11 burials was excavated in an area close to a Late Volosovo dwelling pit.

In **burial 8** (individual ID I8407) the dead person was laid in a crouched position, oriented to the northwest. No grave goods were found (Kostyleva and Utkin, 2010). There is no radiocarbon date for this burial yet.

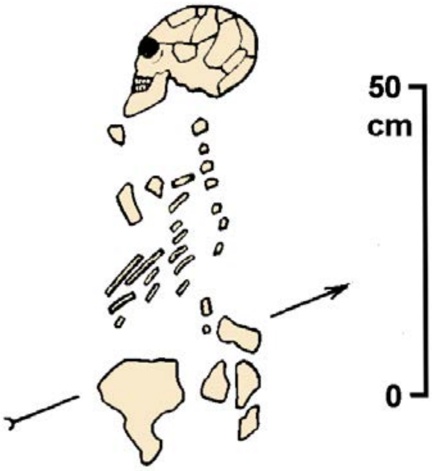

***Fig. S61.*** *Burial 11 from the Sakhtysh-1 site (Kostyleva and Utkin 2010, modified).*

###### *(b) The Sakhtysh-2 settlement, burial 10 (individual ID I8437).*

At the Sakhtysh-2 site, the Volosovo burials were dispersed within the occupational area of the Lyalovo-Volosovo settlement without any discernible order.

**Burial 10** (individual ID I8437) consisted of a skeleton laid in a supine position, with the head oriented to the southwest. The arms were brought together at the elbows behind the back, while the hands were positioned beneath the pelvis. The legs were crossed, with the right leg overlapping the left (Kostyleva and Utkin, 2010). The burial is dated to 3635-2896 calBCE (4540B±160 BP, GIN-6234).

*(c) The Sakhtysh-2 settlement, burial 12 (I12494).*

Collective **burial 12** (individual ID I12494) encompassed four skeletons and was located in the cultural layer at a depth of 40-60 cm. Although the grave pit itself was not discernible, its boundaries were clearly demarcated by an ocher spot. Just above the ocher spot, accumulations of large charcoal pieces were recorded, suggesting the presence of a ritual fire. The individual ID I12494 is labeled as one of the skeletons from burial 12, without specification.

*Skeleton "A"* belonged to a male individual estimated to be 30-35 years old. Positioned in the western part of the ocher spot, this skeleton was laid on the right side in a slightly crouched position, facing the south-southeast. The skull was found in proximity to a severely crushed pelvis. The left leg lacked bones, except for a fragment of the knee joint, while the bones of the right foot were absent, and the thigh was displaced in relation to the lower leg. The left arm extended along the body and fell behind the back, while the right arm was bent with the hand placed near the cervical vertebrae. Beneath the skeleton, a pile of 213 beads made of bird bones was discovered.

*Skeleton "B"*, a female aged 25-30 years old, was positioned in the eastern half of the ocher spot. This skeleton was laid on the right side in a slightly crouched position, with the head facing the north-northwest and the face oriented to the southwest. The legs were spread apart, and the arms were bent at the elbows, with the left arm drawn toward the body and the right arm in the opposite direction. The hands were joined together, and the skull from skeleton "A" rested upon them. Additionally, a layer of thin bird bone beads (215 pieces) was present in the belt area.

*Skeleton "C"* was situated in the northeastern part of the spot, positioned behind the female skeleton and slightly deeper. The anatomical arrangement of this skeleton was severely disrupted, with only portions of the ribs and spine, fragments of an arm and pelvic bones, and a part of a leg (without the lower leg bones and both feet) remaining. The skull was missing. It appears that initially the deceased, estimated to be a child aged 5-8 years old based on bone size, was laid stretched out on the back with the head facing the north-northwest. Later, a slight "shift" toward the northeast caused the skeletal disorder. Three amber pendants, one button, and one bone pendant were found beneath the skeleton in various locations. Additionally, a couple of amber pendants discovered on the eastern outskirts of the ocher spot likely belong to this individual.

*Skeleton "D"* was located between the skeletons of a male and a female, closer to the male and at a slightly greater depth. This skeleton, likely that of a teenager, was entirely fragmented, with only the spine, fragments of the pelvis, and one arm bone preserved. The remaining skeletal parts, including the skull, were missing. Four amber pendants and two buttons were found along and beneath the spine. One button lay on the fragments of the male skeleton's pelvis, another was positioned near the hand of the skeleton, and two pendants were discovered beneath a layer of beads.

It is highly probable that collective burial 12 was formed through a two-stage process. Initially, a teenager and a child were interred in a paired burial, likely positioned supine with their bodies fully extended and their heads oriented to the north-northwest. Subsequently, after a relatively short period of time, the burial was disturbed by the interment of adult individuals. This disturbance resulted in the displacement of certain bones, including the skulls, which were expelled from their original positions. Furthermore, some of the accompanying ornaments fell into the backfill. The adults were buried simultaneously, with deliberate dismemberment observed in the male individual (Kostyleva and Utkin, 2010).

The burial is dated to 3621-3373 calBCE (4705B±25 BP, PSUAMS-9400).

*(d) The Sakhtysh-2 settlement, burial 13 (I12496).*

In **burial 13** (individual ID I12496), a solitary skeleton was found in a supine position. The right hand was placed alongside the body, while the left hand was slightly bent, with the hand resting on the pelvic bones (Kostyleva and Utkin, 2010). There is no radiocarbon date for this burial yet.

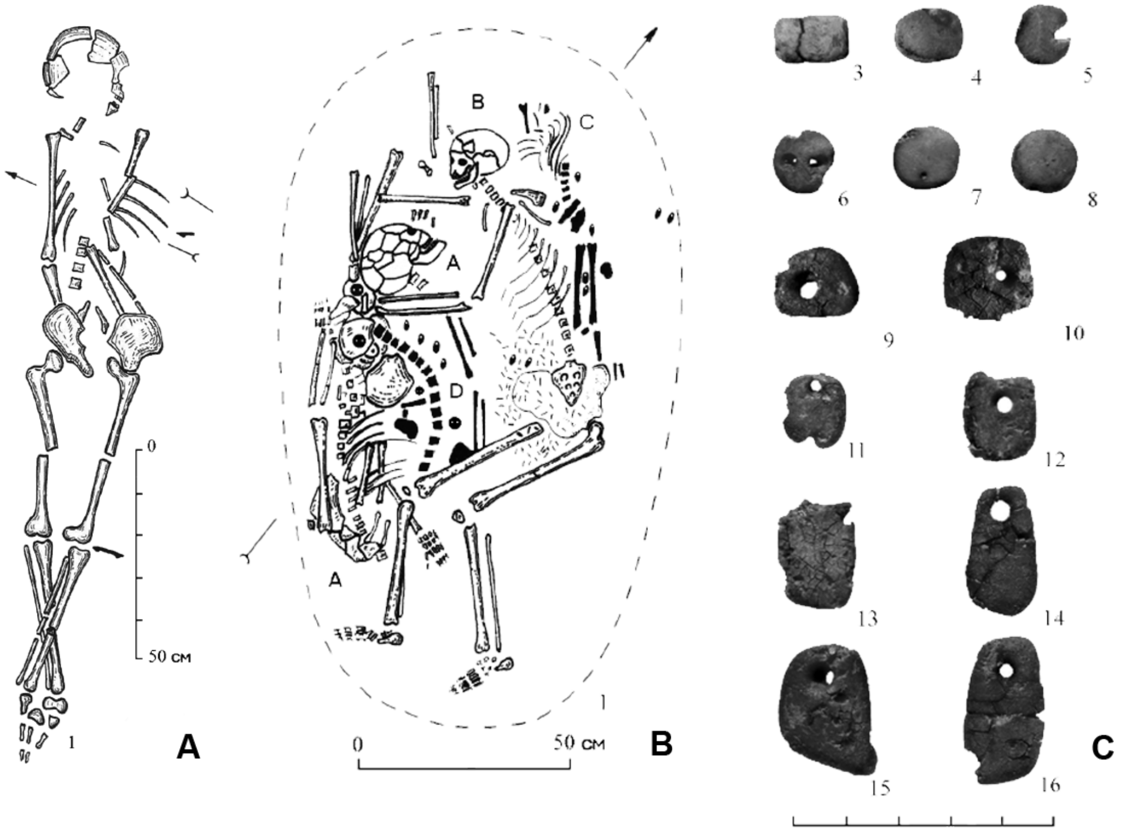

***Fig. S62.*** *Burials 10 (**A**) and 12 (**B**) at the Sakhtysh-2 site, and amber buttons and pendants from burial 12 (Kostyleva and Utkin 2010, modified).*

*(e) The Sakhtysh-2a settlement, burials 9 (individual ID I8420), 11 (I8419), 13 (I12500), 32 (I8404), 33 (I8408), 34 (I12964), 35 (I8414), 39 (I8418), 46 (I8405), 54 (I12962), 56 (I12498), 58 (I8417), and an unidentified burial (I8409)*

All of the burials discovered at the Lyalovo-Volosovo site were situated within the cultural layer of the settlement, characterized by a high concentration of organic material. No discernible grave pits were identified, suggesting that the deceased individuals were likely placed directly on the cleared ground surface and subsequently covered with soil. These Volosovo burials were predominantly individual in nature, and researchers have observed a pattern indicating rows of graves. In close proximity to the burials, two treasure hoards containing amber ornaments and arrowheads were unearthed. Additionally, archaeological interpretations have identified traces of structures believed to be sanctuaries.

In the majority of the burials, the skeletons were found in supine positions, with their arms extended alongside their bodies. Some skeletons exhibited slight bending of the arms at the elbows, with the hands positioned near the abdomen (Kostyleva and Utkin, 2010).

However, **burial 11** (individual I8419) deviated from this norm, as the skeleton was discovered in a crouched position. The deceased individual was laid on their left side, with the left arm extended beneath the body and the right arm bent at the elbow, pressed against the chest. The leg bones also displayed significant bending, as the lower legs were in contact with the stomach, while the upper sections of the legs overlapped the feet, which had shifted due to the pressure exerted by the surrounding soil, leading to ligament rupture. It is possible that the body was originally placed in a swaddled state, indicated by the overall compression of the skeleton, resulting in the bursting of numerous bones unable to withstand mechanical compression (Kostyleva and Utkin, 2010).

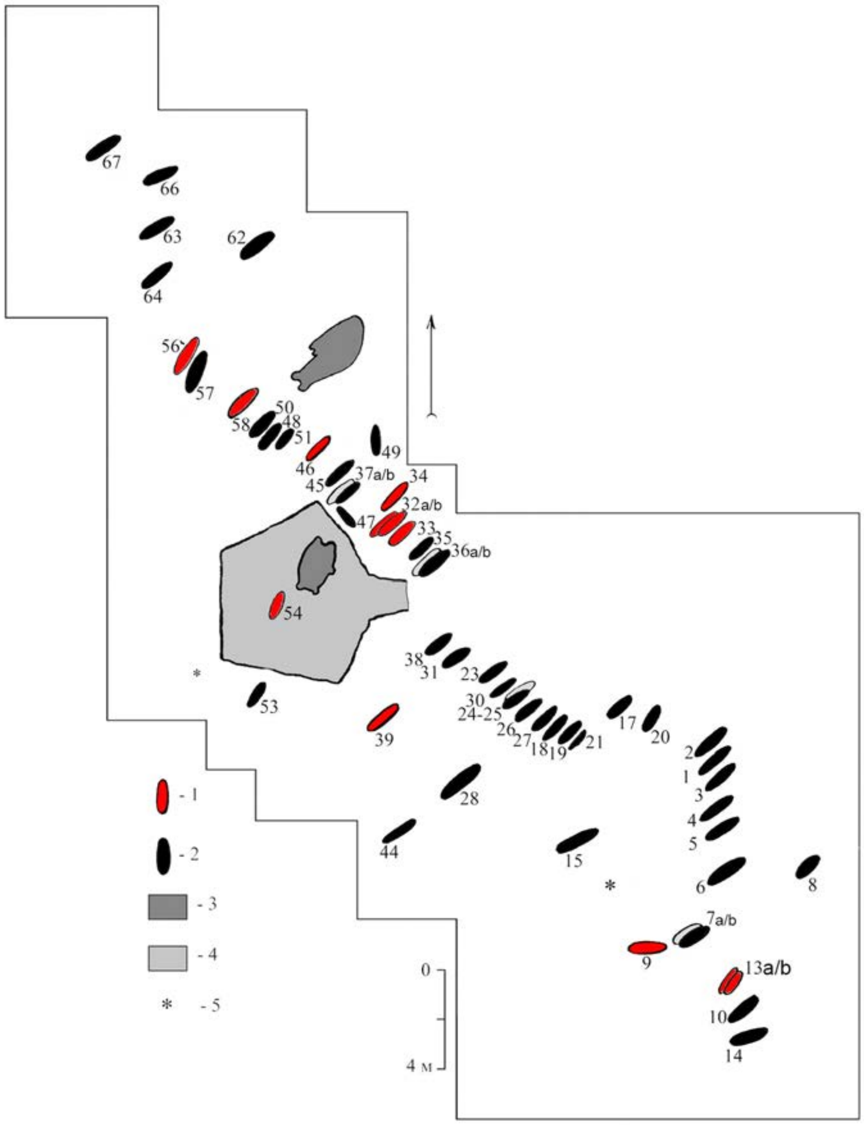

***Fig. S63.*** *Volosovo burials at the Sakhtysh-2a site.* ***1*** *- burials with sequenced individuals;* ***2*** *- other burials;* ***3*** *- ritual (?) pits;* ***4*** *- ritual (?) construction;* ***5*** *- hoards (Kostyleva and Utkin 2010, modified).*

Funerary offerings were not present in all of the burials examined. However, in **burial 39** (individual ID I8418), a beaver's lower jaw was discovered near the right knee, while a bear's tusk was found near the left hip. Various funerary items including jewelry such as beads and pendants, as well as clothing adornments made from materials like stone (serpentine, slate), bones, and animal teeth (bear, elk, beaver, and wolf) were found in **burials 9, 13** (individual ID I12500), **34** (I12964), **39** (I8418), **46** (I8405), **54** (I12962), **56** (I12498), and **58** (I8417). Amber, a highly prized material, was also among the discovered items.

Based on the positioning of the clothing adornments, researchers have been able to reconstruct the attire worn by the Volosovo population. It appears that their costume consisted of a long, closed jacket with a hood. The jacket was intricately embroidered with amber objects, which were found on the chest, the hem's edge, sleeves, and framing of the hood.

In **burial 58** (individual ID I8417), a particularly remarkable discovery was made—a pendant in the form of an anthropomorphic figurine. This unique artifact was located near the neck vertebrae. Crafted from a thin (2 mm) and meticulously polished bone plate, the figurine depicts a standing person when viewed from the front, presenting a full-face image.

The chronological attribution of the Volosovo burials from the Sakhtysh-2a site spans from 4000 to 3000 BCE. Burial 32 is dated to 3952-3792 calBCE (5060B±25 BP, PSUAMS-9086), 34 - to 3630-2904 calBCE (4540B±150 BP, GIN-7276), 35 - to 3283-2049 calBCE (4080B±180 BP, GIN-7273), 54 - to 3768-3649 calBCE (4940B±19 BP) [R_combine: (4900B±30 BP, PSUAMS-9401); (4964B±23 BP, AAR-15051)], 58 - 4342-4248 calBCE (5430B±25 BP, PSUAMS-9088), and the unidentified burial (I8409) is dated to 4047-3961 calBCE (5195B±25 BP, PSUAMS-9087).

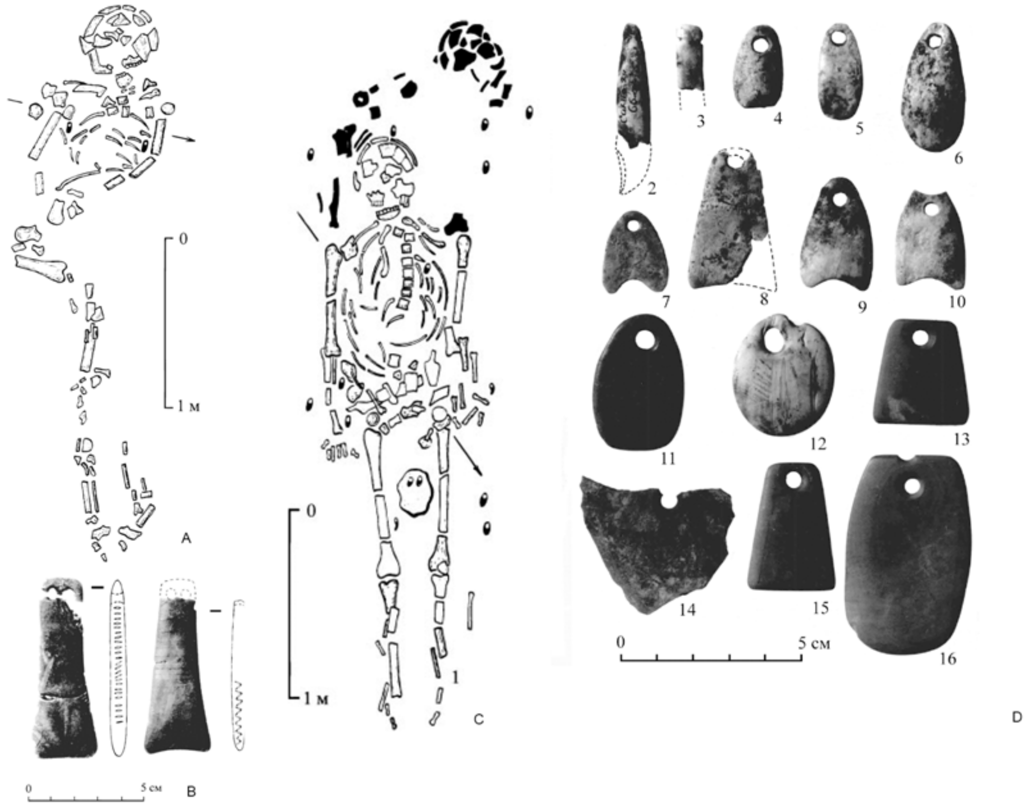

***Fig. S64.*** *Burials 9 and 13a/b at the Sakhtysh-2a site.* ***A*** *- burial 9;* ***B*** *- amber pendants from burial 9;* ***C*** *- burials 13a and 13b;* ***D*** *- pendants made of teeth and amber from burials 13a and 13b (Kostyleva and Utkin 2010, modified).*

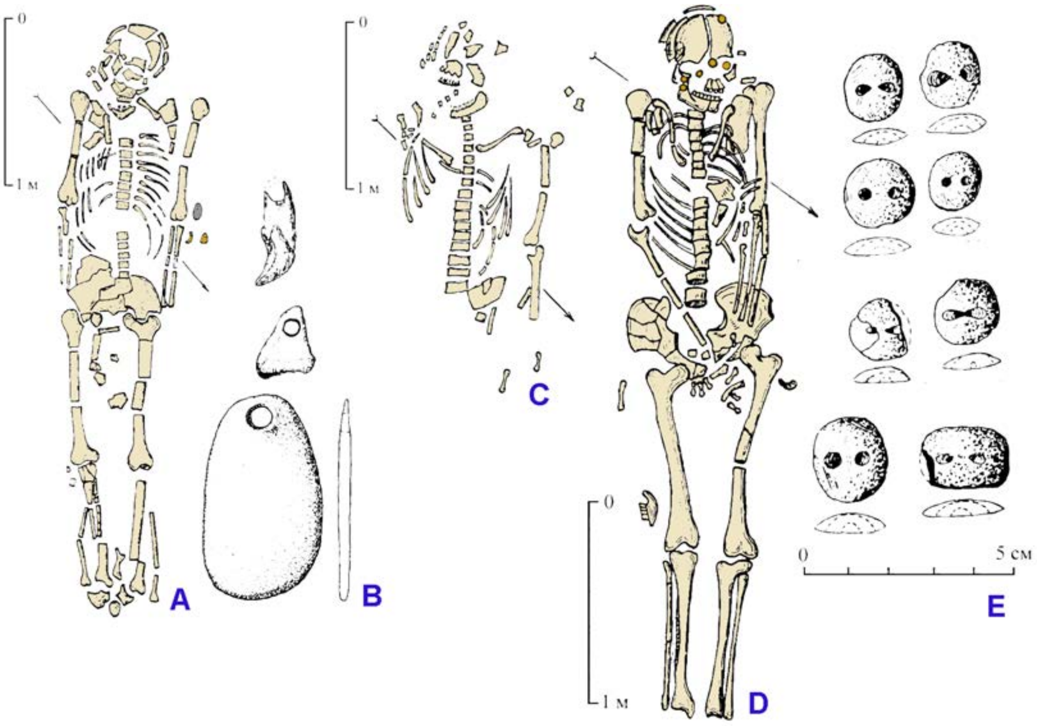

***Fig. S65.*** *Burials 33, 34, 39 at the Sakhtysh-2a site.* ***A*** *- burial 34;* ***B*** *- pendants made of stone and teeth from burial 34;* ***C*** *- burial 33;* ***D*** *- burial 39;* ***E*** *- amber buttons from burial 39 (Kostyleva and Utkin 2010, modified).*

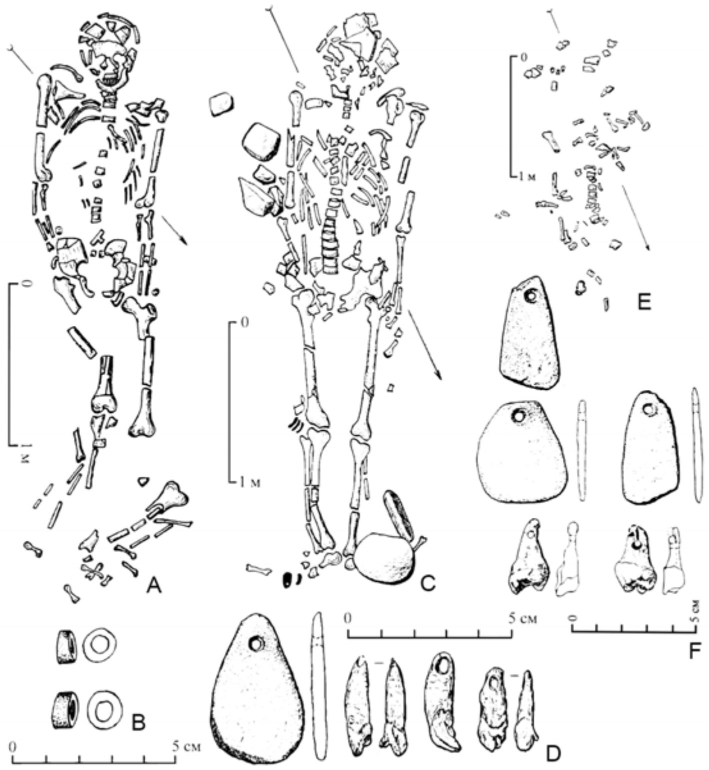

***Fig. S66.*** *Burials 46, 54, 56 at the Sakhtysh-2a site.* ***A*** *- burial 46;* ***B*** *- stone beads from burial 46;* ***C*** *- burial 56;* ***D*** *- pendants made of stone and teeth from burial 56;* ***E*** *- burial 54;* ***F****- pendants made of stone and teeth from burial 54 (Kostyleva and Utkin 2010, modified).*

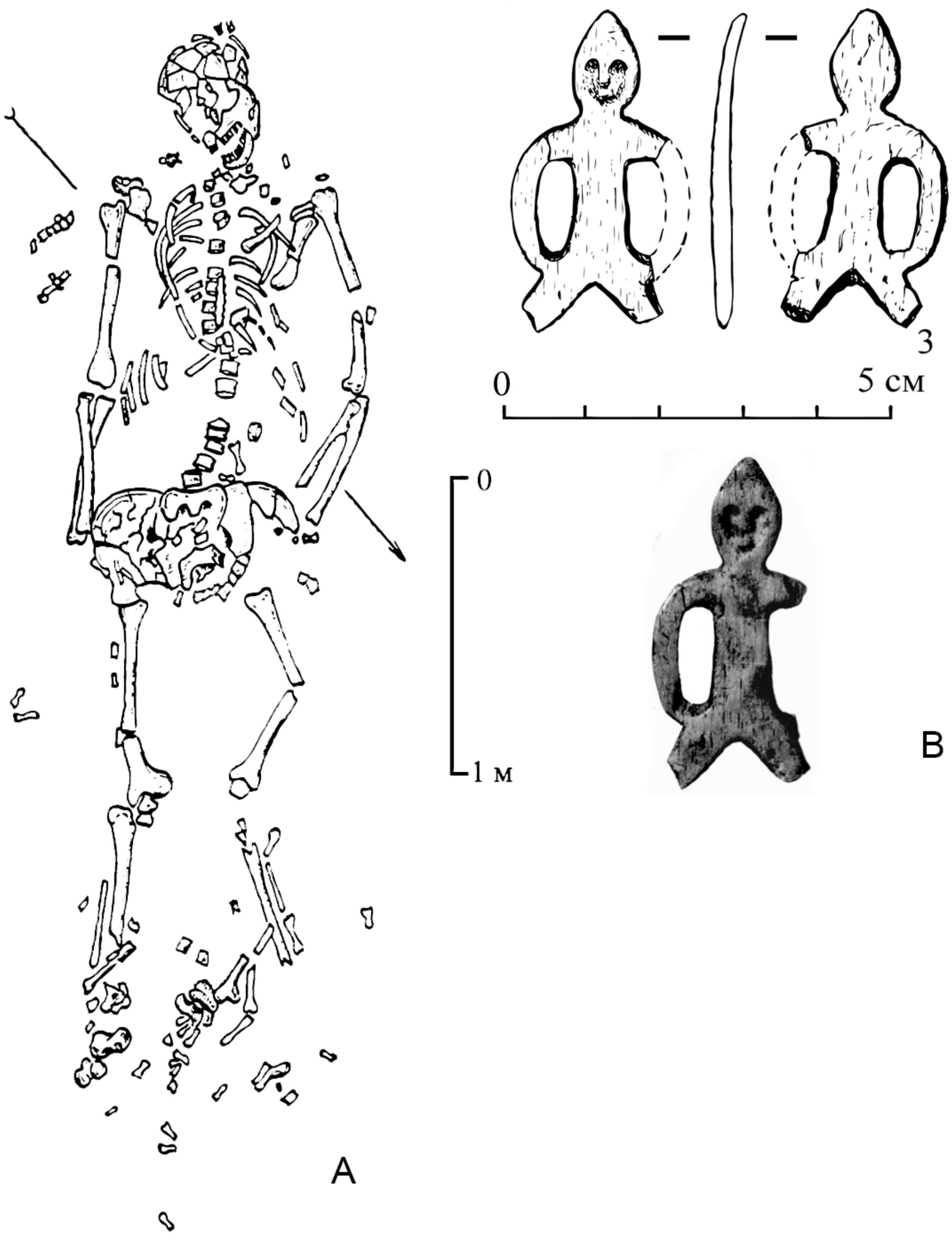

***Fig. S67.*** *Burial 58 at the Sakhtysh-2a site.* ***A*** *- burial 58;* ***B*** *- anthropomorphic figurine pendant from burial 58 (Kostyleva and Utkin 2010, modified).*

##### (2) Imerka-8 occupation site

The Imerka-8 archaeological site is situated on a gentle slope of a sandy dune covered with grass, located on the right bank of the Vad River. The dune reaches a height of 7 meters above the water level. The stratigraphic sequence observed at the site enabled the identification of both early and late Volosovo cultural complexes, which were subsequently overlaid by a layer containing Imerka cultural materials.

The Imerka culture layer superimposes earlier deposits and has yielded a significant number of ceramic fragments, some of which exhibit traces of metalworking activities. These ceramic artifacts are characterized by their distinct originality and lack of analogs in the materials of local cultures. The Imerka culture was first recognized in the mid-1980s through a study of the Imerka-5 settlement by A.A. Vybornov and V.P. Tretyakov. Initially, these materials were attributed to the Volosovo culture, but further research has revealed their later chronological placement, indicating that the Imerka culture succeeded and replaced the Volosovo culture in the Sura and Moksha river basins. It is now understood that the Imerka culture came to an end with the arrival and expansion of populations associated with the Fatyanovo and Balanovo Corded Ware cultures in the Sura and Moksha regions around 2300 BCE.

The Volosovo complex at the site comprises three fully excavated dwellings and an additional five that have been partially explored. Numerous fragments of ceramic vessels and tools have been recovered from these contexts, providing valuable insights into the material culture of the Volosovo people. In addition to the Volosovo cultural remains, pottery associated with the Sredniy Stog culture has also been discovered at the Imerka-8 settlement, indicating possible cultural interactions or influences in the region (Korolev and Stavitskiy, 2021a).

*(a) Unnumbered burial (individual ID I8292)*

The skeletal remains from this **unnumbered burial** (individual ID I8292) are partially preserved and were discovered within the humus layer. The exact boundaries of the grave pit could not be determined. The cultural affiliation of the burial remains unclear, as it could potentially be attributed to either the Volosovo or Imerka population. The chronological placement of the skeleton closely aligns with the dating of animal bones found in the fill of Volosovo structures (Korolev, personal communication).

#### *II.F. Seima-Turbino-period individuals*

##### II.F.i. Overview of the Seima-Turbino phenomenon

The Seima-Turbino assemblage, also known as the Seima-Turbino transcultural phenomenon, refers to a complex cultural development that spread across Northern Eurasia during the transition from the Middle to Late Bronze Age, spanning the last centuries of the third millennium BCE to the first centuries of the second millennium BCE. Artifacts associated with the Seima-Turbino assemblage have been discovered in various locations, ranging from the Sayano-Altai region and Xinjiang to Fennoscandia and the Dniester basin.

The Seima-Turbino assemblage is characterized by a distinctive assemblage of bronze weapons, including spearheads, celts, knives, and daggers with zoomorphic pommels (pommels with animal figurines). These artifacts were crafted using advanced thin-walled casting techniques.

The majority of Seima-Turbino artifacts have been found in burial contexts, with notable sites including Seima, Turbino, and Rostovka. While human burials are infrequently encountered, the focus is primarily on memorial clusters (i.e., cenotaphs, or collections of funerary goods with no associated body). These clusters consist of shallowly buried groups of bronze tools, weapons, and stone arrowheads.

It is believed that the Seima-Turbino cultural complex originated in the region near the Dzhungar Gates. From there, several militarized groups carrying Seima-Turbino traditions migrated eastward, as evidenced by the appearance of Seima-Turbino technologies and artifact forms in cultures such as Elunino, Krotovo, and Odinovo in West Siberia in the Ob-Irtysh basin. From archaeological evidence, it is also believed that the westward spread of Seima-Turbino artefacts also involved forest hunters and fishermen from the Upper Yenisei and Baikal regions, as indicated by the presence of flint and bone arrowheads and knives with bone bases. The influence of Seima-Turbino traditions on the populations of the Sintashta and Abashevo cultures is documented in the Kama and Ural regions. In the forested areas of North Asia, the Seima-Turbino traditions can be observed within the Samus'-Kizhirovo cultural complex (Chernykh and Kuz'minykh, 2015), a trans-cultural analogue of the Seima-Turbino phenomenon with similar but slightly differing bronze artifacts.

#### II.F.ii. Seima-Turbino necropolises (burial sites)

##### II.F.ii.a) Rostovka burial site

The burial ground is located 15 km east of Omsk, on the second left bank terrace of the Omi River, a right tributary of the Irtysh River. The excavation of this site was conducted by V. I. Matyushenko in 1966-1969. Over the course of four seasons, an area of 1376 square meters was excavated, revealing 38 burials and a number of artifact assemblages outside the burials.

Two phases have been identified at the Rostovka burial ground: an early phase associated with the Seima-Turbino phenomenon and a late phase belonging to the Early Iron Age.

The burials of the Seima-Turbino phase were constructed as rectangular pits, ranging in depth from 0.1 to 0.7 meters from the surface, mostly measuring 0.3 to 0.5 meters deep. They were usually oriented from east to west, with slight variations towards the south or north. The burial ritual included: 1) inhumation, sometimes accompanied by partial cremation; 2) off-site cremation; 3) decapitated burials; 4) burials of skulls only. One interesting detail of the burial ritual is the offering of objects to the deceased, left at the edge of the burial pits or placed slightly apart from them. Often, these objects, such as casting molds, are intentionally broken or damaged. In some cases, a ritual is observed where spears and celts are thrust into the bottom, walls, or edges of the burial pit.

Many burials at Rostovka were intentionally and systematically destroyed in the past. In all the disturbed graves, only the bones of the lower limbs, either tibiae and talus bones or only feet, are preserved in anatomical order.

One of the specific features of Rostovka is the presence of burials of metalworkers, which are rare in the Eurasian necropolises of the Bronze Age. At least two such burials have been recorded here (Burial 21 and the burial in square 20Ж). Fragments of casting molds are also found on the surface of the burial ground, although they may originate from disturbed burials.

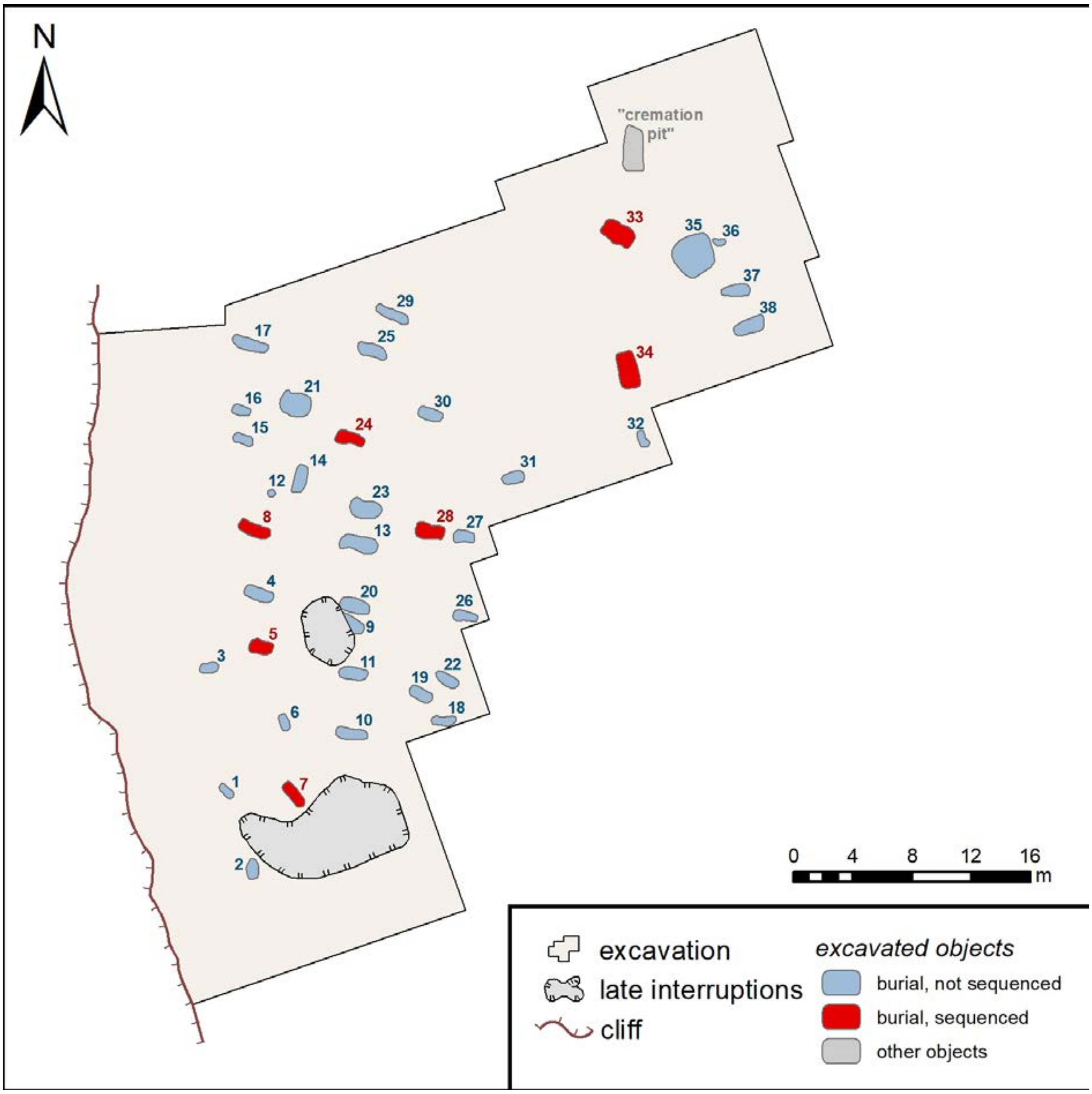

Fig. ***S68***. The Burial Ground at Rostovka, Seima-Turbino burials

The burial ground has yielded a large number of artifacts, including stone arrowheads, knives, plates; bronze celts, spearheads, knives and daggers, awls and chisels, needles; bronze, gold, jade, and lapis lazuli ornaments; bone handles of knives and awls, scales; talc and clay casting molds, as well as fragments of pottery (Chernykh and Kuzminykh, 1989).

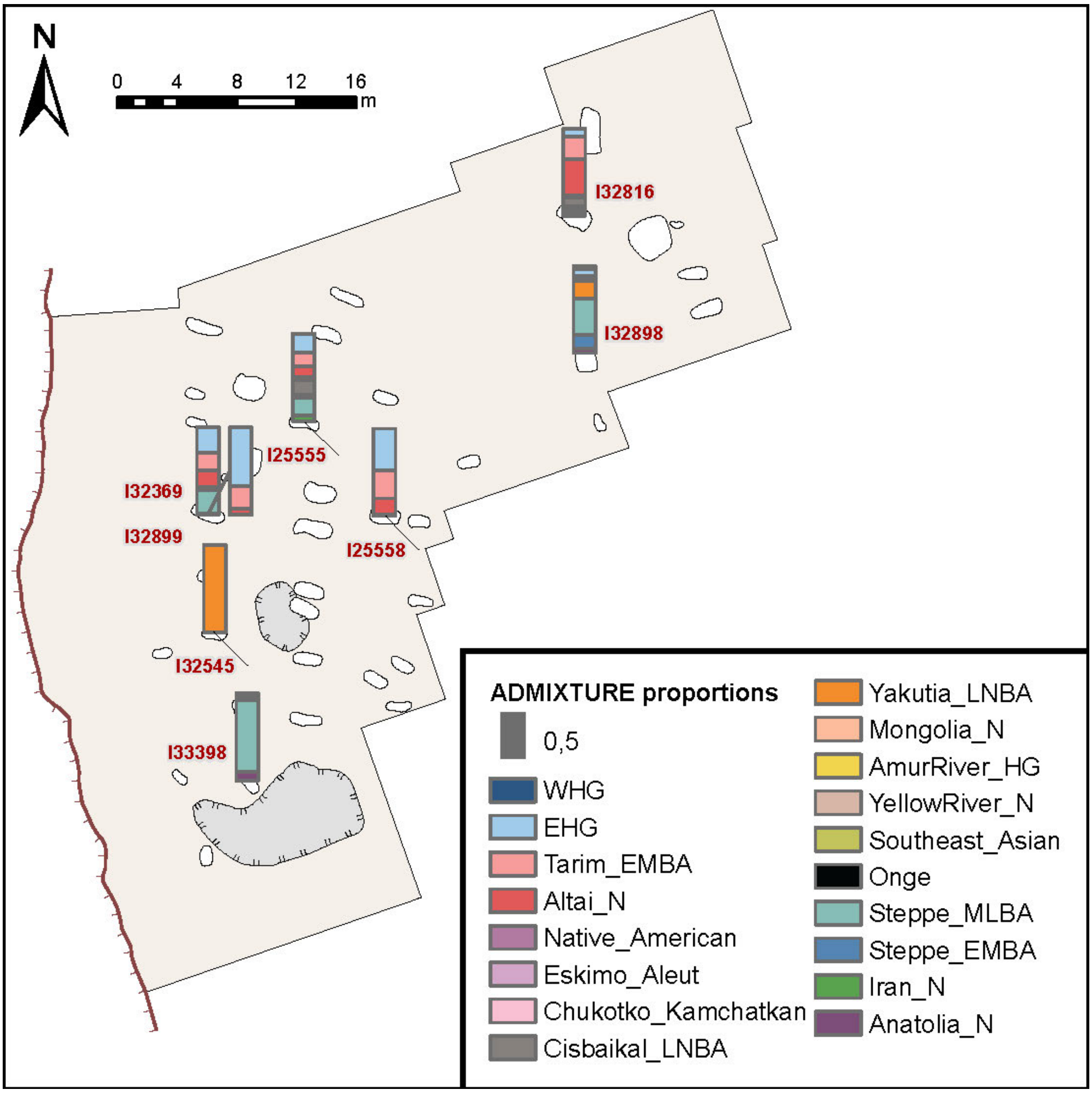

***Fig. S69.*** *The burial ground at Rostovka. The ADMIXTURE proportions of the sequenced Seima-Turbino burials.*

The burials of the Seima-Turbino phase date back to the 22nd-18th centuries BCE. Based on the characteristics of the artifacts and a series of radiocarbon dates, two periods of burials have been identified (Marchenko et al., 2017).

**Table S2**

| Grave number | Genetic ID | Dated material | C14 date lab code | R-date | cal Date BCE | Chronological markers in grave | Period | Reference |
|---|---|---|---|---|---|---|---|---|
| | | human bone | UBA-31379 | 3508±40 | 1938–1700 | | | |

| Grave number | Genetic ID | Dated material | C14 date lab code | R-date | cal Date BCE | Chronological markers in grave | Period | Reference |
|---|---|---|---|---|---|---|---|---|
| 5 | I32545 | tooth | UBA-31396 | 3682±40 | 2197–1951 | spearhead with rhombic cross-section | 2 | Marchenko et al., 2017 |
| | | sample | PSUAMS-7548 | 3585±20 | 2021-1884 | | | |
| 7 | I33398 | sample | PSUAMS-9093 | 3520±20 | 1926-1754 | | | |
| 8 | I32899 | ind. 4 human bone | UBA-31381 | 3709±34 | 2202–1983 | | 1 | Marchenko et al., 2017 |
| | | ind. 4 tooth | UBA-31398 | 3708±30 | 2200–2023 | | | |
| | I32369 | ind. 3 | | | | | | |
| 14 | | | | | | dagger with a longitudinal rib | 2 | |
| 20 | | human bone | IMCES (ИМКЭС)-14C982 | 3634±68 | 2201-1776 | | | Kovtun et al., 2017 |
| 23 | | human bone | UBA-29311 | 3822±32 | 2452–2144 | | | Marchenko et al., 2017 |
| 24 | I25555 | tooth | UBA-29313 | 3564±38 | 2054–1774 | cast mold for spearheads with rhombic cross-section; dagger with a longitudinal rib, spearhead with rhombic cross-section (found in associated assemblage) | 2 | |
| 27 | | human bone | UBA-31383 | 3655±40 | 2141–1918 | | | Marchenko et al., 2017 |
| | | tooth | UBA-31399 | 3635±40 | 2135–1896 | | | |
| 28 | I25558 | sample | PSUAMS-12587 | 3620±20 | 2035-1900 | | | |
| 29 | | human bone | IMCES (ИМКЭС)-14C916 | 3888±105 | 2832-2033 | | | Kovtun et al., 2017 |
| 33 | I32816 | human bone | UBA-31382 | 3640±31 | 2133–1919 | | 1 | Marchenko et al., 2017 |
| 34 | I32898 | human bone | UBA-31380 | 3646±39 | 2137–1919 | | 1 | Marchenko et al., 2017 |
| | | tooth | UBA-31397 | 4066±50 | 2862–2473 | | | |

**Table 2**. *Relative and absolute chronology of the sequenced Rostovka burials.*

#### (1) Burials at the Rostovka Seima-Turbino necropolis

##### *(a) Burial 5 (individual ID I32545)*

**Burial 5** (individual ID I32545) was situated at a depth of 30 cm from the modern surface. It appeared as a rectangular patch of gray earth filled with charcoal and burnt soil, with dimensions measuring 65x150 cm. The orientation of the burial was east-northeast–west-southwest. The skeleton of a presumed male, approximately 25 years old, was found lying on his back with the head facing east. The preserved bones include the ulna and humerus of the arms, the femur and lower leg of the left leg, as well as the pelvic bones, although they were displaced. The skull, chest bones, spine, and phalanges were in a very poor state of preservation. Within the grave, several artifacts were discovered, including a bronze socketed spearhead, four jasper flakes, and stone arrowheads. The spearhead tip was likely not placed within the grave but left after most of the filling took place, and it was found stuck into the wall of the grave.

##### *(a) Burial 7 (individual ID I33398)*

**Burial 7** (individual ID I33398) was located at a depth of 40 cm from the modern surface. It was observed as a rectangular patch of gray earth filled with charcoal and burnt soil. The dimensions of the burial were measured to be 70x220 cm, with the west-northwest–south-southeast orientation. During the excavation, a strip of charcoal was uncovered at the level of the initial filling, running from the northwest to the southeast. Additionally, pieces of charcoal were found in separate clusters, approximately 20 cm above the bottom of the grave, suggesting the presence of remnants of a burnt wooden ceiling or lining. Among the findings within the burial, on the first layer, a roughly worked stone artifact, a fragment of a deer antler possibly used as a handle, and an ellipsoid-shaped bone object with holes at the ends were discovered. The skeletal remains of an adult male, estimated to be around 40 years old, were found in a bent position on the left side with the feet pointing to the west. However, the preservation of the skeleton is poor, with only the left and right femur and tibia intact. Disorganized ribs and a humerus were located beneath the left thigh, while the lower jaw was found under the right ulna. The cluster of finds near burial 7 included an asymmetric quartzite knife, several fragments of bone armor, a fragment of a stone object, and a small stone.

##### *(a) Burial 8, individuals 1, 2, 3 (individual ID I32369) and 4 (individual ID I32899)*

**Burial 8** was situated at a depth of 40 cm. It was identified as a rectangular patch of gray mixed earth, slightly infused with charcoal. The orientation of the burial was noted as east-northeast–west-southwest. Within the grave, the central and northeastern halves contained preserved burnt human bones and skull fragments, indicating a cremation event in this section. Additionally, remnants of burnt trees were discovered in the central part, arranged in two rows within the grave's fill and along its walls. Charcoal and calcined soil were also present in various areas of the burial. It is likely that the deceased's body was cremated on a wooden platform, as the remains of burnt wood and coals were found beneath the burnt human bones.

During the excavation, five stone arrowheads were unearthed at different locations within the grave, specifically at the level of charcoal and human bones. Following the removal of the burnt skeleton, which appeared to belong to a child *(individual 1)*, a clearer picture emerged.

At the bottom of the grave, positioned on their back in an extended posture with the head facing east, was another *skeleton (individual 2)* in reasonably well-preserved condition, presumably that of an adult male. The major bones were intact, although the skull was missing. This particular skeleton was associated with an accumulation of six stone arrowheads and two knife-like plates located in the southwestern part of the grave near the left knee. All arrowheads were oriented towards the west, suggesting the presence of a quiver in that area. Additionally, the same skeleton included a bronze knife and a bronze awl with a bone handle, found in the pelvic region. The knife was pointing southward, while the awl was oriented towards the north.

On top of the second skeleton in the southwestern portion of the grave, a third skeleton, ***individual 3*** (individual ID I32369) was positioned, likely that of a male individual. The skull of this individual was placed above the left half of the second skeleton's pelvis, with the back of the head facing upwards. Adjacent to the skull, the handle of a bone knife was preserved. The pelvic bones (two of them) from this third skeleton were discovered in the eastern half of the grave.

***Individual 4*** (individual ID I32899) exhibited a specific arrangement of remains. Above the leg bones of the second buried individual, situated in the western half of the grave, lay the skeletal remains of a fourth individual, likely a woman aged 40-45 years. The preserved elements from this individual included the skull with the face upward, the left tibia, the right half of the pelvis, the right ulna, and the left clavicle. Following the removal of all the skeletons, a patch of mixed earth was discovered at the bottom of the grave in its eastern half. This patch was deepened by 15 cm from the bottom of the grave. Within the deepest section of this patch, three bronze objects were found. The first was a forked tip of a stake, above which was another similar tip with a spike. On top of these tips lay a bronze one-eared celt adorned with a ladder ornament and hatched triangles, measuring 14 cm in length. Notably, all these bronze items were positioned with their blades facing west. The accumulation of finds associated with burial 8 included the following items: eight stone flakes, a bronze socketed spearhead featuring a long open socket deeply embedded, with its point facing downward into the mainland, a fragment of a spearhead crafted from shale, a chert plate resembling a knife, a blank for a knife made of diorite, and a pot.

*(b) Burial 24 (individual ID I25555)*

**Burial 24** (individual ID I25555) was situated at a depth of 15 cm from modern surface and appeared as a dark patch of mixed earth, measuring 70x200 cm. Pieces of burnt soil were present within the grave's fill. The remains of the deceased were discovered in the grave, albeit in a poorly preserved state. The western corner contained fragments of the thighs, pelvis, tibia, and right hand. In the eastern half of the grave, the lower jaw of a woman, approximately 40 years old, was found. The inventory associated with burial 24 comprised several items. These included a stone arrowhead with a straight base, a fragment of a clay mold used for casting a spearhead, a bronze chisel, and a bronze plate knife. Furthermore, the accumulation of finds within the grave consisted of a flake of clay slate, a bronze forked tip of a stake featuring a spike at the base of the feather. The sleeve of the stake was equipped with an eyelet and four convex rollers. Additionally, a badly corroded bronze two-eared celt was present, featuring a round sleeve and a slightly convex blade without ornamentation. Moreover, a bushed bronze spearhead, cast in a double-leaf mold and possessing two through holes in the bushing, was also part of the assemblage. All these artifacts were found together, in close contact with each other, with their points facing downward and stuck into the ground. In proximity to these items, a fragment of an arrowhead made of clay shale and a fragment of rock crystal were also discovered.

*(c) Burial 28 (individual ID I25558)*

**Burial 28** (individual ID I25558) was situated at a depth of 60 cm from modern surface and appeared as a patch of dark mixed soil, measuring 80x180 cm. Its orientation was from east to west. Within the grave, the remains of a man aged 30-35 years were discovered. The preserved bones included the leg bones, which were found in an orderly manner in the western half of the grave, as well as fragments of the skull, ribs, and the left ulna. It appears that the deceased individual was laid with their head facing east. Notably, the position of the bones in the right foot suggests that the foot was turned towards the left heel during the person's lifetime. Curiously, one of the bones ended up on top of all the other bones, although no evidence of disturbance in the grave was observed at this location.

The collection of artifacts associated with burial 28 consisted of the following items: three knife-shaped flint blades, a bronze tetrahedral awl, a flint blade exhibiting traces of secondary processing, six flint flakes, one fragment of ceramics, an accumulation of ceramics, a fragment of a knife-shaped flint blade, two additional flint flakes, and two fragments of a single object displaying a transverse groove on one side and a longitudinal groove on the other.

*(d) Burial 33 (individual ID I32816)*

**Burial 33** (individual ID I32816) was located at a depth of 25 cm from modern surface. It was identified by a patch of dark coaly earth with an indefinite shape. The dimensions of the grave were measured as 25x200 cm. At a distance of 30 cm from the southeastern wall, a forked spear was discovered, with its tip inserted into the ground. Within the southeastern half of the grave, a significant accumulation of bone armor plates was found. These plates were in a disordered pile, and among them, a portion of a woman's skull, estimated to be aged 20-25 years, was also discovered. The lower layer of plates rested at the bottom of the grave, extending in some cases towards the bottom or the wall. In the northwestern half of the grave, tibia bones and a tibia were found. Based on their position, it can be inferred that the skeleton was lying on its back in an extended position, with the head facing southeast. Among the armor, a golden ring was found closer to the south-eastern wall, while a second golden ring of larger diameter was found closer to the center of the grave. Two abrasives were located in the southern corner of the grave. Within the armor, a fragment of an unfinished bone puncture was also discovered. In the center of the grave, a stone arrowhead with a straight base was present. The filling material of the grave yielded additional artifacts, including the second half of a bone puncture, another stone arrowhead with a straight base, and a bone knife handle with a hole for the blade. The handle was adorned with a delicate pink geometric ornament. The collection of findings within grave 33 also included a flint flake, fragments of ceramics, a stone flake, two more ceramic fragments, fragments of animal bones, another stone flake, additional ceramic fragments, an accumulation of ceramic fragments, and further fragments of animal bones.

*(e) Burial 34 (individual ID I32898)*

**Burial 34** (individual ID I32898) was situated at a depth of 65 cm from modern surface. Prior to its discovery, an indistinct spot was identified at the lower horizon of the chernozem. Within this spot, a flake and two stone arrowheads with straight bases were found. The dimensions of the grave were 90x280 cm, and it was oriented from southeast to east-northwest. Inside the grave, the well-preserved remains of a man, aged between 25 and 30 years, were uncovered. The individual was lying on his back in an extended position with his head facing south-southeast. Adjacent to the right elbow of the deceased, a bronze celt was discovered, pointing towards the eastern wall. The celt's sleeve contained remnants of a wooden handle. Beneath the celt, a substantial bronze spear with a forked rod and a hook below a feather was found. Golden earrings in the form of rings with open edges were situated near the temples of the skull. On the pelvic bones, there was a lamellar bronze knife with traces of a wooden scabbard. Adjacent to the pelvic bones on the left side, a stone arrowhead with a straight base was positioned, while two similar points were located near the left knee joint. A bone puncture was observed near the left hand. On the left side of the skull, a bronze point-awl with a bone handle and the remains of a bronze artifact in the form of an awl inserted at an angle into the bone handle were discovered. In the southern corner of the grave, to the left of the skull, a large fragment of ceramics was recovered from the filling material of the grave, near the level of the frontal part of the skull.

##### II.F.ii.b) Satyga-16 burial ground

The burial ground is located on the northern shore of the Satyginsky Tuman Lake, a little to the east of the place where the Gorodishe River flows into it. The Evra River flows into the lake from the northwestern side, and the Sumpanya River from the south. Satyga-16 is a multi-phase site, confined to a small oval-shaped hill, elongated in the latitudinal direction by 40-45 m with a width of up to 20 m, located at the edge of the lake terrace. The height of the hill is up to 2.5-3.0 m from the water's edge (as of July-August 2001), the slopes are gentle.

The site was excavated in 1986 by O. N. Korochkova, V. I. Stefanov, A. A. Pogodin, and E. M. Bezprozvanny. In total, during three field seasons, an area of 529 sq. m and 41 burials of the Bronze Age were investigated. The Satyga-16 burial ground has been explored for the most part. A feature of burial grounds of the Seima-Turbino type, which is present at Satyga-16, is abundance of finds in inter-grave space (Trufanov, 2011).

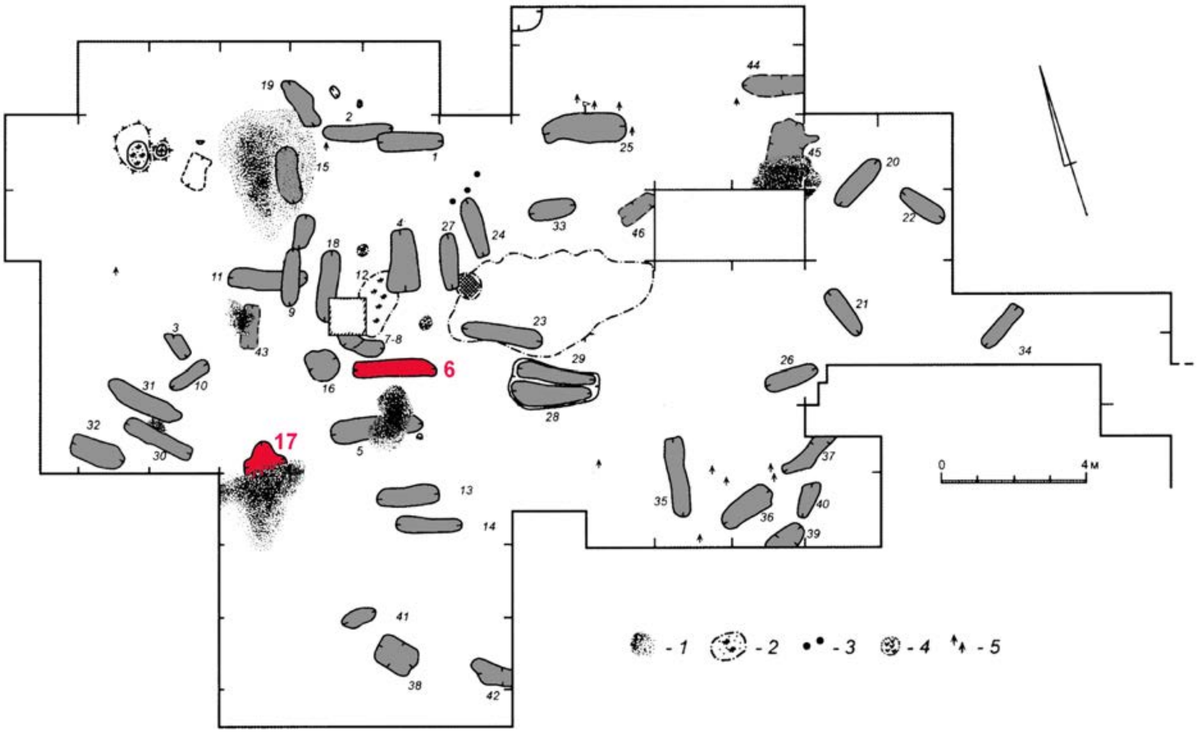

***Fig. S70.*** *Cemetery Satyga-16 at the Satyga-16a occupation site.* ***1*** *- uprootings;* ***2*** *- calcinated bones;* ***3*** *- postholes;* ***4*** *- ocher;* ***5*** *- arrowhwads. The sequenced burials are labeled in red (Trufanov et al., 2011, modified).*

#### (1) Burials at the Satyga-16 Seima-Turbino necropolis

##### *(a) Burial 6 (individual ID I32552)*

**Burial 6** (individual ID I32552). The grave pit is subrectangular in plan with rounded corners, 220x46 cm in size, oriented along the west-east line. The eastern wall is cut by a small rounded pit, 35 cm in diameter. The walls of the grave pit are even, gently descending to the bottom, noted at a depth of 10-18 cm in the mainland. The eastern half of the pit is 5-10 cm deeper than the western one. The filling is brown sandy loam, covering a thin (3-6 cm) layer of pale yellow sandy loam in the bottom part, saturated with fine coals and remains of wood decay. Two carbonaceous strips (134x12x2-10 cm) located parallel to each other were unearthed 10-20 cm above the bottom along the long axis of the grave. Twelve fragments of pottery were found above the burial. At the bottom of the pit near the western wall, a skull and teeth of an adult male 30-40 years old were found. Next to the skull lay a bronze dagger in a birch bark sheath. Small fragments of ceramics were found in different places of the grave (Trufanov, 2011).

Burial 6 is dated to 2571-2348 calBCE (3960±25 BP, PSUAMS-12559).

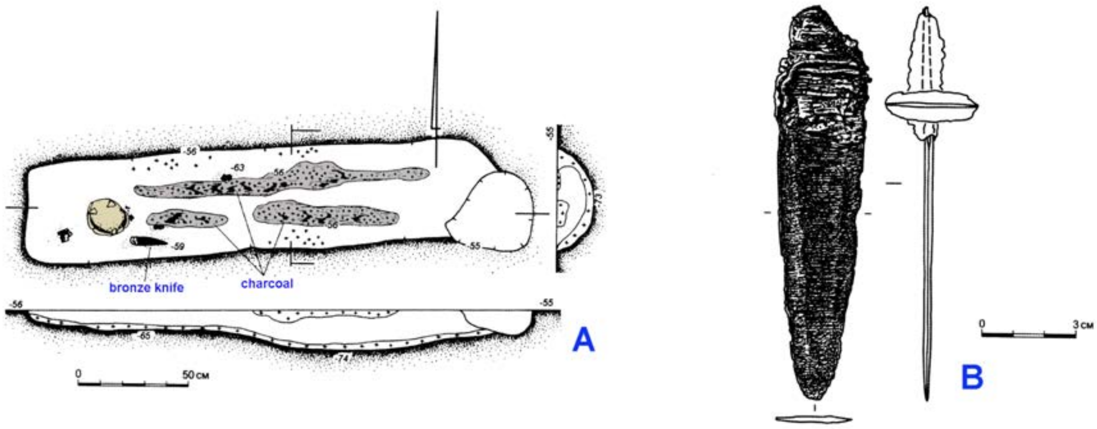

*__Fig. S71.__ Burial 6 at the Satyga-16 site. __A__ - burial; __B__ - grave goods.*

*(b) Burial 17 (individual ID I32551)*

**Burial 17** (individual ID I32551) suffered significant damage due to disturbance by an uprooting, making it difficult to reconstruct its shape and parameters. It is believed that two graves were originally present at this location, oriented at an angle to each other. This is supported by the preserved cruciform outlines of the shared depression, as well as the presence of one skull within the pit and the discovery of another individual's skull not far from its eastern wall.

Presumably, a burial (17a) was initially constructed in this spot, aligned along the west-southwest axis, containing the remains of a 25-30-year-old man.

Later, another pit (17b) was dug above it, oriented along the NW-SSE line, approximately 0.5 m wide, in which a woman of about 35 years old was buried.

The sloping walls of this grave descended steeply to the bottom, reaching a depth of 0.2 m into the subsoil. In the redeposited layer of the uprooting pit, a bronze scraper, a bronze object resembling a chisel with remnants of a bone handle, a stone arrowhead, and a tool on a plate were found. Above the graves, an accumulation of jar vessel fragments was recorded, while fragments of a third vessel were uncovered to the south of the burials (Trufanov, 2011).

Burial 17 is dated to 2288-2058 calBCE (3770±25 BP, PSUAMS-12647).

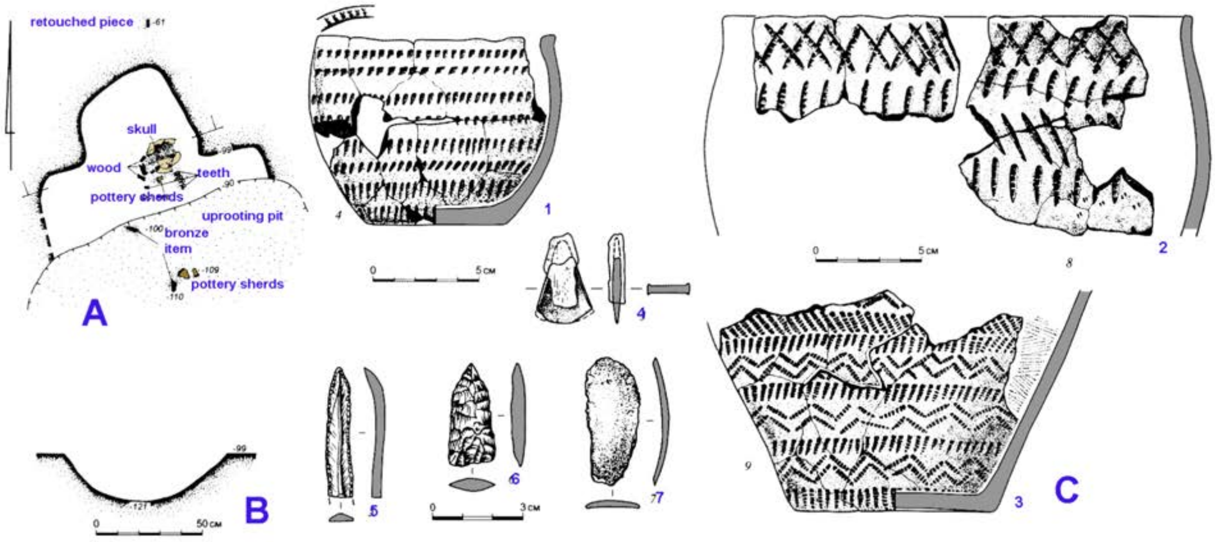

***Fig. S72.*** *Burial 17 at the Satyga-16 site.* ***A*** *- burial;* ***B*** *- a section of the grave pit;* ***C*** *- grave goods.*

##### II.F.iii. Peripheral Area of the Seima-Turbino Phenomenon

###### II.F.iii.a) Chernoozerye-1 burial site

At the Chernoozerye-1 burial site (described previously in SI Section II.C.ii.b.4)), a grave possibly dated to the Seima-Turbino horizon among others that are from the Eneolithic and Late Bronze age periods was excavated. The presence of the Seima-Turbino phase is also evidenced by a find of an anthropomorhic figurine, which finds parallels in Rostovka and Galich hoard (Stefanov, 2004).

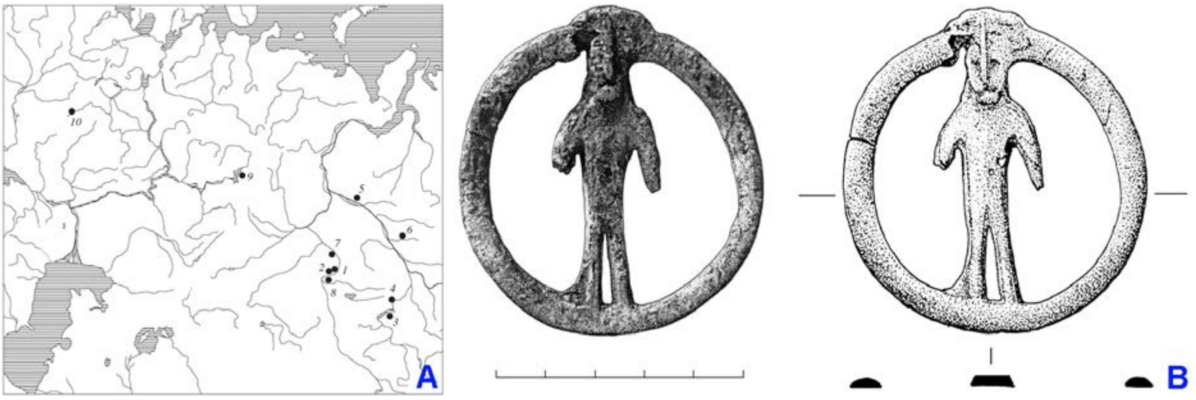

***Fig. S73. A****: anthropomorphic figurines in the Bronze Age: 1 - Chernoozerye-1, 2 - Borovyanka-17, 3 - Zavyalovo-1A, 4 - Krokhalevka 13, 5 - Saigatino-6, 6 - Novy-Napas, 7 - Chudskaya-Gora, 8 - Rostovka, 9 - Bor-Lyonva, 10 - Galich treasure.* ***B****: antrhopomorhic figurine found at Chernoozerye-1 (Stefanov, 2004, modified).*

*(c) Burial 132 (individual ID I6787)*

**Burial 132** (individual ID I6787). The grave is aligned along the northwest-southeast axis, measuring approximately 230x75x30 cm. The southeastern part of the grave was disturbed by a ditch associated with an Early Iron Age mound, resulting in the displacement of the skull onto the chest bones. The deceased individual, presumed to be a woman and estimated to be around 40 years old, is laid in a supine position with the head facing southeast. No grave goods accompanied this burial (Gening and Stefanova, 1994). The burial is dated to 2916-2884 calBCE (4285B±20 BP, PSUAMS-2920).

The date looks too old for the Seima-Turbino period, but this fact can be explained by a freshwater reservoir offset of the radiocarbon dating. The dated individual had a fish-based diet, which is proved by the observed δ13C/δ13N ratio. The offsets in radiocarbon dates from the Seima-Turbino period sites are well observed in the Rostovka cemetery, where some dates demonstrate even older chronological attribution (see Table S2, dates on burials 29 and 34). Based on this observation and on the results of the genetic analysis, which revealed ancestry proportions characteristical for the Seimo-Turbino period individuals, we assume an Early-Middle Bronze Age chronological attribution of burial 132 at the Chernoozerye-1 site.

##### II.F.iii.b) Tatarka Hill site

In 1998, a flat burial ground on Tatarka Hill was discovered by A.S. Vdovin and S.A. Krasnolutsky. Excavations were conducted between 1998 and 2000 by the same archaeologists (Kuz'minykh, 2011).

The burial ground is situated on an elevated platform called Tatarka Hill, located on the western outskirts of the village of Bolshoye Ozero, in the Sharypovsky District of the Krasnoyarsk Krai, Russia. The estimated area of the Tatarka burial ground is approximately 700-800 square meters. However, it is likely that a significant portion of the site was destroyed during the extraction of stone for road backfilling. Therefore, the original size of the site might have been around 1,500-2,000 square meters. The burials, comprising at least 50 individuals, are arranged in rows, closely spaced at intervals of several meters, located on a small elevated area that terminates in cliffs on the southwestern and southern sides. The orientation of the skeletons is towards the west-northwest. The primary grave goods found at the site include bronze celt axes, knives, awls, ornaments, jade rings, and stone arrowheads. Except for one flat-bottomed vessel, pottery is absent from the burials.

Thecemetery at the Tatarka Hill contains graves of various period, the latest phase dates from the Middle Ages.

The most part of the burials dates back to the Late Bronze Age. The funeral rite and material culture of these burials reveal similarities with the monuments of the forest-steppe “andronoid” cultures of Western Siberia (such as Elovka 2, Sopka-2, Chernoozerye-1). Among the metal artifacts listed are single-edged and, less commonly, double-edged knives, “horned” bracelets, rings covered with gold foil, various plate decorations, etc. In burials 14 and 15, two false-eared celts of type K–54 of the Samus-Kizhirovo tradition were found.

Another group of burials is characterized by connections with the cultures of the Baikal region, as indicated by stone burial structures and jade rings (Kuz'minykh, 2011).

A group of burials in shallow grave pits, oriented in the northwest to southeast direction, contained individuals laid in supine positions with their heads to the southeast. The grave pits were partially covered by stone slabs. The burials lack grave goods, and their chronology could be estimated only based on radiocarbon datings.

We sequenced four individuals from this group of burials and dated two of them. The radiocarbon dates (2288-2058 calBCE (3770±25 BP, PSUAMS-9048) and 2021–1884 calBCE (3585 BP±20, PSUAMS-7545)) reveal the attribution of these burials to the Seima-Turbino period and distinguish them from the total mass of the Late Bronze Age Andronovo-like burials. The freshwater reservoir offset of these dates might have explained the incongruity of the C14 data and the archaeological context, but the isotopic data reveals the highest values of δ13C among all our analyzed samples from Siberia and a relatively low value of δ13N. The δ13C/δ13N ratio better supports a foraging-hunting diet than the intensive fish consumption of the studied individuals. Based on this observation, we assume a specific Early-Middle Bronze Age phase at the Tatarka-Hill cemetery.

We found the closest parallels in the burial rites, observed in those Early-Middle Bronze Age graves of the Tatarka-Hill site, in burials of the Nefteprovod-1 and 2 cemeteries of the Anzhevskiy ensemble located in Kansk forest-steppe. Alongside the geographical proximity, the Tatarka-Hill and the EMBA burials of the Anzhevski ensemble share attribution to the same chronological period.

#### (1) Burials at the Tatarka Hill Site

##### *(a) Burial 2 (individual ID I20305)*

**Burial 3** (individual ID I20305). The grave pit is covered with stone slabs, and additional slabs are placed along the sides of the grave pit. The shape of the grave pit is elongated and oval, extending in the northwest to southeast direction.

At a depth of 50-60 cm from the surface, the remains of a single individual, estimated to be 25-35 years old, were discovered. The skeleton was found in a supine position. The skeletal remains are in a poorly preserved condition, with damage to the outer layer (cortical layer) of the bones. The skull has been restored, and only small fragments of the pelvis, ribs, vertebrae, as well as the bones of the feet and hands, have survived, as the epiphyses of the long bones of the postcranial skeleton are all destroyed.

Dento-jaw pathologies observed include caries, evidence of tooth loss during the individual's lifetime, and abscesses on the upper and lower jaws, along with chips on the teeth. Arthrosis is present in the thoracic and lumbar spine, as well as in the elbows and hands of the upper extremities.

No associated grave goods or additional items were found in the burial.

##### *(b) Burial 26 (individual ID I10998)*

**Burial 26** (individual ID I10998). The grave pit of burial 26 exhibits an elongated oval shape in the northwest-southeast direction. The interred individual's remains were found in a supine position, lying on their back. The skeleton belonged to a woman estimated to be 20-25 years old, and it was discovered at a depth of 35-40 cm below the soil surface. Based on the overall orientation of the remains, the deceased was positioned with her head towards the south-southeast. The skeletal preservation is very poor, with fragments of the skull (occipital part of the braincase and frontal bone showing outer cortical layer destruction, along with a lower jaw displaying damaged condyles). The diaphyses of long bones in the postcranial skeleton show evidence of destruction of the epiphyses. Additionally, fragments of the pelvis, shoulder blades, ribs, vertebrae, as well as bones from the hands and feet were recovered.

The accompanying grave goods consist of a fang, possibly of a wolf or dog. The burial is dated to 2021–1884 calBCE (3585 BP±20, PSUAMS-7545).

##### *(c) Burial 27A (individual ID I20547)*

**Burial 27A** (individual ID I20547). The grave pit of burial 27a was uncovered at a depth of 25-30 cm from the contemporary surface. It has an elongated oval shape in the northwest-southeast direction and is partially covered by stone slabs. The interred individual, a woman estimated to be 35-40 years old, was found in a supine position with her head towards the southeast. The skeletal remains are highly deteriorated, with only small fragments of the skull remaining (due to poor preservation, the skull was excavated as a monolith with the surrounding soil, making restoration impossible). The lower jaw exhibits a destroyed left condyle. Among the postcranial skeleton, fragments of the humerus, femur, tibia, one fibula, small fragments of the pelvic bones, and the first two cervical vertebrae were recovered. Signs of arthrosis are evident in the cervical spine.

No associated grave goods or additional items were found in the burial.

*(d) Burial 26 (individual ID I20549)*

**Burial 55** (individual ID I20549). The grave pit of burial 55 was discovered at a depth of 35-40 cm from the present surface. It possesses an elongated-trapezoidal shape oriented in the north-south direction and is partially covered with stone slabs. The interred individual, a woman estimated to be 30-35 years old, was found in an extended supine position. Observations on the skeleton include missing teeth, an abscess in the region of the lower second premolar, and evidence of periodontal disease. Arthrosis is apparent in various areas, including the cervical, thoracic, lumbar, lumbosacral, and sacroiliac regions, as well as the left elbow, wrist, and hand.

No associated grave goods or additional items were found in the burial.

##### II.F.iii.c) The Nefteprovod-1 and Nefteprovod-2 sites (Anzhevskiy complex), archaeologically related to the site of Tatarka Hill

The archaeological sites Nefteprovod-1 and Nefteprovod-2 of the Anzhevsky ensemble were discovered in 1972 by N.A. Savelyev. The ensemble is located in the Kansk forest steppe, on the right bank of the Kan River, in Ilansky district of the Krasnoyarsk Krai (Vybornov et al., 2015). The excavations of 2015-2016 carried out at the sites Nefteprovod-1 and Nefteprovod-2 revealed 29 burials of the multi-phase burial ground. The burials were preliminary attributed to several phases, the Neolithic, the Late Bronze Age (Krasnoyarsk culture), the Bronze Age (taiga cultural circle), Early Iron Age, and the Middle Ages (Timoshchenko, Vybornov, Rybin, 2018).

The first burial at the Nefteprovod-2 site was excavated in 2011 by E.V. Knyazeva [Mandryka et al., 2018] in a survey excavation block. The skeleton was laid in a supine position with his head to the southeast, no grave goods were associated with the burial. The **buried individual** was studied genetically (individual ID kra001) and dated to 4280-4085 calBP (Kılınç et al., Sci. Adv. 2021; 7:eabc4587; p. 2).

Further excavations of 2011 and 2015 revealed other burials dated to the same period and located around the first one or in other areas in the Nefteprovod 1-2 sites.

A group of burials around the first one were numbered from 1 to 8. They are located on the low terrace at heights 216.49 - 217.65 m (ASL), 12-13 m above the river water level, in an area 22x21 m. The graves were placed in three rows, elongated from south to north (up the slope from the river bank). The eastern row includes graves 1, 2, 5, 6, and 8, the middle row – the grave, excavated in 2011 and burial 4, and the western one includes burial 7.

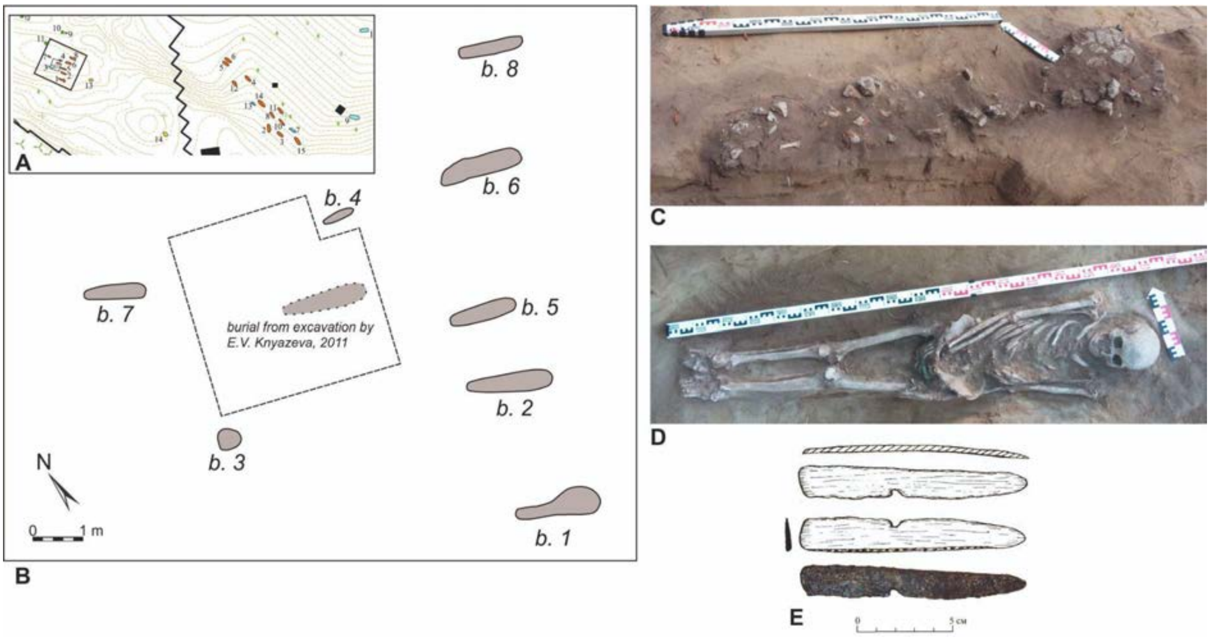

*Fig. S74. Anzhevsky ensemble. **A:** layout of burials at the archaeological sites Nefteprovod-1 and Nefteprovod-2. **B:** layout of the burials of the lower group of burials at the site Nefteprovod-2. **C:** masonry over burial 2 at the Nefteprovod-2. **D:** skeleton in burial 2 at the site Nefteprovod-2. **E:** bronze knife in burial 2 at the site Nefteprovod-2.*

All these graves were sunk from the top of the second lithological layer that contained Neolithic-Bronze Age artifacts. The gravepits were amorphous and their boundaries were hard to recognize. The skeletons were put in the anatomical order in supine positions with their heads oriented upstream, to the southeast. The faces were oriented upwards, or slightly tilted to the right, when recognizable. Traces of fire were recognized on the bones of burial 5, and charred birch bark in burials 5 and 8. Elongated rubbleworks masonry 1.03–1.95 × 0.22–0.43 m in area were recorded in burials 1, 2, 6, 7, 8. Separate stones were found above burial 5. In burials 2, 6, and 8 the skeletons were complete, and in burials 1, 4, 5, 7 some bones were missing. Burial 4 had no rubblework and its gravepit was relatively smaller, 0.67 × 0.18 m, it contained remains of a 5-6 year old child.

Grave goods include bronze knives from burials 2 and 4, bronze beads from burial 5, and fragments of bronze items on a skeleton in burial 8. No grave goods were recorded in burials 1, 6, 7.

Another group of burials dated to the same period as kra001 was discovered 120 m east of it, on the edge of the upper terrace, at the Nefteprovod-1 site, in 2015-2016. It includes graves 2-6, 8, 10-12, and 14-15. The graves of this group are located along the edge of the terrace in three rows, elongated from the northwest to the southeast, at the heights of 219.3 - 223.74 m (ASL), 15-19 m above the river water level. The graves are elongated along the northwest-southeast line, the skeletons were put in supine positions, their heads are oriented to the southeast. Above graves 2 and 3 some stones were found, above graves 5, 6, 14 an elongated rubblework was recorded, and above grave 8 was recorded a round-shaped one. In grave 10, a bronze knife and a bronze piece were found near the skull.

Radiocarbon dates have been obtained for two burials of this group of graves, burial 14 (4248-3902 calBP (3733±58 BP, GV-02873)) and burial 15 (3886-3574 calBP (3473±59 BP, GV-02874)).

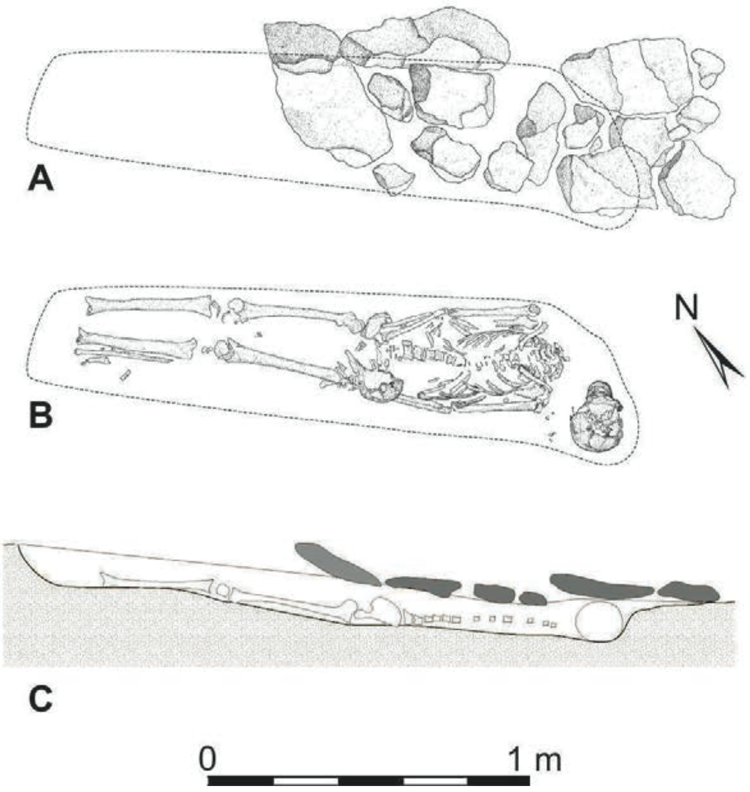

*Fig. S75. Anzhevsky ensemble.* ***A:*** *rubblework over burial 14 at the Nefteprovod-1.* ***B:*** *skeleton in burial 14 at the site Nefteprovod-1.* ***C:*** *longitudinal section of burial 14 at the site Nefteprovod-1.*

In conclusion, we identify a group of burials at the Nefteprovod 1 and 2 sites that share common details in burial rites and are chronologically dated to the period 4200-3900 calBP.

At the Nefteprovod-2 site, those are burials in shallow and narrow grave pits containing skeletons put in supine positions with their skulls in the southeastern parts of grave pits. Above some burials, elongated rubbleworks were recorded. In some cases, scarce grave goods were found, including bronze knives in the hands of the dead and beads in the head areas. In two cases, there were traces of fire on the bones and fragments of charred birch bark. One of the burials of this group, located in the middle row and explored in 2011 by E.V. Knyazeva has a radiocarbon date of 4280-4085 cal BP [Kılınç et al., Sci. Adv. 2021). Apparently, the entire group of similar burials should be dated to the same period.

At the Nefteprovod-1 site, a group of burials could be associated with the same population as the kra001. They share such burial rites details as shallow grave pits, the supine position of the dead, orientation to the southeast, and presence of rubble works covering the graves. Some differences were also distinguished in the planigraphy of the rows of graves, the location of bronze knives, and the absence of any traces of fire on the bones of the buried persons.

#### *II.G. Krasnoyarsk Cisbaikal_LNBA-rich outliers*

The Late Bronze Age period in the Minusinsk Basin has a long history of study and interpretation. The first steps of it are dated back to the 1920s century when S.A. Teploukhov described them as the Karasuk culture. Several decades after the accumulation of data revealed the nonuniformity of the Karasuk-associated assemblage, and M.P. Gryaznov was the first researcher who in the 1960s reattributed some "non-typical" Karasuk artifacts as a specific phase of the cultural development called Kamenny-Log. Later, the chronological definitions of the "Classic Karasuk" and the "Kamenny-Log Karasuk" were elaborated, and the last one was attributed to 1000-800 BCE while the "classic" one got an earlier date, 1300-1000 BCE (Gryaznov 1979).

Although this version became conventional among most archaeologists, being supported by G.A. Maksimenkov, B.N. Pyatkin, and others (Maksimenkov 1975), some researchers were inclined to interpret "atypical" assemblage as a specific cultural group, and not as a stage of linear development. This opinion was most clearly articulated by N.L. Chlenova, who described them as a specific Lugavskaya culture, which, she postulated, developed from local Neolithic and Okunev traditions and later co-existed with the Karasuk one in the Minusinsk Basin during 1400/1300-700/600 BCE (Chlenova 1972).

Later studies expounded one of these two competing models in one way or another. In the most recent publication, a compromising model was suggested by A.V. Polyakov (2022).

He followed the idea of determination of several stages in the Late Bronze Age period in the Minusinsk basin into several successive stages (calendar dates are approximate):

- LBA-I (Karasuk) with substages A and B - 1450-1250 BCE
- LBA-II (Karasuk-Lugavskaya) - 1300-1075 BCE
- LBA-III (Lugavskaya) with substages A, B, and C - 1025-825 BCE
- LBA-IV (Bainov) with substages A and B - 850-750 BCE

This chronological scheme, however, does not repeat uncritically the old models of stadial cultural development. Polyakov and Lazaretov (2008) assume the gradual replacement of the Karasuk population by newcomers.

In their model, the formation of the Karasuk assemblage of the LBA-I stage was a result of influence (or migration) of the Andronovo Alakul/Fedorovka population. The homogeneous "classic" Karasuk material culture of the LBA-I stage experienced extra-cultural influences starting from the LBA-II period. Those influences are reflected in the dispersal of "atypical" pottery, grave pits instead of stone boxes, and some categories of artifacts. In the LBA-III period, the innovations completely replaced the initial Karasuk tradition, recording transformation to the Lugavskaya cultural assemblage. Polyakov calls the changes in material culture "barbarisation", because of the disappearance of advanced bronze casting technologies, but notices the increase in the number of sites. He considered the reasons for cultural transitions in migration processes, resulting in the first continuous cohabitance of groups of different origins and after that a gradual replacement of the Karasuk population with the newcomers.

The origin area of the migration that led to the formation of the Lugavskaya culture is under discussion. Polyakov assumes an infiltration of a new group of the population that bore Late Kazakhstan Andronovo traits in pottery and Central Asian bronzes across West Sayan. Earlier investigators found some parallels of the Lugavskaya pottery in the southern taiga zone of the Middle Yenisei (Chlenova 1972). At the moment, archaeological data lacks evidence for the robust reconstruction of the routes of migrations, that impacted transformations of the material culture in the Minusinsk Basin during the LBA.

The published archaeogenetic data on the Minusinsk LBA populations (Allentoft et al. 2015) mostly derives from the “classic” Karasuk sites of the LBA-I stage. Besides, some samples from the Arban-1 site were collected frob the graves attributed to the Lugavskaya cultural assemblage (Savinov and Polyakov 2007, Polyakov 2019). We list them below with their radiocarbon dates and their ADMIXTURE results for the Steppe_MLBA and Cisbaikal_LNBA components at K=18 (SI Section IV, Extended Data Figure 10).

**RISE496** - <u>burial 6</u> - 1416-1260 calBCE (3070±28 BP, OxA-31213): *Steppe_MLBA 75%, Cisbaikal_LNBA 4%*

This individual was attributed as a "Classic" Karasuk dating to the LBA-I stage.

**RISE495.SG** - <u>burial 27</u> – undated: *Steppe_MLBA 44%, Cisbaikal_LNBA 36%*

This burial, according to Polyakov (2007, 2022) belongs to a small group of "atypical" burials with specific pottery ("cordoned ware"), which started to appear at the very end of the LBA-I-B stage and later shaped the material culture of a transitional (Karasuk->Lugavskaya) LBA-II period.

**RISE497.SG** - <u>burial 55</u> – undated: *Steppe_MLBA 0%, Cisbaikal_LNBA 83%*

This burial belongs to the latest, the LBA-III-A (Lugavskaya) stage of the Arban-1 site. The burials of this stage are placed aside from the earlier group of burials on the site.

The presence of the Cisbaikal_LNBA component was not observed in other individuals attributed to the Karasuk population. Their archaeological context relates them to the cultural transformations in the Minusinsk Basin LBA, which resulted in the replacement of the Karasuk culture with the Lugavskaya one. This observation supports the archaeological reconstructions that consider the migration-driven changes in burial rites and artifactual assemblage. The Cisbaikal_LNBA may have participated in that migration as at least one of the population groups involved in this transformation.

Unfortunately, due to the poor state of the records, we were unable to locate archaeological documentation for the other Cisbaikal_LNBA-rich outlier, the previously-published individual labeled RISE554.SG. However, the radiocarbon date and location data provided in the previous publication indicate that this individual is likely to represent part of the same phenomenon as described in this section.

#### II.H. References (for section II)

Agapov, S. A. (Ed.) *Khvalynsk Copper Age cemeteries and Khvalynsk Copper Age cultures: The researches of materials* **2** (Ofort-Press, 2010).

Agapov, S. A., Vasil'yev, I. B. & Pestrikova, V. I. *Khvalynsk Eneolithic Burial Site* (Saratov St. Univ. publ. house, 1990).

Alekseev, A. N. *The Stone Age of the Olyokma* (Irkutsk Univ. publ. house, 1987).

Alekseev, A. N. *Ancient Yakutia: the Neolithic and the Bronze Age* (Institute of Archaeology and Ethnography SB RAS publ. house, 1996).

Alekseev, A. N., Dyakonov, V. M., Solovyova, E. N., Nikolaev, E. N. & Boeskorov, G. G. The burial Ogonyok in the middle Lena River region: a new site of the Bel'kachi Culture. *Bulletin of Archeology, Anthropology, and Ethnography* **1** (**56**), 2071-0437-2022-56-1-6 (2022).

Alekseev, A.N. & Dyakonov, V. Radiocarbon Chronology of Neolithic and Bronze Age Cultures in Yakutia. *Archeology, Ethnography and Anthropology of Eurasia* **3** (**39**), 26-40 (2009).

Alekseyev, V. P. Eneolithic Skull from Krasnoyarsk (on the Issue of Southern Admixture in the Population of the Altai-Sayan Upland). *Brief Comm. Inst. Arch.* **34**, 79-85 (1960).

Alekseyev, V. P. in *Anthropological Collection* **3**: *Proceedings of the Institute of Ethnography* **71**, 107-206 (1961a).

Alekseyev, V. P. in *Questions of the history of Siberia and the Far East* (ed. Dulov, V. I.) 377-385 (Siberian Branch of the Acad. Scien., 1961b).

Alekseev, A. N., Zhirkov, E. K., Stepanov, A. D., Sharaborin, A. K. & Alekseeva, L. L. The burial of Ymyiakhtakh warriors in Kyordyughen, Yakutia. *Archaeology, Ethnology & Anthropology of Eurasia* **2** (**26**), 45-52 (2006).

Andreev, K. M., Berezina, N. S., Vybornov, A. A. & Stavitskiy, V. V. in *Archaeology of the Volga-Ural Region* (ed. Sitdikov, A. G., Galimova, M. Sh.) **1**, 250-260 (AS of the Republ. Tatarstan publ. house, 2021).

Andreyev, K. M., Vasil'yeva, I. N. & Vybornov, A. A. Neolithic pottery assemblage of the Chekalino-4 site: morphology, technology, chronology. *The Volga River Region Archaeology* **1** (**23**), 8-27 (2018).

Argunov, V. G. & Pestereva, K. A. Exploration of the Yakut Arctic: Questions of ancient migration and mobility in the Circumpolar world. *Society: history, philosophy, culture* **3**, fik.2018.3.6 (2018).

Aseyev, I. V. The Kitoi culture in the Neolithic of the Baikal region and adjacent territories: issues of chronology, migration areas of its carriers. *Archeology, Ethnography and Anthropology of Eurasia* **2** (**10**), 59-70 (2002).

Aseyev, I. V. in *Problems of Archaeology, Ethnography, and Anthropology of Siberia and Adjacent Territories* (ed. Derevyanko, A. P. & Molodin V. I.) **15**, 34-38 (2009).

Bader, O. N. The Neolithic of the Urals. *Materials and Research in Archaeology of the USSR* **166**, 157-171 (1970).

Bagashev, A. N. & Solodovnikov, K. N. in *In this connection: Collection of articles dedicated to the anniversary of Margarita Mikhailovna Gerasimova* 100-140 (Buki Vedi, 2019).

Bazaliyskiy, V. I. Funeral complexes of the Late Mesolithic - Neolithic period of the Baikal Siberia: burial traditions, absolute age. *News of the Laboratory of Ancient Technologies* **1** (**9**), 43-101 (2012).

Berdnikov, I. M., Makarov, N. P. & Savenkova, T. M. in Study of the ancient history of Northern and Central Asia: from origins to the present: abstracts of the International Scientific Conference dedicated to the 300th anniversary of D.G. Messerschmidt's expedition (ed. Molodin, V. I.) 73-76 (SB RAS publ. house, 2022).

Berezina, N. S., Vybornov, A .A., Nikitin, V. V. & Stavitsky, V. V. in Archaeology of the Volga-Ural Region (ed. Sitdikov, A.G., Chizhevskiy, A.A.) **1**, 297-315 (AS Republic Tatarstan publ. house, 2021).

Bobrov, V. V. On the Problem of Interethnic Relations in South Siberia in the Third and Early Second Millennia B.C. *Arctic Anthropology* **25** (**2**), 30–46 (1988).

Borodaev V. B., Kiryushin K. Y., Kuzmenkin D. V. & Solodovnikov K. N. Ornaments Made from Unio Shells in a Neolithic Burial at Ust-Aleika-5, Barnaul, Southwestern Siberia. *Archaeology, Ethnology & Anthropology of Eurasia* **50** (**1**) 1563-0102.2022.50.1.048-056 (2022).

Borodkin, Y. M. Works of visual art from the Neolithic burials of the Vaskovo burial ground. *News of the Laboratory of Ancient Technologies* **7**, 99-109 (1976).

Chernykh, E. N. & Kuzminykh, S.V. in: *Archeology of the USSR. The Bronze Era in the Forest Belt of the USSR* (ed. Bader, O. N., Kraynov, D. A. & Kosarev, M. F.) 84–105 (Nauka, 1987).

Chernykh, E. N. & Kuzminykh, S. V. *Ancient Metallurgy of Northern Eurasia (the Seima-Turbino phenomenon)*. (Nauka, 1989).

Chernykh, E. N. & Kuzminikh, S. V. In: The Great Russian Encyclopedia (ed. Osipov U. S. et al) **29**, 675 (2015).

Chizhevskiy, A. A. in Proceedings of the 2 (18) All-Russian Archaeological Congress in Suzdal (ed. Derevyanko, A. P. & Makarov, N. A.) **1**, 367-37 (Archeology Institute publ. house, 2008).

Chizhevskiy, A. A. & Shipilov, A. V. in *Ural Archaeological Meeting, dedicated to the 85th anniversary of G. I. Matveyeva and the 70th anniversary of I.B. Vasil'yev* (ed. Vybornov, A. A. et al) **21**, 80-84 (Porto-Print, 2018).

Chizhevskiy, A. A. & Shipilov, A. V. in *Archaeology of the Volga-Ural Region* (ed. Sitdikov, A.G., Chizhevskiy, A.A.) **2**, 140-163 (AS Republic Tatarstan publ. house, 2021).

Chizhevsky, A. A., Lyganov, A. V. & Kuzminykh, S. V. Early (atabaevo) Stage of Maklasheevka culture. *Archaeology of the Eurasian Steppes* **2**, 99-123 (2019).

Chlenova, N. L. Chronology of monuments of the Karasuk epoch. Materials and research on the archaeology of the USSR **182** (Nauka, 1972).

Debets, G. F. Paleoanthropology of the USSR: *Proceedings of the Institute of Ethnography* **4** (AS USSR publ. house, 1948).

Dromov, V. A. in *Chronology and cultural affiliation of monuments from the Stone and Bronze Ages of Southern Siberia* (ed. Derevyanko, A. P. et al) 40-43 (Altai Univ. publ. house, 1988).

Dudarek, S. P. & Lokhov, D. N. Burial complexes of the Bronze Age in the Northern Angara Region. Chronology problems and cultural identity. *News of the Irkutsk St. Univ. Series: Geoarchaeology. Ethnology. Anthropology* **7**, 54-80 (2014).

Dus, F. Deux cranes neolithigues siberiens. Anthropologiai Fuzetok (Anthropologia Hungarica), E.I., **1-3**, 72-78 (1923).

Dyakonov, V. M., Shpakova E. G., Chikisheva T. A. & Pozdnyakov D. V. in *Ancient cultures of Northeast Asia. Astroarchaeology. Paleoinformatics* (ed. Alexeev, A. N.) 65-90 (Nauka, 2003).

Dyakonov, V. M., Kuzmin, Ya. V. & Hogins, G. W. L. Burial of the Vilyuyskoye Highway in Yakutsk: problems of determining the absolute age and cultural affiliation. *News of the Laboratory of Ancient Technologies* **9**, 122-128 (2012).

Dyakonov, V. & Takase, K. in *Man and the North: Anthropology, Archaeology, Ecology. Proceedings of the All-Russian Scientific Conference* (ed. Bagashev, A.N., et al.) **4**, 73-77 (Tyumen Scientific Center of the SB of the RAS, 2018).

Engovatova, A. V. in *The Great Russian Encyclopedia* (ed. Osipov, U. S. et al) **18**, 208 (2011).

Engovatova, A. V. in *The Great Russian Encyclopedia* (ed. Osipov, U. S. et al) **35**, 707 (2017).

Epimakhov, A. V. & Mosin, V. S. Chronology of the Trans-Urals Eneolithic. *Bulletin of Archeology, Anthropology and Ethnography* **4** (**31**), 27-37 (2015).

Everestov, S. I. in *Ancient cultures of Northeast Asia. Astroarchaeology. Paleoinformatics* (ed. Alexeev, A. N.) 54-64 (Nauka, 1999).

Everestov, S. I. Some parallels in cultures of ancient Ymyyakhtakhians and Yukaghirs of the 17th-19th centuries. *Arctic and North* **15**, 152-159 (2014).

Fedorova, S. A. *Genetic portraits of the peoples of the Republic of Sakha (Yakutia): analysis of mitochondrial DNA and Y-chromosome lines* (Yakut SC SB of RAS, 2008).

Fedoseeva, S. A. *Ancient cultures of Upper Vilyui* (ed. Okladnikov, A. P.) (Nauka, 1968).

Fedoseeva, S. A. *The Ymyyakhtakh culture of Northeast Asia* (ed. Larichev, V. E.) (Nauka, 1980).

Fedoseeva, S. A. in *Archeology of Yakutia* (ed. Alexeev, A. N. et al) 79-98 (Yakut St. Univ .publ. house, 1988).

Fedoseeva, S. A. in *Archaeological research in Yakutia* (ed. Mochanov, Yu. A.) 64-105 (Nauka, 1992).

Fribus, A. V. & Grushin, S. P. in *Man and the North: Anthropology, Archaeology, Ecology: Proceedings of the All-Russian Conference* (ed. Bagashev, A. N. et al) **3**, 196–199 (Inst. of Probl. of Devel. of the North SB RAS publ. house, 2015).

Fribus, A. V. & Grushin, S. P. in *Proceedings of the 5 (21) All-Russian Archaeological Congress in Barnaul-Belokurikha* (ed. Derevianko A. P. & Tishkin A. A.) **1**, 178-182 (Altai Univ. publ. house, 2017a).

Fribus, A. V. & Grushin, S. P. Bird feathers in the makeup of a Neolithic costume (based on the materials of the Chumysh-Perekat cemetery in the south of West Siberia). *Notes of the Institute of the History of Material Culture* (ed. Nosov E. N.) **16**, 20-34. (2017b).

Fribus, A., Grushin, S. in *Radiocarbon in Archeology and Paleoecology: Past, Present, Future. Proceedings of the International Conference* (ed. Burova, N. D. et al) 978-5-91867-213-6-103-104 (Porto-print, 2020).

Fribus, A. V., Grushin, S. P., Zaybert, V. O. & Trusova, Ye. V. in *Modern Solutions to Current Issues of Eurasian Archaeology* (ed. Tishkin A. A.) **2**, 42-47 (Altai Univ. publ. house, 2018).

Fribus, A. V. et al. in *Interdisciplinary Study of Archaeology of Western Siberia and Altai* (ed. Derevyanko, A.P. & Kiryushin, Yu. F.) **1**, 107-108 (Altai Univ. publ. house, 2014).

Garkusha, Yu. N., Marchenko, Zh. V. & Grishin, A. E. Burials of “Pashina site” (the Northern Angara River basin): Problems of defining of chronological and cultural background. *Bulletin of Novosibirsk St. Univ. Series History, Philology* **12** (**7**), 186-197 (2013).

Gening, V. F., & Stefanova, N. K. *Chernoozer'ye-1 - Bronze Age Cemetery of the Middle Irtysh Region.* Preprint (Ural St. Univ., 1994).

Gening, V. F. in *Monuments of Ancient History of Eurasia* (ed. Kojin P. M. et al) 211-220 (Nauka, 1975).

Georgiyevskaya, G. M. *The Kitoi culture of the Transbaikal region* (Nauka, 1989).

Gerasimov, M. M., Chernykh, E. N. in *Prehistoric archaeology of Siberia* (ed. Mandel’shtam, A. M.) 23–48 (Nauka, 1975).

Gerasimova, M. M. Neolithic Burials at Lake Dolgoye (Kansk). *Voprosy antropologii* **18**, 135-142 (1964).

Glusskaya, Z. K. in *Materials and Research on Archaeology, Ethnography, and History of the Krasnoyarsk Region* (ed. Lipsky A. N.) 29-37 (Krasnoyarsk Book publ. house, 1963).

Goldina R. D., Chernykh E. M. Archaeological Map of the Karakulinsky District, Udmurt Republic: *Materials and Research of the Kama-Vyatka Archaeological Expedition* **18** (Udmurtia St. Univ. publ. house, 2011).

Goryunova O. I. *Ancient Burial Grounds of the Baikal Region (Neolithic-Bronze Age)* (Irkutsk St. Univ. publ. house, 2002).

Goryunova, O. I., Novikov, A. G., Turkin, G. V. & Veber, A. V. Results of Study and Dating of Early Neolithic Burial Complexes in the Lake Baikal Region. *Bulletin of the Irkutsk State University. Geoarchaeology, Ethnology, and Anthropology Series* **23**, 44-70 (2018).

Grebenyuk, P. S., Fedorchenko, A. Yu., Lebedintsev, A. I. & Malyarchuk, B. A. The ancient cultures of the extreme Northeast Asia and ethnogenetic reconstructions. *Tomsk Journal of Linguistics and Anthropology* **2** (**24**), 2307-6119-2019-2-110-136 (2019).

Grevtsov, Y. A., Lysenko, D. N. & Galukhin, L. L. in *Problems of Archeology, Ethnography, Anthropology of Siberia and adjacent territories* (ed. Derevyanko, A. P. & Molodin, V. I.) **16**, 509-514 (2010).

Gurina, N. N. & Kraynov, D. A. in: *Neolithic of Northern Eurasia* (ed. Oshibkina, S.V.) 173-180 (Nauka, 1996).

Grushin, S. P. & Fribus, A. V. in *Prehistoric art in the context of cultural and historical processes of Eurasia: to the 300th anniversary of the scientific discovery of the Tomskaya Pisanitsa. Proceedings of the International Conference* (ed. Muhareva, A. N. & Rusakova, I. D.) 318-323 (KRIPKiPRO, 2021).

Gryaznov, M.P. (ed.) *Complex of archaeological sites near the Tepsei Mountain on the Yenisei* (Nauka, 1979).

Herman P. V. & Leontiev S. N. Materials from the burials of Sergushkin-3 ground cemetery in the Northern Angara region: on the issue of chronology and geneses of the Tsepan Culture. *Bulletin of Archeology, Anthropology and Ethnography* **1** (**56**), 101-114 (2022).

Hlobystin, L. P. *Ancient history of the Taimyr Arctic* (Dm.Bulanin, 1998).

Ivashina, L. G. On the problem of the Kitoi culture (on the history of researching the problem). *News of the Laboratory of Ancient Technologies* **1** (**4**), 9-14 (2006).

Kashin, V. A. *The Neolithic of the Middle Kolyma* (ed. Bravina, R. I.) (Nauka, 2013).

Kashin, V. A. & Kalinina, V. V. *Pomazkino archaeological site complex as a part of the circumpolar culture* (Severoved, 1997).

Khohlov, A. A. & Nechvoloda, A. I. A human cranium, buried on the territory of the ancient settlement Gladunino-3. *Questions of the Archeology of the Urals* (ed. Kovaleva, V. T.). **24**, 192-201 (Ural Univ. publ. house, 2002).

Khvostov, V. A. in *Problems of Study of the Neolithic of Western Siberia* (ed. Zakh, V. A.) 134-139 (Inst. of Probl. of Devel. of the North SB RAS publ. house, 2001).

Kiryak, M. A. *An archeology of the Western Chukotka: in connection with the Yukaghir problem* (Nauka, 1993).

Kiryushin, K. Yu. Modern Concepts of the Cultural Affiliation of Neolithic Settlement Complexes in Altai. *Bulletin of Altay St. Univ.* **4-2**, 78-84 (2008).

Kiryushin, K. Yu. et al. On the relative and absolute chronology of early burials at the Firsovo-XI burial ground (Barnaul, Ob River region). *Bulletin of Archeology, Anthropology and Ethnography* Bulletin of Archeology, Anthropology and Ethnography **3** (**54**), 2071-0437-2021-54-3-2 (2021).

Kiryushin, K. Yu., Kiryushin, Yu. V. & Frolov, Ya. V. Tourist potential of funeral complexes of the Final Mesolithic - Neolithic ground burial ground Firsovo-XI (Barnaul Priobie). *Science and Tourism: Interaction Strategies* **13**, 51-67 (2021).

Kiryushin, K. Yu., Kiryushin, Yu. F., Solodovnikov, K. N., Frolov, Y. V. & Schmidt A. V. Results of Radiocarbon Dating of Early Burials in the Firsovo Archaeological Area, Barnaul Stretch of the Ob. *Archaeology, Ethnology & Anthropology of Eurasia* **49** (**3**), 1563-0110.2021.49.3.024-031 (2021).

Kiryushin, Yu. F. *Eneolithic and Early Bronze Age of the South of Western Siberia* (Altai Univ. publ. house, 2002).

Kiryushin, Yu. F. & Kiryushin, K. Yu. Problems of cultural belonging of the early burials of the Tuzovsky Mounds-I soil burial ground (One of the versions of historical and cultural interpretation). *Theory and Practice of Archaeological Research*, **12** (**2**), tpai(2015)2(12).-05 (2015).

Kiryshin, Yu. F., Kiryushin, K. Yu., Shmidt, A. V. & Abdulganeev M. T. Ornaments Made from Animal Teeth in Human Burials at Tuzovskiye Bugry-1 and Their Relevance to Ethnic Processes in the Altai, 3rd Millennium BC. *Archaeology, Ethology* & *Anthropology of Eurasia* **3** (**51**), 59-66 (2012).

Kiryushin, Yu. F., Kiryushin, K. Yu., Shmidt, A. V., Kuzmenkin, D. V., & Abdulganeyev, M. T. Mollusk shells in the burials of the Tuzovskiye Bugry-1 site as an indicator of ethnocultural processes in the territory of Southern Siberia and Central Asia. *Archeology, Ethnography and Anthropology of Eurasia* **2**, 37-45 (2011).

Kiryushin, Yu. F., Kungurova, N. Yu., & Kadikov, B. H. *The Oldest Burial Grounds of the Northern Foothills of Altai* (ed. Molodin, V. I) (Altai Univ. publ. house, 2000).

Kiryushin, Yu. F., Shamshin, A. B. & Shmidt, A. V. in *Modern solutions to the actual problems of Eurasian archaeology* (ed. Tishkin, A. A.) 137-140 (Altai Univ. publ. house, 2013).

Kochmar, N. N. *Petroglyphs of Yakutia* (Institute of Archaeology and Ethnography SB RAS publ. house, 1994).

Koksharov, S. F. *The Culture of the Population of the North of Western Siberia in the Bronze Age: Dissertation* (Novosibirsk, 2015).

Konikov, B. A. *Archaeology of Omsk: Illustrated Encyclopedia* (ed. Zhuk, A. V., Gur'yanova, G. G.) (Publisher, 2016).

Konopatsky, A. K. *Ancient cultures of Baikal (Olkhon Island)* (ed. Okladnikov, A. P.) (Nauka, 1982).

Korolev, A. I. in *Archaeology of the Volga-Ural Region* (ed. Sitdikov, A.G., Chizhevskiy, A.A.) **2**, 106-126 (AS Republic Tatarstan publ. house, 2021).

Korolev, A. I. & Shalapinin, A. A. The Chronology and Periodization of Forest Steppe and Steppe Eneolit in Volga Region. *Proceedings of the Samara Scientific Center of the Russian Academy of Sciences* **16** (**3-1**), 266-275 (2014).

Korolev, A. I. & Shalapinin, A. A. in *Proceedings of the VI (XXII) All-Russian Archaeological Congress in Samara* (ed. Derevyanko, A. P., Makarov, N. A. & Mochalov, O. D.) **1**, 162-164 (Samara state Socio-ped. Univ. publ. house, 2020).

Korolev, A.I. & Stavitskiy, V.V. in *Archaeology of the Volga-Ural Region* (ed. Sitdikov, A.G., Chizhevskiy, A.A.) **2**, 208-221 (AS Republic Tatarstan publ. house, 2021a).

Korolev, A.I. & Stavitskiy, V.V. in *Archaeology of the Volga-Ural Region* (ed. Sitdikov, A. G., Chizhevskiy, A. A.) **2**, 40-60 (AS Republic Tatarstan publ. house, 2021b).

Korolev, A. I., Kochkina, A. F. & Stashenkov, D. A. in *Proceedings of the VI (XXII) All-Russian Archaeological Congress in Samara* (ed. Derevyanko, A. P., Makarov, N. A. & Mochalov, O. D.) **1**, 159-161 (Samara state Socio-ped. Univ. publ. house, 2020).

Korolev, A. I., Myshkin, V. N. & Shalapinin, A. A. Results of Work at the Maximovka Barrow Cemetery and in the Forest-Steppe of the Volga Region in 2018. *Oriental Studies* **14** (**3**), 515-530 (2021).

Kostyleva E. L. & Utkin A. V. *Neo-Eneolithic burial grounds of the Upper Volga region and the Volga-Oka interfluve: Planigraphic and chronological structures* (TAUS, 2010).

Kostyleva, Y. L. & Utkin, A. V. The history of excavations of Fatyanovo culture burial mounds of the Bronze Age in Central Russia (On the 110th anniversary of D. A. Kraynov's birth). *Bulletin of the Ivanovo St. Univ. Series: Humanities* **4**, 62-68 (2014).

Kostyleva, E. L., Utkin, A. V. & Macane, A. in T*ver archaeological collection* (ed. Chernykh, I. N.) **11**, 565-577 (Triada Publishers Ltd, 2018).

Kovaleva, V. T. & Zyryanova, S. YU. *Neolithic of the Middle Trans-Urals: Boborykino Culture* (Center Educational book, 2010).

Kovalevskiy, S. A. (2016). *Irmen Antiquities of the South of Western Siberia: History of Study and Research Concepts* (Dissertation, Barnaul, 2016).

Kozintsev, A. G. Did Negroes Penetrate into Siberia in Ancient Times? *Voprosy antropologii* **47**, 191-196 (1974).

Kraynov, D. A. (1964). Volosovo-Danilovskiy Burial Ground of the Fatyanovo Culture. *Soviet Archaeology* **4**, 70 (1964).

Krenke, N. Radiocarbon chronology of the Fatyanovo culture. *Russian Archaeology* **2**, S086960630004830-2 (2019).

Kungurova, N. Ju. *Solontsy 5 Burial Ground. Culture of Neolithic Burials in Altai* (Barnaul Law institute, 2005).

Kungurova, N. YU. Materials from the Solontsy-5 Burial Ground (on the Issue of Ethnocultural Connections of the Bearers of the Kuznetsk-Altai Culture). *Archeology, Ethnography and Anthropology of Eurasia* **2** (**14**), 30-40 (2003).

Kuptsova, L. V., Zaretskaya, N. E., Morgunova, N. L. & Hohlov, A. A. The oldest burials in the Orenburg region (concerning two graves of the burial mound near Labazy village). *Vestnik Moskovskogo Universiteta. Seria 23. Antropologia* **1**, 131-139 (2019).

Kuzminykh, S. V. New data on the problem of the Seima-Turbino. *Brief Reports of the Institute of Archeology RAS* **225**, 240-263 (2011).

Kuzminykh, S. V. in *Mobility and Migration: Concepts, Methods and Results* (ed. Polos'mak, N. V. et al.) 89–103 (SB RAS publ. house, 2019).

Kuzminykh, S. V. in *The Great Russian Encyclopedia* (ed. Osipov, U. S. et al) **5**, 670-671 (2006).

Kuznetsov, P. in *Radiocarbon in Archaeology and Paleoecology: Past, Present, Future. Proceedings of the International Conference dedicated to the 80th anniversary of the Senior Researcher of IIMK RAS, Candidate of Chemical Sciences Ganna Ivanovna Zaytseva* (ed. Burova, N. D., Vybornov, A. A. & Kul'kova, M. A.) 978-5-91867-213-6-47-50 (Porto-print, 2020).

Lazaretov, I. P. & Poliakov, A. V. in Ethnocultural processes in the Upper Ob and adjacent regions at the end of the Bronze Age (ed. Shamshin, A. B.) 33-55 (Concept, 2008).

Lbova, L. V. & Zhambaltarova, E. D. Characteristics of the Burial Rites of the Early Neolithic in Western Trans-Baikal. *Bulletin of Novosibirsk State University. Series: History. Philology* **3** (**8**), 89-97 (2009).

Lbova, L. V., Zhambaltarova, E. D. & Konev, V. P. *The funeral complexes of the Neolithic – early Bronze Age of Trans-Baikal (the formation of archetypes of primitive culture)* (SB RAS publ. house, 2008).

Lychagina, E. L. On the Chronology and Periodization of the Neolithic in the Upper and Middle Kama Region. *Archaeology, Ethnography and Anthropology of Eurasia* **1** (**45**), 28-33 (2011).

Makarov, N. P. Chronology and Periodization of the Neolithic and Bronze Age in the Krasnoyarsk Forest-Steppe Zone. *News of the Laboratory of Ancient Technologies* **3**, 149-171 (2005).

Makarov, N. P. The Krasnoyask and Kansk forest steppe and its place in the cultural dialogue of Neolithic period and Bronze Age in Western and Eastern Siberia. *Bulletin of Tomsk State University. History* **4** (**42**), 94-97 (2016).

Maksimenkov, G. A. The Burial Ground near the Dolgoye Lake in Kansk. *Questions of Anthropology* **18**, 132-134 (1964).

Mamonov, A. Ye. in *Ancient Cultures of the Forest-Steppe Region of the Volga* (ed. Vasil'yev, I. B.) 3–25 (Samara state Ped. Univ. publ. house, 1995)

Mamonova, N. N. & Sulerzhitsky, L. D. The experience of dating according to C14 burials Pri-Baikal of the Holocene Epoch. *Soviet Archaeology* **1**, 19-33 (1989).

Mandryka, P. V. *Bronze and Early Iron Age in the Southern Taiga of the Middle Yenisei and Lower Angara. Dissertation* (Barnaul, 2018).

Mandryka, P. V. in Archaeology of North and Central Asia: new discoveries and results of interdisciplinary research (ed. Tishkin, A. A.) 978-5-7904-2526-4.2021.7 (2021).

Mandryka, P. V. & Senotrusova, P. O. New materials of the late Bronze Age from the Kansk-Rybinsk forest-steppe depression. *News of the Laboratory of Ancient Technologies* **14.1**, 2415-8739-2018-1-56-69 (2018).

Marchenko, Z. V. *Cultural affiliation, chronology, and periodization of archaeological sites in the middle course of the Tara River (Neolithic and Bronze Age). Dissertation* (Novosibirsk, 2009).

Marchenko, Zh. V., Svyatko, S. V., Grishin, A. E. & Rykun, M. P. in Proceedings of the V (XXI) All-Russian Archaeological Congress in Barnaul-Belokurikha (ed. Derevianko, A. P. & Tishkin, A. A.) **1**, 287-291 (Altai Univ. publ. house, 2017).

Marochkin, A. G. The Cultural and Historical Concept of the Neolithic and Early Metal of the Upper Ob Basin in the National Historiography of the 1970s – the Beginnings of the 2010s. *Bulletin of the Tomsk St. Ped. Univ.* **7** (**135**), 84-90 (2013).

Marochkin, A. G. *Burial practice of the Upper Ob region population during the Neolithic and Eneolithic periods (history of study, structural analysis and typology, problems of cultural and chronological interpretation). Dissertation* (Kemerovo, 2014).

Matyushchenko, V. I. (Ed.). The Lower Tara archaeological microdistrict (Nauka, 2001).

Maksimenkov, G. A. in Prehistory archeology of Siberia (ed. Mandelshtam, A. M.) 48-58 (Nauka, 1975).

Metel, I. V. History of the study of the complex archaeological site "Omsk Site". *Magistra Vitae: Electronic Journal of Historical Sciences and Archaeology* **2**, 2542-0275-2016-00016 (2016).

Mochanov, Yu. A. The Early Neolithic Aldan. *Soviet Archaeology* **2**, 126-136 (1966).

Mochanov, Yu. A. *Multiphase site Bel'kachi 1 and a periodization of the Stone Age of Yakutia* (ed. Okladnikov, A. P.) (Nauka, 1969).

Mochanov, Yu. A. *Oldest stages of human settlement in Northeast Asia* (Nauka, 1977).

Mochanov, Yu. A. & Fedoseeva, S. A. In: *Beringia in Cenozoic. Materials of the All-Union Symposium "The Bering Land and Its Significance for the Development of Holarctic Flora and Fauna in the Cenozoic"* (ed. Kontrimavichus, V. L.) 515-539 (ESC AS USSR publ. house, 1976).

Mochanov, Yu. A. & Fedoseeva, S. A. *Essays on the pre-written history of Yakutia. Stone Age* **2** (DaniAlmas, 2013).

Moiseyev, V.G. et al. A Metric Analysis of a Human Cranium from the Khatystyr Cave, Republic of Sakha (Yakutia). *Archaeology, Ethnology and Anthropology of Eurasia* **2** (**51**), 1563-0110.2023.51.2.142-152 (2023).

Molodin, V. I. *The Neolithic and Bronze Age of the forest-steppe region of the Ob-Irtysh area* (Nauka, 1977).

Molodin, V. I. *Archaeological site of Sopka-2 on the Omi River* **1** (Institute of Archaeology and Ethnography SB RAS publ. house, 2001).

Molodin, V. I. *Archaeological site of Sopka-2 on the Omi River* **3** (Institute of Archaeology and Ethnography SB RAS publ. house, 2012).

Molodin, V. I. The late Krotovo (Cherno-Ozerye) culture in the Irtysh forest-steppe, Western Siberia. *Archaeology, Ethnology and Anthropology of Eurasia* **1** (**57**), 49-54 (2014).

Molodin, V. I., Mylnikova, L. N. & Nesterova, M. S. The Vengerovo-2A Neolithic Cemetery, Southwestern Siberia: Results of a Multidisciplinary Study. *Archaeology, Ethnology and Anthropology of Eurasia* **2** (**44**), 1563-0102.2016.44.2.030-046 (2016).

Molodin, V. I., Mylnikova, L. N., Nesterova, M. S., Kobeleva, L. S. & Nenakhov, D. A. Baraba Culture of Early Neolithic Period. *Vestnik NSU. Series: History and Philology* **7** (**19**), 1818-7919-2020-19-7-69-93 (2020).

Molodin, V. I. et al. *The interdisciplinary research of Baraba forest-steppe populations (IV-I millennium BC): archaeological, paleogenetic and anthropological perspectives* (SB RAS publ. house, 2013).

Morgunova, N. in *Radiocarbon in Archaeology and Paleoecology: Past, Present, Future. Proceedings of the International Conference dedicated to the 80th anniversary of the Senior Researcher of IIMK*

*RAS, Candidate of Chemical Sciences Ganna Ivanovna Zaytseva* (ed. Burova, N. D., Vybornov, A. A. & Kul'kova, M. A.) 978-5-91867-213-6-59-61 (Porto-print, 2020).

Mosin, V. S. Central Ural in the 4th-8th millennia BC. Questions of the Archeology of the Urals **25**, 159-174 (2008).

Mosin, V. S. & Epimakhov, A. V. Radiocarbon chronology of the Trans-Urals Neolithic: spatial aspect. *Ural Historical Journal* **60** (**3**), 1728-9718-2018-3(60)-108-114 (2018).

Mosin, V. S., Bobrov, V. V. & Marochkin, A. G. New absolute dates for the Trans-Uralian and western Siberian Neolithic. *Archaeology, Ethnology and Anthropology of Eurasia* **4** (**45**), 1563-0102.2017.45.4.065-073 (2017).

Motuzaite Matuzeviciute et al. Climatic or dietary change? Stable isotope analysis of Neolithic-Bronze Age populations from the Upper Ob and Tobol River basins. *The Holocene* **26**, 0959683616646843 (2016).

Narasimhan, V. M. et al. The formation of human populations in South and Central Asia. *Science* **365** (**6457**), eaat7487 (2019).

Nikitin, V. V. in *Archaeology of the Volga-Ural Region* (ed. Sitdikov, A.G., Chizhevskiy, A.A.) **2**, 164-186 (AS Republic Tatarstan publ. house, 2021).

Okladnikov, A. P. *The antiquities of the Lena* **2** (Yakut State Printing House, 1946).

Okladnikov, A. P. Neolithic burials on Afontova Gora. *Brief Reports Of The Institute Of The History Of Material Culture* **25**, 7-13 (1949).

Okladnikov, A. P. *The antiquities of the Lena* **3** (Yakut State Printing House, 1950).

Okladnikov, A. P. *History of the Yakut ASSR* **1** (Yakut State Printing House, 1955).

Okladnikov, A. P. Neolithic and Bronze Age of the Baikal region. Part 3 (Glazkovo time): *Materials and Research in Archaeology of the USSR* **43** (AS USSR publ. house, 1955).

Okladnikov, A. P. From the history of ethnic and cultural ties of the Neolithic tribes of the Middle Yenisei (To the question of the origin of the Samoyed tribes). *Soviet Archaeology* **1**, 26-55 (1957).

Okladnikov, A. P. & Zaporojskaya, V. D. *The antiquities of the Middle Lena* (Nauka, 1972).

Okladnikov, A. P. *Neolithic Sites of the Angara* (Nauka, 1974).

Okladnikov, A. P. *Neolithic Sites of the Middle Angara: From the Mouth of the Beloy River to Ust-Uda* (SB Nauka, 1975)

Okladnikov, A. P. *Neolithic Sites of the Lower Angara: From Serovo to Bratsk* (Nauka, 1976).

Okladnikov, A.P. Neolithic and Bronze Age of the Cis-Baikal region: *Materials and Research in Archaeology of the USSR* **18** (AS USSR Publ, 1950).

Pestereva, K. A., Stepanov, A. D. & Dyakonov, V. M. in *Ancient cultures of Mongolia, Baikal Siberia and Northern China* (ed. Mandryka, P. V. et al) **1**, 74-82 (Siberian Federal University, 2016).

Petrov, A. I. *The Late Neolithic and Early Bronze Age in the Middle Irtish Region* (Omsk St. Univ. publ. house, 2014).

Petrova, N. A., Smotrova, V. I., Smotrova, G. I., Drozdov, N. I. & Larchenko, S. I. in *Archaeological discoveries of 1975* (ed. Rybakov, B. A.) 272-273 (Nauka, 1976).

Pitulko, V. V. & Kasparov, A. K. Archaeological dogs from the Early Holocene Zhokhov site in the Eastern Siberian Arctic. *Journal of Archaeological Science: Reports* **13**, j.jasrep.2017.04.003 (2017).

Pitulko, V. V. & Pavlova, E. Yu. Radiocarbon dating of cultural deposits of the Zhokhov site (New Siberia archipelago, Siberian Arctic). *Notes of the Institute of the History of Material Culture* **12**, 27-55 (2015).

Pitulko, V. V. et al. The ancient anthropological finds of the high latitude Arctic (the Zhokhov site, New Siberian islands). *Ural Historical Journal* **2**, 61-72 (2015).

Polyakov, A. V., Svyatko, S. V. & Stepanova, N. F. (2019). in *Phenomena of cultures in the Early Bronze Age of the steppe and forest-steppe belt of Eurasia: paths of cultural interaction in the V-III millennium BCE* (ed. Morgunova, N. L.) 181-187 (Orenburg State Ped. Univ. publ. house).

Poliakov, A. V. & Lazaretov, I. P. in The Past of Humankind as seen by the Petersburg Archaeologists at the Dawn of the Millenium (to the Centennial of the Russian Academic Archaeology) (ed. Vinogradov, Yu. A. et al) 188-202 (St. Petersburg Centre for Oriental Studies Publishers, 2019).

Poliakov, A. V. Chronology and cultural genesis of the Paleometal epoch sites in Minusinsk basins (Institute for the History of Material Culture RAS publ. house, 2022).

Poliakov, A. V. Review of the results of the initial stage of paleogenetic studies of the Bronze Age population of the Minusinsk Basins. *Theory and practice of archaeological research* **26 (2)**, tpai(2019)2(26).-08 (2019).

Polos'mak, N. V., Chikisheva, T. A. & Baluyeva, T. S. *Neolithic Burial Grounds of Northern Baraba* (Nauka, 1989).

Privalikhin, V. I. (1998). Sosnovy Mys - 2 - a new multi-temporal burial ground of Northern Priangary. *Siberian Inter-museum Collection* (ed. Paramonov, V. I.) 72-87 (1998).

Privalikhin, V. I. in *Yeniseyskaya provintsiya* (ed. Vdovin, A. S. et al). **4**, 300-310 (2009).

Privalikhin, V. I., Drozdov, N. I. & Makulov, V. I. in *Archaeological research of the antiquities of the Lower Angara and adjacent territories* (ed. Karnauhova, L. L.) 42-56 (2013).

Sal'nikov, K. V. A new variant of the Early Bronze Age culture in the Trans-Urals. *Brief Reports Of The Institute Of Archaeology* **85**, 3-10 (1961).

Sal'nikov, K. V. An attempt to classify the ceramics of the forest-steppe zone of the Trans-Urals. *Soviet Archaeology* **2**, 37-48 (1961).

Sal'nikov, K. V. in *Archeology and Ethnography of Bashkiria* (ed. Kuzeev, R. G. & Sal'nikov, K. V.) **1**, 16-58 (BB AS USSA publ. house, 1962).

Saveliev, N. A., Dudarek, S. P., Timoshchenko, A. A. & Lokhov, D. N. Sosnovyi Mys Burial Ground in the Northern Angara Region: Analysis of the Mortuary Tradition and the Results of Radiocarbon Dating. *Bulletin of the Irkutsk State University. Geoarchaeology, Ethnology, and Anthropology Series* **34**, 2227-2380.2020.34.3 (2020).

Savenkova, T. M. & Reys, E. S. in *Physical anthropology: methods, databases, scientific results* (ed. Groomov, A.V.) 49-59 (Lema, 2014).

Savenkova, T. M. & Makarov, N. P. in *Ancient cultures of Mongolia, Southern Siberia and Northern China: Transactions of the 9 International Conference* (ed. Bazarov B. V. & Kradin N. N) **1**, 158-162. (Buryat branch SO RAS publ. house, 2018).

Serikov, Yu. B. in *Questions of the Archeology of the Urals* (ed. Kosinskaya, L. L.) **21**, 120-143 (Ural St. Univ. publ. house, 1993).

Serikov, Yu. B.  in *Questions of the Archeology of the Urals* (ed. Kovaleva V. T.) **23**, 29-47 (Ural St. Univ. publ. house, 1998).

Shalapinin, A. A. Late Eneolithic Ceramic Complexes of the Forest-Steppe Trans-Volga Region. *Archeology, Ethnography and Anthropology of Eurasia* **1** (**46**), 27-33 (2018).

Sherstobitova, O.S. in *Integration of Archaeological and Ethnographic Research* (ed. Tataurov, S. F. & Tataurova, L. V. et al) 328-331 (Omsk state Ped. Univ. publ. house, 2007).

Sherstobitova, O. S. in *Ural Archaeological Meeting* (ed. Trufanov A. Ya.). Yekaterinburg-Surgut: **12**, 178-180 (Magellan, 2007).

Shilov, S. N. & Maslyuzhenko, D. N. in *Questions of the Archeology of the Urals* (ed. Kovaleva, V. T. et al) **24**, 165-191 (Ural St. Univ. publ. house, 2002).

Shirokov, V. N. *Ural Rock Art: Southern Ural* (AMB publ. house, 2009).

Shishlina, N., Turetsky, M. A. & van der Plicht, J. Radiocarbon dating of samples from the Lebyazhinka burial ground in the Eneolithic period: verification and interpretation of data. *News of the Samara Scientific Center RAS* **19-3**, 196-202 (2017).

Shmidt, A. V. Chronological interpretation of the Neolithic and Eneolithic burial complexes in the forest-steppe and foothill Altai. *Eurasia in the Cenozoic. Stratigraphy, paleoecology, cultures* **7**, 266-273 (2018).

Shorin, A. F. & Nokhrina, T. I. Funerary ritual of the Eneolithic population of the Urals. *Archeology, Ethnography and Anthropology of Eurasia* **2** (**2**), 96-106 (2000).

Shorin, A. F. & Shorina, A. A. Historiography of the Neolithic Trans-Urals: the Kozlov and Poludenskaya Cultures. *Bulletin of Archeology, Anthropology and Ethnography* **2** (**57**), 5-16 (2022).

Shorin, A. F., Vilisov, E. V. & Shorina, A. A. Basyanovo archaeological complex of the late Neolithic age: the foundation for allotting. *Russian Archaeology* **1**, 5-18 (2015).

Shunkov M. V. (Ed.). History of Siberia **1** (Inst. of Archaeology and Ethnography SB RAS publ. house, 2022).

Sikora, M., Pitulko, V. V., Sousa V. C. et al. The population history of northeastern Siberia since the Pleistocene. *Nature* **570** (**7760**), s41586-019-1279-z (2019).

Sokolov, V. N. in *Material culture of the ancient population of Eastern Siberia* (ed. Generalov, A. G.) 101-116 (Irkutsk Univ. publ. house, 1982).

Solodovnikov, K. N., Bagashev, A. N., & Savenkova, T. M. Areas of anthropological communities of the Neolithic population in the southern regions of Western and Central Siberia. *Bulletin of Tomsk State University. History* **68**, 158-167 (2020).

Solodovnikov, K., Bagashev, A., Tur, S., Gromov, A. V., Nechvoloda, A. & Kravchenko, G. Neolithic-Eneolithic paleoanthropological sources from the middle Irtysh area. *Bulletin of Archeology, Anthropology, and Ethnography* **3** (**46**), 2071-0437-2019-46-3-116-136 (2019).

Stefanov, V. I. The Forgotten Find from the Chernoozer'e Cemetery. *Russian Archaeology* **4**, 114-118 (2004).

Stepanov, A. Early Iron Age Dyupsya burial, Central Yakutia. *Archaeology, Ethnology and Anthropology of Eurasia* **1** (**38**), j.aeae.2010.05.004 (2010).

Stepanov, A. D., Kuzmin, Ya. V., Hodgins, G.W.L. & Jull A.J.T. Kyordyughen site, Ymyiakhtakh culture, Yakutia: An interpretation of the burial rite. *Archaeology Ethnology & Anthropology of Eurasia* **4** (**52**), 51–61 (2012).

Stepanov, A. D., Kuzmin, Y. V. & Jull, A. J. T. New chronological data of Early Iron age in Yakutia. *News of the Irkutsk St. Univ. Series: Geoarchaeology. Ethnology. Anthropology* **7**, 106-112 (2014).

Timoshchenko, A. A., Vybornov, A. V. & Rybin, E. P. Anzhevsky complex of archaeological site - new asynchronous burial in the territory of the Kansk-Rybinsk basin. Preliminary data. *Theory and practice of archaeological research* **2** (**22**), 99-116 (2018).

Timoshchenko, A. A. The Neolithic and the Bronze Age of the Kansk-Rybinsk Basin: Dissertation (Kemerovo, 2013).

Tishkin, A. A. (Ed.) History of Altai **1** (Altay Univ. publ. house, Constant., 2019).

Tolpeko, I. V. Some problems in the study of complex archaeological sites in the Tatar Uvala D. Okunevo. *Bulletin of Omsk University. Series: Historical Sciences* **3**, 71-74 (2015).

Troitskaya, T. N. in *Issues of the archaeology of Siberia* (ed. Troitskaya, T. N.) 84-101 (Novosibirsk ped. inst. publ. house, 1973).

Tsibin, V. A. & Shalapinin, A. A. The Eneolithic Burial of Maksimovka I flat Burial Ground from the Samara Trans-Volga Region. *Bulletin of Volgograd State University. History. Regional studies. International relations* **3** (**24**), jvolsu4.2019.3.1 (2019).

Ulanov, I. V. & Berdnikov, I. M. Ceramic complexes of Ust-Belaya: systematization, chronometry, chronology. *Bulletin of Irkutsk St. Univ. Geoarchaeology, Ethnology, Anthropology* **12**, 47–80 (2015).

Usacheva, I. V. Sosnovy Ostrov culture: identification of pottery and issues of chronology. *Bulletin of Archeology, Anthropology and Ethnographyi* **4** (**35**), 5-15 (2016).

Vdovin, A. S., & Makarov, N. P. in *Essays to the 80th Anniversary of German Ivanovich Medvedev* (ed. Berdnikov, I. M. & Lipnina, Ye. A.) 339-348. (Irkutsk St. Univ. publ. house, 2016).

Veber, A. V. & Link, D. V. Neolithic of the Baikal region: results and prospects of study. *Archeology, Ethnography and Anthropology of Eurasia* **1** (**5**), 135-146 (2001).

Viktorova, V. D. Sosnovy Ostrov - a settlement of the Neolithic and Bronze Age of the Middle Trans-Urals. *Soviet Archaeology* **4**, 161-173 (1968).

Vokhmentsev, M. P. *Eneolithic of the forest-steppe zone of the Transbaikal region: author's abstract of the Dissertation* (Moscow, 2000).

Volynkina, A. in *54 Ural-Volga Archaeological Student Conference* (ed. Vasil'yev, D. V.) 13-15. (Sorokin, 2022).

Vybornov, A. A., Kolev, Yu. I. & Mamonov, A. Ye. (Ed.). *History of the Samara Volga Region from Ancient Times to the Present Day: Stone Age* (Samara Scientific Center of RAS publ. house, 2000).

Vybornov, A. A., Mosin, V. S. & Epimakhov, A. V. Chronology of the Urals Neolithic. *Archeology, Ethnography and Anthropology of Eurasia* **1**, 33-48 (2014).

Vybornov, A., Lychagina, E., Vasilyeva, I. N., Melnichuk, A. & Kulkova, M. A. New data on the periodization and chronology of the Novoilinka, Garino, and Bor sites of the Kama region. Bulletin of Perm Univ. History **1** (**44**), 34-47 (2019).

Vybornov, A. A., Lebedeva (Ovchinnikova) N. V., Ovchinnikov, A. Yu. & Khokhlov, A. A. in *Aere perennius. More lasting than bronze. Essays in honour of Valentin Dergachev on the occasion of his 80th birthday* (ed. Dergacheva, L. V.) SL23VAD649 (Stratum Plus, 2023).

Weber, A. The Neolithic and Early Bronze Age of the Lake Baikal Region: A Review of Recent Research. *Journal of World Prehistory* **1** (**9**), 99–165 (1995).

Zaika, A. L. Neolithic burial in the mouth of the Kamenka River on the lower Angara. *News of the Laboratory of Ancient Technologies* **1** (**7**), 60-72 (2009).

Zakh, V. A. in *Western Siberia in Antiquity and the Middle Ages* (ed. Vasil'evsky, R. S.) 23-29 (Tumen St. Univ. publ. house, 1985).

Zakh, V.A. *Chronostratigraphy of the Neolithic and Early Metal Ages in the Forested Tobol-Ishim Region* (Nauka, 2009).

Zhambaltarova, E. D. & Volkov, P. V. Antler Spoons of the Fofanovo Cemetery. *Bulletin of the Novosibirsk St. Univ. Series: History. Philology* **15** (**3**), 32-40 (2016).

Zubova, A. V., Batanina, O. V., Panov, V. S., Stepanov, A. D. & Kishkurno M. S. Matta, a Neolithic burial in Central Yakutia. A result of the anthropological analysis. *Bulletin of Archeology, Anthropology and Ethnography* **4** (**39**), 2071-0437-2017-39-4-079-089 (2017).

### III. Population Genetic Analysis: PCA

***Data Merging and processing.*** For this analysis, we used a merged dataset consisting of 1957 present-day individuals genotyped on the Affymetrix Human Origins array [1], 398 shotgun-sequenced present-day individuals (CHB.SG, CHS.SG, CDX.SG and FIN.SG) from the 1000 genomes project dataset [2], 2 shotgun-sequenced present-day individuals (Mari.SG and Saami.SG) from [3] and [4], and 1347 ancient sequences, 281 of which were shotgun-sequenced (for which population labels end with *.SG*) and 1066 of which were produced from hybridization capture (*1240K dataset*; total data from [1–3,3–51]). The merged dataset contains 3304 sequences with sequence data at ~0.59 million positions.

***Method.*** All PCAs were carried out using the smartpca package of EIGENSOFT [52]. We used default parameters with the addition of two options (lsqproject:YES and numoutlieriter:0) to project the ancient individuals onto the PCA space. Three PCAs were constructed for projection, to highlight different dimensions of genetic variation among the studied populations. Positions without sequence data were treated as missing genotypes.

***Figure 1 of main text, center: Eurasian and American PCA.*** We used as a basis for the projection 1957 present-day West Eurasian, East Asian and Native American individuals from 122 populations genotyped on the Affymetrix Human Origins Array [1], specifically chosen to possess minimal recent sub-Saharan African and Oceanian admixture, and to all share the same sequencing and processing pipeline. All ancient individuals, as well as shotgun-sequenced present-day individuals, were projected onto this basis.

We find that the PCA separates three distinct groups of present-day populations: West Eurasians, East Asians, and Native Americans, with Siberian, Beringian and Athabaskan populations stretching between the East Asian and Native American positions on the PCA. We find noteworthy that:

1. The forest-steppe-hunter-gatherer cline (henceforth *FSHG cline*) is readily apparent as a continent-spanning cline stretching from present-day West Eurasians to present-day East Asians, but does not overlap similar clines that bridge West and East that are formed by present-day populations, because the center of the *FSHG* cline is deflected toward representatives of the *Ancient North Eurasian* (henceforth *ANE*) population, represented here by MA1_HG.SG and AfontovaGora3/AG3.
2. Some groupings of individuals from Eastern Siberia that do not fall genetically and geographically within the *FSHG* cline, such as Kolyma_M_10.1kya, Dzhilinda1_M_N_8.4kya, Syalakh-Belkachi, and *Yakutia_LNBA* are shifted off-cline in the direction of Native Americans.
3. The *Yakutia_LNBA* and Russia_Tatarka_BA populations fall close to each other and also close to present-day Nganasans. Uralic populations form a cline stretching from present-day West Eurasians to the *Yakutia_LNBA* and Samusakaya populations.

Populations used in underlying projection are: *Adygei, Altaian_Chelkan, Tubalar, Altaian, Nivh, Evenk_FarEast, Negidal, Nanai, Armenian_Hemsheni, Azeri, Bashkir, Buryat, Khamnegan, Circassian, Tabasaran, Darginian, Kubachinian, Lak, Avar, Kaitag, Evenk_Transbaikal, Ingushian, Kabardinian, Karelian, Veps, Abazin, Shor_Khakassia, Khakass, Khakass_Kachin, Kurd, Kazakh, Mordovian, Nogai_Astrakhan, Nogai_Stavropol, Ossetian, Russian_Archangelsk_Krasnoborsky, Russian_Archangelsk_Leshukonsky, Russian, Russian_Archangelsk_Pinezhsky, Shor_Mountain, Tatar_Siberian, Tatar_Siberian_Zabolotniye, Tatar_Kazan, Tatar_Mishar, Chuvash, Tuvinian, Todzin, Udmurt, Besermyan, Yukagir_Forest, Yukagir_Tundra, Chukchi, Eskimo_Naukan, Nganasan, Chukchi1, Ulchi, Even, Eskimo_ChaplinSireniki, Koryak, Itelmen, Tlingit, Yakut, Balkar, Chechen, Lezgin, Nogai_Karachay_Cherkessia, Kumyk, Kalmyk, Dolgan, Selkup, Ukrainian, Mansi, Aleut, Enets, Ket, Tofalar, Han, Tujia, Yi, Miao, Oroqen, Daur, Mongola, Hezhen, Xibo, Uyghur, Dai, China_Lahu, She, Naxi, Tu, Tibetan, Maonan, Zhuang, Kazakh_China, Gelao, Bonan, Yugur, Kyrgyz_China, Li, Dong, Mulam, Qiang, Salar, Tibetan_Yunnan, Dongxiang, Mongol, Finnish, Estonian, Norwegian, Belarusian, Kyrgyz_Kyrgyzstan, Karakalpak, Uzbek, Dungan, Turkmen, Iran_Zoroastrian, Mayan, Pima, Surui, Karitiana.*

***Extended Data Figure 4: PCA projected on ancient individuals rich in ANE ancestry.*** To further illuminate the role that levels of *ANE* ancestry plays among the populations we analyze, we use as a basis for another projection 71 shotgun-sequenced ancient individuals, of which a large proportion are enriched in *ANE* ancestry and fall outside the range of present-day variation (e.g. individuals from populations like *Tyumen_HG.SG* or *Kazakhstan_Botai.SG*; for a full list, see next paragraph). Of these, a number of individuals serving as representatives for *EEF* (Early European Farmer), *Iran_N* (Neolithic Iranian agropastoralists), *WHG* (*Western Hunter-gatherer*) and East Asian ancestries were also included. All other ancient and present-day individuals were projected onto this basis, with the hybridization capture versions of the individuals included in the basis (i.e. samples duplicated through different sequencing pipelines) excluded from the set of projected individuals; the PCA does not restrict to positions covered on the 71 individuals only. We find that the PCA separates Iran_N, EEF, *WHG*, *EHG* and *ANE* and East Asian ancestry, with each of these groups occuppying a different part of the plot. The forest-steppe-hunter-gatherer cline now forms a curved arc stretching from *EHG* populations to present-day East Asians; the center of the arc is strongly moved toward the positive direction in PC2, dominated by populations rich in *ANE* ancestry. Indeed, the individual furthest along the positive direction in PC2 is the AG3 individual, a relatively recent (~16kya) representative of near-unadmixed *ANE* ancestry and the closest representative of the population that contributed *ANE* admixture to later groups [3].

The basis was constructed using 71 shotgun-sequenced ancient individuals, with a large fraction of individuals with high levels of *ANE* ancestry. Genotyped versions of the same individuals were excluded from the set of projected individuals. The PCA positions of the basis individuals are not displayed. Individuals used in underlying projection are: *Bon001.SG, Bon002.SG, Bon004.SG, Bon005.SG, Tep002.SG, Tep003.SG, Tep004.SG, Bar31.SG, Bar8.SG, AH1.SG, AH2.SG, AH4.SG, WC1.SG, MA1.SG, I0061.SG, Hum1.SG, Hum2.SG, DA245.SG, DA246.SG, DA247.SG, DA248.SG, DA249.SG, DA250.SG, DA251.SG, DA252.SG, DA253.SG, DA340.SG, DA341.SG, DA357.SG, DA359.SG, DA362.SG, Sidelkino.SG, I0707.SG, I0708.SG, I0709.SG, I0745.SG, I0996.SG, I1098.SG, I1580.SG, I1581.SG, I1583.SG, I1763.SG, I1819.SG, I1947.SG, I1959.SG, I1960.SG, I1961.SG, I1962.SG, I1965.SG, I2134.SG, I2137.SG, I2139.SG, NEO236.SG, NEO238.SG, NEO239.SG, NEO240.SG, Yana_old.SG, Yana_old2.SG, I1290.SG, I12961.SG, I13097.SG, I2142.SG, I6413.SG, I6916.SG, brn001.SG, brn008.SG, cta016.SG, irk007.SG, Dzhilinda1_M_N_8.4kya, N2a.SG, yak025.SG.*

***Extended Data Figure 5: East Eurasian PCA.*** The basis was constructed using only present-day Eurasian and Native American populations genotyped on the Affymetrix Human Origins array **(Patterson et. al. 2012)**; the East Asian populations were chosen to have minimal West Eurasian admixture. Populations used in underlying projection: *Nivh, Evenk_FarEast, Negidal, Nanai, Evenk_Transbaikal, Eskimo_Naukan, Nganasan, Chukchi, Ulchi, Even, Koryak, Itelmen, Yakut, Han, Tujia, Yi, Miao, Oroqen, Daur, Mongola, Hezhen, Xibo, Dai, China_Lahu, She, Naxi, Tu, Tibetan, Maonan, Zhuang, Gelao, Bonan, Li, Dong, Mulam, Qiang, Tibetan_Yunnan, Norwegian.*

### IV. Population Genetic Analysis: ADMIXTURE

All relatives and shotgun sequences were excluded from ADMIXTURE analysis. For relative pairs or groups, the lower-coverage individual was excluded.

We used ADMIXTURE after pruning for linkage disequilibrium, after which 455,788 autosomal snps were retained. We used K=18 as the first K value where Yakutia_LNBA and Cisbaikal_LNBA were clearly separated from East Asian components characteristic of FSHG populations (e.g. the components maximized in Mongolia_N_North and AmurRiver_14K).

**{Figure here}**

**Extended Data Figure 10 | ADMIXTURE results**

### V. Population Genetic Analysis: Formal Grouping of Individuals in Northeastern Siberia into Populations

***Grouping Procedure.*** We use a similar method as that used in Kilinc. et. al. (2020) to group individuals in Eastern Siberia and the Cisbaikal region into populations, by running all f4-statistics [1] of the form f4(O, X1, X2, X3), where X1, X2 and X3 are individuals among the 22 and O is an outgroup population equally related to all (in this case *Ethiopia_4500_published.SG*, from [53]). When two individuals from the same population are placed in X1 and X2, and an individual who is not from that population is placed in X3, the statistic is expected to be negative, as members of the same population are expected to share drift exclusive to that group. Therefore, a valid clustering of individuals into populations should maximize negative statistics of the form f4(O, X1, X2, X3) where X1 and X2 are drawn from the same population, and X3 is drawn from another population. We perform the grouping procedure separately in two groups of individuals: first, all individuals in Siberia Northeast of Lake Baikal (i.e. the Transbaikal region and Eastern Siberia), and secondly all individuals in the Cisbaikal region (i.e. just west of lake Baikal). The two oldest individuals from Siberia, *Khairygas_16.7kya* (~16.7kya) and *Ust_Kyakhta_14kya* (~14kya), as well as another individual from close to the Bering Strait (*Kolyma_M_10.1kya*) from ~10kya are left ungrouped. We proceed to introduce all population labels produced by this analysis (which may in some cases apply to only a single ungrouped individual, but usually applies to multiple) that form the basis for further work on the deep population history of Siberia, starting with the two ungrouped individuals.

***Dataset and Method.*** For this analysis, we used a merged dataset consisting of 105 individuals from Siberia Northeast of Lake Baikal and the Cisbaikal region, just west of Lake Baikal; the dataset consists of 66 shotgun-sequenced samples and 39 hybridization-capture samples (*33*)[7–9,16,41,51] with data at ~1.24 million snps. All f4-statistics were performed using the *qpDstat* package of *ADMIXTOOLS* with the `f4mode: YES` parameter.

### *V.A. Ungrouped pre-Holocene individuals from Siberia*

***Khairygas_16.7kya and Ust_Kyakhta_14kya.*** Two of the oldest individuals from Siberia are left ungrouped: *Khairygas_16.7kya* from Khaiyrgas cave from ~16,700 BP, in the middle reaches of the Lena river in Southern Yakutia, and *Ust_Kyakhta_14kya* from ~14,000 BP, from just South of Lake Baikal. In PCAs, these individuals fall in a position indicating that they have broadly East Eurasian affinities, but contain significant amounts of ancestry related to *ANE* and to Native Americans (Main Text Fig. 1, center; Extended Data Figures 4, 5, 6). In ADMIXTURE, the two individuals behave similarly, with ~25% *Altai_N*- or *Tarim_EMBA*-related components and another ~25% of Native-American- or Beringian-related components, with the rest of their ancestry (~50%) modeled as being drawn from East Asian components (Extended Data Figure 10; Extended Data Figure 7).

***Kolyma_M_10.1kya.*** Another individual, *Kolyma_M_10.1kya* from the Kolyma River Valley close to the Arctic Sea and Beringia ~10,000 BP, is also left ungrouped. This individual clusters close to *Ust_Kyakhta_14kya* in PCAs, but is shifted further in the direction of Native Americans and Beringians (Main Text Fig. 1, center; Extended Data Figures 4, 5, 6). Fittingly, this individual when compared to *Khairygas_16.7kya* and *Ust_Kyakhta_14kya* has more (~40%) of its ancestry modeled as deriving from the Native American (peaking in South Americans), Beringian (peaking in Neo-Eskimos and present-day Inuit), and Chukotko-Kamchatkan (peaking in Itelmen and Chukchi) components, and less from East Asian and *ANE*-related components (Extended Data Figure 10; Extended Data Fig. 7).

The patterns displayed by *Khairygas_16.7kya*, *Ust_Kyakhta_14kya* and *Kolyma_M_10.1kya* are repeated in the later *Dzhilinda1_M_N_8.4kya* and *Syalakh-Belkachi* populations. In the main text, we define these populations as being rich in "Ancient Paloesiberian" (*APS*) ancestry—a term first used by [41] to refer to an ancestry source related to but deeply diverged from that found in Native Americans.

#### V.B. Grouping individuals from the Transbaikal region and Eastern Siberia

We present results for the grouping of Siberian individuals North and East of Lake Baikal in Table V. A-1. Geographic and temporal details are also list for each individual. Aside from newly sequenced samples, we also include samples from [16] from this region, for a total of 22 individuals.

***Results.*** We find that the clustering of individuals presented in Table V. A-1, under the column "Grouping", produces lists of statistics of the form f4(O, X1, X2, X3) where all significant statistics are negative when X1 and X2 come from the same group (Data S4 Table 2). The clusters are compatible with the results of sections III and IV, as individuals grouped into the same population are also extremely similar in PCA and ADMIXTURE (Extended Data Figure 10; Main Text Fig. 1, center; Extended Data Figures 7, 4, 5, 6). The grouping that results includes two unclustered individuals *KhatystyrCave_M_10.2kya* and *Dzhilinda1_M_N_8.4kya,* as well as three groups, which we have chosen to name *Syalakh-Belkachi*, *Transbaikal_EMN*, and *Yakutia_LNBA*. We recover all the groups also recovered in [16] but in the case of *Yakutia_LNBA* and *Syalakh-Belkachi* the groups are augmented by the newly-sequenced individuals in our study. All $f_4$ statistics produced in this analysis are accessible in Data S4 Table 2.

**Table S3**

| Individual ID | Source | Grouping | Details |
|---|---|---|---|
| KhatystyrCave_M_10.2kya | Newly sequenced, this study | *KhatystyrCave_M_10.2kya/Yakutia_Mesolithic* | ~10150 years BP, Southern Yakutia, on the Aldan River leading to the Middle Lena Basin |
| Dzhilinda1_M_N_8.4kya | Kilinc. et. al. (2020) | *Dzhilinda1_M_N_8.4kya* | ~8400 years BP, Northern Buryatia adjoining Southern Yakutia |
| N2a.SG | Kilinc. et. al. (2020) | *Syalakh-Belkachi* | ~6800 years BP, Middle Lena Basin, Central Yakutia |
| N5a.SG | Kilinc. et. al. (2020) | *Syalakh-Belkachi* | ~6200 years BP, Middle Lena Basin, Central Yakutia |
| I23448 | Newly sequenced, this study | *Syalakh-Belkachi* | ~5800 years BP, Middle Lena Basin, Central Yakutia |
| I15934 | Newly sequenced, this study | *Syalakh-Belkachi* | ~6300 years BP, Middle Lena Basin, Central Yakutia |
| brn001.SG | Kilinc. et. al. (2020) | *Transbaikal_EMN* | ~7300 years BP, Shilka Valley, Buryatia, Transbaikal Region |
| brn002.SG | Kilinc. et. al. (2020) | *Transbaikal_EMN* | ~6100 years BP, Onon/Upper Shilka Valley, Southern Buryatia, Transbaikal Region |
| brn003.SG | Kilinc. et. al. (2020) | *Transbaikal_EMN* | ~6600 years BP, Ingoda/Upper Shilka Valley, Buryatia, Transbaikal Region |
| brn008.SG | Kilinc. et. al. (2020) | *Transbaikal_EMN* | ~7400 years BP, Shilka Valley, Buryatia, Transbaikal Region |
| brn012.SG | Kilinc. et. al. (2020) | *Transbaikal_EMN* | ~6100 years BP, Onon Valley, Southern Buryatia, Transbaikal Region |
| cta016.SG | Kilinc. et. al. (2020) | *Transbaikal_EMN* | ~8300 years BP, Argun Valley, Buryatia, Transbaikal Region |
| irk007.SG | Kilinc. et. al. (2020) | *Transbaikal_EMN* | ~8800 years BP, Upper Lena, Cibaikal Region |
| kra001.SG | Kilinc. et. al. (2020) | *Yakutia_LNBA* | ~4200 years BP, Kan tributary of the Yenisei, far Southeastern Krasnoyarsk |

| | | | |
|---|---|---|---|
| N4a1.SG | Kilinc. et. al. (2020) | *Yakutia_LNBA* | ~4500 years BP, Middle Lena Basin, Central Yakutia |
| N4b2.SG | Kilinc. et. al. (2020) | *Yakutia_LNBA* | ~4200 years BP, Middle Lena Basin, Central Yakutia |
| yak021.SG | Kilinc. et. al. (2020) | *Yakutia_LNBA* | ~3200 years BP, Kolyma River Basin, Far Northeastern Yakutia |
| yak022.SG | Kilinc. et. al. (2020) | *Yakutia_LNBA* | ~3800 years BP, Kolyma River Basin, Far Northeastern Yakutia |
| yak024.SG | Kilinc. et. al. (2020) | *Yakutia_LNBA* | ~3500 years BP, Kolyma River Basin, Far Northeastern Yakutia |
| I20528 | Newly sequenced, this study | *Yakutia_LNBA* | ~3800 years BP, Middle Lena Basin, Central Yakutia |
| yak023.SG | Kilinc. et. al. (2020) | Not included in grouping procedure (first-degree relative of individual *yak024.SG* in this group), but falls into in *Yakutia_LNBA* | ~3700 years BP, Kolyma River Basin, Far Northeastern Yakutia |

***Populations produced from grouping procedure.*** We introduce all the populations produced by the grouping procedure here.

*KhatystyrCave_M_10.2kya.* This individual is found in the middle reaches of the Lena river, in Southern Yakutia ~10,100 BP. PCA indicates that this individual's ancestry is mostly East Asian. ADMIXTURE additionally indicates that this individual has very small proportions of Native-American-, Beingian-, or *Altai_N*-related ancestry (<10%), and that most of this individual's ancestry is Amur-River-related (>60% Amur-River-related component; Box 1), in line with its high level of shared drift with the *AmurRiver_LPaleolithic_14K* population in f4 statistics (Fig. 4B).

*Dzhilinda1_M_N_8.4kya.* This individual is found in Buryatia south of the Lena river, ~8400 BP. PCA indicates that this individual, like Khairygas_16.7kya, *Ust_Kyakhta_14kya*, and *Kolyma_M_10.1kya* has a significant amount of *ANE* ancestry. Like *Kolyma_M_10.1kya*, this individual is close to *Ust_Kyakhta_14kya* in all PCAs, but is shifted further to Native Americans and Beringians, yet not as much as *Kolyma_M_10.1kya* (Main text Fig 1, Center; Extended Data Figure 6). ADMIXTURE indicates that this individual has about 20% of *Altai_N/Tarim_EMBA*-related components and almost ~40% of Native-American/Bering Straits-related components, with the rest being East Asian-related, similar to *Kolyma_M.10kya* but with more East Asian ancestry (Extended Data Figure 7).

*Transbaikal_EMN.* In this group, two individuals are somewhat older than the rest: cta016.SG and irk007.SG, dating to ~8300 and ~8800 years BP respectively; the rest of the group dates to ~7300-6100 years BP. While the group is named after the Transbaikal region from which its members tend to be recovered, the irk007.SG individual was in fact recovered from the Western shores of Lake Baikal in the Cisbaikal region, while the individual cta016.SG was recovered from the Argun River valley, close to the Amur River. The two oldest individuals are thus the furthest West and East among the members of this cluster, respectively. The rest of this cluster, who are younger, are recovered from the river valleys of tributaries of the Shilka River, in Central-Southern Buryatia in the center of the Transbaikal region. The Shilka river catchment stretches from just east of Lake Baikal to merge with the Amur River further east. Individuals in this cluster are members of the *FSHG* cline in all PCAs (Main text Figure 1, center; Extended Data Figures 4, 5, 6). They fall close to other East Asians but are shifted away from them in the direction of *ANE*-rich populations of the *FSHG* cline, such as *Altai_N*, indicating that they have a small quantity of *ANE* ancestry and do not share very much drift with Native Americans and Beringians, unlike populations recovered in this grouping analysis from Northeast of Lake Baikal. In

ADMIXTURE, these individuals have high levels (~60%) of a component centered on *Mongolia_N*, and low levels of the *Yakutia_LNBA* and *AmurRiver_N*-centered components (~20%; Box 1).

*Syalakh-Belkachi.* All individuals in the *Syalakh-Belkachi* cluster come from the Middle to Lower Lena valley in Central Yakutia ~6800-6300 years BP. Individuals in the *Syalakh-Belkachi* cluster fall close to *Dzhilinda1_M_N_8.4kya* in PCAs, and therefore share the broad affinities of other individuals rich in APS ancestry (*Khairygas_16.7kya*, *Ust_Kyakhta_14kya*, *Kolyma_M_10.1kya* and *Dzhilinda1_M_N_8.4kya*) and are among the more East-Asian admixed among this group. However, individuals in this cluster are pulled away from *Dzhilinda1_M_N_8.4kya* in a straight line in the direction of *Yakutia_LNBA* in PCAs. Indeed, individuals in this cluster score ~10% in *Tarim_EMBA*- or *Altai-N*-related components and ~15% in the Beringian-related component, but also demonstrate a steadily increasing fraction of the *Yakutia_LNBA*-related component with time (~25% to ~40%), with the remainder derived from other East Asian components (Box 1).

*Yakutia_LNBA.* All individuals in the *Yakutia_LNBA* cluster date to ~4500-3200 years BP and fall in a region stretching from the Middle to Lower Lena valley in Central Yakutia to the Kolyma River basin in far Northeastern Yakutia close to Beringia, except for the Bronze Age individual *kra001.SG*, who instead was sampled from the Kan river valley in the Southeastern Krasnoyarsk region ~4200 years BP, far to the southwest. Individuals in this cluster have PCA positions indicating that they are broadly East Asian and have a low level of *ANE* ancestry and are closest to Nganasans among present-day populations (Main Text Fig. 1, center; Extended Data Figures 4, 5). In PCAs constructed with Native Americans and Beringians alongside Eurasians, individuals in *Yakutia_LNBA* are projected onto the cline formed by Native Americans, Beringians, and Siberians and East Asians, specifically in a spot between Beringians and Siberians close to where Nganasans fall (Main Text Fig. 1, center). In PCAs constructed with East Asian and West Eurasian populations, individuals in *Yakutia_LNBA* are projected onto a position between Nganasans and other Siberians, closer to Nganasans (Extended Data Fig. 5). Notably, Uralic populations form a cline that terminates on *Yakutia_LNBA* in all PCAs (Main Text Fig. 1, center; Extended Data Figures 4, 5). In ADMIXTURE, *Yakutia_LNBA* dominates a component that occurs in high levels among Uralic populations (Extended Data Fig. 7; Main text Fig. 3C; Extended Data Figure 10).

***Testing group homogeneity.*** We further test for evidence for intragroup heterogeneity within each of the three populations that are not singletons. Using *qpAdm*, we attempt to model each individual from the three groups as admixtures between other individuals of the group and a set of sources using an outgroup rotation framework (Harney et. al. 2021). Here, when a source is not used, it is left in the set of references (i.e., references are rotated into the sources), and all possible combinations of references rotated into the sources are tested.

The set of references/sources include *China_Paleolithic* (a combination of the Tianyuan individual from North China, ~45,000 BP, with the Upper Paleolithic individual from the Amur River Basin at ~33,000 BP reported in **Mao et. al. 2020)***, Japan_HG_Jomon, MA1_HG.SG, EHG, Russia_AfontovaGora3, Peru_Laramate_900BP, China_SEastAsia_Coastal_EN* (the Qihe individual from Southeast Coastal China, ~8,400 years BP)*, China_NEastAsia_Coastal_EN* (a group comprising the Bianbian, Boshan, Xiaojingshan and Xiaogao individuals from Northeast coastal China from ~9,700-7,800 years BP, from **Yang et. al. 2020)***, China_NEastAsia_Inland_EN* (the *Yumin* individual from inland Northern China, ~8,400 years BP from **Yang et. al. 2020)***,* and populations from the time transect of the Amur region in **Mao et. al. (2020)**: *China_AmurRiver_LPaleolithic_19k, China_AmurRiver_LPaleolithic_14k, China_AmurRiver_Mesolithic,* and *China_AmurRiver_EarlyN*.

We discover in almost all cases that individuals in each population may be successfully modeled ($p > 0.05$) in 1-way models with complete descent from another member of the same population, even with the extensive set of reference populations listed above. Individuals that cannot be modeled in 1-

way models with another member of their population tend to be successfully modeled in 2-way models with near-complete descent from another member of the population, plus a non-significant contribution from the source that has been rotated in (i.e., with admixture proportions much smaller than the standard error). This suggests that the grouping procedure has identified populations which are for the most part cladal to the exclusion of a wide range of other populations across Eurasia. All models for this analysis can be found in SI Data S4 Table 4.

#### *V.C. Grouping individuals from the Cisbaikal region and Western Siberia*

***Rationale.*** While all populations of the Cisbaikal region fall on the *FSHG* cline between the *ANE*-rich populations of the Altai and the East Asian populations of the Transbaikal region, ADMIXTURE analysis at higher K (K = 18, SI Section IV) suggests that a distinct component of ancestry predominates among Late Neolithic, Eneolithic, and Bronze Age groups of the Cisbaikal region that we term *Cisbaikal_LNBA*, that was not present in the early Neolithic groups of the Cisbaikal region on the *FSHG* cline. This ancestry appears at very high levels in an individual of the Middle Angara River valley by the early Bronze Age, and two outliers of the Karasuk horizon in the Minusinsk basin in the late Bronze Age. This component, when it appears, also shows a restricted distribution among modern populations, peaking in speakers of Yeniseian (Kets), Samoyedic, and South Siberian Turkic languages. The appearance of this component suggests marked discontinuity in the population history of the Cisbaikal region, during the Early to Late Neolithic, when *Cisbaikal_EN* was replaced by *Cisbaikal_LNBA* (Extended Data Figure 1, Extended Data Figure 7).

Because ADMIXTURE results should not be overinterpreted [54], we test using formal methods the inference that individuals with high levels of the *Cisbaikal_LNBA* component in admixture in the Late Neolithic-Bronze Age periods are especially closely related. We therefore re-run the clustering analysis for individuals from the Cisbaikal region from the EN to BA (Bronze Age) periods.

***Grouping Procedure.*** We included individuals of broadly East Eurasian affinity from the Cisbaikal region, just west of Lake Baikal in our clustering analysis (78 individuals), from [7–9,41,51], in addition to 4 newly-sampled individuals and one merger of two previously-published sequences. Details for each individual are listed in Table V. B-2. We use the same method as that used in the previous section (V. A) to group individuals.

***Results.*** We find that the grouping presented in Table IV. B-1 produces lists of statistics of the form f4(O, X1, X2, X3) within which all significant statistics are negative when X1 and X2 come from the same group. All f-4 statistics produced in this analysis are accessible in SI Data S4, Table 3.

**Table S4**

| Individual ID | Source | Grouping | Details |
|---|---|---|---|
| DA340.SG | *Cisbaikal_EN* | Damgaard et. al. (2018a) | ~7000 years BP, site of Lokomotiv within city of Irkutsk on the headwaters of the Angara River |
| DA341.SG | *Cisbaikal_EN* | Damgaard et. al. (2018a) | ~7600 years BP, site of Lokomotiv within city of Irkutsk on the headwaters of the Angara River |
| DA357.SG | *Cisbaikal_EN* | Damgaard et. al. (2018a) | ~7800 years BP, site of Lokomotiv within city of Irkutsk on the headwaters of the Angara River |
| DA359.SG | *Cisbaikal_EN* | Damgaard et. al. (2018a) | ~7500 years BP, site of Lokomotiv within city of Irkutsk on the headwaters of the Angara River |
| DA245.SG | *Cisbaikal_EN* | Damgaard et. al. (2018a) | ~8000 years BP, site of Shamanka II on the southern tip of Lake Baikal |

| | | | |
|---|---|---|---|
| DA246.SG | *Cisbaikal_EN* | Damgaard et. al. (2018a) | ~7700 years BP, site of Shamanka II on the southern tip of Lake Baikal |
| DA247.SG | *Cisbaikal_EN* | Damgaard et. al. (2018a) | ~7700 years BP, site of Shamanka II on the southern tip of Lake Baikal |
| DA248.SG | *Cisbaikal_EN* | Damgaard et. al. (2018a) | ~7600 years BP, site of Shamanka II on the southern tip of Lake Baikal |
| DA249.SG | *Cisbaikal_EN* | Damgaard et. al. (2018a) | ~7800 years BP, site of Shamanka II on the southern tip of Lake Baikal |
| DA250.SG | *Cisbaikal_EN* | Damgaard et. al. (2018a) | ~7400 years BP, site of Shamanka II on the southern tip of Lake Baikal |
| DA251.SG | *Cisbaikal_EN* | Damgaard et. al. (2018a) | ~7300 years BP, site of Shamanka II on the southern tip of Lake Baikal |
| DA252.SG | *Cisbaikal_EN* | Damgaard et. al. (2018a) | ~7300 years BP, site of Shamanka II on the southern tip of Lake Baikal |
| DA253.SG | *Cisbaikal_EN* | Damgaard et. al. (2018a) | ~7200 years BP, site of Shamanka II on the southern tip of Lake Baikal |
| DA362.SG | *Cisbaikal_EN* | Damgaard et. al. (2018a) | ~7200 years BP, site of Shamanka II on the southern tip of Lake Baikal |
| I7759 | *Cisbaikal_EN* | Flegontov et. al. (2019) | ~7000 years BP, site of Ust-Belaya, mouth of the Belaya river at the headwaters of the Angara River |
| irk051.SG | *Cisbaikal_EN* | Kilinc et. al. (2020) | ~8000 years BP, within city of Irkutsk on the headwaters of the Angara River |
| I0999 | *Cisbaikal_EN* | Newly sequenced, this study | ~7700 years BP, Kachug district, just West of Lake Baikal |
| IUO001 | *Cisbaikal_EN* | Yu et. al. (2020) | ~6900 years BP, headwaters of the Lena river, just West of Lake Baikal |
| KAG001 | *Cisbaikal_EN* | Yu et. al. (2020) | ~6900 years BP, Kachug district, just West of Lake Baikal |
| STB002 | *Cisbaikal_EN* | Yu et. al. (2020) | ~6580 years BP, headwaters of the Lena River just West of Lake Baikal |
| ANG001 | Unclustered, but close to *Cisbaikal_EN* | Yu et. al. (2020) | ~7300 years BP, headwaters of the Angara river, just West of Lake Baikal |
| NEO229.SG | Unclustered, but close to *Cisbaikal_EN* | SikoraNature2019 | ~6600 years BP, site of Ust-Balaya, mouth of the Belaya river at the headwaters of the Angara River |
| DA343.SG | *Cisbaikal_LNBA* | Damgaard et. al. (2018b) | ~4900 years BP, site of Ust-Ida on the Upper Angara river |
| DA353.SG | *Cisbaikal_LNBA* | Damgaard et. al. (2018b) | ~4300 years BP, site of Ust-Ida on the Upper Angara river |
| DA356.SG | *Cisbaikal_LNBA* | Damgaard et. al. (2018b) | ~4300 years BP, site of Ust-Ida on the Upper Angara river |
| DA361.SG | *Cisbaikal_LNBA* | Damgaard et. al. (2018b) | ~4100 years BP, site of Ust-Ida on the Upper Angara river |
| DA358.SG | *Cisbaikal_LNBA* | Damgaard et. al. (2018a) | ~4700 years BP, site of Kurma XI on western short of Lake Baikal |
| DA360.SG | *Cisbaikal_LNBA* | Damgaard et. al. (2018a) | ~4700 years BP, site of Kurma XI on western short of Lake Baikal |
| DA334.SG | *Cisbaikal_LNBA* | Damgaard et. al. (2018a) | ~4100 years BP, site of Shamanka II on the southern tip of Lake Baikal |
| DA336.SG | *Cisbaikal_LNBA* | Damgaard et. al. (2018a) | ~4200 years BP, site of Shamanka II on the southern tip of Lake Baikal |
| DA337.SG | *Cisbaikal_LNBA* | Damgaard et. al. (2018a) | ~4300 years BP, site of Shamanka II on the southern tip of Lake Baikal |

| | | | |
|---|---|---|---|
| DA339.SG | *Cisbaikal_LNBA* | Damgaard et. al. (2018a) | ~4000 years BP, site of Shamanka II on the southern tip of Lake Baikal |
| I1526 | *Cisbaikal_LNBA* | Flegontov et. al. (2019) | ~4800 years BP, site of Ust-Belaya, mouth of the Belaya river at the headwaters of the Angara River |
| I7779 | *Cisbaikal_LNBA* | Flegontov et. al. (2019) | ~4800 years BP, site of Ust-Belaya, mouth of the Belaya river at the headwaters of the Angara River |
| I7782 | *Cisbaikal_LNBA* | Flegontov et. al. (2019) | ~4600 years BP, site of Ust-Belaya, mouth of the Belaya river at the headwaters of the Angara River |
| I7335 | *Cisbaikal_LNBA* | Flegontov et. al. (2019) | ~4500 years BP, site of Ust-Belaya, mouth of the Belaya river at the headwaters of the Angara River |
| I8295_published | *Cisbaikal_LNBA* | Flegontov et. al. (2019) | ~4500 years BP, site of Ust-Belaya, mouth of the Belaya river at the headwaters of the Angara River |
| I8296_published | *Cisbaikal_LNBA* | Flegontov et. al. (2019) | ~4700 years BP, site of Ust-Belaya, mouth of the Belaya river at the headwaters of the Angara River |
| I7780 | *Cisbaikal_LNBA* | Merger of sequences from Flegontov et. al. (2019) and Harney et. al. (2021) | ~4600 years BP, site of Ust-Belaya, mouth of the Belaya river at the headwaters of the Angara River |
| irk025.SG | *Cisbaikal_LNBA* | Kilinc et. al. (2020) | ~4350 years BP, Upper Angara river |
| irk017.SG | *Cisbaikal_LNBA* | Kilinc et. al. (2020) | ~5100 years BP, headwaters of the Lena river just West of Lake Baikal |
| irk022.SG | *Cisbaikal_LNBA* | Kilinc et. al. (2020) | ~5100 years BP, Upper Angara river |
| irk033.SG | *Cisbaikal_LNBA* | Kilinc et. al. (2020) | ~4300 years BP, Upper Angara river |
| irk036.SG | *Cisbaikal_LNBA* | Kilinc et. al. (2020) | ~4700 years BP, Upper Angara river |
| irk061.SG | *Cisbaikal_LNBA* | Kilinc et. al. (2020) | ~4300 years BP, headwaters of the Lena river just West of Lake Baikal |
| irk068.SG | *Cisbaikal_LNBA* | Kilinc et. al. (2020) | ~4500 years BP, headwaters of the Lena river just West of Lake Baikal |
| irk071.SG | *Cisbaikal_LNBA* | Kilinc et. al. (2020) | ~5100 years BP, western shore of Lake Baikal |
| irk075.SG | *Cisbaikal_LNBA* | Kilinc et. al. (2020) | ~5400 years BP, western shore of Lake Baikal |
| irk034.SG | *Cisbaikal_LNBA* | Kilinc et. al. (2020) | ~4700 years BP, Upper Angara river |
| irk040.SG | *Cisbaikal_LNBA* | Kilinc et. al. (2020) | ~4700 years BP, Upper Angara river |
| irk057.SG | *Cisbaikal_LNBA* | Kilinc et. al. (2020) | ~4700 years BP, Upper Angara river |
| irk008.SG | *Cisbaikal_LNBA* | Kilinc et. al. (2020) | ~3800 years BP, headwaters of the Lena river just West of Lake Baikal |
| mak026.SG | *Cisbaikal_LNBA* | Kilinc et. al. (2020) | ~4800 years BP, headwaters of the Lena river just West of Lake Baikal |
| I1001 | *Cisbaikal_LNBA* | Newly sequenced, this study | ~4900 years BP, Kachug district, just West of Lake Baikal |
| I1000 | *Cisbaikal_LNBA* | Newly sequenced, this study | ~4600 years BP, Kachug district, just West of Lake Baikal |
| I2135 | *Cisbaikal_LNBA* | Newly sequenced, this study | ~4600 years BP, western shore of Lake Baikal |
| I0998 | *Cisbaikal_LNBA* | Newly sequenced, this study | ~4500 years BP, Orkhon island, off the western shore of Lake Baikal |

| | | | |
|---|---|---|---|
| KAG002 | *Cisbaikal_LNBA* | Yu et. al. (2020) | ~3700 years BP, headwaters of the Lena River just west of Lake Baikal |
| KPT001 | *Cisbaikal_LNBA* | Yu et. al. (2020) | ~3800 years BP, headwaters of the Lena River just west of Lake Baikal |
| KPT002_petrous | *Cisbaikal_LNBA* | Yu et. al. (2020) | ~3900 years BP, headwaters of the Lena River just west of Lake Baikal |
| KPT003 | *Cisbaikal_LNBA* | Yu et. al. (2020) | ~3900 years BP, headwaters of the Lena River just west of Lake Baikal |
| KPT004 | *Cisbaikal_LNBA* | Yu et. al. (2020) | ~3800 years BP, headwaters of the Lena River just west of Lake Baikal |
| KPT006 | *Cisbaikal_LNBA* | Yu et. al. (2020) | ~3600 years BP, headwaters of the Lena River just west of Lake Baikal |
| STB001 | *Cisbaikal_LNBA* | Yu et. al. (2020) | ~4600 years BP, headwaters of the Lena River just west of Lake Baikal |
| ZPL001 | *Cisbaikal_LNBA* | Yu et. al. (2020) | ~4000 years BP, headwaters of the Lena River just west of Lake Baikal |
| ZPL002 | *Cisbaikal_LNBA* | Yu et. al. (2020) | ~4200 years BP, headwaters of the Lena River just west of Lake Baikal |
| DA342.SG | Unclustered, but close to *Cisbaikal_LNBA* | Damgaard et. al. (2018a) | ~5600 years BP, site of Ust-Belaya, mouth of the Belaya river at the headwaters of the Angara River |
| DA344.SG | Unclustered, but close to *Cisbaikal_LNBA* | Damgaard et. al. (2018a) | ~5600 years BP, site of Ust-Belaya, mouth of the Belaya river at the headwaters of the Angara River |
| DA345.SG | Unclustered, but close to *Cisbaikal_LNBA* | Damgaard et. al. (2018a) | ~5400 years BP, site of Ust-Belaya, mouth of the Belaya river at the headwaters of the Angara River |
| I8298_published | Clustered with irk030.SG, close to *Cisbaikal_LNBA*. Treated as a set of outliers to be modeled in section VI. A. 6. | Flegontov et. al. (2019) | ~5700 years BP, site of Ust-Belaya, mouth of the Belaya river at the headwaters of the Angara River |
| irk030.SG | Clustered with I8298_published, but close to *Cisbaikal_LNBA*. Treated as a set of outliers to be modeled in section VI. A. 6. | Kilinc et. al. (2020) | ~6000 years BP, headwaters of the Lena River just west of Lake Baikal |
| DA335.SG | Unclustered, but close to *Cisbaikal_LNBA* | Damgaard et. al. (2018a) | ~4200 years BP, site of Shamanka II on the southern tip of Lake Baikal |
| GLZ003 | Unclustered, but close to *Cisbaikal_LNBA* | Yu et. al. (2020) | ~4500 years BP, site of Glazkovskoe Predmestie within city of Irkutsk on the headwaters of the Angara River |
| irk050.SG | Unclustered, but close to *Cisbaikal_LNBA* | Kilinc et. al. (2020) | ~4350 years BP, Upper Angara river |
| GLZ001 | Clustered with GLZ002, but close to *Cisbaikal_LNBA*. Treated as a set of outliers to be modeled in section VI. A. 6. | Yu et. al. (2020) | ~4600 years BP, site of Glazkovskoe Predmestie within city of Irkutsk on the headwaters of the Angara River |
| GLZ002 | Clustered with GLZ001, but close | Yu et. al. (2020) | ~4400 years BP, site of Glazkovskoe Predmestie within city of Irkutsk on the headwaters of the Angara River |

| | | | |
|---|---|---|---|
| | to *Cisbaikal_LNBA*. Treated as a set of outliers to be modeled in section VI. A. 6. | | |
| DA355.SG | Unclustered, but close to *Cisbaikal_LNBA* | Damgaard et. al. (2018a) | ~5500 years BP, site of Ust-Belaya, mouth of the Belaya river at the headwaters of the Angara River |

***Populations produced from grouping procedure.*** The clustering analysis produces two temporally separated groups: *Cisbaikal_EN,* whose individuals span ~8,000 to ~6,850 years BP, and *Cisbaikal_LNBA*, whose individuals span ~5,100 to ~3,800 years BP. The rest of the individuals, who are mostly singletons, are close to the two groups. Further description of the singleton individuals is provided in Data S4 Table 1.

*Cisbaikal_EN.* All individuals in the *Cisbaikal_EN* (~8,000 to ~6,850 years BP) fall in a region stretching from the western shores of Lake Baikal to the upper reaches of the Lena River just west of it. In all PCAs, individuals in this cluster are projected onto the *FSHG* cline, and help to bridge between 1) populations in the Krasnoyarsk region and the Altai region further west, and 2) populations in the *Transbaikal_EMN* cluster from the eponymous region covered in the previous section. In ADMIXTURE, individuals in the *Cisbaikal_EN* cluster are once again similar to each other, but demonstrate some heterogeneity. Samples with more East Asian ancestry tend toward increasing levels of the components found in *Transbaikal_EMN*, while populations with less East Asian ancestry tend to toward increasing levels of those components also found among APSs, with the difference that the *Altai_N*-related and *Cisbaikal_LNBA*-related components increase preferentially compared to their Native American and Beringian-related components.

*Cisbaikal_LNBA.* All individuals in the *Cisbaikal_LNBA* cluster (~5,100 to ~3,800 years BP) also fall in a region stretching from the western shores of Lake Baikal to the upper reaches of the Lena River just west of it. In most PCAs, individuals in this cluster are projected onto the *FSHG* cline, and form a tight cluster close to Cisbaikal_EN, indicating that this population has broadly East Asian affinities with some *ANE* ancestry. In ADMIXTURE, individuals in the *Cisbaikal_LNBA* cluster are extremely homogeneous and dominate a component that appears at moderate levels among Kets, Samoyeds, and South Siberian Turkic populations.

### VI. Population Genetic Analysis: qpAdm & Formal Statistics

#### *VI.A. The Deep Population History of Eastern Siberia*

##### VI.A.i. Modelling Approach and Parameters

We use a chronology-dependent sequential modelling approach to uncover the interrelationships between the populations in Northeastern Siberia uncovered by the grouping procedure used in the previous section. At the broadest level, we split populations in Northeastern Siberia into tranches by time period, and model the populations in each tranche as admixtures of populations from previous tranches. In other words, we model the populations in each tranche as descending from populations that are likely to have been older.

In our description of the tranches, which follows, all population labels are italicized:

- Tranche 0. This tranche will be used as references and sources that are included in all later analyzes. Populations in this tranche are likely to have diverged very deeply from those that contributed to the genetic history of Northeastern Siberia after the Last Glacial Maximum

~20,000 BP. These include *China_Paleolithic* (a combination of the Tianyuan individual from North China, ~45,000 BP, with the Upper Paleolithic individual from the Amur River Basin at ~33,000 BP reported in [22], *Japan_HG_Jomon, MA1_HG.SG, Russia_AfontovaGora3, Peru_Laramate_900BP, China_SEastAsia_Coastal_EN* (the Qihe individual from Southeast Coastal China, ~8,400BP)*, China_NEastAsia_Coastal_EN* (a group comprising the Bianbian, Boshan, Xiaojingshan and Xiaogao individuals from Northeast coastal China from ~9,700-7,800 BP, from [49]*, China_NEastAsia_Inland_EN* (the *Yumin* individual from inland Northern China, ~8,400 BP from [49]). Populations from the time transect of the Amur region in [22] are also included: *China_AmurRiver_LPaleolithic_19k, China_AmurRiver_LPaleolithic_14k, China_AmurRiver_Mesolithic,* and *China_AmurRiver_EarlyN*, to take into account the fact that ancestry from any of these populations resident in the Amur River region may have contributed to Northeastern Siberians.

- Tranche 1. This tranche includes only the *Khaiyrgas* individual, at ~16,700 BP.
- Tranche 2. This tranche includes only the *Ust_Kyakhta_14kya* individual, at ~14,000 BP.
- Tranche 3. This tranche includes the *Kolyma_M_10.1kya* individual and the KhatystyrCave_M_10.2kya/*Yakutia_Mesolithic* individual, both from ~10,000 BP.
- Tranche 4. This tranche includes a population labeled *Transbaikal_EMN_old*, which encompasses the two individuals (irk007.SG and cta016.SG), at ~8,300 & ~8,800 BP, that were split off from the rest of the *Transbaikal_EMN* cluster (~7,400 to ~6,100 BP) due to their age. The last population is the *Dzhilinda1_M_N_8.4kya* individual. As deducible from our list, all groups in this tranche fall between ~10,000 BP to ~8,000 BP.
- Tranche 5. This tranche included the *Syalakh-Belkachi*, *Yakutia_LNBA*, and *Transbaikal_EMN* groups that resulted from the grouping procedure undertaken in {Supplementary Information 1, section V} (in this document). All individuals in this tranche are younger than ~8,000 BP. As mentioned before, two individuals (cta016.SG and irk007.SG) were removed from the *Transbaikal_EMN* population we will use in *qpAdm* for tranche 5 into *Transbaikal_EMN_old* that we use in tranche 4, leaving only individuals younger than 8,000BP in *Transbaikal_EMN*. This tranche also includes *China_Amur_River_N* (individuals younger than 8,000 BP from the Amur Basin, Mao et. al. 2020) and *Altai_N*. *Altai_N* is a group that unites individuals from *Kemerovo_N, Firsovo_N* and *West_Siberia_N*—populations that are near-identical in unsupervised clustering, including PCAs (Fig. S1-S4) and ADMIXTURE (Fig. S5).

The identities of the individuals included under each population label can be found in Data S1 Table 4.

A table with populations and their initial grouping into tranches is presented below.

**Table S5**

| Tranche Number | Populations Included | Date Range | Notes |
|---|---|---|---|
| Tranche 0 | *China_Paleolithic, Japan_HG_Jomon, MA1_HG.SG, Russia_AfontovaGora3, Peru_Laramate_900BP, China_SEastAsia_Coastal_EN, China_NEastAsia_Coastal_EN, China_NEastAsia_Inland_EN, China_AmurRiver_LPaleolithic_19k, China_AmurRiver_LPaleolithic_14k, China_AmurRiver_Mesolithic,* and *China_AmurRiver_EarlyN,* | -- | Used as references/sources only. Members of other tranches are used as both targets to be modeled, as well as references/sources for models in later tranches |
| Tranche 1 | *Khaiyrgas* | ~16,700 BP | |

| | | |
|---|---|---|
| Tranche 2 | *Ust_Kyakhta_14kya* | ~14,000 BP |
| Tranche 3 | *Kolyma_M_10.1kya, Yakutia_Mesolithic* | ~10,000 BP |
| Tranche 4 | *Transbaikal_EMN_old, Dzhilinda1_M_N_8.4kya* | Between 10,000 BP, and 8,000 BP |
| Tranche 5 | *Syalakh-Belkachi, Cisbaikal_EN, Mongolia_N,* (*13*, *24*, *66*)*Transbaikal_EMN, Amur_River_N, Altai_N* | Between 8,000 BP and 6,000 BP |
| Tranche 6 | *Yakutia_LNBA, Cisbaikal_LNBA* | Between 6,000 BP and 3,500 BP |

For any population in each tranche, we test two sets of models using outgroup rotation (Harney et. al. 2021):

1. A model with populations in previous tranches in the references, rotating into the sources. We term this set of models ‘A’.
2. A model with previous tranches and all populations in the current tranche other than the target in the references, rotating into the sources. We term this set of models ‘B’.

Except in cases where some populations in the references/sources are unlikely to be direct contributors to the target (for example, the Tianyuan-like *China_Paleolithic* population from 45-33kya, or *ANE* ancestry from the pre-LGM *AG3,* are not likely contributors to a 3kya-year-old sample), the sets of models ‘A’ and ‘B’ rotate all eligible populations in the current or previous tranches into the sources in in all possible 1-, 2-, 3- or 4-way admixture combinations. Therefore, these sets exhaust all possible models of the target population as admixtures of current or previous populations. In the upcoming sections, we provide tables that list all passing models for any given target and detail in captions, under the tables, the references and sources that were rotated from the reference set for that table of *qpAdm*. Note that these tables list only the simplest set of models that pass at the p = 0.05 level. In other words, if some 2-way models pass with p>0.05, we present only these models, even if multiple passing 3-way and 4-way models exist. We will also specify which file contains the corresponding list of all models (both passing and failing) for a particular will set.

We use a special procedure to handle the case where no passing models are found for a target population in both the A and B model sets. In such cases, we move the target population to the tranche that is immediately older and repeat this process until we recover a passing model. The target population is then fixed as a member of the tranche where it received a passing model, and all models in the later tranches are run again with that newly-fixed population in the references, due to its identity as a new population in an older tranche than the target. To sum up, our procedure can be illustrated in the following pseudocode:

```
For currentTrancheNumber in [1, 2, 3, 4, 5]:
    For population in tranche(currentTrancheNumber):
            passing_A_Models[population] = Model_A(population, tranche = currentTrancheNumber)
            passing_B_Models[population] = Model_B(population, tranche = currentTrancheNumber)
                                     If    (passing_A_Models[population]    =    []    and
            passing_B_Models[population] = []):
                    add(tranche(currentTrancheNumber-1), population)
                    remove(tranche(currentTrancheNumber), population)
                    currentTrancheNumber = currentTrancheNumber-1
            continue
```

This procedure accounts for the fact that some populations that fall into younger tranches due to their dating may in fact be descended from a population that also existed earlier in time, but whose earlier periods are not represented in our archaeogenetic record. If such a population is modeled in a younger tranche, its potential genetic connections to the older populations in the references would cause all models to fail. We expect such populations to be modeled successfully only in an older tranche, despite the young ages of its samples in the archaeogenetic record.

To see this, suppose a population, X, is modeled with the following reference populations: R0, R1, R2, R3… RN. Populations R5, R6 … RN are younger than X, and X is in turn younger than R0, R1, R2, R3 and R4. A population genetically similar to X but older than it so it preceded R1, R2, R3 and R4 in time, X', contributed to the ancestry of R1, R2, R3 and R4. All models with X as the target and any one of R1, R2, R3 and R4 in the references are then expected to fail. As we move X up tranches, we eventually exclude R1, R2, R3, and R4, upon which X gets a passing model in a model with a reference set R0, R5, R6, … RN. Now, we model R4, R3, R2 and R1 in downstream analyses with X in the reference set, allowing for the possibility that an older population genetically similar to X, X', contributed to later populations, as is actually the case in reality.

We found one case of a population that had to be moved into an older tranche: *Cisbaikal_LNBA,* which was moved from Tranche 6 into Tranche 3. In the following sections, we present models as they would have appeared as we implemented the pseudocode; i.e., starting with naïve models without populations moved to any other tranche, from oldest to youngest, followed by models for Cisbaikal_LNBA as members of tranches 5, 4, and 3, followed by models for Tranche 4, 5, and 6 with *Cisbaikal_LNBA* moved into Tranche 3 (i.e. moved into the references).

***Dataset and method.*** All models were run using ADMIXTOOLS2[55]. Precalculated $f_2$-statistics, used to speed up the process of *f4*-ratio estimation central to *qpAdm*, were performed allowing for maximal missingness = .99 over a dataset of 192 shotgun-sequenced and hybridization-captured ancient individuals, listed in Data S3 Table 1.

#### VI.A.ii. Naïve Models

##### VI.A.ii.a) Tranche 1: Khaiyrgas_16.7kya

***Khairygas_16.7kya.*** We find, with the populations in tranche 0 in the set of references/sources, that *Khairygas_16.7kya* from ~16,700 BP can be modeled as being derived entirely from a population related to Native Americans, represented by *Peru_Laramate_900BP*, with a passing P-value (p=0.092). A slightly better model is of *Khairygas_16.7kya* is as a 2-way admixture with majority descent (~93%) from a Native-American-related lineage and minor ancestry (~7%) from an *ANE*-related lineage represented by *Russia_AfontovaGora3* (p=0.293). No simpler models pass. For this set of models, *KhatystyrCave_M_10.2kya*/*Yakutia_Mesolithic* was included in the reference set alongside other populations from tranche 0 to account for the possibility that the Mesolithic Southern Yakutian is most closely related to a source of East-Asian ancestry in *Khairygas_16.7kya*. All models, including passing 3- and 4-way models, can be found in Data S3, Table 2. Because there is only one target population in this tranche, there are no models in the B set. As these models are not superseded in later analyses, they contribute to the summary of population relationships depicted in Main Text Fig. 2C.

**Table S6**

| Target | Model | P | Source | Proportion | SE | Z |
|---|---|---|---|---|---|---|
| **Khaiyrgas** | Peru_Laramate_900BP | 0.092 | Peru_Laramate_900BP | 1.000 | 0.000 | 3.13E+13 |
| **Khaiyrgas** | Russia_AfontovaGora3 + Peru_Laramate_900BP | 0.293 | Russia_AfontovaGora3 | 0.065 | 0.026 | 2.47E+00 |
| **Khaiyrgas** | Russia_AfontovaGora3 + Peru_Laramate_900BP | 0.293 | Peru_Laramate_900BP | 0.935 | 0.026 | 3.56E+01 |

**Table S6 |** Passing 2-way model for Khaiyrgas. No other 2-way models pass. **References:** [Tranche 0 + *KhatystyrCave_M_10.2kya*] **Sources:** [*Japan_HG_Jomon, Russia_AfontovaGora3, Peru_Laramate_900BP, China_SEastAsia_Coastal_EN, China_AmurRiver_LPaleolithic, China_AmurRiver_Mesolithic, China_AmurRiver_EarlyN, China_NEastAsia_Coastal_EN, China_NEastAsia_Inland_EN, KhatystyrCave_M_10.2kya*]

We found it intriguing and surprising that *Khairygas_16.7kya* could be modeled as forming a clade with Native Americans, and therefore investigated this further (section VI.A.iv) by switching the position of the Native American population and *Khairgas.SG* in our *qpAdm*s. We found that all passing models for the Native American target as a member of Tranche 1 have them demonstrating complete or near-complete descent from *Khairygas_16.7kya* (Table VI.A-21). *Khairygas_16.7kya* is therefore likely to be a near-unadmixed descendant of a population closely related to the founding population of the Americas, a fact we discuss in further detail in section VI.A.iv. Our *qpAdm* suggest that Khairygas_16.7kya may be a near-unadmixed representative of Ancient Paleosiberian (APS) ancestry, and may have mediated this ancestry to later populations in Siberia.

##### VI.A.ii.b) Tranche 2: Ust_Kyakhta_14kya

***Ust_Kyakhta_14kya.*** We find, using the populations used in the previous set plus *Khairygas_16.7kya* (i.e. tranches 0 + 1) as references/sources, that *Ust_Kyakhta_14kya* can be modeled as a 2-way admixture with majority descent from *Khairygas_16.7kya* or *Peru_Laramate_900BP*, with a minority fraction of East Asian admixture whose source is poorly resolved. *Khaiyrgas* or *Peru_Laramate_900BP* is present in all models at ~53-69%. No simpler models pass. For this set of models, as before, *KhatystyrCave_M_10.2kya*/*Yakutia_Mesolithic* was included in the reference to account for the possibility that the Mesolithic Southern Yakutian is most closely related to the source of East-Asian ancestry in *Ust_Kyakhta_14kya*. All models, including passing 3-way models, can be found in Data S3, Table 3. Because there is only one target population in this tranche, there are no models in the B set.

**Table S7**

| Target | Model | P | Source | Proportion | SE | Z |
|---|---|---|---|---|---|---|
| Ust_Kyakhta_14kya | China_AmurRiver_LPaleolithic_14K + Khaiyrgas | 0.349 | China_AmurRiver_LPaleolithic_14K | 0.371 | 0.031 | 1.19E+01 |
| Ust_Kyakhta_14kya | China_AmurRiver_LPaleolithic_14K + Khaiyrgas | 0.349 | Khaiyrgas | 0.629 | 0.031 | 2.01E+01 |
| Ust_Kyakhta_14kya | Peru_Laramate_900BP + KhatystyrCave_M_10.2kya | 0.258 | Peru_Laramate_900BP | 0.635 | 0.040 | 1.58E+01 |
| Ust_Kyakhta_14kya | Peru_Laramate_900BP + KhatystyrCave_M_10.2kya | 0.258 | KhatystyrCave_M_10.2kya | 0.365 | 0.040 | 9.10E+00 |
| Ust_Kyakhta_14kya | KhatystyrCave_M_10.2kya + Khaiyrgas | 0.095 | KhatystyrCave_M_10.2kya | 0.467 | 0.046 | 1.01E+01 |
| Ust_Kyakhta_14kya | KhatystyrCave_M_10.2kya + Khaiyrgas | 0.095 | Khaiyrgas | 0.533 | 0.046 | 1.16E+01 |
| Ust_Kyakhta_14kya | Peru_Laramate_900BP + China_AmurRiver_LPaleolithic_14K | 0.087 | Peru_Laramate_900BP | 0.693 | 0.030 | 2.32E+01 |
| Ust_Kyakhta_14kya | Peru_Laramate_900BP + China_AmurRiver_LPaleolithic_14K | 0.087 | China_AmurRiver_LPaleolithic_14K | 0.307 | 0.030 | 1.03E+01 |
| Ust_Kyakhta_14kya | Peru_Laramate_900BP + China_NEastAsia_Inland_EN | 0.069 | Peru_Laramate_900BP | 0.636 | 0.034 | 1.86E+01 |
| Ust_Kyakhta_14kya | Peru_Laramate_900BP + China_NEastAsia_Inland_EN | 0.069 | China_NEastAsia_Inland_EN | 0.364 | 0.034 | 1.07E+01 |

**Table S1-VI. A3 |** Passing 2-way models for Ust_Kyakhta_14kya. No simpler models pass. **References:** [Tranche 0 + Tranche 1 + *KhatystyrCave_M_10.2kya/Yakutia_Mesolithic*]
**Sources:** {*Japan_HG_Jomon, Russia_AfontovaGora3, Peru_Laramate_900BP, China_SEastAsia_Coastal_EN, China_AmurRiver_LPaleolithic, China_AmurRiver_Mesolithic, China_AmurRiver_EarlyN, China_NEastAsia_Coastal_EN, China_NEastAsia_Inland_EN, KhatystyrCave_M_10.2kya, Khaiyrgas*]

##### VI.A.ii.c) Tranche 3: KhatystyrCave_M_10.2kya, Kolyma_M_10.1kya

***KhatystyrCave_M_10.2kya/Yakutia_Mesolithic.*** We find, using the populations used in tranche 0 plus *Khaiyrgas* and *Ust_Kyakhta_14kya* (i.e., tranches 0 + 1 + 2), that *KhatystyrCave_M_10.2kya*/*Yakutia_Mesolithic* from Southern Yakutia ~10,000 BP can be modeled as a 2-way admixture of *Khaiyrgas/Ust_Kyakhta_14kya* + *China_AmurRiver_Mesolithic* in set A with a marginally passing p-value (p = 0.056). We therefore display all passing 3-way models: *China_AmurRiver_Mesolithic* is present in all passing models at ~57-78. The second and third ancestry sources in passing 3-way models tend to be a population from further south in East Asia: *China_NEastAsia_Inland_EN* or *China_SEastAsia_Coastal_EN*, or an *ANE*-rich population, usually *Ust_Kyakhta_14kya*. As a population from the Amur River Region from ~10,000 BP, *China_AmurRiver_Mesolithic* is near-contemporaneous with KhatystyrCave_M_10.2kya/*Yakutia_Mesolithic*, and its identity as the major ancestry source in the Mesolithic Southern Yakutian fits well with temporal and geographical considerations. In set B, with co-member of this tranche *Kolyma_M_10.1kya* added to the references/sources, additional models involving *Khairygas_16.7kya* in place of *Ust_Kyakhta_14kya* pass, in keeping with the close relation between *Ust_Kyakhta_14kya* and *Khairygas_16.7kya*.

***Kolyma_M_10.1kya.*** In set A, *Kolyma_M_10.1kya* from the Kolyma River Basin in Northeastern Siberia ~10,000 BP, can be modeled as forming a clade with Ust_Kyakhta_14kya with a marginally passing p-value (p = 0.055). We therefore also present 2-way models, which have Kolyma_M_10.1kya as admixed between either *Khairygas_16.7kya, Ust_Kyakhta_14kya,* or *Peru_Laramate_900BP*, and either one of East Asian ancestry, *AG3*, or ancestry from another APS population. *Peru_Laramate_900BP* or a APS-related source is required in all passing models. In set B when *KhatystyrCave_M_10.2kya* is added to the sources and references, an additional model with *Khaiyrgas* + *KhatystyrCave_M_10.2kya*/*Yakutia_Mesolithic* passes. While models for Kolyma_M_10.1kya are quite varied, they are highly consistent in suggesting that this population is mostly descended from Ancient Paleosiberians, and is less admixed with East Asian ancestry compared to Ust_Kyakhta_14kya (approximately two-thirds to three-fourths of the East Asian admixture in Ust_Kyakhta_14kya).

All models in this tranche, including passing 3- and 4-way models, can be found in Tables 4A (Set A) and 4B (Set B) in Data S3.

**Table S8**

| Target | Model | P | Source | Proportion | SE | Z |
|---|---|---|---|---|---|---|
| KhatystyrCave_M_10.2kya | Ust_Kyakhta_14kya + China_AmurRiver_Mesolithic | 0.056 | Ust_Kyakhta_14kya | 0.325 | 0.064 | 5.10E+00 |
| KhatystyrCave_M_10.2kya | Ust_Kyakhta_14kya + China_AmurRiver_Mesolithic | 0.056 | China_AmurRiver_Mesolithic | 0.675 | 0.064 | 1.06E+01 |
| KhatystyrCave_M_10.2kya | Ust_Kyakhta_14kya + China_AmurRiver_Mesolithic + China_SEastAsia_Coastal_EN | 0.118 | Ust_Kyakhta_14kya | 0.210 | 0.099 | 2.11E+00 |
| KhatystyrCave_M_10.2kya | Ust_Kyakhta_14kya + China_AmurRiver_Mesolithic + China_SEastAsia_Coastal_EN | 0.118 | China_AmurRiver_Mesolithic | 0.613 | 0.077 | 7.93E+00 |
| KhatystyrCave_M_10.2kya | Ust_Kyakhta_14kya + China_AmurRiver_Mesolithic + China_SEastAsia_Coastal_EN | 0.118 | China_SEastAsia_Coastal_EN | 0.177 | 0.119 | 1.49E+00 |
| KhatystyrCave_M_10.2kya | Russia_AfontovaGora3 + China_AmurRiver_Mesolithic + China_SEastAsia_Coastal_EN | 0.093 | Russia_AfontovaGora3 | 0.072 | 0.026 | 2.81E+00 |
| KhatystyrCave_M_10.2kya | Russia_AfontovaGora3 + China_AmurRiver_Mesolithic + China_SEastAsia_Coastal_EN | 0.093 | China_AmurRiver_Mesolithic | 0.748 | 0.086 | 8.72E+00 |
| KhatystyrCave_M_10.2kya | Russia_AfontovaGora3 + China_AmurRiver_Mesolithic + China_SEastAsia_Coastal_EN | 0.093 | China_SEastAsia_Coastal_EN | 0.180 | 0.100 | 1.80E+00 |
| KhatystyrCave_M_10.2kya | Russia_AfontovaGora3 + Ust_Kyakhta_14kya + China_AmurRiver_Mesolithic | 0.079 | Russia_AfontovaGora3 | 0.023 | 0.064 | 3.61E-01 |
| KhatystyrCave_M_10.2kya | Russia_AfontovaGora3 + Ust_Kyakhta_14kya + China_AmurRiver_Mesolithic | 0.079 | Ust_Kyakhta_14kya | 0.280 | 0.212 | 1.32E+00 |
| KhatystyrCave_M_10.2kya | Russia_AfontovaGora3 + Ust_Kyakhta_14kya + China_AmurRiver_Mesolithic | 0.079 | China_AmurRiver_Mesolithic | 0.697 | 0.152 | 4.58E+00 |

| | | | | | | |
|---|---|---|---|---|---|---|
| KhatystyrCave _M_10.2kya | Ust_Kyakhta_14kya + China_AmurRiver_Mesolithic + China_NEastAsia_Inland_EN | 0.057 | Ust_Kyakhta_14kya | 0.256 | 0.131 | 1.95E+00 |
| KhatystyrCave _M_10.2kya | Ust_Kyakhta_14kya + China_AmurRiver_Mesolithic + China_NEastAsia_Inland_EN | 0.057 | China_AmurRiver_Mesolithic | 0.571 | 0.170 | 3.35E+00 |
| KhatystyrCave _M_10.2kya | Ust_Kyakhta_14kya + China_AmurRiver_Mesolithic + China_NEastAsia_Inland_EN | 0.057 | China_NEastAsia_Inland_EN | 0.173 | 0.272 | 6.35E-01 |
| Kolyma_M_10 .1kya | Ust_Kyakhta_14kya | 0.055 | Ust_Kyakhta_14kya | 1.000 | 0.000 | 3.13E+13 |
| Kolyma_M_10 .1kya | Khaiyrgas + China_AmurRiver_LPaleolithic_14K | 0.357 | Khaiyrgas | 0.739 | 0.032 | 2.33E+01 |
| Kolyma_M_10 .1kya | Khaiyrgas + China_AmurRiver_LPaleolithic_14K | 0.357 | China_AmurRiver_LPaleolithic _14K | 0.261 | 0.032 | 8.21E+00 |
| Kolyma_M_10 .1kya | Russia_AfontovaGora3 + Ust_Kyakhta_14kya | 0.308 | Russia_AfontovaGora3 | 0.054 | 0.019 | 2.86E+00 |
| Kolyma_M_10 .1kya | Russia_AfontovaGora3 + Ust_Kyakhta_14kya | 0.308 | Ust_Kyakhta_14kya | 0.946 | 0.019 | 5.03E+01 |
| Kolyma_M_10 .1kya | Khaiyrgas + China_AmurRiver_Mesolithic | 0.248 | Khaiyrgas | 0.735 | 0.031 | 2.39E+01 |
| Kolyma_M_10 .1kya | Khaiyrgas + China_AmurRiver_Mesolithic | 0.248 | China_AmurRiver_Mesolithic | 0.265 | 0.031 | 8.61E+00 |
| Kolyma_M_10 .1kya | Khaiyrgas + Ust_Kyakhta_14kya | 0.225 | Khaiyrgas | 0.213 | 0.096 | 2.22E+00 |
| Kolyma_M_10 .1kya | Khaiyrgas + Ust_Kyakhta_14kya | 0.225 | Ust_Kyakhta_14kya | 0.787 | 0.096 | 8.21E+00 |
| Kolyma_M_10 .1kya | Peru_Laramate_900BP + Ust_Kyakhta_14kya | 0.217 | Peru_Laramate_900BP | 0.312 | 0.125 | 2.50E+00 |
| Kolyma_M_10 .1kya | Peru_Laramate_900BP + Ust_Kyakhta_14kya | 0.217 | Ust_Kyakhta_14kya | 0.688 | 0.125 | 5.52E+00 |
| Kolyma_M_10 .1kya | Khaiyrgas + China_NEastAsia_Inland_EN | 0.157 | Khaiyrgas | 0.655 | 0.041 | 1.59E+01 |
| Kolyma_M_10 .1kya | Khaiyrgas + China_NEastAsia_Inland_EN | 0.157 | China_NEastAsia_Inland_EN | 0.345 | 0.041 | 8.39E+00 |
| Kolyma_M_10 .1kya | Peru_Laramate_900BP + China_NEastAsia_Inland_EN | 0.157 | Peru_Laramate_900BP | 0.757 | 0.041 | 1.84E+01 |
| Kolyma_M_10 .1kya | Peru_Laramate_900BP + China_NEastAsia_Inland_EN | 0.157 | China_NEastAsia_Inland_EN | 0.243 | 0.041 | 5.90E+00 |

#### Table S9

| Target | Model | P | Source | Proportion | SE | Z |
|---|---|---|---|---|---|---|
| KhatystyrCave _M_10.2kya | Ust_Kyakhta_14kya + China_AmurRiver_Mesolithic | 0.068 | Ust_Kyakhta_14kya | 0.327 | 0.064 | 5.13E+00 |
| KhatystyrCave _M_10.2kya | Ust_Kyakhta_14kya + China_AmurRiver_Mesolithic | 0.068 | China_AmurRiver_Mesolithic | 0.673 | 0.064 | 1.06E+01 |
| KhatystyrCave _M_10.2kya | Ust_Kyakhta_14kya + China_AmurRiver_Mesolithic + China_SEastAsia_Coastal_EN | 0.103 | Ust_Kyakhta_14kya | 0.246 | 0.095 | 2.59E+00 |
| KhatystyrCave _M_10.2kya | Ust_Kyakhta_14kya + China_AmurRiver_Mesolithic + China_SEastAsia_Coastal_EN | 0.103 | China_AmurRiver_Mesolithic | 0.630 | 0.076 | 8.28E+00 |
| KhatystyrCave _M_10.2kya | Ust_Kyakhta_14kya + China_AmurRiver_Mesolithic + China_SEastAsia_Coastal_EN | 0.103 | China_SEastAsia_Coastal_EN | 0.124 | 0.112 | 1.10E+00 |
| KhatystyrCave _M_10.2kya | Ust_Kyakhta_14kya + China_AmurRiver_Mesolithic + China_NEastAsia_Inland_EN | 0.066 | Ust_Kyakhta_14kya | 0.275 | 0.126 | 2.18E+00 |
| KhatystyrCave _M_10.2kya | Ust_Kyakhta_14kya + China_AmurRiver_Mesolithic + China_NEastAsia_Inland_EN | 0.066 | China_AmurRiver_Mesolithic | 0.591 | 0.168 | 3.52E+00 |
| KhatystyrCave _M_10.2kya | Ust_Kyakhta_14kya + China_AmurRiver_Mesolithic + China_NEastAsia_Inland_EN | 0.066 | China_NEastAsia_Inland_EN | 0.134 | 0.264 | 5.08E-01 |
| KhatystyrCave _M_10.2kya | Khaiyrgas + China_AmurRiver_Mesolithic + China_NEastAsia_Inland_EN | 0.051 | Khaiyrgas | 0.152 | 0.077 | 1.98E+00 |

| | | | | | | |
|---|---|---|---|---|---|---|
| KhatystyrCave _M_10.2kya | Khaiyrgas + China_AmurRiver_Mesolithic + China_NEastAsia_Inland_EN | 0.051 | China_AmurRiver_Mesolithic | 0.583 | 0.189 | 3.09E+00 |
| KhatystyrCave _M_10.2kya | Khaiyrgas + China_AmurRiver_Mesolithic + China_NEastAsia_Inland_EN | 0.051 | China_NEastAsia_Inland_EN | 0.265 | 0.248 | 1.07E+00 |
| KhatystyrCave _M_10.2kya | Khaiyrgas + China_AmurRiver_Mesolithic + China_SEastAsia_Coastal_EN | 0.050 | Khaiyrgas | 0.190 | 0.062 | 3.05E+00 |
| KhatystyrCave _M_10.2kya | Khaiyrgas + China_AmurRiver_Mesolithic + China_SEastAsia_Coastal_EN | 0.050 | China_AmurRiver_Mesolithic | 0.740 | 0.078 | 9.46E+00 |
| KhatystyrCave _M_10.2kya | Khaiyrgas + China_AmurRiver_Mesolithic + China_SEastAsia_Coastal_EN | 0.050 | China_SEastAsia_Coastal_EN | 0.069 | 0.112 | 6.20E-01 |
| Kolyma_M_10 .1kya | Ust_Kyakhta_14kya | 0.078 | Ust_Kyakhta_14kya | 1.000 | 0.000 | 3.13E+13 |
| Kolyma_M_10 .1kya | Khaiyrgas + China_AmurRiver_LPaleolithic_14K | 0.306 | Khaiyrgas | 0.738 | 0.032 | 2.33E+01 |
| Kolyma_M_10 .1kya | Khaiyrgas + China_AmurRiver_LPaleolithic_14K | 0.306 | China_AmurRiver_LPaleolithic_14K | 0.262 | 0.032 | 8.27E+00 |
| Kolyma_M_10 .1kya | Khaiyrgas + Ust_Kyakhta_14kya | 0.292 | Khaiyrgas | 0.211 | 0.094 | 2.24E+00 |
| Kolyma_M_10 .1kya | Khaiyrgas + Ust_Kyakhta_14kya | 0.292 | Ust_Kyakhta_14kya | 0.789 | 0.094 | 8.40E+00 |
| Kolyma_M_10 .1kya | Peru_Laramate_900BP + Ust_Kyakhta_14kya | 0.282 | Peru_Laramate_900BP | 0.311 | 0.125 | 2.50E+00 |
| Kolyma_M_10 .1kya | Peru_Laramate_900BP + Ust_Kyakhta_14kya | 0.282 | Ust_Kyakhta_14kya | 0.689 | 0.125 | 5.52E+00 |
| Kolyma_M_10 .1kya | Khaiyrgas + China_AmurRiver_Mesolithic | 0.275 | Khaiyrgas | 0.733 | 0.031 | 2.39E+01 |
| Kolyma_M_10 .1kya | Khaiyrgas + China_AmurRiver_Mesolithic | 0.275 | China_AmurRiver_Mesolithic | 0.267 | 0.031 | 8.71E+00 |
| Kolyma_M_10 .1kya | Peru_Laramate_900BP + China_NEastAsia_Inland_EN | 0.186 | Peru_Laramate_900BP | 0.762 | 0.041 | 1.87E+01 |
| Kolyma_M_10 .1kya | Peru_Laramate_900BP + China_NEastAsia_Inland_EN | 0.186 | China_NEastAsia_Inland_EN | 0.238 | 0.041 | 5.85E+00 |
| Kolyma_M_10 .1kya | Khaiyrgas + KhatystyrCave_M_10.2kya | 0.107 | Khaiyrgas | 0.702 | 0.038 | 1.86E+01 |
| Kolyma_M_10 .1kya | Khaiyrgas + KhatystyrCave_M_10.2kya | 0.107 | KhatystyrCave_M_10.2kya | 0.298 | 0.038 | 7.90E+00 |
| Kolyma_M_10 .1kya | Khaiyrgas + China_NEastAsia_Inland_EN | 0.104 | Khaiyrgas | 0.657 | 0.041 | 1.60E+01 |
| Kolyma_M_10 .1kya | Khaiyrgas + China_NEastAsia_Inland_EN | 0.104 | China_NEastAsia_Inland_EN | 0.343 | 0.041 | 8.32E+00 |

**Table S1-VI. A-4 & 5 |** Passing 2-way models for KhatystyrCave_M_10.2kya/*Yakutia_Mesolithic* & *Kolyma_M_10.1kya*. No simpler models pass. **References:** [Tranche 0 + Tranche 1 + Tranche 2 + *KhatystyrCave_M_10.2kya/Yakutia_Mesolithic* (in Set B) + *Kolyma_M_10.1kya* (in Set B)]

**Sources:** [*Peru_Laramate_900BP, Russia_AfontovaGora3, Khaiyrgas, Ust_Kyakhta_14kya, China_AmurRiver_LPaleolithic_19K, China_AmurRiver_LPaleolithic_14K, China_AmurRiver_Mesolithic, China_NEastAsia_Coastal_EN, China_NEastAsia_Inland_EN, China_SEastAsia_Coastal_EN, KhatystyrCave_M_10.2kya* (in set B), *Kolyma_M_10.1kya* (in set B)]

##### VI.A.ii.d) Tranche 4: Dzhilinda1_M_N_8.4kya, Transbaikal_EMN_old, Altai_N_old

***Dzhilinda1_M_N_8.4kya.*** We find, using the populations in tranche 0, 1, 2 and 3 as references, that the *Dzhilinda1_M_N_8.4kya* individual from Northern Buryatia at ~8,400 BP, can be modeled as having all its ancestry coming from *Ust_Kyakhta_14kya*, but that this model fails with Transbaikal_EMN placed in the references in set B. There, three models pass with *Dzhilinda1_M_N_8.4kya* as an admixture of *Kolyma_M_10.1kya* or *Khaiyrgas* and an East-Asian-related source with much ancestry from *KhatystyrCave_M_10.2kya KhatystyrCave_M_10.2kya* or *Transbaikal_EMN_old*.

***Transbaikal_EMN_old.*** *Transbaikal_EMN_old* is a combination of the individuals irk007.SG and cta016.SG, who are from the *Transbaikal_EMN* cluster defined by the grouping process in section V of this document, but who are notably older than the rest, at ~8,300 & ~8,800 BP, where the rest of the individuals span the interval from ~7,400 to ~6,100 BP. Only one passing model was found for *Transbaikal_EMN_old* as admixed between the Yumin individual (*China_NEastAsia_Inland_EN*) and *KhatystyrCave_M_10.2kya* was found using the outgroup set for this tranche for both set A and set B.

***Altai_N_old.*** The oldest individuals under the *Altai_N* group label (uniting *Kemerovo_N*, *Firsovo_N*, and *West_Siberia_N*—extremely similar populations found in the Altai region) older than ~8000 BP (including individuals I2137 and I13098), can be successfully modeled as a 2-way admixture between a minor fraction of ancestry from *AG3* and a major fraction of ancestry from the three APS populations *Khaiyrgas*, *Ust_Kyakhta_14kya, Kolyma_M_10.1kya*. The same models pass in set B; no simpler models pass.

All models in this tranche can be found in Tables 5A (Set A) and 5B (Set B) in Data S3.

**Table S10**

| Target | Model | P | Source | Prop. | SE | Z |
|---|---|---|---|---|---|---|
| Dzhilinda1_M_N_8.4kya | Ust_Kyakhta_14kya | 0.096 | Ust_Kyakhta_14kya | 1.000 | 0.000 | 3.13E+13 |
| Transbaikal_EMN_old | China_NEastAsia_Inland_EN + KhatystyrCave_M_10.2kya | 0.215 | China_NEastAsia_Inland_EN | 0.566 | 0.138 | 4.11E+00 |
| Transbaikal_EMN_old | China_NEastAsia_Inland_EN + KhatystyrCave_M_10.2kya | 0.215 | KhatystyrCave_M_10.2kya | 0.434 | 0.138 | 3.15E+00 |
| Altai_N_old | Russia_AfontovaGora3 + Ust_Kyakhta_14kya | 0.574 | Russia_AfontovaGora3 | 0.377 | 0.016 | 2.38E+01 |
| Altai_N_old | Russia_AfontovaGora3 + Ust_Kyakhta_14kya | 0.574 | Ust_Kyakhta_14kya | 0.623 | 0.016 | 3.94E+01 |
| Altai_N_old | Russia_AfontovaGora3 + Kolyma_M_10.1kya | 0.296 | Russia_AfontovaGora3 | 0.337 | 0.018 | 1.89E+01 |
| Altai_N_old | Russia_AfontovaGora3 + Kolyma_M_10.1kya | 0.296 | Kolyma_M_10.1kya | 0.663 | 0.018 | 3.70E+01 |
| Altai_N_old | Russia_AfontovaGora3 + Khaiyrgas | 0.130 | Russia_AfontovaGora3 | 0.195 | 0.026 | 7.41E+00 |
| Altai_N_old | Russia_AfontovaGora3 + Khaiyrgas | 0.130 | Khaiyrgas | 0.805 | 0.026 | 3.07E+01 |

**Table S11**

| Target | Model | P | Source | Prop. | SE | Z |
|---|---|---|---|---|---|---|
| Dzhilinda1_M_N_8.4kya | Kolyma_M_10.1kya + Transbaikal_EMN_old | 0.252 | Kolyma_M_10.1kya | 0.834 | 0.054 | 1.55E+01 |
| Dzhilinda1_M_N_8.4kya | Kolyma_M_10.1kya + Transbaikal_EMN_old | 0.252 | Transbaikal_EMN_old | 0.166 | 0.054 | 3.08E+00 |
| Dzhilinda1_M_N_8.4kya | Khaiyrgas + KhatystyrCave_M_10.2kya | 0.137 | Khaiyrgas | 0.584 | 0.040 | 1.45E+01 |
| Dzhilinda1_M_N_8.4kya | Khaiyrgas + KhatystyrCave_M_10.2kya | 0.137 | KhatystyrCave_M_10.2kya | 0.416 | 0.040 | 1.03E+01 |
| Dzhilinda1_M_N_8.4kya | KhatystyrCave_M_10.2kya + Kolyma_M_10.1kya | 0.113 | KhatystyrCave_M_10.2kya | 0.183 | 0.053 | 3.45E+00 |
| Dzhilinda1_M_N_8.4kya | KhatystyrCave_M_10.2kya + Kolyma_M_10.1kya | 0.113 | Kolyma_M_10.1kya | 0.817 | 0.053 | 1.53E+01 |
| Transbaikal_EMN_old | China_NEastAsia_Inland_EN + KhatystyrCave_M_10.2kya | 0.190 | China_NEastAsia_Inland_EN | 0.482 | 0.129 | 3.74E+00 |
| Transbaikal_EMN_old | China_NEastAsia_Inland_EN + KhatystyrCave_M_10.2kya | 0.190 | KhatystyrCave_M_10.2kya | 0.518 | 0.129 | 4.02E+00 |
| Altai_N_old | Russia_AfontovaGora3 + Ust_Kyakhta_14kya | 0.562 | Russia_AfontovaGora3 | 0.375 | 0.016 | 2.41E+01 |
| Altai_N_old | Russia_AfontovaGora3 + Ust_Kyakhta_14kya | 0.562 | Ust_Kyakhta_14kya | 0.625 | 0.016 | 4.01E+01 |
| Altai_N_old | Russia_AfontovaGora3 + Kolyma_M_10.1kya | 0.302 | Russia_AfontovaGora3 | 0.336 | 0.018 | 1.90E+01 |
| Altai_N_old | Russia_AfontovaGora3 + Kolyma_M_10.1kya | 0.302 | Kolyma_M_10.1kya | 0.664 | 0.018 | 3.76E+01 |
| Altai_N_old | Russia_AfontovaGora3 + Khaiyrgas | 0.206 | Russia_AfontovaGora3 | 0.193 | 0.026 | 7.56E+00 |
| Altai_N_old | Russia_AfontovaGora3 + Khaiyrgas | 0.206 | Khaiyrgas | 0.807 | 0.026 | 3.15E+01 |

**Table S1-VI. A-6 & 7 |** Passing 2-way models for *Dzhilinda1_M_N_8.4kya*. No passing models were found for either A or B for *Transbaikal_EMN_old*; the listed models fail because of a small nonsignificant negative contribution from *Ust_Kyakhta_14kya*. **References:** [Tranche 0 + Tranche 1 + Tranche 2 + Tranche 4]

**Sources:** [*Peru_Laramate_900BP*, *AG3*, *Khairygas_16.7kya, Ust_Kyakhta_14kya, Kolyma.SG, China_AmurRiver_Paleolithic_19K, China_AmurRiver_LPaleolithic_14K, China_AmurRiver_Mesolithic, China_NEastAsia_Coastal_EN, China_NEastAsia_Inland_EN, China_SEastAsia_Coastal_EN, KhatystyrCave_M_10.2kya, (irk001.SG, Alta_N_old, Transbaikal_EMN_old* included in Set B)]

#### VI.A.ii.e) Tranche 5: Altai_N, Transbaikal_EMN, China_AmurRiver_N, Cisbaikal_EN, Mongolia_N_North, Syalakh-Belkachi

***Altai_N.*** This group, which comprises all individuals in the *Altai_N* group label (all individuals labeled *Kemerovo_N*, *Firsovo_N*, and *West_Siberia_N*) younger than 8000 BP, can be modeled as descending completely from the older individuals in this cluster (*Altai_N_old*). The same model passes in set B.

***Transbaikal_EMN.*** In set A, *Transbaikal_EMN,* consisting of younger individuals in the *Transbaikal_EMN* cluster defined in section V found in the Shilka river catchment between the Baikal and the Amur River that are between ~8000-6000 years BP, can be modeled as descending completely in a 1-way model from *Transbaikal_EMN_old*, even with an extensive outgroup set combining tranches 0, 1, 2, 3 and 4. The same 1-way model passes in set B.

***China_AmurRiver_N.*** This population, consisting of individuals from the Amur River Basin region younger than ~8000 BP, can be modeled as descending almost entirely (~77-98%) from prior populations of the Amur River region (> 8,000 years BP) in multiple 2-way models. Most passing 2-way models feature near complete descent from *China_AmurRiver_EarlyN* or *China_AmurRiver_Mesolithic* with the addition of a small contribution from one other, typically Northeast Asian or APS source, testifying to genetic continuity but with low levels of gene flow in the Amur River region. A 1-way model of *China_AmurRiver_N* as 100% *China_AmurRiver_EarlyN* fails with weak statistical significance (p-value = 0.019). A similar set of 2-way models with near-complete descent from *China_AmurRiver_EarlyN/China_AmurRiver_Mesolithic* passes in set B as well.

***Cisbaikal_EN.*** In set A, models of *Cisbaikal_EN*, which includes individuals from the Cisbaikal region from ~8000BP to ~6800BP, with major contributions from *Transbaikal_EMN_old* (~65%-79%), and minor contributions from an APS population (*Ust_Kyakhta_14kya* ~35% or *Khairygas_16.7kya* ~31%) pass marginally (0.05 < p < 0.069). We therefore also list passing 3-way models with higher p-values, which appear to require very small additional contributions from populations from further south in East Asia proper, or from *AG3*, at levels within the standard error. No models pass for Cisbaikal_EN in set B.

***Mongolia_N_North.*** A two-way model for *Mongolia_N_North*, from the northern region of the Mongolian plateau just south of Lake Baikal approximately 7500BP, as an admixture between *Transbaikal_EMN_old* (~65%) + *China_NEastAsia_Inland_EN* (~35%) passes marginally (p = 0.079) in set A. We therefore also present 3-way models with higher p-values, all of which retain *Transbaikal_EMN_old* and *China_NEastAsia_Inland_EN* in nearly the same proportions as in the 2-way model, with the addition of a small fraction (<7%) of ancestry from an *ANE*-rich source (either *Altai_N_old* or from a APS-related population such as *Khairygas_16.7kya, Ust_Kyakhta_14kya*, *Dzhilinda1_M_N_8.4kya* or *Kolyma_M_10.1kya*). No models pass in set B.

***Syalakh-Belkachi.*** In Set A, the *Syalakh-Belkachi* population in the Middle Lena river valley in Central Yakutia from 6800-6300 BP can be modeled as descending entirely from the Mesolithic Northern Buryatian *Dzhilinda1_M_N_8.4kya*, with a marginally passing p-value (p = 0.066). We also list passing 2-way models with higher p-values, where *Syalakh-Belkachi* continues to demonstrate majority descent from *Dzhilinda1_M_N_8.4kya* with a minor contribution (~10-20%) from a broadly East-Asian-related population whose identity is poorly resolved. In set B, this is reduced to three passing models, with a major contribution from *Dzhilinda1_M_N_8.4kya* (~76%) and a minor contribution from a population rich in Inland Northeast Asian ancestry (represented by the Yumin individual under the group label *China_NEastAsia_Inland_EN*), either *Cisbaikal_EN*, *Transbaikal_EMN_old* or *Mongolia_N_North* (~24%).

All models in this set are found in Table 6A (Set A) or Table 6B (Set B) in Data S3.

**Table S12**

| Target | Model | P | Source | Prop. | SE | Z |
|---|---|---|---|---|---|---|
| Altai_N | Altai_N_old | 0.690 | Altai_N_old | 1.000 | 0.000 | 3.13E+13 |
| Transbaikal_EMN | Transbaikal_EMN_old | 0.540 | Transbaikal_EMN_old | 1.000 | 0.000 | 3.13E+13 |
| China_AmurRiver_N | China_AmurRiver_EarlyN + Transbaikal_EMN_old | 0.528 | China_AmurRiver_EarlyN | 0.817 | 0.046 | 1.78E+01 |
| China_AmurRiver_N | China_AmurRiver_EarlyN + Transbaikal_EMN_old | 0.528 | Transbaikal_EMN_old | 0.183 | 0.046 | 3.99E+00 |
| China_AmurRiver_N | China_AmurRiver_EarlyN + China_NEastAsia_Inland_EN | 0.257 | China_AmurRiver_EarlyN | 0.877 | 0.043 | 2.03E+01 |
| China_AmurRiver_N | China_AmurRiver_EarlyN + China_NEastAsia_Inland_EN | 0.257 | China_NEastAsia_Inland_EN | 0.123 | 0.043 | 2.83E+00 |
| China_AmurRiver_N | China_AmurRiver_EarlyN + Dzhilinda1_M_N_8.4kya | 0.227 | China_AmurRiver_EarlyN | 0.942 | 0.020 | 4.62E+01 |
| China_AmurRiver_N | China_AmurRiver_EarlyN + Dzhilinda1_M_N_8.4kya | 0.227 | Dzhilinda1_M_N_8.4kya | 0.058 | 0.020 | 2.87E+00 |
| China_AmurRiver_N | China_AmurRiver_EarlyN + China_NEastAsia_Coastal_EN | 0.128 | China_AmurRiver_EarlyN | 0.895 | 0.049 | 1.81E+01 |
| China_AmurRiver_N | China_AmurRiver_EarlyN + China_NEastAsia_Coastal_EN | 0.128 | China_NEastAsia_Coastal_EN | 0.105 | 0.049 | 2.13E+00 |
| China_AmurRiver_N | China_AmurRiver_Mesolithic + Transbaikal_EMN_old | 0.122 | China_AmurRiver_Mesolithic | 0.775 | 0.041 | 1.88E+01 |
| China_AmurRiver_N | China_AmurRiver_Mesolithic + Transbaikal_EMN_old | 0.122 | Transbaikal_EMN_old | 0.225 | 0.041 | 5.45E+00 |
| China_AmurRiver_N | China_AmurRiver_EarlyN + KhatystyrCave_M_10.2kya | 0.088 | China_AmurRiver_EarlyN | 0.875 | 0.067 | 1.31E+01 |
| China_AmurRiver_N | China_AmurRiver_EarlyN + KhatystyrCave_M_10.2kya | 0.088 | KhatystyrCave_M_10.2kya | 0.125 | 0.067 | 1.87E+00 |
| China_AmurRiver_N | Ust_Kyakhta_14kya + China_AmurRiver_EarlyN | 0.082 | Ust_Kyakhta_14kya | 0.058 | 0.021 | 2.73E+00 |
| China_AmurRiver_N | Ust_Kyakhta_14kya + China_AmurRiver_EarlyN | 0.082 | China_AmurRiver_EarlyN | 0.942 | 0.021 | 4.43E+01 |
| China_AmurRiver_N | Khaiyrgas + China_AmurRiver_EarlyN | 0.072 | Khaiyrgas | 0.034 | 0.014 | 2.45E+00 |
| China_AmurRiver_N | Khaiyrgas + China_AmurRiver_EarlyN | 0.072 | China_AmurRiver_EarlyN | 0.966 | 0.014 | 6.95E+01 |
| China_AmurRiver_N | China_AmurRiver_EarlyN + Kolyma_M_10.1kya | 0.070 | China_AmurRiver_EarlyN | 0.954 | 0.019 | 5.09E+01 |
| China_AmurRiver_N | China_AmurRiver_EarlyN + Kolyma_M_10.1kya | 0.070 | Kolyma_M_10.1kya | 0.046 | 0.019 | 2.45E+00 |
| China_AmurRiver_N | China_AmurRiver_Mesolithic + China_NEastAsia_Inland_EN | 0.065 | China_AmurRiver_Mesolithic | 0.830 | 0.043 | 1.91E+01 |
| China_AmurRiver_N | China_AmurRiver_Mesolithic + China_NEastAsia_Inland_EN | 0.065 | China_NEastAsia_Inland_EN | 0.170 | 0.043 | 3.92E+00 |
| China_AmurRiver_N | Altai_N_old + China_AmurRiver_EarlyN | 0.059 | Altai_N_old | 0.022 | 0.012 | 1.93E+00 |
| China_AmurRiver_N | Altai_N_old + China_AmurRiver_EarlyN | 0.059 | China_AmurRiver_EarlyN | 0.978 | 0.012 | 8.44E+01 |
| Cisbaikal_EN | Ust_Kyakhta_14kya + Transbaikal_EMN_old | 0.069 | Ust_Kyakhta_14kya | 0.342 | 0.023 | 1.47E+01 |
| Cisbaikal_EN | Ust_Kyakhta_14kya + Transbaikal_EMN_old | 0.069 | Transbaikal_EMN_old | 0.658 | 0.023 | 2.83E+01 |
| Cisbaikal_EN | Khaiyrgas + Transbaikal_EMN_old | 0.051 | Khaiyrgas | 0.205 | 0.016 | 1.27E+01 |
| Cisbaikal_EN | Khaiyrgas + Transbaikal_EMN_old | 0.051 | Transbaikal_EMN_old | 0.795 | 0.016 | 4.95E+01 |
| Cisbaikal_EN | Ust_Kyakhta_14kya + China_SEastAsia_Coastal_EN + Transbaikal_EMN_old | 0.215 | Ust_Kyakhta_14kya | 0.335 | 0.026 | 1.27E+01 |
| Cisbaikal_EN | Ust_Kyakhta_14kya + China_SEastAsia_Coastal_EN + Transbaikal_EMN_old | 0.215 | China_SEastAsia_Coastal_EN | 0.023 | 0.028 | 8.01E-01 |

| | | | | | | |
|---|---|---|---|---|---|---|
| Cisbaikal_EN | Ust_Kyakhta_14kya + China_SEastAsia_Coastal_EN + Transbaikal_EMN_old | 0.215 | Transbaikal_EMN_old | 0.643 | 0.027 | 2.35E+01 |
| Cisbaikal_EN | Russia_AfontovaGora3 + Ust_Kyakhta_14kya + Transbaikal_EMN_old | 0.113 | Russia_AfontovaGora3 | 0.031 | 0.023 | 1.37E+00 |
| Cisbaikal_EN | Russia_AfontovaGora3 + Ust_Kyakhta_14kya + Transbaikal_EMN_old | 0.113 | Ust_Kyakhta_14kya | 0.212 | 0.097 | 2.19E+00 |
| Cisbaikal_EN | Russia_AfontovaGora3 + Ust_Kyakhta_14kya + Transbaikal_EMN_old | 0.113 | Transbaikal_EMN_old | 0.756 | 0.075 | 1.01E+01 |
| Cisbaikal_EN | Ust_Kyakhta_14kya + China_NEastAsia_Inland_EN + Transbaikal_EMN_old | 0.104 | Ust_Kyakhta_14kya | 0.334 | 0.025 | 1.36E+01 |
| Cisbaikal_EN | Ust_Kyakhta_14kya + China_NEastAsia_Inland_EN + Transbaikal_EMN_old | 0.104 | China_NEastAsia_Inland_EN | 0.105 | 0.078 | 1.35E+00 |
| Cisbaikal_EN | Ust_Kyakhta_14kya + China_NEastAsia_Inland_EN + Transbaikal_EMN_old | 0.104 | Transbaikal_EMN_old | 0.560 | 0.072 | 7.79E+00 |
| Cisbaikal_EN | Khaiyrgas + Ust_Kyakhta_14kya + Transbaikal_EMN_old | 0.084 | Khaiyrgas | 0.152 | 0.213 | 7.13E-01 |
| Cisbaikal_EN | Khaiyrgas + Ust_Kyakhta_14kya + Transbaikal_EMN_old | 0.084 | Ust_Kyakhta_14kya | 0.090 | 0.361 | 2.48E-01 |
| Cisbaikal_EN | Khaiyrgas + Ust_Kyakhta_14kya + Transbaikal_EMN_old | 0.084 | Transbaikal_EMN_old | 0.758 | 0.149 | 5.09E+00 |
| Cisbaikal_EN | Altai_N_old + Ust_Kyakhta_14kya + Transbaikal_EMN_old | 0.077 | Altai_N_old | 0.096 | 0.093 | 1.03E+00 |
| Cisbaikal_EN | Altai_N_old + Ust_Kyakhta_14kya + Transbaikal_EMN_old | 0.077 | Ust_Kyakhta_14kya | 0.140 | 0.206 | 6.79E-01 |
| Cisbaikal_EN | Altai_N_old + Ust_Kyakhta_14kya + Transbaikal_EMN_old | 0.077 | Transbaikal_EMN_old | 0.764 | 0.114 | 6.73E+00 |
| *Mongolia_N_North* | China_NEastAsia_Inland_EN + Transbaikal_EMN_old | 0.079 | China_NEastAsia_Inland_EN | 0.348 | 0.110 | 3.18E+00 |
| *Mongolia_N_North* | China_NEastAsia_Inland_EN + Transbaikal_EMN_old | 0.079 | Transbaikal_EMN_old | 0.652 | 0.110 | 5.95E+00 |
| *Mongolia_N_North* | Altai_N_old + China_NEastAsia_Inland_EN + Transbaikal_EMN_old | 0.505 | Altai_N_old | 0.017 | 0.013 | 1.26E+00 |
| *Mongolia_N_North* | Altai_N_old + China_NEastAsia_Inland_EN + Transbaikal_EMN_old | 0.505 | China_NEastAsia_Inland_EN | 0.377 | 0.099 | 3.80E+00 |
| *Mongolia_N_North* | Altai_N_old + China_NEastAsia_Inland_EN + Transbaikal_EMN_old | 0.505 | Transbaikal_EMN_old | 0.606 | 0.095 | 6.39E+00 |
| *Mongolia_N_North* | Ust_Kyakhta_14kya + China_NEastAsia_Inland_EN + Transbaikal_EMN_old | 0.290 | Ust_Kyakhta_14kya | 0.063 | 0.028 | 2.21E+00 |
| *Mongolia_N_North* | Ust_Kyakhta_14kya + China_NEastAsia_Inland_EN + Transbaikal_EMN_old | 0.290 | China_NEastAsia_Inland_EN | 0.304 | 0.106 | 2.88E+00 |
| *Mongolia_N_North* | Ust_Kyakhta_14kya + China_NEastAsia_Inland_EN + Transbaikal_EMN_old | 0.290 | Transbaikal_EMN_old | 0.633 | 0.099 | 6.43E+00 |
| *Mongolia_N_North* | Khaiyrgas + China_NEastAsia_Inland_EN + Transbaikal_EMN_old | 0.192 | Khaiyrgas | 0.032 | 0.018 | 1.79E+00 |
| *Mongolia_N_North* | Khaiyrgas + China_NEastAsia_Inland_EN + Transbaikal_EMN_old | 0.192 | China_NEastAsia_Inland_EN | 0.302 | 0.105 | 2.87E+00 |
| *Mongolia_N_North* | Khaiyrgas + China_NEastAsia_Inland_EN + Transbaikal_EMN_old | 0.192 | Transbaikal_EMN_old | 0.666 | 0.099 | 6.73E+00 |
| *Mongolia_N_North* | China_NEastAsia_Inland_EN + Kolyma_M_10.1kya + Transbaikal_EMN_old | 0.164 | China_NEastAsia_Inland_EN | 0.281 | 0.105 | 2.66E+00 |
| *Mongolia_N_North* | China_NEastAsia_Inland_EN + Kolyma_M_10.1kya + Transbaikal_EMN_old | 0.164 | Kolyma_M_10.1kya | 0.044 | 0.024 | 1.79E+00 |
| *Mongolia_N_North* | China_NEastAsia_Inland_EN + Kolyma_M_10.1kya + Transbaikal_EMN_old | 0.164 | Transbaikal_EMN_old | 0.676 | 0.100 | 6.78E+00 |
| *Mongolia_N_North* | China_NEastAsia_Inland_EN + Transbaikal_EMN_old + Dzhilinda1_M_N_8.4kya | 0.130 | China_NEastAsia_Inland_EN | 0.274 | 0.127 | 2.16E+00 |
| *Mongolia_N_North* | China_NEastAsia_Inland_EN + Transbaikal_EMN_old + Dzhilinda1_M_N_8.4kya | 0.130 | Transbaikal_EMN_old | 0.679 | 0.125 | 5.42E+00 |

| | | | | | | |
|---|---|---|---|---|---|---|
| *Mongolia_N_North* | China_NEastAsia_Inland_EN + Transbaikal_EMN_old + Dzhilinda1_M_N_8.4kya | 0.130 | Dzhilinda1_M_N_8.4kya | 0.047 | 0.028 | 1.66E+00 |
| Syalakh-Belkachi | Dzhilinda1_M_N_8.4kya | 0.066 | Dzhilinda1_M_N_8.4kya | 1.000 | 0.000 | 3.13E+13 |
| Syalakh-Belkachi | China_AmurRiver_LPaleolithic_14K + Dzhilinda1_M_N_8.4kya | 0.919 | China_AmurRiver_LPaleolithic_14K | 0.144 | 0.037 | 3.91E+00 |
| Syalakh-Belkachi | China_AmurRiver_LPaleolithic_14K + Dzhilinda1_M_N_8.4kya | 0.919 | Dzhilinda1_M_N_8.4kya | 0.856 | 0.037 | 2.33E+01 |
| Syalakh-Belkachi | China_NEastAsia_Coastal_EN + Dzhilinda1_M_N_8.4kya | 0.845 | China_NEastAsia_Coastal_EN | 0.194 | 0.042 | 4.59E+00 |
| Syalakh-Belkachi | China_NEastAsia_Coastal_EN + Dzhilinda1_M_N_8.4kya | 0.845 | Dzhilinda1_M_N_8.4kya | 0.806 | 0.042 | 1.91E+01 |
| Syalakh-Belkachi | China_AmurRiver_Mesolithic + Dzhilinda1_M_N_8.4kya | 0.834 | China_AmurRiver_Mesolithic | 0.154 | 0.035 | 4.46E+00 |
| Syalakh-Belkachi | China_AmurRiver_Mesolithic + Dzhilinda1_M_N_8.4kya | 0.834 | Dzhilinda1_M_N_8.4kya | 0.846 | 0.035 | 2.44E+01 |
| Syalakh-Belkachi | China_AmurRiver_EarlyN + Dzhilinda1_M_N_8.4kya | 0.765 | China_AmurRiver_EarlyN | 0.145 | 0.035 | 4.13E+00 |
| Syalakh-Belkachi | China_AmurRiver_EarlyN + Dzhilinda1_M_N_8.4kya | 0.765 | Dzhilinda1_M_N_8.4kya | 0.855 | 0.035 | 2.43E+01 |
| Syalakh-Belkachi | Transbaikal_EMN_old + Dzhilinda1_M_N_8.4kya | 0.691 | Transbaikal_EMN_old | 0.206 | 0.051 | 4.02E+00 |
| Syalakh-Belkachi | Transbaikal_EMN_old + Dzhilinda1_M_N_8.4kya | 0.691 | Dzhilinda1_M_N_8.4kya | 0.794 | 0.051 | 1.55E+01 |
| Syalakh-Belkachi | China_AmurRiver_LPaleolithic_19K + Dzhilinda1_M_N_8.4kya | 0.337 | China_AmurRiver_LPaleolithic_19K | 0.138 | 0.046 | 3.03E+00 |
| Syalakh-Belkachi | China_AmurRiver_LPaleolithic_19K + Dzhilinda1_M_N_8.4kya | 0.337 | Dzhilinda1_M_N_8.4kya | 0.862 | 0.046 | 1.89E+01 |
| Syalakh-Belkachi | KhatystyrCave_M_10.2kya + Dzhilinda1_M_N_8.4kya | 0.315 | KhatystyrCave_M_10.2kya | 0.149 | 0.056 | 2.64E+00 |
| Syalakh-Belkachi | KhatystyrCave_M_10.2kya + Dzhilinda1_M_N_8.4kya | 0.315 | Dzhilinda1_M_N_8.4kya | 0.851 | 0.056 | 1.51E+01 |
| Syalakh-Belkachi | China_NEastAsia_Inland_EN + Dzhilinda1_M_N_8.4kya | 0.262 | China_NEastAsia_Inland_EN | 0.163 | 0.057 | 2.85E+00 |
| Syalakh-Belkachi | China_NEastAsia_Inland_EN + Dzhilinda1_M_N_8.4kya | 0.262 | Dzhilinda1_M_N_8.4kya | 0.837 | 0.057 | 1.47E+01 |
| Syalakh-Belkachi | China_SEastAsia_Coastal_EN + Dzhilinda1_M_N_8.4kya | 0.120 | China_SEastAsia_Coastal_EN | 0.085 | 0.042 | 2.02E+00 |
| Syalakh-Belkachi | China_SEastAsia_Coastal_EN + Dzhilinda1_M_N_8.4kya | 0.120 | Dzhilinda1_M_N_8.4kya | 0.915 | 0.042 | 2.18E+01 |
| Syalakh-Belkachi | KhatystyrCave_M_10.2kya + Kolyma_M_10.1kya | 0.065 | KhatystyrCave_M_10.2kya | 0.328 | 0.044 | 7.45E+00 |
| Syalakh-Belkachi | KhatystyrCave_M_10.2kya + Kolyma_M_10.1kya | 0.065 | Kolyma_M_10.1kya | 0.672 | 0.044 | 1.52E+01 |

#### Table S13

| Target | Model | P | Source | Prop. | SE | Z |
|---|---|---|---|---|---|---|
| Altai_N | Altai_N_old | 0.150 | Altai_N_old | 1.000 | 0.000 | 3.13E+13 |
| Transbaikal_EMN | Transbaikal_EMN_old | 0.066 | Transbaikal_EMN_old | 1.000 | 0.000 | 3.13E+13 |
| China_AmurRiver_N | China_AmurRiver_EarlyN + China_NEastAsia_Inland_EN | 0.137 | China_AmurRiver_EarlyN | 0.921 | 0.037 | 2.51E+01 |
| China_AmurRiver_N | China_AmurRiver_EarlyN + China_NEastAsia_Inland_EN | 0.137 | China_NEastAsia_Inland_EN | 0.079 | 0.037 | 2.17E+00 |
| China_AmurRiver_N | China_AmurRiver_EarlyN + Transbaikal_EMN_old | 0.113 | China_AmurRiver_EarlyN | 0.921 | 0.031 | 2.97E+01 |
| China_AmurRiver_N | China_AmurRiver_EarlyN + Transbaikal_EMN_old | 0.113 | Transbaikal_EMN_old | 0.079 | 0.031 | 2.57E+00 |
| China_AmurRiver_N | China_AmurRiver_EarlyN + Dzhilinda1_M_N_8.4kya | 0.108 | China_AmurRiver_EarlyN | 0.957 | 0.020 | 4.81E+01 |
| China_AmurRiver_N | China_AmurRiver_EarlyN + Dzhilinda1_M_N_8.4kya | 0.108 | Dzhilinda1_M_N_8.4kya | 0.043 | 0.020 | 2.17E+00 |
| China_AmurRiver_N | China_AmurRiver_EarlyN + China_NEastAsia_Coastal_EN | 0.106 | China_AmurRiver_EarlyN | 0.893 | 0.049 | 1.81E+01 |
| China_AmurRiver_N | China_AmurRiver_EarlyN + China_NEastAsia_Coastal_EN | 0.106 | China_NEastAsia_Coastal_EN | 0.107 | 0.049 | 2.16E+00 |

| | | | | | | |
|---|---|---|---|---|---|---|
| China_AmurRiver_N | China_AmurRiver_EarlyN + Transbaikal_EMN | 0.090 | China_AmurRiver_EarlyN | 0.920 | 0.031 | 2.94E+01 |
| China_AmurRiver_N | China_AmurRiver_EarlyN + Transbaikal_EMN | 0.090 | Transbaikal_EMN | 0.080 | 0.031 | 2.54E+00 |
| China_AmurRiver_N | China_AmurRiver_EarlyN + *Mongolia_N_North* | 0.083 | China_AmurRiver_EarlyN | 0.928 | 0.027 | 3.38E+01 |
| China_AmurRiver_N | China_AmurRiver_EarlyN + *Mongolia_N_North* | 0.083 | *Mongolia_N_North* | 0.072 | 0.027 | 2.64E+00 |
| China_AmurRiver_N | China_AmurRiver_EarlyN + Syalakh-Belkachi | 0.082 | China_AmurRiver_EarlyN | 0.939 | 0.023 | 4.06E+01 |
| China_AmurRiver_N | China_AmurRiver_EarlyN + Syalakh-Belkachi | 0.082 | Syalakh-Belkachi | 0.061 | 0.023 | 2.62E+00 |
| China_AmurRiver_N | China_AmurRiver_EarlyN + Altai_N | 0.080 | China_AmurRiver_EarlyN | 0.976 | 0.010 | 9.53E+01 |
| China_AmurRiver_N | China_AmurRiver_EarlyN + Altai_N | 0.080 | Altai_N | 0.024 | 0.010 | 2.34E+00 |
| China_AmurRiver_N | China_AmurRiver_EarlyN + Cisbaikal_EN | 0.072 | China_AmurRiver_EarlyN | 0.923 | 0.028 | 3.25E+01 |
| China_AmurRiver_N | China_AmurRiver_EarlyN + Cisbaikal_EN | 0.072 | Cisbaikal_EN | 0.077 | 0.028 | 2.72E+00 |
| China_AmurRiver_N | China_AmurRiver_Mesolithic + Transbaikal_EMN | 0.061 | China_AmurRiver_Mesolithic | 0.832 | 0.028 | 2.94E+01 |
| China_AmurRiver_N | China_AmurRiver_Mesolithic + Transbaikal_EMN | 0.061 | Transbaikal_EMN | 0.168 | 0.028 | 5.94E+00 |
| China_AmurRiver_N | Khaiyrgas + China_AmurRiver_EarlyN | 0.054 | Khaiyrgas | 0.029 | 0.014 | 2.17E+00 |
| China_AmurRiver_N | Khaiyrgas + China_AmurRiver_EarlyN | 0.054 | China_AmurRiver_EarlyN | 0.971 | 0.014 | 7.16E+01 |
| Syalakh-Belkachi | Dzhilinda1_M_N_8.4kya + Cisbaikal_EN | 0.209 | Dzhilinda1_M_N_8.4kya | 0.762 | 0.051 | 1.50E+01 |
| Syalakh-Belkachi | Dzhilinda1_M_N_8.4kya + Cisbaikal_EN | 0.209 | Cisbaikal_EN | 0.238 | 0.051 | 4.70E+00 |
| Syalakh-Belkachi | Dzhilinda1_M_N_8.4kya + *Mongolia_N_North* | 0.193 | Dzhilinda1_M_N_8.4kya | 0.757 | 0.037 | 2.02E+01 |
| Syalakh-Belkachi | Dzhilinda1_M_N_8.4kya + *Mongolia_N_North* | 0.193 | *Mongolia_N_North* | 0.243 | 0.037 | 6.49E+00 |
| Syalakh-Belkachi | Transbaikal_EMN_old + Dzhilinda1_M_N_8.4kya | 0.168 | Transbaikal_EMN_old | 0.240 | 0.041 | 5.87E+00 |
| Syalakh-Belkachi | Transbaikal_EMN_old + Dzhilinda1_M_N_8.4kya | 0.168 | Dzhilinda1_M_N_8.4kya | 0.760 | 0.041 | 1.86E+01 |

**Table S1-VI. A-8 & VI. A-9 |** Passing models for the populations in Tranche 5. For *Mongolia_N_North* and *Syalakh-Belkachi* in Set A, both passing 2-way models and 3-way models are presented, as the 2-way model is marginally passing. For all other targets no simpler models pass. **References:** [Tranche 0 + Tranche 1 + Tranche 2 + Tranche 3 + Tranche 4 + Tranche 5(only for Set B)]

**Sources:** [*Peru_Laramate_900BP*, *AG3, Khairygas_16.7kya, Ust_Kyakhta_14kya, Kolyma_M_10.1kya, Dzhilinda1_M_N_8.4kya, China_NEastAsia_Inland_EN*, *China_SEastAsia_Coastal_EN*, *China_NEastAsia_Coastal_EN*, *China_AmurRiver_Paleolithic_19K, China_AmurRiver_LPaleolithic_14K, China_AmurRiver_Mesolithic, KhatystyrCave_M_10.2kya, Transbaikal_EMN_old, Altai_N_old (Syalakh_Belkachi, Yakutia_LNBA, Transbaikal_EMN, Mongolia_N_North, China_AmurRiver_N are only included in set B)*]

#### VI.A.ii.f) Tranche 6: Yakutia_LNBA, Cisbaikal_LNBA

***Yakutia_LNBA.*** *Yakutia_LNBA,* comprising individuals dating to ~4500-3200 years BP and from a region stretching from the Kan river valley in the Southeastern Krasnoyarsk region, to the Middle to Lower Lena valley in Central Yakutia and the Kolyma River basin in far Northeastern Yakutia close to Beringia, can be modeled as a three-way admixture between *Syalakh-Belkachi* (~50%), *Transbaikal_EMN* (~42%), and a source related to Amur River-related populations (~8%; p = 0.07). 4-way models with a contribution from *Dzhilinda1_M_N_8.4kya* or *KhatystyrCave_M_10.2kya* that is within the standard error pass with higher p-values. No models pass in set B, but a 4-way model retaining the high levels of admixture from *Syalakh-Belkachi* and *Transbaikal_EMN* found in passing models in set A comes close.

***Cisbaikal_LNBA.*** No models pass for *Cisbaikal_LNBA* in either set A or B. We proceed to move Cisbaikal_LNBA to older tranches in the following sections.

All models in this set are found in Table 6B (Set A) or Table 7A (Set B) in Data S3.

**Table S14**

| Target | Model | P | Source | Prop. | SE | Z |
|---|---|---|---|---|---|---|
| *Yakutia_LN BA* | Syalakh-Belkachi + Transbaikal_EMN + China_AmurRiver_N | 0.069 | Syalakh-Belkachi | 0.503 | 0.028 | 1.80E+01 |
| *Yakutia_LN BA* | Syalakh-Belkachi + Transbaikal_EMN + China_AmurRiver_N | 0.069 | Transbaikal_EMN | 0.418 | 0.034 | 1.23E+01 |
| *Yakutia_LN BA* | Syalakh-Belkachi + Transbaikal_EMN + China_AmurRiver_N | 0.069 | China_AmurRiver_N | 0.079 | 0.026 | 3.02E+00 |
| *Yakutia_LN BA* | China_AmurRiver_Mesolithic + Syalakh-Belkachi + Transbaikal_EMN | 0.051 | China_AmurRiver_Mesolithic | 0.081 | 0.023 | 3.46E+00 |
| *Yakutia_LN BA* | China_AmurRiver_Mesolithic + Syalakh-Belkachi + Transbaikal_EMN | 0.051 | Syalakh-Belkachi | 0.495 | 0.028 | 1.80E+01 |
| *Yakutia_LN BA* | China_AmurRiver_Mesolithic + Syalakh-Belkachi + Transbaikal_EMN | 0.051 | Transbaikal_EMN | 0.424 | 0.032 | 1.31E+01 |

**Table S15**

| Target | Model | P | Source | Prop. | SE | Z |
|---|---|---|---|---|---|---|
| *Yakutia_LN BA* | Ust_Kyakhta_14kya + KhatystyrCave_M_10.2kya + Syalakh-Belkachi + Transbaikal_EMN | 0.022 | Ust_Kyakhta_14kya | 0.081 | 0.032 | 2.50E+00 |
| *Yakutia_LN BA* | Ust_Kyakhta_14kya + KhatystyrCave_M_10.2kya + Syalakh-Belkachi + Transbaikal_EMN | 0.022 | KhatystyrCave_M_10.2kya | 0.163 | 0.103 | 1.59E+00 |
| *Yakutia_LN BA* | Ust_Kyakhta_14kya + KhatystyrCave_M_10.2kya + Syalakh-Belkachi + Transbaikal_EMN | 0.022 | Syalakh-Belkachi | 0.364 | 0.066 | 5.47E+00 |
| *Yakutia_LN BA* | Ust_Kyakhta_14kya + KhatystyrCave_M_10.2kya + Syalakh-Belkachi + Transbaikal_EMN | 0.022 | Transbaikal_EMN | 0.393 | 0.075 | 5.21E+00 |

**Table S1-VI. A-10 & VI. A-11 |** Passing models, or the best failing model, for the populations in Tranche 6. No simpler models pass. **References:** [Tranche 0 + Tranche 1 + Tranche 2 + Tranche 3 + Tranche 4 + Tranche 5 (+ *Cisbaikal_LNBA* in set B)]
**Sources:** [*Peru_Laramate_900BP, AG3, Khairygas_16.7kya, Ust_Kyakhta_14kya, China_NEastAsia_Coastal_EN, China_NEastAsia_Inland_EN, China_SEastAsia_Coastal_EN, China_AmurRiver_Paleolithic_19K, China_AmurRiver_LPaleolithic_14K, China_AmurRiver_Mesolithic, China_AmurRiver_EarlyN, KhatystyrCave_M_10.2kya, Kolyma_M_10.1kya, Transbaikal_EMN_old, Dzhilinda1_M_N_8.4kya, Altai_N_old, Syalakh_Belkachi, Transbaikal_EMN, Mongolia_N_North, China_AmurRiver_N, Cisbaikal_EN, (Cisbaikal_LNBA in set B)*]

#### VI.A.iii. Models with Cisbaikal_LNBA in Tranche 3

##### VI.A.iii.a) Finding a Tranche for Cisbaikal_LNBA

***Finding a tranche for Cisbaikal_LNBA.*** As *Cisbaikal_LNBA* cannot be modeled in tranche 6, we attempt to model this population in successively older tranches. *Cisbaikal_LNBA* cannot be modeled as a member of Tranche 5 or 4, though 2- and 3-way models in Tranches 5 and 4 only fail marginally. We list the marginally failing models in Set A for *Cisbaikal_LNBA* in Tranches 5 and 4 in VI. A-12 and 13. We note that a model with a high p-value for *Cisbaikal_LNBA* in tranche 4 has *Cisbaikal_LNBA* with ancestry from *Ust_Kyakhta_14kya* and *China_NEastAsia_Inland_EN* and a small negative contribution from *KhatystyrCave_M_10.2kya* just above the standard error; *KhatystyrCave_M_10.2kya* has high levels of admixture from Amur-related populations in previous *qpAdm*. That this *qpAdm* model passes may indicate that *Cisbaikal_LNBA* received its East Asian ancestry from a source even further in the Inland direction on the cline between Inland Northaast Asians and Amur River Basin populations than the Yumin individual was itself.

All models in this for Cisbaikal_LNBA as a member of Tranches 4 or 5 are found in Table 8 in Data S3.

**Table S16**

| Target | Model | P | Source | Prop. | SE | Z |
|---|---|---|---|---|---|---|
| Cisbaikal_LNBA | Ust_Kyakhta_14kya + Transbaikal_EMN_old | 0.003 | Ust_Kyakhta_14kya | 0.882 | 0.035 | 2.49E+01 |

| | | | | | | |
|---|---|---|---|---|---|---|
| Cisbaikal_LNBA | Ust_Kyakhta_14kya + Transbaikal_EMN_old | 0.003 | Transbaikal_EMN_old | 0.118 | 0.035 | 3.34E+00 |
| Cisbaikal_LNBA | Altai_N_old + Ust_Kyakhta_14kya + Transbaikal_EMN_old | 0.005 | Altai_N_old | 0.135 | 0.143 | 9.45E-01 |
| Cisbaikal_LNBA | Altai_N_old + Ust_Kyakhta_14kya + Transbaikal_EMN_old | 0.005 | Ust_Kyakhta_14kya | 0.571 | 0.315 | 1.81E+00 |
| Cisbaikal_LNBA | Altai_N_old + Ust_Kyakhta_14kya + Transbaikal_EMN_old | 0.005 | Transbaikal_EMN_old | 0.295 | 0.173 | 1.70E+00 |

#### Table S17

| Target | Model | P | Source | Prop. | SE | Z |
|---|---|---|---|---|---|---|
| Cisbaikal_LNBA | Ust_Kyakhta_14kya + China_NEastAsia_Inland_EN | 0.036 | Ust_Kyakhta_14kya | 0.855 | 0.041 | 2.10E+01 |
| Cisbaikal_LNBA | Ust_Kyakhta_14kya + China_NEastAsia_Inland_EN | 0.036 | China_NEastAsia_Inland_EN | 0.145 | 0.041 | 3.57E+00 |
| Cisbaikal_LNBA | Russia_AfontovaGora3 + Ust_Kyakhta_14kya + China_NEastAsia_Inland_EN | 0.023 | Russia_AfontovaGora3 | 0.024 | 0.028 | 8.31E-01 |
| Cisbaikal_LNBA | Russia_AfontovaGora3 + Ust_Kyakhta_14kya + China_NEastAsia_Inland_EN | 0.023 | Ust_Kyakhta_14kya | 0.758 | 0.136 | 5.58E+00 |
| Cisbaikal_LNBA | Russia_AfontovaGora3 + Ust_Kyakhta_14kya + China_NEastAsia_Inland_EN | 0.023 | China_NEastAsia_Inland_EN | 0.218 | 0.109 | 2.00E+00 |
| Cisbaikal_LNBA | Ust_Kyakhta_14kya + China_NEastAsia_Inland_EN + KhatystyrCave_M_10.2kya | 0.241 | Ust_Kyakhta_14kya | 0.853 | 0.040 | 2.11E+01 |
| Cisbaikal_LNBA | Ust_Kyakhta_14kya + China_NEastAsia_Inland_EN + KhatystyrCave_M_10.2kya | 0.241 | China_NEastAsia_Inland_EN | 0.225 | 0.069 | 3.25E+00 |
| Cisbaikal_LNBA | Ust_Kyakhta_14kya + China_NEastAsia_Inland_EN + KhatystyrCave_M_10.2kya | 0.241 | KhatystyrCave_M_10.2kya | -0.078 | 0.062 | -1.26E+00 |

**Table S1-VI. A-12 & VI. A-13 |** Best failing models for *Cisbaikal_LNBA* as a member of tranche 5 and 4 respectively.
**References:** [Tranche 0 + Tranche 1 + Tranche 2 + Tranche 3], [Tranche 0 + Tranche 1 + Tranche 2 + Tranche 3 + Tranche 4]
**Sources:** [*Russia_AfontovaGora, Peru_Laramate_900BP, Khaiyrgas, Ust_Kyakhta_14kya, China_AmurRiver_LPaleolithic_19K, China_AmurRiver_LPaleolithic_14K, China_AmurRiver_Mesolithic, China_NEastAsia_Coastal_EN, China_NEastAsia_Inland_EN, China_SEastAsia_Coastal_EN, KhatystyrCave_M_10.2kya, Kolyma_M_10.1kya, Transbaikal_EMN_old, Altai_N_old*], [*Russia_AfontovaGora, Peru_Laramate_900BP, Khaiyrgas, Ust_Kyakhta_14kya, China_AmurRiver_LPaleolithic_19K, China_AmurRiver_LPaleolithic_14K, China_AmurRiver_Mesolithic, China_NEastAsia_Coastal_EN, China_NEastAsia_Inland_EN, China_SEastAsia_Coastal_EN, KhatystyrCave_M_10.2kya, Kolyma_M_10.1kya*]

***Cisbaikal_LNBA as a member of Tranche 3.*** We find that Cisbaikal_LNBA can be successfully modeled as a member of Tranche 3. In both sets A and B, a model of Cisbaikal_LNBA as deriving most of its ancestry (~86%) from Ust_Kyakhta_14kya, with some ancestry from Inland Northeast Asians (~14%), is the only model passing with a high p-value. A model with Coastal Northeast Asians from the Bohai Sea region ~9000-8300BP passes marginally in set A. We suggest that the model with ancestry from Inland Northeast Asians that passes in both set A and B is more plausible as similar models fit the best in the previous tranches, and as Cisbaikal_LNBA possesses more affinity to Inland Northeast Asians than can be explained by admixture between *Ust_Kyakhta_14kya* and Coastal Northeast Asians in f4-statistics (Main Text Fig. 2B); in fact, *Cisbaikal_LNBA* possess more affinity to Inland Northeast Asians than can be explained by an admixture event between *Ust_Kyakhta_14kya* and the preceding population of the Cisbaikal region (i.e. a model of *Ust_Kykahta* + *Cisbaikal_EN*; Fig. 2B).

All models in this for Cisbaikal_LNBA as a member of Tranche 3 are found in Table 4A (Set A) and Table 4C (Set B) in Data S3.

**Table S18**

| Target | Model | P | Source | Prop. | SE | Z |
|---|---|---|---|---|---|---|
| Cisbaikal_LNBA | Ust_Kyakhta_14kya + China_NEastAsia_Inland_EN | 0.481 | Ust_Kyakhta_14kya | 0.866 | 0.040 | 2.19E+01 |
| Cisbaikal_LNBA | Ust_Kyakhta_14kya + China_NEastAsia_Inland_EN | 0.481 | China_NEastAsia_Inland_EN | 0.134 | 0.040 | 3.40E+00 |
| Cisbaikal_LNBA | Ust_Kyakhta_14kya + China_NEastAsia_Coastal_EN | 0.062 | Ust_Kyakhta_14kya | 0.913 | 0.034 | 2.66E+01 |
| Cisbaikal_LNBA | Ust_Kyakhta_14kya + China_NEastAsia_Coastal_EN | 0.062 | China_NEastAsia_Coastal_EN | 0.087 | 0.034 | 2.55E+00 |

**Table S19**

| Target | Model | P | Source | Prop. | SE | Z |
|---|---|---|---|---|---|---|
| Cisbaikal_LNBA | Ust_Kyakhta_14kya + China_NEastAsia_Inland_EN | 0.103 | Ust_Kyakhta_14kya | 0.863 | 0.041 | 2.12E+01 |
| Cisbaikal_LNBA | Ust_Kyakhta_14kya + China_NEastAsia_Inland_EN | 0.103 | China_NEastAsia_Inland_EN | 0.137 | 0.041 | 3.38E+00 |

**Table S1-VI. A-14 & Table S1-VI. A-15 |** Passing 2-way models for *Cisbaikal_LNBA*. **References:** [Tranche 0 + Tranche 1 + Tranche 2] **Sources:** [*Russia_AfontovaGora, Peru_Laramate_900BP, Khaiyrgas, Ust_Kyakhta_14kya, China_AmurRiver_LPaleolithic_19K, China_AmurRiver_LPaleolithic_14K, China_AmurRiver_Mesolithic, China_NEastAsia_Coastal_EN, China_NEastAsia_Inland_EN, China_SEastAsia_Coastal_EN*, (*KhatystyrCave_M_10.2kya, Kolyma_M_10.1kya* only for Set B)]

##### VI.A.iii.b) Tranche 3: KhatystyrCave_M_10.2kya & Kolyma_M_10.1kya

***Rest of tranche 3 with the addition of Cisbaikal_LNBA to the references.***

As the models in set A for KhatystyrCave_M_10.2kya/*Yakutia_Mesolithic* and *Kolyma_M_10.1kya* are identical when we include *Cisbaikal_LNBA* in tranche 3, we do not repeat these models; however, set B is modified as *Cisbaikal_LNBA* will appear in the references/sources for the models in B.

***KhatystyrCave_M_10.2kya/Yakutia_Mesolithic.*** With a reference/source set including tranches 0 + 1 + 2 plus *Cisbaikal_LNBA*, we find that only a single 3-way model passes for the Mesolithic Southern Yakutian I25033_d from ~10,000 BP: ~24% *Cisbaikal_LNBA* + ~61% *China_AmurRiver_Mesolithic* and a small fraction of ancestry from a Southeast Asian-related source represented by *China_SEastAsia_Coastal_EN,* similar to the models in set A, except that *Cisbaikal_LNBA,* itself having most of its ancestry from APSs, has replaced the APS source.

***Kolyma_M_10.1kya.*** We find once again that passing models involve a 2-way admixture of a Native American- or APS-related source (~72-77%) and an East-Asian-related source; however, the number of passing models has fallen when compared to the models in the previous section (Tables VI. A-4 & VI. A-5).

All models in this tranche can be found in Table 4C in Data S3.

**Table S20**

| Target | Model | P | Source | Prop. | SE | Z |
|---|---|---|---|---|---|---|
| KhatystyrCave_M_10.2kya | China_AmurRiver_Mesolithic + China_SEastAsia_Coastal_EN + Cisbaikal_LNBA | 0.068 | China_AmurRiver_Mesolithic | 0.611 | 0.076 | 8.04E+00 |

| | | | | | | |
|---|---|---|---|---|---|---|
| KhatystyrCave _M_10.2kya | China_AmurRiver_Mesolithic + China_SEastAsia_Coastal_EN + Cisbaikal_LNBA | 0.068 | China_SEastAsia_Coastal_EN | 0.147 | 0.093 | 1.59E+00 |
| KhatystyrCave _M_10.2kya | China_AmurRiver_Mesolithic + China_SEastAsia_Coastal_EN + Cisbaikal_LNBA | 0.068 | Cisbaikal_LNBA | 0.241 | 0.077 | 3.14E+00 |
| Kolyma_M_10 .1kya | Peru_Laramate_900BP + China_NEastAsia_Inland_EN | 0.141 | Peru_Laramate_900BP | 0.771 | 0.041 | 1.87E+01 |
| Kolyma_M_10 .1kya | Peru_Laramate_900BP + China_NEastAsia_Inland_EN | 0.141 | China_NEastAsia_Inland_EN | 0.229 | 0.041 | 5.57E+00 |
| Kolyma_M_10 .1kya | Khaiyrgas + China_AmurRiver_LPaleolithic_14K | 0.110 | Khaiyrgas | 0.726 | 0.031 | 2.34E+01 |
| Kolyma_M_10 .1kya | Khaiyrgas + China_AmurRiver_LPaleolithic_14K | 0.110 | China_AmurRiver_LPaleolithic_14K | 0.274 | 0.031 | 8.82E+00 |
| Kolyma_M_10 .1kya | Khaiyrgas + China_AmurRiver_Mesolithic | 0.068 | Khaiyrgas | 0.722 | 0.030 | 2.41E+01 |
| Kolyma_M_10 .1kya | Khaiyrgas + China_AmurRiver_Mesolithic | 0.068 | China_AmurRiver_Mesolithic | 0.278 | 0.030 | 9.29E+00 |

**Table 20 |** Passing models for the populations in Tranche 3, set B, with *Cisbaikal_LNBA*. No simpler models pass. **References:** [Tranche 0 + Tranche 1 + Tranche 2 + Cisbaikal_LNBA (+ Tranche 3 only for Set B)]
**Sources:** [*Peru_Laramate_900BP, Russia_AfontovaGora3, Khaiyrgas, Ust_Kyakhta_14kya, China_AmurRiver_LPaleolithic_19K, China_AmurRiver_LPaleolithic_14K, China_AmurRiver_Mesolithic, China_NEastAsia_Coastal_EN, China_NEastAsia_Inland_EN, China_SEastAsia_Coastal_EN (KhatystyrCave_M_10.2kya, Kolyma_M_10.1kya, Cisbaikal_LNBA included in set B)*]

##### VI.A.iii.c) Tranche 4: Dzhilinda1_M_N_8.4kya, Altai_N_old & Transbaikal_EMN_old

***Dzhilinda1_M_N_8.4kya. Dzhilinda1_M_N_8.4kya.*** The model of *Dzhilinda1_M_N_8.4kya* with complete descent from *Ust_Kyakhta_14kya* now fails, as do most 2-way models previously listed. A 2-way model of *Dzhilinda1_M_N_8.4kya* as admixed between *Khairygas_16.7kya* and *KhatystyrCave_M_10.2kya* passes marginally (p = 0.052) in set A. 3-way models in Set A with higher p-values now incorporate multiple APS sources, one of which is always *Kolyma_M_10.1kya* and the other of which is *Ust_Kyakhta_14kya* or *Cisbaikal_LNBA*, with the third source of ancestry attributed to an East Asian population from the Amur River region; we list them below. The same marginally passing 2-way model passes in set B.

***Altai_N_old.*** The model of *Altai_N_old* as deriving all its APS-related ancestry from *Khairygas_16.7kya* with some admixture from *AG3* now fails. All passing models for *Altai_N_old* require a 3-way admixture between a much larger quantity of ancestry from *AG3* (~26-43%), one of either *Ust_Kyakhta_14kya* or *Cisbaikal_LNBA* (~26-43%), and a minor contribution from an additional source which may be either East Asian or APS-related. A similar set of models passes in set B.

***Transbaikal_EMN_old.*** The previously presented model of *Transbaikal_EMN_old* as a 2-way admixture between *China_NEastAsia_Inland_EN* and *KhatystyrCave_M_10.2kya* fails in set A but passes in set B. In set A, *Cisbaikal_LNBA* must be moved into the sources, but contributes a very small proportion that falls below the standard error.

All models in this tranche can be found in Table 5C (Set A) and Table 5D (Set B) in Data S3.

Table S21

| Target | Model | P | Source | Prop. | SE | Z |
|---|---|---|---|---|---|---|
| Dzhilinda1_M_N_8.4k ya | Khaiyrgas + KhatystyrCave_M_10.2kya | 0.052 | Khaiyrgas | 0.604 | 0.047 | 1.29E+01 |
| Dzhilinda1_M_N_8.4k ya | Khaiyrgas + KhatystyrCave_M_10.2kya | 0.052 | KhatystyrCave_M_10.2ky a | 0.396 | 0.047 | 8.43E+00 |
| Dzhilinda1_M_N_8.4k ya | Ust_Kyakhta_14kya + KhatystyrCave_M_10.2kya + Kolyma_M_10.1kya | 0.498 | Ust_Kyakhta_14kya | 0.244 | 0.059 | 4.12E+00 |

| | | | | | | |
|---|---|---|---|---|---|---|
| Dzhilinda1_M_N_8.4kya | Ust_Kyakhta_14kya + KhatystyrCave_M_10.2kya + Kolyma_M_10.1kya | 0.498 | KhatystyrCave_M_10.2kya | 0.102 | 0.048 | 2.13E+00 |
| Dzhilinda1_M_N_8.4kya | Ust_Kyakhta_14kya + KhatystyrCave_M_10.2kya + Kolyma_M_10.1kya | 0.498 | Kolyma_M_10.1kya | 0.654 | 0.067 | 9.80E+00 |
| Dzhilinda1_M_N_8.4kya | KhatystyrCave_M_10.2kya + Kolyma_M_10.1kya + Cisbaikal_LNBA | 0.403 | KhatystyrCave_M_10.2kya | 0.132 | 0.052 | 2.53E+00 |
| Dzhilinda1_M_N_8.4kya | KhatystyrCave_M_10.2kya + Kolyma_M_10.1kya + Cisbaikal_LNBA | 0.403 | Kolyma_M_10.1kya | 0.858 | 0.118 | 7.28E+00 |
| Dzhilinda1_M_N_8.4kya | KhatystyrCave_M_10.2kya + Kolyma_M_10.1kya + Cisbaikal_LNBA | 0.403 | Cisbaikal_LNBA | 0.010 | 0.119 | 8.33E-02 |
| Dzhilinda1_M_N_8.4kya | Ust_Kyakhta_14kya + China_AmurRiver_LPaleolithic_14K + Kolyma_M_10.1kya | 0.154 | Ust_Kyakhta_14kya | 0.266 | 0.061 | 4.39E+00 |
| Dzhilinda1_M_N_8.4kya | Ust_Kyakhta_14kya + China_AmurRiver_LPaleolithic_14K + Kolyma_M_10.1kya | 0.154 | China_AmurRiver_LPaleolithic_14K | 0.079 | 0.038 | 2.07E+00 |
| Dzhilinda1_M_N_8.4kya | Ust_Kyakhta_14kya + China_AmurRiver_LPaleolithic_14K + Kolyma_M_10.1kya | 0.154 | Kolyma_M_10.1kya | 0.656 | 0.077 | 8.52E+00 |
| Dzhilinda1_M_N_8.4kya | China_AmurRiver_EarlyN + Kolyma_M_10.1kya + Cisbaikal_LNBA | 0.112 | China_AmurRiver_EarlyN | 0.073 | 0.042 | 1.73E+00 |
| Dzhilinda1_M_N_8.4kya | China_AmurRiver_EarlyN + Kolyma_M_10.1kya + Cisbaikal_LNBA | 0.112 | Kolyma_M_10.1kya | 0.901 | 0.124 | 7.30E+00 |
| Dzhilinda1_M_N_8.4kya | China_AmurRiver_EarlyN + Kolyma_M_10.1kya + Cisbaikal_LNBA | 0.112 | Cisbaikal_LNBA | 0.025 | 0.123 | 2.07E-01 |
| Dzhilinda1_M_N_8.4kya | Ust_Kyakhta_14kya + China_AmurRiver_EarlyN + Kolyma_M_10.1kya | 0.103 | Ust_Kyakhta_14kya | 0.281 | 0.062 | 4.55E+00 |
| Dzhilinda1_M_N_8.4kya | Ust_Kyakhta_14kya + China_AmurRiver_EarlyN + Kolyma_M_10.1kya | 0.103 | China_AmurRiver_EarlyN | 0.048 | 0.039 | 1.22E+00 |
| Dzhilinda1_M_N_8.4kya | Ust_Kyakhta_14kya + China_AmurRiver_EarlyN + Kolyma_M_10.1kya | 0.103 | Kolyma_M_10.1kya | 0.672 | 0.079 | 8.55E+00 |
| Dzhilinda1_M_N_8.4kya | Khaiyrgas + KhatystyrCave_M_10.2kya + Kolyma_M_10.1kya | 0.064 | Khaiyrgas | 0.593 | 0.326 | 1.82E+00 |
| Dzhilinda1_M_N_8.4kya | Khaiyrgas + KhatystyrCave_M_10.2kya + Kolyma_M_10.1kya | 0.064 | KhatystyrCave_M_10.2kya | 0.399 | 0.155 | 2.57E+00 |
| Dzhilinda1_M_N_8.4kya | Khaiyrgas + KhatystyrCave_M_10.2kya + Kolyma_M_10.1kya | 0.064 | Kolyma_M_10.1kya | 0.009 | 0.471 | 1.84E-02 |
| Dzhilinda1_M_N_8.4kya | Ust_Kyakhta_14kya + China_AmurRiver_Mesolithic + Kolyma_M_10.1kya | 0.057 | Ust_Kyakhta_14kya | 0.287 | 0.062 | 4.65E+00 |
| Dzhilinda1_M_N_8.4kya | Ust_Kyakhta_14kya + China_AmurRiver_Mesolithic + Kolyma_M_10.1kya | 0.057 | China_AmurRiver_Mesolithic | 0.060 | 0.037 | 1.60E+00 |
| Dzhilinda1_M_N_8.4kya | Ust_Kyakhta_14kya + China_AmurRiver_Mesolithic + Kolyma_M_10.1kya | 0.057 | Kolyma_M_10.1kya | 0.653 | 0.077 | 8.47E+00 |
| Dzhilinda1_M_N_8.4kya | China_AmurRiver_Mesolithic + Kolyma_M_10.1kya + Cisbaikal_LNBA | 0.054 | China_AmurRiver_Mesolithic | 0.087 | 0.041 | 2.14E+00 |
| Dzhilinda1_M_N_8.4kya | China_AmurRiver_Mesolithic + Kolyma_M_10.1kya + Cisbaikal_LNBA | 0.054 | Kolyma_M_10.1kya | 0.879 | 0.120 | 7.31E+00 |
| Dzhilinda1_M_N_8.4kya | China_AmurRiver_Mesolithic + Kolyma_M_10.1kya + Cisbaikal_LNBA | 0.054 | Cisbaikal_LNBA | 0.034 | 0.121 | 2.82E-01 |
| Altai_N_old | Russia_AfontovaGora3 + Ust_Kyakhta_14kya + Kolyma_M_10.1kya | 0.561 | Russia_AfontovaGora3 | 0.361 | 0.016 | 2.27E+01 |
| Altai_N_old | Russia_AfontovaGora3 + Ust_Kyakhta_14kya + Kolyma_M_10.1kya | 0.561 | Ust_Kyakhta_14kya | 0.366 | 0.038 | 9.50E+00 |
| Altai_N_old | Russia_AfontovaGora3 + Ust_Kyakhta_14kya + Kolyma_M_10.1kya | 0.561 | Kolyma_M_10.1kya | 0.273 | 0.043 | 6.28E+00 |
| Altai_N_old | Russia_AfontovaGora3 + Kolyma_M_10.1kya + Cisbaikal_LNBA | 0.526 | Russia_AfontovaGora3 | 0.359 | 0.018 | 2.01E+01 |
| Altai_N_old | Russia_AfontovaGora3 + Kolyma_M_10.1kya + Cisbaikal_LNBA | 0.526 | Kolyma_M_10.1kya | 0.428 | 0.095 | 4.52E+00 |
| Altai_N_old | Russia_AfontovaGora3 + Kolyma_M_10.1kya + Cisbaikal_LNBA | 0.526 | Cisbaikal_LNBA | 0.213 | 0.085 | 2.51E+00 |
| Altai_N_old | Russia_AfontovaGora3 + Khaiyrgas + Ust_Kyakhta_14kya | 0.238 | Russia_AfontovaGora3 | 0.278 | 0.032 | 8.62E+00 |
| Altai_N_old | Russia_AfontovaGora3 + Khaiyrgas + Ust_Kyakhta_14kya | 0.238 | Khaiyrgas | 0.457 | 0.092 | 4.95E+00 |

| | | | | | | |
|---|---|---|---|---|---|---|
| Altai_N_old | Russia_AfontovaGora3 + Khaiyrgas + Ust_Kyakhta_14kya | 0.238 | Ust_Kyakhta_14kya | 0.265 | 0.065 | 4.07E+00 |
| Altai_N_old | Russia_AfontovaGora3 + Ust_Kyakhta_14kya + China_AmurRiver_Mesolithic | 0.233 | Russia_AfontovaGora3 | 0.430 | 0.015 | 2.92E+01 |
| Altai_N_old | Russia_AfontovaGora3 + Ust_Kyakhta_14kya + China_AmurRiver_Mesolithic | 0.233 | Ust_Kyakhta_14kya | 0.431 | 0.030 | 1.43E+01 |
| Altai_N_old | Russia_AfontovaGora3 + Ust_Kyakhta_14kya + China_AmurRiver_Mesolithic | 0.233 | China_AmurRiver_Mesolithic | 0.139 | 0.023 | 6.01E+00 |
| Altai_N_old | Russia_AfontovaGora3 + Ust_Kyakhta_14kya + China_NEastAsia_Coastal_EN | 0.205 | Russia_AfontovaGora3 | 0.424 | 0.014 | 2.98E+01 |
| Altai_N_old | Russia_AfontovaGora3 + Ust_Kyakhta_14kya + China_NEastAsia_Coastal_EN | 0.205 | Ust_Kyakhta_14kya | 0.427 | 0.032 | 1.33E+01 |
| Altai_N_old | Russia_AfontovaGora3 + Ust_Kyakhta_14kya + China_NEastAsia_Coastal_EN | 0.205 | China_NEastAsia_Coastal_EN | 0.148 | 0.027 | 5.47E+00 |
| Altai_N_old | Russia_AfontovaGora3 + Khaiyrgas + Cisbaikal_LNBA | 0.186 | Russia_AfontovaGora3 | 0.262 | 0.050 | 5.25E+00 |
| Altai_N_old | Russia_AfontovaGora3 + Khaiyrgas + Cisbaikal_LNBA | 0.186 | Khaiyrgas | 0.546 | 0.168 | 3.26E+00 |
| Altai_N_old | Russia_AfontovaGora3 + Khaiyrgas + Cisbaikal_LNBA | 0.186 | Cisbaikal_LNBA | 0.192 | 0.121 | 1.59E+00 |
| Altai_N_old | Russia_AfontovaGora3 + Ust_Kyakhta_14kya + China_NEastAsia_Inland_EN | 0.179 | Russia_AfontovaGora3 | 0.431 | 0.014 | 2.98E+01 |
| Altai_N_old | Russia_AfontovaGora3 + Ust_Kyakhta_14kya + China_NEastAsia_Inland_EN | 0.179 | Ust_Kyakhta_14kya | 0.358 | 0.042 | 8.61E+00 |
| Altai_N_old | Russia_AfontovaGora3 + Ust_Kyakhta_14kya + China_NEastAsia_Inland_EN | 0.179 | China_NEastAsia_Inland_EN | 0.211 | 0.036 | 5.92E+00 |
| Altai_N_old | Russia_AfontovaGora3 + Ust_Kyakhta_14kya + China_AmurRiver_LPaleolithic_14K | 0.156 | Russia_AfontovaGora3 | 0.430 | 0.015 | 2.94E+01 |
| Altai_N_old | Russia_AfontovaGora3 + Ust_Kyakhta_14kya + China_AmurRiver_LPaleolithic_14K | 0.156 | Ust_Kyakhta_14kya | 0.435 | 0.030 | 1.43E+01 |
| Altai_N_old | Russia_AfontovaGora3 + Ust_Kyakhta_14kya + China_AmurRiver_LPaleolithic_14K | 0.156 | China_AmurRiver_LPaleolithic_14K | 0.136 | 0.023 | 5.77E+00 |
| Altai_N_old | Russia_AfontovaGora3 + Ust_Kyakhta_14kya + China_AmurRiver_LPaleolithic_19K | 0.118 | Russia_AfontovaGora3 | 0.419 | 0.014 | 2.95E+01 |
| Altai_N_old | Russia_AfontovaGora3 + Ust_Kyakhta_14kya + China_AmurRiver_LPaleolithic_19K | 0.118 | Ust_Kyakhta_14kya | 0.434 | 0.030 | 1.44E+01 |
| Altai_N_old | Russia_AfontovaGora3 + Ust_Kyakhta_14kya + China_AmurRiver_LPaleolithic_19K | 0.118 | China_AmurRiver_LPaleolithic_19K | 0.146 | 0.025 | 5.79E+00 |
| Altai_N_old | Russia_AfontovaGora3 + Ust_Kyakhta_14kya + China_AmurRiver_EarlyN | 0.071 | Russia_AfontovaGora3 | 0.434 | 0.015 | 2.92E+01 |
| Altai_N_old | Russia_AfontovaGora3 + Ust_Kyakhta_14kya + China_AmurRiver_EarlyN | 0.071 | Ust_Kyakhta_14kya | 0.433 | 0.032 | 1.34E+01 |
| Altai_N_old | Russia_AfontovaGora3 + Ust_Kyakhta_14kya + China_AmurRiver_EarlyN | 0.071 | China_AmurRiver_EarlyN | 0.134 | 0.025 | 5.36E+00 |
| Transbaikal_EMN_old | China_NEastAsia_Inland_EN + KhatystyrCave_M_10.2kya + Cisbaikal_LNBA | 0.057 | China_NEastAsia_Inland_EN | 0.509 | 0.195 | 2.62E+00 |
| Transbaikal_EMN_old | China_NEastAsia_Inland_EN + KhatystyrCave_M_10.2kya + Cisbaikal_LNBA | 0.057 | KhatystyrCave_M_10.2kya | 0.477 | 0.162 | 2.94E+00 |
| Transbaikal_EMN_old | China_NEastAsia_Inland_EN + KhatystyrCave_M_10.2kya + Cisbaikal_LNBA | 0.057 | Cisbaikal_LNBA | 0.014 | 0.064 | 2.21E-01 |
| Transbaikal_EMN_old | China_AmurRiver_LPaleolithic_14K + China_NEastAsia_Inland_EN + KhatystyrCave_M_10.2kya + Cisbaikal_LNBA | 0.109 | China_AmurRiver_LPaleolithic_14K | 0.414 | 0.169 | 2.45E+00 |
| Transbaikal_EMN_old | China_AmurRiver_LPaleolithic_14K + China_NEastAsia_Inland_EN + KhatystyrCave_M_10.2kya + Cisbaikal_LNBA | 0.109 | China_NEastAsia_Inland_EN | 0.344 | 0.126 | 2.74E+00 |
| Transbaikal_EMN_old | China_AmurRiver_LPaleolithic_14K + China_NEastAsia_Inland_EN + KhatystyrCave_M_10.2kya + Cisbaikal_LNBA | 0.109 | KhatystyrCave_M_10.2kya | 0.071 | 0.176 | 4.04E-01 |
| Transbaikal_EMN_old | China_AmurRiver_LPaleolithic_14K + China_NEastAsia_Inland_EN + KhatystyrCave_M_10.2kya + Cisbaikal_LNBA | 0.109 | Cisbaikal_LNBA | 0.171 | 0.070 | 2.43E+00 |

#### Table S22

| Target | Model | P | Source | Prop. | SE | Z |
|---|---|---|---|---|---|---|
| Dzhilinda1_M_N_8.4kya | Khaiyrgas + KhatystyrCave_M_10.2kya | 0.179 | Khaiyrgas | 0.584 | 0.040 | 1.45E+01 |

| | | | | | | |
|---|---|---|---|---|---|---|
| Dzhilinda1_M_N_8.4kya | Khaiyrgas + KhatystyrCave_M_10.2kya | 0.179 | KhatystyrCave_M_10.2kya | 0.416 | 0.040 | 1.03E+01 |
| Altai_N_old | Russia_AfontovaGora3 + Kolyma_M_10.1kya + Cisbaikal_LNBA | 0.746 | Russia_AfontovaGora3 | 0.358 | 0.018 | 2.04E+01 |
| Altai_N_old | Russia_AfontovaGora3 + Kolyma_M_10.1kya + Cisbaikal_LNBA | 0.746 | Kolyma_M_10.1kya | 0.421 | 0.089 | 4.72E+00 |
| Altai_N_old | Russia_AfontovaGora3 + Kolyma_M_10.1kya + Cisbaikal_LNBA | 0.746 | Cisbaikal_LNBA | 0.222 | 0.080 | 2.78E+00 |
| Altai_N_old | Russia_AfontovaGora3 + Ust_Kyakhta_14kya + Kolyma_M_10.1kya | 0.620 | Russia_AfontovaGora3 | 0.358 | 0.016 | 2.29E+01 |
| Altai_N_old | Russia_AfontovaGora3 + Ust_Kyakhta_14kya + Kolyma_M_10.1kya | 0.620 | Ust_Kyakhta_14kya | 0.353 | 0.036 | 9.68E+00 |
| Altai_N_old | Russia_AfontovaGora3 + Ust_Kyakhta_14kya + Kolyma_M_10.1kya | 0.620 | Kolyma_M_10.1kya | 0.289 | 0.041 | 7.10E+00 |
| Altai_N_old | Russia_AfontovaGora3 + Ust_Kyakhta_14kya + Dzhilinda1_M_N_8.4kya | 0.550 | Russia_AfontovaGora3 | 0.375 | 0.015 | 2.47E+01 |
| Altai_N_old | Russia_AfontovaGora3 + Ust_Kyakhta_14kya + Dzhilinda1_M_N_8.4kya | 0.550 | Ust_Kyakhta_14kya | 0.305 | 0.044 | 6.91E+00 |
| Altai_N_old | Russia_AfontovaGora3 + Ust_Kyakhta_14kya + Dzhilinda1_M_N_8.4kya | 0.550 | Dzhilinda1_M_N_8.4kya | 0.320 | 0.047 | 6.85E+00 |
| Altai_N_old | Russia_AfontovaGora3 + Ust_Kyakhta_14kya + Cisbaikal_LNBA | 0.467 | Russia_AfontovaGora3 | 0.377 | 0.019 | 1.99E+01 |
| Altai_N_old | Russia_AfontovaGora3 + Ust_Kyakhta_14kya + Cisbaikal_LNBA | 0.467 | Ust_Kyakhta_14kya | 0.586 | 0.275 | 2.13E+00 |
| Altai_N_old | Russia_AfontovaGora3 + Ust_Kyakhta_14kya + Cisbaikal_LNBA | 0.467 | Cisbaikal_LNBA | 0.037 | 0.264 | 1.41E-01 |
| Altai_N_old | Russia_AfontovaGora3 + Khaiyrgas + Ust_Kyakhta_14kya | 0.305 | Russia_AfontovaGora3 | 0.269 | 0.031 | 8.73E+00 |
| Altai_N_old | Russia_AfontovaGora3 + Khaiyrgas + Ust_Kyakhta_14kya | 0.305 | Khaiyrgas | 0.484 | 0.087 | 5.57E+00 |
| Altai_N_old | Russia_AfontovaGora3 + Khaiyrgas + Ust_Kyakhta_14kya | 0.305 | Ust_Kyakhta_14kya | 0.247 | 0.062 | 4.02E+00 |
| Altai_N_old | Russia_AfontovaGora3 + Dzhilinda1_M_N_8.4kya + Cisbaikal_LNBA | 0.284 | Russia_AfontovaGora3 | 0.386 | 0.015 | 2.57E+01 |
| Altai_N_old | Russia_AfontovaGora3 + Dzhilinda1_M_N_8.4kya + Cisbaikal_LNBA | 0.284 | Dzhilinda1_M_N_8.4kya | 0.347 | 0.076 | 4.57E+00 |
| Altai_N_old | Russia_AfontovaGora3 + Dzhilinda1_M_N_8.4kya + Cisbaikal_LNBA | 0.284 | Cisbaikal_LNBA | 0.266 | 0.072 | 3.68E+00 |
| Altai_N_old | Russia_AfontovaGora3 + Khaiyrgas + Cisbaikal_LNBA | 0.274 | Russia_AfontovaGora3 | 0.259 | 0.052 | 5.01E+00 |
| Altai_N_old | Russia_AfontovaGora3 + Khaiyrgas + Cisbaikal_LNBA | 0.274 | Khaiyrgas | 0.558 | 0.173 | 3.22E+00 |
| Altai_N_old | Russia_AfontovaGora3 + Khaiyrgas + Cisbaikal_LNBA | 0.274 | Cisbaikal_LNBA | 0.183 | 0.125 | 1.47E+00 |
| Altai_N_old | Russia_AfontovaGora3 + Ust_Kyakhta_14kya + Transbaikal_EMN_old | 0.155 | Russia_AfontovaGora3 | 0.448 | 0.015 | 2.89E+01 |
| Altai_N_old | Russia_AfontovaGora3 + Ust_Kyakhta_14kya + Transbaikal_EMN_old | 0.155 | Ust_Kyakhta_14kya | 0.325 | 0.046 | 7.05E+00 |
| Altai_N_old | Russia_AfontovaGora3 + Ust_Kyakhta_14kya + Transbaikal_EMN_old | 0.155 | Transbaikal_EMN_old | 0.227 | 0.038 | 6.05E+00 |
| Altai_N_old | Russia_AfontovaGora3 + Ust_Kyakhta_14kya + China_NEastAsia_Inland_EN | 0.121 | Russia_AfontovaGora3 | 0.433 | 0.015 | 2.97E+01 |
| Altai_N_old | Russia_AfontovaGora3 + Ust_Kyakhta_14kya + China_NEastAsia_Inland_EN | 0.121 | Ust_Kyakhta_14kya | 0.348 | 0.040 | 8.72E+00 |
| Altai_N_old | Russia_AfontovaGora3 + Ust_Kyakhta_14kya + China_NEastAsia_Inland_EN | 0.121 | China_NEastAsia_Inland_EN | 0.219 | 0.033 | 6.53E+00 |
| Altai_N_old | Russia_AfontovaGora3 + Ust_Kyakhta_14kya + China_AmurRiver_Mesolithic | 0.065 | Russia_AfontovaGora3 | 0.431 | 0.015 | 2.90E+01 |
| Altai_N_old | Russia_AfontovaGora3 + Ust_Kyakhta_14kya + China_AmurRiver_Mesolithic | 0.065 | Ust_Kyakhta_14kya | 0.426 | 0.030 | 1.40E+01 |
| Altai_N_old | Russia_AfontovaGora3 + Ust_Kyakhta_14kya + China_AmurRiver_Mesolithic | 0.065 | China_AmurRiver_Mesolithic | 0.143 | 0.023 | 6.22E+00 |
| Altai_N_old | Russia_AfontovaGora3 + China_AmurRiver_Mesolithic + Cisbaikal_LNBA | 0.050 | Russia_AfontovaGora3 | 0.447 | 0.016 | 2.79E+01 |
| Altai_N_old | Russia_AfontovaGora3 + China_AmurRiver_Mesolithic + Cisbaikal_LNBA | 0.050 | China_AmurRiver_Mesolithic | 0.144 | 0.033 | 4.43E+00 |
| Altai_N_old | Russia_AfontovaGora3 + China_AmurRiver_Mesolithic + Cisbaikal_LNBA | 0.050 | Cisbaikal_LNBA | 0.409 | 0.043 | 9.45E+00 |

| | | | | | | |
|---|---|---|---|---|---|---|
| Transbaikal_EMN_old | China_NEastAsia_Inland_EN + KhatystyrCave_M_10.2kya | 0.083 | China_NEastAsia_Inland_EN | 0.446 | 0.163 | 2.74E+00 |
| Transbaikal_EMN_old | China_NEastAsia_Inland_EN + KhatystyrCave_M_10.2kya | 0.083 | KhatystyrCave_M_10.2kya | 0.554 | 0.163 | 3.41E+00 |

**Table 21 & 22 |** Passing models for the populations in Tranche 4, with *Cisbaikal_LNBA* in tranche 2. **References:** [Tranche 0 + Tranche 1 + Tranche 2 + Tranche 3 + Cisbaikal_LNBA (+ Tranche 4 only for Set B)]
**Sources:** [*Peru_Laramate_900BP, Russia_AfontovaGora3, Khaiyrgas, Ust_Kyakhta_14kya, China_AmurRiver_LPaleolithic_19K, China_AmurRiver_LPaleolithic_14K, China_AmurRiver_Mesolithic, China_NEastAsia_Coastal_EN, China_NEastAsia_Inland_EN, China_SEastAsia_Coastal_EN, KhatystyrCave_M_10.2kya, Kolyma_M_10.1kya, Cisbaikal_LNBA, (Dzhilinda1_M_N_8.4kya, Altai_N_old and Transbaikal_EMN included in set B)*]

##### VI.A.iii.d) Tranche 5: Altai_N, Transbaikal_EMN, China_AmurRiver_N, Cisbaikal_EN, Mongolia_N_North & Syalakh-Belkachi

***Tranche 5 with Cisbaikal_LNBA in the references.***

***Altai_N.*** As before, *Altai_N* can be modeled as descending completely from the older individuals in this cluster (*Altai_N_old*). The same model passes in set B.

***Transbaikal_EMN.*** A simple model of Transbaikal_EMN with complete descent from *Transbaikal_EMN_old* no longer passes in either set A or B. Instead, a wide variety of 2- and 3-way models pass where *Transbaikal_EMN* continues to draw almost all its ancestry from *Transbaikal_EMN_old*, suggesting low levels of gene flow from adjacent populations in Northeast Asia and Siberia into a population otherwise demonstrating a high level of continuity with the preceding population of the Transbaikal region.

***China_AmurRiver_N.*** As before, passing 2-way models feature near complete descent from *China_AmurRiver_EarlyN* or *China_AmurRiver_Mesolithic* with the addition of a small contribution from one other, typically Northeast Asian or APS source, testifying to genetic continuity but with low levels of gene flow in the Amur River region.

***Cisbaikal_EN.*** The prior 2-way admixture model of *Cisbaikal_EN* with ancestry from Transbaikal_EMN_old and one APS source (*Ust_Kyakhta_14kya* or *Khairygas_16.7kya*) no longer pass in set A, but a similar set of 3-way models pass in Set A that retain a major contribution from *Transbaikal_EMN_old* in similar proportions, plus minor contributions from two APS-related sources– one of which is always *Ust_Kyakhta_14kya* or *Cisbaikal_LNBA*. No models pass for Cisbaikal_EN in set B.

***Mongolia_N_North.*** The two-way model for *Mongolia_N_North* as an admixture between *Transbaikal_EMN_old* and *China_NEastAsia_Inland_EN* no longer passes; however, similar 3-way models with the small proportion of *ANE*-rich ancestry substituted by *Cisbaikal_LNBA* now pass in set A. No models pass in set B.

***Syalakh-Belkachi.*** In Set A, the model drawing the ancestry of *Syalakh-Belkachi* a single source, the Mesolithic Northern Buryatian *Dzhilinda1_M_N_8.4kya*, now fails. However, the 2-way models that passed in the previous set of *qpAdm* in set A, where *Syalakh-Belkachi* continues to demonstrate majority descent from *Dzhilinda1_M_N_8.4kya* with a minor contribution (~10-20%) from a broadly East-Asian-related population, continue to pass. In set B, the same three passing models as passed before, which contained a major contribution from *Dzhilinda1_M_N_8.4kya* (~76%) and a minor contribution from a population rich in Inland Northeast Asian ancestry (either *Cisbaikal_EN*, *Transbaikal_EMN_old* or *Mongolia_N_North*), continues to pass.

All models in this tranche can be found in Table 6C (Set A) and Table 6D (Set B) in Data S3.

#### Table S23

| Target | Model | P | Source | Prop. | SE | Z |
|---|---|---|---|---|---|---|
| Altai_N | Altai_N_old | 0.737 | Altai_N_old | 1.000 | 0.000 | 3.13E+13 |
| Transbaikal_EMN | China_NEastAsia_Coastal_EN + Transbaikal_EMN_old + Cisbaikal_LNBA | 0.445 | China_NEastAsia_Coastal_EN | 0.075 | 0.072 | 1.05E+00 |
| Transbaikal_EMN | China_NEastAsia_Coastal_EN + Transbaikal_EMN_old + Cisbaikal_LNBA | 0.445 | Transbaikal_EMN_old | 0.921 | 0.083 | 1.11E+01 |
| Transbaikal_EMN | China_NEastAsia_Coastal_EN + Transbaikal_EMN_old + Cisbaikal_LNBA | 0.445 | Cisbaikal_LNBA | 0.003 | 0.032 | 9.83E-02 |
| China_AmurRiver_N | China_AmurRiver_EarlyN + Transbaikal_EMN_old | 0.589 | China_AmurRiver_EarlyN | 0.824 | 0.040 | 2.04E+01 |
| China_AmurRiver_N | China_AmurRiver_EarlyN + Transbaikal_EMN_old | 0.589 | Transbaikal_EMN_old | 0.176 | 0.040 | 4.34E+00 |
| China_AmurRiver_N | China_AmurRiver_EarlyN + China_NEastAsia_Inland_EN | 0.296 | China_AmurRiver_EarlyN | 0.870 | 0.041 | 2.14E+01 |
| China_AmurRiver_N | China_AmurRiver_EarlyN + China_NEastAsia_Inland_EN | 0.296 | China_NEastAsia_Inland_EN | 0.130 | 0.041 | 3.20E+00 |
| China_AmurRiver_N | China_AmurRiver_EarlyN + Dzhilinda1_M_N_8.4kya | 0.193 | China_AmurRiver_EarlyN | 0.939 | 0.020 | 4.62E+01 |
| China_AmurRiver_N | China_AmurRiver_EarlyN + Dzhilinda1_M_N_8.4kya | 0.193 | Dzhilinda1_M_N_8.4kya | 0.061 | 0.020 | 3.02E+00 |
| China_AmurRiver_N | China_AmurRiver_Mesolithic + Transbaikal_EMN_old | 0.159 | China_AmurRiver_Mesolithic | 0.778 | 0.036 | 2.19E+01 |
| China_AmurRiver_N | China_AmurRiver_Mesolithic + Transbaikal_EMN_old | 0.159 | Transbaikal_EMN_old | 0.222 | 0.036 | 6.23E+00 |
| China_AmurRiver_N | China_AmurRiver_EarlyN + China_NEastAsia_Coastal_EN | 0.133 | China_AmurRiver_EarlyN | 0.894 | 0.049 | 1.81E+01 |
| China_AmurRiver_N | China_AmurRiver_EarlyN + China_NEastAsia_Coastal_EN | 0.133 | China_NEastAsia_Coastal_EN | 0.106 | 0.049 | 2.15E+00 |
| China_AmurRiver_N | China_AmurRiver_EarlyN + KhatystyrCave_M_10.2kya | 0.115 | China_AmurRiver_EarlyN | 0.869 | 0.063 | 1.38E+01 |
| China_AmurRiver_N | China_AmurRiver_EarlyN + KhatystyrCave_M_10.2kya | 0.115 | KhatystyrCave_M_10.2kya | 0.131 | 0.063 | 2.08E+00 |
| China_AmurRiver_N | Ust_Kyakhta_14kya + China_AmurRiver_EarlyN | 0.101 | Ust_Kyakhta_14kya | 0.051 | 0.017 | 2.97E+00 |
| China_AmurRiver_N | Ust_Kyakhta_14kya + China_AmurRiver_EarlyN | 0.101 | China_AmurRiver_EarlyN | 0.949 | 0.017 | 5.50E+01 |
| China_AmurRiver_N | China_AmurRiver_EarlyN + Cisbaikal_LNBA | 0.092 | China_AmurRiver_EarlyN | 0.945 | 0.020 | 4.63E+01 |
| China_AmurRiver_N | China_AmurRiver_EarlyN + Cisbaikal_LNBA | 0.092 | Cisbaikal_LNBA | 0.055 | 0.020 | 2.69E+00 |
| China_AmurRiver_N | Khaiyrgas + China_AmurRiver_EarlyN | 0.077 | Khaiyrgas | 0.035 | 0.014 | 2.54E+00 |
| China_AmurRiver_N | Khaiyrgas + China_AmurRiver_EarlyN | 0.077 | China_AmurRiver_EarlyN | 0.965 | 0.014 | 6.96E+01 |
| China_AmurRiver_N | China_AmurRiver_Mesolithic + China_NEastAsia_Inland_EN | 0.066 | China_AmurRiver_Mesolithic | 0.813 | 0.039 | 2.06E+01 |
| China_AmurRiver_N | China_AmurRiver_Mesolithic + China_NEastAsia_Inland_EN | 0.066 | China_NEastAsia_Inland_EN | 0.187 | 0.039 | 4.73E+00 |
| Cisbaikal_EN | Khaiyrgas + Transbaikal_EMN_old + Cisbaikal_LNBA | 0.151 | Khaiyrgas | 0.144 | 0.030 | 4.86E+00 |
| Cisbaikal_EN | Khaiyrgas + Transbaikal_EMN_old + Cisbaikal_LNBA | 0.151 | Transbaikal_EMN_old | 0.735 | 0.027 | 2.73E+01 |
| Cisbaikal_EN | Khaiyrgas + Transbaikal_EMN_old + Cisbaikal_LNBA | 0.151 | Cisbaikal_LNBA | 0.121 | 0.049 | 2.48E+00 |
| Cisbaikal_EN | Altai_N_old + Transbaikal_EMN_old + Cisbaikal_LNBA | 0.106 | Altai_N_old | 0.111 | 0.022 | 4.98E+00 |
| Cisbaikal_EN | Altai_N_old + Transbaikal_EMN_old + Cisbaikal_LNBA | 0.106 | Transbaikal_EMN_old | 0.766 | 0.031 | 2.50E+01 |
| Cisbaikal_EN | Altai_N_old + Transbaikal_EMN_old + Cisbaikal_LNBA | 0.106 | Cisbaikal_LNBA | 0.123 | 0.048 | 2.56E+00 |
| Cisbaikal_EN | Ust_Kyakhta_14kya + Kolyma_M_10.1kya + Transbaikal_EMN_old | 0.100 | Ust_Kyakhta_14kya | 0.273 | 0.024 | 1.15E+01 |
| Cisbaikal_EN | Ust_Kyakhta_14kya + Kolyma_M_10.1kya + Transbaikal_EMN_old | 0.100 | Kolyma_M_10.1kya | 0.057 | 0.023 | 2.49E+00 |
| Cisbaikal_EN | Ust_Kyakhta_14kya + Kolyma_M_10.1kya + Transbaikal_EMN_old | 0.100 | Transbaikal_EMN_old | 0.670 | 0.021 | 3.26E+01 |
| Cisbaikal_EN | Khaiyrgas + Ust_Kyakhta_14kya + Transbaikal_EMN_old | 0.074 | Khaiyrgas | 0.059 | 0.022 | 2.64E+00 |
| Cisbaikal_EN | Khaiyrgas + Ust_Kyakhta_14kya + Transbaikal_EMN_old | 0.074 | Ust_Kyakhta_14kya | 0.249 | 0.031 | 8.15E+00 |
| Cisbaikal_EN | Khaiyrgas + Ust_Kyakhta_14kya + Transbaikal_EMN_old | 0.074 | Transbaikal_EMN_old | 0.693 | 0.019 | 3.65E+01 |
| Cisbaikal_EN | Ust_Kyakhta_14kya + Transbaikal_EMN_old + Cisbaikal_LNBA | 0.074 | Ust_Kyakhta_14kya | 0.097 | 0.389 | 2.49E-01 |
| Cisbaikal_EN | Ust_Kyakhta_14kya + Transbaikal_EMN_old + Cisbaikal_LNBA | 0.074 | Transbaikal_EMN_old | 0.621 | 0.053 | 1.18E+01 |

| | | | | | | |
|---|---|---|---|---|---|---|
| Cisbaikal_EN | Ust_Kyakhta_14kya + Transbaikal_EMN_old + Cisbaikal_LNBA | 0.074 | Cisbaikal_LNBA | 0.282 | 0.437 | 6.46E-01 |
| Cisbaikal_EN | Altai_N_old + Ust_Kyakhta_14kya + Transbaikal_EMN_old | 0.073 | Altai_N_old | 0.050 | 0.017 | 2.98E+00 |
| Cisbaikal_EN | Altai_N_old + Ust_Kyakhta_14kya + Transbaikal_EMN_old | 0.073 | Ust_Kyakhta_14kya | 0.242 | 0.030 | 7.98E+00 |
| Cisbaikal_EN | Altai_N_old + Ust_Kyakhta_14kya + Transbaikal_EMN_old | 0.073 | Transbaikal_EMN_old | 0.708 | 0.020 | 3.50E+01 |
| Cisbaikal_EN | Ust_Kyakhta_14kya + China_NEastAsia_Inland_EN + Transbaikal_EMN_old | 0.054 | Ust_Kyakhta_14kya | 0.310 | 0.019 | 1.64E+01 |
| Cisbaikal_EN | Ust_Kyakhta_14kya + China_NEastAsia_Inland_EN + Transbaikal_EMN_old | 0.054 | China_NEastAsia_Inland_EN | 0.141 | 0.073 | 1.93E+00 |
| Cisbaikal_EN | Ust_Kyakhta_14kya + China_NEastAsia_Inland_EN + Transbaikal_EMN_old | 0.054 | Transbaikal_EMN_old | 0.549 | 0.073 | 7.49E+00 |
| *Mongolia_N_North* | China_NEastAsia_Inland_EN + Transbaikal_EMN_old + Cisbaikal_LNBA | 0.323 | China_NEastAsia_Inland_EN | 0.303 | 0.101 | 3.00E+00 |
| *Mongolia_N_North* | China_NEastAsia_Inland_EN + Transbaikal_EMN_old + Cisbaikal_LNBA | 0.323 | Transbaikal_EMN_old | 0.620 | 0.098 | 6.34E+00 |
| *Mongolia_N_North* | China_NEastAsia_Inland_EN + Transbaikal_EMN_old + Cisbaikal_LNBA | 0.323 | Cisbaikal_LNBA | 0.078 | 0.028 | 2.80E+00 |
| *Mongolia_N_North* | China_NEastAsia_Inland_EN + KhatystyrCave_M_10.2kya + Cisbaikal_LNBA | 0.055 | China_NEastAsia_Inland_EN | 0.619 | 0.137 | 4.51E+00 |
| *Mongolia_N_North* | China_NEastAsia_Inland_EN + KhatystyrCave_M_10.2kya + Cisbaikal_LNBA | 0.055 | KhatystyrCave_M_10.2kya | 0.311 | 0.124 | 2.52E+00 |
| *Mongolia_N_North* | China_NEastAsia_Inland_EN + KhatystyrCave_M_10.2kya + Cisbaikal_LNBA | 0.055 | Cisbaikal_LNBA | 0.069 | 0.040 | 1.71E+00 |
| Syalakh-Belkachi | Transbaikal_EMN_old + Dzhilinda1_M_N_8.4kya | 0.516 | Transbaikal_EMN_old | 0.213 | 0.051 | 4.17E+00 |
| Syalakh-Belkachi | Transbaikal_EMN_old + Dzhilinda1_M_N_8.4kya | 0.516 | Dzhilinda1_M_N_8.4kya | 0.787 | 0.051 | 1.55E+01 |
| Syalakh-Belkachi | China_AmurRiver_LPaleolithic_14K + Dzhilinda1_M_N_8.4kya | 0.157 | China_AmurRiver_LPaleolithic _14K | 0.091 | 0.037 | 2.43E+00 |
| Syalakh-Belkachi | China_AmurRiver_LPaleolithic_14K + Dzhilinda1_M_N_8.4kya | 0.157 | Dzhilinda1_M_N_8.4kya | 0.909 | 0.037 | 2.43E+01 |
| Syalakh-Belkachi | China_NEastAsia_Coastal_EN + Dzhilinda1_M_N_8.4kya | 0.139 | China_NEastAsia_Coastal_EN | 0.143 | 0.043 | 3.29E+00 |
| Syalakh-Belkachi | China_NEastAsia_Coastal_EN + Dzhilinda1_M_N_8.4kya | 0.139 | Dzhilinda1_M_N_8.4kya | 0.857 | 0.043 | 1.97E+01 |
| Syalakh-Belkachi | China_AmurRiver_Mesolithic + Dzhilinda1_M_N_8.4kya | 0.118 | China_AmurRiver_Mesolithic | 0.111 | 0.036 | 3.11E+00 |
| Syalakh-Belkachi | China_AmurRiver_Mesolithic + Dzhilinda1_M_N_8.4kya | 0.118 | Dzhilinda1_M_N_8.4kya | 0.889 | 0.036 | 2.48E+01 |
| Syalakh-Belkachi | KhatystyrCave_M_10.2kya + Dzhilinda1_M_N_8.4kya | 0.104 | KhatystyrCave_M_10.2kya | 0.143 | 0.062 | 2.30E+00 |
| Syalakh-Belkachi | KhatystyrCave_M_10.2kya + Dzhilinda1_M_N_8.4kya | 0.104 | Dzhilinda1_M_N_8.4kya | 0.857 | 0.062 | 1.38E+01 |
| Syalakh-Belkachi | China_AmurRiver_EarlyN + Dzhilinda1_M_N_8.4kya | 0.097 | China_AmurRiver_EarlyN | 0.103 | 0.036 | 2.84E+00 |
| Syalakh-Belkachi | China_AmurRiver_EarlyN + Dzhilinda1_M_N_8.4kya | 0.097 | Dzhilinda1_M_N_8.4kya | 0.897 | 0.036 | 2.47E+01 |
| Syalakh-Belkachi | China_NEastAsia_Inland_EN + Dzhilinda1_M_N_8.4kya | 0.068 | China_NEastAsia_Inland_EN | 0.152 | 0.061 | 2.50E+00 |
| Syalakh-Belkachi | China_NEastAsia_Inland_EN + Dzhilinda1_M_N_8.4kya | 0.068 | Dzhilinda1_M_N_8.4kya | 0.848 | 0.061 | 1.40E+01 |

#### Table S24

| Target | Model | P | Source | Prop. | SE | Z |
|---|---|---|---|---|---|---|
| Altai_N | Altai_N_old | 0.150 | Altai_N_old | 1.000 | 0.000 | 3.13E+13 |
| Transbaikal_EMN | Transbaikal_EMN_old | 0.066 | Transbaikal_EMN_old | 1.000 | 0.000 | 3.13E+13 |
| Transbaikal_EMN | Transbaikal_EMN_old + Cisbaikal_EN | 0.518 | Transbaikal_EMN_old | 0.992 | 0.081 | 1.23E+01 |
| Transbaikal_EMN | Transbaikal_EMN_old + Cisbaikal_EN | 0.518 | Cisbaikal_EN | 0.008 | 0.081 | 9.75E-02 |
| Transbaikal_EMN | China_NEastAsia_Inland_EN + Transbaikal_EMN_old | 0.481 | China_NEastAsia_Inland_EN | 0.222 | 0.066 | 3.37E+00 |
| Transbaikal_EMN | China_NEastAsia_Inland_EN + Transbaikal_EMN_old | 0.481 | Transbaikal_EMN_old | 0.778 | 0.066 | 1.18E+01 |
| Transbaikal_EMN | China_AmurRiver_LPaleolithic_19K + Transbaikal_EMN_old | 0.202 | China_AmurRiver_LPaleolithic _19K | 0.071 | 0.034 | 2.07E+00 |
| Transbaikal_EMN | China_AmurRiver_LPaleolithic_19K + Transbaikal_EMN_old | 0.202 | Transbaikal_EMN_old | 0.929 | 0.034 | 2.72E+01 |
| Transbaikal_EMN | Altai_N_old + Transbaikal_EMN_old | 0.180 | Altai_N_old | 0.016 | 0.012 | 1.34E+00 |

| | | | | | | |
|---|---|---|---|---|---|---|
| Transbaikal_EMN | Altai_N_old + Transbaikal_EMN_old | 0.180 | Transbaikal_EMN_old | 0.984 | 0.012 | 8.25E+01 |
| Transbaikal_EMN | China_NEastAsia_Coastal_EN + Transbaikal_EMN_old | 0.159 | China_NEastAsia_Coastal_EN | 0.080 | 0.036 | 2.23E+00 |
| Transbaikal_EMN | China_NEastAsia_Coastal_EN + Transbaikal_EMN_old | 0.159 | Transbaikal_EMN_old | 0.920 | 0.036 | 2.56E+01 |
| Transbaikal_EMN | China_SEastAsia_Coastal_EN + Transbaikal_EMN_old | 0.113 | China_SEastAsia_Coastal_EN | 0.044 | 0.024 | 1.82E+00 |
| Transbaikal_EMN | China_SEastAsia_Coastal_EN + Transbaikal_EMN_old | 0.113 | Transbaikal_EMN_old | 0.956 | 0.024 | 3.91E+01 |
| Transbaikal_EMN | Transbaikal_EMN_old + Altai_N | 0.091 | Transbaikal_EMN_old | 0.999 | 0.013 | 7.96E+01 |
| Transbaikal_EMN | Transbaikal_EMN_old + Altai_N | 0.091 | Altai_N | 0.001 | 0.013 | 6.98E-02 |
| Transbaikal_EMN | Kolyma_M_10.1kya + Transbaikal_EMN_old | 0.084 | Kolyma_M_10.1kya | 0.023 | 0.021 | 1.11E+00 |
| Transbaikal_EMN | Kolyma_M_10.1kya + Transbaikal_EMN_old | 0.084 | Transbaikal_EMN_old | 0.977 | 0.021 | 4.68E+01 |
| Transbaikal_EMN | China_AmurRiver_Mesolithic + Transbaikal_EMN_old | 0.084 | China_AmurRiver_Mesolithic | 0.043 | 0.030 | 1.43E+00 |
| Transbaikal_EMN | China_AmurRiver_Mesolithic + Transbaikal_EMN_old | 0.084 | Transbaikal_EMN_old | 0.957 | 0.030 | 3.17E+01 |
| Transbaikal_EMN | Russia_AfontovaGora3 + Transbaikal_EMN_old | 0.069 | Russia_AfontovaGora3 | 0.006 | 0.007 | 8.14E-01 |
| Transbaikal_EMN | Russia_AfontovaGora3 + Transbaikal_EMN_old | 0.069 | Transbaikal_EMN_old | 0.994 | 0.007 | 1.36E+02 |
| Transbaikal_EMN | Ust_Kyakhta_14kya + Transbaikal_EMN_old | 0.065 | Ust_Kyakhta_14kya | 0.022 | 0.027 | 8.27E-01 |
| Transbaikal_EMN | Ust_Kyakhta_14kya + Transbaikal_EMN_old | 0.065 | Transbaikal_EMN_old | 0.978 | 0.027 | 3.66E+01 |
| Transbaikal_EMN | Khaiyrgas + Transbaikal_EMN_old | 0.064 | Khaiyrgas | 0.011 | 0.017 | 6.68E-01 |
| Transbaikal_EMN | Khaiyrgas + Transbaikal_EMN_old | 0.064 | Transbaikal_EMN_old | 0.989 | 0.017 | 5.94E+01 |
| Transbaikal_EMN | Transbaikal_EMN_old + Dzhilinda1_M_N_8.4kya | 0.060 | Transbaikal_EMN_old | 0.979 | 0.026 | 3.74E+01 |
| Transbaikal_EMN | Transbaikal_EMN_old + Dzhilinda1_M_N_8.4kya | 0.060 | Dzhilinda1_M_N_8.4kya | 0.021 | 0.026 | 8.03E-01 |
| Transbaikal_EMN | China_AmurRiver_EarlyN + Transbaikal_EMN_old | 0.056 | China_AmurRiver_EarlyN | 0.034 | 0.034 | 1.01E+00 |
| Transbaikal_EMN | China_AmurRiver_EarlyN + Transbaikal_EMN_old | 0.056 | Transbaikal_EMN_old | 0.966 | 0.034 | 2.88E+01 |
| Transbaikal_EMN | Transbaikal_EMN_old + China_AmurRiver_N | 0.055 | Transbaikal_EMN_old | 0.971 | 0.035 | 2.77E+01 |
| Transbaikal_EMN | Transbaikal_EMN_old + China_AmurRiver_N | 0.055 | China_AmurRiver_N | 0.029 | 0.035 | 8.40E-01 |
| China_AmurRiver_N | China_AmurRiver_EarlyN + China_NEastAsia_Inland_EN | 0.137 | China_AmurRiver_EarlyN | 0.921 | 0.037 | 2.51E+01 |
| China_AmurRiver_N | China_AmurRiver_EarlyN + China_NEastAsia_Inland_EN | 0.137 | China_NEastAsia_Inland_EN | 0.079 | 0.037 | 2.17E+00 |
| China_AmurRiver_N | China_AmurRiver_EarlyN + Transbaikal_EMN_old | 0.113 | China_AmurRiver_EarlyN | 0.921 | 0.031 | 2.97E+01 |
| China_AmurRiver_N | China_AmurRiver_EarlyN + Transbaikal_EMN_old | 0.113 | Transbaikal_EMN_old | 0.079 | 0.031 | 2.57E+00 |
| China_AmurRiver_N | China_AmurRiver_EarlyN + Dzhilinda1_M_N_8.4kya | 0.108 | China_AmurRiver_EarlyN | 0.957 | 0.020 | 4.81E+01 |
| China_AmurRiver_N | China_AmurRiver_EarlyN + Dzhilinda1_M_N_8.4kya | 0.108 | Dzhilinda1_M_N_8.4kya | 0.043 | 0.020 | 2.17E+00 |
| China_AmurRiver_N | China_AmurRiver_EarlyN + China_NEastAsia_Coastal_EN | 0.106 | China_AmurRiver_EarlyN | 0.893 | 0.049 | 1.81E+01 |
| China_AmurRiver_N | China_AmurRiver_EarlyN + China_NEastAsia_Coastal_EN | 0.106 | China_NEastAsia_Coastal_EN | 0.107 | 0.049 | 2.16E+00 |
| China_AmurRiver_N | China_AmurRiver_EarlyN + Transbaikal_EMN | 0.090 | China_AmurRiver_EarlyN | 0.920 | 0.031 | 2.94E+01 |
| China_AmurRiver_N | China_AmurRiver_EarlyN + Transbaikal_EMN | 0.090 | Transbaikal_EMN | 0.080 | 0.031 | 2.54E+00 |
| China_AmurRiver_N | China_AmurRiver_EarlyN + *Mongolia_N_North* | 0.083 | China_AmurRiver_EarlyN | 0.928 | 0.027 | 3.38E+01 |
| China_AmurRiver_N | China_AmurRiver_EarlyN + *Mongolia_N_North* | 0.083 | *Mongolia_N_North* | 0.072 | 0.027 | 2.64E+00 |
| China_AmurRiver_N | China_AmurRiver_EarlyN + Syalakh-Belkachi | 0.082 | China_AmurRiver_EarlyN | 0.939 | 0.023 | 4.06E+01 |
| China_AmurRiver_N | China_AmurRiver_EarlyN + Syalakh-Belkachi | 0.082 | Syalakh-Belkachi | 0.061 | 0.023 | 2.62E+00 |
| China_AmurRiver_N | China_AmurRiver_EarlyN + Altai_N | 0.080 | China_AmurRiver_EarlyN | 0.976 | 0.010 | 9.53E+01 |
| China_AmurRiver_N | China_AmurRiver_EarlyN + Altai_N | 0.080 | Altai_N | 0.024 | 0.010 | 2.34E+00 |
| China_AmurRiver_N | China_AmurRiver_EarlyN + Cisbaikal_EN | 0.072 | China_AmurRiver_EarlyN | 0.923 | 0.028 | 3.25E+01 |
| China_AmurRiver_N | China_AmurRiver_EarlyN + Cisbaikal_EN | 0.072 | Cisbaikal_EN | 0.077 | 0.028 | 2.72E+00 |
| China_AmurRiver_N | China_AmurRiver_Mesolithic + Transbaikal_EMN | 0.061 | China_AmurRiver_Mesolithic | 0.832 | 0.028 | 2.94E+01 |
| China_AmurRiver_N | China_AmurRiver_Mesolithic + Transbaikal_EMN | 0.061 | Transbaikal_EMN | 0.168 | 0.028 | 5.94E+00 |
| China_AmurRiver_N | Khaiyrgas + China_AmurRiver_EarlyN | 0.054 | Khaiyrgas | 0.029 | 0.014 | 2.17E+00 |
| China_AmurRiver_N | Khaiyrgas + China_AmurRiver_EarlyN | 0.054 | China_AmurRiver_EarlyN | 0.971 | 0.014 | 7.16E+01 |

| | | | | | | |
|---|---|---|---|---|---|---|
| Syalakh-Belkachi | Dzhilinda1_M_N_8.4kya + Cisbaikal_EN | 0.209 | Dzhilinda1_M_N_8.4kya | 0.762 | 0.051 | 1.50E+01 |
| Syalakh-Belkachi | Dzhilinda1_M_N_8.4kya + Cisbaikal_EN | 0.209 | Cisbaikal_EN | 0.238 | 0.051 | 4.70E+00 |
| Syalakh-Belkachi | Dzhilinda1_M_N_8.4kya + *Mongolia_N_North* | 0.193 | Dzhilinda1_M_N_8.4kya | 0.757 | 0.037 | 2.02E+01 |
| Syalakh-Belkachi | Dzhilinda1_M_N_8.4kya + *Mongolia_N_North* | 0.193 | *Mongolia_N_North* | 0.243 | 0.037 | 6.49E+00 |
| Syalakh-Belkachi | Transbaikal_EMN_old + Dzhilinda1_M_N_8.4kya | 0.168 | Transbaikal_EMN_old | 0.240 | 0.041 | 5.87E+00 |
| Syalakh-Belkachi | Transbaikal_EMN_old + Dzhilinda1_M_N_8.4kya | 0.168 | Dzhilinda1_M_N_8.4kya | 0.760 | 0.041 | 1.86E+01 |

**Table S1-VI. A-15 & VI. A-16 |** Passing models for the populations in Tranche 5 with the addition of *Cisbaikal_LNBA* as a new member of this tranche, as well as *Transbaikal_EMN_old* in tranche 3, *Altai_N_old* in tranche 4. No simpler models pass. **References:** [Tranche 0 + Tranche 1 + Tranche 2 + Tranche 3 + Tranche 4 + Cisbaikal_LNBA (+ Tranche 5 in set B)]
**Sources:** [*Peru_Laramate_900BP, AG3, Khairygas_16.7kya, Ust_Kyakhta_14kya, China_NEastAsia_Coastal_EN, China_NEastAsia_Inland_EN, China_SEastAsia_Coastal_EN, China_AmurRiver_Paleolithic_19K, China_AmurRiver_LPaleolithic_14K, China_AmurRiver_Mesolithic, China_AmurRiver_EarlyN, KhatystyrCave_M_10.2kya, Kolyma_M_10.1kya, Cisbaikal_LNBA, Transbaikal_EMN_old, Dzhilinda1_M_N_8.4kya, Altai_N_old, (Syalakh_Belkachi, Transbaikal_EMN, Mongolia_N_North, China_AmurRiver_N, Cisbaikal_EN in set B)*]

##### VI.A.iii.e) Tranche 6: Yakutia_LNBA

Because Cisbaikal_LNBA was already in the references in set B of the previous set of *qpAdm* for Yakutia_LNBA in Table VI. A-11, we do not repeat the set of models here.

#### VI.A.iv. Summary of Population Relationships in Siberia

##### VI.A.iv.a) Investigating the genetic peculiarities of Cisbaikal_LNBA

Because *Cisbaikal_LNBA* has to be moved to a much older tranche, we suggest that either 1) *Cisbaikal_LNBA* contains ancestry from sources that contributed significantly to populations much older than itself, such that these populations cannot be placed in the references without reducing the p-value for models for *Cisbaikal_LNBA*, or 2) *Cisbaikal_LNBA* contains unusual ancestry. The fact that individuals from *Cisbaikal_LNBA* also form their own cluster in PCA (Extended Data Fig. 6) also motivated us to investigate the ancestry of *Cisbaikal_LNBA* more deeply.

First, we note that while *Cisbaikal_EN* can be modeled as having a large amount of ancestry from *Transbaikal_EMN*-related populations, *Cisbaikal_LNBA* cannot be modeled in this way and requires a direct contribution from Inland Northeast Asians (*China_NEastAsia_Inland_EN*). Indeed, a comparison of f4-statistics (Main text Figure 2B) indicate that *Cisbaikal_LNBA* could not have gained its level of shared drift with Inland Northeast Asians through admixture between *Cisbaikal_EN* and *Ust_Kyakhta_14kya*. We also note that the movement of *Cisbaikal_LNBA* into Tranche 3–and therefore into the references for *qpAdm* targeting later populations from Tranches 4 and 5–caused either *Ust_Kyakhta_14kya*, or the closely related *Cisbaikal_LNBA* itself (~85% *Ust_Kyakhta_14kya* + ~15% *China_NEastAsia_Inland_EN*, Tables VI. A-14 and 15), to be required as a source in passing models for many populations in Tranches 4 and 5, where they were not required before. Specifically, we find that the *Ust_Kyakhta_14kya*-related source, or *Cisbaikal_LNBA* itself, now appears in models for many populations from the early Holocene (~8.5-7kya) from regions that surround the Cisbaikal region, such as *Dzhilinda1_M_N_8.4kya* (~1-28%, ~8.4kya) from Southern Yakutia, *Altai_N_old* (~22-58%, ~8.2kya) from the Altai mountains, *Mongolia_N_North* (~7%, ~7kya) from the Mongolian Plateau, or the preceding population of the Cisbaikal region, *Cisbaikal_EN* (~12-37%, ~8-7kya). This may suggest that *Cisbaikal_LNBA* was especially related to an unsampled *Ust_Kyakhta_14kya*-related source that contributed to numerous early Holocene populations from areas adjacent to the the Cisbaikal region, and that it is this connection causes the models targeting *Cisbaikal_LNBA* to fail when it is placed in Tranches later than Tranche 3.

The oldest individuals in the *Cisbaikal_LNBA* cluster dates to ~4.9kya, which is much more recent than those populations in Tranche 4 and 5 (dating to ~8-7kya) that may have been affected by this unsampled *Cisbaikal_LNBA/Ust_Kyakhta_14kya*-like source. These early Holocene populations in Tranches 4 and 5 are also temporally much later than the *Ust_Kyakhta_14kya* individual itself, who was recovered from just south of Lake Baikal ~14kya. We suggest that an APS population similar to *Ust_Kyakhta_14kya* may have persisted from ~14kya down to ~8kya northwest of the regions of Siberia from which our samples indicate persistence of APS ancestry. For example, such a population may have persisted in the Taiga region north of the Krasnoyarsk region–from which we do not have samples from the early Holocene–and maintained gene flow with numerous populations around the Cisbaikal region throughout this period, admixing with Inland Northeast Asian ancestry to form the *Cisbaikal_LNBA* population before ~4.9kya, prior to the appearance of *Cisbaikal_LNBA* individuals in the Baikal region at that date.

##### VI.A.iv.b) The Khaiyrgas_16.7kya is a near-unadmixed representative of an "Ancient Paleosiberian" (APS) lineage that retains broad affinities to Native Americans

The oldest individual among our targets, *Khairygas_16.7kya* (~16,700 BP), from far Southwestern Yakutia on the middle reaches of the Lena River, has previously been defined as part of an "Ancient Paleosiberian" (APS) lineage related closely related to Native Americans [16,41]**.** Our models corroborate its close affinity to Native Americans (*Khairygas_16.7kya* = 100% *Peru_Laramate_900BP*, p = 0.092; *Khairygas_16.7kya* = ~93% *Peru_Laramate_900BP* + ~7% *AG3*, p = 0.293, Table VI. A-2). This is also supported by the fact that a model for *Peru_Laramate_900BP* with complete descent from *Khairygas_16.7kya* marginally passes in Tranche 1 when the positions of *Peru_Laramate_900BP* in the references and *Khairygas_16.7kya* in the targets are reversed (Table VI. A-21). Indeed, all passing models for Native Americans as a target in Tranche 1 have them derive almost all their ancestry from a *Khairygas_16.7kya*-related population (Table VI. A-21; Data S3 Table 2). However, the slight East Asian admixture (~6-9%) required in passing models with high P-values, when interpreted together with the slight *ANE* admixture in 2-way models for *Khairygas_16.7kya* when Native Americans are the other source, may indicate that *Khairygas_16.7kya.SG* has slightly more *ANE* ancestry than Native American populations, a fact supported by f4-statistics (F4(*Khaiyrgas, Peru_Laramate_900BP; Russia_MA1_HG.SG, China_AmurRiver_LPaleolithic_19K*) = 0.00211, Z = 2.081). All *qpAdm*s for Peru_Laramate_900BP as a member of Tranche 1 can be found in Table 3 of Data S3.

**Table S25**

| Target | Model | P | Source | Proportion | SE | Z |
|---|---|---|---|---|---|---|
| Peru_Laramate_900BP | Khaiyrgas | 0.068 | Khaiyrgas | 1.000 | 0.000 | 3.13E+13 |
| Peru_Laramate_900BP | Khaiyrgas + China_NEastAsia_Coastal_EN | 0.172 | Khaiyrgas | 0.926 | 0.044 | 2.10E+01 |
| Peru_Laramate_900BP | Khaiyrgas + China_NEastAsia_Coastal_EN | 0.172 | China_NEastAsia_Coastal_EN | 0.074 | 0.044 | 1.69E+00 |
| Peru_Laramate_900BP | Khaiyrgas + KhatystyrCave_M_10.2kya | 0.106 | Khaiyrgas | 0.941 | 0.053 | 1.79E+01 |
| Peru_Laramate_900BP | Khaiyrgas + KhatystyrCave_M_10.2kya | 0.106 | KhatystyrCave_M_10.2kya | 0.059 | 0.053 | 1.13E+00 |
| Peru_Laramate_900BP | Khaiyrgas + China_NEastAsia_Inland_EN | 0.097 | Khaiyrgas | 0.911 | 0.046 | 1.99E+01 |
| Peru_Laramate_900BP | Khaiyrgas + China_NEastAsia_Inland_EN | 0.097 | China_NEastAsia_Inland_EN | 0.089 | 0.046 | 1.95E+00 |
| Peru_Laramate_900BP | Khaiyrgas + China_AmurRiver_LPaleolithic_19K | 0.069 | Khaiyrgas | 0.930 | 0.042 | 2.23E+01 |
| Peru_Laramate_900BP | Khaiyrgas + China_AmurRiver_LPaleolithic_19K | 0.069 | China_AmurRiver_LPaleolithic_19K | 0.070 | 0.042 | 1.69E+00 |

| | | | | | | |
|---|---|---|---|---|---|---|
| Peru_Laramate_900BP | Khaiyrgas + China_SEastAsia_Coastal_EN | 0.065 | Khaiyrgas | 0.942 | 0.048 | 1.98E+01 |
| Peru_Laramate_900BP | Khaiyrgas + China_SEastAsia_Coastal_EN | 0.065 | China_SEastAsia_Coastal_EN | 0.058 | 0.048 | 1.21E+00 |

**Table S1-VI. A-21 |** Passing models for Peru_Laramate_900BP modeled with references/sources used for tranche 1.
**References:** [Tranche 0]
**Sources:** [Japan_HG_Jomon, Russia_AfontovaGora3, Khaiyrgas, China_SEastAsia_Coastal_EN, China_AmurRiver_LPaleolithic_19K, China_NEastAsia_Coastal_EN, China_NEastAsia_Inland_EN, KhatystyrCave_M_10.2kya]

##### VI.A.iv.c) Ancient Paleosiberian (APS) ancestry persists in Ust_Kyakhta_14kya and Kolyma_M_10.1kya, but with East Asian admixture and possible Native American backflow

Other individuals having high levels of APS-related ancestry include *Ust_Kyakhta_14kya* from ~14,000 years BP from just South of Lake Baikal (~56-59% *Khairygas_16.7kya-* or *Peru_Laramate_900BP*-related ancestry among 5 passing models when the remainder is from an East Asian source, Table VI. A-3), and to *Kolyma_M_10.1kya* from ~10,000 years BP from far Northeastern Siberia in the Kolyma River Valley (~65-73% *Khairygas_16.7kya-* or *Peru_Laramate_900BP*-related ancestry when the remainder is from an East Asian source, Table VI. A-5). The rest of their ancestry is broadly East Asian and can be drawn from the Mesolithic Southern Yakutian *KhatystyrCave_M_10.2kya*, the Inland Northeast Asian population represented by the *Yumin* individual from Inner Mongolia (*China_NEastAsia_Inland_EN* from ~8,400 years BP), or a population from the Amur Basin. The identity of the East Asian source among *Ust_Kyakhta_14kya* and *Kolyma_M_10.1kya* is difficult to identify with precision from currently available populations, and may await further sampling.

Unlike *Khairygas_16.7kya*, *Kolyma_M_10.1kya* and *Ust_Kyakhta_14kya* cannot be modeled as forming a clade with Native Americans in our *qpAdm* models, and require substantial admixture from an East Asian source. However, these individuals share equal or more drift with Native Americans as *Khairygas_16.7kya* in $f_4$-statistics (E.g. Main text Fig. 2B). These results are consistent with previous analyses that directly compared the relative affinities of *Ust_Kyakhta_14kya*/*Kolyma_M_10.1kya* and *Khairygas_16.7kya* with Native Americans, where *Kolyma_M_10.1kya* followed by *Ust_Kyakhta_14kya* exceed Khairygas_16.7kya in their affinity with Native Americans (for example, Z=2.1 for $f_4$(*Yoruba, Anzick-1.SG, Khairygas_16.7kya, Ust_Kyakhta_14kya)*, [16]; see also [51]). The aforementioned $f_4$-statistics may indicate that *Kolyma_M_10.1kya* and *Ust_Kyakhta_14kya* may derive their Native-American-related ancestry from backflow from the Americas that increased the Native American affinity of the two later individuals despite their mixed ancestry. However, a test for asymmetry between Northern Native Americans (NNA) and Southern Native Americans (SNA; [27]) reveals that *Khaiyrgas*, *Ust_Kyakhta_14kya* and *Kolyma_M_10.1kya* are not more related to NNA than SNA populations ($f_4$(*Ethiopia_4500BP.SG, Khaiyrgas/Ust_Kyakhta_14kya/Kolyma_M_10.1kya; NNA, SNA*) approximately 0; SI Data S3 Table 9), a result that is unexpected if their affinity to Native Americans was increased by movements from the New World. We suggest that APS populations were structured, and that *Ust_Kyakhta_14kya* and *Kolyma_M_10.1kya* may simply derive their APS ancestry from an APS population that was even more closely related to Native Americans than the population represented by *Khairygas_16.7kya*.

##### VI.A.iv.d) Ancient Paleosiberian (APS) ancestry persisted into late Holocene populations of Northeastern Eurasia and the Americas along two routes.

Our *qpAdm* analyses indicate that APS ancestry left a legacy in later populations in Northeastern Siberia through two routes.

- The first route involves a population closely related to *Ust_Kyakhta_14kya* and *Cisbaikal_LNBA*. The best models for *Cisbaikal_LNBA* demonstrate a very high level of continuity from an *Ust_Kyakhta_14kya*-related population under all *qpAdm* setups (~86% *Ust_Kyakhta_14kya* in Table VI. A-16; see also Tables VI. A-14 and 15), with the remainder drawn from a *China_NEastAsia_Inland_EN*-related population. An *Ust_Kyakhta_14kya*-related population closely related to the source that contributed to *Cisbaikal_LNBA*, or *Cisbaikal_LNBA* itself, also contributed smaller fractions of ancestry to early Holocene (~8.5-7kya) populations from areas adjacent to the Cisbaikal region, such *Dzhilinda1_M_N_8.4kya* (~1-28%, ~8.4kya), *Altai_N_old* (~22-58%, ~8.2kya), *Cisbaikal_EN* (~12-37%, ~8-7kya), and *Mongolia_N_North* (~7%, ~7kya); all passing models for these populations require either admixture from *Ust_Kyakhta_14kya* or *Cisbaikal_LNBA* once *Cisbaikal_LNBA* is moved into Tranche 3. As covered in later sections, populations rich in *Cisbaikal_LNBA* ancestry appear along the Middle Angara and Upper Yenisei Rivers in the Late Bronze and Iron Ages (Section VI. D. ii.) and *Cisbaikal_LNBA* appears to have left a genetic legacy among present-day Yeniseians and other present-day populations who admixed with historically Yeniseian-speaking groups (Section VI.D.iii.). We also find suggestive evidence that a *Cisbaikal_LNBA* or *Ust_Kyakhta_14kya*-related lineage contributed a small quantity of ancestry to ancient Athabaskans (Section VI.B.i).

Related to the events above, an admixture event between APS ancestry (most of which is related to the *Ust_Kyakhta_14kya* or *Cisbaikal_LNBA*-like source) and *ANE* ancestry produced an *ANE*-rich population (*Altai_N*) that survived in the Altai mountains down to the Eneolithic period (Section VI.D.2). The admixture event involves a contribution from an *AG3*-related source and ancestry from two APS-rich ancestry sources, at least one of which is derived from an *Ust_Kyakhta_14kya*- or *Cisbaikal_LNBA*-related population that mediates most of the APS-related ancestry (Table VI. A-17, VI. A-18). This admixed population (*Altai_N_old*) appeared at the latest in the Altai region by ~8,200 years BP, and later populations from the same region (*Altai_N*) down to ~6,200 years BP demonstrate complete genetic continuity with their predecessors. This group contributed at low levels to populations of the Steppe and Forest-Steppe in the Bronze and Iron Ages down to present-day populations of the same regions (Section VI.C.3).

- The second route involves a sequence of populations that emerged in Northern Buryatia and Yakutia proper, in the Kolyma and Middle-to-Lower Lena River basins in Northeastern Siberia (*Kolyma_M_10.1kya, Dzhilinda1_M_N_8.4kya, Syalakh-Belkachi*, and *Yakutia_LNBA*). Populations in this route (but not the APS-rich populations in the previously mentioned routes) show special affinity to ancient and present-day groups on either side of the Bering Straits, in Arctic North America, and parts of Siberia (but not the main Native American lineage that is the sole ancestry in non-Na-Dene-speaking Native Americans and is represented in these analyzes by *Peru_Laramate_900BP*). Ancestry from populations in this route may be responsible for the genetic affinities that these present-day groups share with each other.

  The second population in this sequence, the Mesolithic Northern Buryatian *Dzhilinda1_M_N_8.4kya* from ~8,400 years BP, derives much of its ancestry from either *Khairygas_16.7kya* or *Kolyma.M.SG* (from the Kolyma River Valley close to the Bering Strait from ~10kya)*;* the simplest 2-way models with *Khairygas_16.7kya* sees *Dzhilinda1_M_N_8.4kya* drawing ~60% of its ancestry from this source (Table VI. A-17 and 18), though more complex models with multiple sources of APS ancestry, one of which is always *Kolyma_M_10.1kya*, also pass. *Dzhilinda1_M_N_8.4kya* is an important contributor in turn to the third and fourth groups that appear in this sequence, which are increasingly admixed with East Asian sources: *Syalakh-Belkachi*, which derives most of its ancestry from a group related to *Dzhilinda1_M_N_8.4kya* (~75%-90%, Tables VI. A-19 and 20), and in turn to *Yakutia_LNBA* through *Syalakh-Belkachi* (~50% *Syalakh-Belkach* + ~50% *Transbaikal_EMN*, Table VI. A-10).

Kilinc et. al. [16] and Yu et. al. [51] find that *Kolyma_M_10.1kya* is more closely related to ancient and present-day Beringian Arctic and far Northeastern Siberian populations when compared with *Ust_Kyakhta_14kya* and *Khairygas_16.7kya*. We find that, when compared to other groups rich in APS ancestry (such as *Khairygas_16.7kya*, *Ust_Kyakhta_14kya*, *Cisbaikal_LNBA* and many groups along the *FSHG* cline), the other three populations in the second route (*Dzhilinda1_M_N_8.4kya, Syalakh-Belkachi* and *Yakutia_LNBA*) also consistently exhibit increased affinity for later populations from Beringia, far Northeastern Siberia, and the North American Arctic, such as the *Saqqaq.SG* Paleo-Eskimo from Greenland ~3800BP, the *OldBeringSea_Ekven* and *OldBeringSea_Uelen* Neo-Eskimo populations from the Asian shore of the Bering Straits ~2000BP, the *Magadan_BA* population from the Tokarev culture of the Northern shore of the Okhotsk Sea ~3000BP, Paleo- and Neo-Aleuts from the Aleutian Islands from ~2000-200BP, an Iron Age Yakutian from the Lena River Valley from ~2600BP, and ancient Athabaskans from inland Alaska from ~1100BP (Fig. S77) Indeed, present-day populations from Beringia, Arctic North America, and far Northeastern Siberia consistently minimize statistics of the form F4(O, Target, *Route2, Route1*) where *Route2* is an APS-rich population from the second route (*Kolyma_M_10.1kya*, *Dzhilinda1_M_N_8.4kya*, *Syalakh-Belkachi* and *Yakutia_LNBA*), and *Route1* are *Ust_Kyakhta_14kya* or *Cisbaikal_LNBA* (Fig. S6); the populations that maximize this statistic include Eskimo-Aleut groups, Yukaghiric groups, Chukotko-Kamchatkans, and Samoyeds, and to a lesser extent Tungusic populations from the shores of the Okhotsk Sea and Northeastern Siberia. Such statistics indicate that the populations in the second route are distinguished from other APS groups by a special affinity to populations along the Bering Straits and adjacent regions. The statistic is not confounded by the ratio of West to East Eurasian ancestry, or relative affinity to Native American populations, because the pattern persists when Route1 is *Dzhilinda1_M_N_8.4kya* and Route2 is *Ust_Kyakhta_14kya*, or when Route2 is *Syalakh-Belkachi* and Route1 is *Cisbaikal_LNBA*; these population pairs have similar levels of West Eurasian ancestry and similar levels of affinity to Native American populations (Fig. S76).

*qpAdm* in later sections indicate that this affinity can be explained by how subsequent populations of the Bering Straits and surrounding regions (that may have been ancestral to present-day Chukotko-Kamchatkan-, Eskimo-Aleut-, and Yukaghiric-speaking populations) derive some of or all their APS ancestry from populations in the second route (SI Section VI.B). Additionally, we present the novel finding that *Yakutia_LNBA* plays an important role in the genomic formation of Uralic-speaking populations in general (i.e. not just Samoyeds) and is the primary or even the only source for the East Asian ancestry of Uralic populations as far west as the Baltic Sea; this will be investigated in detail in a later section (Sections VI. C. i., ii. and iii.). We conclude that populations in the second route may have transmitted APS ancestry into a wide range of present-day populations in Beringia, Arctic North America, and Northeastern Siberia, and through *Yakutia_LNBA* to Uralic populations stretching westward into Europe.

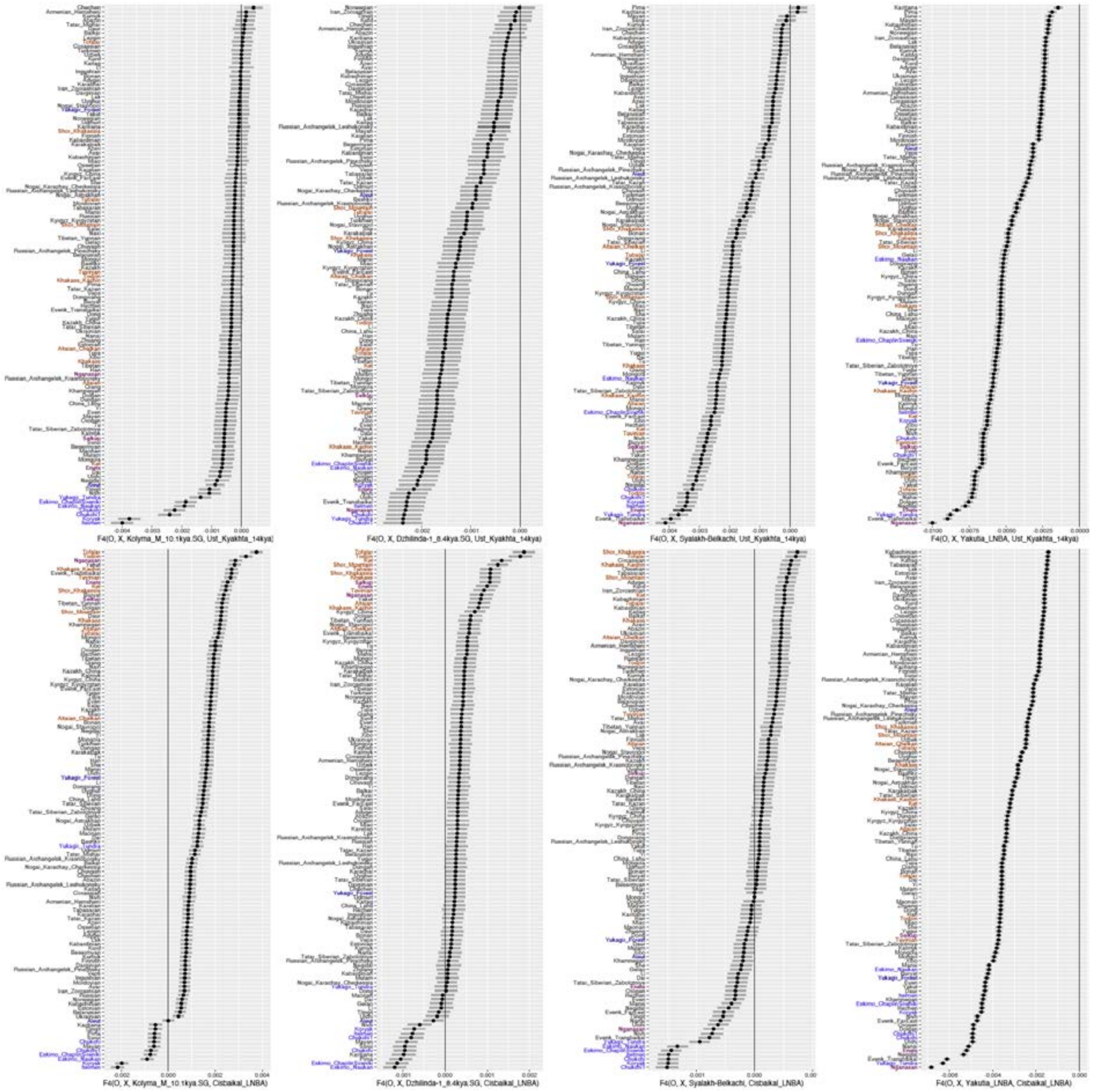

**Figure S76| Statistics of the form f4(Ethiopia_4500BP.SG, Target, Kolyma_M_10.1kya/Dzhilinda1_M_N_8.4kya, Ust_Kyakhta_14kya/Cisbaikal_LNBA).** Central Siberian populations from the Yenisei Basin (including Kets and South Siberian Turks) are highlighted in brown, while Arctic North American and Asian populations on either side of the Bering Straits populations are highlighted in blue. Despite the similarity of the APS-rich populations in this comparison (all being admixtures between APS ancestry and East Asian ancestry), present-day groups of the Bering Straits are generally closer to groups with "Route 2" APS ancestry (i.e., Kolyma_M_10.1kya → Dzhilinda1_8.4kya → Syalakh-Belkachi → Yakutia_LNBA), while Central Siberian populations of the Yenisei Basin are always closer to Cisbaikal_LNBA and Ust_Kyakhta_14kya. Bering Straits populations that are heavily European-admixed (Aleut and Yukagir_forest) are colored dark blue, while Samoyedic populations (Enets, Selkup, and Nganasan) are colored violet.

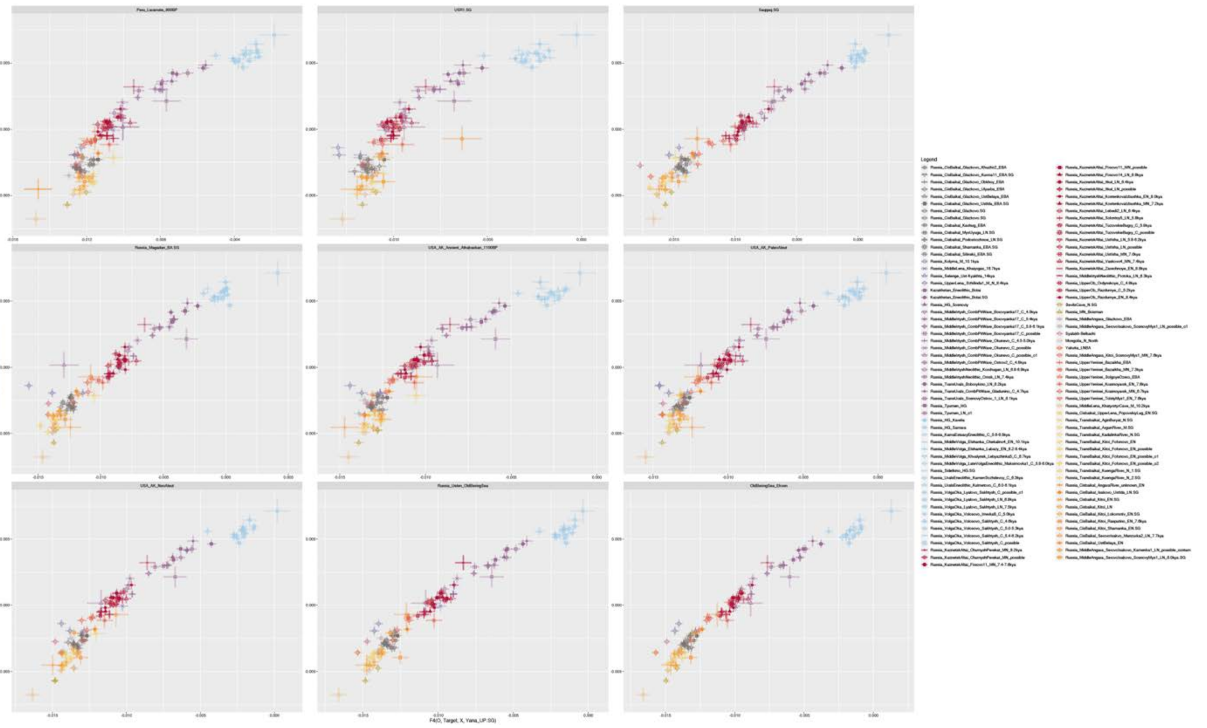

**Figure S77** | Statistics of the form f4(Ethiopia_4500BP.SG, Target, China_Paleolithic, Yana_UP.SG) on the y-axis, plotted against f4(Ethiopia_4500BP.SG, Target, X, Yana_UP.SG), where X are ancient Native Americans (USR1.SG, Peru_Laramate_900BP) and wide range of ancient populations from Beringia (*Saqqaq.SG, OldBeringSea_Ekven, OldBeringSea_Uelen, USA_AK_PaleoAleut, USA_AK_NeoAleut, USA_AK_Ancient_Athabaskan_1100BP*, and *Russia_Magadan_BA*). Notice that ancient populations in the second route: *Dzhilinda1_M_N_8.4kya, Kolyma_M_10.1kya, Syalakh-Belkachi,* and *Yakutia_LNBA* are consistently shifted to the left of the curve formed by other *FSHG* populations as they share more drift with ancient Beringian populations than similar populations on the *FSHG* cline. We removed the following group labels in the *FSHG* cline from this analysis, due to their being single individuals with low coverage (< 0.1):, resulting in wide confidence intervals and high variability in the F4-statistics: *Russia_AngaraRiver_EN/ANG001, Russia_Okunevsky_Eneolithic_o1/I6950, Russia_KuangaRiver_N_1.SG/brn001.SG, Russia_MiddleAngara_N_possible/I10900_d, Russia_Zhuravlevo_N_possible/I25462, Russia_unknown_N/I7616, Russia_Popovo_HG/Popovo2, Russia_Omskay_MN/I11449,* and *Russia_Tyumen_LN_o1/I1958.*

#### VI.A.iv.e) East-Asian ancestry Admixed into Siberian populations during the Holocene along two routes

The oldest non-APS population in Siberia—i.e., the oldest population we have in our sample with only a minority of ancestry from *Khairygas_16.7kya* or *Peru_Laramate_900BP*—is the Mesolithic Southern Yakutian *KhatystyrCave_M_10.2kya* at ~10,150 BP, uncovered on the banks of the Aldan tributary leading to the Middle Lena. This individual is geographically further north than the Mesolithic Buryatian *Dzhilinda1_M_N_8.4kya* ~2,000 from years later, suggesting that the process by which East-Asian-related populations ("Neosiberians" in the terminology of [41] displaced APS populations was not smooth and that populations with greater APS affinities could have displaced populations of East Asian affinity in some time periods. However, our analyses indicate that "Neosiberian" or *ANA* ancestry was not a homogeneous entity.

The increased East Asian ancestry of the *KhatystyrCave_M_10.2kya* individual ~10,000BP from southern Yakutia appears to come mostly from Amur basin-related populations (~61% ancestry from *China_AmurRiver_Mesolithic* + ~24% ancestry from *Cisbaikal_LNBA* + ~15% from *China_SEastAsia_Coastal_EN*; Tables VI. A-16). On the other hand, two individuals (*irk007.SG* and *cta016.SG*) from the Baikal and Buryatia regions from ~8,200–8,000 BP who are part of the *Transbaikal_EMN_old* cluster demonstrate high levels of admixture from an Inland Northeast Asian-related population, represented here by an Early Neolithic Inner Mongolian (i.e. the *Yumin* individual, under the population label *China_NEastAsia_Inland_EN*). A total of ~34% to ~51% of their ancestry comes from *China_NEastAsia_Inland_EN* in two- or three-way models when the rest of ancestry comes from Amur basin-related populations (Table VI. A-17 and 18). This ancestry is retained in *Transbaikal_EMN* in Buryatia (~7,400 years BP to ~6,000 years BP), which demonstrates near-complete continuity with *Transbaikal_EMN_old*, and is augmented by further admixture in *Mongolia_N_North*, which can be modeled as *Transbaikal_EMN_old* with additional admixture from Inland Northeast Asians (Table VI. A-19). Among populations with high levels of APS ancestry, the East-Asian-related ancestry in *Syalakh-Belkachi* (which can be modeled as admixed between another APS-related indifividual, *Dzhilinda1_M_N_8.4kya* and an East-Asian-related source), also appears to be derived from an East Asian source rich in Inland Northeast Asian ancestry (either *Cisbaikal_EN*, *Mongolia_N_North*, or *Transbaikal_EMN_old*, but not others; Tables VI. A-20). This phenomenon also affects *Yakutia_LNBA*, who derives approximately half its ancestry from populations of the Transbaikal region and the rest from *Syalakh-Belkachi* (Table VI. A-10).

We find the same phenomenon in F4-statistics (Fig. S78); as affinity to *ANE* or to Native Americans declines and affinity to East Asians increases, populations become increasingly heterogeneous in their affinity to Inland or Amur-river-related populations, suggesting similarly heterogeneous sources for the East Asian ancestry in ancient groups from East Siberia. We note that agriculturalist populations along the Yellow River ~5,300-4,000BP likewise increase in affinity to Inland Neolithic East Asians as we move westward; such affinity peaks in a population of the Upper Yellow River (*China_Upper_YR_LN*). This suggests that populations related to the Yumin individual may have extended further west between the North China Plain and Mongolian Steppes, and been the source of the increased non-agriculturalist *Ancient North Asian/ANA* admixture other studies have found among agriculturalists of the Upper Yellow River valley [30]In summary, populations of the Amur River Basin are important contributors to genetic turnovers in Eastern Siberia, contributing ~50% to populations in the Transbaikal region and more to the Mesolithic Yakutian *KhatystyrCave_M_10.2kya*. However, populations of Inland Northeast Asia also appear to play an important role as well, possibly traversing the Mongolian Plateau through an inland route to contribute another type of “Neosiberian” ancestry across many periods in the Holocene.

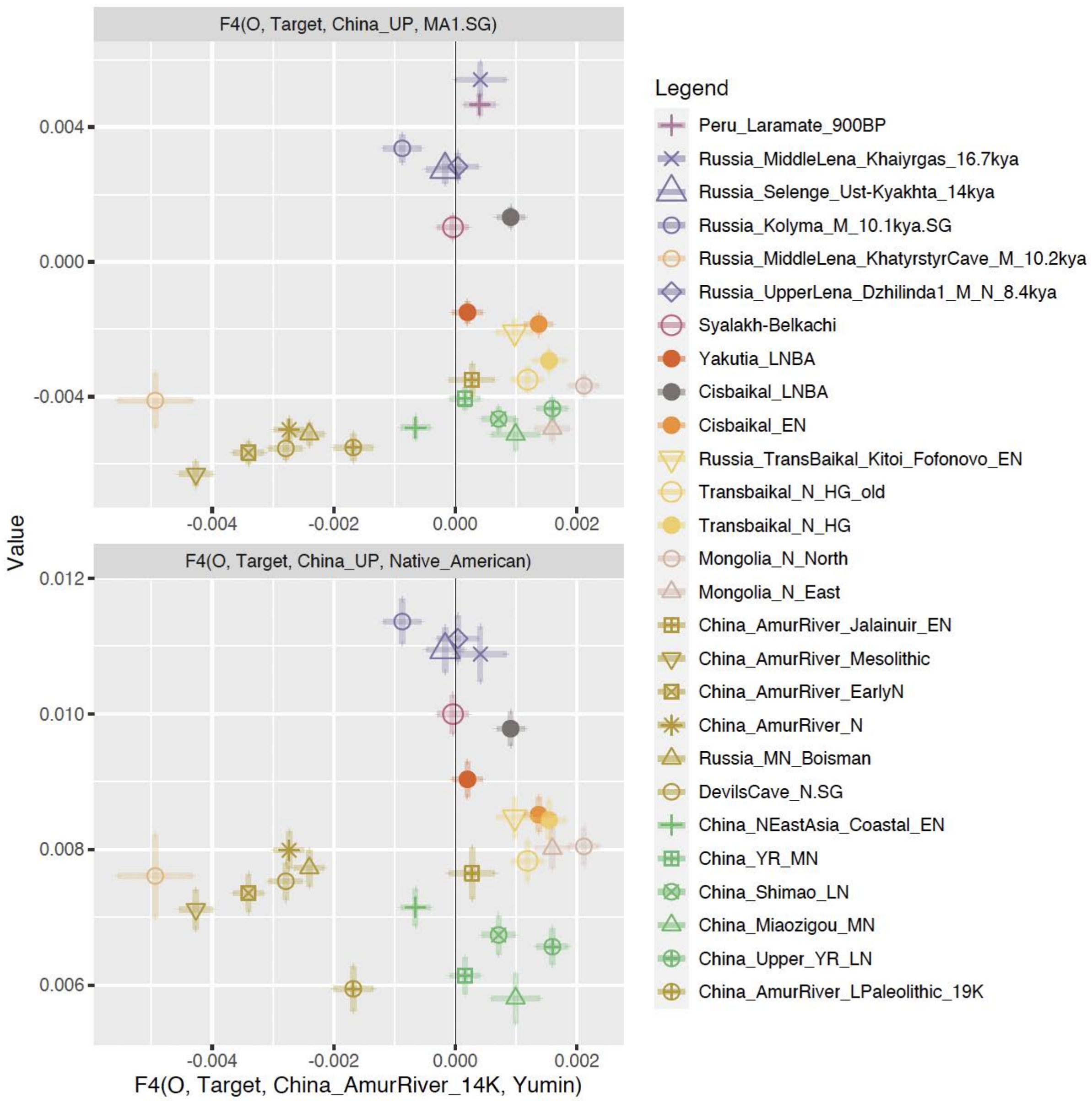

**Figure S78** | Statistics of the form F4(Ethiopia_4500BP.SG, X, China_NEastAsia_Inland_EN, China_AmurRiver_Mesolithic 14K), plotted against F4(Ethiopia_4500BP.SG, X, China_Paleolithic, MA1_HG.SG) (top) and F4(Ethiopia_4500BP.SG, X, China_Paleolithic, Peru_Laramate_900BP) (bottom), where X are ancient populations in Northeast Asia and Siberia. These statistics indicate that substructure exists within the East Asian ancestry of admixed groups in Northeast East Asia and Siberia, with differentiation between an Inland East Asian-related source represented by the *Yumin* hunter-gatherer under the population label *China_NEastAsia_Inland_EN*, and an Amur-River-related source represented by *China_AmurRiver_Mesolithic_14K*. Populations from the Amur River region throughout its history share high affinity with *China_AmurRiver_Mesolithic_14K*, while populations on the Mongolian Plateau and the Baikal area share more affinity with the Yumin hunter-gatherer. A geographic exception to this rule is the individual *KhatystyrCave_M_10.2kya*, which is extremely Amur-River-related despite occurring northeast of the Baikal region in Southern Yakutia. Interestingly, affinity to *China_NEastAsia_Inland_EN* also increases in agriculturalist populations of the Yellow River Valley as one moves west, suggesting that this population was more widely distributed. Groups rich in APS ancestry, such as *Dzhilinda1_M_N_8.4kya*, *Kolyma_M_10.1kya*, and *Syalakh-Belkachi*, tend to be almost equally related to both groups, with the exception of *Cisbaikal_LNBA*, which is strongly Inland Northeast Asian-related.

#### *VI.B. Persistence of Ancient Paleosiberian ancestry into later populations along the Bering Straits*

To further investigate the close relationship between the APS populations in the second route (*Kolyma_M_10.1kya, Dzhilinda1_M_N_8.4kya, Syalakh-Belkachi, and Yakutia_LNBA,* VI.A.iv.) and populations on either side of the Bering Sea, we attempt to model a selection of later populations from Beringia and the surrounding regions that linguists and archaeologists have connected to present-day groups. We model these selected populations using the same or a similar *qpAdm* setup as was used in the previous section—except that these Beringian populations are treated as a final tranche, with all other populations (Tranche 0 + 1 + 2 + 3 + 4 + 5 + 6) in the references/sources.

The Beringian populations selected for this analysis include:

- The *Saqqaq.SG* Paleo-Eskimo, from Greenland ~3,800BP, with unclear affiliations;
- A Late Dorset individual (*Canada_LateDorset.SG*) from an archaeological culture that continued Paleo-Eskimo traditions as recently as ~800BP, from the Canadian Arctic Islands [33];
- The *OldBeringSea_Ekven* and *OldBeringSea_Uelen* Neo-Eskimo populations from the Asian shore of the Bering Straits ~2,000BP, that have been connected to present-day Inuit [9];
- A *Thule.SG* individual from the Canadian Arctic Islands ~600BP, also connected present-day Inuit [32];
- The *Magadan_BA* population from the Northern shore of the Okhotsk Sea ~3,000BP that is extremely similar and is most closely related to present-day Chukotko-Kamchatkans [41];
- An Iron Age Yakutian (*Russia_Yakutia_IA.SG*) from the Lena River Valley from ~2600BP, from a time and place that would likely have connected his population with present-day Yukaghirs [16];
- Three ancient Athabaskan individuals (*USA_AK_Ancient_Athabaskan_1100BP*) from inland Alaska from ~1100BP [9].

Because there is only one Native-American-related population (*Peru_Laramate_900BP*) in the original set of Tranches (Tranche 0 + 1 + 2 + 3 + 4 + 5 + 6), we noticed that Native-American-related ancestry proportions were poorly estimated when that population was moved into the sources, such that models for some targets with *Peru_Laramate_900BP* in the sources always had negative admixture coefficients. We therefore also ran a second set of *qpAdm* analyses including a source for Northern Native American (*NNA*) ancestry and a reference for Southern Native American (*SNA*) ancestry (**Scheib et. al. 2018**); here, only the NNA population was moved into the sources, so that the SNA population in the references could be used to constrain estimates of the putatively North-American-related gene flow into our selected Beringian populations. The identity of the individuals included in all group labels in this analysis, including those under the Northern Native American (*NNA_10000BP-400_BP*) and Southern Native American (*SNA_12000-400BP*) populations, can be found in Table 1 of Data S5.

***Modelling approach and Parameters.*** We use the same set of references and settings for `admixr` in the previous section (Section VI. A. 1.), and a maximal set of populations in the references (i.e. with tranche 0 + 1 + 2 + 3 + 4 + 5 + 6 in the references/sources). We recalculate the $f_2$-statistics used by admixr over the same individuals as used in the previous section, with the addition of our targeted Beringian populations.

We first use this setup to model the oldest individual in our dataset, the Saqqaq Paleo-Eskimo. Then, we move Saqqaq Paleo-Eskimo into the references for all later populations that we model from the Beringian region (i.e. with tranche 0 + 1 + 2 + 3 + 4 + 5 + 6 + *Saqqaq.SG* in the references/sources), as Saqqaq.SG ancestry has been argued in previous publications to have played an important role in all populations in the New World that received gene flow from Holocene migrations from the Old World, including Paleo- and Neo-Eskimos and Athabaskans.

In subsequent models, we find that having a single Native American reference population (*Peru_Laramate_900BP*) that may be moved into the sources causes Native American admixture to be poorly estimated whenever *Peru_Laramate_900BP* is a source; many such models have negative ancestry coefficients, or very high standard errors. We therefore replace the single

*Peru_Laramate_900BP* with two Native American populations, one each in the references and sources respectively. The first is a population constituting 42 individuals representing *Southern Native American* ancestry (*SNA*, also known as *ANC_A* [40]*),* added to in the references, and the second a population constituting 7 individuals representing *Northern Native American* ancestry (*NNA*, also known as *ANC_B*) added to the sources. We used these populations under the assumption that all Native American admixture in Beringian groups comes from *Northern Native Americans*/*NNA*, while also aiming to maximize coverage by using large numbers of shotgun-sequenced individuals. We find that Native American admixture in ancient Beringians is much better estimated with this new setup. We present both sets of results for all the populations that follow after the *Saqqaq.SG* individual (i.e. the first set with tranche 0 + 1 + 2 + 3 + 4 + 5 + 6 + *Saqqaq.SG* in the references and sources, which means *Peru_Laramate_900BP* is the single Native American population in the references and sources, and the second set with tranche 0 + 1 + 2 + 3 + 4 + 5 + 6 + *Saqqaq.SG* but with *Peru_Laramate_900BP* replaced by *SNA* in the references and *NNA* in the sources). Because of the large number of references which creates a multiple hypothesis testing burden, we list all models with p > 0.01. We find passing models for all populations at this level of significance.

##### VI.B.i. Connection between the Syalakh-Belkachi Population and the Saqqaq Paleo-Eskimo

***Close connection between the Syalakh-Belkachi population and the ancient Saqqaq Paleo-Esimo.*** We find an especially close relationship exists between the *Syalakh-Belkachi* population and the Paleo-Inuit individual from the Saqqaq site in Greenland, replicating a result in [16]. Only 3-way models with *Syalakh-Belkachi* (~64% - ~79%), *Kolyma_M_10.1kya*, and an East Asian source rich in Inland Northeast Asian ancestry pass; *Syalakh-Belkachi* and *Kolyma_M_10.1kya* are rotated into the sources from the references in every single passing model (Table VI. B-1). The results indicate that the Paleo-Inuit individual from Greenland is modeled with most of its ancestry from people of the *Syalakh-Belkachi* culture of Siberia and a minor contribution from Mesolithic populations close to the Bering Strait, which is sensible in light of temporal and geographic considerations. All models can be found in Table 1 of Data S5.

The Syalakh and Belkachi cultures are similar Mid-Holocene archaeological cultures that occupied extensive areas of the Lena basin. They marked the first appearance of ceramics and polished stone tools in the region and are considered Early and Middle Neolithic cultures within the local chronology [56]. They have been connected repeatedly (based on archaeological evidence) to the emergence of the Paleo-Eskimo culture and its associated Arctic Small Tool tradition of the North American Arctic region in the literature ([56] and references therein), in line with the genomic connections between *Syalakh-Belkachi* samples and the *Saqqaq.SG* individual discussed here and in [16]. While [9] suggest that this trans-Beringian connection is linked to the dispersal of Dene languages into North America by way of the *Saqqaq.SG* -related contribution to present-day Athabaskans, [57] instead connect this to the spread of the Eskimo-Aleut languages from an Asian location into North America—a dispersal that accounts for connections between the Eskimo-Aleut family and language groups of the Old World, such as the Yukagir and Uralic language families. Further investigation and discussion of the place of *Saqqaq.SG* and *Syalakh-Belkachi* in the Beringian genomic landscape in light of our results takes place in SI section VI. B. iii.

All models in this section can be found in Table 2, Data S5.

**Table S26**

| Target | Model | P | Source | Prop. | SE | Z |
|---|---|---|---|---|---|---|

| | | | | | | |
|---|---|---|---|---|---|---|
| Greenland_Saqqaq.SG | Kolyma_M_10.1kya + Syalakh-Belkachi + Cisbaikal_EN | 0.144 | Kolyma_M_10.1kya | 0.183 | 0.067 | 2.72E+00 |
| Greenland_Saqqaq.SG | Kolyma_M_10.1kya + Syalakh-Belkachi + Cisbaikal_EN | 0.144 | Syalakh-Belkachi | 0.640 | 0.105 | 6.09E+00 |
| Greenland_Saqqaq.SG | Kolyma_M_10.1kya + Syalakh-Belkachi + Cisbaikal_EN | 0.144 | Cisbaikal_EN | 0.178 | 0.063 | 2.80E+00 |
| Greenland_Saqqaq.SG | Kolyma_M_10.1kya + Syalakh-Belkachi + *Mongolia_N_North* | 0.134 | Kolyma_M_10.1kya | 0.188 | 0.067 | 2.81E+00 |
| Greenland_Saqqaq.SG | Kolyma_M_10.1kya + Syalakh-Belkachi + *Mongolia_N_North* | 0.134 | Syalakh-Belkachi | 0.653 | 0.097 | 6.75E+00 |
| Greenland_Saqqaq.SG | Kolyma_M_10.1kya + Syalakh-Belkachi + *Mongolia_N_North* | 0.134 | *Mongolia_N_North* | 0.158 | 0.050 | 3.18E+00 |
| Greenland_Saqqaq.SG | Kolyma_M_10.1kya + Syalakh-Belkachi + Transbaikal_EMN | 0.121 | Kolyma_M_10.1kya | 0.211 | 0.072 | 2.94E+00 |
| Greenland_Saqqaq.SG | Kolyma_M_10.1kya + Syalakh-Belkachi + Transbaikal_EMN | 0.121 | Syalakh-Belkachi | 0.621 | 0.115 | 5.38E+00 |
| Greenland_Saqqaq.SG | Kolyma_M_10.1kya + Syalakh-Belkachi + Transbaikal_EMN | 0.121 | Transbaikal_EMN | 0.169 | 0.059 | 2.84E+00 |
| Greenland_Saqqaq.SG | Kolyma_M_10.1kya + Syalakh-Belkachi + Cisbaikal_LNBA | 0.104 | Kolyma_M_10.1kya | 0.012 | 0.075 | 1.58E-01 |
| Greenland_Saqqaq.SG | Kolyma_M_10.1kya + Syalakh-Belkachi + Cisbaikal_LNBA | 0.104 | Syalakh-Belkachi | 0.783 | 0.069 | 1.13E+01 |
| Greenland_Saqqaq.SG | Kolyma_M_10.1kya + Syalakh-Belkachi + Cisbaikal_LNBA | 0.104 | Cisbaikal_LNBA | 0.205 | 0.073 | 2.83E+00 |
| Greenland_Saqqaq.SG | China_NEastAsia_Inland_EN + Kolyma_M_10.1kya + Syalakh-Belkachi | 0.096 | China_NEastAsia_Inland_EN | 0.094 | 0.047 | 1.99E+00 |
| Greenland_Saqqaq.SG | China_NEastAsia_Inland_EN + Kolyma_M_10.1kya + Syalakh-Belkachi | 0.096 | Kolyma_M_10.1kya | 0.109 | 0.064 | 1.71E+00 |
| Greenland_Saqqaq.SG | China_NEastAsia_Inland_EN + Kolyma_M_10.1kya + Syalakh-Belkachi | 0.096 | Syalakh-Belkachi | 0.797 | 0.083 | 9.63E+00 |
| Greenland_Saqqaq.SG | Kolyma_M_10.1kya + Transbaikal_EMN_old + Syalakh-Belkachi | 0.065 | Kolyma_M_10.1kya | 0.187 | 0.074 | 2.53E+00 |
| Greenland_Saqqaq.SG | Kolyma_M_10.1kya + Transbaikal_EMN_old + Syalakh-Belkachi | 0.065 | Transbaikal_EMN_old | 0.125 | 0.056 | 2.24E+00 |
| Greenland_Saqqaq.SG | Kolyma_M_10.1kya + Transbaikal_EMN_old + Syalakh-Belkachi | 0.065 | Syalakh-Belkachi | 0.688 | 0.113 | 6.09E+00 |

**Table 25 |** Passing models for the Saqqaq individual with all populations from Siberia and Northeast Asia in the references. **References:** [Tranche 0 + Tranche 1 + Tranche 2 + Tranche 3 + Tranche 4 + Tranche 5]

**Sources:** [*Peru_Laramate_900BP, Russia_AfontovaGora3, Khaiyrgas, Ust_Kyakhta_14kya, China_AmurRiver_LPaleolithic_19K, China_AmurRiver_LPaleolithic_14K, China_AmurRiver_Mesolithic, China_NEastAsia_Coastal_EN, China_NEastAsia_Inland_EN, China_SEastAsia_Coastal_EN*, *KhatystyrCave_M_10.2kya, Kolyma_M_10.1kya, Transbaikal_EMN_old, Dzhilinda1_M_N_8.4kya, Altai_N_old, Syalakh_Belkachi, Yakutia_LNBA, Transbaikal_EMN, Mongolia_N_North, China_AmurRiver_N*]

#### VI.B.ii. *qpAdm* models indicate that Ancient Paleosiberian ancestry persisted on both sides of the Bering Strait, especially through *Saqqaq.SG*

***Canada_LateDorset.SG.*** The Late Dorset individual from ~800BP, from North Devon Island (among the Canadian Arctic Islands stretching between the North American mainland and Greenland), can be modeled under both sets of *qpAdm* as having complete descent from *Saqqaq.SG*, in conformity with archaeological deductions that the Dorset culture as being among the last of the Paleo-Eskimo archaeological traditions. There are no other passing models for either set of *qpAdm*.

***Canada_Thule.SG.*** The Thule individual from ~600BP, from Southampton Island also in the Canadian Arctic, is very similar to and demonstrates substantial continuity with present-day Inuit [3]. It can be modeled under both sets of *qpAdm* as retaining partial ancestry from *Saqqaq.SG* (~60-54%) with the remainder drawn from Native American-related populations (~40-46%). All models with p>0.03 contain Saqqaq.SG, including all passing 3- and 4-way models; the lowest p-value of any model with

*Saqqaq.SG* in the sources is p=0.084, compared to the single passing model without *Saqqaq.SG* at p=0.025.

***Russia_OldBeringSea_Uelen* & *Russia_OldBeringSea_Ekven*.** The populations of the Uelen and the Ekven sites of the Old Bering Sea culture (from ~2000-700BP), which spread along the Asian coast of the Bering straits and which archaeologists have described as the oldest culture with Thule traditions, can be modeled, like *Canada_Thule.SG,* as having much of its ancestry drawn from a combination of *Saqqaq.SG* and a Native-American-related source. While this is true for both sets of *qpAdm* analyses, standard errors are very large under the reference set containing only *Peru_Laramate_900BP*. This is much improved in the *qpAdm* with *NNA* and *SNA* populations. Here, both Ekven and Uelen populations are modeled as having most ancestry (~46-50%) from the NNA source, substantial ancestry from Saqqaq.SG (~25-43%), and the small remainder drawn from one or more sources rich in East Asian ancestry (*Cisbaikal_EN, Dzhilinda1_M_N_8.4kya*, *Yakutia_LNBA* or *KhatystyrCave_M_10.2kya*). However, the models for *Russia_OldBeringSea_Ekven* now marginally fail under the *qpAdm* with NNA and SNA populations. All passing models with a larger number of sources than those listed here, as well as all models with p>0.001 for *Russia_OldBeringSea_Ekven* under the *qpAdm* with NNA and SNA populations, involve both *Saqqaq.SG* and a Native-American-related population as sources that account for most of the ancestry of these two populations.

***Russia_Magadan_BA.*** This population, from the Ol'skaia site of the Tokarev culture on the northern shores of the Okhotsk sea ~3000BP, is extremely similar to and shares a very high level of genetic drift with present-day Chukotko-Kamchatkan-speaking populations [41]. We find that this population has relatively complex ancestry; no models pass under the *qpAdm* with *Peru_Laramate_900BP* in the sources and references, and, while 4-way models pass under the *qpAdm* with NNA and SNA populations (p<0.03), only 5-way models pass at the p>0.05 level (p = 0.13). We find that 4- and 5-way models in the first set of *qpAdm* with *Peru_Laramate_900BP* as the Native American-related ancestry source (which the other set of *qpAdm* involving *NNA* and *SNA* populations indicate is required to model this population correctly) tend to fail due to the presence of negative admixture coefficients, which also attests to the complex ancestry of this population.

The passing models in the set with NNA and SNA ancestries indicate that, like *Canada_Thule.SG* and the Old Bering Sea populations, *Russia_Magadan_BA* requires ancestry from a Native-American-related source and *Saqqaq.SG*; but unlike them, all passing models for Russia_Magadan_BA also require *Kolyma_M_10.1kya* as a source. This three-way combination of sources (a Native American-related source, *Saqqaq.SG*, and *Kolyma_M_10.1kya*) is present in all passing models. The remainder of their ancestry (<~25%) can be accounted for by minor admixture from one or two sources with high levels of East Asian ancestry (*KhatystyrCave_M_10.2kya, Cisbaikal_LNBA*, *Ust_Kyakhta_14kya,* or *China_NEastAsia_Inland_EN*).

***Russia_Yakutia_IA.SG.*** This population from the Central Lena River Valley ~2,600BP, which is extremely similar to present-day Yukaghirs and Nganasans, clustering with them in PCA and ADMIXTURE (Main Text Fig. 1, Center; Extended Data Figure 6; Extended Data Figure 10), and can be modeled with complete descent from *Yakutia_LNBA.*

All models in this section can be found in Table 3A (models with *Peru_Laramate_900BP*) and 3B (models with *NNA_10000BP-400_BP* and *SNA_12000-400BP*), Data S5.

**Table S27**

| Target | Model | P | Source | Prop. | SE | Z |
|---|---|---|---|---|---|---|
| Canada_LateDorset.SG | Greenland_Saqqaq.SG | 0.052 | Greenland_Saqqaq.SG | 1.00 | 3.20E-14 | 3.13E+13 |
| Canada_Thule.SG | Peru_Laramate_900BP + Greenland_Saqqaq.SG | 0.145 | Peru_Laramate_900BP | 0.41 | 6.79E-02 | 5.99E+00 |

| | | | | | | |
|---|---|---|---|---|---|---|
| Canada_Thule.SG | Peru_Laramate_900BP + Greenland_Saqqaq.SG | 0.145 | Greenland_Saqqaq.SG | 0.59 | 6.79E-02 | 8.74E+00 |
| Canada_Thule.SG | Peru_Laramate_900BP + Syalakh-Belkachi | 0.026 | Peru_Laramate_900BP | 0.29 | 6.59E-02 | 4.39E+00 |
| Canada_Thule.SG | Peru_Laramate_900BP + Syalakh-Belkachi | 0.026 | Syalakh-Belkachi | 0.71 | 6.59E-02 | 1.08E+01 |
| Russia_OldBeringSea_Uelen | Dzhilinda1_M_N_8.4kya + Peru_Laramate_900BP + Greenland_Saqqaq.SG | 0.060 | Dzhilinda1_M_N_8.4kya | 0.64 | 2.01E-01 | 3.15E+00 |
| Russia_OldBeringSea_Uelen | Dzhilinda1_M_N_8.4kya + Peru_Laramate_900BP + Greenland_Saqqaq.SG | 0.060 | Peru_Laramate_900BP | 0.23 | 7.11E-02 | 3.25E+00 |
| Russia_OldBeringSea_Uelen | Dzhilinda1_M_N_8.4kya + Peru_Laramate_900BP + Greenland_Saqqaq.SG | 0.060 | Greenland_Saqqaq.SG | 0.13 | 1.45E-01 | 9.23E-01 |
| Russia_OldBeringSea_Ekven | Kolyma_M_10.1kya + Peru_Laramate_900BP + Yakutia_LNBA + Greenland_Saqqaq.SG | 0.073 | Kolyma_M_10.1kya | 0.24 | 2.33E-01 | 1.01E+00 |
| Russia_OldBeringSea_Ekven | Kolyma_M_10.1kya + Peru_Laramate_900BP + Yakutia_LNBA + Greenland_Saqqaq.SG | 0.073 | Peru_Laramate_900BP | 0.35 | 1.79E-01 | 1.95E+00 |
| Russia_OldBeringSea_Ekven | Kolyma_M_10.1kya + Peru_Laramate_900BP + Yakutia_LNBA + Greenland_Saqqaq.SG | 0.073 | Yakutia_LNBA | 0.14 | 8.99E-02 | 1.58E+00 |
| Russia_OldBeringSea_Ekven | Kolyma_M_10.1kya + Peru_Laramate_900BP + Yakutia_LNBA + Greenland_Saqqaq.SG | 0.073 | Greenland_Saqqaq.SG | 0.27 | 8.40E-02 | 3.24E+00 |
| Russia_CentralYakutia_IA.SG | Yakutia_LNBA | 0.667 | Yakutia_LNBA | 1.00 | 2.31E-13 | 4.33E+12 |

**Table S1-VI. B-2 |** Passing models for ancient Beringians and Arctic North Americans with all populations from Siberia and Northeast Asia in the references. **References:** [Tranche 0 + Tranche 1 + Tranche 2 + Tranche 3 + Tranche 4 + Tranche 5 + Tranche 6 + *Saqqaq.SG*] **Sources:** [*Peru_Laramate_900BP, Russia_AfontovaGora3, Khaiyrgas, Ust_Kyakhta_14kya, China_AmurRiver_LPaleolithic_19K, China_AmurRiver_LPaleolithic_14K, China_AmurRiver_Mesolithic, China_NEastAsia_Coastal_EN, China_NEastAsia_Inland_EN, China_SEastAsia_Coastal_EN*, *KhatystyrCave_M_10.2kya, Kolyma_M_10.1kya, Transbaikal_EMN_old, Dzhilinda1_M_N_8.4kya, Altai_N_old, Syalakh_Belkachi, Yakutia_LNBA, Transbaikal_EMN, Mongolia_N_North, China_AmurRiver_N, Saqqaq.SG*]

#### Table S28

| Target | Model | P | Source | Prop. | SE | Z |
|---|---|---|---|---|---|---|
| Canada_LateDorset.SG | Greenland_Saqqaq.SG | 0.044 | Greenland_Saqqaq.SG | 1.000 | 3.20E-14 | 3.13E+13 |
| Canada_Thule.SG | NNA_10000-400BP.SG + Greenland_Saqqaq.SG | 0.098 | NNA_10000-400BP.SG | 0.461 | 2.62E-02 | 1.76E+01 |
| Canada_Thule.SG | NNA_10000-400BP.SG + Greenland_Saqqaq.SG | 0.098 | Greenland_Saqqaq.SG | 0.539 | 2.62E-02 | 2.06E+01 |
| Russia_OldBeringSea_Uelen | NNA_10000-400BP.SG + Yakutia_LNBA + Greenland_Saqqaq.SG | 0.224 | NNA_10000-400BP.SG | 0.505 | 1.43E-02 | 3.52E+01 |
| Russia_OldBeringSea_Uelen | NNA_10000-400BP.SG + Yakutia_LNBA + Greenland_Saqqaq.SG | 0.224 | Yakutia_LNBA | 0.112 | 5.97E-02 | 1.88E+00 |
| Russia_OldBeringSea_Uelen | NNA_10000-400BP.SG + Yakutia_LNBA + Greenland_Saqqaq.SG | 0.224 | Greenland_Saqqaq.SG | 0.382 | 6.65E-02 | 5.75E+00 |
| Russia_OldBeringSea_Uelen | Dzhilinda1_M_N_8.4kya + NNA_10000-400BP.SG + Greenland_Saqqaq.SG | 0.016 | Dzhilinda1_M_N_8.4kya | 0.285 | 2.05E-01 | 1.39E+00 |
| Russia_OldBeringSea_Uelen | Dzhilinda1_M_N_8.4kya + NNA_10000-400BP.SG + Greenland_Saqqaq.SG | 0.016 | NNA_10000-400BP.SG | 0.460 | 2.19E-02 | 2.10E+01 |
| Russia_OldBeringSea_Uelen | Dzhilinda1_M_N_8.4kya + NNA_10000-400BP.SG + Greenland_Saqqaq.SG | 0.016 | Greenland_Saqqaq.SG | 0.255 | 1.88E-01 | 1.36E+00 |
| Russia_OldBeringSea_Uelen | KhatystyrCave_M_10.2kya + NNA_10000-400BP.SG + Greenland_Saqqaq.SG | 0.012 | KhatystyrCave_M_10.2kya | 0.074 | 4.36E-02 | 1.69E+00 |
| Russia_OldBeringSea_Uelen | KhatystyrCave_M_10.2kya + NNA_10000-400BP.SG + Greenland_Saqqaq.SG | 0.012 | NNA_10000-400BP.SG | 0.494 | 1.45E-02 | 3.41E+01 |
| Russia_OldBeringSea_Uelen | KhatystyrCave_M_10.2kya + NNA_10000-400BP.SG + Greenland_Saqqaq.SG | 0.012 | Greenland_Saqqaq.SG | 0.432 | 5.08E-02 | 8.50E+00 |
| Russia_OldBeringSea_Ekven | China_NEastAsia_Coastal_EN + NNA_10000-400BP.SG + Cisbaikal_EN + Yakutia_LNBA + Greenland_Saqqaq.SG | 0.005 | China_NEastAsia_Coastal_EN | 0.061 | 2.83E-02 | 2.16E+00 |

| | | | | | | |
|---|---|---|---|---|---|---|
| Russia_OldBeringSea_Ekven | China_NEastAsia_Coastal_EN + NNA_10000-400BP.SG + Cisbaikal_EN + Yakutia_LNBA + Greenland_Saqqaq.SG | 0.005 | NNA_10000-400BP.SG | 0.475 | 1.29E-02 | 3.70E+01 |
| Russia_OldBeringSea_Ekven | China_NEastAsia_Coastal_EN + NNA_10000-400BP.SG + Cisbaikal_EN + Yakutia_LNBA + Greenland_Saqqaq.SG | 0.005 | Cisbaikal_EN | 0.022 | 5.39E-02 | 4.01E-01 |
| Russia_OldBeringSea_Ekven | China_NEastAsia_Coastal_EN + NNA_10000-400BP.SG + Cisbaikal_EN + Yakutia_LNBA + Greenland_Saqqaq.SG | 0.005 | Yakutia_LNBA | 0.016 | 9.33E-02 | 1.76E-01 |
| Russia_OldBeringSea_Ekven | China_NEastAsia_Coastal_EN + NNA_10000-400BP.SG + Cisbaikal_EN + Yakutia_LNBA + Greenland_Saqqaq.SG | 0.005 | Greenland_Saqqaq.SG | 0.426 | 6.43E-02 | 6.62E+00 |
| Russia_Magadan_BA | China_NEastAsia_Inland_EN + Kolyma_M_10.1kya + NNA_10000-400BP.SG + Greenland_Saqqaq.SG | 0.024 | China_NEastAsia_Inland_EN | 0.093 | 4.05E-02 | 2.29E+00 |
| Russia_Magadan_BA | China_NEastAsia_Inland_EN + Kolyma_M_10.1kya + NNA_10000-400BP.SG + Greenland_Saqqaq.SG | 0.024 | Kolyma_M_10.1kya | 0.158 | 7.97E-02 | 1.98E+00 |
| Russia_Magadan_BA | China_NEastAsia_Inland_EN + Kolyma_M_10.1kya + NNA_10000-400BP.SG + Greenland_Saqqaq.SG | 0.024 | NNA_10000-400BP.SG | 0.123 | 2.16E-02 | 5.67E+00 |
| Russia_Magadan_BA | China_NEastAsia_Inland_EN + Kolyma_M_10.1kya + NNA_10000-400BP.SG + Greenland_Saqqaq.SG | 0.024 | Greenland_Saqqaq.SG | 0.626 | 7.43E-02 | 8.43E+00 |
| Russia_Magadan_BA | KhatystyrCave_M_10.2kya + Kolyma_M_10.1kya + NNA_10000-400BP.SG + Greenland_Saqqaq.SG | 0.018 | KhatystyrCave_M_10.2kya | 0.139 | 6.37E-02 | 2.18E+00 |
| Russia_Magadan_BA | KhatystyrCave_M_10.2kya + Kolyma_M_10.1kya + NNA_10000-400BP.SG + Greenland_Saqqaq.SG | 0.018 | Kolyma_M_10.1kya | 0.226 | 8.88E-02 | 2.54E+00 |
| Russia_Magadan_BA | KhatystyrCave_M_10.2kya + Kolyma_M_10.1kya + NNA_10000-400BP.SG + Greenland_Saqqaq.SG | 0.018 | NNA_10000-400BP.SG | 0.118 | 2.22E-02 | 5.30E+00 |
| Russia_Magadan_BA | KhatystyrCave_M_10.2kya + Kolyma_M_10.1kya + NNA_10000-400BP.SG + Greenland_Saqqaq.SG | 0.018 | Greenland_Saqqaq.SG | 0.518 | 1.14E-01 | 4.54E+00 |
| Russia_Magadan_BA | Ust_Kyakhta_14kya + Kolyma_M_10.1kya + NNA_10000-400BP.SG + Greenland_Saqqaq.SG | 0.016 | Ust_Kyakhta_14kya | 0.119 | 4.19E-02 | 2.85E+00 |
| Russia_Magadan_BA | Ust_Kyakhta_14kya + Kolyma_M_10.1kya + NNA_10000-400BP.SG + Greenland_Saqqaq.SG | 0.016 | Kolyma_M_10.1kya | 0.106 | 9.16E-02 | 1.15E+00 |
| Russia_Magadan_BA | Ust_Kyakhta_14kya + Kolyma_M_10.1kya + NNA_10000-400BP.SG + Greenland_Saqqaq.SG | 0.016 | NNA_10000-400BP.SG | 0.123 | 2.19E-02 | 5.64E+00 |
| Russia_Magadan_BA | Ust_Kyakhta_14kya + Kolyma_M_10.1kya + NNA_10000-400BP.SG + Greenland_Saqqaq.SG | 0.016 | Greenland_Saqqaq.SG | 0.652 | 6.58E-02 | 9.90E+00 |
| Russia_Magadan_BA | Kolyma_M_10.1kya + NNA_10000-400BP.SG + Cisbaikal_LNBA + Greenland_Saqqaq.SG | 0.014 | Kolyma_M_10.1kya | 0.126 | 9.34E-02 | 1.35E+00 |
| Russia_Magadan_BA | Kolyma_M_10.1kya + NNA_10000-400BP.SG + Cisbaikal_LNBA + Greenland_Saqqaq.SG | 0.014 | NNA_10000-400BP.SG | 0.134 | 2.28E-02 | 5.88E+00 |
| Russia_Magadan_BA | Kolyma_M_10.1kya + NNA_10000-400BP.SG + Cisbaikal_LNBA + Greenland_Saqqaq.SG | 0.014 | Cisbaikal_LNBA | 0.115 | 6.84E-02 | 1.68E+00 |
| Russia_Magadan_BA | Kolyma_M_10.1kya + NNA_10000-400BP.SG + Cisbaikal_LNBA + Greenland_Saqqaq.SG | 0.014 | Greenland_Saqqaq.SG | 0.625 | 7.04E-02 | 8.88E+00 |
| Russia_Magadan_BA | KhatystyrCave_M_10.2kya + Kolyma_M_10.1kya + NNA_10000-400BP.SG + Cisbaikal_LNBA + Greenland_Saqqaq.SG | 0.129 | KhatystyrCave_M_10.2kya | 0.162 | 5.53E-02 | 2.93E+00 |
| Russia_Magadan_BA | KhatystyrCave_M_10.2kya + Kolyma_M_10.1kya + NNA_10000-400BP.SG + Cisbaikal_LNBA + Greenland_Saqqaq.SG | 0.129 | Kolyma_M_10.1kya | 0.191 | 8.90E-02 | 2.15E+00 |
| Russia_Magadan_BA | KhatystyrCave_M_10.2kya + Kolyma_M_10.1kya + NNA_10000-400BP.SG + Cisbaikal_LNBA + Greenland_Saqqaq.SG | 0.129 | NNA_10000-400BP.SG | 0.135 | 2.23E-02 | 6.03E+00 |
| Russia_Magadan_BA | KhatystyrCave_M_10.2kya + Kolyma_M_10.1kya + NNA_10000-400BP.SG + Cisbaikal_LNBA + Greenland_Saqqaq.SG | 0.129 | Cisbaikal_LNBA | 0.098 | 6.43E-02 | 1.52E+00 |

| | | | | | | |
|---|---|---|---|---|---|---|
| Russia_Magadan_BA | KhatystyrCave_M_10.2kya + Kolyma_M_10.1kya + NNA_10000-400BP.SG + Cisbaikal_LNBA + Greenland_Saqqaq.SG | 0.129 | Greenland_Saqqaq.SG | 0.414 | 1.01E-01 | 4.12E+00 |
| Russia_CentralYakutia_IA.SG | Yakutia_LNBA | 0.602 | Yakutia_LNBA | 1.000 | 2.31E-13 | 4.33E+12 |

**Table S1-VI. B-3 |** Passing models for ancient Beringians and Arctic North Americans with all populations from Siberia and Northeast Asia in the references. Peru_Laramate has been dropped from the references and sources; *SNA_12000-400BP* has been added to the references, and *NNA_10000-400BP.SG* has been added to the sources. The model for *Russia_OldBeringSea_Ekven* marginally fails, as highlighted in orange, while the 5-way model for *Russia_Magadan_BA* passes with a much higher p-value, highlighted in green.

**References:** [Tranche 0 + Tranche 1 + Tranche 2 + Tranche 3 + Tranche 4 + Tranche 5 + Tranche 6 + *Saqqaq.SG*], but without *Peru_Laramate_900BP* and with *SNA_12000-400BP*

**Sources:** [*NNA_10000-400BP.SG, Russia_AfontovaGora3, Khaiyrgas, Ust_Kyakhta_14kya, China_AmurRiver_LPaleolithic_19K, China_AmurRiver_LPaleolithic_14K, China_AmurRiver_Mesolithic, China_NEastAsia_Coastal_EN, China_NEastAsia_Inland_EN, China_SEastAsia_Coastal_EN, KhatystyrCave_M_10.2kya, Kolyma_M_10.1kya, Transbaikal_EMN_old, Dzhilinda1_M_N_8.4kya, Altai_N_old, Syalakh_Belkachi, Yakutia_LNBA, Transbaikal_EMN, Mongolia_N_North, China_AmurRiver_N, Saqqaq.SG*]

#### VI.B.iii. *qpAdm* models provide evidence for a distinct stream of Ancient Paleosiberian ancestry among ancient Athabaskans

***USA_AK_Ancient_Athabaskan_1100BP.*** To our surprise, we find that two ancient Athabaskan/Dene individuals from inland Alaska ~1100BP could be modeled without *Saqqaq.SG* (i.e. with *Saqqaq.SG* retained in the references), in contrast with other populations from the Beringian, which always require significant ancestry from *Saqqaq.SG* in all passing models (VI.B.ii.).

In our first set of *qpAdm*, a marginally passing 2-way model (p = 0.024) has the ancient Athabaskans drawing their APS affinities from a large fraction of ancestry related to *Kolyma_M_10.1kya*, with the remainder drawn from a Native American-related source. The p-value is much improved (p = ~0.1) in 3-way models that draw a minor fraction of ancestry (~10%) from a third source related to *Cisbaikal_LNBA* or *Ust_Kyakhta_14kya*. Other Siberian sources produce passing 3-way models with lower p-values. In our second set of *qpAdm*, with *NNA* and *SNA* sources, all passing models require *Cisbaikal_LNBA* or *Ust_Kyakhta_14kya* among the sources, though the ancestry fractions and standard errors are poorly estimated. Once again, *Saqqaq.SG* is not required as a source for passing models.

All models in this section can be found in Table 3A (models with *Peru_Laramate_900BP*) and 3B (models with *NNA_10000BP-400_BP* and *SNA_12000-400BP*), Data S5.

##### Table S29

| Target | Model | P | Source | Prop. | SE | Z |
|---|---|---|---|---|---|---|
| USA_AK_Ancient_Athabaskan_1100BP | Peru_Laramate_900BP + Kolyma_M_10.1kya | 0.024 | Peru_Laramate_900BP | 0.23 | 1.47E-01 | 1.57E+00 |
| USA_AK_Ancient_Athabaskan_1100BP | Peru_Laramate_900BP + Kolyma_M_10.1kya | 0.024 | Kolyma_M_10.1kya | 0.77 | 1.47E-01 | 5.23E+00 |
| USA_AK_Ancient_Athabaskan_1100BP | Peru_Laramate_900BP + Kolyma_M_10.1kya + Cisbaikal_LNBA | 0.132 | Peru_Laramate_900BP | 0.294 | 0.121 | 2.44E+00 |
| USA_AK_Ancient_Athabaskan_1100BP | Peru_Laramate_900BP + Kolyma_M_10.1kya + Cisbaikal_LNBA | 0.132 | Kolyma_M_10.1kya | 0.612 | 0.146 | 4.20E+00 |
| USA_AK_Ancient_Athabaskan_1100BP | Peru_Laramate_900BP + Kolyma_M_10.1kya + Cisbaikal_LNBA | 0.132 | Cisbaikal_LNBA | 0.094 | 0.056 | 1.66E+00 |
| USA_AK_Ancient_Athabaskan_1100BP | Peru_Laramate_900BP + Ust_Kyakhta_14kya + Kolyma_M_10.1kya | 0.097 | Peru_Laramate_900BP | 0.252 | 0.125 | 2.02E+00 |
| USA_AK_Ancient_Athabaskan_1100BP | Peru_Laramate_900BP + Ust_Kyakhta_14kya + Kolyma_M_10.1kya | 0.097 | Ust_Kyakhta_14kya | 0.110 | 0.039 | 2.87E+00 |

| | | | | | | |
|---|---|---|---|---|---|---|
| USA_AK_Ancient_Athabaskan_1100BP | Peru_Laramate_900BP + Ust_Kyakhta_14kya + Kolyma_M_10.1kya | 0.097 | Kolyma_M_10.1kya | 0.637 | 0.129 | 4.95E+00 |
| USA_AK_Ancient_Athabaskan_1100BP | Peru_Laramate_900BP + Kolyma_M_10.1kya + Syalakh-Belkachi | 0.061 | Peru_Laramate_900BP | 0.520 | 0.359 | 1.45E+00 |
| USA_AK_Ancient_Athabaskan_1100BP | Peru_Laramate_900BP + Kolyma_M_10.1kya + Syalakh-Belkachi | 0.061 | Kolyma_M_10.1kya | 0.282 | 0.519 | 5.44E-01 |
| USA_AK_Ancient_Athabaskan_1100BP | Peru_Laramate_900BP + Kolyma_M_10.1kya + Syalakh-Belkachi | 0.061 | Syalakh-Belkachi | 0.198 | 0.164 | 1.20E+00 |
| USA_AK_Ancient_Athabaskan_1100BP | Peru_Laramate_900BP + Russia_AfontovaGora3 + Kolyma_M_10.1kya | 0.050 | Peru_Laramate_900BP | 0.172 | 0.308 | 5.58E-01 |
| USA_AK_Ancient_Athabaskan_1100BP | Peru_Laramate_900BP + Russia_AfontovaGora3 + Kolyma_M_10.1kya | 0.050 | Russia_AfontovaGora3 | 0.034 | 0.039 | 8.70E-01 |
| USA_AK_Ancient_Athabaskan_1100BP | Peru_Laramate_900BP + Russia_AfontovaGora3 + Kolyma_M_10.1kya | 0.050 | Kolyma_M_10.1kya | 0.794 | 0.273 | 2.91E+00 |
| USA_AK_Ancient_Athabaskan_1100BP | Peru_Laramate_900BP + Altai_N_old + Kolyma_M_10.1kya | 0.037 | Peru_Laramate_900BP | 0.00 | 2.65E-01 | 1.37E-02 |
| USA_AK_Ancient_Athabaskan_1100BP | Peru_Laramate_900BP + Altai_N_old + Kolyma_M_10.1kya | 0.037 | Altai_N_old | 0.11 | 7.46E-02 | 1.51E+00 |
| USA_AK_Ancient_Athabaskan_1100BP | Peru_Laramate_900BP + Altai_N_old + Kolyma_M_10.1kya | 0.037 | Kolyma_M_10.1kya | 0.88 | 2.07E-01 | 4.28E+00 |
| USA_AK_Ancient_Athabaskan_1100BP | Peru_Laramate_900BP + Kolyma_M_10.1kya + Dzhilinda1_M_N_8.4kya | 0.036 | Peru_Laramate_900BP | 0.48 | 3.26E-01 | 1.46E+00 |
| USA_AK_Ancient_Athabaskan_1100BP | Peru_Laramate_900BP + Kolyma_M_10.1kya + Dzhilinda1_M_N_8.4kya | 0.036 | Kolyma_M_10.1kya | 0.25 | 6.29E-01 | 3.96E-01 |
| USA_AK_Ancient_Athabaskan_1100BP | Peru_Laramate_900BP + Kolyma_M_10.1kya + Dzhilinda1_M_N_8.4kya | 0.036 | Dzhilinda1_M_N_8.4kya | 0.27 | 3.09E-01 | 8.82E-01 |
| USA_AK_Ancient_Athabaskan_1100BP | Peru_Laramate_900BP + Kolyma_M_10.1kya + Cisbaikal_EN | 0.021 | Peru_Laramate_900BP | 0.31 | 1.58E-01 | 1.95E+00 |
| USA_AK_Ancient_Athabaskan_1100BP | Peru_Laramate_900BP + Kolyma_M_10.1kya + Cisbaikal_EN | 0.021 | Kolyma_M_10.1kya | 0.64 | 1.89E-01 | 3.37E+00 |
| USA_AK_Ancient_Athabaskan_1100BP | Peru_Laramate_900BP + Kolyma_M_10.1kya + Cisbaikal_EN | 0.021 | Cisbaikal_EN | 0.06 | 4.57E-02 | 1.21E+00 |
| USA_AK_Ancient_Athabaskan_1100BP | Peru_Laramate_900BP + Kolyma_M_10.1kya + *Mongolia_N_North* | 0.018 | Peru_Laramate_900BP | 0.30 | 1.61E-01 | 1.89E+00 |
| USA_AK_Ancient_Athabaskan_1100BP | Peru_Laramate_900BP + Kolyma_M_10.1kya + *Mongolia_N_North* | 0.018 | Kolyma_M_10.1kya | 0.65 | 1.85E-01 | 3.53E+00 |
| USA_AK_Ancient_Athabaskan_1100BP | Peru_Laramate_900BP + Kolyma_M_10.1kya + *Mongolia_N_North* | 0.018 | *Mongolia_N_North* | 0.04 | 3.65E-02 | 1.21E+00 |
| USA_AK_Ancient_Athabaskan_1100BP | Peru_Laramate_900BP + Kolyma_M_10.1kya + Transbaikal_EMN_old | 0.018 | Peru_Laramate_900BP | 0.27 | 1.95E-01 | 1.40E+00 |
| USA_AK_Ancient_Athabaskan_1100BP | Peru_Laramate_900BP + Kolyma_M_10.1kya + Transbaikal_EMN_old | 0.018 | Kolyma_M_10.1kya | 0.69 | 2.27E-01 | 3.06E+00 |
| USA_AK_Ancient_Athabaskan_1100BP | Peru_Laramate_900BP + Kolyma_M_10.1kya + Transbaikal_EMN_old | 0.018 | Transbaikal_EMN_old | 0.03 | 4.27E-02 | 7.81E-01 |
| USA_AK_Ancient_Athabaskan_1100BP | Peru_Laramate_900BP + Kolyma_M_10.1kya + Transbaikal_EMN | 0.017 | Peru_Laramate_900BP | 0.33 | 1.72E-01 | 1.92E+00 |
| USA_AK_Ancient_Athabaskan_1100BP | Peru_Laramate_900BP + Kolyma_M_10.1kya + Transbaikal_EMN | 0.017 | Kolyma_M_10.1kya | 0.63 | 2.05E-01 | 3.06E+00 |
| USA_AK_Ancient_Athabaskan_1100BP | Peru_Laramate_900BP + Kolyma_M_10.1kya + Transbaikal_EMN | 0.017 | Transbaikal_EMN | 0.04 | 4.34E-02 | 9.42E-01 |
| USA_AK_Ancient_Athabaskan_1100BP | Peru_Laramate_900BP + Kolyma_M_10.1kya + Yakutia_LNBA | 0.013 | Peru_Laramate_900BP | 0.38 | 3.58E-01 | 1.07E+00 |

| | | | | | | |
|---|---|---|---|---|---|---|
| USA_AK_Ancient_Athabaskan_1100BP | Peru_Laramate_900BP + Kolyma_M_10.1kya + Yakutia_LNBA | 0.013 | Kolyma_M_10.1kya | 0.55 | 4.79E-01 | 1.15E+00 |
| USA_AK_Ancient_Athabaskan_1100BP | Peru_Laramate_900BP + Kolyma_M_10.1kya + Yakutia_LNBA | 0.013 | Yakutia_LNBA | 0.06 | 1.25E-01 | 5.15E-01 |

**Table S1-VI. B-4 |** Passing models for ancient Athabaskans with all populations from Siberia and Northeast Asia in the references.
**References:** [Tranche 0 + Tranche 1 + Tranche 2 + Tranche 3 + Tranche 4 + Tranche 5 + Tranche 6 + *Saqqaq.SG*]
**Sources:** [*Peru_Laramate_900BP, Russia_AfontovaGora3, Khaiyrgas, Ust_Kyakhta_14kya, China_AmurRiver_LPaleolithic_19K, China_AmurRiver_LPaleolithic_14K, China_AmurRiver_Mesolithic, China_NEastAsia_Coastal_EN, China_NEastAsia_Inland_EN, China_SEastAsia_Coastal_EN, KhatystyrCave_M_10.2kya, Kolyma_M_10.1kya, Transbaikal_EMN_old, Dzhilinda1_M_N_8.4kya, Altai_N_old, Syalakh_Belkachi, Yakutia_LNBA, Transbaikal_EMN, Mongolia_N_North, China_AmurRiver_N, Saqqaq.SG*]

#### Table S30

| Target | Model | P | Source | Prop. | SE | Z |
|---|---|---|---|---|---|---|
| USA_AK_Ancient_Athabaskan_1100BP | Kolyma_M_10.1kya + NNA_10000-400BP.SG + Cisbaikal_LNBA + Greenland_Saqqaq.SG | 0.026 | Kolyma_M_10.1kya | 0.097 | 8.56E-02 | 1.14E+00 |
| USA_AK_Ancient_Athabaskan_1100BP | Kolyma_M_10.1kya + NNA_10000-400BP.SG + Cisbaikal_LNBA + Greenland_Saqqaq.SG | 0.026 | NNA_10000-400BP.SG | 0.707 | 2.28E-02 | 3.10E+01 |
| USA_AK_Ancient_Athabaskan_1100BP | Kolyma_M_10.1kya + NNA_10000-400BP.SG + Cisbaikal_LNBA + Greenland_Saqqaq.SG | 0.026 | Cisbaikal_LNBA | 0.050 | 6.19E-02 | 8.00E-01 |
| USA_AK_Ancient_Athabaskan_1100BP | Kolyma_M_10.1kya + NNA_10000-400BP.SG + Cisbaikal_LNBA + Greenland_Saqqaq.SG | 0.026 | Greenland_Saqqaq.SG | 0.146 | 6.74E-02 | 2.17E+00 |
| USA_AK_Ancient_Athabaskan_1100BP | Ust_Kyakhta_14kya + Kolyma_M_10.1kya + NNA_10000-400BP.SG + Greenland_Saqqaq.SG | 0.021 | Ust_Kyakhta_14kya | 0.126 | 3.44E-02 | 3.67E+00 |
| USA_AK_Ancient_Athabaskan_1100BP | Ust_Kyakhta_14kya + Kolyma_M_10.1kya + NNA_10000-400BP.SG + Greenland_Saqqaq.SG | 0.021 | Kolyma_M_10.1kya | 0.012 | 8.69E-02 | 1.41E-01 |
| USA_AK_Ancient_Athabaskan_1100BP | Ust_Kyakhta_14kya + Kolyma_M_10.1kya + NNA_10000-400BP.SG + Greenland_Saqqaq.SG | 0.021 | NNA_10000-400BP.SG | 0.708 | 2.21E-02 | 3.20E+01 |
| USA_AK_Ancient_Athabaskan_1100BP | Ust_Kyakhta_14kya + Kolyma_M_10.1kya + NNA_10000-400BP.SG + Greenland_Saqqaq.SG | 0.021 | Greenland_Saqqaq.SG | 0.154 | 6.29E-02 | 2.45E+00 |
| USA_AK_Ancient_Athabaskan_1100BP | Kolyma_M_10.1kya + NNA_10000-400BP.SG + Syalakh-Belkachi + Cisbaikal_LNBA | 0.020 | Kolyma_M_10.1kya | 0.094 | 8.58E-02 | 1.09E+00 |
| USA_AK_Ancient_Athabaskan_1100BP | Kolyma_M_10.1kya + NNA_10000-400BP.SG + Syalakh-Belkachi + Cisbaikal_LNBA | 0.020 | NNA_10000-400BP.SG | 0.703 | 2.27E-02 | 3.10E+01 |
| USA_AK_Ancient_Athabaskan_1100BP | Kolyma_M_10.1kya + NNA_10000-400BP.SG + Syalakh-Belkachi + Cisbaikal_LNBA | 0.020 | Syalakh-Belkachi | 0.196 | 6.07E-02 | 3.23E+00 |
| USA_AK_Ancient_Athabaskan_1100BP | Kolyma_M_10.1kya + NNA_10000-400BP.SG + Syalakh-Belkachi + Cisbaikal_LNBA | 0.020 | Cisbaikal_LNBA | 0.007 | 5.30E-02 | 1.34E-01 |

**Table S1-VI. B-5 |** Passing models for ancient populations of Beringia and Arctic North Americans with all populations from Siberia and Northeast Asia in the references.
**References:** [Tranche 0 + Tranche 1 + Tranche 2 + Tranche 3 + Tranche 4 + Tranche 5 + Tranche 6 + *Saqqaq.SG*]
**Sources:** [*Peru_Laramate_900BP, Russia_AfontovaGora3, Khaiyrgas, Ust_Kyakhta_14kya, China_AmurRiver_LPaleolithic_19K, China_AmurRiver_LPaleolithic_14K, China_AmurRiver_Mesolithic, China_NEastAsia_Coastal_EN, China_NEastAsia_Inland_EN, China_SEastAsia_Coastal_EN, KhatystyrCave_M_10.2kya, Kolyma_M_10.1kya, Transbaikal_EMN_old, Dzhilinda1_M_N_8.4kya, Altai_N_old, Syalakh_Belkachi, Yakutia_LNBA, Transbaikal_EMN, Mongolia_N_North, China_AmurRiver_N, Saqqaq.SG*]

#### VI.B.iv. Summary of the role of *Ancient Paleosiberian* ancestry in population relationships on either side of the Bering Straits

***Populations in the second route through which APS ancestry persisted contributed ancestry to later groups along the Bering Strait.*** We find that later populations on either side of the Bering Straits, including later groups very similar to present-day Eskimo-Aleuts, Chukotko-Kamchatkans, Yukaghirs, and Samoyeds demonstrate partial population continuity with of the populations in the second route through which APS ancestry persisted (*Kolyma_M_10.1kya*, *Dzhilinda1_M_N_8.4kya*, *Syalakh-Belkachi*, and *Yakutia_LNBA*). Specifically, we find that:

- *Syalakh-Belkachi*-related ancestry persists through admixture from a *Saqqaq.SG*-related source in all later putatively Eskimo-Aleut-speaking ancient groups in Beringia and Arctic North America. The *Saqqaq.SG* Paleo-Eskimo can be modeled with ~62-79% of its ancestry from a *Syalakh-Belkachi*-related population, with the remainder being drawn from Mesolithic Beringia (*Kolyma_M_10.1kya*) and an Inland East Asian-related source. The large genetic contribution from a *Syalakh-Belkachi*-related source is in conformity with archaeological inferences about the origin of elements of Paleo-Eskimo archaeological traditions from the Syalakh and Belkachi horizons in Siberia ([56] and references therein). The fact that further East Asian-related ancestry (from a source rich in Inland Northeast Asian ancestry) is present in the *Saqqaq.SG* Paleo-Eskimo may indicate that the direct contributor of this ancestry to the *Saqqaq.SG* population came from further south in Siberia in the Syalakh-Belkachi horizon, perhaps close to the Baikal region. *Saqqaq.SG* ancestry is present in near-unadmixed form in a Late Dorset individual down to ~800BP.

- *Saqqaq.SG* ancestry is present in all populations putatively connected to ancient Eskimo-Aleut speakers. It is present at high levels in individuals from Old Bering Sea sites (~ up to ~43%) and in an individual from the classic Thule culture (up to ~59%), who is very similar to present-day Inuit [32]. *Saqqaq.SG* ancestry and a Native American-related source is present in passing models for all of these groups, which in the case of the Old Bering Sea culture suggests back-migration from the New World to Chukotka.

- *Saqqaq.SG* ancestry also persists along the Northern shores of the Sea of Okhotsk in a population from the Tokarev culture from ~3000 BP (*Magadan_BA*). This population is very similar to present-day Chukotko-Kamchatkans [41], and has very complex ancestry, with contributions from at least a *Kolyma_M_10.1kya*-related source and a Native American-related source on top of *Saqqaq.SG* ancestry; the remainder is drawn from East Asian-related sources.

- Ancestry related to that found in *Syalakh-Belkachi* also through persists in an Iron Age Yakutian (~2600BP) from the Middle Lena River Valley, who can be modeled as cladal with *Yakutia_LNBA*. This individual is extremely similar to present-day Yukaghirs and Nganasans, and clusters with them in PCA and ADMIXTURE (Main Text Fig. 1, Center; Extended Data Figure 6; Extended Data Figure 10).

The strong affinity between APS populations in the second route (*Kolyma_M_10.1kya, Dzhilinda1_M_N_8.4kya, Syalakh-Belkachi and Yakutia_LNBA*) and present-day Samoyed, Yukaghiric, Chukotko-Kamchatkan and Eskimo-Aleut individuals that we find in Section VI.A.iv (Fig. S77) can therefore be explained by extensive admixture from such APS populations specifically into the ancestors of these present-day groups, mediated extensively through *Saqqaq.SG-, Yakutia_LNBA-, *and to a lesser extent *Kolyma_M_10.1kya*-related populations.

Previous work [9] has indicated that Paleo-Eskimo ancestry akin to that found in *Saqqaq.SG* is present in ancient and present-day Athabaskan individuals, as well as Eskimo-Aleut and Chukotko-Kamchatkan-speakers. On the other hand, our work (in SI Section VI.D.iii) suggests a close connection between the prehistoric movements of Yeniseian speakers and the spread of *Cisbaikal_LNBA* ancestry. This immediately suggests—if ancestry from *Saqqaq.SG* is indeed associated with the spread of Athabaskan languages in North America—that we investigate the relationship between the *Saqqaq.SG*

Paleo-Eskimo and *Cisbaikal_LNBA*. Our set of *qpAdm* for *Saqqaq.SG* do not indicate a special affinity between this individual and *Cisbaikal_LNBA* (Table VI. B-1). On the other hand, some linguists have suggested that the Paleo-Eskimo migration may instead be related to the expansion of ancient Eskimo-Aleut speakers and not with the movements of ancient Dene groups, which may instead have come from a separate mid-Holocene migration from the Old World [57]. They cite the lack of loanwords supporting extensive contact between the prehistoric speakers of Dene- and Eskimo-Aleut languages (which is unexpected if the ancestors of both linguistic groups traveled together in a *Saqqaq.SG*-related population), and the lack of linguistic evidence for an Arctic coastal foraging stage among the linguistic ancestors of Dene-speakers, to support their deductions. It has also been difficult to find archaeological links between the Arctic-coastal material culture of the Paleo-Eskimos and cultures linked to ancient Athabaskans [58]. We therefore also attempt to model ancient Athabaskans from ~1100 years BP using our *qpAdm* framework (S1 Table VI. B-4 & 5).

While all ancient populations from Beringia and Arctic North America require a *Saqqaq.SG*-related source for passing models, we find, to our surprise, that Ancient Athabaskans do not share in this pattern. Our models indicate that ancient Athabaskans can be modeled without any ancestry from *Saqqaq.SG* (i.e., with *Saqqaq.SG* retained in the reference populations); rather, in a 3-way model where most ancestry comes from a Native-American-related and a *Kolyma_M_10.1kya*-related source, model p-values are maximized by a minor (~9-13%) contribution from a third source that may be either *Cisbaikal_LNBA* or the closely-related *Ust_Kyakhta_14kya* (Table VI. B-4). In the second set of *qpAdm* models with an *NNA* population in the sources and a *SNA* population in the references, we find that either *Cisbaikal_LNBA* or *Ust_Kyakhta_14kya* are also required as the sources in all passing models, though the standard errors are high. We therefore obtain some evidence that ancient Athabaskan populations may have been genetically isolated from Paleo-Eskimo populations like *Saqqaq.SG*, and weak–but suggestive–evidence, that populations related to *Cisbaikal_LNBA* contributed to Athabaskan speakers of the New World (Section VI.D.).

As a long interval separates any putative migration of Dene speakers and the time when these ancient Athabaskans lived, a high proportion of Native-American-related ancestry that dilutes a significant fraction of presumably intrusive ancestry, which in turn is a combination of a major contribution from Mesolithic ancestry native to Beringia (represented here by *Kolyma_M_10.1kya*) and minor ancestry from linguistically related groups in the Old World (represented here by *Cisbaikal_LNBA* or the closely-related *Ust_Kyakhta_14kya*) makes sense geographically and linguistically. Our results, on their face, may appear to contradict those of [9], who used rare allele analysis to show that present-day Dene-speakers and Eskimo-Aleuts share a non-Native American ancestry source best represented by Saqqaq.SG. However, Saqqaq.SG was the only ancient sample with substantial quantities of APS ancestry available to researchers during the time when this manuscript was being prepared (the earliest version of that manuscript was posted in 2016 [59]**,** and the manuscript was received 2017**)**, and it is therefore no surprise that any connections between Athabaskan populations with other ancient APS-rich populations of Siberia or the Bering Straits could not be accounted for. Our results are more compatible with those of [41] who showed that the Siberian ancestry source for Athabaskan populations was more closely related to *Kolyma_M_10.1kya* than with *Saqqaq.SG* using chromosome painting and qpGraph, and [28], who showed that the Siberian ancestry source for Athabaskan populations was more closely related to Koryaks than with *Saqqaq.SG* using qpGraph, and more recently [60], using an MCMC algorithm sampling over admixture graph topologies. These are in line with our finding that *Kolyma_M_10.1kya* contributed ancestry to both ancient Athabaskans and ancient populations related to Chukotko-Kamchatkans, but *Saqqaq.SG* only to Chukotko-Kamchatkans and Eskimo-Aleuts but not Athabaskans.

Circumstantial evidence for a link between the *Cisbaikal_LNBA* or *Ust_Kyakhta_14kya* populations and Dene-speakers of North America may be seen in the distribution of Y-chromosomal haplogroup C-P39, which is almost exclusively found at high frequencies in North America among Athabaskan-speaking populations from Inland Alaska down to New Mexico and Arizona, and is markedly absent in Eskimo-Aleut groups of the Arctic and non-Dene Native American populations further south. The only ancient individuals found to carry this Y-chromosomal haplogroup so far are the *Ust_Kyakhta_14kya* individual himself, and 2 outlier individuals (GLZ001 and GLZ002, ~4500BP) from the Glazkovo Predmestie site of the Glazkovo culture in the Cisbaikal region in the Early Bronze Age, who carry very high levels of *Cisbaikal_LNBA* ancestry, and are surrounded by main population (i.e. the non-outlier) individuals that all fall into the *Cisbaikal_LNBA* cluster. No other ancient individuals published thus far carry this haplogroup.

Our evidence, while suggestive, is weak. Further investigation of the relationship between *Cisbaikal_LNBA*-related populations and the migration of Dene speakers would require more powerful analytic techniques and intensive sampling of ancient individuals from Alaska older than ~1100BP.

#### VI.C. Affinities of Yakutia_LNBA

Before we review evidence from formal statistics, we first highlight what unsupervised methods indicate about the distribution of *Yakutia_LNBA* ancestry among later populations and present-day groups. ADMIXTURE indicates that the *Yakutia_LNBA*-centered component (orange at K = 18, Fig. S5) occurs widely across Northern and Inner Eurasia among populations admixed between West and East Eurasian ancestry, but especially among Uralic populations. PCAs constructed to differentiate sources of East Asian ancestry likewise suggests that Uralic populations form a cline between West Eurasian populations and *Yakutia_LNBA* (Extended Data Figure 4, 5). To explore these signals further, we analyze the East Asian affinities of a set of 116 Admixed Inner Asian populations (*AIEAs*) which included Late Bronze Age and Iron Age Steppe populations, Scythians and Sarmatians, and ancient and modern Turkic, Tungusic, Mongolic, Yukaghiric, Yeniseian, and Uralic populations, using formal statistics. These groups are all the product of admixture between West and East Eurasian populations, with the West Eurasian source well-proxied by *Steppe_MLBA/Europe_LNBA* ancestries [5,29]. The situation with the East Eurasian ancestries is considerably more complex, and has been less systematically investigated (though see [51]). We first use *f4*-statistics to explore how affinity to *Yakutia_LNBA* varies among *AIEA*s, especially when compared to other types of East Asian ancestry. We especially try to find if any later populations share a specific affinity to *Yakutia_LNBA* over other East Asian ancestries. We then turn to *qpAdm* to quantify the contribution of *Yakutia_LNBA* ancestry to *AIEA*s. Our results indicate that, out of all *AIEA*s, *Yakutia_LNBA* shares a very specific connection to ancient and present-day Uralic-speaking populations.

##### VI.C.i. *f4*-statistics demonstrate that *Yakutia_LNBA* is especially related to Uralic-speaking populations

***Dataset and Methods.*** We assembled a dataset of 1408 individuals. This dataset included 1146 present-day individuals, most of which were genotyped on the Affymetric Human Origins array, the exception being 98 individuals three of which were shotgun-sequenced (Saami.DG and Mari.SG from **Raghavan et. al. (2014)** and **Lamnidis et. al. (2018)**) and the rest of which were high-coverage sequences (FIN.SG) from the 1000 genomes project study (**1000 genomes project consortium, 2015**). Of these 262 were ancient individuals, 74 of which were shotgun-sequenced and the rest of which were the result of hybridization-capture. The dataset contains information at ~0.5 million SNPs for 116 *AIEA* populations from a wide array of linguistic groups stretching back to the Middle-to-Late Bronze Age, when East Asian and West Eurasian ancestries began to mix on the Eurasian steppes [5,12,17,18,29,45]. All *F4*-statistics were calculated with the *qpDstat* package of ADMIXTOOLS with the `f4mode: YES` parameter. All individuals in this analysis, their group labels and their publications can be found in Data S1, Table 3 Column M.

***F4-statistics indicate that Uralic populations, out of all Admixed Inner Eurasian populations at any level of East Asian admixture, always share the most affinity with Yakutia_LNBA.*** We calculated statistics of the form $f_4$(O, *AIEA*, *Yakutia_LNBA*, EA) where O is the genome of the Mota hunter-gatherer (Ethiopia_4500BP.SG), *AIEA* is any Admixed Inner Eurasian population, and EA is an East Asian population from the following set: [*Yakutia_LNBA, Cisbaikal_LNBA, Syalakh-Belkachi, Mongolia_N_North, Transbaikal_EMN, China_AmurRiver_N, China_YR_LBIA*]. *Mongolia_N_North* are ~7.5 kya hunter-gatherers from South of Lake Baikal, *China_AmurRiver_N* are ~8.5-6.2kya hunter-gatherers from the Amur River Basin, and *China_YR_LBIA* are ~3.2-2.2kya farmers from Henan province in China. This statistic measures how closely related the *AIEA* population is to *Yakutia_LNBA* against a wide variety of other predominantly East Asian populations. We plotted this statistic against f4(O, *AIEA*, *China_Paleolithic*, *AG3*) where *China_Paleolithic* are two individuals from the Tianyuan-related meta-population in China prior to 33kya **(Ning et. al.)**, and *AG3* is the ~16kya old *AfontovaGora3* hunter-gatherer individual; this statistic is meant to measure the ratio of West Eurasian to East Asian ancestry in the *AIEA* population. We find that, at any ratio of East Asian to West Eurasian ancestry, the population most closely related to *Yakutia_LNBA* among *AIEA*s is always a Uralic population.

Interestingly, we find that this statistic distinguishes some Uralic populations from their otherwise very similar non-Uralic neighbours. Yeniseian-speaking Kets resemble adjacent populations speaking Uralic Samoyedic languages (Enets, Nenets and Selkup), and Yukaghiric populations resemble another Samoyedic-speaking population–Nganasans (SI Section III, Extended Data Figure 10). However in this analysis the Uralic Samoyedic-speaking populations just mentioned are clearly distinguished from populations speaking Yukaghiric and Yeniseian languages in their closer affinity to *Yakutia_LNBA*. This analysis demonstrates that Uralic-speaking populations, compared to other groups, derive the greatest relative fraction of their East Asian ancestry (or possibly even all their East Asian ancestry) from *Yakutia_LNBA*.

***PCAs of f4-statistics indicate that Uralic populations share a large amount of specific drift with Yakutia_LNBA among other East Asian populations.*** High levels of shared drift with *Yakutia_LNBA* may be caused by higher levels of shared ancestry with groups related to *Yakutia_LNBA* than with *Yakutia_LNBA* itself. Using the same dataset as before, to further investigate if Uralic populations share a high level of drift specific to *Yakutia_LNBA* over other, possibly correlated East Asian ancestries, we calculated statistics of the form f4(O, *AIEA*, *AG3*, EA) where O is the genome of the Mota hunter-gatherer (*Ethiopia_4500BP.SG*), *AIEA* is any Admixed Inner Eurasian population, *AG3* is the AfontovaGora3 hunter-gatherer, and EA is an East Asian population from the following set: [*Yakutia_LNBA, Cisbaikal_LNBA, Mongolia_N_North, China_AmurRiver_N, Transbaikal_EMN*]. We ran PCA on these statistics to reduce the influence of shared drift between different sources of East Asian ancestry, and to highlight shared drift specific to particular types of East Asian ancestry. We find that Uralic populations, again, tend to be placed on the PCA in the direction of the biplot indicating increased affinity to *Yakutia_LNBA*.

##### VI.C.ii. *qpAdm* demonstrates that Uralic-speaking populations uniquely among Inner Eurasian populations draw their East Asian ancestry almost exclusively from *Yakutia_LNBA*

Our *f4*-statistics indicate a special relationship between Uralic populations and *Yakutia_LNBA*. To quantify bounds on the level the amount of shared ancestry, we used *qpAdm* [1] to estimate the proportion of *Yakutia_LNBA* ancestry among 99 *AIEA* populations. We tested all combinations of up to seven admixture sources (one of which was always *Russia_Srubnaya* to represent *Steppe_MLBA* or *Europe_LNBA* ancestry) drawn from out of ten possible ancestry sources for each of 99 *AIEA* populations. In the list of ten possible sources, the single *Steppe_MLBA/Europe_LNBA* source was complemented by other predominantly West Eurasian sources: *Germany_EN_LBK* (a proxy for European or Anatolian farmer ancestry), *BMAC* (a proxy for ancestry from South-Central Asian agriculturalists), *Hungary_Koros_EN* (a proxy for *WHG* ancestry), *EHG*, *Tyumen_N*, and *Altai_N*. The East Asian sources included *Yakutia_LNBA*, *Mongolia_N* (a proxy for what has been termed *Ancient North Asian* or *ANA* ancestry, from the Mongolian Plateau and the Western Amur region), and *China_YR_MN* (a proxy for East Asian agriculturalist ancestry).

To distinguish between this large number of ancestry sources, we use the following set of 13 references: *China_Paleolithic* (Upper Paleolithic hunter-gatherers from China from before 33kya, including the *Tianyuan* individual), *Iran_GanjDareh_N* (early Iranian agropastoralists), *Morocco_Iberomaurusian* (hunter-gatherers from Northwest Africa), *Altai_N_old* (individuals closely related to *Altai_N* but older than 8,000 BP), *Russia_AfontovaGora3, Italy_South_HG_Ostuni* (Pre-LGM hunter-gatherers from Italy ~27kya), *Turkey_Boncuklu_N* (early Anatolia farmers), *Peru_Laramate_900BP* (a proxy for Native American ancestry), *China_SEastAsia_Island_EN, China_NEastAsia_Inland_EN* (the *Yumin* individual from Mongolia ~8,000 BP), and three populations that may help to distinguish the significant levels of APS ancestry in *Yakutia_LNBA* from other East Asian ancestries: *Syalakh-Belkachi, USA_AK_PaleoAleut,* and *USA_AK_NeoAleut*. All individuals under each population label are listed in Data S1, Table 3, Column N. We present our results in visual form by graphing the simplest model (i.e., the model that has the fewest sources) that passes for each of the 99 *AIEA* populations, except when that model is only marginally passing ($0.05 < p < 0.07$), in which case we present the simplest model that passes with a p-value > 0.07.

***Dataset and Method.*** We used a merged dataset consisting of 1781 individuals, including 99 of the *AIEA* populations mentioned in the previous section on F4-statistics (VI.C.i), and an additional number of ancient individuals used in the reference and source sets for *qpAdm* modeling. All individuals under each population label are listed in Data S1, Table 3, Column N. All models were run using `admixr` [55]. Precalculated $f_2$-statistics, used to speed up the process of f4-ratio estimation central to *qpAdm*, were used allowing for maximal missingness = .99.

***Results.*** We find passing models for 89/99 populations at the p>0.01 level, with all populations except *Nogai_Stavropol* requiring 5 sources or fewer for passing models. We find that the East Asian sources for *AIEA* populations are strikingly varied. Populations from the LBA-Scythosarmatian period typically do not require *China_YR_MN* ancestry for passing models, while most present-day Mongolic and Tungusic populations do. Mongolic, Tungusic, Turkic and LBA-Scythosarmatian populations tend to have high levels of ANA ancestry, as represented by contributions from *Mongolia_N_North*. The most striking trend in our results is that, unlike members of every other linguistic group, almost all Uralic populations can have all of their East Asian ancestry modeled as deriving from *Yakutia_LNBA*, with no contributions from *Mongolia_N_North* or *China_YR_MN*, represented in Fig. S79 by the lower row of orange dots above each bar. At the same time, almost all Uralic populations require *Yakutia_LNBA* to be present in the sources for the production of passing models. In other words, *every single passing model* (p > 0.01) for almost all Uralic populations produced by the exhaustive combinatorial generation of up to seven-way models from 10 possible sources contains *Yakutia_LNBA* within the sources. This is represented by the upper row of orange dots in Fig. S79.

Our results show that, except for the two Uralic populations that do not have passing models (*Komi_Zyrian* and *Finland_Levanluhta*), all Uralic populations both 1) require *Yakutia_LNBA* to be present within the sources for passing models and 2) can be modeled with *Yakutia_LNBA* as the sole source of their East Asian ancestry–facts that are true for no other group. All models generated from this process are listed in label are listed in Data S6, Table 1A.

To ensure that our results are robust to sequencing biases, we repeated this analysis with all shotgun sequences removed from the references and the sources [61]. Specifically, we removed all shotgun-sequenced individuals from all population labels, so that all groups in the references and sources were produced through similar enrichment-based pipelines. We find that this does not alter the results significantly, except that p-values are raised slightly, so that 91/99 populations now pass. One of these populations is *Finland_Levanluhta*, which like other Uralic populations requires *Yakutia_LNBA* in all passing models and can be modeled with *Yakutia_LNBA* as the sole East Asian source. All models in this analysis that removed all shotgun samples are listed in Data S6, Table 1B. We also modified this analysis by adding a fourth source of East Asian ancestry: *China_AmurRiver_N*, and also add *China_AmurRiver_14K* to the references, to see if the introduction of a fourth major source of East Asian ancestry on the steppes from the Amur River Basin alters our results. We find that our major conclusions are unchanged, except for some populations from the Central and Eastern Steppes and some Tungusic-speaking groups from Eastern Siberia, whose passing models now require low levels of *China_AmurRiver_N* in addition to the other three East Asian ancestries (SI Data S6, Table 2).

##### VI.C.iii. Modeling the spread of *Yakutia_LNBA* ancestry to the Volga-Ural region

We noticed that four individuals from the *Russia_Tatarka_BA* culture ~4kya just Northwest of the Minusinsk Basin, on the edge of the Baraba steppes, appear to cluster together with *Yakutia_LNBA* individuals in PCA (Main text Fig. 1, Center; Extended Data Fig. 4, 5) and ADMIXTURE (Main Test Fig. 4B). If these individuals carry high levels of *Yakutia_LNBA* ancestry, they would represent the earliest individuals with *Yakutia_LNBA* ancestry to be found so far West and South, and would signify a movement of populations carrying *Yakutia_LNBA* ancestry from the Taiga zone southwest to reach the Baraba Steppes, just northwest of the Altai mountains. We analyze these individuals using *qpAdm* to confirm that most of their ancestry derives from *Yakutia_LNBA*. Due to how similar these individuals appear to be in terms of ancestry, we group them into one population for the *qpAdm* analysis.

**Dataset and method.** We used the same *qpAdm* setup, precalculated $f_2$-statistics, and parameters as in the set of *qpAdm* used to analyze populations in Siberia in section VI.B. (i.e., Tranche 1 + 2 + 3 + 4 + 5 + 6)

**Results.** We find that the individuals from the site of Tatarka Hill can be modeled as deriving almost all their ancestry from *Yakutia_LNBA*. Passing models appear at the p>0.01 level with 2-way admixture models, but at the p>0.05 level with 4-way admixture models with small fractions of ancestry from various Siberian sources that tend to possess Inland Northeast Asian ancestry (*China_NEastAsia_Inland_EN, Cisbaikal_LNBA*), or APS ancestry from the first route (*Ust_Kyakhta_14kya, Cisbaikal_LNBA*). This suggests that the Tatarka Hill population derives almost all its ancestry from *Yakutia_LNBA*, but also experienced a low level of genetic exchange with surrounding populations in the Krasnoyarsk region and the Yenisei Basin relative to *Yakutia_LNBA*. All models in this section are found in SI Data S6, Table 3.

Continuity between the Tatarka Hill and *Yakutia_LNBA* populations is also suggested by the Y-chromosomal haplogroups, which tend to be clades of haplogroup N-L708 and its subclades N-M2126 and N-L1026. The same subclades are carried by *Yakutia_LNBA* individuals *I10855, Kra001.SG, N2a.SG*, and *N4a1.SG*. These subclades of haplogroup N are found at high frequencies among present-day Uralic speakers as well [62,63]. Our results suggest that individuals carrying *Yakutia_LNBA* ancestry reached the steppes west of the Altai mountains by the Middle Bronze Age, ~4kya, and were involved in cultures engaged in the Seima-Turbino phenomenon. Such an involvement may have opened the way to interactions and migrations that continued to spread this ancestry westward into Central Siberia and in turn west of the Urals, into European Russia.

**Table S31**

| Target | Model | P | Source | Proportion | SE | Z |
|---|---|---|---|---|---|---|
| *Russia_Samus_Tatarka_BA* | Russia_AfontovaGora3 + Yakutia_LNBA | 0.019 | Russia_AfontovaGora3 | 0.012 | 7.13E-03 | 1.66E+00 |
| *Russia_Samus_Tatarka_BA* | Russia_AfontovaGora3 + Yakutia_LNBA | 0.019 | *Yakutia_LNBA* | 0.988 | 7.13E-03 | 1.39E+02 |
| *Russia_Samus_Tatarka_BA* | Ust_Kyakhta_14kya + Yakutia_LNBA | 0.018 | Ust_Kyakhta_14kya | 0.064 | 2.31E-02 | 2.79E+00 |
| *Russia_Samus_Tatarka_BA* | Ust_Kyakhta_14kya + Yakutia_LNBA | 0.018 | *Yakutia_LNBA* | 0.936 | 2.31E-02 | 4.06E+01 |
| *Russia_Samus_Tatarka_BA* | Cisbaikal_LNBA + Yakutia_LNBA | 0.018 | Cisbaikal_LNBA | 0.078 | 3.47E-02 | 2.24E+00 |
| *Russia_Samus_Tatarka_BA* | Cisbaikal_LNBA + Yakutia_LNBA | 0.018 | *Yakutia_LNBA* | 0.922 | 3.47E-02 | 2.66E+01 |
| *Russia_Samus_Tatarka_BA* | Dzhilinda1_M_N_8.4kya + Cisbaikal_LNBA + Yakutia_LNBA | 0.032 | Dzhilinda1_M_N_8.4kya | 0.014 | 6.17E-02 | 2.22E-01 |
| *Russia_Samus_Tatarka_BA* | Dzhilinda1_M_N_8.4kya + Cisbaikal_LNBA + Yakutia_LNBA | 0.032 | Cisbaikal_LNBA | 0.063 | 5.01E-02 | 1.25E+00 |
| *Russia_Samus_Tatarka_BA* | Dzhilinda1_M_N_8.4kya + Cisbaikal_LNBA + Yakutia_LNBA | 0.032 | *Yakutia_LNBA* | 0.924 | 4.34E-02 | 2.13E+01 |
| *Russia_Samus_Tatarka_BA* | Russia_AfontovaGora3 + China_NEastAsia_Inland_EN + Cisbaikal_LNBA + Yakutia_LNBA | 0.056 | Russia_AfontovaGora3 | 0.001 | 1.06E-02 | 9.75E-02 |
| *Russia_Samus_Tatarka_BA* | Russia_AfontovaGora3 + China_NEastAsia_Inland_EN + Cisbaikal_LNBA + Yakutia_LNBA | 0.056 | China_NEastAsia_Inland_EN | 0.026 | 4.31E-02 | 5.92E-01 |
| *Russia_Tatarka_BA* | Russia_AfontovaGora3+China_NEastAsia_Inland_EN+Cisbaikal_LNBA+Yakutia_LNBA | 0.0561291 | Cisbaikal_LNBA | 0.0885458 | 0.04939687 | 1.79253888 |
| *Russia_Tatarka_BA* | Russia_AfontovaGora3+China_NEastAsia_Inland_EN+Cisbaikal_LNBA+Yakutia_LNBA | 0.0561291 | Yakutia_LNBA | 0.88489824 | 0.05179417 | 17.0849019 |
| *Russia_Tatarka_BA* | Russia_AfontovaGora3+Altai_N_old+Ust_Kyakhta_14kya+Yakutia_LNBA | 0.05004386 | Russia_AfontovaGora3 | 9.05E-06 | 0.01975125 | 4.58E-04 |
| *Russia_Tatarka_BA* | Russia_AfontovaGora3+Altai_N_old+Ust_Kyakhta_14kya+Yakutia_LNBA | 0.05004386 | Altai_N_old | 0.00115176 | 0.03712041 | 0.03102775 |
| *Russia_Samus_Tatarka_BA* | Russia_AfontovaGora3 + Altai_N_old + Ust_Kyakhta_14kya + Yakutia_LNBA | 0.050 | Ust_Kyakhta_14kya | 0.072 | 2.94E-02 | 2.43E+00 |
| *Russia_Samus_Tatarka_BA* | Russia_AfontovaGora3 + Altai_N_old + Ust_Kyakhta_14kya + Yakutia_LNBA | 0.050 | *Yakutia_LNBA* | 0.927 | 2.64E-02 | 3.51E+01 |

**Table S1-VI. A-21 |** Passing models for *Russia_Samus_Tatarka_BA* modeled with with all populations from Siberia and Northeast Asia in the references.

**References:** [Tranche 0 + Tranche 1 + Tranche 2 + Tranche 3 + Tranche 4 + Tranche 5 + Tranche 6]

**Sources:** [*Peru_Laramate_900BP, Russia_AfontovaGora3, Khaiyrgas, Ust_Kyakhta_14kya, China_AmurRiver_LPaleolithic_19K, China_AmurRiver_LPaleolithic_14K, China_AmurRiver_Mesolithic, China_NEastAsia_Coastal_EN, China_NEastAsia_Inland_EN, China_SEastAsia_Coastal_EN, KhatystyrCave_M_10.2kya, Kolyma_M_10.1kya, Transbaikal_EMN_old, Dzhilinda1_M_N_8.4kya, Altai_N_old, Syalakh_Belkachi, Yakutia_LNBA, Transbaikal_EMN, Mongolia_N_North, China_AmurRiver_N*]

### *VI.D. Affinities of Cisbaikal_LNBA*

We first highlight what unsupervised methods indicate about the distribution of *Cisbaikal_LNBA* ancestry among later populations and present-day groups. ADMIXTURE indicates that the *Cisbaikal_LNBA*-centered component (grey at K=18, (Extended Data Figure 10) occurs at non-negligible, but low levels only among present-day Kets, South-Siberian-Turkic speaking groups (Tuvinian, Tofalar, Tubalar, Altaian, Altaian_Chelkan, Kakhass, Kakhass_Kachin, Shor,

Shor_Mountain, and Todzin), and Samoyedic groups (Enets and Selkup), with the Tofalar and Ket having the highest and second-highest levels of this component. All these populations are located within the Yenisei basin. The component also appears among a scattering of ancient populations from the Bronze to Iron Ages, including an early Bronze Age individual from the Middle Angara River, and two East Asian outliers from the late Bronze Age Karasuk period, possibly from the Lugavskaya culture (SI Section II.G) from within the Minusinsk Basin on the upper reaches of the Yenisei river.

We found the geographic and temporal distribution of this component (spreading from the Cisbaikal region in the Eneolithic and Early Bronze Age, to the Middle Angara river and the Upper Yenisei by the Middle Bronze Age and Iron Ages, and finally peaking among present-day populations of the Yenisei basin), to be intriguing, as it forms a very close match to the migrations linguists have inferred that prehistoric Yeniseian-speaking populations undertook, which also involved a major movement from the Cisbaikal region to the Middle and Upper Yenisei at some point before the common era, followed 1) by assimilation of Southern Yeniseian populations into South Siberian Turkic populations in a process that began from the first millennium AD down to the Russian colonial period and 2) a more recent migration of some Yeniseian-speakers to the Central Yenisei where the Ket now live [64,65]. We therefore explored this phenomenon further using formal statistics. We first use *F4*-statistics to explore how affinity to *Cisbaikal_LNBA* varies among *AIEA*s, especially when compared to other types of East Asian ancestry. We then turn to *qpAdm* to confirm the distribution of *Cisbaikal_LNBA* ancestry among *AIEA*s suggested by ADMIXTURE. Our results indicate that, out of all *AIEA*s, *Cisbaikal_LNBA* shares very specific connections to particular ancient and present-day populations that closely match to the reconstructed ancient migrations, historical distributions, and inferred contact histories of Yeniseian-speaking populations.

##### VI.D.i. $f_4$-statistics demonstrate that *Cisbaikal_LNBA* ancestry plays an important role among Yeniseian, South Siberian Turkic, and Samoyedic populations

***PCAs of f4-statistics indicate that a subset of present-day and ancient AIEA populations centered around the Yenisei region share high levels of shared drift with Cisbaikal_LNBA.*** Using the same PCA over *f4*-statistics to reduce the influence of shared drift between different sources of East Asian ancestry, and to highlight shared drift specific to particular types of East Asian ancestry, we find that the same individuals and populations highlighted as having high levels of the *Cisbaikal_LNBA*-related component in ADMIXTURE possess high levels of drift shared specifically with *Cisbaikal_LNBA* out of multiple East Asian ancestries (the others of which are *Mongolia_N_North, China_AmurRiver_N, Transbaikal_EMN, Yakutia_LNBA*) (Main Text Fig. 3A, right).

We find that PC1 is dominated by variation in West vs East Eurasian ancestry. The biplots indicate that PC2 is driven by relative affinity to *Yakutia_LNBA* vs *Mongolia_N_North*, while PC3 highlights relative affinity to *Cisbaikal_LNBA* vs *China_AmurRiver_N*. Among ancient individuals, we find that the "Karasuk outlier" individual from the Late Bronze Age (*Karasuk_o1.SG,* individual ID *RISE497.SG*), that our research (SI Section II.G) reveals is from the Lugavskaya culture of the Northern Minusinsk Basin along the Upper Yenisei, far exceeds any present-day *AIEA* population in shared drift with *Cisbaikal_LNBA*, followed by an East-Asian-shifted outlier individual from Krasnoyarsk from the Late Bronze Age, also from the Minusinsk Basin (*Krasnoyarsk_LBA_o1.SG*, individual ID *RISE554.SG*). Among present-day populations, Kets, Selkups, and South Siberian Turkic populations of the Yenisei river valley are shifted toward *Cisbaikal_LNBA*. All populations that receive a high level of the *Cisbaikal_LNBA* component in ADMIXTURE tend to be shifted in the direction indicated by the biplot to imply a high level of drift shared specifically with *Cisbaikal_LNBA* out of East Asian populations (Main Text Fig. 3A, right).

##### VI.D.ii. *qpAdm* models demonstrate that *Cisbaikal_LNBA* ancestry plays an important role among Yeniseian, South Siberian Turkic, and Samoyedic populations

Our *f4*-statistics indicate a special relationship between Late Bronze Age East Asian outliers of the Yenisei Basin, as well as present-day populations of the Yenisei Basin, and *Cisbaikal_LNBA*. To further corroborate this relationship, we used *qpAdm* to check if *Cisbaikal_LNBA* is required for passing models, and to estimate the level of *Cisbaikal_LNBA* ancestry, among 99 *AIEA* populations. We use the same setup, dataset, and method as used for *qpAdm* in section VI.C.ii., except that we add *Cisbaikal_LNBA* as an additional population in the references, to be rotated into the sources. Populations that have significant levels of *Cisbaikal_LNBA* ancestry should now require *Cisbaikal_LNBA* to be rotated into the sources for all passing models.

***Results.*** We find again a striking concordance between linguistic and genetic boundaries, as Kets, Siberian Turkic, and Samoyedic populations consistently require ancestry from *Cisbaikal_LNBA* for passing models–i.e., tevery single passing model includes *Cisbaikal_LNBA* among the sources. This is not the case for other Turkic or Uralic populations, or for other *AIEA* populations. Two populations among the Siberian Turks and Samoyeds—Nganansans and Yakuts—appear to have lower levels of or even no *Cisbaikal_LNBA* ancestry as suggested by ADMIXTURE analysis and the PCA of *f4*-statistics, but nonetheless appear to require low levels of ancestry from *Cisbaikal_LNBA* in *qpAdm*.

The two ancient outliers from the Late Bronze Age Minusinsk Basin from the period of the Karasuk culture that ADMIXTURE and *F4*-statistics suggest have high levels of ancestry from *Cisbaikal_LNBA* also require such ancestry in *qpAdm*; interestingly, the population that succeeds the Karasuk and Lugavskaya cultures in the region, the Tagar culture (*Russia_Tagar.SG*), also requires ancestry from *Cisbaikal_LNBA* in *qpAdm* for all passing models. In addition, Mongolic-speaking Kalmyks and *Kazakhstan_CentralKazakhSteppe_Saka* also require such ancestry for passing models in this *qpAdm* setup, but no other populations from their ethnolinguistic or cultural categories behave similarly. All *qpAdm* models in this section are listed in SI Data 6, Table 4.

The proportions of *Cisbaikal_LNBA* estimated by *qpAdm* for the populations that require it suggests again that all populations with relatively high *Cisbaikal_LNBA* ancestry come from the Yenisei Basin. However, while our results corroborate the inferences from *F4*-statistics and ADMIXTURE, we caution against interpreting the estimated proportions at face value, as no passing *qpAdm* models for our *AIEA* populations contain both *Cisbaikal_LNBA* and *ANE*-rich sources such as *Altai_N* or *Tyumen_N_HG* in the sources simult*ANE*ously. When more than one of these sources are included, either standard errors increase to beyond acceptable levels, or some proportions become negative, suggesting that our references cause the estimates of *Cisbaikal_LNBA* ancestry to be inflated by conflation of ancestry from *Altai_N* or *Tyumen_N_HG* with *Cisbaikal_LNBA*. The fact that the *qpAdm* estimates may be affected by a suboptimal choice of reference populations is also supported by the fact that the relative affinity of *AIEA* populations to *Cisbaikal_LNBA* in the PCA of F4-statistics closely reflects the level of *Cisbaikal_LNBA* component estimated for each *AIEA* population in ADMIXTURE, but this correlation does not extend to the *qpAdm* results presented here. For example, the present-day populations most shifted in the direction of *Cisbaikal_LNBA* as indicated by the biplot in the PCA of *F4*-statistics are Tubalar, Todzin, and the Kets, and these are also the three present-day populations with the highest levels of the *Cisbaikal_LNBA* component in ADMIXTURE, but this pattern does not repeat in the *qpAdm* results. We therefore treat the *qpAdm* results as a diagnostic test for the presence of *Cisbaikal_LNBA* ancestry among *AIEA* populations, and less as a method for estimating for its true level.

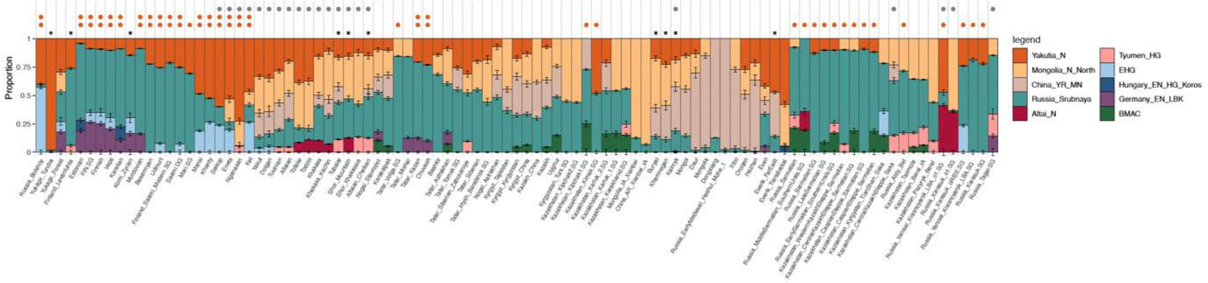

**Figure S79 | *qpAdm* results for Admixed Inner Asian populations.** One orange dot indicates that all East Asian ancestry can be modeled as deriving from *Yakutia_LNBA*; two orange dots indicate that all passing models include *Yakutia_LNBA* among the sources. A grey dot indicates that all passing models include *Cisbaikal_LNBA* in the sources when *Cisbaikal_LNBA* was placed in the references, with the option of being rotated into the sources. All models displayed are of the simplest model that passes with p>0.01; if multiple models pass, the model with the highest p-value is plotted. Populations without models passing at the p>0.01 level are indicated with an asterisk; in such cases, the model with the highest p-value is plotted.

**References:** [*China_Paleolithic, Iran_GanjDareh_N, Morocco_Iberomaurusian, Altai_N_old, Russia_AfontovaGora3, Italy_South_HG_Ostuni, Turkey_Boncuklu_N, Peru_Laramate_900BP, China_SEastAsia_Island_EN, China_NEastAsia_Inland_EN, Syalakh-Belkachi, USA_AK_PaleoAleut, USA_AK_NeoAleut*]

**Sources:** [*Russia_Srubnaya, Hungary_Koros_EN, EHG, Tyumen_N, Altai_N, Yakutia_LNBA, Mongolia_N, China_YR_MN, Germany_EN_LBK, BMAC*]

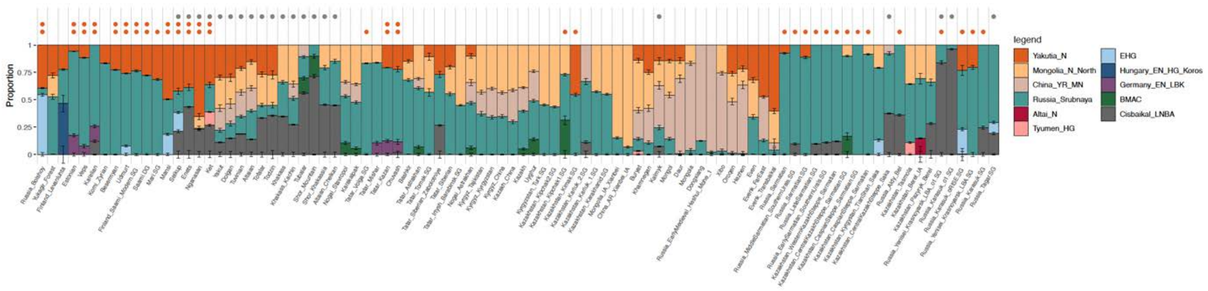

**Fig. S80| *qpAdm* results for Admixed Inner Asian populations, with estimated proportions of *Cisbaikal_LNBA* ancestry.** Note that no passing model here contains both *Cisbaikal_LNBA* and an *ANE*-rich source such as *Altai_N* and *Tyumen_N_HG*, which makes us caution against taking the estimated levels of *Cisbaikal_LNBA* ancestry at face value.

**References:** [*China_Paleolithic, Iran_GanjDareh_N, Morocco_Iberomaurusian, Altai_N_old, Russia_AfontovaGora3, Italy_South_HG_Ostuni, Turkey_Boncuklu_N, Peru_Laramate_900BP, China_SEastAsia_Island_EN, China_NEastAsia_Inland_EN, Syalakh-Belkachi, USA_AK_PaleoAleut, USA_AK_NeoAleut, Cisbaikal_LNBA*]

**Sources:** [*Russia_Srubnaya, Hungary_Koros_EN, EHG, Tyumen_N, Altai_N, Yakutia_LNBA, Mongolia_N, China_YR_MN, Germany_EN_LBK, BMAC, Cisbaikal_LNBA*]

##### VI.D.iii. Cisbaikal_LNBA ancestry expands into the Middle Angara and Krasnoyarsk regions in the Bronze Age

We noticed that three individuals along the Middle Angara River (*Russia_MiddleAngara_EBA*, constituted by the individuals I5280, I2133 and I2143) during the early Bronze Age (~4800-4500BP) appear to cluster with *Cisbaikal_LNBA* in PCA and ADMIXTURE (Fig. S1-4, Fig. S5). As the Angara river flows from its source in Lake Baikal to empty into the Middle Yenisei, these individuals may indicate that the spread of *Cisbaikal_LNBA* ancestry from the Cisbaikal region west to reach its present-day distribution along the Yenisei Basin took place during the Early to Late Bronze Age. As part of this process, the two aforementioned individuals (*Russia_Karasuk_o1.SG/RISE497.SG* and *Russia_Yenisei_Krasnoyarsk_LBA_o1.SG/RISE554.SG*) along the Upper Yenisei River from the Northern and Southern Minusinsk Basin respectively in the late Bronze Age (~3000BP and ~2800BP respectively), that possess especially high levels of shared drift with *Cisbaikal_LNBA* in the PCA of *f4*-statistics (Main Text Fig. 3A), may also be relevant. We therefore model these individuals specifically.

**Dataset and method.** To model *Russia_MiddleAngara_LBA.SG*, we used the same *qpAdm* setup, precalculated $f_2$-statistics, and parameters as used in the set of *qpAdm* for later Siberian or Beringian populations in section VI.B.i (i.e. Tranche 1 + 2 + 3 + 4 + 5 + 6). To model the two outliers from the Minusinsk Basin (*Russia_Karasuk_o1.SG/RISE497.SG* and *Russia_Yenisei_Krasnoyarsk_LBA_o1.SG/RISE554.SG*), we used the same *qpAdm* setup, precalculated $f_2$-statistics, and parameters as in the set of *qpAdm* used to analyze *AIEA*s in section VI.D.ii.

**Results.** We find that the Early Bronze Age individuals along the Middle Angara river can be modeled in 3-way models with a high proportion (>50%) of ancestry from *Cisbaikal_LNBA*, with the remainder from adjacent populations (*Altai_N_old/Altai_N*, and *Transbaikal_EMN_old/Transbaikal_EMN*). *Cisbaikal_LNBA* ancestry must be moved into the sources for all passing models. All *qpAdm* models for this population are listed in SI Data 3, Table 9.

We also find that the two Late Bronze Age individuals from the Minusinsk Basin can be modeled in simple 2-way admixture models between a *Steppe_MLBA*-related source and a *Cisbaikal_LNBA*-related source; *Cisbaikal_LNBA* is again moved into the sources for every passing model. The *Russia_Karasuk_o1.SG* individual can be modeled with almost complete descent from *Cisbaikal_LNBA* (~96%), while the *Russia_Yenisei_Krasnoyarsk_LBA_o1.SG* individual has slightly more *Steppe_MLBA* admixture (~15%). All *qpAdm* models for this population are listed in SI Data 6, Table 4.

Our models indicate that populations with *Cisbaikal_LNBA* ancestry reached the Middle Angara region by the Early Bronze Age from the Cisbaikal region. Additionally, the two individuals from the Minusinsk Basin demonstrate that subsequent contact between *Cisbaikal_LNBA* populations and *Steppe_MLBA* populations took place in the Karasuk period (~3000-2800BP), but that high levels of *Cisbaikal_LNBA* admixture was confined to various East Asian outliers at this time, which may originate from the non-Steppe, non-Karasuk, Taiga-dwelling populations of the region that may have influenced the Lugavskaya culture (SI Section II.G). If this interpretation is correct, *Cisbaikal_LNBA* populations may have effected an almost complete population turnover among the forest or forest-steppe populations of the Minusinsk basin of this period, but its influence on pastoralist populations on the Steppe proper was limited. However, the succeeding population of the Iron Age Tagar culture (*Russia_Tagar.SG*) in the Minusinsk Basin, which like the Karasuk population contains high levels of *Steppe_MLBA* ancestry, nevertheless differs from the Karasuk population by requiring low levels of admixture from a *Cisbaikal_LNBA*-related source for all of its passing models (Section VI.D.ii; Fig. S80), suggesting that admixture continued into succeeding populations of the region.

**Table S32**

| Target | Model | P | Source | Proportion | SE | Z |
|---|---|---|---|---|---|---|
| Russia_MiddleAngara_EBA | Altai_N_old + Transbaikal_EMN_old + Cisbaikal_LNBA | 0.059 | Altai_N_old | 0.185 | 2.62E-02 | 7.05E+00 |
| Russia_MiddleAngara_EBA | Altai_N_old + Transbaikal_EMN_old + Cisbaikal_LNBA | 0.059 | Transbaikal_EMN_old | 0.303 | 3.97E-02 | 7.64E+00 |
| Russia_MiddleAngara_EBA | Altai_N_old + Transbaikal_EMN_old + Cisbaikal_LNBA | 0.059 | Cisbaikal_LNBA | 0.512 | 6.11E-02 | 8.38E+00 |
| Russia_MiddleAngara_EBA | Transbaikal_EMN_old + Altai_N + Cisbaikal_LNBA | 0.031 | Transbaikal_EMN_old | 0.272 | 3.73E-02 | 7.29E+00 |
| Russia_MiddleAngara_EBA | Transbaikal_EMN_old + Altai_N + Cisbaikal_LNBA | 0.031 | Altai_N | 0.138 | 2.32E-02 | 5.95E+00 |
| Russia_MiddleAngara_EBA | Transbaikal_EMN_old + Altai_N + Cisbaikal_LNBA | 0.031 | Cisbaikal_LNBA | 0.589 | 5.58E-02 | 1.06E+01 |
| Russia_MiddleAngara_EBA | Altai_N_old + Transbaikal_EMN + Cisbaikal_LNBA | 0.012 | Altai_N_old | 0.181 | 2.70E-02 | 6.71E+00 |
| Russia_MiddleAngara_EBA | Altai_N_old + Transbaikal_EMN + Cisbaikal_LNBA | 0.012 | Transbaikal_EMN | 0.283 | 3.98E-02 | 7.11E+00 |

| | | | | | | |
|---|---|---|---|---|---|---|
| Russia_MiddleAngara_EBA | Altai_N_old + Transbaikal_EMN + Cisbaikal_LNBA | 0.012 | Cisbaikal_LNBA | 0.536 | 6.22E-02 | 8.62E+00 |

**Table S1-VI. D-1 |** Passing models for *Russia_MiddleAngara_EBA* modeled with references/sources used for Siberians and ancient Beringians in section VI.B.i (i.e. Tranche 1 + 2 + 3 + 4 + 5 + 6). No simpler models pass.
**References:** [Tranche 0 + Tranche 1 + Tranche 2 + Tranche 3 + Tranche 4 + Tranche 5 + Tranche 6]
**Sources:** [*Peru_Laramate_900BP, Russia_AfontovaGora3, Khaiyrgas, Ust_Kyakhta_14kya, China_AmurRiver_LPaleolithic_19K, China_AmurRiver_LPaleolithic_14K, China_AmurRiver_Mesolithic, China_NEastAsia_Coastal_EN, China_NEastAsia_Inland_EN, China_SEastAsia_Coastal_EN, KhatystyrCave_M_10.2kya, Kolyma_M_10.1kya, Transbaikal_EMN_old, Dzhilinda1_M_N_8.4kya, Altai_N_old, Syalakh_Belkachi, Yakutia_LNBA, Transbaikal_EMN, Mongolia_N_North, China_AmurRiver_N*]

##### Table S33

| Target | Model | P | Source | Proportion | SE | Z |
|---|---|---|---|---|---|---|
| Russia_Yenisei_Krasnoyarsk_LBA_o1.SG | Cisbaikal_LNBA + Russia_Srubnaya | 0.129 | Cisbaikal_LNBA | 0.838 | 3.19E-02 | 2.63E+01 |
| Russia_Yenisei_Krasnoyarsk_LBA_o1.SG | Cisbaikal_LNBA + Russia_Srubnaya | 0.129 | Russia_Srubnaya | 0.162 | 3.19E-02 | 5.08E+00 |
| Russia_Karasuk_o1.SG | Cisbaikal_LNBA + Russia_Srubnaya | 0.543 | Cisbaikal_LNBA | 0.961 | 1.59E-02 | 6.06E+01 |
| Russia_Karasuk_o1.SG | Cisbaikal_LNBA + Russia_Srubnaya | 0.543 | Russia_Srubnaya | 0.039 | 1.59E-02 | 2.43E+00 |

**Table S1-VI. D-1 |** Passing models for *Russia_Yenisei_Krasnoyarsk_o1.SG* & *Russia_Karasuk_o1.SG* modeled with references/sources used for *AIEA* populations in section VI.D.ii. No simpler models pass.
**References:** [*China_Paleolithic, Iran_GanjDareh_N, Morocco_Iberomaurusian, Altai_N_old, Russia_AfontovaGora3, Italy_South_HG_Ostuni, Turkey_Boncuklu_N, Peru_Laramate_900BP, China_SEastAsia_Island_EN, China_NEastAsia_Inland_EN, Syalakh-Belkachi, USA_AK_PaleoAleut, USA_AK_NeoAleut, Cisbaikal_LNBA*]
**Sources:** [*Russia_Srubnaya, Hungary_Koros_HG, Germany_EN_LBK, BMAC, EHG, Tyumen_N, Altai_N, Cisbaikal_LNBA, Yakutia_LNBA, Mongolia_N_North, China_YR_LBIA*]

### *VI.E. Modeling the FSHG cline*

#### VI.E.i. Simple *qpAdm* models demonstrate that *FSHG* populations can be modeled as a cline of admixed populations grading between East Asian ancestry, *ANE*, *EHG* and *WHG*, with post-LGM ancestry sources

Our PCAs suggest that *FSHG* populations are varying admixtures of *WHG*, *EHG*, *ANE* and East Asian-associated ancestries. To investigate this further, as well as to quantify the proportion of each ancestry source, we used *qpAdm* [1], including a source representing each of these ancestries. As mentioned in the main text, because there is unlikely to be an unadmixed source of *ANE* ancestry for the Holocene populations of the *FSHG* cline, and also because ADMIXTURE identifies an ancestry component linked to *Tarim_EMBA* populations [66] among populations of the Trans-Ural region of the *FSHG* cline, we also ran a separate set of *qpAdm* to check if the *FSHG* cline could be modeled using a set of younger, Holocene populations high in *ANE* ancestry. Because work in previous sections (VI.A) has shown that populations in Eastern Siberia (i.e. East of the Altai region) are very complex, with contribution from Ancient Paleosiberian ancestry (APS), and also because the *ANE* ancestry in East Siberians appears to derive primarily from APS sources and is thus accounted for, we do not model them using the proximal *qpAdm* framework described here, concentrating instead on the part of the *FSHG* cline including all populations around the Altai mountains and west of it.

For the set of *qpAdm* quantifying *WHG*, *EHG*, *ANE* and East Asian-associated ancestry proportions, we tested all combinations of up to four admixture sources (*Romania_IronGates_HG* representing *WHG* ancestry; *Russia_HG_Elshanks*, an individual from the Volga region ~10,100 years BP representing

*EHG* ancestry; *Russia_AfontovaGora3* representing *ANE* ancestry; and *China_AmurRiver_19K* representing East Asian ancestry). For the set of *qpAdm* quantifying input from more recent sources of ancestry, we tested all combinations of up to five ancestry sources (the same sources for *WHG* and *EHG*, with the addition of *Altai_N, Tarim_EMBA* from [66]*, Iran_N*, and *CHG.SG*). Two older individuals out of the Altai region (I2137 and I13098, >8,000 BP) with radiocarbon dates were chosen out of the *FSHG* cline to constitute the *Altai_N* population. Similarly to the *qpAdm* in Section VI.A but different from those in Section VI.C and D, in models where a particular source is not included, it is moved into the references, in an implementation of the outgroup rotation approach to *qpAdm* [61].

To distinguish between these ancestry sources, we use the following highly discriminating set of at least 12 references: *Sweden_Motala_HG, Russia_Sidelkino_HG.SG, Ukraine_Mesolithic, WestEurasian_UP* (a merged population consisting of Upper Paleolithic West Eurasian individuals from the Kostenki [67] and Sunghir sites [42]), *Yana_UP.SG, Morocco_Iberomaurusian, Italy_North_Villabruna_HG, China_Paleolithic* (a combination of the *Tianyuan* and *China_AmurRiver_Paleolithic_33K* Upper Paleolithic samples [22]), *Spain_ElMiron, Israel_Natufian, Russia_Ust_Ishim_HG*, and *India_GreatAndaman_1000BP.SG*). In the second *qpAdm* set testing for proximal sources of *ANE* ancestry, we also include *Khairygas_16.7kya* and *Ust_Kyakhta_14kya*, for a set of at least 14 references.

***Dataset and Method.*** We used a merged dataset consisting of 541 individuals. All individuals under each population label are listed in Data S2, Table 1. All models were run using `admixr` [55]. Precalculated $f_2$-statistics, used to speed up the process of f4-ratio estimation central to *qpAdm*, were used allowing for maximal missingness = .99.

***Results.*** We find passing models for 84/93 *FSHG* populations at the p>0.01 level as up to four-way admixtures between *WHG*, *EHG*, *ANE* and East Asian ancestries. Hunter-gatherers from west of the Urals trace most of their ancestry from *EHG*, with some minor admixture from *WHG* (<20%). Lower levels of *WHG* ancestry are detected as we move closer to the Urals. *ANE* and East Asian ancestry begins to appear at low levels (<30%) near the Ural foothills. In the Trans-Ural region, *EHG* ancestry continues to decline, while *ANE* and East Asian ancestry increases up to ~80% among populations at the Eastern end of the Trans-Ural steppes. Populations around the Altai region are very homogeneous, deriving almost half their ancestry from East Asian sources, ~45% from an *ANE*-related source, and ~10% from an *EHG*-related source. Populations from the Krasnoyarsk region to the Transbaikal region appear to form a smooth cline where *ANE* ancestry declines and East Asian ancestry increases to ~100% in the Transbaikal region. Our models suggest that *FSHG* populations derive all their ancestry from post-Last Glacial Maximum (LGM) populations (<20 kya) with no ancestry from Upper Paleolithic West or East Eurasian populations such as *Sunghir, Vestonice* or *Tianyuan/China_AmurRiver_Paleolithic_33K*. All models in this *qpAdm* set can be found in Data S2, Table 2.

We find passing models for 57/61 *FSHG* populations west of the Altai region at the p>0.01 level as up to four-way admixtures between *WHG*, *EHG*, *Tarim_EMBA*, and *Altai_N*. Some populations additionally require small ancestry proportions from *Iran_N, CHG.SG*, or East Asian ancestry (represented once again by China_AmurRiver_19K). Importantly, 60/61 populations do not require additional ancestry from *AG3, MA1_HG.SG* or *Yana_UP.SG* in this *qpAdm* setup (the exception being *Russia_Bashkotorstan_Kulmetovo_LN* with a ~12% ancestry proportion from *AG3*) even with these high-*ANE* populations in the references, suggesting that all the *ANE* ancestry of *FSHG* populations can be accounted for by *EHG*, *Tarim_EMBA* and *Altai_N* with no further contribution from older populations. The small quantities of ancestry from *Iran_N* and *CHG.SG* may suggest a low level of gene flow between *FSHG* populations and groups to the South. More importantly, we find that 12/19 populations of the Trans-Ural region and 2 populations from the foothills of the Urals just east of the Trans-Ural region (*Russia_Urals_EBA* and *Russia_Bashkotorstan_Kulmetovo_LN*) require admixture from

*Tarim_EMBA* in all passing models; additionally, models with high p-values for the remaining populations from the Trans-Ural region also tend to involve ancestry from *Tarim_EMBA*. Our results indicate that a *Tarim_EMBA*-related population experienced gene flow with populations from the Trans-Ural region of the *FSHG* cline, and are in line with suggestions that such a hunter-gatherer population with affinities to *Tarim_EMBA* may have occupied regions of southern Central Asia prior to the onset of Neolithic and Bronze Age population movements [14]. Lastly, we find that no populations of the western part of the *FSHG* cline require ancestry from *Khairygas_16.7kya* and *Ust_Kyakhta_14kya*, which suggests that no members of the *FSHG* cline experienced independent gene flow from Northeastern Siberian populations rich in APS ancestry and that all their APS-related ancestry was mediated through the geographically adjacent *Altai_N* (Section V.A). All models in this *qpAdm* set can be found in Data S2, Table 3.

### VII. Population Genetic Analysis: Individuals from Seima-Turbino-period Sites

Our unsupervised analyses indicate that the Seima-Turbino individuals from the sites of Rostovka and Satyga XVI are extremely heterogeneous (SI Section III-IV); however, they also indicate that three major sources of ancestry may suffice to account for the genetic profile of most of the individuals (10/12): 1) *Steppe_MLBA*, *Yakutia_LNBA a*nd ancestry from the *FSHG* cline (SI Sections III, IV; Figure 4E). Because the *FSHG* ancestry of these individuals were not homogeneous (ratios of components maximized in *EHG* individuals and Neolithic individuals of the Altai region vary), the *FSHG* ancestry within the ST individuals at Satyga and Rostovka are unlikely to have derived from a single population. In addition, two other individuals (I32816 and I25555) from Rostovka appear to have low levels of East Asian-associated admixture from sources not closely related to *Yakutia_LNBA*.

We used *qpAdm* [1] to estimate the proportion of *Yakutia_LNBA*, *Steppe_MLBA*, ancestry from the *FSHG* cline, and other West Eurasian and East Asian ancestries among the ST individuals. Due to the heterogeneity of the ST group, we analyzed each individual as a separate target for *qpAdm*. Because individuals vary greatly in coverage, and this exerts an effect on model quality (with simple models passing when individuals have low coverage), we highlight low-coverage individuals in our discussion of the results whenever possible.

We analyzed the ST individuals under two qpAdm setups, which we term the *Distal* and *Proximal qpAdm* setups respectively.

***Distal qpAdms.*** In the first, which we will term the *Distal setup*, we use the same *qpAdm* setup that we used in Section VI.C.ii for *AIEA* individuals, but dropping two sources (*BMAC* and *Germany_EN_LBK*) due to lack of evidence for substantial contribution from these sources in our ADMIXTURE and qpAdm analyses (Figure 4E). In other words, we tested all combinations of up to seven admixture sources drawn from eight possible admixture sources. The eight possible admixture sources are *Steppe_MLBA/Europe_LNBA*, *Hungary_Koros_EN* (a proxy for *WHG* ancestry), *EHG*, *Tyumen_N*, and *Altai_N*. The East Asian sources included *Yakutia_LNBA*, *Mongolia_N* (a proxy for what has been termed *Ancient North Asian* or *ANA* ancestry, from the Mongolian Plateau and the Western Amur region), and *China_YR_MN* (a proxy for East Asian agriculturalist ancestry). We use the same set of 13 references as in Section VI.C.ii: *China_Paleolithic* (Upper Paleolithic hunter-gatherers from China from before 33kya, including the *Tianyuan* individual)*, Iran_GanjDareh_N* (early Iranian agropastoralists)*, Morocco_Iberomaurusian* (hunter-gatherers from Northwest Africa)*, Altai_N_old* (individuals closely related to *Altai_N* but older than 8,000 BP)*, Russia_AfontovaGora3, Italy_South_HG_Ostuni* (Pre-LGM hunter-gatherers from Italy ~27kya)*, Turkey_Boncuklu_N* (early Anatolia farmers)*, Peru_Laramate_900BP* (a proxy for Native American ancestry)*, China_SEastAsia_Island_EN, China_NEastAsia_Inland_EN* (the *Yumin* individual from Mongolia

~8,000 BP), *Syalakh-Belkachi, USA_AK_PaleoAleut,* and *USA_AK_NeoAleut*. As in section VI.C.ii, individuals under each population label for the sources and references are listed in SI Data S7, Table 1. All results for this set of *qpAdm* are listed under SI Data S7, Table 2.

We split the ST individuals into two groups; the first group (including the individuals I32551, I32552, I33398, I25555, and I32369) show substantial fractions of Steppe_MLBA-related ancestry in ADMIXTURE (SI section III, Extended Data Figure 10; Main Text Fig. 4B), and so one of the sources drawn from out of ten possible ancestry sources was always *Russia_Srubnaya* (to represent *Steppe_MLBA* or *Europe_LNBA* ancestry). The rest of the individuals in the second group (I6787, I32545, I32816, I32899, I25558) do not have this restriction, and models are tested which do not include *Russia_Srubnaya* as a source.

***Proximal qpAdms.*** In the second qpAdm, we test for the suitability of *FSHG* ancestry sources from the Trans-Ural region, from localities geographically close to the ST sites, as contributors to the ST individuals. Because of the variation in *FSHG* ancestry source suggested by our unsupervised analyses, we included multiple Late Neolithic and Eneolithic populations from the Trans-Ural steppes radiocarbon dated to ~6.2-4.7 kya (*Russia_Ural_Kulmetovo_C_6.0-6.1kya, Russia_Tobol_Bobrykino_LN_6.2kya, Russia_CombPitWare_Gladunino_C_4.7kya, Russia_MiddleIrtysh_Omsk_C_4.8kya, Russia_CombitPitWare_Borovyanka17_C_5.8-6.1kya*) as sources. Note once again that the C14 dates may appear artificially old due to reservoir effects, and these populations may in reality only shortly precede or even be contemporaneous with the ST individuals (see SI Section VIII). To see if additional ancestry from the FSHG cline apart from these proximal sources is still necessary, *Altai_N* and *Tyumen_N_HG* are now moved into the reference set, rotated in only as part of outgroup rotation.

We also wished to determine the source of the *Yakutia_LNBA*-related ancestry among our ST individuals, as well as to differentiate more clearly between *Yakutia_LNBA* ancestry and other East Asian ancestries among those individuals who require the latter. To that end, we also included *Russia_Tatarka_BA* as a source, moving *Yakutia_LNBA* into the references, rotated in only as part of outgroup rotation.

We list all passing models including Russia_Tatarka_BA and at least one of the proximal FSHG sources in our discussion of the results below. All results for this set of *qpAdm* are listed under SI Data S7, Table 3A.

***Dataset and Method.*** We used a merged dataset consisting of 1648 individuals for *qpAdm* modeling. All individuals under each population label are listed in Data SI Table 3, Column M. All models were run using `admixr` [55]. Precalculated $f_2$-statistics, used to speed up the process of $f_4$-ratio estimation central to *qpAdm*, were used allowing for maximal missingness = .99.

***Results.*** We find passing models for 8/11 ST individuals at the p>0.05 level and 10/11 at the p>0.01 level for the first set of *qpAdm* (without Eneolithic or Early Bronze Age populations from the Trans-Ural Steppes). When these Eneolithic/Early Bronze Age *FSHG* populations are included in the second set of *qpAdm* analyses, we find passing models for all individuals at the p>0.01 level, including for the individual for which no model is found in the first set (I32899). In the following section, we present results for each individual, grouped by notable characteristics in their ancestry. We present results for each set of *qpAdm* analyses in Tables VII. A, VII. B, and VII.C below, restricting models to the simplest model or set of models (fewest sources) that pass at the p>0.05 level, and failing this, at the p>0.01 level.

**Table S34**

| Target | Model | P | Source | Prop | SE | Z |
|---|---|---|---|---|---|---|

| | | | | | | |
|---|---|---|---|---|---|---|
| I32369 | Russia_Srubnaya+Altai_N+Tyumen_HG+Yakutia_LNBA | 0.21 | Altai_N | 0.20 | 0.09 | 2.38E+00 |
| I32369 | Russia_Srubnaya+Altai_N+Tyumen_HG+Yakutia_LNBA | 0.21 | Russia_Srubnaya | 0.44 | 0.03 | 1.49E+01 |
| I32369 | Russia_Srubnaya+Altai_N+Tyumen_HG+Yakutia_LNBA | 0.21 | Tyumen_HG | 0.31 | 0.08 | 3.81E+00 |
| I32369 | Russia_Srubnaya+Altai_N+Tyumen_HG+Yakutia_LNBA | 0.21 | Yakutia_LNBA | 0.05 | 0.03 | 1.48E+00 |
| I32545 | EHG+Yakutia_LNBA | 0.04 | EHG | 0.03 | 0.01 | 2.49E+00 |
| I32545 | EHG+Yakutia_LNBA | 0.04 | Yakutia_LNBA | 0.97 | 0.01 | 7.52E+01 |
| I32551 | Russia_Srubnaya+Yakutia_LNBA | 0.04 | Russia_Srubnaya | 0.71 | 0.04 | 1.96E+01 |
| I32551 | Russia_Srubnaya+Yakutia_LNBA | 0.04 | Yakutia_LNBA | 0.29 | 0.04 | 8.07E+00 |
| I32552 | Russia_Srubnaya+Tyumen_HG+Yakutia_LNBA | 0.66 | Russia_Srubnaya | 0.21 | 0.02 | 1.01E+01 |
| I32552 | Russia_Srubnaya+Tyumen_HG+Yakutia_LNBA | 0.66 | Tyumen_HG | 0.33 | 0.02 | 1.34E+01 |
| I32552 | Russia_Srubnaya+Tyumen_HG+Yakutia_LNBA | 0.66 | Yakutia_LNBA | 0.46 | 0.02 | 2.99E+01 |
| I32552 | Russia_Srubnaya+EHG+Yakutia_LNBA | 0.44 | EHG | 0.46 | 0.03 | 1.44E+01 |
| I32552 | Russia_Srubnaya+EHG+Yakutia_LNBA | 0.44 | Russia_Srubnaya | 0.05 | 0.03 | 1.62E+00 |
| I32552 | Russia_Srubnaya+EHG+Yakutia_LNBA | 0.44 | Yakutia_LNBA | 0.49 | 0.01 | 3.35E+01 |
| I32816 | Altai_N+EHG+Yakutia_LNBA | 0.10 | Altai_N | 0.54 | 0.05 | 1.04E+01 |
| I32816 | Altai_N+EHG+Yakutia_LNBA | 0.10 | EHG | 0.31 | 0.03 | 9.92E+00 |
| I32816 | Altai_N+EHG+Yakutia_LNBA | 0.10 | Yakutia_LNBA | 0.15 | 0.03 | 5.29E+00 |
| I32898 | Russia_Srubnaya+Tyumen_HG+Yakutia_LNBA | 0.33 | Russia_Srubnaya | 0.70 | 0.02 | 3.17E+01 |
| I32898 | Russia_Srubnaya+Tyumen_HG+Yakutia_LNBA | 0.33 | Tyumen_HG | 0.10 | 0.02 | 4.42E+00 |
| I32898 | Russia_Srubnaya+Tyumen_HG+Yakutia_LNBA | 0.33 | Yakutia_LNBA | 0.19 | 0.02 | 1.25E+01 |
| I32898 | Russia_Srubnaya+EHG+Yakutia_LNBA | 0.26 | EHG | 0.13 | 0.03 | 4.21E+00 |
| I32898 | Russia_Srubnaya+EHG+Yakutia_LNBA | 0.26 | Russia_Srubnaya | 0.67 | 0.03 | 2.14E+01 |
| I32898 | Russia_Srubnaya+EHG+Yakutia_LNBA | 0.26 | Yakutia_LNBA | 0.21 | 0.01 | 1.48E+01 |
| I32898 | Russia_Srubnaya+Altai_N+Yakutia_LNBA | 0.15 | Altai_N | 0.10 | 0.03 | 3.75E+00 |
| I32898 | Russia_Srubnaya+Altai_N+Yakutia_LNBA | 0.15 | Russia_Srubnaya | 0.74 | 0.02 | 3.98E+01 |
| I32898 | Russia_Srubnaya+Altai_N+Yakutia_LNBA | 0.15 | Yakutia_LNBA | 0.16 | 0.02 | 7.58E+00 |
| I33398 | Russia_Srubnaya | 0.05 | Russia_Srubnaya | 1.00 | 0.00 | 9.70E+12 |
| I25555 | Russia_Srubnaya+Tyumen_HG+Mongolia_N_North | 0.76 | Mongolia_N_North | 0.22 | 0.02 | 8.97E+00 |
| I25555 | Russia_Srubnaya+Tyumen_HG+Mongolia_N_North | 0.76 | Russia_Srubnaya | 0.33 | 0.04 | 8.50E+00 |
| I25555 | Russia_Srubnaya+Tyumen_HG+Mongolia_N_North | 0.76 | Tyumen_HG | 0.45 | 0.05 | 9.71E+00 |
| I25555 | Russia_Srubnaya+Tyumen_HG+Yakutia_LNBA | 0.48 | Russia_Srubnaya | 0.35 | 0.04 | 8.85E+00 |
| I25555 | Russia_Srubnaya+Tyumen_HG+Yakutia_LNBA | 0.48 | Tyumen_HG | 0.43 | 0.05 | 8.54E+00 |
| I25555 | Russia_Srubnaya+Tyumen_HG+Yakutia_LNBA | 0.48 | Yakutia_LNBA | 0.23 | 0.03 | 7.47E+00 |
| I25555 | Russia_Srubnaya+EHG+Mongolia_N_North | 0.21 | EHG | 0.58 | 0.05 | 1.05E+01 |
| I25555 | Russia_Srubnaya+EHG+Mongolia_N_North | 0.21 | Mongolia_N_North | 0.27 | 0.02 | 1.22E+01 |
| I25555 | Russia_Srubnaya+EHG+Mongolia_N_North | 0.21 | Russia_Srubnaya | 0.16 | 0.05 | 2.87E+00 |
| I25555 | Russia_Srubnaya+EHG+Yakutia_LNBA | 0.10 | EHG | 0.53 | 0.06 | 9.36E+00 |
| I25555 | Russia_Srubnaya+EHG+Yakutia_LNBA | 0.10 | Russia_Srubnaya | 0.19 | 0.05 | 3.49E+00 |
| I25555 | Russia_Srubnaya+EHG+Yakutia_LNBA | 0.10 | Yakutia_LNBA | 0.28 | 0.03 | 1.11E+01 |
| I25558 | Tyumen_HG+China_YR_MN | 0.05 | China_YR_MN | 0.06 | 0.02 | 2.62E+00 |
| I25558 | Tyumen_HG+China_YR_MN | 0.05 | Tyumen_HG | 0.94 | 0.02 | 4.34E+01 |
| I6787 | Altai_N+Tyumen_HG+Hungary_EN_HG_Koros+Yakutia_LNBA | 0.20 | Altai_N | 0.36 | 0.09 | 4.17E+00 |
| I6787 | Altai_N+Tyumen_HG+Hungary_EN_HG_Koros+Yakutia_LNBA | 0.20 | Tyumen_HG | 0.22 | 0.08 | 2.78E+00 |
| I6787 | Altai_N+Tyumen_HG+Hungary_EN_HG_Koros+Yakutia_LNBA | 0.20 | Hungary_EN_HG_Koros | 0.26 | 0.03 | 9.87E+00 |

| | | | | | | |
|---|---|---|---|---|---|---|
| I6787 | Altai_N+Tyumen_HG+Hungary_EN_HG_Koros+Yakutia_LNBA | 0.20 | Yakutia_LNBA | 0.17 | 0.04 | 4.68E+00 |
| I6787 | Altai_N+Hungary_EN_HG_Koros+EHG+Yakutia_LNBA | 0.19 | Altai_N | 0.38 | 0.07 | 5.49E+00 |
| I6787 | Altai_N+Hungary_EN_HG_Koros+EHG+Yakutia_LNBA | 0.19 | Hungary_EN_HG_Koros | 0.15 | 0.04 | 3.60E+00 |
| I6787 | Altai_N+Hungary_EN_HG_Koros+EHG+Yakutia_LNBA | 0.19 | EHG | 0.29 | 0.07 | 3.92E+00 |
| I6787 | Altai_N+Hungary_EN_HG_Koros+EHG+Yakutia_LNBA | 0.19 | Yakutia_LNBA | 0.17 | 0.04 | 4.67E+00 |
| I32899 | Tyumen_HG+EHG+Yakutia_LNBA | 0.008 | EHG | 0.58 | 0.06 | 9.74E+00 |
| I32899 | Tyumen_HG+EHG+Yakutia_LNBA | 0.008 | Tyumen_HG | 0.41 | 0.06 | 6.47E+00 |
| I32899 | Tyumen_HG+EHG+Yakutia_LNBA | 0.008 | Yakutia_LNBA | 0.00 | 0.01 | 1.50E-01 |

**Table S1-VII. A |** Passing models for ST individuals with references used for AIEA populations in section VI.C.ii, and a subset of the sources used for AIEA populations in section VI.C.ii (dropping *BMAC* and *Germany_EN_LBK*). For individuals with models passing at the p>0.05 level, all models are listed; failing that, models with 0.01 < p < 0.05 is listed (I32545). No model passes for I32899 at the p>0.01 level; the model with the greatest p-value is listed. No simpler models pass.
**References:** [*China_Paleolithic, Iran_GanjDareh_N, Morocco_Iberomaurusian, Altai_N_old, Russia_AfontovaGora3, Italy_South_HG_Ostuni, Turkey_Boncuklu_N, Peru_Laramate_900BP, China_SEastAsia_Island_EN, China_NEastAsia_Inland_EN, Syalakh-Belkachi, USA_AK_PaleoAleut, USA_AK_NeoAleut*]
**Sources:** [*Russia_Srubnaya, Hungary_Koros_EN, EHG, Tyumen_N, Altai_N, Yakutia_LNBA, Mongolia_N, China_YR_MN*]

#### Table S35

| Target | Model | P | Source | Prop | SE | Z |
|---|---|---|---|---|---|---|
| I32369 | Russia_Srubnaya + Altai_N + Russia_Tatarka_BA + Russia_Tobol_Boborykino_LN_6.2kya | 0.33 | Altai_N | 0.10 | 0.06 | 1.73E+00 |
| I32369 | Russia_Srubnaya + Altai_N + Russia_Tatarka_BA + Russia_Tobol_Boborykino_LN_6.2kya | 0.33 | Russia_Tobol_Boborykino_LN_6.2kya | 0.51 | 0.06 | 8.79E+00 |
| I32369 | Russia_Srubnaya + Altai_N + Russia_Tatarka_BA + Russia_Tobol_Boborykino_LN_6.2kya | 0.33 | Russia_Tatarka_BA | 0.09 | 0.02 | 4.13E+00 |
| I32369 | Russia_Srubnaya + Altai_N + Russia_Tatarka_BA + Russia_Tobol_Boborykino_LN_6.2kya | 0.33 | Russia_Srubnaya | 0.30 | 0.03 | 9.36E+00 |
| I32369 | Russia_Srubnaya + EHG + Russia_Tatarka_BA + Russia_CombPitWare_Borovyanka17_C_5.8-6.1kya | 0.13 | EHG | 0.25 | 0.10 | 2.40E+00 |
| I32369 | Russia_Srubnaya + EHG + Russia_Tatarka_BA + Russia_CombPitWare_Borovyanka17_C_5.8-6.1kya | 0.13 | Russia_CombPitWare_Borovyanka17_C_5.8-6.1kya | 0.38 | 0.07 | 5.25E+00 |
| I32369 | Russia_Srubnaya + EHG + Russia_Tatarka_BA + Russia_CombPitWare_Borovyanka17_C_5.8-6.1kya | 0.13 | Russia_Tatarka_BA | 0.06 | 0.02 | 3.94E+00 |
| I32369 | Russia_Srubnaya + EHG + Russia_Tatarka_BA + Russia_CombPitWare_Borovyanka17_C_5.8-6.1kya | 0.13 | Russia_Srubnaya | 0.31 | 0.05 | 6.27E+00 |
| I32369 | Russia_Srubnaya + Altai_N + Russia_Tatarka_BA + Russia_MiddleIrtysh_Omsk_C_4.8kya | 0.07 | Altai_N | 0.13 | 0.05 | 2.50E+00 |
| I32369 | Russia_Srubnaya + Altai_N + Russia_Tatarka_BA + Russia_MiddleIrtysh_Omsk_C_4.8kya | 0.07 | Russia_MiddleIrtysh_Omsk_C_4.8kya | 0.43 | 0.05 | 8.54E+00 |
| I32369 | Russia_Srubnaya + Altai_N + Russia_Tatarka_BA + Russia_MiddleIrtysh_Omsk_C_4.8kya | 0.07 | Russia_Tatarka_BA | 0.05 | 0.02 | 2.70E+00 |
| I32369 | Russia_Srubnaya + Altai_N + Russia_Tatarka_BA + Russia_MiddleIrtysh_Omsk_C_4.8kya | 0.07 | Russia_Srubnaya | 0.38 | 0.02 | 1.54E+01 |
| I32545 | Hungary_EN_HG_Koros + Russia_Tatarka_BA | 0.06 | Hungary_EN_HG_Koros | 0.03 | 0.01 | 2.70E+00 |
| I32545 | Hungary_EN_HG_Koros + Russia_Tatarka_BA | 0.06 | Russia_Tatarka_BA | 0.97 | 0.01 | 8.95E+01 |
| I32551 | Russia_Srubnaya + Russia_Tatarka_BA | 0.07 | Russia_Tatarka_BA | 0.28 | 0.03 | 9.44E+00 |
| I32551 | Russia_Srubnaya + Russia_Tatarka_BA | 0.07 | Russia_Srubnaya | 0.72 | 0.03 | 2.43E+01 |
| I32552 | Russia_Srubnaya + Russia_Tatarka_BA + Russia_Tobol_Boborykino_LN_6.2kya | 0.60 | Russia_Tobol_Boborykino_LN_6.2kya | 0.38 | 0.03 | 1.36E+01 |
| I32552 | Russia_Srubnaya + Russia_Tatarka_BA + Russia_Tobol_Boborykino_LN_6.2kya | 0.60 | Russia_Tatarka_BA | 0.48 | 0.01 | 3.63E+01 |
| I32552 | Russia_Srubnaya + Russia_Tatarka_BA + Russia_Tobol_Boborykino_LN_6.2kya | 0.60 | Russia_Srubnaya | 0.15 | 0.03 | 5.44E+00 |
| I32552 | Russia_Srubnaya + Russia_Tatarka_BA + Russia_Ural_Kulmetovo_C_6.0-6.1kya | 0.10 | Russia_Ural_Kulmetovo_C_6.0-6.1kya | 0.36 | 0.03 | 1.35E+01 |
| I32552 | Russia_Srubnaya + Russia_Tatarka_BA + Russia_Ural_Kulmetovo_C_6.0-6.1kya | 0.10 | Russia_Tatarka_BA | 0.48 | 0.01 | 3.62E+01 |
| I32552 | Russia_Srubnaya + Russia_Tatarka_BA + Russia_Ural_Kulmetovo_C_6.0-6.1kya | 0.10 | Russia_Srubnaya | 0.16 | 0.03 | 6.45E+00 |

| | | | | | | |
|---|---|---|---|---|---|---|
| I32552 | Russia_Srubnaya + Russia_Tatarka_BA + Russia_CombPitWare_Gladunino_C_4.7kya | 0.06 | Russia_CombPitWare_Gladunino_C_4.7kya | 0.38 | 0.03 | 1.32E+01 |
| I32552 | Russia_Srubnaya + Russia_Tatarka_BA + Russia_CombPitWare_Gladunino_C_4.7kya | 0.06 | Russia_Tatarka_BA | 0.45 | 0.01 | 3.20E+01 |
| I32552 | Russia_Srubnaya + Russia_Tatarka_BA + Russia_CombPitWare_Gladunino_C_4.7kya | 0.06 | Russia_Srubnaya | 0.17 | 0.03 | 6.37E+00 |
| I32816 | Altai_N + Russia_Tatarka_BA + Russia_Tobol_Boborykino_LN_6.2kya | 0.31 | Altai_N | 0.45 | 0.05 | 8.53E+00 |
| I32816 | Altai_N + Russia_Tatarka_BA + Russia_Tobol_Boborykino_LN_6.2kya | 0.31 | Russia_Tobol_Boborykino_LN_6.2kya | 0.37 | 0.04 | 1.03E+01 |
| I32816 | Altai_N + Russia_Tatarka_BA + Russia_Tobol_Boborykino_LN_6.2kya | 0.31 | Russia_Tatarka_BA | 0.19 | 0.02 | 8.23E+00 |
| I32816 | Russia_Srubnaya + Yakutia_LNBA + Russia_CombPitWare_Borovyanka17_C_5.8-6.1kya | 0.29 | Russia_CombPitWare_Borovyanka17_C_5.8-6.1kya | 0.73 | 0.03 | 2.89E+01 |
| I32816 | Russia_Srubnaya + Yakutia_LNBA + Russia_CombPitWare_Borovyanka17_C_5.8-6.1kya | 0.29 | Russia_Srubnaya | 0.04 | 0.02 | 1.92E+00 |
| I32816 | Russia_Srubnaya + Yakutia_LNBA + Russia_CombPitWare_Borovyanka17_C_5.8-6.1kya | 0.29 | Yakutia_LNBA | 0.23 | 0.02 | 1.29E+01 |
| I32816 | Altai_N + Russia_Tatarka_BA + Russia_MiddleIrtysh_Omsk_C_4.8kya | 0.21 | Altai_N | 0.41 | 0.06 | 7.29E+00 |
| I32816 | Altai_N + Russia_Tatarka_BA + Russia_MiddleIrtysh_Omsk_C_4.8kya | 0.21 | Russia_MiddleIrtysh_Omsk_C_4.8kya | 0.41 | 0.04 | 1.03E+01 |
| I32816 | Altai_N + Russia_Tatarka_BA + Russia_MiddleIrtysh_Omsk_C_4.8kya | 0.21 | Russia_Tatarka_BA | 0.18 | 0.02 | 8.07E+00 |
| I32816 | Russia_Srubnaya + Russia_Tatarka_BA + Russia_CombPitWare_Borovyanka17_C_5.8-6.1kya | 0.19 | Russia_CombPitWare_Borovyanka17_C_5.8-6.1kya | 0.74 | 0.02 | 3.09E+01 |
| I32816 | Russia_Srubnaya + Russia_Tatarka_BA + Russia_CombPitWare_Borovyanka17_C_5.8-6.1kya | 0.19 | Russia_Tatarka_BA | 0.21 | 0.01 | 1.50E+01 |
| I32816 | Russia_Srubnaya + Russia_Tatarka_BA + Russia_CombPitWare_Borovyanka17_C_5.8-6.1kya | 0.19 | Russia_Srubnaya | 0.05 | 0.02 | 2.35E+00 |
| I32816 | Hungary_EN_HG_Koros + Russia_Tatarka_BA + Russia_CombPitWare_Borovyanka17_C_5.8-6.1kya | 0.18 | Hungary_EN_HG_Koros | 0.04 | 0.02 | 2.19E+00 |
| I32816 | Hungary_EN_HG_Koros + Russia_Tatarka_BA + Russia_CombPitWare_Borovyanka17_C_5.8-6.1kya | 0.18 | Russia_CombPitWare_Borovyanka17_C_5.8-6.1kya | 0.75 | 0.02 | 3.38E+01 |
| I32816 | Hungary_EN_HG_Koros + Russia_Tatarka_BA + Russia_CombPitWare_Borovyanka17_C_5.8-6.1kya | 0.18 | Russia_Tatarka_BA | 0.21 | 0.01 | 1.48E+01 |
| I32816 | EHG + Russia_Tatarka_BA + Russia_CombPitWare_Borovyanka17_C_5.8-6.1kya | 0.08 | EHG | 0.05 | 0.04 | 1.19E+00 |
| I32816 | EHG + Russia_Tatarka_BA + Russia_CombPitWare_Borovyanka17_C_5.8-6.1kya | 0.08 | Russia_CombPitWare_Borovyanka17_C_5.8-6.1kya | 0.74 | 0.05 | 1.56E+01 |
| I32816 | EHG + Russia_Tatarka_BA + Russia_CombPitWare_Borovyanka17_C_5.8-6.1kya | 0.08 | Russia_Tatarka_BA | 0.22 | 0.01 | 1.46E+01 |
| I32898 | Russia_Srubnaya + Russia_Tatarka_BA + Russia_Ural_Kulmetovo_C_6.0-6.1kya | 0.22 | Russia_Ural_Kulmetovo_C_6.0-6.1kya | 0.10 | 0.02 | 4.03E+00 |
| I32898 | Russia_Srubnaya + Russia_Tatarka_BA + Russia_Ural_Kulmetovo_C_6.0-6.1kya | 0.22 | Russia_Tatarka_BA | 0.21 | 0.01 | 1.61E+01 |
| I32898 | Russia_Srubnaya + Russia_Tatarka_BA + Russia_Ural_Kulmetovo_C_6.0-6.1kya | 0.22 | Russia_Srubnaya | 0.70 | 0.02 | 2.82E+01 |
| I32898 | Russia_Srubnaya + Russia_Tatarka_BA + Russia_CombPitWare_Gladunino_C_4.7kya | 0.19 | Russia_CombPitWare_Gladunino_C_4.7kya | 0.10 | 0.03 | 3.92E+00 |
| I32898 | Russia_Srubnaya + Russia_Tatarka_BA + Russia_CombPitWare_Gladunino_C_4.7kya | 0.19 | Russia_Tatarka_BA | 0.20 | 0.01 | 1.49E+01 |
| I32898 | Russia_Srubnaya + Russia_Tatarka_BA + Russia_CombPitWare_Gladunino_C_4.7kya | 0.19 | Russia_Srubnaya | 0.70 | 0.03 | 2.78E+01 |
| I32898 | Russia_Srubnaya + Russia_Tatarka_BA + Russia_Tobol_Boborykino_LN_6.2kya | 0.19 | Russia_Tobol_Boborykino_LN_6.2kya | 0.10 | 0.02 | 3.97E+00 |
| I32898 | Russia_Srubnaya + Russia_Tatarka_BA + Russia_Tobol_Boborykino_LN_6.2kya | 0.19 | Russia_Tatarka_BA | 0.21 | 0.01 | 1.63E+01 |
| I32898 | Russia_Srubnaya + Russia_Tatarka_BA + Russia_Tobol_Boborykino_LN_6.2kya | 0.19 | Russia_Srubnaya | 0.69 | 0.03 | 2.71E+01 |
| I32898 | Russia_Srubnaya + Russia_Tatarka_BA + Russia_MiddleIrtysh_Omsk_C_4.8kya | 0.18 | Russia_MiddleIrtysh_Omsk_C_4.8kya | 0.09 | 0.02 | 3.95E+00 |
| I32898 | Russia_Srubnaya + Russia_Tatarka_BA + Russia_MiddleIrtysh_Omsk_C_4.8kya | 0.18 | Russia_Tatarka_BA | 0.20 | 0.01 | 1.55E+01 |
| I32898 | Russia_Srubnaya + Russia_Tatarka_BA + Russia_MiddleIrtysh_Omsk_C_4.8kya | 0.18 | Russia_Srubnaya | 0.71 | 0.02 | 3.21E+01 |
| I32898 | Russia_Srubnaya + Russia_Tatarka_BA + Russia_CombPitWare_Borovyanka17_C_5.8-6.1kya | 0.14 | Russia_CombPitWare_Borovyanka17_C_5.8-6.1kya | 0.08 | 0.02 | 3.89E+00 |
| I32898 | Russia_Srubnaya + Russia_Tatarka_BA + Russia_CombPitWare_Borovyanka17_C_5.8-6.1kya | 0.14 | Russia_Tatarka_BA | 0.19 | 0.01 | 1.39E+01 |

| | | | | | | |
|---|---|---|---|---|---|---|
| I32898 | Russia_Srubnaya + Russia_Tatarka_BA + Russia_CombPitWare_Borovyanka17_C_5.8-6.1kya | 0.14 | Russia_Srubnaya | 0.72 | 0.02 | 3.63E+01 |
| I32899 | EHG + Russia_Tatarka_BA + Russia_Ural_Kulmetovo_C_6.0-6.1kya | 0.05 | EHG | 0.21 | 0.14 | 1.51E+00 |
| I32899 | EHG + Russia_Tatarka_BA + Russia_Ural_Kulmetovo_C_6.0-6.1kya | 0.05 | Russia_Ural_Kulmetovo_C_6.0-6.1kya | 0.77 | 0.14 | 5.56E+00 |
| I32899 | EHG + Russia_Tatarka_BA + Russia_Ural_Kulmetovo_C_6.0-6.1kya | 0.05 | Russia_Tatarka_BA | 0.03 | 0.01 | 2.40E+00 |
| I33398 | Russia_Srubnaya | 0.05 | Russia_Srubnaya | 1.00 | 0.00 | 9.70E+12 |
| I25555 | Russia_Srubnaya + China_YR_MN + Russia_CombPitWare_Gladunino_C_4.7kya | 0.84 | China_YR_MN | 0.23 | 0.03 | 9.06E+00 |
| I25555 | Russia_Srubnaya + China_YR_MN + Russia_CombPitWare_Gladunino_C_4.7kya | 0.84 | Russia_CombPitWare_Gladunino_C_4.7kya | 0.54 | 0.05 | 1.12E+01 |
| I25555 | Russia_Srubnaya + China_YR_MN + Russia_CombPitWare_Gladunino_C_4.7kya | 0.84 | Russia_Srubnaya | 0.23 | 0.04 | 5.02E+00 |
| I25558 | EHG + Russia_Tatarka_BA + Russia_MiddleIrtysh_Omsk_C_4.8kya | 0.25 | EHG | 0.18 | 0.11 | 1.74E+00 |
| I25558 | EHG + Russia_Tatarka_BA + Russia_MiddleIrtysh_Omsk_C_4.8kya | 0.25 | Russia_MiddleIrtysh_Omsk_C_4.8kya | 0.77 | 0.11 | 7.00E+00 |
| I25558 | EHG + Russia_Tatarka_BA + Russia_MiddleIrtysh_Omsk_C_4.8kya | 0.25 | Russia_Tatarka_BA | 0.05 | 0.02 | 2.75E+00 |
| I6787 | Hungary_EN_HG_Koros + Russia_Tatarka_BA + Russia_CombPitWare_Borovyanka17_C_5.8-6.1kya | 0.09 | Hungary_EN_HG_Koros | 0.24 | 0.02 | 1.25E+01 |
| I6787 | Hungary_EN_HG_Koros + Russia_Tatarka_BA + Russia_CombPitWare_Borovyanka17_C_5.8-6.1kya | 0.09 | Russia_Tatarka_BA | 0.20 | 0.02 | 1.23E+01 |
| I6787 | Hungary_EN_HG_Koros + Russia_Tatarka_BA + Russia_CombPitWare_Borovyanka17_C_5.8-6.1kya | 0.09 | Russia_CombPitWare_Borovyanka17_C_5.8-6.1kya | 0.57 | 0.02 | 2.51E+01 |
| I6787 | Hungary_EN_HG_Koros + Yakutia_LNBA + Russia_CombPitWare_Borovyanka17_C_5.8-6.1kya | 0.06 | Hungary_EN_HG_Koros | 0.23 | 0.02 | 1.20E+01 |
| I6787 | Hungary_EN_HG_Koros + Yakutia_LNBA + Russia_CombPitWare_Borovyanka17_C_5.8-6.1kya | 0.06 | Yakutia_LNBA | 0.21 | 0.02 | 1.06E+01 |
| I6787 | Hungary_EN_HG_Koros + Yakutia_LNBA + Russia_CombPitWare_Borovyanka17_C_5.8-6.1kya | 0.06 | Russia_CombPitWare_Borovyanka17_C_5.8-6.1kya | 0.56 | 0.02 | 2.35E+01 |
| I6787 | Altai_N + Hungary_EN_HG_Koros + Russia_Tatarka_BA + Russia_MiddleIrtysh_Omsk_C_4.8kya | 0.55 | Altai_N | 0.36 | 0.05 | 6.59E+00 |
| I6787 | Altai_N + Hungary_EN_HG_Koros + Russia_Tatarka_BA + Russia_MiddleIrtysh_Omsk_C_4.8kya | 0.55 | Hungary_EN_HG_Koros | 0.24 | 0.02 | 1.11E+01 |
| I6787 | Altai_N + Hungary_EN_HG_Koros + Russia_Tatarka_BA + Russia_MiddleIrtysh_Omsk_C_4.8kya | 0.55 | Russia_Tatarka_BA | 0.16 | 0.02 | 7.17E+00 |
| I6787 | Altai_N + Hungary_EN_HG_Koros + Russia_Tatarka_BA + Russia_MiddleIrtysh_Omsk_C_4.8kya | 0.55 | Russia_MiddleIrtysh_Omsk_C_4.8kya | 0.24 | 0.05 | 4.96E+00 |
| I6787 | Altai_N + Hungary_EN_HG_Koros + Russia_Tatarka_BA + Russia_Ural_Kulmetovo_C_6.0-6.1kya | 0.46 | Altai_N | 0.38 | 0.05 | 7.42E+00 |
| I6787 | Altai_N + Hungary_EN_HG_Koros + Russia_Tatarka_BA + Russia_Ural_Kulmetovo_C_6.0-6.1kya | 0.46 | Hungary_EN_HG_Koros | 0.22 | 0.02 | 8.83E+00 |
| I6787 | Altai_N + Hungary_EN_HG_Koros + Russia_Tatarka_BA + Russia_Ural_Kulmetovo_C_6.0-6.1kya | 0.46 | Russia_Tatarka_BA | 0.16 | 0.02 | 7.05E+00 |
| I6787 | Altai_N + Hungary_EN_HG_Koros + Russia_Tatarka_BA + Russia_Ural_Kulmetovo_C_6.0-6.1kya | 0.46 | Russia_Ural_Kulmetovo_C_6.0-6.1kya | 0.24 | 0.05 | 4.84E+00 |
| I6787 | Altai_N + Hungary_EN_HG_Koros + Russia_Tatarka_BA + Russia_Tobol_Boborykino_LN_6.2kya | 0.28 | Altai_N | 0.37 | 0.06 | 6.39E+00 |
| I6787 | Altai_N + Hungary_EN_HG_Koros + Russia_Tatarka_BA + Russia_Tobol_Boborykino_LN_6.2kya | 0.28 | Hungary_EN_HG_Koros | 0.21 | 0.03 | 7.62E+00 |
| I6787 | Altai_N + Hungary_EN_HG_Koros + Russia_Tatarka_BA + Russia_Tobol_Boborykino_LN_6.2kya | 0.28 | Russia_Tatarka_BA | 0.17 | 0.03 | 6.89E+00 |
| I6787 | Altai_N + Hungary_EN_HG_Koros + Russia_Tatarka_BA + Russia_Tobol_Boborykino_LN_6.2kya | 0.28 | Russia_Tobol_Boborykino_LN_6.2kya | 0.25 | 0.06 | 4.55E+00 |
| I6787 | Altai_N + Hungary_EN_HG_Koros + Russia_Tatarka_BA + Russia_CombPitWare_Gladunino_C_4.7kya | 0.19 | Altai_N | 0.38 | 0.06 | 6.20E+00 |
| I6787 | Altai_N + Hungary_EN_HG_Koros + Russia_Tatarka_BA + Russia_CombPitWare_Gladunino_C_4.7kya | 0.19 | Hungary_EN_HG_Koros | 0.22 | 0.03 | 7.93E+00 |

| | | | | | | |
|---|---|---|---|---|---|---|
| I6787 | Altai_N + Hungary_EN_HG_Koros + Russia_Tatarka_BA + Russia_CombPitWare_Gladunino_C_4.7kya | 0.19 | Russia_Tatarka_BA | 0.15 | 0.02 | 6.45E+00 |
| I6787 | Altai_N + Hungary_EN_HG_Koros + Russia_Tatarka_BA + Russia_CombPitWare_Gladunino_C_4.7kya | 0.19 | Russia_CombPitWare_Gladunino_C_4.7kya | 0.25 | 0.06 | 4.02E+00 |

**Table S1-VII. B |** Passing models for ST individuals with a modified list of the references used for AIEA populations in section VI.C.ii (adding *Altai_N*, *Tyumen_N* and *Yakutia_LNBA*), and modified list of the sources used for AIEA populations in section VI.C.ii (dropping *BMAC* and *Germany_EN_LBK*, and adding the Samus' population and 5 FSHG sources from the Trans-Ural region). For individuals with models passing at the p>0.05 level, all models are listed; failing that, the models with the highest p value < 0.05 is listed (I32899). Passing models are found for all individuals. No simpler models pass.

**References:** [*China_Paleolithic, Iran_GanjDareh_N, Morocco_Iberomaurusian, Altai_N_old, Russia_AfontovaGora3, Italy_South_HG_Ostuni, Turkey_Boncuklu_N, Peru_Laramate_900BP, China_SEastAsia_Island_EN, China_NEastAsia_Inland_EN, Syalakh-Belkachi, USA_AK_PaleoAleut, USA_AK_NeoAleut, Altai_N, Tyumen_N, Yakutia_LNBA*]

**Sources:** [*Russia_Srubnaya, Hungary_Koros_EN, EHG, Tyumen_N, Altai_N, Yakutia_LNBA, Mongolia_N, China_YR_MN, Russia_Tatarka_BA, Russia_Tobol_Boborykino_LN_6.2kya, Russia_CombPitWare_Gladunino_C_4.7kya, Russia_MiddleIrtysh_Omsk_C_4.8kya, Russia_CombPitWare_Borovyanka17_C_5.8-6.1kya, Russia_Ural_Kulmetovo_C_6.0-6.1kya*]

***Discussion.*** We find that the *qpAdm* results are broadly congruent with those of unsupervised analyses (PCA and ADMIXTURE) and discuss them in context. First, we find in all cases that ST individuals can derive all their *Yakutia_LNBA* ancestry from a Samus'-related population, even with *Yakutia_LNBA* itself moved into the set of references. This implies that *Yakutia_LNBA* ancestry in the ST individuals could be mediated by a source closely related to the population in the Samus' culture, without direct admixture from populations with such ancestry from further further afield in East Siberia. Second, we find that FSHG sources from the Trans-Ural region suffice to account for the FSHG ancestry among many ST individuals, with at least one model passing for most individuals that has FSHG ancestry deriving near-completely from the local FSHG sources we used in the proximal qpAdm setup. Lastly, we find connections further afield to the far west (WHG ancestry) and the far east (non-Yakutia_LNBA East Asian ancestry).

We group individuals into 7 groups: unadmixed representatives of three major sources of ancestry (*Yakutia_LNBA*, *Steppe_MLBA*, *FSHG*), admixed individuals representing 2-way (between *FSHG* and *Steppe_MLBA*) and 3-way (between *FSHG*, *Steppe_MLBA*, and *Yakutia_LNBA*) admixtures, individuals requiring other East Asian-associated ancestry sources than *Yakutia_LNBA*, and the individual (I6787) requiring *WHG* ancestry.

***Individuals with near-unadmixed Yakutia_LNBA ancestry.***

***I32545.*** We find that this individual can be modeled with near-unadmixed ancestry from *Yakutia_LNBA* (~97% *Yakutia_LNBA*) at the p>0.01 level; this is improved with the addition of *Russia_Tatarka_BA* in the second *qpAdm* setup, even with *Yakutia_LNBA* moved to the references (~97% *Russia_Tatarka_BA*, p=0.06). All passing models for this individual include a *Yakutia_LNBA*-related source.

***Individuals with near-unadmixed Steppe_MLBA ancestry.***

***I33398.*** We find that this individual can be modeled as an unadmixed representative of a *Steppe_MLBA*-related source (100% *Russia_Srubnaya*, p=0.053), in keeping with his behavior in PCA and ADMIXTURE (Main Text Fig. 1 center, 4B; Extended Data Fig. 4, 5).

***Indivduals with near-unadmixed FSHG ancestry.***

***I25558.*** We find that this individual can be modeled with near-complete descent from *Tyumen_N_HG* (~93%); when populations from the Eneolithic Trans-Ural steppe are added, this individual can be modeled with complete descent from *Russia_CombPitWare_Gladunino_C_4.7kya* (~100%, p=0.25), in keeping with her behavior in PCA and ADMIXTURE (Main Text Fig. 1 center, 4B; Extended Data Fig. 4, 5).

***I32899.*** No models pass for this individual in the first set of *qpAdm*s (highest p=0.007), but when populations of the Eneoltihic Trans-Ural steppe are included, a model with 77% *Russia_Ural_Kulmetovo_C_6.0-6.1kya* + 23% *EHG* (p = 0.25) passes.

***Individuals with 2-way admixture between FSHG ancestry and Steppe_MLBA ancestry.***

***I32369.*** This individual can be modeled as a 3-way admixture with major fractions of ancestry from a *Steppe_MLBA*-related source (~30-44%) and *FSHG* ancestry (~51-61%) and minor contributions from an East-Asian related source, in keeping with his behavior in PCA and ADMIXTURE (Main Text Fig. 1 center, 4B; Extended Data Fig. 4, 5).

***Individuals with 3-way admixture between FSHG ancestry, Steppe_MLBA ancestry, and Yakutia_MLBA ancestry.***

***I32551_d.*** This individual can be modeled as a product of a two-way admixture (~30% *Yakutia_LNBA* + ~70% *Srubnaya_LNBA*) in the first set of *qpAdm*s at the p>0.01 level (p=0.04). A similar model passes when *FSHG* populations of the Eneolithic Trans-Ural steppe are added, with *Russia_Tatarka_BA* as the *Yakutia_LNBA*-related source (p=0.07). All passing models for this individual include a *Yakutia_LNBA*-related source.

***I32552.*** In the first set of *qpAdm*, this individual can be modeled as three-way admixtures between ~47% *Yakutia_LNBA* + ~5-20% *Srubnaya_LNBA*, with the other two sources drawn from the *FSHG* cline (p>0.44). These models are in keeping with the behavior of this individual in ADMIXTURE (Fig. 4B, Extended Data Figure 10). In the second set of *qpAdm*s, with populations of the Eneolithic/Early Bronze Age Trans-Ural Steppe as sources, 3-way admixture models pass (p>0.10), with ~40% each of *Russia_Tatarka_BA*-related and *FSHG*-related ancestry, and ~15-20% ancestry from the *Steppe_MLBA*-related source. All passing models for this individual include a *Yakutia_LNBA*-related source.

***I32898.*** In the first set of *qpAdm*, a set of 3-way admixture models pass (p>0.15), with sources drawn from the *FSHG* cline contributing ~10%, *Yakutia_LNBA* ancestry sources contributing ~20%, and *Steppe_MLBA* ancestry contributing ~70% of ancestry. A similar set of models pass in the second set of qpAdms, with ~10% ancestry from the *FSHG* cline, East Asian ancestry from *Russia_Tatarka_BA* (~20%), with the remainder coming from a *Steppe_MLBA*-related source (~70%; p>0.14).

***Individuals with other East Asian ancestry.***

***I25555.*** 4 3-way models pass for I25555 in the first set of *qpAdm*, involving a substantial fraction of ancestry from the FSHG cline (either ~45%Tyumen_HG or ~55% EHG), Steppe_MLBA ancestry (~15-35%) and either Mongolia_N or Yakutia_LNBA ancestry (~22-28%; p>0.10). A wide range of 4-way models pass in the proximal qpAdm, with either 2 FSHG sources or 2 East Asian sources; *Yakutia_LNBA*-related sources, *Mongolia_N_North*, and *China_YR_MN* can individually or in combination account for the East Asian ancestry of this individual. However, the range of models that pass contrasts with results of ADMIXTURE (Extended Data Figure 10, Fig. 4B), where this individual appears to possess ancestry from the *FSHG* cline alongside a complex mix of East Asian ancestries, which may be due to this individual's low coverage (0.04x). As the main East Asian source for this individual in ADMIXTURE is *Cisbaikal_LNBA*, we repeated the proximal qpAdm for all ST individuals with *Cisbaikal_LNBA* added to the references, rotated into the sources. We find a wide range of passing models for I25555, but once again with either *Cisbaikal_LNBA*, *Mongolia_N_North* or *Yakutia_LNBA* sufficing for this individual's East Asian ancestry. We suggest once again that this may be due to the poor coverage. All models containing *Cisbaikal_LNBA* in the sources and references can be found in SI Data S7, Table 3B.

***I32816.*** A 3-way model passes for I32816 in the first set of *qpAdm* (~54% *Altai_N* + ~30% *EHG* + ~15% *Yakutia_LNBA*; p=0.09); this model is congruent with the behavior of this individual in

ADMIXTURE (Fig. 4B). A similar set of models pass in the proximal qpAdms, with multiple proximal FSHG sources equally sufficing for all of this individual's FSHG ancestry. However, as ADMIXTURE also shows that this individual possesses significant *Cisbaikal_LNBA* ancestry, we repeated the proximal qpAdm with *Cisbaikal_LNBA* added to the references, rotated into the sources. We find a wide range of passing 3-way and 4-way models for I32816, but now all passing models involve *Cisbaikal_LNBA* ancestry rotated into the sources at ~15-20%. We selected the simplest (4-way) qpAdm model containing ancestry from both *Russia_Tatarka_BA* and *Cisbaikal_LNBA* ancestry with the highest p-value for Figure 4B. All models containing *Cisbaikal_LNBA* in the sources and references can be found in SI Data S7, Table 3B.

***An Individual with WHG ancestry.***

***I6787.*** We find in the proximal setup that the simplest passing qpAdm models involved 4 sources, all including two sources of FSHG ancestry (*EHG* and *Tyumen_N* or *Altai_N* and *Tyumen_N*), a source of *Yakutia_LNBA* ancestry, and surprisingly a source of *WHG* ancestry (*Hungary_EN_HG_Koros*)—a result in line with ADMIXTURE (Figure 4B). Under the proximal setup, the same phenomenon recurs, but the presence of proximal sources of FSHG ancestry allows for simpler 3-way models to pass where a single FSHG source suffices for all the individual's FSHG ancestry. The presence of WHG ancestry suggests connections far to the West, to regions close to the Baltic sea at this time.

### VIII. Y-chromosomes Haplogroups

In the ancient DNA record during the time period that our study covers (~10-3.5 kya), the phylogenetically closest Individuals upstream of the Y-chromosomal haplogroup found at high levels among modern Kets (Q-YP1691, with a TMRCA ~2.0kya) [68,69] are individuals carrying the subclades Q-YP1102 and Q-Y11236, which have a TMRCA in the mid-Holocene ~4.0kya [70]. Three individuals falling within the time period of our study carry these subclades: irk022.SG (Russia_Cisbaikal_Glazkovo.SG), I7335 (Russia_Cisbaikal_Glazkovo_Ustbelaya_EBA) and DA339.SG (Russia_Shamanka_EBA.SG; SI Data 1 Table 2, Column N). All three are members of the Cisbaikal_LNBA grouping produced by the clustering procedure (SI Section V.C) and hail from sites of the Glazkovo culture [8,9,16].

Individuals of the Transbaikal_EMN cluster uniformly carry subclades of haplogroup N (including N-TAT, N-F1360, and N-L666, TRMCAs > 10kya) whose descendants are today found in a wide range of populations across Northern Eurasia, from Yukagiric-speaking populations and other populations of the Bering Straits in the East, to Uralic-speaking populations of the Baltic Sea region in the West (SI Data 1 Table 2, Column N). A subset of these are also found among individuals of the Yakutia_LNBA cluster and at Tatarka, where subclades of N-TAT more closely related to those found today among Uralic speakers (N-F1419 and N-M2126) are represented. At Rostovka, a specific subclade of N-M2126, N-Z1936 (TRMCA ~4.0kya), is found; unlike the aforementioned subclades, this clade is found at high frequency only among Uralic-speaking populations and is most prevalent among Finnic-speaking populations around the Baltic Sea. [63]

### IX. Radiocarbon and Stable Isotope Analysis

In total, we publish 76 new radiocarbon dates. At least some of these dates do not coincide with the generally accepted chronology of the archaeological context of the dated individuals. Such disparity of radiocarbon age and archaeologically-based chronology has been repeatedly reported for prehistoric populations in Northern Eurasia and are attributed to the freshwater reservoir effect (FRE). FRE is manifested in human bone samples when the population practices a fish-based diet and consumes aquatic resources enriched with non-atmospheric (i.e., aquatic) carbon [71].

The assessment of the likelihood of age discrepancies in the acquired radiocarbon dates, therefore, requires a reconstruction of dietary and determination of a possible freshwater reservoir offset (FRO).

***Dietary analysis via Stable Isotopes***

Carbon (δ13C) and nitrogen (δ15N) stable isotope ratios are extensively employed for reconstructions of diet of past human populations. The most impressive results were obtained in distinguishing marine and terrestrial diets, but lower δ13C with elevated δ15N values in human collagen from continental areas are also supposed to reflect the consumption of freshwater fish, including among pastoral and nomadic communities [72]. Although we do have not enough comparative faunal isotopic data to perform full-scale dietary research, we can elaborate on some assumptions regarding the role of freshwater resources consumption in populations studied, based on published data on δ13C and δ13N isotopes in river fish and prehistoric humans from various parts of Siberia [72,73].

The human samples in our dataset show a range from -24,5 to -16,9‰ in δ13C and from 11,6 to 16,4‰ in δ15N.

The observed range of δ15N characterizes the studied populations as non-agricultural consumers of terrestrial and aquatic fauna with a high trophic position. Privat et al. [74] consider δ15N above 13,5 ‰ as a marker of freshwater fish consumption for the Bronze Age population of the Baraba forest steppe.

The fish samples from the rivers of Northern Eurasia show decreased values of δ13C varying mostly from -30,6‰ to -22,1‰ [72,73,75]. In lake fish, the δ13C values vary in a greater range and are higher on average than in river fish, from -29,6‰ to -8,3‰, but the analyzed excavated fish samples demonstrate values vary from -29,6 to -20,2‰, which match with river fish better (ibid.). For Altai and Minusinsk herbivores, δ13C varies from -18 to -22 ‰ [76]

Based on the observed values, we assume the individuals from our dataset who demonstrate δ13N values greater than 13,5‰ as having had a significant (or even prevailing) proportion of fish in their diet. Respectively, the fish diet of the individuals with less than -22‰ δ13C was oriented mostly on river fish consumption while the others with higher δ13C values might include more recourses from lakes and other inland waterbodies.

***Freshwater reservoir offset (FRO) estimation***

Recent comprehensive observations of the freshwater reservoir effect recorded in human bones from Eurasian riparian populations [75] demonstrate high inequality of biases derived from the FRO.

The 14C dating of paired terrestrial and aquatic animals from archaeological records revealed the FRO measured up to ~1000 radiocarbon years in the fish from the Ob and the Irtysh as well as from several smaller rivers and lakes of the Kazakh Uplands [76].

Special observation on paired human-fauna 14C dating of the Baikal region revealed a positive correlation between δ13N and expected FRO [77]. Another study in the Upper Lena region revealed a moderate but significant negative correlation between human δ13C values and human-animal offsets in 14C years and no relationship between the 14C offset and δ15N [78]. Marchenko et al. [72] noticed greater values of FRO in samples from the big rivers, the Ob, and the Irtysh, relative to small tributaries and lakes.

All observations demonstrate the regional character of the recorded relations between carbon and nitrogen isotopic values and the FRO. We see no possibility at the moment to elaborate on any general regression equation for the trans-regional estimation of the expected FRO. Also, the observed values of carbon and nitrogen isotopes in our dataset do not exclude the possibility of the FRO for any individual. However, we can assess the relative probability of FRE-driven aging of the 14C dates of studied individuals based on the observed δ13N-δ13C ratio.

We expect relatively highest FRO in dated individuals with river fish-based diet, from the sites Satyga-16, Neolithic-Early Bronze Age burials at Okunevo, Tuzovskie Bugry-1 (Vasino-5), Sosnovy Island, Firsovo-11, and Chernoozerye-1 in the Ob River basin, and Maksimovka-1, Sakhtysh-2a, and Murzikha-2 in the Volga and Kama river basins. The dates of individuals from these sites require an alternative confirmation, which can be obtained from archaeological context or additional paired human-fauna dating.

The lowest probability of the FRO we expect in the individuals with the lowest values of δ15N and highest values of δ13C. This makes dates from the sites Chuiya, Khatystyr-Cave, and Manzurka-2 in the Lena basin, Boborykino, Gladunino, and Tatarka-Hill in the Ob basin, Kulmetovskiy-Grot, and Sakhtysh-2 in the Volga-Kama basin, Afontova-Gora, Bazaikha, Krasnoyarsk finds, Rasputino, Tolsty-Mys-1, and Ust'-Shamanka-1.

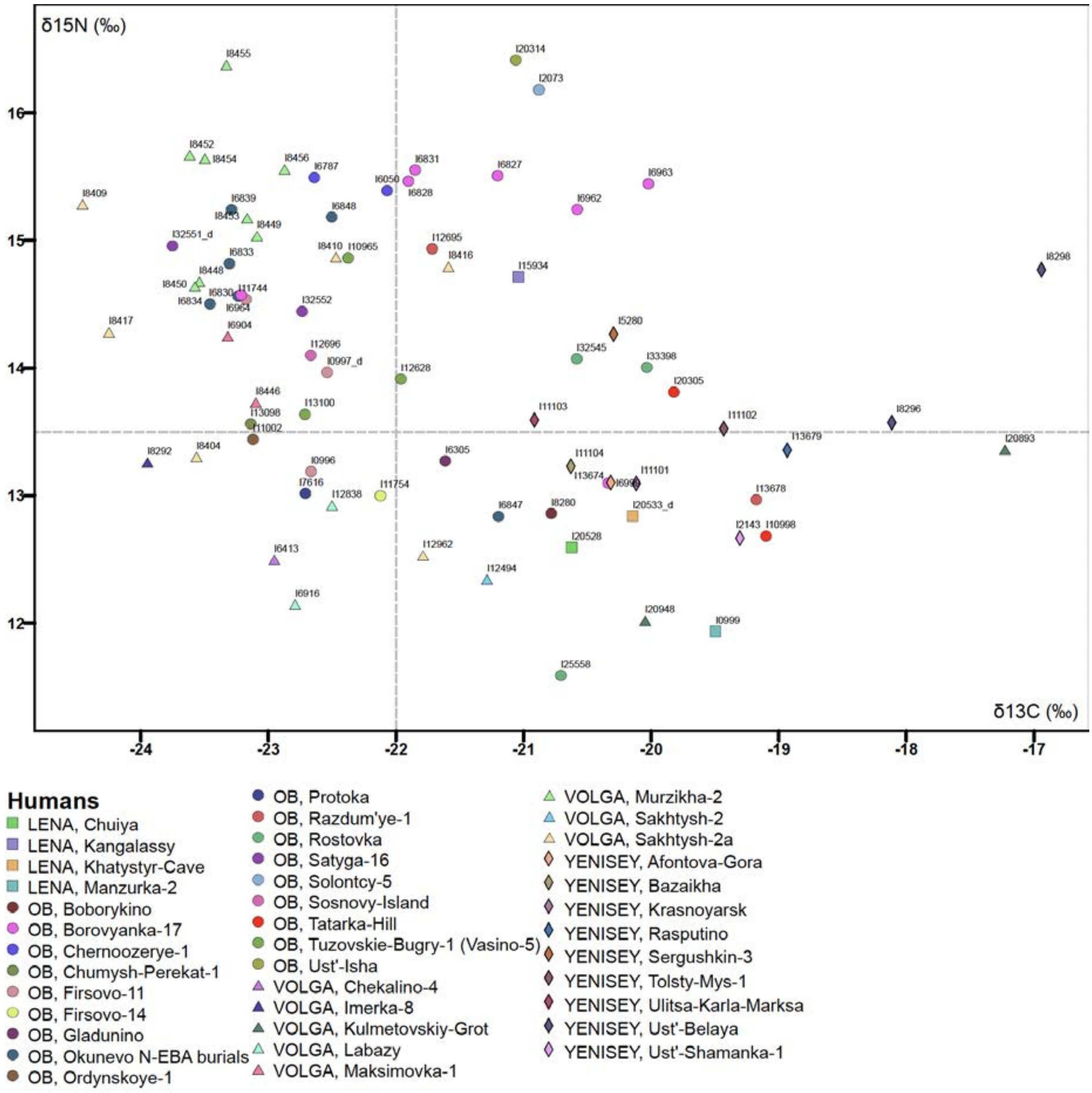

**Figure S81 | δ13C and δ15N isotopic collagen values of newly dated human bone samples**

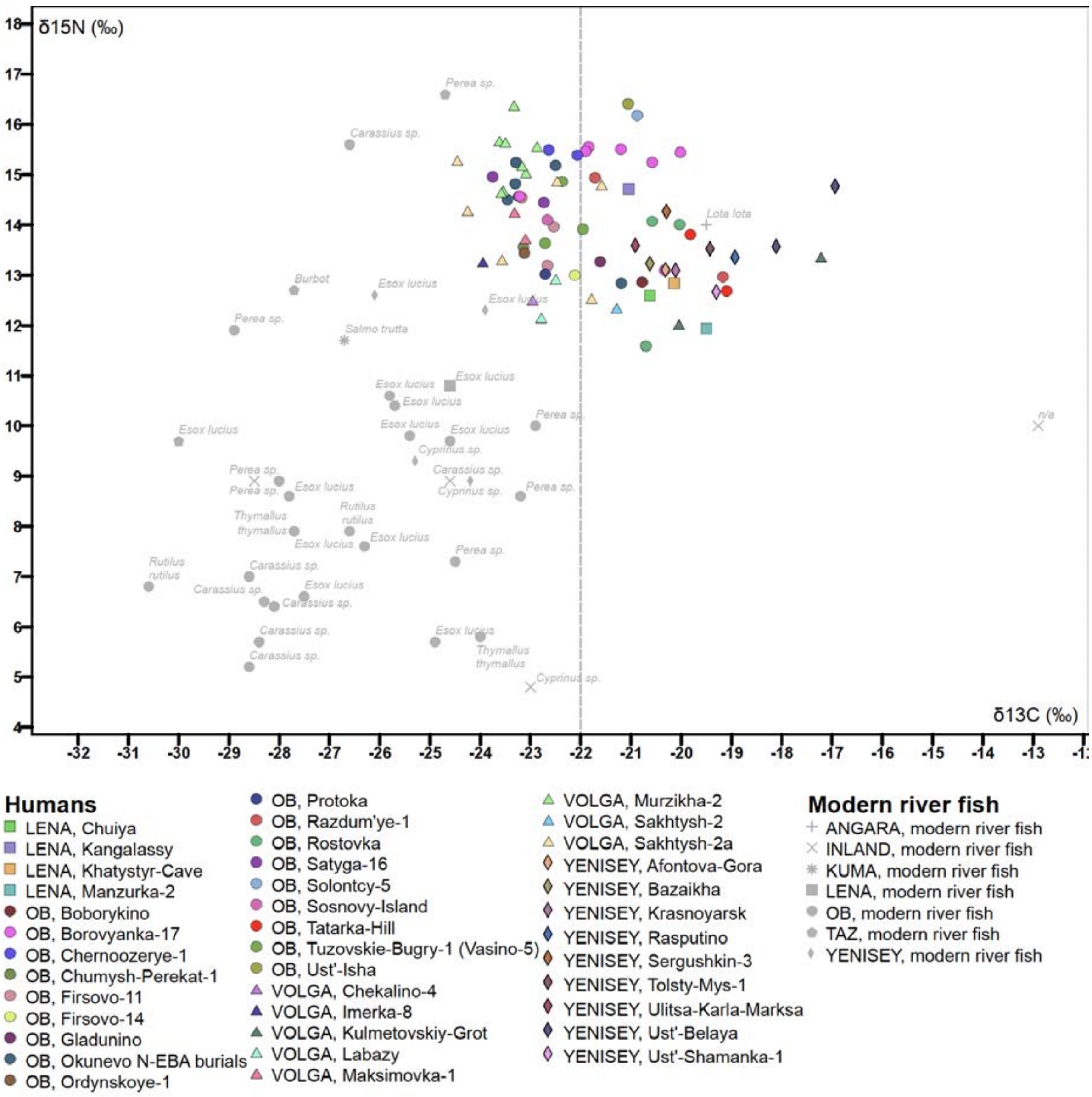

**Figure S82| δ13C and δ15N isotopic collagen values of human bone samples against modern fish from Northern Eurasia**
